# Considerable escape of SARS-CoV-2 variant Omicron to antibody neutralization

**DOI:** 10.1101/2021.12.14.472630

**Authors:** Delphine Planas, Nell Saunders, Piet Maes, Florence Guivel-Benhassine, Cyril Planchais, Julian Buchrieser, William-Henry Bolland, Françoise Porrot, Isabelle Staropoli, Frederic Lemoine, Hélène Péré, David Veyer, Julien Puech, Julien Rodary, Guy Baela, Simon Dellicour, Joren Raymenants, Sarah Gorissen, Caspar Geenen, Bert Vanmechelen, Tony Wawina-Bokalanga, Joan Martí-Carrerasi, Lize Cuypers, Aymeric Sève, Laurent Hocqueloux, Thierry Prazuck, Félix Rey, Etienne Simon-Lorrière, Timothée Bruel, Hugo Mouquet, Emmanuel André, Olivier Schwartz

**Author notes:** co-first authors (Delphine Planas, Nell Saunders, Piet Maes). co-last authors (Timothée Bruel, Hugo Mouquet, Emmanuel André, Olivier Schwartz). Corresponding authors &.

## Abstract

The SARS-CoV-2 Omicron variant was first identified in November 2021 in Botswana and South Africa^1,2^. It has in the meantime spread to many countries and is expected to rapidly become dominant worldwide. The lineage is characterized by the presence of about 32 mutations in the Spike, located mostly in the N-terminal domain (NTD) and the receptor binding domain (RBD), which may enhance viral fitness and allow antibody evasion. Here, we isolated an infectious Omicron virus in Belgium, from a traveller returning from Egypt. We examined its sensitivity to 9 monoclonal antibodies (mAbs) clinically approved or in development^3^, and to antibodies present in 90 sera from COVID-19 vaccine recipients or convalescent individuals. Omicron was totally or partially resistant to neutralization by all mAbs tested. Sera from Pfizer or AstraZeneca vaccine recipients, sampled 5 months after complete vaccination, barely inhibited Omicron. Sera from COVID-19 convalescent patients collected 6 or 12 months post symptoms displayed low or no neutralizing activity against Omicron. Administration of a booster Pfizer dose as well as vaccination of previously infected individuals generated an anti-Omicron neutralizing response, with titers 5 to 31 fold lower against Omicron than against Delta. Thus, Omicron escapes most therapeutic monoclonal antibodies and to a large extent vaccine-elicited antibodies.

---

In less than three weeks following its discovery, the Omicron variant has been detected in dozens of countries. The WHO has classified this lineage (previously known as Pango lineage B.1.1.529) as a Variant of Concern (VOC) on November 26, 2021 ^1^. Preliminary estimates of its doubling time range between 1.2 and 3.6 days, in populations with high rate of SARS-CoV-2 immunity ^2,4^. Omicron is expected to supplant the currently dominant Delta lineage in the next weeks or months. Little is known about its sensitivity to the humoral immune response. Recent preprints indicated a reduced sensitivity of Omicron to certain monoclonal and polyclonal antibodies ^5 6^, whereas CD8+ T cell epitopes previously characterized in other variants seem to be conserved in Omicron ^7^.

## Isolation and characterization of the Omicron variant

We isolated an Omicron variant from a nasopharyngeal swab of an unvaccinated individual that developed moderate symptoms eleven days after returning to Belgium from Egypt. The virus was amplified by one passage on Vero E6 cells. Sequences of the swab and the outgrown virus were identical and identified the Omicron variant (Pango lineage BA.1, GISAID accession ID: (EPI_ISL_6794907 and EPI_ISL_7413964 respectively) (Fig. 1a). The Spike protein contained 32 changes, when compared to the D614G strain (belonging to the basal B.1 lineage) used here as a reference, including 7 changes in the N terminal domain (NTD), with substitutions, deletions and a three amino-acid insertion (A67V,*Δ69-70, T95I, G142D, Δ141-143, Δ211L212I, Ins214EPE*), 15 mutations in the RBD (G339D, S371L, S373P, S375F, K417N, N440K, G446S, S477N, T478K, E484A, Q493R, G496S, Q498R and N501Y, Y505H) the T574K mutation, 3 mutations close to the furin cleavage site (H655Y, N679K and P681H) and 6 in the S2 region (*N764K, D796Y, N856K, Q954H, N969, L981F*) (Fig. 1a). This extensive constellation of changes is unique, but includes at least 11 modifications observed in other lineages and VOCs or at sites mutated in other variants (Fig. 1a). Viral stocks were titrated using S-Fuse reporter cells and Vero cells. S-Fuse cells become GFP+ upon infection, allowing rapid assessment of infectivity and the measurement of neutralizing antibody levels ^8–10^. Syncytia were observed in Omicron-infected S-Fuse cells (not shown).

**Fig. 1.**
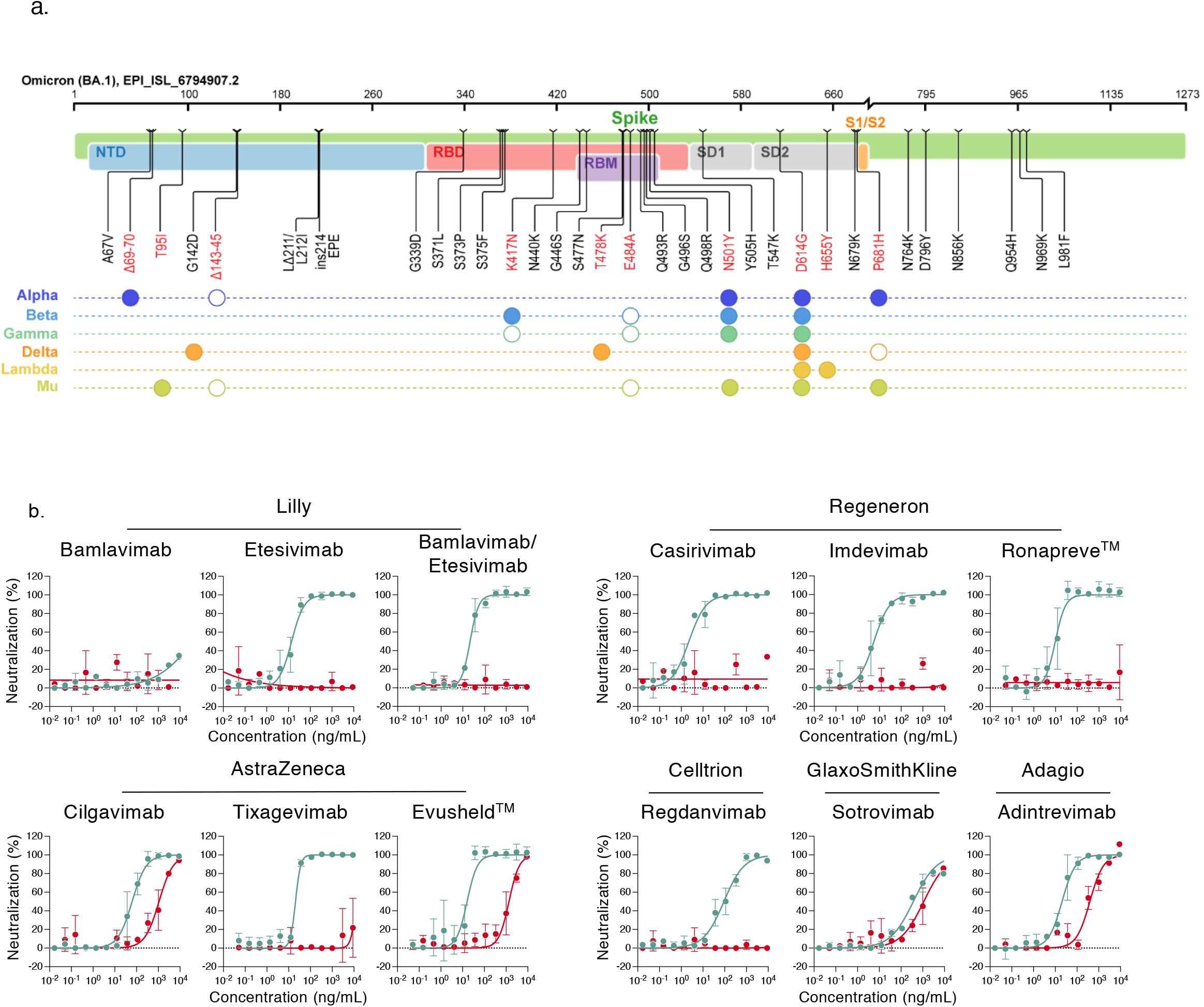
Neutralization of SARS-CoV-2 variants Delta and Omicron by clinical and pre-clinical mAbs. **a. Mutational landscape of the Omicron Spike**. The amino acid modifications are indicated in comparison to the ancestral Wuhan-Hu-1 sequence (NC_045512). Consensus sequences of the Spike protein were built with the Sierra tool ^33^. The Omicron sequence corresponds to the viral strain isolated in Belgium and used in the study (GISAID accession ID: (EPI_ISL_6794907). Mutations are compared to some preexisting variants of concern and variants of interest. Filled circles: change identical to Omicron. Open circles: different substitution at the same position. **b. Neutralization curves of mAbs**. Dose response analysis of the neutralization by clinical or pre-clinical mAbs (Bamlanivimab, Etesivimab, Casirivimab, Imdevimab, Adintrevimab, Cligavimab, Tixagevimab, Regdanvimab, Sotrovimab) and the indicated combinations (Bamlanivimab/Etesivimab, Casirivimab/Imdevimab [Ronapreve™], Cligavimab/Tixagevimab [Evusheld™]) on Delta (turquoise) and Omicron (red) variants. Data are mean±SD of 2 to 3 independent experiments. For each antibody, the IC50 are presented in Extended table 1.

## Phylogenetic analysis of the Omicron lineage

We inferred a global phylogeny subsampling SARS-CoV-2 sequences available on the GISAID EpiCoV database. To better contextualize the isolated virus genome, we performed a focused phylogenetic analysis using as background all Omicron samples deposited on GISAID on December 6, 2021, (Extended Data Fig. 1). The tree topology indicates that the Omicron lineage does not directly derive from any of the previously described VOCs. The very long branch of the Omicron lineage in the time-calibrated tree (Extended Fig. 1) might reflect a cryptic and potentially complex evolutionary history. At the time of writing, no Omicron genomic sequences from Egypt were available on GISAID, nor do we know of any sequences of travellers that used the same planes. The isolated strain genome showed no close connection to other Belgian Omicron infections. Follow-up analyses with additional genomic data will improve phylogenetic resolution to determine whether the patient was infected before or after returning to Belgium.

## Mutational landscape in variant Omicron

We highlighted the 29 amino acid substitutions, 3 amino-acid deletions and a 3-residue insertion present in the Omicron Spike, with respect to the Wuhan strain, in a 3D model of the protein (Extended Figure 2a). The 15 mutations in the RBD cluster in particular around the trimer interface. The RBD is the target of the most potently neutralizing monoclonal antibodies (mAbs) against SARS-CoV-2, which have been divided into four classes depending of the location of their epitope ^11^ (Extended Figure 2b). mAbs in classes 1 and 2 compete for hACE2 binding, whereas those from classes 3 and 4 bind away from the hACE2 interaction surface (Extended Figure 2b). The epitopes of the class 2 and 3 mAbs are exposed irrespective of the conformation of the RBD on the spike (Up or Down configuration ^12^), while those of classes 1 and 4 require an RBD in the Up conformation. Whereas the previous VOCs displayed mutations only in the region targeted by class 1 and 2 mAbs, Omicron mutations are located within the epitopes of all four classes of mAbs. In the NTD, the mutations, insertion and deletions might also impact recognition of this domain by antibodies.

**Fig. 2.**
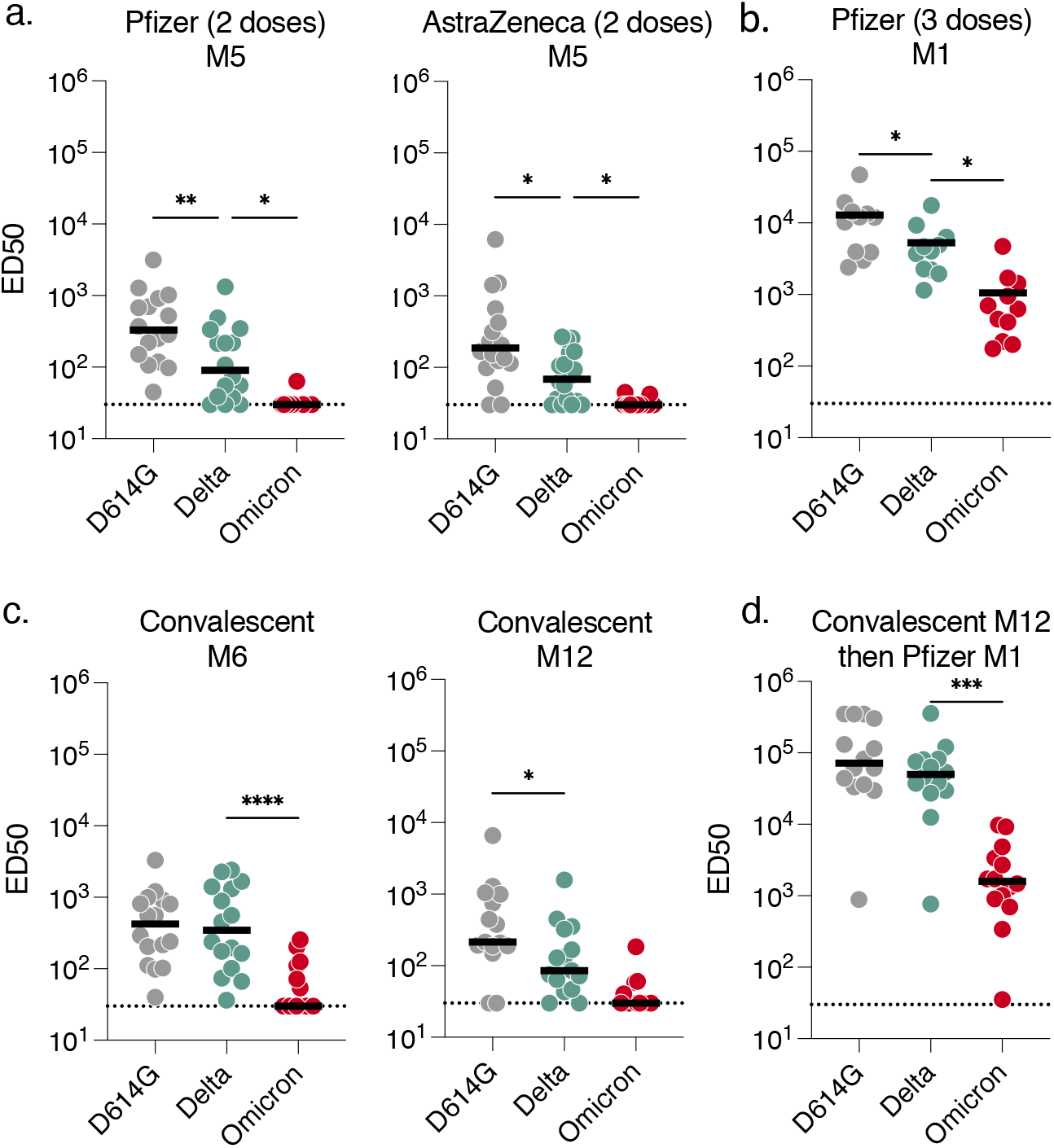
Sensitivity of SARS-CoV-2 variants D614G, Delta and Omicron to sera from vaccinated, convalescent or infected then vaccinated individuals. Neutralization titers of the sera against the three indicated viral variants are expressed as ED50 (Effective Dose 50%). a. Neutralizing activity of sera from AstraZeneca (n=18) (left panel) and Pfizer (n=16) (right panel) vaccinated recipients sampled at 5 months post-second dose. b. Neutralizing activity of sera from Pfizer vaccinated recipients sampled one month after the 3^rd^ injection (n=11). The dotted line indicates the limit of detection (ED50=30). c. Neutralizing activity of sera from convalescent individuals, sampled at 6 (n=16) months post onset of symptoms (right panel). Neutralizing activity of sera from convalescent individuals (n=15), sampled at 12 months post onset of symptoms (left panel). d. Neutralizing activity of sera from infected then vaccinated individuals, sampled one month after the 1^st^ injection (n=14). In each panel, data are mean from 2 to 3 independent experiments. Two-sided Friedman test with Dunn’s multiple comparison was performed to compare D614G and Omicron to the Delta variant. *p< 0.05, **p< 0.01, ***p< 0.001.

## Neutralization of variant Delta by monoclonal antibodies

We then assessed the sensitivity of Omicron to a panel of human mAbs using the S-Fuse assay. We tested 9 antibodies in clinical use or in development ^13,14 15 16 17 18,19^. Neutralizing mAbs targeting the RBD can be classified into 4 main classes depending on their binding epitope ^3,11,20^. Bamlanivimab and Etesevimab (class 2 and class 1, respectively) are mixed in the Lilly cocktail. Casirivimab and Imdevimab (class 1 and class 3, respectively) form the REGN-COV2 cocktail from Regeneron and Roche (Ronapreve™). Cilgavimab and Tixagevimab (class 2 and class 1, respectively) from AstraZeneca are also used in combination (Evusheld™). Regdanvimab (Regkirona™) (Celltrion) is a class 1 antibody. Sotrovimab (Xevudy™) by GlaxoSmithKline and Vir Biotechnology is a class 3 antibody that displays activity against diverse coronaviruses. It targets an RBD epitope outside the receptor binding motif, which includes N343-linked glycans. Adintrevimab (ADG20) developed by Adagio binds to an epitope located in between the class 1 and class 4 sites.

We measured the activity of the 9 antibodies described above against Omicron and included the Delta variant for comparison purposes (Fig. 1b). As previously reported, Bamlanivimab did not neutralize Delta ^10 21 22^. The other antibodies neutralized Delta with IC50 (Inhibitory Concentration 50%) varying from 2.2 to 369 ng/mL (Fig. 1b and Extented table 1). Six antibodies (Bamlanivimab, Etesevimab, Casirivimab, Imdevimab, Tixagevimab and Regdanvimab) lost antiviral activity against Omicron. The three other antibodies displayed a 3 to 20-fold increase of IC50 (ranging from 391 to 1114 ng/ml) against Omicron. Sotrovimab was the only antibody displaying a rather similar activity against both strains, with a IC50 of 369 and 1114 ng/mL against Delta and Omicron, respectively. We also tested the antibodies in combination, to mimic the therapeutic cocktails. Bamlanivimab/Etesevimab (Lilly) or Casirivimab/Imdevimab (Ronapreve™) are inactive against Omicron. Cilgavimab/Tixagevimab (Evusheld™) neutralized Omicron with an IC50 of 1355 ng/mL.

We next examined by flow cytometry the binding of each mAb to Vero cells infected with Delta and Omicron variants. Five out of six clinical antibodies that lost antiviral activity (Bamlanivimab, Etesevimab, Casirivimab, Imdevimab and Regdanvimab) no longer recognized Omicron infected cells (table S1). The other antibodies still bound to Omicron-infected cell (table S1).

Thus, Omicron escapes neutralization by the tested antibodies to various extents. Our results are in line with results from a recent preprint, obtained using Spike-coated pseudoviruses ^6^.

## Sensitivity of variant Omicron to sera from vaccine recipients

We next asked whether vaccine-elicited antibodies neutralized Omicron. To this aim, we randomly selected 36 individuals from a cohort established in the French city of Orléans, composed of vaccinated subjects that were not previously infected with SARS-CoV-2. The characteristics of vaccinees are depicted in extented table 2a. 16 individuals received the Pfizer two-dose vaccine regimen and 18 the Astrazeneca two-dose vaccine regimen. 11 individuals vaccinated with Pfizer received a booster dose. We measured the potency of their sera against the Delta and Omicron strains. We used as a control the D614G ancestral strain (belonging to the basal B.1 lineage) (Fig. 2a). We calculated the ED50 (Effective Dose 50%) for each combination of serum and virus. Sera were first sampled 5 months after the full two-dose vaccination. With the Pfizer vaccine, the levels of neutralizing antibodies were relatively low against D614G and Delta (median ED50 of neutralization of 329 and 91), reflecting the waning of the humoral response ^10^ (Fig. 2a). We did not detect any neutralization against the Omicron variant with these sera (Fig. 2a).

A similar pattern was observed with the AstraZeneca vaccine. Five months after vaccination, the levels of antibodies neutralizing Delta were low (ED50 of 187 and 68 against D614G and Delta, respectively), and no antiviral activity was observed against Omicron (Fig. 2a).

We next examined the impact of a Pfizer booster dose, administrated 8 months after Pfizer vaccination. The sera were collected one month after the third dose. The booster dose enhanced neutralization titers against D614G and Delta by 33 and 56 fold (ED50 12873 and 5280, respectively). It was also associated with strong increase of the neutralization activity against Omicron (ED50 of 1050) (Fig. 2b).

Altogether, these results indicate that Omicron is poorly or not neutralized by vaccinees’ sera sampled 5 months after vaccination. The booster dose triggered a detectable cross-neutralization activity against Omicron. However, even after the booster dose the variant displayed a reduction of ED50 of 12- and 5-fold, when compared to D614G and Delta, respectively.

## Sensitivity of variant Omicron to sera from convalescent individuals

We subsequentely examined the neutralization ability of sera from convalescent subjects. We randomly selected 45 longitudinal samples from 36 donors in a cohort of infected individuals from Orléans. Individuals were diagnosed with SARS-CoV-2 infection by RT-qPCR (Extended table 2b). We previously studied the potency of these sera against D614G, Alpha, Beta and Delta isolates ^9 10^. We analyzed individuals sampled at a median of 6 and 12 months (M6 and M12) post onset of symptoms (POS). With the D614G and Delta variants, the neutralization titers slightly decreased overtime (422 and 215 for D614G, 344 and 85 for Delta, at M6 and M12, respectively) ^9^ (Fig. 2c). The convalescent sera barely neutralized Omicron at these time points.

Fourteen individuals were vaccinated at M12 with a Pfizer dose. Sera sampled one month after vaccination showed a drastic increase in neutralizing antibody titers against the D614G and Delta variants, reaching a median ED50 of 71555 and 49778, respectively (Fig. 2d). These sera also neutralized Omicron, with a median ED50 of 1598 (Fig. 2d). Therefore, as shown with other variants ^23,24 9^ a single dose of vaccine boosts cross-neutralizing antibody responses to Omicron in previously infected individuals. The neutralization titers are however reduced by 44 and 31 fold, when compared to D614G and Delta, respectively.

## Discussion

The Omicron variant has opened a new chapter in the COVID-19 pandemic ^2,25^. The principal concerns about this variant include its high transmissibility, as underlined by its rapid spread in different countries, and the presence of over 55 mutations spanning the whole viral genome. Omicron contains 32 mutations in the Spike, lying in the NTD, RBD and in vicinity of the furin cleavage site. Some mutations were already present in other VOCs and VOIs, and have been extensively characterized ^25–27^. Due to their position, they are expected to affect the binding of natural or therapeutic antibodies, to increase affinity to ACE2 and to enhance the fusogenic activity of the Spike. Future work will help determining how this association of mutations impacts viral fitness in culture systems and their contribution to the high transmissibility of the variant.

Here, we studied the cross-reactivity of clinical or pre-clinical mAbs, as well as 90 sera from vaccine recipients and long-term convalescent individuals against an infectious Omicron isolate. We report that among nine mAb in clinical use or in development, six (Bamlanivimab, Etesevimab, Casirivimab, Imdevimab Tixagevimab and Regdanvimab) were inactive against Omicron. Two other antibodies (Cilgavimab, Andintrevimab) displayed about a 20-fold increase of IC50. Sotrovimab was less affected by Omicron’s mutations, with IC50 increased by only 3 fold. We also show that Omicron was barely neutralized by sera from vaccinated individuals sampled 5 months after administration of two doses of Pfizer or AstraZeneca vaccine. Sera from convalescent individuals at 6 or 12 months post infection barely neutralized or did not detectably neutralize Omicron.

The decrease of antibody efficacy helps explaining the high number of breakthrough infections and reinfection cases, and the spread of Omicron in both non-immune and immune individuals ^28^. There is currently no evidence of increased disease severity associated with Omicron compared with Delta, either among naïve or immunized individuals. It is likely that even if pre-existing SARS-CoV-2 antibodies may poorly prevent Omicron infection, anamnestic responses and cellular immunity will be operative to prevent severe forms of the disease ^29^.

We further report that a booster dose of Pfizer vaccine, as well as vaccination of previously infected individuals, strongly increased overall levels of anti-SARS-CoV-2 neutralizing antibodies, well above a threshold allowing inhibition of Omicron. Affinity maturation of antibodies is known to improve the efficacy of the humoral anti-SARS-CoV-2 response overtime ^30,31^. This process helps explaining the efficacy of booster doses in immune patients. However, sera with high antibody levels displayed a 5 to 31 fold reduction in neutralization efficacy against Omicron, when compared to the currently predominant Delta strain.

Potential limitations of our work include a low number of vaccine recipients and convalescents sera analyzed and the lack of characterization of cellular immunity, which is known to be more cross-reactive than the humoral response. Our results may therefore partly underestimate the residual protection offered by vaccines and previous infections against Omicron infection, in particular with regard to the severity of disease. We only analyzed sera sampled 1 month after the booster dose, or after vaccination of infected individuals. Future work with more individuals and longer survey periods will help characterize the duration of the humoral response against Omicron. We focused on immune responses elicited by Pfizer and AstraZeneca vaccination. It will be worth determining the potency of other vaccines against this variant.

Our results have important public health consequences regarding the use of therapeutic mAbs and vaccines. Clinical indications of mAbs include pre-exposure prophylaxis in individuals unable to mount an immune response, as well as prevention of COVID-19 in infected individuals at high risk for evolution towards severe disease. Antibody-based treatment strategies need to be rapidy adapted to Omicron. Experiments in preclinical models or clinical trials are warranted to assess whether the drops in IC50 are translated into impaired clinical efficacy of the mAbs that retain efficacy against Omicron. Most of the low-income countries display a weak vaccination rate, a situation that likely facilitates SARS-CoV-2 spread and continuous evolution. A booster dose significantly improves the quality and the level of the humoral immune response, and is associated with a strong protection against severe forms of the disease ^32^. An accelerated deployment of vaccines and boosters throughout the world is necessary to counteract viral spread. Our results also suggest that there is a need to update and complete the current pharmacopoeia, in particular with regard to vaccines and mAbs.

## Methods

No statistical methods were used to predetermine sample size. The experiments were not randomized and the investigators were not blinded to allocation during experiments and outcome assessment. Our research complies with all relevant ethical regulation.

### Orléans Cohort of convalescent and vaccinated individuals

Since August 27, 2020, a prospective, monocentric, longitudinal, interventional cohort clinical study enrolling 170 SARS-CoV-2-infected individuals with different disease severities, and 59 non-infected healthy controls is on-going, aiming to describe the persistence of specific and neutralizing antibodies over a 24-months period. This study was approved by the ILE DE FRANCE IV ethical committee. At enrolment, written informed consent was collected and participants completed a questionnaire which covered sociodemographic characteristics, virological findings (SARS-CoV-2 RT-PCR results, including date of testing), clinical data (date of symptom onset, type of symptoms, hospitalization), and data related to anti-SARS-CoV-2 vaccination if ever (brand product, date of first and second doses). Serological status of participants was assessed every 3 months. Those who underwent anti-SARS-CoV-2 vaccination had regular blood sampling after first dose of vaccine (ClinicalTrials.gov Identifier: NCT04750720). The primary outcome was the presence of antibodies to SARS-CoV-2 Spike protein as measured with the S-Flow assay. The secondary outcome was the presence of neutralizing antibodies as measured with the S-Fuse assay. For the present study, we selected 36 convalescent and 36 vaccinated participants. Some individuals were sampled multiple times. We analyzed a total of 90 sera. Study participants did not receive any compensation.

### Phylogenetic analysis

To contextualize the isolated Omicron genome, all SARS-CoV-2 sequences available on the GISAID EpiCov™ database as of December 06, 2021 were retrieved. A subset of complete and high coverage sequences, as indicated in GISAID, assigned to lineages B.1.529 or BA.1 and BA.2 were randomly subsampled. This subset was included in a global SARS-CoV-2 phylogeny reconstructed with augur and visualized with auspice as implemented in the Nextstrain pipeline (https://github.com/nextstrain/ncov, version from May 06, 2021) ^34^. Within Nextstrain, a random subsampling approach capping a maximum number of sequences per global region was used. The acknowledgment of contributing and originating laboratories for all sequences used in the analysis is provided in Supplementary Table 3.

### 3D representation of mutations on B1.617.2 and other variants to the Spike surface

Panels in Fig. 1 were prepared with The PyMOL Molecular Graphics System, Version 2.1 Schrödinger, LLC. The atomic model used (PDB:6XR8) has been previously described ^36^.

### S-Fuse neutralization assay

U2OS-ACE2 GFP1-10 or GFP 11 cells, also termed S-Fuse cells, become GFP+ when they are productively infected by SARS-CoV-2 ^8,9^. Cells were tested negative for mycoplasma. Cells were mixed (ratio 1:1) and plated at 8×10^3^ per well in a μClear 96-well plate (Greiner Bio-One). The indicated SARS-CoV-2 strains were incubated with serially diluted mAb or sera for 15 minutes at room temperature and added to S-Fuse cells. The sera were heat-inactivated 30 min at 56°C before use. 18 hours later, cells were fixed with 2% PFA, washed and stained with Hoechst (dilution 1:1,000, Invitrogen). Images were acquired with an Opera Phenix high content confocal microscope (PerkinElmer). The GFP area and the number of nuclei were quantified using the Harmony software (PerkinElmer). The percentage of neutralization was calculated using the number of syncytia as value with the following formula: 100 x (1 – (value with serum – value in “non-infected”)/(value in “no serum” – value in “non-infected”)). Neutralizing activity of each serum was expressed as the half maximal effective dilution (ED50). ED50 values (in μg/ml for mAbs and in dilution values for sera) were calculated with a reconstructed curve using the percentage of the neutralization at the different concentrations.

### Characteristics of the patient infected with Omicron

The 32-year-old woman was unvaccinated and developed moderate symptoms on November 22, 2021, 11 days after returning to Belgium from Egypt via Turkey (stop-over to switch flights, without having left the airport). She did not display any risk factor for severe COVID-19 and rapidly recovered. She transmitted the virus to her husband but not to their children. She provided informed written consent to use the swab for future studies. The nasopharyngeal swab tested positive for SARS-CoV-2 on this date. The leftover material of the sample was used in this study after performing routine diagnostics, within the context of the mandate that was provided to UZ/KU Leuven as National Reference Center (NRC) of respiratory pathogens, as described in detail in the Belgian Royal Decree of 09/02/2011.

### Virus strains

The reference D614G strain (hCoV-19/France/GE1973/2020) was supplied by the National Reference Centre for Respiratory Viruses hosted by Institut Pasteur (Paris, France) and headed by Pr. S. van der Werf. This viral strain was supplied through the European Virus Archive goes Global (Evag) platform, a project that has received funding from the European Union’s Horizon 2020 research and innovation program under grant agreement n° 653316. The variant strains were isolated from nasal swabs using Vero E6 cells and amplified by one or two passages. Delta was isolated from a nasopharyngeal swab of a hospitalized patient returning from India. The swab was provided and sequenced by the laboratory of Virology of Hopital Européen Georges Pompidou (Assistance Publique – Hopitaux de Paris). The Omicron-positive sample was cultured on Vero E6 cells as previously described ^37^ Viral growth was confirmed by RT-qPCR 3 days post-infection (p.i.). At day 6 p.i., a cytopathic effect (CPE) was detected and a full-length sequencing of the virus was performed. The Omicron strain was supplied and sequenced by the NRC UZ/KU Leuven (Leuven, Belgium). Both patients provided informed consent for the use of the biological materials. Titration of viral stocks was performed on Vero E6, with a limiting dilution technique allowing a calculation of TCID50, or on S-Fuse cells. Viruses were sequenced directly on nasal swabs, and after one or two passages on Vero cells. Sequences were deposited on GISAID immediately after their generation, with the following IDs: D614G: EPI_ISL_414631; Delta ID: EPI_ISL_2029113; Omicron ID: EPI_ISL_6794907.

### Flow Cytometry

Vero cells were infected with the indicated viral strains at a multiplicity of infection (MOI) of 0.1. Two days after, cells were detached using PBS-EDTA and transferred into U-bottom 96-well plates (50,000 cell/well). Cells were then incubated for 15-30 min at RT with the indicated mAbs (1 μg/mL) in PBS, 1% BSA, 0.05% sodium azide, and 0.05% Saponin. Cells were washed with PBS and stained using anti-IgG AF647 (1:600 dilution) (ThermoFisher). Stainings were also performed on control uninfected cells. Cells were then fixed in 4% PFA for 15-30 min at RT. Data were acquired on an Attune Nxt instrument using Attune Nxt Software v3.2.1 (Life Technologies) and analysed with FlowJo 10.7.1 (Becton Dickinson).

### Antibodies

Four clinically available antibodies (Bamlavimab, Casirivimab, Etesevimab and Imedvimab) were kindly provided by CHR Orleans. The other human SARS-CoV-2 anti-RBD neutralizing antibodies (ADG2 or Adintrevimab, AZD1061 (COV2-2130) or Cilgavimab, AZD8895 (COV2-2196) or Tixagevimab, CT-P59 or Regdanvimab, LY-CoV016 (CB6) or Etesevimab, LY-CoV555 or Bamlanivimab, REGN10933 or Casirivimab, REGN10987 or Imdevimab, and VIR-7831 (S309) or Sotrovimab ^13 14 15 16 17 18,19^ were produced as followed. DNA fragments coding for their IgH and IgL variable domains were synthetized (Life Technologies, Thermo Fisher Scientific). Purified digested DNA fragments were cloned into human Igγ1- and Ig κ-/ Igλ-expressing vectors ^38^ and recombinant IgG1 antibodies were produced by transient co-transfection of Freestyle™ 293-F suspension cells (Thermo Fisher Scientific) using PEI-precipitation method as previously described ^39^. IgG1 antibodies were purified by batch/gravity-flow affinity chromatography using protein G sepharose 4 fast flow beads (Cytivia) according to the manufacturer’s instructions, dialyzed against PBS using Slide-A-Lyzer® dialysis cassettes (Thermo Fisher Scientific), quantified using NanoDrop 2000 instrument (Thermo Fisher Scientific) and checked for purity and quality on a silver-stained SDS-PAGE gel (3-8% Tris-Acetate Novex, Thermo Fisher Scientific). The anti-NTD neutralizing antibody NTD-18 and the pan-coronavirus anti-S2 non-neutralizing antibody Ab-10 were previously described ^9,10^.

### Statistical analysis

Flow cytometry data were analyzed with FlowJo v10 software (TriStar). Calculations were performed using Excel 365 (Microsoft). Figures were drawn on Prism 9 (GraphPad Software). Statistical analysis was conducted using GraphPad Prism 9. Statistical significance between different groups was calculated using the tests indicated in each figure legend.

## Acknowledgments

We thank Stewart Cole for his help in initiating the collaboration between Institut Pasteur and KU Leuven We thank Nicoletta Casartelli for critical reading of the manuscript. We thank patients who participated to this study, members of the Virus and Immunity Unit and other teams for discussions and help, Nathalie Aulner and the UtechS Photonic BioImaging (UPBI) core facility (Institut Pasteur), a member of the France BioImaging network, for image acquisition and analysis. The Opera system was co-funded by Institut Pasteur and the Région ile de France (DIM1Health). We thank the KU Leuven University authorities and Jef Arnout, Bruno Lambrecht, Chris Van Geet and Luc Sels for their support. We thank Laurent Belec, Nicolas Robillard and Madelina Saliba for their help with sequencing. We thank Fabienne Peira, Vanessa Legros and Laura Courtellemont for their help with the cohorts.

## Author contributions

Experimental strategy design, experiments: DP, NS, FGB, CP, JB, WHB, FP, IS, FR, ESL, TB, OS

Vital materials: PM, CP, FL, HP, DV, JP, JR, GB, SD, JR, SG, CG, BV, TWB, JMC, LC, AS, LH, TP, HM, EA

Phylogenetic analysis: GB, ESL

Manuscript writing: DP, FR, ESL, TB, HM, EA, OS

Manuscript editing: DP, NS, PM, GB, LC, FR, ESL, TB, HM, EA, OS

## Data availability

All data supporting the findings of this study are available within the article or from the corresponding authors upon request. Source data are provided with this paper. Viral sequences are available upon request and were deposited at GISAID (https://www.gisaid.org/) under the following numbers: D614G: EPI_ISL_414631; Delta ID: EPI_ISL_2029113; Omicron ID: EPI_ISL_6794907.

**Extended data Fig. 1.**
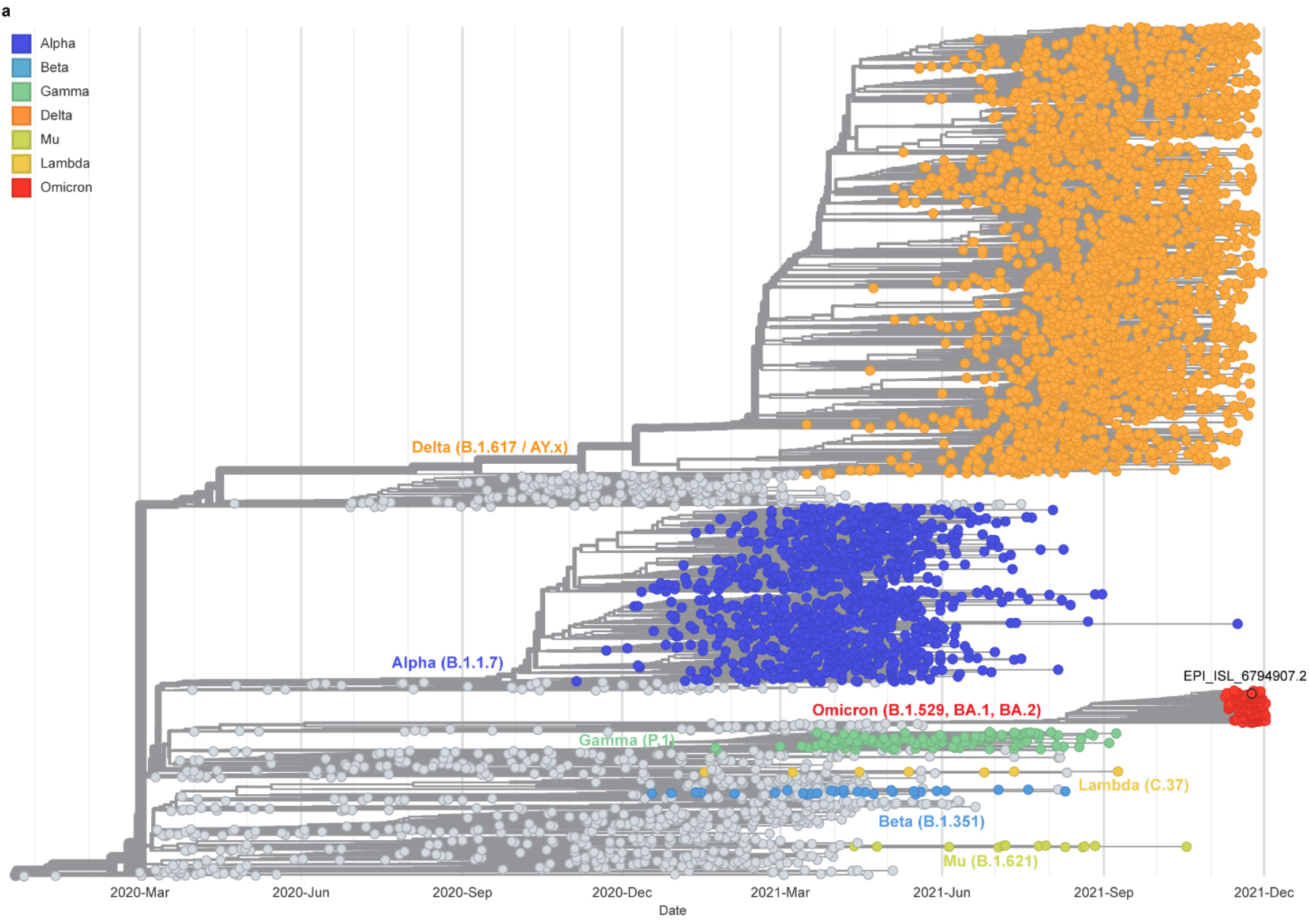
Global phylogeny of SARS-CoV-2 highlighting the Omicron lineage. Time calibrated global SARS-CoV-2 phylogeny available from the Nextstrain platform (https://nextstrain.org/ncov/gisaid/global) ^34^. The position of the isolated Omicron variant is highlighted, and the VOCs (Alpha, Beta, Gamma, Delta and Omicron) and VOIs (Lambda, Mu) are colored as indicated in the legend.

**Extended data Fig. 2.**
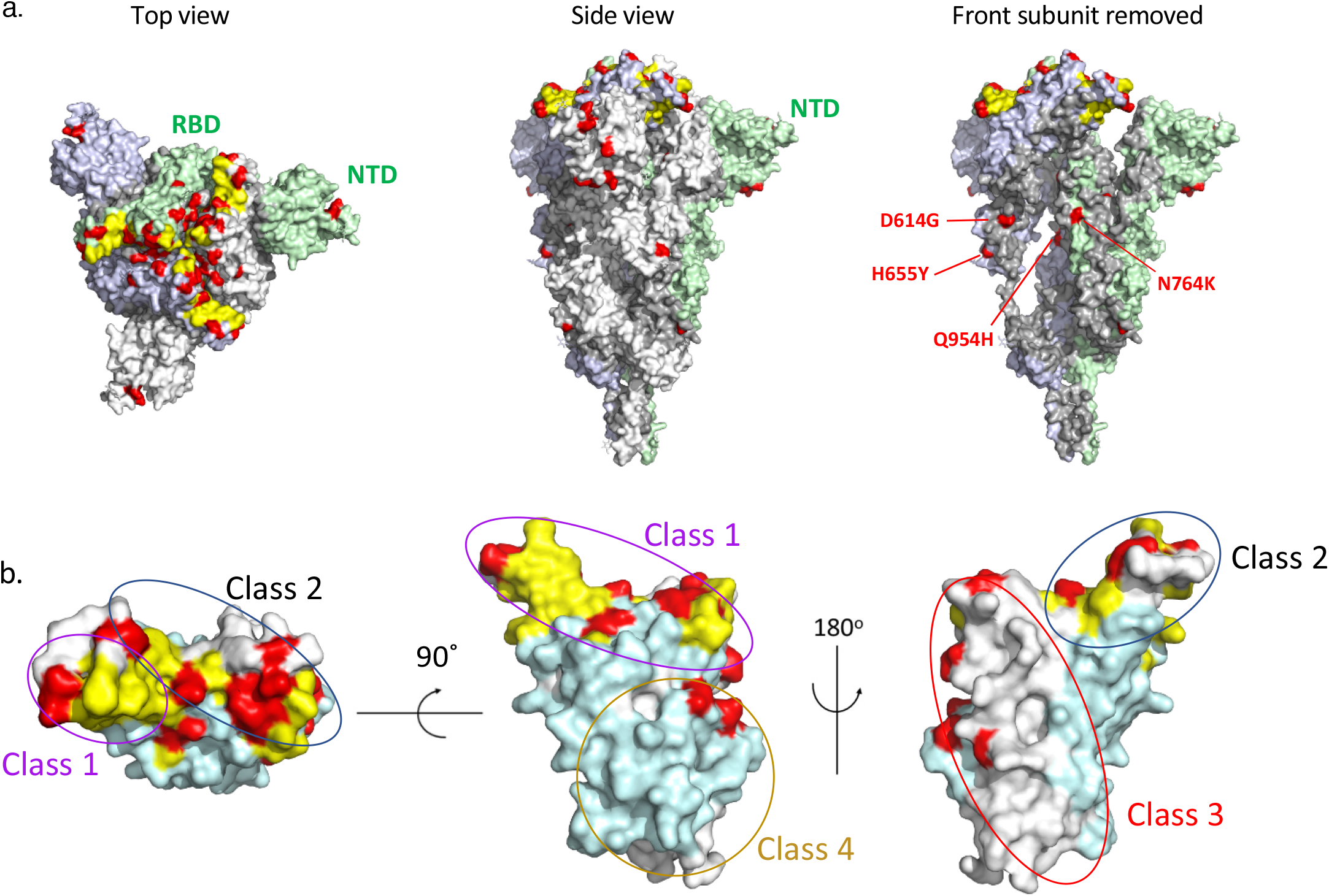
Mapping of the mutations present in Omicron to the Spike’s surface. **a**. The spike shown in top (left panel) and in side view (right panels). The spike trimer is shown in surface representation with the three protomers colored in light grey, light blue and light green. N-terminal and the receptor-binding (NTD and RBD) domains are labeled for the protomer in green only. The represented spike (PDB: 6XR8) is in the closed conformation, i.e., with all three RBDs in the “Down” conformation ^35^. The RBD surface of interaction with hACE2 (which is partially occluded in a closed spike) is colored in yellow. The amino acid differences in the spike of the Omicron variant with respect to the initial Wuhan sequence are marked in red. In the right panel, the front subunit was removed to show changes in S2 and in the C-terminal segment of S1 (labeled) that map to the trimer interface, which could impact the stability of the spike trimer. **b**. The RBD view down the hACE2 binding surface (left panel) and in two other orthogonal orientations (middle and right panel), as indicated. The hACE2 binding surface is colored in yellow and the residues altered in Omicron are in red. The RBD surfaces that are buried and exposed in a closed spike are colored in light cyan and white, respectively. The ovals outline the location of the epitopes of neutralizing antibodies of the various classes that have been described ^11^.

**Extended Table 1.** Inhibitory Concentrations (IC50) of mAbs against Delta and Omicron variants. The IC50 of the indicated mAbs (Bamlanivimab, Etesivimab, Casirivimab, Imdevimab, Adintrevimab, Cligavimab, Tixagevimab, Regdanvimab, Sotrovimab) and their combinations (Bamlanivimab/Etesivimab, Casirivimab/Imdevimab [Ronapreve™], Cligavimab/Tixagevimab [Evusheld™]) were calculated from the neutralization curves displayed in Fig. 1b. Results are in ng/mL. Color code: Grey: inactive mAbs. Green: mAbs displaying a neutralizing activity. The binding activity was measured by flow cytometry on Vero cells infected with the indicated variants. +: binding detected. -: no binding detected.

|  |  | Netrailization (IC50 ng/ml) |  | Binding |  |
| --- | --- | --- | --- | --- | --- |
|  |  | Delta | Omicron | Delta | Omicron |
| Lilly | Bamlavimab | >9000 | >9000 | +/- | - |
|  | Etesivimab | 14 | >9000 | + | - |
|  | Bamla/Ete | 21.7 | >9000 | + | ND |
| Regeneron | Casirivimab | 2.2 | >9000 | + | - |
|  | Imdevimab | 5.7 | >9000 | + | - |
|  | Ronapreve™ | 98 | >9000 | + | ND |
| AstraZeneca | Cilgavimab | 62 | 1079 | + | + |
|  | Tixagevimab | 20 | >9000 | + | + |
|  | Evusheld™ | 16 | 1355 | + | ND |
| Celltrion | Regdanvimab | 92 | >9000 | + | - |
| GlaxoSmithKline | Sotrovimab | 369 | 1114 | + | + |
| Adagio | Adintrevimab | 20 | 391 | + | + |

**Extended Table 2.** **Characteristics of the two cohorts of vaccinated and convalescent individuals. Supplementary Table 1. Contributing and originating laboratories for all sequences used in extended data Fig. 1.**

a. Vaccines
|  |  | **AstraZeneca**<br>**(2 doses)** | **Pfizer**<br>**(2 doses)** | **Pfizer**<br>**(3 doses)** |
| --- | --- | --- | --- | --- |
|  |  | **n=18** | **n=16** | **n=11** |
| Sex |  |  |  |  |
|  | Female | 14 | 8 | 3 |
|  | Male | 4 | 10 | 8 |
| Age (median; range) |  | 60 (55;78) | 59 (34;74) | 65 (48;74) |
| Immune deficiency |  | 0 | 0 | 0 |
| Previous COVID-19 |  | 0 | 0 | 0 |
| Anti-N |  | 0 | 0 | 0 |
| 1st shot |  | Feb 5 – April 7, 2021 | Jan 4- Feb 4, 2021 | Jan 4- Feb 4, 2021 |
| 2nd shot |  | May3-19, 2021 | Jan 29 - March 3, 2021 | Jan 29 - March 3, 2021 |
| 3rd shot |  | - | - | July 1 – Sept 30, 2021 |
| Sampling days post-2<sup>nd</sup><br>dose; M5 (median; range) |  | 150<br>(110-178) | 160<br>(153-176) | - |
| Sampling days post-3<sup>rd</sup><br>dose; M1 (median; range) |  | - | - | 36<br>(28-89) |

b. Convalescents
|  |  | **6M POS**<br>**n=16** | **12M POS**<br>**n=15** | **12M POS**<br>**than vaccinated**<br>**n=14** |
| --- | --- | --- | --- | --- |
| Sex |  |  |  |  |
|  | Female | 10 | 6 | 4 |
|  | Male | 6 | 9 | 10 |
| Age (Median; range) |  | 56 (32;77) | 52 (45;82) | 56 (45 ;82) |
| Severity |  |  |  |  |
|  | Severe | 8 | 10 | 9 |
|  | Mild-Moderate | 7 | 5 | 4 |
|  | Asymptomatic | 1 | 0 | 1 |
| PCR |  | 15 | 14 | 12 |
| Anti-S (S-Flow) |  | 16 | 15 | 14 |
| Sampling days POS (median;<br>range) |  | 184<br>(129;195) | 363<br>(344-383) | - |
| Days between the vaccination<br>and onset of symptoms (median;<br>range) |  | - | - | 392<br>(365-444) |
| Sampling days post-vaccination<br>(median; range) |  | - | - | 33<br>(31-47) |

