## supplementary table 1 for "Considerable escape of SARS-CoV-2 variant Omicron to antibody neutralization"

| gisaid_epi_isl | originating_lab | submitting_lab | authors |
|---|---|---|---|
| EPI_ISL_3568176 | Area of Virology, Serology and Virology Division (SAViD), New South Wales Health Pathology Randwick | Virology Research Laboratory; Area of Virology, Serology and Virology Division (SAViD), New South Wales Health Pathology Randwick | Foster et al |
| EPI_ISL_591803 | Microbiological Diagnostic Unit - Public Health Laboratory (MDU-PHL) | MDU-PHL | Seemann T. et al |
| EPI_ISL_3242859 | AZDelta | AZ Delta Medical Laboratories in Roeselare, Belgium | Geert Martens et al |
| EPI_ISL_520836 | Victorian Infectious Diseases Reference Laboratory (VIDRL) | VIDRL and MDU-PHL | Caly L. et al |
| EPI_ISL_7130344 | AGES-Institute for medical Microbiology and Hygiene Vienna | AGES-Institute for medical Microbiology and Hygiene Vienna | Kathrin Lippert et al |
| EPI_ISL_6891760 | AGES-Institute for medical Microbiology and Hygiene Vienna | AGES-Institute for medical Microbiology and Hygiene Vienna | Kathrin Lippert et al |
| EPI_ISL_5946824 | ZOL | Jessa | Severine Berden et al. on behalf of the Jessa_cmdLab et al |
| EPI_ISL_4992862 | Dutch COVID-19 response team | National Institute for Public Health and the Environment (RIVM) | Adam Meijer et al |
| EPI_ISL_7041396 | AZDelta | AZ Delta Medical Laboratories in Roeselare, Belgium | Geert Martens et al |
| EPI_ISL_4556534 | Australian Clinical Labs (formerly Healthscope Pathology) | NSW Health Pathology - Institute of Clinical Pathology and Medical Research; Westmead Hospital; University of Sydney | Arnott A. et al |
| EPI_ISL_5113803 | Cliniques universitaires Saint-Luc | UCLouvain/IREC/MBLG | Jean Ruelle et al |
| EPI_ISL_7130189 | AGES-Institute for medical Microbiology and Hygiene Vienna | AGES-Institute for medical Microbiology and Hygiene Vienna | Marion Blaschitz et al |
| EPI_ISL_2828082 | Histopath | NSW Health Pathology - Institute of Clinical Pathology and Medical Research; Westmead Hospital; University of Sydney | CIDM-PH et al. |
| EPI_ISL_521786 | Microbiological Diagnostic Unit - Public Health Laboratory (MDU-PHL) | MDU-PHL | Seemann T. et al |
| EPI_ISL_477128 | Child Health Research Foundation | Child Health Research Foundation | Senjuti Saha et al |
| EPI_ISL_3184992 | Laverty Pathology | NSW Health Pathology - Institute of Clinical Pathology and Medical Research; Westmead Hospital; University of Sydney | CIDM-PH et al. |
| EPI_ISL_4032415 | Lifebrain Covid Labor GmbH | Dept. of Laboratory Medicine | Harald Esterbauer et al |

| | | | |
|---|---|---|---|
| EPI_ISL_4404919 | Instituto Nacional de Epidemiología "Dr. Jara" | Área de Secuenciación del Laboratorio de Virología del Hospital de Niños Dr. Ricardo Gutierrez on behalf of 'Proyecto Argentino Interinstitucional de genomica de SARS-CoV-2' (PAIS Consortium) | Nabaes Jodar et al |
| EPI_ISL_6386622 | PHV-FSS | PHV-FSS | Son Nguyen et al. |
| EPI_ISL_4647889 | CLINIQUE N-D DE GRACE | Institut de Pathologie et Genetique (IPG) | Pascale Hilbert et al |
| EPI_ISL_7016910 | ACT Pathology | Schwessinger Lab | Austin Bird et al |
| EPI_ISL_4804713 | Lab voor klinische biologie | Lab voor klinische biologie | May-Linh Truong et al |
| EPI_ISL_512719 | PathWest Laboratory Medicine WA | PathWest Laboratory Medicine WA Microbial Surveillance Unit | PathWest Laboratory Medicine WA Microbial Surveillance Unit et al |
| EPI_ISL_1621212 | Plateforme de testing Namuroise | Plateforme de testing Namuroise | Céline Maschietto et al |
| EPI_ISL_3389799 | Dutch COVID-19 response team | National Institute for Public Health and the Environment (RIVM) | Adam Meijer et al |
| EPI_ISL_6490577 | Heilig hart Lier | Imelda Ziekenhuis | Johan Frans et al |
| EPI_ISL_6698202 | Plateforme de testing Namuroise | Plateforme de testing Namuroise | Gilliard Nicolas et al |
| EPI_ISL_583559 | Universitaetsklinik für Innere Medizin II Innsbruck | Bergthaler laboratory, CeMM Research Center for Molecular Medicine of the Austrian Academy of Sciences | Alexandra Popa et al |
| EPI_ISL_1014619 | Dutch COVID-19 response team | National Institute for Public Health and the Environment (RIVM) | Adam Meijer et al |
| EPI_ISL_6958955 | New South Wales Health Pathology Royal Prince Alfred Hospital | Microbiology RPAH | Foster et al |
| EPI_ISL_1904587 | AZDelta | AZ Delta Medical Laboratories in Roeselare, Belgium | Geert Martens et al |
| EPI_ISL_4884008 | Microbiological Diagnostic Unit - Public Health Laboratory (MDU-PHL) | Microbiological Diagnostic Unit - Public Health Laboratory (MDU-PHL) | Seemann et al |
| EPI_ISL_6829577 | New South Wales Health Pathology Royal Prince Alfred Hospital | Microbiology RPAH | Foster et al |
| EPI_ISL_3800616 | New South Wales Health Pathology Royal Prince Alfred Hospital | Microbiology RPAH | Foster et al |
| EPI_ISL_3050794 | Microbiological Diagnostic Unit - Public Health Laboratory (MDU-PHL) | MDU-PHL | Seemann T. et al |
| EPI_ISL_5941593 | Capital Pathology | Schwessinger Lab | Carolina Correa Ospina et al |
| EPI_ISL_3502291 | Histopath | NSW Health Pathology - Institute of Clinical Pathology and Medical Research; Westmead Hospital; University of Sydney | Arnott A. et al |

| | | | |
|---|---|---|---|
| EPI_ISL_6902052 | Institute of Virology, Department of Hygiene, Microbiology and Public Health at Innsbruck Medical University | Institute of Virology, Department of Hygiene, Microbiology and Public Health at Innsbruck Medical University | David Bante et al |
| EPI_ISL_3835094 | ZOL | Jessa | Marijke Raymaekers et al. on behalf of the Jessa_cmdLab et al |
| EPI_ISL_3918637 | Dutch COVID-19 response team | National Institute for Public Health and the Environment (RIVM) | Adam Meijer et al |
| EPI_ISL_6902053 | Institute of Virology, Department of Hygiene, Microbiology and Public Health at Innsbruck Medical University | Institute of Virology, Department of Hygiene, Microbiology and Public Health at Innsbruck Medical University | David Bante et al |
| EPI_ISL_6972689 | Dept. of Laboratory Medicine | Dept. of Laboratory Medicine | Harald Esterbauer et al |
| EPI_ISL_663334 | Microbiological Diagnostic Unit - Public Health Laboratory (MDU-PHL) | MDU-PHL | Seemann T. et al |
| EPI_ISL_792332 | Instituto de Investigaciones Biomédicas en Retrovirus y SIDA | Área de Secuenciación del Laboratorio de Virología del Hospital de Niños Dr. Ricardo Gutierrez on behalf of 'Proyecto Argentino Interinstitucional de genomica de SARS-CoV-2' (PAIS Consortium) | Nabaes Jodar et al |
| EPI_ISL_5305648 | Pathology West - NSW Health Pathology | NSW Health Pathology - Institute of Clinical Pathology and Medical Research; Westmead Hospital; University of Sydney | Arnott A. et al |
| EPI_ISL_4003468 | Laboratorio Central, Ministerio de Salud Córdoba | Instituto de Patologia Vegetal (CIAP-INTA) on behalf of 'Proyecto Argentino Interinstitucional de genomica de SARS-CoV-2' (PAIS Consortium) | Fernández et al |
| EPI_ISL_3714168 | Plateforme de testing Namuroise | Plateforme de testing Namuroise | Nicolas Gillard et al |
| EPI_ISL_663632 | Microbiological Diagnostic Unit - Public Health Laboratory (MDU-PHL) | MDU-PHL | Seemann T. et al |
| EPI_ISL_522236 | Victorian Infectious Diseases Reference Laboratory (VIDRL) | VIDRL and MDU-PHL | Caly L. et al |
| EPI_ISL_3120301 | AZDelta | AZ Delta Medical Laboratories in Roeselare, Belgium | Geert Martens et al |
| EPI_ISL_5655968 | Servicio Virosis Respiratorias-Departamento Virología-INEI | Instituto Nacional Enfermedades Infecciosas C.G.Malbran | Baumeister E. et al |
| EPI_ISL_563803 | Microbiological Diagnostic Unit - Public Health Laboratory (MDU-PHL) | MDU-PHL | Seemann et al |
| EPI_ISL_518616 | Microbiological Diagnostic Unit - Public Health Laboratory (MDU-PHL) | MDU-PHL | Seemann T. et al |

| | | | |
|---|---|---|---|
| EPI_ISL_3535296 | AZDelta | AZ Delta Medical Laboratories in Roeselare, Belgium | Geert Martens et al |
| EPI_ISL_5533904 | Histopath | NSW Health Pathology - Institute of Clinical Pathology and Medical Research; Westmead Hospital; University of Sydney | Arnott A. et al |
| EPI_ISL_3911543 | ZOL | Jessa | Marijke Raymaekers et al. on behalf of the Jessa_cmdLab et al |
| EPI_ISL_4884068 | Microbiological Diagnostic Unit - Public Health Laboratory (MDU-PHL) | Microbiological Diagnostic Unit - Public Health Laboratory (MDU-PHL) | Seemann et al |
| EPI_ISL_518909 | Microbiological Diagnostic Unit - Public Health Laboratory (MDU-PHL) | MDU-PHL | Seemann T. et al |
| EPI_ISL_427701 | Centre for Infectious Diseases and Microbiology Public Health | NSW Health Pathology - Institute of Clinical Pathology and Medical Research; Westmead Hospital; University of Sydney | Draper J et al |
| EPI_ISL_1756022 | New South Wales Health Pathology Royal Prince Alfred Hospital | Microbiology RPAH | Foster et al |
| EPI_ISL_2037425 | Arlon | Plateforme de testing Namuroise | Lesly Nyinkeu Kemamen et al |
| EPI_ISL_6864915 | Pathology West - NSW Health Pathology | NSW Health Pathology - Institute of Clinical Pathology and Medical Research; Westmead Hospital; University of Sydney | Arnott A. et al |
| EPI_ISL_520954 | Microbiological Diagnostic Unit - Public Health Laboratory (MDU-PHL) | MDU-PHL | Seemann T. et al |
| EPI_ISL_3089051 | Cliniques universitaires Saint-Luc | UCLouvain/IREC/MBLG | Jean Ruelle et al |
| EPI_ISL_5936495 | Histopath | NSW Health Pathology - Institute of Clinical Pathology and Medical Research; Westmead Hospital; University of Sydney | Arnott A. et al |
| EPI_ISL_6825551 | Territory Pathology | Territory Pathology | Ella Meumann et al |
| EPI_ISL_1041979 | Jessa | Jessa | Cruys et al. on behalf of the Jessa_cmdLab et al |
| EPI_ISL_5433221 | Lab voor klinische biologie | Lab voor klinische biologie | May-Linh Truong et al |
| EPI_ISL_5158904 | Servicio Virosis Respiratorias-Departamento Virología-INEI | Instituto Nacional Enfermedades Infecciosas C.G.Malbran | Baumeister E. et al |
| EPI_ISL_4356680 | Austrian Agency for Health and Food Safety (AGES) | Bergthaler laboratory, CeMM Research Center for Molecular Medicine of the Austrian Academy of Sciences | Lukas Endler et al |
| EPI_ISL_4520287 | Capital Pathology | Schwessinger Lab | Carolina Correa Ospina et al |
| EPI_ISL_426872 | Victorian Infectious Diseases Reference Laboratory (VIDRL) | Microbiological Diagnostic Unit Public Health Laboratory and Victorian Infectious Diseases Reference Laboratory, Doherty Institute | Caly L. et al |

| | | | |
|---|---|---|---|
| EPI_ISL_4788935 | Microbiological Diagnostic Unit - Public Health Laboratory (MDU-PHL) | Microbiological Diagnostic Unit - Public Health Laboratory (MDU-PHL) | Seemann et al |
| EPI_ISL_519222 | Microbiological Diagnostic Unit - Public Health Laboratory (MDU-PHL) | MDU-PHL | Seemann T. et al |
| EPI_ISL_2955593 | The Caribbean Public Health Agency | Carrington Lab, Department of Preclinical Sciences, Faculty of Medical Sciences, The University of the West Indies, St Augustine Campus | Nikita S. D. Sahadeo et al |
| EPI_ISL_4461057 | Austrian Agency for Health and Food Safety (AGES) | Bergthaler laboratory, CeMM Research Center for Molecular Medicine of the Austrian Academy of Sciences | Lukas Endler et al |
| EPI_ISL_3869175 | Lab voor klinische biologie | Lab voor klinische biologie | Marija Janevska et al |
| EPI_ISL_6829575 | New South Wales Health Pathology Royal Prince Alfred Hospital | Microbiology RPAH | Foster et al |
| EPI_ISL_7130486 | AGES-Institute for medical Microbiology and Hygiene Vienna | AGES-Institute for medical Microbiology and Hygiene Vienna | Marion Blaschitz et al |
| EPI_ISL_5942022 | ACT Pathology | Schwessinger Lab | Carolina Correa Ospina et al |
| EPI_ISL_3374156 | New South Wales Health Pathology Royal Prince Alfred Hospital | Microbiology RPAH | Foster et al |
| EPI_ISL_1180768 | Austrian Agency for Health and Food Safety (AGES) | Bergthaler laboratory, CeMM Research Center for Molecular Medicine of the Austrian Academy of Sciences | Lukas Endler et al |
| EPI_ISL_1557235 | Marche en Famenne | Plateforme de testing Namuroise | Céline Maschietto et al |
| EPI_ISL_5941894 | ACT Pathology | Schwessinger Lab | Carolina Correa Ospina et al |
| EPI_ISL_4254989 | New South Wales Health Pathology Royal Prince Alfred Hospital | Microbiology RPAH | Foster et al |
| EPI_ISL_4636700 | ACT Pathology | Schwessinger Lab | Carolina Correa Ospina et al |
| EPI_ISL_3186110 | St Vincent's Pathology (SydPath) | NSW Health Pathology - Institute of Clinical Pathology and Medical Research; Westmead Hospital; University of Sydney | CIDM-PH et al. |
| EPI_ISL_3868935 | Lab voor klinische biologie | Lab voor klinische biologie | Marija Janevska et al |
| EPI_ISL_2192527 | Lab voor klinische biologie | Lab voor klinische biologie | Marija Janevska et al |
| EPI_ISL_6268044 | 4Cyte Pathology | NSW Health Pathology - Institute of Clinical Pathology and Medical Research; Westmead Hospital; University of Sydney | Arnott A. et al |
| EPI_ISL_5433299 | Lab voor klinische biologie | Lab voor klinische biologie | May-Linh Truong et al |
| EPI_ISL_3655567 | The Caribbean Public Health Agency | Carrington Lab, Department of Preclinical Sciences, Faculty of Medical Sciences, The University of the West Indies, St Augustine Campus | Nikita S. D. Sahadeo et al |

| | | | |
|---|---|---|---|
| EPI_ISL_3568368 | Area of Virology, Serology and Virology Division (SAViD), New South Wales Health Pathology Randwick | Virology Research Laboratory; Area of Virology, Serology and Virology Division (SAViD), New South Wales Health Pathology Randwick | Foster et al |
| EPI_ISL_5778641 | Histopath | NSW Health Pathology - Institute of Clinical Pathology and Medical Research; Westmead Hospital; University of Sydney | Arnott A. et al |
| EPI_ISL_6589116 | Microbiological Diagnostic Unit - Public Health Laboratory (MDU-PHL) | Microbiological Diagnostic Unit - Public Health Laboratory (MDU-PHL) | Seemann et al |
| EPI_ISL_563599 | Microbiological Diagnostic Unit - Public Health Laboratory (MDU-PHL) | MDU-PHL | Seemann et al |
| EPI_ISL_3867784 | Lab voor klinische biologie | Lab voor klinische biologie | Marija Janevska et al |
| EPI_ISL_1383235 | South Eastern Area Laboratory Services (SEALS) | NSW Health Pathology - Institute of Clinical Pathology and Medical Research; Westmead Hospital; University of Sydney | CIDM-PH et al. |
| EPI_ISL_5881525 | Plateforme de testing Namuroise | Plateforme de testing Namuroise | Gilliard Nicolas et al |
| EPI_ISL_6825546 | Institute of Virology, Department of Hygiene, Microbiology and Public Health at Innsbruck Medical University | Institute of Virology, Department of Hygiene, Microbiology and Public Health at Innsbruck Medical University | David Bante et al |
| EPI_ISL_6774084 | Botswana Harvard HIV Reference Laboratory | Botswana Harvard HIV Reference Laboratory | Sikhulile Moyo et al |
| EPI_ISL_6901960 | LATE - Laboratório de Técnicas Especiais - Hospital Israelita Albert Einstein | LATE - Laboratório de Técnicas Especiais - Hospital Israelita Albert Einstein | Deyvid Amgarten et al |
| EPI_ISL_2017397 | HLAGYN - Laboratorio de Imunologia de Transplantes de Goias | HLAGYN - Laboratorio de Imunologia de Transplantes de Goias | Fernando Antonio Vinhal dos Santos et al |
| EPI_ISL_3235235 | Laboratorio Central de Saude Publica do Estado da Bahia (LACEN/BA) | Laboratory of Respiratory Viruses and Measles, Oswaldo Cruz Institute, FIOCRUZ | Paola Resende et al |
| EPI_ISL_1795363 | UNIDADE BASICA DE SAUDE DE PIQUETE | Instituto Butantan / ESALQ-Piracicaba | Instituto Butantan: Alexander Roberto Precioso et al |
| EPI_ISL_5366221 | KU Leuven, Rega Institute, Clinical and Epidemiological Virology | KU Leuven, Rega Institute, Clinical and Epidemiological Virology | Tony Wawina-Bokalanga et al |
| EPI_ISL_1963349 | Platform BIS UZA/UAntwerpen | Labo Klinische Biologie, UZA | Jasmine Coppens et al |
| EPI_ISL_6780130 | Platform BIS UZA/UAntwerpen | Labo Klinische Biologie, UZA | Jasmine Coppens et al |
| EPI_ISL_5146209 | Central Public Health Laboratory - LACEN -Bahia, Salvador, Brazil | Central Public Health Laboratory - LACEN -Bahia, Salvador, Brazil | Stephane Tosta et al |
| EPI_ISL_6752027 | Botswana Harvard HIV Reference Laboratory | Botswana Harvard HIV Reference Laboratory | Sikhulile Moyo et al |
| EPI_ISL_2861702 | Platform BIS UZA/UAntwerpen | Labo Klinische Biologie, UZA | Marie Le Mercier et al |
| EPI_ISL_5429659 | Dutch COVID-19 response team | National Institute for Public Health and the Environment (RIVM) | Adam Meijer et al |

| | | | |
|---|---|---|---|
| EPI_ISL_3799190 | CHECK-UP - Medicina e Diagnóstico - Matriz | HLAGYN - Laboratorio de Imunologia de Transplantes de Goias | Fernando Antonio Vinhal dos Santos et al |
| EPI_ISL_3050543 | Laboratório Central de Saúde Pública do Amazonas - LACEN-AM | Laboratorio de Ecologia de Doencas Transmissiveis na Amazonia, Instituto Leonidas e Maria Deane - Fiocruz Amazonia | Valdinete Nascimento et al |
| EPI_ISL_1469707 | Hospital Bom Jesus | Epiclin | Fernando Hayashi Sant'Anna et al |
| EPI_ISL_2196325 | Labortorio Central de Saude Publica do Estado do Rio de Janeiro (LACEN/RJ) | Laboratory of Respiratory Viruses and Measles, Oswaldo Cruz Institute, FIOCRUZ | Paola Resende et al |
| EPI_ISL_6774093 | Botswana Harvard HIV Reference Laboratory | Botswana Harvard HIV Reference Laboratory | Sikhulile Moyo et al |
| EPI_ISL_2362595 | Platform BIS UZA/UAntwerpen | Labo Klinische Biologie, UZA | Marie Le Mercier et al |
| EPI_ISL_1966812 | USF JARDIM SAO JOAO SCRPARDO | Instituto Butantan / Mendelics | Instituto Butantan: Dimas Tadeu Covas et al |
| EPI_ISL_1795295 | USF GUACUANO | Instituto Butantan / ESALQ-Piracicaba | Instituto Butantan: Alexander Roberto Precioso et al |
| EPI_ISL_3664131 | DASA | DASA | Rodrigo Guarischi et al |
| EPI_ISL_5385636 | KU Leuven, Rega Institute, Clinical and Epidemiological Virology | KU Leuven, Rega Institute, Clinical and Epidemiological Virology | Tony Wawina-Bokalanga et al |
| EPI_ISL_4299854 | Palapye Primary Hospital Laboratory | Botswana Harvard HIV Reference Laboratory | Sikhulile Moyo et al |
| EPI_ISL_6774087 | Botswana Harvard HIV Reference Laboratory | Botswana Harvard HIV Reference Laboratory | Sikhulile Moyo et al |
| EPI_ISL_4414250 | DASA | DASA | Rodrigo Guarischi et al |
| EPI_ISL_2017276 | HLAGYN - Laboratorio de Imunologia de Transplantes de Goias | HLAGYN - Laboratorio de Imunologia de Transplantes de Goias | Fernando Antonio Vinhal dos Santos et al |
| EPI_ISL_6640919 | Botswana Harvard HIV Reference Laboratory | Botswana Harvard HIV Reference Laboratory | Sikhulile Moyo et al |
| EPI_ISL_2241608 | Laboratório Central de Saúde Pública do Rio Grande do Norte | Coordenação Geral de Laboratórios de Saúde Pública (CGLAB/DAEVS/SVS/MS) | Vagner Fonseca et al |
| EPI_ISL_7074135 | National Platform bis COVID ULB-IBC | National Platform bis COVID ULB-IBC | Ricardo De Mendonça et al |
| EPI_ISL_5391251 | KU Leuven, Rega Institute, Clinical and Epidemiological Virology | KU Leuven, Rega Institute, Clinical and Epidemiological Virology | Tony Wawina-Bokalanga et al |
| EPI_ISL_4037272 | SYNLAB | GIGA Medical Genomics | Keith Durkin et al |
| EPI_ISL_7121195 | Botswana Harvard HIV Reference Laboratory | Botswana Harvard HIV Reference Laboratory | Sikhulile Moyo et al |
| EPI_ISL_5647428 | INSIDE DIAGNOSTICOS | Instituto Butantan | Dimas Tadeu Covas et al |
| EPI_ISL_1380949 | KU Leuven, Rega Institute, Clinical and Epidemiological Virology | KU Leuven, Rega Institute, Clinical and Epidemiological Virology | Tony Wawina-Bokalanga et al |
| EPI_ISL_6173267 | Laboratorio Central de Saude Publica do Estado da Paraiba (LACEN/PB) | Laboratory of Respiratory Viruses and Measles, Oswaldo Cruz Institute, FIOCRUZ | Paola Resende et al |

| | | | |
|---|---|---|---|
| EPI_ISL_6752026 | Botswana Harvard HIV Reference Laboratory | Botswana Harvard HIV Reference Laboratory | Sikhulile Moyo et al |
| EPI_ISL_3134658 | LACEN/PE | WallauLab on behalf of Fiocruz COVID-19 Genomic Surveillance Network | Marcelo Henrique dos Santos Paiva et al |
| EPI_ISL_420359 | KU Leuven, Clinical and Epidemiological Virology | KU Leuven, Clinical and Epidemiological Virology | Joan Marti-Carreras et al |
| EPI_ISL_4211050 | Platform BIS UZA/UAntwerpen | Labo Klinische Biologie, UZA | Marie Le Mercier et al |
| EPI_ISL_2661786 | Laboratorio Central de Saude Publica do Estado do Rio Grande do Sul (LACEN-RS) | Laboratory of Respiratory Viruses and Measles, Oswaldo Cruz Institute, FIOCRUZ | Paola Resende et al |
| EPI_ISL_2493600 | CENTRO MEDICO SOCIAL COMUNITARIO JANUARIO TEODORO DE SOUZA | Instituto Butantan | Dimas Tadeu Covas et al |
| EPI_ISL_4080875 | Laboratorio Central de Saude Publica do Estado de Goias (LACEN/GO) | Laboratory of Respiratory Viruses and Measles, Oswaldo Cruz Institute, FIOCRUZ | Paola Resende et al |
| EPI_ISL_6262940 | SYNLAB | GIGA Medical Genomics | Keith Durkin et al |
| EPI_ISL_5366241 | KU Leuven, Rega Institute, Clinical and Epidemiological Virology | KU Leuven, Rega Institute, Clinical and Epidemiological Virology | Tony Wawina-Bokalanga et al |
| EPI_ISL_6116175 | Platform BIS UZA/UAntwerpen | Labo Klinische Biologie, UZA | Jasmine Coppens et al |
| EPI_ISL_734689 | UZ Leuven, National Reference Laboratory for Coronaviruses, Laboratory Medicine, Leuven, Belgium | KU Leuven, Rega Institute, Clinical and Epidemiological Virology | Tony Wawina-Bokalanga et al |
| EPI_ISL_6231642 | Dutch COVID-19 response team | National Institute for Public Health and the Environment (RIVM) | Adam Meijer et al |
| EPI_ISL_2386160 | Botswana Harvard HIV Reference Laboratory | Botswana Harvard HIV Reference Laboratory | Sikhulile Moyo et al |
| EPI_ISL_6474171 | HOSPITAL MUNICIPAL JOSE GONCALVES ROSA | ACME Lab, Oswaldo Cruz Foundation, FIOCRUZ/CE | Fabio Miyajima et al |
| EPI_ISL_2422215 | Platform BIS UZA/UAntwerpen | Labo Klinische Biologie, UZA | Marie Le Mercier et al |
| EPI_ISL_935304 | KU Leuven, Rega Institute, Clinical and Epidemiological Virology | KU Leuven, Rega Institute, Clinical and Epidemiological Virology | Tony Wawina-Bokalanga et al |
| EPI_ISL_6774088 | Botswana Harvard HIV Reference Laboratory | Botswana Harvard HIV Reference Laboratory | Sikhulile Moyo et al |
| EPI_ISL_2864588 | KU Leuven, Rega Institute, Clinical and Epidemiological Virology | KU Leuven, Rega Institute, Clinical and Epidemiological Virology | Tony Wawina-Bokalanga et al |
| EPI_ISL_524470 | Hospital do Servidor Público Estadual Francisco Morato de Oliveira | Instituto Adolfo Lutz, Interdiciplinary Procedures Center, Strategic Laboratory | Claudio Tavares Sacchi et al |
| EPI_ISL_1826694 | SYNLAB | GIGA Medical Genomics | Keith Durkin et al |
| EPI_ISL_2385647 | Laboratorio Central Noel Nutels | Bioinformatics Laboratory / LNCC | Luiz G P de Almeida et al |
| EPI_ISL_2777670 | Laboratório Central de Saúde Pública do Amazonas - LACEN-AM | Laboratorio de Ecologia de Doencas Transmissiveis na Amazonia, Instituto Leonidas e Maria Deane - Fiocruz Amazonia | Valdinete Nascimento et al |

| | | | |
|---|---|---|---|
| EPI_ISL_4271111 | Laboratory of Respiratory Viruses and Measles, Oswaldo Cruz Institute, FIOCRUZ | Laboratory of Respiratory Viruses and Measles, Oswaldo Cruz Institute, FIOCRUZ | Paola Resende et al |
| EPI_ISL_1381280 | Platform BIS UZA/UAntwerpen | UAntwerp, Laboratory of Medical Microbiology | Basil Britto Xavier et al |
| EPI_ISL_3912280 | UBS DO BAIRRO SAO JOSE | Analytical Competence Molecular Epidemiology Lab/ACME, Oswaldo Cruz Foundation, Ceara (FIOCRUZ CE) | Fabio Miyajima et al |
| EPI_ISL_6913917 | Grupo CR Diagnosticos | Instituto Adolfo Lutz Strategic Laboratory | Claudio Tavares Sacchi et al |
| EPI_ISL_5414661 | KU Leuven, Rega Institute, Clinical and Epidemiological Virology | KU Leuven, Rega Institute, Clinical and Epidemiological Virology | Tony Wawina-Bokalanga et al |
| EPI_ISL_2345438 | AMBULATORIO DE ESPECIALIDADE V E MOGI MIRIM | Instituto Butantan / ESALQ-Piracicaba | Dimas Tadeu Covas et al |
| EPI_ISL_6774085 | Botswana Harvard HIV Reference Laboratory | Botswana Harvard HIV Reference Laboratory | Sikhulile Moyo et al |
| EPI_ISL_6774083 | Botswana Harvard HIV Reference Laboratory | Botswana Harvard HIV Reference Laboratory | Sikhulile Moyo et al |
| EPI_ISL_6722289 | LACEN - Laboratório Central de Saúde Pública de Pernambuco | Evandro Chagas Institute | Mirleide Cordeiro dos Santos et al |
| EPI_ISL_2964887 | KU Leuven, Rega Institute, Clinical and Epidemiological Virology | KU Leuven, Rega Institute, Clinical and Epidemiological Virology | Tony Wawina-Bokalanga et al |
| EPI_ISL_5645889 | INSIDE DIAGNOSTICOS | Instituto Butantan | Dimas Tadeu Covas et al |
| EPI_ISL_3190382 | Laboratorio Central de Saude Publica do Estado do Parana (LACEN/PR) | Laboratory of Respiratory Viruses and Measles, Oswaldo Cruz Institute, FIOCRUZ | Paola Resende et al |
| EPI_ISL_1966350 | CENTRO DE SAUDE DR RENATO DE CARVALHO RIBEIRO ANGATUBA | Instituto Butantan / Mendelics | Instituto Butantan: Dimas Tadeu Covas et al |
| EPI_ISL_3235279 | Laboratorio Central de Saude Publica do Estado do Rio Grande do Sul (LACEN-RS) | Laboratory of Respiratory Viruses and Measles, Oswaldo Cruz Institute, FIOCRUZ | Paola Resende et al |
| EPI_ISL_1103562 | Algemeen Medisch Laboratorium | UAntwerp, Laboratory of Medical Microbiology | Basil Britto Xavier et al |
| EPI_ISL_6640916 | Botswana Harvard HIV Reference Laboratory | Botswana Harvard HIV Reference Laboratory | Sikhulile Moyo et al |
| EPI_ISL_6794907 | KU Leuven, Rega Institute, Clinical and Epidemiological Virology | KU Leuven, Rega Institute, Clinical and Epidemiological Virology | Tony Wawina-Bokalanga et al |
| EPI_ISL_6670244 | Botswana Harvard HIV Reference Laboratory | Botswana Harvard HIV Reference Laboratory | Sikhulile Moyo et al |
| EPI_ISL_1966110 | CENTRO DE SAUDE II DR GABRIEL MESQUITA VARGEM GDE DO SUL | Instituto Butantan / Mendelics | Instituto Butantan: Dimas Tadeu Covas et al |

| | | | |
|---|---|---|---|
| EPI_ISL_3536210 | HEMOCE CENTRO DE HEMATOLOGIA E HEMOTERAPIA DO CEARA | Analytical Competence Molecular Epidemiology Lab/ACME, Oswaldo Cruz Foundation, Ceara (FIOCRUZ CE) | Fabio Miyajima et al |
| EPI_ISL_7021517 | National Platform bis COVID ULB-IBC | National Platform bis COVID ULB-IBC | Ricardo De Mendonça et al |
| EPI_ISL_5915327 | AMALAB/FACISA/UFRN | WallauLab on behalf of Fiocruz COVID-19 Genomic Surveillance Network | Allan Roberto Dias Nunes et al |
| EPI_ISL_560385 | National Health Laboratory | Botswana Institute for Technology Research and Innovation | Kefentse Arnold Tumedi et al |
| EPI_ISL_5658370 | Laboratório Translacional, Centro de Pesquisa em Medicina Tropical, CEPEM/RO | Laboratório de Ecologia de Doenças Transmissíveis na Amazônia, Instituto Leônidas e Maria Deane - Fiocruz Amazônia | Dhelio Pereira et al |
| EPI_ISL_1967303 | SECRETARIA MUNICIPAL DE SAUDE | Instituto Butantan / Mendelics | Instituto Butantan: Dimas Tadeu Covas et al |
| EPI_ISL_7121080 | Diagnofirm Medical Laboratories | Botswana Harvard HIV Reference Laboratory | Sikhulile Moyo et al |
| EPI_ISL_1967058 | CENTRO DE SAUDE ESF IV ZONA RURAL DOMINGOS DE S SJRIOPARDO | Instituto Butantan / ESALQ-USP (Piracicaba) | Instituto Butantan: Dimas Tadeu Covas et al |
| EPI_ISL_925479 | Department of Clinical Microbiology | GIGA Medical Genomics | Keith Durkin et al |
| EPI_ISL_1963373 | Platform BIS UZA/UAntwerpen | Labo Klinische Biologie, UZA | Jasmine Coppens et al |
| EPI_ISL_5795541 | LACEN ALAGOAS | Instituto Butantan | Dimas Tadeu Covas et al |
| EPI_ISL_3536538 | HLAGYN - Laboratorio de Imunologia de Transplantes de Goias | HLAGYN - Laboratorio de Imunologia de Transplantes de Goias | Fernando Antonio Vinhal dos Santos et al |
| EPI_ISL_7121204 | Botswana Harvard HIV Reference Laboratory | Botswana Harvard HIV Reference Laboratory | Sikhulile Moyo et al |
| EPI_ISL_5414814 | KU Leuven, Rega Institute, Clinical and Epidemiological Virology | KU Leuven, Rega Institute, Clinical and Epidemiological Virology | Tony Wawina-Bokalanga et al |
| EPI_ISL_6969264 | Laboratorio Central Noel Nutels | Bioinformatics Laboratory / LNCC | Luiz G P de Almeida et al |
| EPI_ISL_4414916 | Laboratorio Central de Saude Publica do Estado de Santa Catarina (LACEN/SC) | Laboratory of Respiratory Viruses and Measles, Oswaldo Cruz Institute, FIOCRUZ | Paola Resende et al |
| EPI_ISL_7042669 | Department of Clinical Microbiology | GIGA Medical Genomics | Keith Durkin et al |
| EPI_ISL_7063764 | National Platform bis UMONS/Jolimont | National Platform bis UMONS/Jolimont | Guillaume Bayon-Vicente et al |
| EPI_ISL_4489274 | SYNLAB | GIGA Medical Genomics | Keith Durkin et al |
| EPI_ISL_6691407 | Unidade de apoio ao diagnostico da COVID - UNADIG | Bioinformatics Laboratory / LNCC | Luiz G P de Almeida et al |
| EPI_ISL_5390557 | Laboratorio Central de Saude Publica do Estado do Espirito Santo (LACEN/ES) | Laboratory of Respiratory Viruses and Measles, Oswaldo Cruz Institute, FIOCRUZ | Paola Resende et al |
| EPI_ISL_4414459 | DASA | DASA | Rodrigo Guarischi et al |

| | | | |
|---|---|---|---|
| EPI_ISL_6909001 | Laboratório Central de Saúde Pública - LACEN/BA | Plataforma de Vigilancia Molecular (PVM) - FIOCRUZ/BA | Laise de Moraes et al |
| EPI_ISL_2272674 | Platform BIS UZA/UAntwerpen | Labo Klinische Biologie, UZA | Marie Le Mercier et al |
| EPI_ISL_6929785 | University of Liège COVID-19 testing center | GIGA Medical Genomics | Keith Durkin et al |
| EPI_ISL_1379938 | Platform BIS UZA/UAntwerpen | UZA, Clinical Biology | Marie Le Mercier et al |
| EPI_ISL_2557361 | Lboratorio Central de Saude Publica do Estado do Parana (LACEN/PR) | Laboratory of Respiratory Viruses and Measles, Oswaldo Cruz Institute, FIOCRUZ | Paola Resende et al |
| EPI_ISL_3320585 | Platform BIS UZA/UAntwerpen | Labo Klinische Biologie, UZA | Marie Le Mercier et al |
| EPI_ISL_6512795 | PZU “Eurofarm - Centar Poliklinika” | University of Sarajevo, Veterinary Faculty, Laboratory for Molecular Diagnostic and Research | Goletic S et al |
| EPI_ISL_4601910 | National Platform bis UMONS/Jolimont | National Platform bis UMONS/Jolimont | Guillaume Bayon-Vicente et al |
| EPI_ISL_2833362 | KU Leuven, Rega Institute, Clinical and Epidemiological Virology | KU Leuven, Rega Institute, Clinical and Epidemiological Virology | Tony Wawina-Bokalanga et al |
| EPI_ISL_7121205 | Botswana Harvard HIV Reference Laboratory | Botswana Harvard HIV Reference Laboratory | Sikhulile Moyo et al |
| EPI_ISL_3827923 | Laboratorio Central de Saude Publica do Estado do Espirito Santo (LACEN/ES) | Laboratory of Respiratory Viruses and Measles, Oswaldo Cruz Institute, FIOCRUZ | Paola Resende et al |
| EPI_ISL_6640917 | Botswana Harvard HIV Reference Laboratory | Botswana Harvard HIV Reference Laboratory | Sikhulile Moyo et al |
| EPI_ISL_3245263 | Laboratorio Central Noel Nutels | Bioinformatics Laboratory / LNCC | Luiz G P de Almeida et al |
| EPI_ISL_4880316 | Laboratorio Central de Saude Publica do Estado do Maranhao (LACEN-MA) | Laboratory of Respiratory Viruses and Measles, Oswaldo Cruz Institute, FIOCRUZ | Paola Resende et al |
| EPI_ISL_6940676 | Laboratorio Central de Saude Publica do Estado do Rio Grande do Sul (LACEN-RS) | Laboratory of Respiratory Viruses and Measles, Oswaldo Cruz Institute, FIOCRUZ | Paola Resende et al |
| EPI_ISL_912411 | KU Leuven, Rega Institute, Clinical and Epidemiological Virology | KU Leuven, Rega Institute, Clinical and Epidemiological Virology | Tony Wawina-Bokalanga et al |
| EPI_ISL_1533727 | Hospital Universitario da USP Sao Paulo | Instituto Adolfo Lutz, Interdiciplinary Procedures Center, Strategic Laboratory | Claudio Tavares Sacchi et al |
| EPI_ISL_6774091 | Botswana Harvard HIV Reference Laboratory | Botswana Harvard HIV Reference Laboratory | Sikhulile Moyo et al |
| EPI_ISL_6721336 | LACEN - Laboratório Central de Saúde Pública do Rio Grande do Norte | Evandro Chagas Institute | Mirleide Cordeiro dos Santos et al |
| EPI_ISL_6774081 | Botswana Harvard HIV Reference Laboratory | Botswana Harvard HIV Reference Laboratory | Sikhulile Moyo et al |
| EPI_ISL_1671976 | SYNLAB | GIGA Medical Genomics | Keith Durkin et al |
| EPI_ISL_6992912 | DASA | DASA | Rodrigo Guarischi et al |
| EPI_ISL_2863845 | Laboratorio Central de Saude Publica do Estado do Para (LACEN/PA) | Laboratory of Respiratory Viruses and Measles, Oswaldo Cruz Institute, FIOCRUZ | Paola Resende et al |

| | | | |
|---|---|---|---|
| EPI_ISL_541380 | Laboratório Central de Saúde Pública do Estado de Sergipe (LACEN-SE) | Laboratory of Respiratory Viruses and Measles, Oswaldo Cruz Institute, FIOCRUZ | Paola Resende et al |
| EPI_ISL_641580 | Department of Clinical Microbiology | GIGA Medical Genomics | Keith Durkin et al |
| EPI_ISL_2726019 | KU Leuven, Rega Institute, Clinical and Epidemiological Virology | KU Leuven, Rega Institute, Clinical and Epidemiological Virology | Tony Wawina-Bokalanga et al |
| EPI_ISL_6774089 | Botswana Harvard HIV Reference Laboratory | Botswana Harvard HIV Reference Laboratory | Sikhulile Moyo et al |
| EPI_ISL_6901961 | LATE - Laboratório de Técnicas Especiais - Hospital Israelita Albert Einstein | LATE - Laboratório de Técnicas Especiais - Hospital Israelita Albert Einstein | Deyvid Amgarten et al |
| EPI_ISL_4358275 | KU Leuven, Rega Institute, Clinical and Epidemiological Virology | KU Leuven, Rega Institute, Clinical and Epidemiological Virology | Tony Wawina-Bokalanga et al |
| EPI_ISL_6989250 | KU Leuven, Rega Institute, Clinical and Epidemiological Virology | KU Leuven, Rega Institute, Clinical and Epidemiological Virology | Tony Wawina-Bokalanga et al |
| EPI_ISL_5425393 | KU Leuven, Rega Institute, Clinical and Epidemiological Virology | KU Leuven, Rega Institute, Clinical and Epidemiological Virology | Tony Wawina-Bokalanga et al |
| EPI_ISL_1967301 | SECRETARIA MUNICIPAL DE SAUDE | Instituto Butantan / Mendelics | Instituto Butantan: Dimas Tadeu Covas et al |
| EPI_ISL_6774090 | Botswana Harvard HIV Reference Laboratory | Botswana Harvard HIV Reference Laboratory | Sikhulile Moyo et al |
| EPI_ISL_5336765 | SYNLAB | GIGA Medical Genomics | Keith Durkin et al |
| EPI_ISL_5676841 | NUCLEO SAUDE III N HABITACIONAL PRESIDENTE GEISEL BAURU | Instituto Butantan | Dimas Tadeu Covas et al |
| EPI_ISL_4561958 | Edmonton Provincial Lab | Public Health Agency of Canada (PHAC) National Microbiology Laboratory | Buss et al |
| EPI_ISL_3601474 | British Columbia Centre For Disease Control | BCCDC Public Health Laboratory | Prystajecky Natalie et al |
| EPI_ISL_4731417 | National Center of Infectious and Parasitic Diseases | National Center of Infectious and Parasitic Diseases | Alexiev et al |
| EPI_ISL_854541 | Alberta Precision Labs (APL) | Alberta Precision Labs (APL) | Gordon P et al |
| EPI_ISL_6949582 | Edmonton Provincial Lab | Alberta Precision Labs (APL) | Buss et al |
| EPI_ISL_6664360 | Edmonton Provincial Lab | Public Health Agency of Canada (PHAC) National Microbiology Laboratory | Buss et al |
| EPI_ISL_806113 | Alberta Precision Labs (APL) | Alberta Precision Labs (APL) | Gordon P et al |
| EPI_ISL_3116712 | Edmonton Provincial Lab | Public Health Agency of Canada (PHAC) National Microbiology Laboratory | Buss et al |
| EPI_ISL_4278654 | British Columbia Centre For Disease Control | BCCDC Public Health Laboratory | Prystajecky Natalie et al |
| EPI_ISL_3599290 | British Columbia Centre For Disease Control | BCCDC Public Health Laboratory | Prystajecky Natalie et al |
| EPI_ISL_4430823 | British Columbia Centre For Disease Control | BCCDC Public Health Laboratory | Prystajecky Natalie et al |

| | | | |
|---|---|---|---|
| EPI_ISL_4546701 | National Center of Infectious and Parasitic Diseases | National Center of Infectious and Parasitic Diseases | Alexiev et al |
| EPI_ISL_4420871 | British Columbia Centre For Disease Control | BCCDC Public Health Laboratory | Prystajecky Natalie et al |
| EPI_ISL_7123666 | Edmonton Provincial Lab | Alberta Precision Labs (APL) | Buss et al |
| EPI_ISL_4747188 | HOSPITAL MUNICIPAL DR JOSE DE CARVALHO FLORENCE | Instituto Butantan | Dimas Tadeu Covas et al |
| EPI_ISL_4558884 | Edmonton Provincial Lab | Public Health Agency of Canada (PHAC) National Microbiology Laboratory | Buss et al |
| EPI_ISL_5651283 | SECRETARIA MUNICIPAL DE SAUDE DE ANDRADINA | Instituto Butantan | Dimas Tadeu Covas et al |
| EPI_ISL_5651110 | SANTA CASA DE MISERICORDIA DE UBATUBA | Instituto Butantan | Dimas Tadeu Covas et al |
| EPI_ISL_4278536 | British Columbia Centre For Disease Control | BCCDC Public Health Laboratory | Prystajecky Natalie et al |
| EPI_ISL_4829061 | British Columbia Centre For Disease Control | BCCDC Public Health Laboratory | Prystajecky Natalie et al |
| EPI_ISL_4421460 | British Columbia Centre For Disease Control | BCCDC Public Health Laboratory | Prystajecky Natalie et al |
| EPI_ISL_4434028 | British Columbia Centre For Disease Control | BCCDC Public Health Laboratory | Prystajecky Natalie et al |
| EPI_ISL_3115853 | Edmonton Provincial Lab | Public Health Agency of Canada (PHAC) National Microbiology Laboratory | Buss et al |
| EPI_ISL_4255114 | Centre Hospitalier Universitaire Yalgado OUEDRAOGO | Africa Centre for Excellence for Genomics of Infectious Diseases (ACEGID), Redeemer’s University | Olawoye et al |
| EPI_ISL_4419885 | British Columbia Centre For Disease Control | BCCDC Public Health Laboratory | Prystajecky Natalie et al |
| EPI_ISL_3483015 | British Columbia Centre For Disease Control | BCCDC Public Health Laboratory | Prystajecky Natalie et al |
| EPI_ISL_3485991 | British Columbia Centre For Disease Control | BCCDC Public Health Laboratory | Prystajecky Natalie et al |
| EPI_ISL_6897079 | SECRETARIA DE SAUDE | Instituto Butantan | Dimas Tadeu Covas et al |
| EPI_ISL_4831120 | British Columbia Centre For Disease Control | BCCDC Public Health Laboratory | Prystajecky Natalie et al |
| EPI_ISL_4427975 | British Columbia Centre For Disease Control | BCCDC Public Health Laboratory | Prystajecky Natalie et al |
| EPI_ISL_5801946 | Sms Secretaria Municipal De Saude De Boituva | Instituto Butantan | Dimas Tadeu Covas et al |
| EPI_ISL_2167559 | Alberta Precision Labs (APL) | Public Health Agency of Canada (PHAC) National Microbiology Laboratory | Buss et al |
| EPI_ISL_5630787 | Edmonton Provincial Lab | Public Health Agency of Canada (PHAC) National Microbiology Laboratory | Buss et al |
| EPI_ISL_5417428 | National Center of Infectious and Parasitic Diseases | National Center of Infectious and Parasitic Diseases | Alexiev et al |
| EPI_ISL_5679078 | UBS CELESTINO CARNIELO | Instituto Butantan | Dimas Tadeu Covas et al |
| EPI_ISL_4469605 | BIOFAST | Instituto Butantan | Dimas Tadeu Covas et al |

| | | | |
|---|---|---|---|
| EPI_ISL_6101749 | National Center of Infectious and Parasitic Diseases | National Center of Infectious and Parasitic Diseases | Alexiev et al |
| EPI_ISL_2163526 | Alberta Precision Labs (APL) | Public Health Agency of Canada (PHAC) National Microbiology Laboratory | Buss et al |
| EPI_ISL_4830766 | British Columbia Centre For Disease Control | BCCDC Public Health Laboratory | Prystajecky Natalie et al |
| EPI_ISL_805720 | Alberta Precision Labs (APL) | Alberta Precision Labs (APL) | Gordon P et al |
| EPI_ISL_5021305 | CENTRO DE SAUDE DE MARACAI | Instituto Butantan | Dimas Tadeu Covas et al |
| EPI_ISL_4053812 | Edmonton Provincial Lab | Public Health Agency of Canada (PHAC) National Microbiology Laboratory | Buss et al |
| EPI_ISL_4833122 | British Columbia Centre For Disease Control | BCCDC Public Health Laboratory | Prystajecky Natalie et al |
| EPI_ISL_6570154 | INSIDE DIAGNOSTICOS | Instituto Butantan | Dimas Tadeu Covas et al |
| EPI_ISL_2180134 | National Center of Infectious and Parasitic Diseases | National Center of Infectious and Parasitic Diseases | Alexiev et al |
| EPI_ISL_5688704 | INSIDE DIAGNOSTICOS | Instituto Butantan | Dimas Tadeu Covas et al |
| EPI_ISL_4420914 | British Columbia Centre For Disease Control | BCCDC Public Health Laboratory | Prystajecky Natalie et al |
| EPI_ISL_4277584 | British Columbia Centre For Disease Control | BCCDC Public Health Laboratory | Prystajecky Natalie et al |
| EPI_ISL_4430654 | British Columbia Centre For Disease Control | BCCDC Public Health Laboratory | Prystajecky Natalie et al |
| EPI_ISL_5801467 | Psf Do Parque Dos Pinheiros | Instituto Butantan | Dimas Tadeu Covas et al |
| EPI_ISL_4560818 | Edmonton Provincial Lab | Public Health Agency of Canada (PHAC) National Microbiology Laboratory | Buss et al |
| EPI_ISL_2478599 | Edmonton Provincial Lab | Public Health Agency of Canada (PHAC) National Microbiology Laboratory | Buss et al |
| EPI_ISL_3596578 | British Columbia Centre For Disease Control | BCCDC Public Health Laboratory | Prystajecky Natalie et al |
| EPI_ISL_3483359 | British Columbia Centre For Disease Control | BCCDC Public Health Laboratory | Prystajecky Natalie et al |
| EPI_ISL_4374554 | AFIP | Instituto Butantan | Dimas Tadeu Covas et al |
| EPI_ISL_6660684 | Edmonton Provincial Lab | Public Health Agency of Canada (PHAC) National Microbiology Laboratory | Buss et al |
| EPI_ISL_3602521 | British Columbia Centre For Disease Control | BCCDC Public Health Laboratory | Prystajecky Natalie et al |
| EPI_ISL_2169431 | Alberta Precision Labs (APL) | Public Health Agency of Canada (PHAC) National Microbiology Laboratory | Buss et al |
| EPI_ISL_3598491 | British Columbia Centre For Disease Control | BCCDC Public Health Laboratory | Prystajecky Natalie et al |
| EPI_ISL_4829036 | British Columbia Centre For Disease Control | BCCDC Public Health Laboratory | Prystajecky Natalie et al |
| EPI_ISL_4433859 | British Columbia Centre For Disease Control | BCCDC Public Health Laboratory | Prystajecky Natalie et al |

| | | | |
|---|---|---|---|
| EPI_ISL_1786365 | National Center of Infectious and Parasitic Diseases | National Center of Infectious and Parasitic Diseases | Alexiev et al |
| EPI_ISL_5668624 | VIGILANCIA SANITARIA E VIG EPIDEMIOLOGICA DE ITAPEVI | Instituto Butantan | Dimas Tadeu Covas et al |
| EPI_ISL_4425971 | British Columbia Centre For Disease Control | BCCDC Public Health Laboratory | Prystajecky Natalie et al |
| EPI_ISL_3482089 | British Columbia Centre For Disease Control | BCCDC Public Health Laboratory | Prystajecky Natalie et al |
| EPI_ISL_3489303 | British Columbia Centre For Disease Control | BCCDC Public Health Laboratory | Prystajecky Natalie et al |
| EPI_ISL_2169650 | Alberta Precision Labs (APL) | Public Health Agency of Canada (PHAC) National Microbiology Laboratory | Buss et al |
| EPI_ISL_2180421 | National Center of Infectious and Parasitic Diseases | National Center of Infectious and Parasitic Diseases | Alexiev et al |
| EPI_ISL_3387387 | Virology Unit, Institut Pasteur du Cambodge | Virology Unit, Institut Pasteur du Cambodge | Leakhena Pum et al |
| EPI_ISL_3599090 | British Columbia Centre For Disease Control | BCCDC Public Health Laboratory | Prystajecky Natalie et al |
| EPI_ISL_4423696 | British Columbia Centre For Disease Control | BCCDC Public Health Laboratory | Prystajecky Natalie et al |
| EPI_ISL_2163934 | Alberta Precision Labs (APL) | Public Health Agency of Canada (PHAC) National Microbiology Laboratory | Buss et al |
| EPI_ISL_7123680 | Edmonton Provincial Lab | Alberta Precision Labs (APL) | Buss et al |
| EPI_ISL_5661686 | UNIDADE REFERENCIAL SUDOESTE | Instituto Butantan | Dimas Tadeu Covas et al |
| EPI_ISL_4833581 | British Columbia Centre For Disease Control | BCCDC Public Health Laboratory | Prystajecky Natalie et al |
| EPI_ISL_6813143 | HOSPITAL MUNICIPAL DE ILHABELA GOV MARIO COVAS JR | Instituto Butantan | Dimas Tadeu Covas et al |
| EPI_ISL_2940873 | Edmonton Provincial Lab | Public Health Agency of Canada (PHAC) National Microbiology Laboratory | Buss et al |
| EPI_ISL_4426263 | British Columbia Centre For Disease Control | BCCDC Public Health Laboratory | Prystajecky Natalie et al |
| EPI_ISL_2941124 | Edmonton Provincial Lab | Public Health Agency of Canada (PHAC) National Microbiology Laboratory | Buss et al |
| EPI_ISL_3116011 | Edmonton Provincial Lab | Public Health Agency of Canada (PHAC) National Microbiology Laboratory | Buss et al |
| EPI_ISL_2169774 | Alberta Precision Labs (APL) | Public Health Agency of Canada (PHAC) National Microbiology Laboratory | Buss et al |
| EPI_ISL_1401330 | National Center of Infectious and Parasitic Diseases | National Center of Infectious and Parasitic Diseases | Alexiev et al |
| EPI_ISL_4168652 | UNIDADE RESPIRATORIA NOVA HORTOLANDIA | Instituto Butantan | Dimas Tadeu Covas et al |
| EPI_ISL_5650453 | ESF NOVA TANABI II | Instituto Butantan | Dimas Tadeu Covas et al |

| | | | |
|---|---|---|---|
| EPI_ISL_5653509 | HOSPITAL DE CAMPANHA COVID 19 CAIEIRAS | Instituto Butantan | Dimas Tadeu Covas et al |
| EPI_ISL_4834455 | British Columbia Centre For Disease Control | BCCDC Public Health Laboratory | Prystajecky Natalie et al |
| EPI_ISL_4829694 | British Columbia Centre For Disease Control | BCCDC Public Health Laboratory | Prystajecky Natalie et al |
| EPI_ISL_2165250 | Alberta Precision Labs (APL) | Public Health Agency of Canada (PHAC) National Microbiology Laboratory | Buss et al |
| EPI_ISL_4054095 | Edmonton Provincial Lab | Public Health Agency of Canada (PHAC) National Microbiology Laboratory | Buss et al |
| EPI_ISL_2170646 | Alberta Precision Labs (APL) | Public Health Agency of Canada (PHAC) National Microbiology Laboratory | Buss et al |
| EPI_ISL_5653515 | HOSP E MATERNIDADE MUNICIPAL N SRA MONTE SERRAT | Instituto Butantan | Dimas Tadeu Covas et al |
| EPI_ISL_2165619 | Alberta Precision Labs (APL) | Public Health Agency of Canada (PHAC) National Microbiology Laboratory | Buss et al |
| EPI_ISL_7123692 | Edmonton Provincial Lab | Alberta Precision Labs (APL) | Buss et al |
| EPI_ISL_5596830 | National Public Health Laboratory, Ministry of Public Health of Cameroon | CERI, Centre for Epidemic Response and Innvoation, Stellenbosch University and KRISP, KZN Research Innovation and Sequencing Platform, UKZN. | Dr Essomba Rene Ghislain et al |
| EPI_ISL_5662980 | BIOFAST | Instituto Butantan | Dimas Tadeu Covas et al |
| EPI_ISL_4277837 | British Columbia Centre For Disease Control | BCCDC Public Health Laboratory | Prystajecky Natalie et al |
| EPI_ISL_2648086 | National Center of Infectious and Parasitic Diseases | National Center of Infectious and Parasitic Diseases | Alexiev et al |
| EPI_ISL_5802363 | Ubs Jardim Caiuby | Instituto Butantan | Dimas Tadeu Covas et al |
| EPI_ISL_2523061 | BCCDC Public Health Laboratory | BCCDC Public Health Laboratory | Prystajecky Natalie et al |
| EPI_ISL_2523826 | BCCDC Public Health Laboratory | BCCDC Public Health Laboratory | Prystajecky Natalie et al |
| EPI_ISL_2653375 | BCCDC Public Health Laboratory | BCCDC Public Health Laboratory | Prystajecky Natalie et al |
| EPI_ISL_579565 | QEII Health Sciences Centre | National Microbiology Laboratory (NML) | Anna Majer et al |
| EPI_ISL_6753566 | British Columbia Centre For Disease Control | BCCDC Public Health Laboratory | Prystajecky Natalie et al |
| EPI_ISL_3487783 | British Columbia Centre For Disease Control | BCCDC Public Health Laboratory | Prystajecky Natalie et al |
| EPI_ISL_4846265 | British Columbia Centre For Disease Control | BCCDC Public Health Laboratory | Prystajecky Natalie et al |
| EPI_ISL_961364 | Toronto Invasive Bacterial Diseases Network | McMaster University | Allison McGeer et al |
| EPI_ISL_6740283 | British Columbia Centre For Disease Control | BCCDC Public Health Laboratory | Prystajecky Natalie et al |

| | | | |
|---|---|---|---|
| EPI_ISL_2530508 | BCCDC Public Health Laboratory | BCCDC Public Health Laboratory | Prystajecky Natalie et al |
| EPI_ISL_2523944 | BCCDC Public Health Laboratory | BCCDC Public Health Laboratory | Prystajecky Natalie et al |
| EPI_ISL_6826714 | Dynacare | National Microbiology Laboratory (NML) | Anna Majer et al |
| EPI_ISL_2527110 | BCCDC Public Health Laboratory | BCCDC Public Health Laboratory | Prystajecky Natalie et al |
| EPI_ISL_6989667 | Dynacare | National Microbiology Laboratory (NML) | Anna Majer et al |
| EPI_ISL_2529272 | BCCDC Public Health Laboratory | BCCDC Public Health Laboratory | Prystajecky Natalie et al |
| EPI_ISL_2655457 | BCCDC Public Health Laboratory | BCCDC Public Health Laboratory | Prystajecky Natalie et al |
| EPI_ISL_3480709 | British Columbia Centre For Disease Control | BCCDC Public Health Laboratory | Prystajecky Natalie et al |
| EPI_ISL_6738297 | British Columbia Centre For Disease Control | BCCDC Public Health Laboratory | Prystajecky Natalie et al |
| EPI_ISL_6826713 | Dynacare | National Microbiology Laboratory (NML) | Anna Majer et al |
| EPI_ISL_2528667 | BCCDC Public Health Laboratory | BCCDC Public Health Laboratory | Prystajecky Natalie et al |
| EPI_ISL_5684385 | British Columbia Centre For Disease Control | BCCDC Public Health Laboratory | Prystajecky Natalie et al |
| EPI_ISL_2524353 | BCCDC Public Health Laboratory | BCCDC Public Health Laboratory | Prystajecky Natalie et al |
| EPI_ISL_6746316 | British Columbia Centre For Disease Control | BCCDC Public Health Laboratory | Prystajecky Natalie et al |
| EPI_ISL_6742971 | British Columbia Centre For Disease Control | BCCDC Public Health Laboratory | Prystajecky Natalie et al |
| EPI_ISL_2720833 | BCCDC Public Health Laboratory | BCCDC Public Health Laboratory | Prystajecky Natalie et al |
| EPI_ISL_7008190 | British Columbia Centre For Disease Control | BCCDC Public Health Laboratory | Prystajecky Natalie et al |
| EPI_ISL_2528215 | BCCDC Public Health Laboratory | BCCDC Public Health Laboratory | Prystajecky Natalie et al |
| EPI_ISL_2656491 | BCCDC Public Health Laboratory | BCCDC Public Health Laboratory | Prystajecky Natalie et al |
| EPI_ISL_4494340 | MB-Cadham Provincial laboratory | National Microbiology Laboratory (NML) | Anna Majer et al |
| EPI_ISL_6738131 | British Columbia Centre For Disease Control | BCCDC Public Health Laboratory | Prystajecky Natalie et al |
| EPI_ISL_975531 | BCCDC Public Health Laboratory | BCCDC Public Health Laboratory | Prystajecky Natalie et al |
| EPI_ISL_1739999 | Ontario's COVID-19 Genomics Rapid Response Coalition | McMaster University | Allison McGeer et al |
| EPI_ISL_974478 | BCCDC Public Health Laboratory | BCCDC Public Health Laboratory | Prystajecky Natalie et al |
| EPI_ISL_6739192 | British Columbia Centre For Disease Control | BCCDC Public Health Laboratory | Prystajecky Natalie et al |
| EPI_ISL_5221046 | British Columbia Centre For Disease Control | BCCDC Public Health Laboratory | Prystajecky Natalie et al |

| | | | |
|---|---|---|---|
| EPI_ISL_6751212 | British Columbia Centre For Disease Control | BCCDC Public Health Laboratory | Prystajecky Natalie et al |
| EPI_ISL_853578 | Toronto Invasive Bacterial Diseases Network | Ontario Institute for Cancer Research | Allison McGeer et al |
| EPI_ISL_6157845 | NB-Hôpital Georges L. Dumont | New Brunswick - Vitalité Health Network | Chacko S. et al |
| EPI_ISL_6725727 | British Columbia Centre For Disease Control | BCCDC Public Health Laboratory | Prystajecky Natalie et al |
| EPI_ISL_968264 | BCCDC Public Health Laboratory | BCCDC Public Health Laboratory | Prystajecky Natalie et al |
| EPI_ISL_6738520 | British Columbia Centre For Disease Control | BCCDC Public Health Laboratory | Prystajecky Natalie et al |
| EPI_ISL_4835853 | British Columbia Centre For Disease Control | BCCDC Public Health Laboratory | Prystajecky Natalie et al |
| EPI_ISL_975156 | BCCDC Public Health Laboratory | BCCDC Public Health Laboratory | Prystajecky Natalie et al |
| EPI_ISL_6740963 | British Columbia Centre For Disease Control | BCCDC Public Health Laboratory | Prystajecky Natalie et al |
| EPI_ISL_6743352 | British Columbia Centre For Disease Control | BCCDC Public Health Laboratory | Prystajecky Natalie et al |
| EPI_ISL_5685621 | British Columbia Centre For Disease Control | BCCDC Public Health Laboratory | Prystajecky Natalie et al |
| EPI_ISL_7001040 | British Columbia Centre For Disease Control | BCCDC Public Health Laboratory | Prystajecky Natalie et al |
| EPI_ISL_5316729 | Shared Hospital Laboratory | Shared Hospital Laboratory | Patryk Aftanas et al |
| EPI_ISL_4846251 | British Columbia Centre For Disease Control | BCCDC Public Health Laboratory | Prystajecky Natalie et al |
| EPI_ISL_4025812 | NB-Hôpital Georges L. Dumont | New Brunswick - Vitalité Health Network | Chacko S. et al |
| EPI_ISL_2719733 | BCCDC Public Health Laboratory | BCCDC Public Health Laboratory | Prystajecky Natalie et al |
| EPI_ISL_5219106 | British Columbia Centre For Disease Control | BCCDC Public Health Laboratory | Prystajecky Natalie et al |
| EPI_ISL_5685552 | British Columbia Centre For Disease Control | BCCDC Public Health Laboratory | Prystajecky Natalie et al |
| EPI_ISL_935791 | Cadham Provincial laboratory | National Microbiology Laboratory (NML) | Anna Majer et al |
| EPI_ISL_4947780 | NB-Hôpital Georges L. Dumont | New Brunswick - Vitalité Health Network | Chacko S. et al |
| EPI_ISL_961576 | Hôpital Georges L. Dumont | National Microbiology Laboratory (NML) | Anna Majer et al |
| EPI_ISL_4834972 | British Columbia Centre For Disease Control | BCCDC Public Health Laboratory | Prystajecky Natalie et al |
| EPI_ISL_591194 | Toronto Invasive Bacterial Diseases Network | McMaster University | Allison McGeer et al |
| EPI_ISL_5224459 | British Columbia Centre For Disease Control | BCCDC Public Health Laboratory | Prystajecky Natalie et al |
| EPI_ISL_2530704 | BCCDC Public Health Laboratory | BCCDC Public Health Laboratory | Prystajecky Natalie et al |
| EPI_ISL_973745 | BCCDC Public Health Laboratory | BCCDC Public Health Laboratory | Prystajecky Natalie et al |

| | | | |
|---|---|---|---|
| EPI_ISL_579662 | QEII Health Sciences Centre | National Microbiology Laboratory (NML) | Anna Majer et al |
| EPI_ISL_4275182 | National Institute of Public Health | State Veterinary Institute Prague | Alexander Nagy et al |
| EPI_ISL_6414029 | Zavod za javno zdravstvo Međimurske županije | Hrvatski zavod za javno zdravstvo | Irena Tabain et al |
| EPI_ISL_6687208 | Shared Hospital Laboratory | Shared Hospital Laboratory | Patryk Aftanas et al |
| EPI_ISL_2821587 | Laboratoire de santé publique du Québec | Laboratoire de santé publique du Québec | Sandrine Moreira et al |
| EPI_ISL_6033882 | Zavod za javno zdravstvo Zagrebačke županije | Hrvatski zavod za javno zdravstvo | Irena Tabain et al |
| EPI_ISL_7015154 | Dynacare | National Microbiology Laboratory (NML) | Anna Majer et al |
| EPI_ISL_3026023 | AREA DE SALUD FORTUNA [San Carlos/ALAJUELA] | Inciensa, Instituto Costarricense de Investigación y Enseñanza en Nutrición y Salud | Francisco Duarte et al |
| EPI_ISL_2674895 | Hrvatski zavod za javno zdravstvo | Hrvatski zavod za javno zdravstvo | Irena Tabain et al |
| EPI_ISL_3320720 | Universidad del Quindio | Instituto Nacional de Salud | Katherine Laiton-Donato et al |
| EPI_ISL_3098329 | Laboratoire de santé publique du Québec | Laboratoire de santé publique du Québec | Sandrine Moreira et al |
| EPI_ISL_5426794 | SK-Roy Romanow Provincial Laboratory | National Microbiology Laboratory (NML) | Anna Majer et al |
| EPI_ISL_2823600 | Laboratoire de santé publique du Québec | Laboratoire de santé publique du Québec | Sandrine Moreira et al |
| EPI_ISL_6454054 | Genetica Molecular and Subdepartamento de Virologia ISP Chile | Instituto de Salud Publica de Chile | Karen Orostica et al |
| EPI_ISL_5530608 | COMPENSAR - LABORATORIO CLINICO LOS COBOS | Instituto Nacional de Salud-Dirección de Investigación en Salud Pública | Katherine Laiton-Donato et al |
| EPI_ISL_5999332 | AREA DE SALUD FORTUNA | Inciensa, Instituto Costarricense de Investigación y Enseñanza en Nutrición y Salud | Francisco Duarte et al |
| EPI_ISL_4900655 | National Institute of Public Health | State Veterinary Institute Prague | Alexander Nagy et al |
| EPI_ISL_5196156 | Hrvatski zavod za javno zdravstvo | Hrvatski zavod za javno zdravstvo | Irena Tabain et al |
| EPI_ISL_3266998 | Ayudas Diagnosticas Sura | Universidad Nacional de Colombia - Laboratorio Genómico One Health | Maria Stella López et al |
| EPI_ISL_2346380 | Institute of Molecular and Translational Medicine / Laboratory of Experimental Medicine, Faculty of Medicine and Dentistry, Palacky University and University Hospital Olomouc | Institute of Molecular and Translational Medicine / Laboratory of Experimental Medicine, Faculty of Medicine and Dentistry, Palacky University | Rastislav Slavkovský et al |
| EPI_ISL_4891921 | Gencore - Universidad de los Andes | Gencore - Universidad de los Andes | Luisa Sacristan et al |
| EPI_ISL_4703618 | Servicios Medicos Olimpus | Laboratorio de Biologia Molecular, Universidad Cooperativa de Colombia, Santa Marta | Andrew S. Muñoz-Gamba et al |

| | | | |
|---|---|---|---|
| EPI_ISL_5843305 | SK-Roy Romanow Provincial Laboratory | Saskatchewan - Roy Romanow Provincial Laboratory (RRPL) | Ryan McDonald et al |
| EPI_ISL_5055447 | UNIVERSIDAD DE ANTIOQUIA - LIME | Instituto Nacional de Salud-Dirección de Investigación en Salud Pública | Katherine Laiton-Donato et al |
| EPI_ISL_5012937 | Hrvatski zavod za javno zdravstvo | Hrvatski zavod za javno zdravstvo | Irena Tabain et al |
| EPI_ISL_5924555 | SangLab Karlovy Vary | Department of Microbiology, University Hospital in Pilsen | Katerina Chudejova et al |
| EPI_ISL_6686668 | Shared Hospital Laboratory | Shared Hospital Laboratory | Patryk Aftanas et al |
| EPI_ISL_5196981 | SK-Roy Romanow Provincial Laboratory | Saskatchewan - Roy Romanow Provincial Laboratory (RRPL) | Ryan McDonald et al |
| EPI_ISL_5010614 | Zavod za javno zdravstvo Osječko-Baranjske županije | Hrvatski zavod za javno zdravstvo | Irena Tabain et al |
| EPI_ISL_2727002 | National Institute of Public Health | National Institute of Public Health | Helena Jirincova et al |
| EPI_ISL_5603748 | Compensar Calle 26 | LSPSDS | Johana Hernandez Toloza et al |
| EPI_ISL_6703485 | Lab Ayudas Diagnosticas SURA-sede Cali | CIAT, Laboratorio de Virologia | Ana M. Leiva et al |
| EPI_ISL_1969869 | Laboratoire de santé publique du Québec | Laboratoire de santé publique du Québec | Sandrine Moreira et al |
| EPI_ISL_4585018 | Institute of Molecular and Translational Medicine / Laboratory of Experimental Medicine, Faculty of Medicine and Dentistry, Palacky University | Institute of Molecular and Translational Medicine / Laboratory of Experimental Medicine, Faculty of Medicine and Dentistry, Palacky University | Rastislav Slavkovský et al |
| EPI_ISL_536183 | Centre hospitalier Anna-Laberge | Laboratoire de santé publique du Québec | Sandrine Moreira et al |
| EPI_ISL_6453616 | Genetica Molecular and Subdepartamento de Virologia ISP Chile | Instituto de Salud Publica de Chile | Karen Orostica et al |
| EPI_ISL_3143439 | Laboratoire de santé publique du Québec | Laboratoire de santé publique du Québec | Sandrine Moreira et al |
| EPI_ISL_4655169 | Genetica Molecular and Subdepartamento de Virologia ISP Chile | Instituto de Salud Publica de Chile | Karen Orostica et al |
| EPI_ISL_5209150 | Dutch COVID-19 response team | National Institute for Public Health and the Environment (RIVM) | Adam Meijer et al |
| EPI_ISL_2970791 | Laboratoire de santé publique du Québec | Laboratoire de santé publique du Québec | Sandrine Moreira et al |
| EPI_ISL_2883647 | SK-Roy Romanow Provincial Laboratory | National Microbiology Laboratory (NML) | Anna Majer et al |
| EPI_ISL_4635597 | Genetica Molecular and Subdepartamento de Virologia ISP Chile | Instituto de Salud Publica de Chile | Karen Orostica et al |
| EPI_ISL_6862005 | Regional Hospital Liberec | Regional Hospital Liberec | Tomas Zajic et al |
| EPI_ISL_2007616 | Laboratoire de santé publique du Québec | Laboratoire de santé publique du Québec | Sandrine Moreira et al |
| EPI_ISL_2824069 | Laboratoire de santé publique du Québec | Laboratoire de santé publique du Québec | Sandrine Moreira et al |

| | | | |
|---|---|---|---|
| EPI_ISL_4221422 | Laboratoire de santé publique du Québec | Laboratoire de santé publique du Québec | Sandrine Moreira et al |
| EPI_ISL_3117775 | Oddělení klinické mikrobiologie | Institute of Medical Microbiology and Virology, University Hospital Carl Gustav Carus, TU Dresden | Marlena Stadtmüller et al |
| EPI_ISL_5834936 | Nemocnice Jihlava | University Hospital Brno, CMBG | Bezdicek Matej et al |
| EPI_ISL_3144428 | Laboratoire de santé publique du Québec | Laboratoire de santé publique du Québec | Sandrine Moreira et al |
| EPI_ISL_2467190 | Clinica INDISA | "Facultad de Ciencias de la Vida, UNAB" | "Claudio Meneses et al |
| EPI_ISL_1365768 | Laboratoire de santé publique du Québec | Laboratoire de santé publique du Québec | Sandrine Moreira et al |
| EPI_ISL_2586909 | SK-Roy Romanow Provincial Laboratory | National Microbiology Laboratory (NML) | Anna Majer et al |
| EPI_ISL_6756545 | SK-Roy Romanow Provincial Laboratory | National Microbiology Laboratory (NML) | Anna Majer et al |
| EPI_ISL_6423450 | Laboratoire de santé publique du Québec | Laboratoire de santé publique du Québec | Sandrine Moreira et al |
| EPI_ISL_2889335 | SK-Roy Romanow Provincial Laboratory | National Microbiology Laboratory (NML) | Anna Majer et al |
| EPI_ISL_6756629 | SK-Roy Romanow Provincial Laboratory | National Microbiology Laboratory (NML) | Anna Majer et al |
| EPI_ISL_2293803 | Laboratoire de santé publique du Québec | Laboratoire de santé publique du Québec | Sandrine Moreira et al |
| EPI_ISL_2386024 | Laboratoire de santé publique du Québec | Laboratoire de santé publique du Québec | Sandrine Moreira et al |
| EPI_ISL_1716639 | Hospital Universitario de Gran Canaria Dr. Negrín | SeqCOVID-SPAIN consortium/IBV(CSIC) | M. Carmen Pérez González et al |
| EPI_ISL_3255724 | SK-Roy Romanow Provincial Laboratory | National Microbiology Laboratory (NML) | Anna Majer et al |
| EPI_ISL_2896147 | Genetica Molecular and Subdepartamento de Virologia ISP Chile | Instituto de Salud Publica de Chile | Karen Orostica et al |
| EPI_ISL_2970566 | Laboratoire de santé publique du Québec | Laboratoire de santé publique du Québec | Sandrine Moreira et al |
| EPI_ISL_5196789 | SK-Roy Romanow Provincial Laboratory | Saskatchewan - Roy Romanow Provincial Laboratory (RRPL) | Ryan McDonald et al |
| EPI_ISL_4632555 | Genetica Molecular and Subdepartamento de Virologia ISP Chile | Instituto de Salud Publica de Chile | Karen Orostica et al |
| EPI_ISL_2467238 | Clinica INDISA | "Facultad de Ciencias de la Vida, UNAB" | "Claudio Meneses et al |
| EPI_ISL_2499765 | Laboratoire de santé publique du Québec | Laboratoire de santé publique du Québec | Sandrine Moreira et al |
| EPI_ISL_5315301 | Shared Hospital Laboratory | Shared Hospital Laboratory | Patryk Aftanas et al |
| EPI_ISL_3332294 | Nemocnice Jihlava | University Hospital Brno, CMBG | Matej Bezdicek et al |
| EPI_ISL_825762 | Laboratoire de santé publique du Québec | Laboratoire de santé publique du Québec | Sandrine Moreira et al |
| EPI_ISL_6687857 | Shared Hospital Laboratory | Shared Hospital Laboratory | Patryk Aftanas et al |
| EPI_ISL_2693570 | Klinika za infektivne bolesti "Dr. Fran Mihaljević" | Hrvatski zavod za javno zdravstvo | Irena Tabain et al |

| | | | |
|---|---|---|---|
| EPI_ISL_3741803 | Zavod za javno zdravstvo Zagrebačke županije | Hrvatski zavod za javno zdravstvo | Irena Tabain et al |
| EPI_ISL_3185437 | Genetica Molecular and Subdepartamento de Virologia ISP Chile | Instituto de Salud Publica de Chile | Karen Orostica et al |
| EPI_ISL_2970339 | Laboratoire de santé publique du Québec | Laboratoire de santé publique du Québec | Sandrine Moreira et al |
| EPI_ISL_2346349 | Institute of Molecular and Translational Medicine / Laboratory of Experimental Medicine, Faculty of Medicine and Dentistry, Palacky University and University Hospital Olomouc | Institute of Molecular and Translational Medicine / Laboratory of Experimental Medicine, Faculty of Medicine and Dentistry, Palacky University | Rastislav Slavkovský et al |
| EPI_ISL_6832215 | Fakultní nemocnice u sv. Anny v Brně | Fakultní nemocnice u sv. Anny v Brně | Milada Dvořáčková. Jan Vrbský et al |
| EPI_ISL_6846326 | Pardubicka nemocnice | University Hospital Hradec Kralove | Marketa Gancarcikova et al |
| EPI_ISL_4926410 | Laboratorio de salud publica departamental del valle de cauca | CIAT, Laboratorio de Virologia | Ana M. Leiva et al |
| EPI_ISL_2388556 | Laboratoire de santé publique du Québec | Laboratoire de santé publique du Québec | Sandrine Moreira et al |
| EPI_ISL_4221481 | Laboratoire de santé publique du Québec | Laboratoire de santé publique du Québec | Sandrine Moreira et al |
| EPI_ISL_2565595 | SK-Roy Romanow Provincial Laboratory | National Microbiology Laboratory (NML) | Anna Majer et al |
| EPI_ISL_2821425 | Laboratoire de santé publique du Québec | Laboratoire de santé publique du Québec | Sandrine Moreira et al |
| EPI_ISL_6073499 | SK-Roy Romanow Provincial Laboratory | National Microbiology Laboratory (NML) | Anna Majer et al |
| EPI_ISL_4221098 | Laboratoire de santé publique du Québec | Laboratoire de santé publique du Québec | Sandrine Moreira et al |
| EPI_ISL_2920020 | Oddělení klinické mikrobiologie | Institute of Medical Microbiology and Virology, University Hospital Carl Gustav Carus, TU Dresden | Marlena Stadtmüller et al |
| EPI_ISL_6413036 | Hrvatski zavod za javno zdravstvo | Hrvatski zavod za javno zdravstvo | Irena Tabain et al |
| EPI_ISL_2824773 | Laboratoire de santé publique du Québec | Laboratoire de santé publique du Québec | Sandrine Moreira et al |
| EPI_ISL_1315058 | National Institute of Public Health | National Institute of Public Health | Helena Jirincova et al |
| EPI_ISL_3508457 | Laboratoire de santé publique du Québec | Laboratoire de santé publique du Québec | Sandrine Moreira et al |
| EPI_ISL_2970327 | Laboratoire de santé publique du Québec | Laboratoire de santé publique du Québec | Sandrine Moreira et al |
| EPI_ISL_3298335 | AREA DE SALUD CARTAGO | Inciensa, Instituto Costarricense de Investigación y Enseñanza en Nutrición y Salud | Francisco Duarte et al |
| EPI_ISL_526273 | Unity Health Toronto | Ontario Institute for Cancer Research | Ramzi Fattouh et al |
| EPI_ISL_791498 | The Hosptial for Sick Children | The Hospital for Sick Children | Aaron Campigotto et al |

| | | | |
|---|---|---|---|
| EPI_ISL_5544303 | Genetica Molecular and Subdepartamento de Virologia ISP Chile | Instituto de Salud Publica de Chile | Karen Orostica et al |
| EPI_ISL_6826328 | Oblastni nemocnice Trutnov a.s. | University Hospital Hradec Kralove | Marketa Gancarcikova et al |
| EPI_ISL_5055359 | UNIVERSIDAD DE ANTIOQUIA | Instituto Nacional de Salud-Dirección de Investigación en Salud Pública | Katherine Laiton-Donato et al |
| EPI_ISL_6604677 | Institute of Molecular and Translational Medicine / Laboratory of Experimental Medicine, Faculty of Medicine and Dentistry, Palacky University | Institute of Molecular and Translational Medicine / Laboratory of Experimental Medicine, Faculty of Medicine and Dentistry, Palacky University | Rastislav Slavkovský et al |
| EPI_ISL_6458753 | Genetica Molecular and Subdepartamento de Virologia ISP Chile | Instituto de Salud Publica de Chile | Karen Orostica et al |
| EPI_ISL_3709743 | Department of Bacteria, Parasites and Fungi, Statens Serum Institut, Copenhagen, Denmark | Statens Serum Institut Bioinformatics and Microbial Genomics | Danish Covid-19 Genome Consortium et al |
| EPI_ISL_4943085 | Department of Bacteria, Parasites and Fungi, Statens Serum Institut, Copenhagen, Denmark | Statens Serum Institut Bioinformatics and Microbial Genomics | Danish Covid-19 Genome Consortium et al |
| EPI_ISL_2963817 | Department of Bacteria, Parasites and Fungi, Statens Serum Institut, Copenhagen, Denmark | Statens Serum Institut Bioinformatics and Microbial Genomics | Danish Covid-19 Genome Consortium et al |
| EPI_ISL_682635 | Department of Virus and Microbiological Special Diagnostics, Statens Serum Institut, Copenhagen, Denmark | Albertsen Lab, Department of Chemistry and Bioscience, Aalborg University, Denmark | Danish Covid-19 Genome Consortium et al |
| EPI_ISL_2668860 | Department of Bacteria, Parasites and Fungi, Statens Serum Institut, Copenhagen, Denmark | Statens Serum Institut Bioinformatics and Microbial Genomics | Danish Covid-19 Genome Consortium et al |
| EPI_ISL_2992823 | Department of Bacteria, Parasites and Fungi, Statens Serum Institut, Copenhagen, Denmark | Statens Serum Institut Bioinformatics and Microbial Genomics | Danish Covid-19 Genome Consortium et al |
| EPI_ISL_3280022 | Department of Bacteria, Parasites and Fungi, Statens Serum Institut, Copenhagen, Denmark | Statens Serum Institut Bioinformatics and Microbial Genomics | Danish Covid-19 Genome Consortium et al |
| EPI_ISL_2838205 | Department of Bacteria, Parasites and Fungi, Statens Serum Institut, Copenhagen, Denmark | Statens Serum Institut Bioinformatics and Microbial Genomics | Danish Covid-19 Genome Consortium et al |

| | | | |
|---|---|---|---|
| EPI_ISL_712459 | Department of Virus and Microbiological Special Diagnostics, Statens Serum Institut, Copenhagen, Denmark | Albertsen Lab, Department of Chemistry and Bioscience, Aalborg University, Denmark | Danish Covid-19 Genome Consortium et al |
| EPI_ISL_3103337 | Department of Bacteria, Parasites and Fungi, Statens Serum Institut, Copenhagen, Denmark | Statens Serum Institut Bioinformatics and Microbial Genomics | Danish Covid-19 Genome Consortium et al |
| EPI_ISL_3040638 | Department of Bacteria, Parasites and Fungi, Statens Serum Institut, Copenhagen, Denmark | Statens Serum Institut Bioinformatics and Microbial Genomics | Danish Covid-19 Genome Consortium et al |
| EPI_ISL_3450517 | Department of Bacteria, Parasites and Fungi, Statens Serum Institut, Copenhagen, Denmark | Statens Serum Institut Bioinformatics and Microbial Genomics | Danish Covid-19 Genome Consortium et al |
| EPI_ISL_3228804 | Department of Bacteria, Parasites and Fungi, Statens Serum Institut, Copenhagen, Denmark | Statens Serum Institut Bioinformatics and Microbial Genomics | Danish Covid-19 Genome Consortium et al |
| EPI_ISL_3917792 | Department of Bacteria, Parasites and Fungi, Statens Serum Institut, Copenhagen, Denmark | Statens Serum Institut Bioinformatics and Microbial Genomics | Danish Covid-19 Genome Consortium et al |
| EPI_ISL_2680855 | Department of Bacteria, Parasites and Fungi, Statens Serum Institut, Copenhagen, Denmark | Statens Serum Institut Bioinformatics and Microbial Genomics | Danish Covid-19 Genome Consortium et al |
| EPI_ISL_3917099 | Department of Bacteria, Parasites and Fungi, Statens Serum Institut, Copenhagen, Denmark | Statens Serum Institut Bioinformatics and Microbial Genomics | Danish Covid-19 Genome Consortium et al |
| EPI_ISL_711472 | Department of Virus and Microbiological Special Diagnostics, Statens Serum Institut, Copenhagen, Denmark | Albertsen Lab, Department of Chemistry and Bioscience, Aalborg University, Denmark | Danish Covid-19 Genome Consortium et al |
| EPI_ISL_3241570 | Department of Bacteria, Parasites and Fungi, Statens Serum Institut, Copenhagen, Denmark | Statens Serum Institut Bioinformatics and Microbial Genomics | Danish Covid-19 Genome Consortium et al |
| EPI_ISL_3839388 | Department of Bacteria, Parasites and Fungi, Statens Serum Institut, Copenhagen, Denmark | Statens Serum Institut Bioinformatics and Microbial Genomics | Danish Covid-19 Genome Consortium et al |
| EPI_ISL_3241127 | Department of Bacteria, Parasites and Fungi, Statens Serum Institut, Copenhagen, Denmark | Statens Serum Institut Bioinformatics and Microbial Genomics | Danish Covid-19 Genome Consortium et al |

| | | | |
|---|---|---|---|
| EPI_ISL_4105474 | Department of Bacteria, Parasites and Fungi, Statens Serum Institut, Copenhagen, Denmark | Statens Serum Institut Bioinformatics and Microbial Genomics | Danish Covid-19 Genome Consortium et al |
| EPI_ISL_3708661 | Department of Bacteria, Parasites and Fungi, Statens Serum Institut, Copenhagen, Denmark | Statens Serum Institut Bioinformatics and Microbial Genomics | Danish Covid-19 Genome Consortium et al |
| EPI_ISL_4626357 | Department of Bacteria, Parasites and Fungi, Statens Serum Institut, Copenhagen, Denmark | Statens Serum Institut Bioinformatics and Microbial Genomics | Danish Covid-19 Genome Consortium et al |
| EPI_ISL_3566895 | Department of Bacteria, Parasites and Fungi, Statens Serum Institut, Copenhagen, Denmark | Statens Serum Institut Bioinformatics and Microbial Genomics | Danish Covid-19 Genome Consortium et al |
| EPI_ISL_3530558 | Department of Bacteria, Parasites and Fungi, Statens Serum Institut, Copenhagen, Denmark | Statens Serum Institut Bioinformatics and Microbial Genomics | Danish Covid-19 Genome Consortium et al |
| EPI_ISL_2684538 | Statens Serum Institut Bioinformatics and Microbial Genomics | Statens Serum Institut Bioinformatics and Microbial Genomics | Danish Covid-19 Genome Consortium et al |
| EPI_ISL_750107 | Department of Virus and Microbiological Special Diagnostics, Statens Serum Institut, Copenhagen, Denmark | Albertsen Lab, Department of Chemistry and Bioscience, Aalborg University, Denmark | Danish Covid-19 Genome Consortium et al |
| EPI_ISL_4170402 | Department of Bacteria, Parasites and Fungi, Statens Serum Institut, Copenhagen, Denmark | Statens Serum Institut Bioinformatics and Microbial Genomics | Danish Covid-19 Genome Consortium et al |
| EPI_ISL_2432500 | Department of Bacteria, Parasites and Fungi, Statens Serum Institut, Copenhagen, Denmark | Statens Serum Institut Bioinformatics and Microbial Genomics | Danish Covid-19 Genome Consortium et al |
| EPI_ISL_4232997 | Department of Bacteria, Parasites and Fungi, Statens Serum Institut, Copenhagen, Denmark | Statens Serum Institut Bioinformatics and Microbial Genomics | Danish Covid-19 Genome Consortium et al |
| EPI_ISL_2992191 | Department of Bacteria, Parasites and Fungi, Statens Serum Institut, Copenhagen, Denmark | Statens Serum Institut Bioinformatics and Microbial Genomics | Danish Covid-19 Genome Consortium et al |
| EPI_ISL_3725120 | Department of Bacteria, Parasites and Fungi, Statens Serum Institut, Copenhagen, Denmark | Statens Serum Institut Bioinformatics and Microbial Genomics | Danish Covid-19 Genome Consortium et al |
| EPI_ISL_4942831 | Department of Bacteria, Parasites and Fungi, Statens Serum Institut, Copenhagen, Denmark | Statens Serum Institut Bioinformatics and Microbial Genomics | Danish Covid-19 Genome Consortium et al |

| | | | |
|---|---|---|---|
| EPI_ISL_2863476 | Department of Bacteria, Parasites and Fungi, Statens Serum Institut, Copenhagen, Denmark | Statens Serum Institut Bioinformatics and Microbial Genomics | Danish Covid-19 Genome Consortium et al |
| EPI_ISL_2431586 | Department of Bacteria, Parasites and Fungi, Statens Serum Institut, Copenhagen, Denmark | Statens Serum Institut Bioinformatics and Microbial Genomics | Danish Covid-19 Genome Consortium et al |
| EPI_ISL_4257205 | Department of Bacteria, Parasites and Fungi, Statens Serum Institut, Copenhagen, Denmark | Statens Serum Institut Bioinformatics and Microbial Genomics | Danish Covid-19 Genome Consortium et al |
| EPI_ISL_3917688 | Department of Bacteria, Parasites and Fungi, Statens Serum Institut, Copenhagen, Denmark | Statens Serum Institut Bioinformatics and Microbial Genomics | Danish Covid-19 Genome Consortium et al |
| EPI_ISL_669883 | Department of Virus and Microbiological Special Diagnostics, Statens Serum Institut, Copenhagen, Denmark | Albertsen Lab, Department of Chemistry and Bioscience, Aalborg University, Denmark | Danish Covid-19 Genome Consortium et al |
| EPI_ISL_3709822 | Department of Bacteria, Parasites and Fungi, Statens Serum Institut, Copenhagen, Denmark | Statens Serum Institut Bioinformatics and Microbial Genomics | Danish Covid-19 Genome Consortium et al |
| EPI_ISL_3530740 | Department of Bacteria, Parasites and Fungi, Statens Serum Institut, Copenhagen, Denmark | Statens Serum Institut Bioinformatics and Microbial Genomics | Danish Covid-19 Genome Consortium et al |
| EPI_ISL_4884175 | Department of Bacteria, Parasites and Fungi, Statens Serum Institut, Copenhagen, Denmark | Statens Serum Institut Bioinformatics and Microbial Genomics | Danish Covid-19 Genome Consortium et al |
| EPI_ISL_3104275 | Department of Bacteria, Parasites and Fungi, Statens Serum Institut, Copenhagen, Denmark | Statens Serum Institut Bioinformatics and Microbial Genomics | Danish Covid-19 Genome Consortium et al |
| EPI_ISL_5592657 | Department of Bacteria, Parasites and Fungi, Statens Serum Institut, Copenhagen, Denmark | Statens Serum Institut Bioinformatics and Microbial Genomics | Danish Covid-19 Genome Consortium et al |
| EPI_ISL_4448492 | Molekylær Medicinsk Afdeling, Aarhus University Hospital, Aarhus, Denmark | Statens Serum Institut Bioinformatics and Microbial Genomics | Danish Covid-19 Genome Consortium et al |
| EPI_ISL_4625922 | Department of Clinical Microbiology, Odense University Hospital, Odense, Denmark | Statens Serum Institut Bioinformatics and Microbial Genomics | Danish Covid-19 Genome Consortium et al |
| EPI_ISL_3709744 | Department of Bacteria, Parasites and Fungi, Statens Serum Institut, Copenhagen, Denmark | Statens Serum Institut Bioinformatics and Microbial Genomics | Danish Covid-19 Genome Consortium et al |

| | | | |
|---|---|---|---|
| EPI_ISL_3806526 | Department of Bacteria, Parasites and Fungi, Statens Serum Institut, Copenhagen, Denmark | Statens Serum Institut Bioinformatics and Microbial Genomics | Danish Covid-19 Genome Consortium et al |
| EPI_ISL_2298562 | Department of Virus and Microbiological Special Diagnostics, Statens Serum Institut, Copenhagen, Denmark | Aalborg University | Danish Covid-19 Genome Consortium et al |
| EPI_ISL_3724758 | Department of Bacteria, Parasites and Fungi, Statens Serum Institut, Copenhagen, Denmark | Statens Serum Institut Bioinformatics and Microbial Genomics | Danish Covid-19 Genome Consortium et al |
| EPI_ISL_3724912 | Department of Bacteria, Parasites and Fungi, Statens Serum Institut, Copenhagen, Denmark | Statens Serum Institut Bioinformatics and Microbial Genomics | Danish Covid-19 Genome Consortium et al |
| EPI_ISL_2683970 | Department of Bacteria, Parasites and Fungi, Statens Serum Institut, Copenhagen, Denmark | Statens Serum Institut Bioinformatics and Microbial Genomics | Danish Covid-19 Genome Consortium et al |
| EPI_ISL_3240032 | Department of Bacteria, Parasites and Fungi, Statens Serum Institut, Copenhagen, Denmark | Statens Serum Institut Bioinformatics and Microbial Genomics | Danish Covid-19 Genome Consortium et al |
| EPI_ISL_3530670 | Department of Bacteria, Parasites and Fungi, Statens Serum Institut, Copenhagen, Denmark | Statens Serum Institut Bioinformatics and Microbial Genomics | Danish Covid-19 Genome Consortium et al |
| EPI_ISL_713942 | Department of Virus and Microbiological Special Diagnostics, Statens Serum Institut, Copenhagen, Denmark | Albertsen Lab, Department of Chemistry and Bioscience, Aalborg University, Denmark | Danish Covid-19 Genome Consortium et al |
| EPI_ISL_2854567 | Department of Bacteria, Parasites and Fungi, Statens Serum Institut, Copenhagen, Denmark | Statens Serum Institut Bioinformatics and Microbial Genomics | Danish Covid-19 Genome Consortium et al |
| EPI_ISL_3664958 | Department of Bacteria, Parasites and Fungi, Statens Serum Institut, Copenhagen, Denmark | Statens Serum Institut Bioinformatics and Microbial Genomics | Danish Covid-19 Genome Consortium et al |
| EPI_ISL_3418400 | Department of Bacteria, Parasites and Fungi, Statens Serum Institut, Copenhagen, Denmark | Statens Serum Institut Bioinformatics and Microbial Genomics | Danish Covid-19 Genome Consortium et al |
| EPI_ISL_3949502 | Department of Bacteria, Parasites and Fungi, Statens Serum Institut, Copenhagen, Denmark | Statens Serum Institut Bioinformatics and Microbial Genomics | Danish Covid-19 Genome Consortium et al |

| | | | |
|---|---|---|---|
| EPI_ISL_3418218 | Department of Bacteria, Parasites and Fungi, Statens Serum Institut, Copenhagen, Denmark | Statens Serum Institut Bioinformatics and Microbial Genomics | Danish Covid-19 Genome Consortium et al |
| EPI_ISL_4886766 | Department of Bacteria, Parasites and Fungi, Statens Serum Institut, Copenhagen, Denmark | Statens Serum Institut Bioinformatics and Microbial Genomics | Danish Covid-19 Genome Consortium et al |
| EPI_ISL_2933089 | Department of Bacteria, Parasites and Fungi, Statens Serum Institut, Copenhagen, Denmark | Statens Serum Institut Bioinformatics and Microbial Genomics | Danish Covid-19 Genome Consortium et al |
| EPI_ISL_3074811 | Department of Bacteria, Parasites and Fungi, Statens Serum Institut, Copenhagen, Denmark | Statens Serum Institut Bioinformatics and Microbial Genomics | Danish Covid-19 Genome Consortium et al |
| EPI_ISL_711540 | Department of Virus and Microbiological Special Diagnostics, Statens Serum Institut, Copenhagen, Denmark | Albertsen Lab, Department of Chemistry and Bioscience, Aalborg University, Denmark | Danish Covid-19 Genome Consortium et al |
| EPI_ISL_3708111 | Department of Bacteria, Parasites and Fungi, Statens Serum Institut, Copenhagen, Denmark | Statens Serum Institut Bioinformatics and Microbial Genomics | Danish Covid-19 Genome Consortium et al |
| EPI_ISL_2684321 | Statens Serum Institut Bioinformatics and Microbial Genomics | Statens Serum Institut Bioinformatics and Microbial Genomics | Danish Covid-19 Genome Consortium et al |
| EPI_ISL_3141707 | Department of Bacteria, Parasites and Fungi, Statens Serum Institut, Copenhagen, Denmark | Statens Serum Institut Bioinformatics and Microbial Genomics | Danish Covid-19 Genome Consortium et al |
| EPI_ISL_3449959 | Department of Bacteria, Parasites and Fungi, Statens Serum Institut, Copenhagen, Denmark | Statens Serum Institut Bioinformatics and Microbial Genomics | Danish Covid-19 Genome Consortium et al |
| EPI_ISL_3053615 | Department of Bacteria, Parasites and Fungi, Statens Serum Institut, Copenhagen, Denmark | Statens Serum Institut Bioinformatics and Microbial Genomics | Danish Covid-19 Genome Consortium et al |
| EPI_ISL_668515 | Department of Virus and Microbiological Special Diagnostics, Statens Serum Institut, Copenhagen, Denmark | Albertsen Lab, Department of Chemistry and Bioscience, Aalborg University, Denmark | Danish Covid-19 Genome Consortium et al |
| EPI_ISL_711690 | Department of Virus and Microbiological Special Diagnostics, Statens Serum Institut, Copenhagen, Denmark | Albertsen Lab, Department of Chemistry and Bioscience, Aalborg University, Denmark | Danish Covid-19 Genome Consortium et al |

| | | | |
|---|---|---|---|
| EPI_ISL_4258319 | Department of Bacteria, Parasites and Fungi, Statens Serum Institut, Copenhagen, Denmark | Statens Serum Institut Bioinformatics and Microbial Genomics | Danish Covid-19 Genome Consortium et al |
| EPI_ISL_3053480 | Department of Bacteria, Parasites and Fungi, Statens Serum Institut, Copenhagen, Denmark | Statens Serum Institut Bioinformatics and Microbial Genomics | Danish Covid-19 Genome Consortium et al |
| EPI_ISL_4257337 | Department of Bacteria, Parasites and Fungi, Statens Serum Institut, Copenhagen, Denmark | Statens Serum Institut Bioinformatics and Microbial Genomics | Danish Covid-19 Genome Consortium et al |
| EPI_ISL_4050385 | Department of Bacteria, Parasites and Fungi, Statens Serum Institut, Copenhagen, Denmark | Statens Serum Institut Bioinformatics and Microbial Genomics | Danish Covid-19 Genome Consortium et al |
| EPI_ISL_2417279 | Department of Bacteria, Parasites and Fungi, Statens Serum Institut, Copenhagen, Denmark | Statens Serum Institut Bioinformatics and Microbial Genomics | Danish Covid-19 Genome Consortium et al |
| EPI_ISL_2670279 | Department of Bacteria, Parasites and Fungi, Statens Serum Institut, Copenhagen, Denmark | Statens Serum Institut Bioinformatics and Microbial Genomics | Danish Covid-19 Genome Consortium et al |
| EPI_ISL_2992171 | Department of Bacteria, Parasites and Fungi, Statens Serum Institut, Copenhagen, Denmark | Statens Serum Institut Bioinformatics and Microbial Genomics | Danish Covid-19 Genome Consortium et al |
| EPI_ISL_2301246 | Department of Virus and Microbiological Special Diagnostics, Statens Serum Institut, Copenhagen, Denmark | Aalborg University | Danish Covid-19 Genome Consortium et al |
| EPI_ISL_4978062 | Department of Bacteria, Parasites and Fungi, Statens Serum Institut, Copenhagen, Denmark | Statens Serum Institut Bioinformatics and Microbial Genomics | Danish Covid-19 Genome Consortium et al |
| EPI_ISL_3464267 | Department of Bacteria, Parasites and Fungi, Statens Serum Institut, Copenhagen, Denmark | Statens Serum Institut Bioinformatics and Microbial Genomics | Danish Covid-19 Genome Consortium et al |
| EPI_ISL_3903946 | Department of Bacteria, Parasites and Fungi, Statens Serum Institut, Copenhagen, Denmark | Statens Serum Institut Bioinformatics and Microbial Genomics | Danish Covid-19 Genome Consortium et al |
| EPI_ISL_3339765 | Department of Bacteria, Parasites and Fungi, Statens Serum Institut, Copenhagen, Denmark | Statens Serum Institut Bioinformatics and Microbial Genomics | Danish Covid-19 Genome Consortium et al |

| | | | |
|---|---|---|---|
| EPI_ISL_4448972 | Department of Bacteria, Parasites and Fungi, Statens Serum Institut, Copenhagen, Denmark | Statens Serum Institut Bioinformatics and Microbial Genomics | Danish Covid-19 Genome Consortium et al |
| EPI_ISL_682541 | Department of Virus and Microbiological Special Diagnostics, Statens Serum Institut, Copenhagen, Denmark | Albertsen Lab, Department of Chemistry and Bioscience, Aalborg University, Denmark | Danish Covid-19 Genome Consortium et al |
| EPI_ISL_5261352 | Department of Bacteria, Parasites and Fungi, Statens Serum Institut, Copenhagen, Denmark | Statens Serum Institut Bioinformatics and Microbial Genomics | Danish Covid-19 Genome Consortium et al |
| EPI_ISL_6099843 | Department of Bacteria, Parasites and Fungi, Statens Serum Institut, Copenhagen, Denmark | Statens Serum Institut Bioinformatics and Microbial Genomics | Danish Covid-19 Genome Consortium et al |
| EPI_ISL_6681529 | Department of Bacteria, Parasites and Fungi, Statens Serum Institut, Copenhagen, Denmark | Statens Serum Institut Bioinformatics and Microbial Genomics | Danish Covid-19 Genome Consortium et al |
| EPI_ISL_1025504 | Department of Virus and Microbiological Special Diagnostics, Statens Serum Institut, Copenhagen, Denmark | Aalborg University | Danish Covid-19 Genome Consortium et al |
| EPI_ISL_748113 | Department of Virus and Microbiological Special Diagnostics, Statens Serum Institut, Copenhagen, Denmark | Albertsen Lab, Department of Chemistry and Bioscience, Aalborg University, Denmark | Danish Covid-19 Genome Consortium et al |
| EPI_ISL_7063588 | Department of Bacteria, Parasites and Fungi, Statens Serum Institut, Copenhagen, Denmark | Statens Serum Institut Bioinformatics and Microbial Genomics | Danish Covid-19 Genome Consortium et al |
| EPI_ISL_928786 | Department of Virus and Microbiological Special Diagnostics, Statens Serum Institut, Copenhagen, Denmark | Aalborg University | Danish Covid-19 Genome Consortium et al |
| EPI_ISL_5883371 | Department of Bacteria, Parasites and Fungi, Statens Serum Institut, Copenhagen, Denmark | Statens Serum Institut Bioinformatics and Microbial Genomics | Danish Covid-19 Genome Consortium et al |
| EPI_ISL_6968136 | Department of Bacteria, Parasites and Fungi, Statens Serum Institut, Copenhagen, Denmark | Statens Serum Institut Bioinformatics and Microbial Genomics | Danish Covid-19 Genome Consortium et al |
| EPI_ISL_6830965 | Department of Bacteria, Parasites and Fungi, Statens Serum Institut, Copenhagen, Denmark | Statens Serum Institut Bioinformatics and Microbial Genomics | Danish Covid-19 Genome Consortium et al |

| | | | |
|---|---|---|---|
| EPI_ISL_6506077 | Department of Bacteria, Parasites and Fungi, Statens Serum Institut, Copenhagen, Denmark | Statens Serum Institut Bioinformatics and Microbial Genomics | Danish Covid-19 Genome Consortium et al |
| EPI_ISL_930446 | Department of Virus and Microbiological Special Diagnostics, Statens Serum Institut, Copenhagen, Denmark | Aalborg University | Danish Covid-19 Genome Consortium et al |
| EPI_ISL_6879899 | Department of Bacteria, Parasites and Fungi, Statens Serum Institut, Copenhagen, Denmark | Statens Serum Institut Bioinformatics and Microbial Genomics | Danish Covid-19 Genome Consortium et al |
| EPI_ISL_817612 | Department of Virus and Microbiological Special Diagnostics, Statens Serum Institut, Copenhagen, Denmark | Albertsen Lab, Department of Chemistry and Bioscience, Aalborg University, Denmark | Danish Covid-19 Genome Consortium et al |
| EPI_ISL_5883800 | Department of Bacteria, Parasites and Fungi, Statens Serum Institut, Copenhagen, Denmark | Statens Serum Institut Bioinformatics and Microbial Genomics | Danish Covid-19 Genome Consortium et al |
| EPI_ISL_1023598 | Department of Virus and Microbiological Special Diagnostics, Statens Serum Institut, Copenhagen, Denmark | Aalborg University | Danish Covid-19 Genome Consortium et al |
| EPI_ISL_6609060 | Department of Bacteria, Parasites and Fungi, Statens Serum Institut, Copenhagen, Denmark | Statens Serum Institut Bioinformatics and Microbial Genomics | Danish Covid-19 Genome Consortium et al |
| EPI_ISL_6438145 | Department of Bacteria, Parasites and Fungi, Statens Serum Institut, Copenhagen, Denmark | Statens Serum Institut Bioinformatics and Microbial Genomics | Danish Covid-19 Genome Consortium et al |
| EPI_ISL_1125372 | Department of Virus and Microbiological Special Diagnostics, Statens Serum Institut, Copenhagen, Denmark | Aalborg University | Danish Covid-19 Genome Consortium et al |
| EPI_ISL_6678712 | Department of Bacteria, Parasites and Fungi, Statens Serum Institut, Copenhagen, Denmark | Statens Serum Institut Bioinformatics and Microbial Genomics | Danish Covid-19 Genome Consortium et al |
| EPI_ISL_5882920 | Department of Bacteria, Parasites and Fungi, Statens Serum Institut, Copenhagen, Denmark | Statens Serum Institut Bioinformatics and Microbial Genomics | Danish Covid-19 Genome Consortium et al |
| EPI_ISL_6934612 | Department of Bacteria, Parasites and Fungi, Statens Serum Institut, Copenhagen, Denmark | Statens Serum Institut Bioinformatics and Microbial Genomics | Danish Covid-19 Genome Consortium et al |

| | | | |
|---|---|---|---|
| EPI_ISL_1876437 | Department of Virus and Microbiological Special Diagnostics, Statens Serum Institut, Copenhagen, Denmark | Aalborg University | Danish Covid-19 Genome Consortium et al |
| EPI_ISL_1124513 | Department of Virus and Microbiological Special Diagnostics, Statens Serum Institut, Copenhagen, Denmark | Aalborg University | Danish Covid-19 Genome Consortium et al |
| EPI_ISL_1861936 | Department of Virus and Microbiological Special Diagnostics, Statens Serum Institut, Copenhagen, Denmark | Aalborg University | Danish Covid-19 Genome Consortium et al |
| EPI_ISL_6966398 | Department of Bacteria, Parasites and Fungi, Statens Serum Institut, Copenhagen, Denmark | Statens Serum Institut Bioinformatics and Microbial Genomics | Danish Covid-19 Genome Consortium et al |
| EPI_ISL_855719 | Department of Virus and Microbiological Special Diagnostics, Statens Serum Institut, Copenhagen, Denmark | Aalborg University | Danish Covid-19 Genome Consortium et al |
| EPI_ISL_928156 | Department of Virus and Microbiological Special Diagnostics, Statens Serum Institut, Copenhagen, Denmark | Aalborg University | Danish Covid-19 Genome Consortium et al |
| EPI_ISL_928821 | Department of Virus and Microbiological Special Diagnostics, Statens Serum Institut, Copenhagen, Denmark | Aalborg University | Danish Covid-19 Genome Consortium et al |
| EPI_ISL_973289 | Department of Virus and Microbiological Special Diagnostics, Statens Serum Institut, Copenhagen, Denmark | Aalborg University | Danish Covid-19 Genome Consortium et al |
| EPI_ISL_620295 | Department of Virus and Microbiological Special Diagnostics, Statens Serum Institut, Denmark | Albertsen lab, Department of Chemistry and Bioscience, Aalborg University, Denmark | Danish Covid-19 Genome Consortia et al |
| EPI_ISL_819061 | Department of Virus and Microbiological Special Diagnostics, Statens Serum Institut, Copenhagen, Denmark | Albertsen Lab, Department of Chemistry and Bioscience, Aalborg University, Denmark | Danish Covid-19 Genome Consortium et al |
| EPI_ISL_6849318 | Department of Bacteria, Parasites and Fungi, Statens Serum Institut, Copenhagen, Denmark | Statens Serum Institut Bioinformatics and Microbial Genomics | Danish Covid-19 Genome Consortium et al |

| | | | |
|---|---|---|---|
| EPI_ISL_1869845 | Department of Virus and Microbiological Special Diagnostics, Statens Serum Institut, Copenhagen, Denmark | Aalborg University | Danish Covid-19 Genome Consortium et al |
| EPI_ISL_619124 | Department of Virus and Microbiological Special Diagnostics, Statens Serum Institut, Denmark | Albertsen lab, Department of Chemistry and Bioscience, Aalborg University, Denmark | Danish Covid-19 Genome Consortia et al |
| EPI_ISL_817919 | Department of Virus and Microbiological Special Diagnostics, Statens Serum Institut, Copenhagen, Denmark | Albertsen Lab, Department of Chemistry and Bioscience, Aalborg University, Denmark | Danish Covid-19 Genome Consortium et al |
| EPI_ISL_1875413 | Department of Clinical Microbiology and Center for Genomic Medicine, Rigshospitalet, Copenhagen, Denmark | Aalborg University | Danish Covid-19 Genome Consortium et al |
| EPI_ISL_844002 | Department of Virus and Microbiological Special Diagnostics, Statens Serum Institut, Copenhagen, Denmark | Albertsen Lab, Department of Chemistry and Bioscience, Aalborg University, Denmark | Danish Covid-19 Genome Consortium et al |
| EPI_ISL_6848425 | Department of Bacteria, Parasites and Fungi, Statens Serum Institut, Copenhagen, Denmark | Statens Serum Institut Bioinformatics and Microbial Genomics | Danish Covid-19 Genome Consortium et al |
| EPI_ISL_1861136 | Department of Virus and Microbiological Special Diagnostics, Statens Serum Institut, Copenhagen, Denmark | Aalborg University | Danish Covid-19 Genome Consortium et al |
| EPI_ISL_6242940 | Department of Bacteria, Parasites and Fungi, Statens Serum Institut, Copenhagen, Denmark | Statens Serum Institut Bioinformatics and Microbial Genomics | Danish Covid-19 Genome Consortium et al |
| EPI_ISL_815523 | Department of Virus and Microbiological Special Diagnostics, Statens Serum Institut, Copenhagen, Denmark | Albertsen Lab, Department of Chemistry and Bioscience, Aalborg University, Denmark | Danish Covid-19 Genome Consortium et al |
| EPI_ISL_6239241 | Department of Bacteria, Parasites and Fungi, Statens Serum Institut, Copenhagen, Denmark | Statens Serum Institut Bioinformatics and Microbial Genomics | Danish Covid-19 Genome Consortium et al |
| EPI_ISL_6680389 | Department of Bacteria, Parasites and Fungi, Statens Serum Institut, Copenhagen, Denmark | Statens Serum Institut Bioinformatics and Microbial Genomics | Danish Covid-19 Genome Consortium et al |

| | | | |
|---|---|---|---|
| EPI_ISL_6243345 | Department of Bacteria, Parasites and Fungi, Statens Serum Institut, Copenhagen, Denmark | Statens Serum Institut Bioinformatics and Microbial Genomics | Danish Covid-19 Genome Consortium et al |
| EPI_ISL_6848500 | Department of Bacteria, Parasites and Fungi, Statens Serum Institut, Copenhagen, Denmark | Statens Serum Institut Bioinformatics and Microbial Genomics | Danish Covid-19 Genome Consortium et al |
| EPI_ISL_6878354 | Department of Bacteria, Parasites and Fungi, Statens Serum Institut, Copenhagen, Denmark | Statens Serum Institut Bioinformatics and Microbial Genomics | Danish Covid-19 Genome Consortium et al |
| EPI_ISL_1878054 | Department of Virus and Microbiological Special Diagnostics, Statens Serum Institut, Copenhagen, Denmark | Aalborg University | Danish Covid-19 Genome Consortium et al |
| EPI_ISL_1874278 | Department of Virus and Microbiological Special Diagnostics, Statens Serum Institut, Copenhagen, Denmark | Aalborg University | Danish Covid-19 Genome Consortium et al |
| EPI_ISL_6243409 | Department of Bacteria, Parasites and Fungi, Statens Serum Institut, Copenhagen, Denmark | Statens Serum Institut Bioinformatics and Microbial Genomics | Danish Covid-19 Genome Consortium et al |
| EPI_ISL_843313 | Department of Virus and Microbiological Special Diagnostics, Statens Serum Institut, Copenhagen, Denmark | Albertsen Lab, Department of Chemistry and Bioscience, Aalborg University, Denmark | Danish Covid-19 Genome Consortium et al |
| EPI_ISL_6131053 | Department of Bacteria, Parasites and Fungi, Statens Serum Institut, Copenhagen, Denmark | Statens Serum Institut Bioinformatics and Microbial Genomics | Danish Covid-19 Genome Consortium et al |
| EPI_ISL_972967 | Department of Virus and Microbiological Special Diagnostics, Statens Serum Institut, Copenhagen, Denmark | Aalborg University | Danish Covid-19 Genome Consortium et al |
| EPI_ISL_617778 | Department of Virus and Microbiological Special Diagnostics, Statens Serum Institut, Denmark | Albertsen lab, Department of Chemistry and Bioscience, Aalborg University, Denmark | Danish Covid-19 Genome Consortia et al |
| EPI_ISL_6438423 | Department of Bacteria, Parasites and Fungi, Statens Serum Institut, Copenhagen, Denmark | Statens Serum Institut Bioinformatics and Microbial Genomics | Danish Covid-19 Genome Consortium et al |

| | | | |
|---|---|---|---|
| EPI_ISL_1858077 | Department of Virus and Microbiological Special Diagnostics, Statens Serum Institut, Copenhagen, Denmark | Aalborg University | Danish Covid-19 Genome Consortium et al |
| EPI_ISL_6878663 | Department of Bacteria, Parasites and Fungi, Statens Serum Institut, Copenhagen, Denmark | Statens Serum Institut Bioinformatics and Microbial Genomics | Danish Covid-19 Genome Consortium et al |
| EPI_ISL_1125604 | Department of Virus and Microbiological Special Diagnostics, Statens Serum Institut, Copenhagen, Denmark | Aalborg University | Danish Covid-19 Genome Consortium et al |
| EPI_ISL_5882003 | Department of Bacteria, Parasites and Fungi, Statens Serum Institut, Copenhagen, Denmark | Statens Serum Institut Bioinformatics and Microbial Genomics | Danish Covid-19 Genome Consortium et al |
| EPI_ISL_1873069 | Department of Virus and Microbiological Special Diagnostics, Statens Serum Institut, Copenhagen, Denmark | Aalborg University | Danish Covid-19 Genome Consortium et al |
| EPI_ISL_7062393 | Department of Bacteria, Parasites and Fungi, Statens Serum Institut, Copenhagen, Denmark | Statens Serum Institut Bioinformatics and Microbial Genomics | Danish Covid-19 Genome Consortium et al |
| EPI_ISL_1872204 | Department of Virus and Microbiological Special Diagnostics, Statens Serum Institut, Copenhagen, Denmark | Aalborg University | Danish Covid-19 Genome Consortium et al |
| EPI_ISL_6880716 | Department of Bacteria, Parasites and Fungi, Statens Serum Institut, Copenhagen, Denmark | Statens Serum Institut Bioinformatics and Microbial Genomics | Danish Covid-19 Genome Consortium et al |
| EPI_ISL_930182 | Department of Virus and Microbiological Special Diagnostics, Statens Serum Institut, Copenhagen, Denmark | Aalborg University | Danish Covid-19 Genome Consortium et al |
| EPI_ISL_1872408 | Department of Virus and Microbiological Special Diagnostics, Statens Serum Institut, Copenhagen, Denmark | Aalborg University | Danish Covid-19 Genome Consortium et al |
| EPI_ISL_1877847 | Department of Virus and Microbiological Special Diagnostics, Statens Serum Institut, Copenhagen, Denmark | Aalborg University | Danish Covid-19 Genome Consortium et al |

| | | | |
|---|---|---|---|
| EPI_ISL_869643 | Department of Virus and Microbiological Special Diagnostics, Statens Serum Institut, Copenhagen, Denmark | Aalborg University | Danish Covid-19 Genome Consortium et al |
| EPI_ISL_6965299 | Department of Bacteria, Parasites and Fungi, Statens Serum Institut, Copenhagen, Denmark | Statens Serum Institut Bioinformatics and Microbial Genomics | Danish Covid-19 Genome Consortium et al |
| EPI_ISL_1873903 | Department of Virus and Microbiological Special Diagnostics, Statens Serum Institut, Copenhagen, Denmark | Aalborg University | Danish Covid-19 Genome Consortium et al |
| EPI_ISL_6878696 | Department of Bacteria, Parasites and Fungi, Statens Serum Institut, Copenhagen, Denmark | Statens Serum Institut Bioinformatics and Microbial Genomics | Danish Covid-19 Genome Consortium et al |
| EPI_ISL_6965921 | Department of Bacteria, Parasites and Fungi, Statens Serum Institut, Copenhagen, Denmark | Statens Serum Institut Bioinformatics and Microbial Genomics | Danish Covid-19 Genome Consortium et al |
| EPI_ISL_6829921 | Department of Bacteria, Parasites and Fungi, Statens Serum Institut, Copenhagen, Denmark | Statens Serum Institut Bioinformatics and Microbial Genomics | Danish Covid-19 Genome Consortium et al |
| EPI_ISL_6130280 | Department of Bacteria, Parasites and Fungi, Statens Serum Institut, Copenhagen, Denmark | Statens Serum Institut Bioinformatics and Microbial Genomics | Danish Covid-19 Genome Consortium et al |
| EPI_ISL_7062087 | Department of Bacteria, Parasites and Fungi, Statens Serum Institut, Copenhagen, Denmark | Statens Serum Institut Bioinformatics and Microbial Genomics | Danish Covid-19 Genome Consortium et al |
| EPI_ISL_759587 | Department of Virus and Microbiological Special Diagnostics, Statens Serum Institut, Copenhagen, Denmark | Albertsen Lab, Department of Chemistry and Bioscience, Aalborg University, Denmark | Danish Covid-19 Genome Consortium et al |
| EPI_ISL_846796 | Department of Virus and Microbiological Special Diagnostics, Statens Serum Institut, Copenhagen, Denmark | Albertsen Lab, Department of Chemistry and Bioscience, Aalborg University, Denmark | Danish Covid-19 Genome Consortium et al |
| EPI_ISL_1864710 | Department of Virus and Microbiological Special Diagnostics, Statens Serum Institut, Copenhagen, Denmark | Aalborg University | Danish Covid-19 Genome Consortium et al |

| | | | |
|---|---|---|---|
| EPI_ISL_970974 | Department of Virus and Microbiological Special Diagnostics, Statens Serum Institut, Copenhagen, Denmark | Aalborg University | Danish Covid-19 Genome Consortium et al |
| EPI_ISL_6968169 | Department of Bacteria, Parasites and Fungi, Statens Serum Institut, Copenhagen, Denmark | Statens Serum Institut Bioinformatics and Microbial Genomics | Danish Covid-19 Genome Consortium et al |
| EPI_ISL_6967619 | Department of Bacteria, Parasites and Fungi, Statens Serum Institut, Copenhagen, Denmark | Statens Serum Institut Bioinformatics and Microbial Genomics | Danish Covid-19 Genome Consortium et al |
| EPI_ISL_6611149 | Department of Bacteria, Parasites and Fungi, Statens Serum Institut, Copenhagen, Denmark | Statens Serum Institut Bioinformatics and Microbial Genomics | Danish Covid-19 Genome Consortium et al |
| EPI_ISL_6773002 | Department of Bacteria, Parasites and Fungi, Statens Serum Institut, Copenhagen, Denmark | Statens Serum Institut Bioinformatics and Microbial Genomics | Danish Covid-19 Genome Consortium et al |
| EPI_ISL_1875649 | Department of Virus and Microbiological Special Diagnostics, Statens Serum Institut, Copenhagen, Denmark | Aalborg University | Danish Covid-19 Genome Consortium et al |
| EPI_ISL_616486 | Department of Virus and Microbiological Special Diagnostics, Statens Serum Institut, Denmark | Albertsen lab, Department of Chemistry and Bioscience, Aalborg University, Denmark | Danish Covid-19 Genome Consortia et al |
| EPI_ISL_1859567 | Department of Virus and Microbiological Special Diagnostics, Statens Serum Institut, Copenhagen, Denmark | Aalborg University | Danish Covid-19 Genome Consortium et al |
| EPI_ISL_2092194 | Lighthouse Lab in Alderley Park | Wellcome Sanger Institute for the COVID-19 Genomics UK (COG-UK) Consortium | Jacquelyn Wynn et al |
| EPI_ISL_1100146 | Lighthouse Lab in Alderley Park | Wellcome Sanger Institute for the COVID-19 Genomics UK (COG-UK) Consortium | Jacquelyn Wynn et al |
| EPI_ISL_878203 | Lighthouse Lab in Alderley Park | Wellcome Sanger Institute for the COVID-19 Genomics UK (COG-UK) Consortium | Jacquelyn Wynn et al |
| EPI_ISL_1896605 | Department of Virus and Microbiological Special Diagnostics, Statens Serum Institut, Copenhagen, Denmark | Aalborg University | Danish Covid-19 Genome Consortium et al |
| EPI_ISL_1100203 | Lighthouse Lab in Alderley Park | Wellcome Sanger Institute for the COVID-19 Genomics UK (COG-UK) Consortium | Jacquelyn Wynn et al |

| | | | |
|---|---|---|---|
| EPI_ISL_886078 | Lighthouse Lab in Alderley Park | Wellcome Sanger Institute for the COVID-19 Genomics UK (COG-UK) Consortium | Jacquelyn Wynn et al |
| EPI_ISL_994499 | Lighthouse Lab in Alderley Park | Wellcome Sanger Institute for the COVID-19 Genomics UK (COG-UK) Consortium | Jacquelyn Wynn et al |
| EPI_ISL_917276 | Lighthouse Lab in Alderley Park | Wellcome Sanger Institute for the COVID-19 Genomics UK (COG-UK) Consortium | Jacquelyn Wynn et al |
| EPI_ISL_931649 | Lighthouse Lab in Alderley Park | Wellcome Sanger Institute for the COVID-19 Genomics UK (COG-UK) Consortium | Jacquelyn Wynn et al |
| EPI_ISL_1256080 | Lighthouse Lab in Alderley Park | Wellcome Sanger Institute for the COVID-19 Genomics UK (COG-UK) Consortium | Jacquelyn Wynn et al |
| EPI_ISL_1176916 | Northumbria University / South Tees Hospitals NHS Foundation Trust / North Cumbria Integrated Care NHS Foundation Trust / North Tees and Hartlepool NHS Foundation Trust / Newcastle Hospitals NHS Foundation Trust | COVID-19 Genomics UK (COG-UK) Consortium | Darren L Smith et al |
| EPI_ISL_464698 | Respiratory Virus Unit, Microbiology Services Colindale, Public Health England | Respiratory Virus Unit, Microbiology Services Colindale, Public Health England | PHE Covid Sequencing Team et al |
| EPI_ISL_1564477 | Lighthouse Lab in Alderley Park | Wellcome Sanger Institute for the COVID-19 Genomics UK (COG-UK) Consortium | Jacquelyn Wynn et al |
| EPI_ISL_1374344 | Lighthouse Lab in Alderley Park | Wellcome Sanger Institute for the COVID-19 Genomics UK (COG-UK) Consortium | Jacquelyn Wynn et al |
| EPI_ISL_2092564 | Lighthouse Lab in Alderley Park | Wellcome Sanger Institute for the COVID-19 Genomics UK (COG-UK) Consortium | Jacquelyn Wynn et al |
| EPI_ISL_1791017 | Lighthouse Lab in Alderley Park | Wellcome Sanger Institute for the COVID-19 Genomics UK (COG-UK) Consortium | Jacquelyn Wynn et al |
| EPI_ISL_1885913 | Department of Virus and Microbiological Special Diagnostics, Statens Serum Institut, Copenhagen, Denmark | Aalborg University | Danish Covid-19 Genome Consortium et al |
| EPI_ISL_1414539 | Lighthouse Lab in Alderley Park | Wellcome Sanger Institute for the COVID-19 Genomics UK (COG-UK) Consortium | Jacquelyn Wynn et al |
| EPI_ISL_1206555 | Lighthouse Lab in Alderley Park | Wellcome Sanger Institute for the COVID-19 Genomics UK (COG-UK) Consortium | Jacquelyn Wynn et al |

| | | | |
|---|---|---|---|
| EPI_ISL_1896644 | Department of Virus and Microbiological Special Diagnostics, Statens Serum Institut, Copenhagen, Denmark | Aalborg University | Danish Covid-19 Genome Consortium et al |
| EPI_ISL_1308080 | Quadram Institute Bioscience | COVID-19 Genomics UK (COG-UK) Consortium | Dave J. Baker et al |
| EPI_ISL_2173590 | Department of Virus and Microbiological Special Diagnostics, Statens Serum Institut, Copenhagen, Denmark | Aalborg University | Danish Covid-19 Genome Consortium et al |
| EPI_ISL_1892013 | Molekylær Medicinsk Afdeling, Aarhus University Hospital, Aarhus, Denmark | Aalborg University | Danish Covid-19 Genome Consortium et al |
| EPI_ISL_1083619 | Lighthouse Lab in Alderley Park | Wellcome Sanger Institute for the COVID-19 Genomics UK (COG-UK) Consortium | Jacquelyn Wynn et al |
| EPI_ISL_466559 | Respiratory Virus Unit, Microbiology Services Colindale, Public Health England | Respiratory Virus Unit, Microbiology Services Colindale, Public Health England | PHE Covid Sequencing Team et al |
| EPI_ISL_1246013 | Lighthouse Lab in Alderley Park | Wellcome Sanger Institute for the COVID-19 Genomics UK (COG-UK) Consortium | Jacquelyn Wynn et al |
| EPI_ISL_791191 | Respiratory Virus Unit, National Infection Service, Public Health England | COVID-19 Genomics UK (COG-UK) Consortium | PHE Covid Sequencing Team et al |
| EPI_ISL_886090 | Lighthouse Lab in Alderley Park | Wellcome Sanger Institute for the COVID-19 Genomics UK (COG-UK) Consortium | Jacquelyn Wynn et al |
| EPI_ISL_1887845 | Department of Virus and Microbiological Special Diagnostics, Statens Serum Institut, Copenhagen, Denmark | Aalborg University | Danish Covid-19 Genome Consortium et al |
| EPI_ISL_886117 | Lighthouse Lab in Alderley Park | Wellcome Sanger Institute for the COVID-19 Genomics UK (COG-UK) Consortium | Jacquelyn Wynn et al |
| EPI_ISL_1536883 | Lighthouse Lab in Alderley Park | Wellcome Sanger Institute for the COVID-19 Genomics UK (COG-UK) Consortium | Jacquelyn Wynn et al |
| EPI_ISL_1894711 | Department of Virus and Microbiological Special Diagnostics, Statens Serum Institut, Copenhagen, Denmark | Aalborg University | Danish Covid-19 Genome Consortium et al |
| EPI_ISL_931734 | Lighthouse Lab in Alderley Park | Wellcome Sanger Institute for the COVID-19 Genomics UK (COG-UK) Consortium | Jacquelyn Wynn et al |

| | | | |
|---|---|---|---|
| EPI_ISL_992117 | Lighthouse Lab in Alderley Park | Wellcome Sanger Institute for the COVID-19 Genomics UK (COG-UK) Consortium | Jacquelyn Wynn et al |
| EPI_ISL_465660 | Respiratory Virus Unit, Microbiology Services Colindale, Public Health England | Respiratory Virus Unit, Microbiology Services Colindale, Public Health England | PHE Covid Sequencing Team et al |
| EPI_ISL_1888931 | Department of Virus and Microbiological Special Diagnostics, Statens Serum Institut, Copenhagen, Denmark | Aalborg University | Danish Covid-19 Genome Consortium et al |
| EPI_ISL_2024546 | Department of Virus and Microbiological Special Diagnostics, Statens Serum Institut, Copenhagen, Denmark | Aalborg University | Danish Covid-19 Genome Consortium et al |
| EPI_ISL_1486821 | Lighthouse Lab in Alderley Park | Wellcome Sanger Institute for the COVID-19 Genomics UK (COG-UK) Consortium | Jacquelyn Wynn et al |
| EPI_ISL_1327146 | Lighthouse Lab in Alderley Park | Wellcome Sanger Institute for the COVID-19 Genomics UK (COG-UK) Consortium | Jacquelyn Wynn et al |
| EPI_ISL_937782 | Lighthouse Lab in Alderley Park | Wellcome Sanger Institute for the COVID-19 Genomics UK (COG-UK) Consortium | Jacquelyn Wynn et al |
| EPI_ISL_1895851 | Department of Virus and Microbiological Special Diagnostics, Statens Serum Institut, Copenhagen, Denmark | Aalborg University | Danish Covid-19 Genome Consortium et al |
| EPI_ISL_1468307 | Lighthouse Lab in Alderley Park | Wellcome Sanger Institute for the COVID-19 Genomics UK (COG-UK) Consortium | Jacquelyn Wynn et al |
| EPI_ISL_1893321 | Department of Virus and Microbiological Special Diagnostics, Statens Serum Institut, Copenhagen, Denmark | Aalborg University | Danish Covid-19 Genome Consortium et al |
| EPI_ISL_1244462 | Lighthouse Lab in Alderley Park | Wellcome Sanger Institute for the COVID-19 Genomics UK (COG-UK) Consortium | Jacquelyn Wynn et al |
| EPI_ISL_1416034 | Lighthouse Lab in Alderley Park | Wellcome Sanger Institute for the COVID-19 Genomics UK (COG-UK) Consortium | Jacquelyn Wynn et al |
| EPI_ISL_885354 | Lighthouse Lab in Alderley Park | Wellcome Sanger Institute for the COVID-19 Genomics UK (COG-UK) Consortium | Jacquelyn Wynn et al |
| EPI_ISL_4496989 | INSPI-CRN DE INFLUENZA Y OTROS VIRUS RESPIRATORIOS | NIC-INSPI | Alfredo Bruno et al |

| | | | |
|---|---|---|---|
| EPI_ISL_881506 | Lighthouse Lab in Alderley Park | Wellcome Sanger Institute for the COVID-19 Genomics UK (COG-UK) Consortium | Jacquelyn Wynn et al |
| EPI_ISL_1485299 | Lighthouse Lab in Alderley Park | Wellcome Sanger Institute for the COVID-19 Genomics UK (COG-UK) Consortium | Jacquelyn Wynn et al |
| EPI_ISL_1410050 | Lighthouse Lab in Alderley Park | Wellcome Sanger Institute for the COVID-19 Genomics UK (COG-UK) Consortium | Jacquelyn Wynn et al |
| EPI_ISL_1415659 | Lighthouse Lab in Alderley Park | Wellcome Sanger Institute for the COVID-19 Genomics UK (COG-UK) Consortium | Jacquelyn Wynn et al |
| EPI_ISL_1517635 | Lighthouse Lab in Alderley Park | Wellcome Sanger Institute for the COVID-19 Genomics UK (COG-UK) Consortium | Jacquelyn Wynn et al |
| EPI_ISL_614826 | Department of Virus and Microbiological Special Diagnostics, Statens Serum Institut, Denmark | Albertsen lab, Department of Chemistry and Bioscience, Aalborg University, Denmark | Danish Covid-19 Genome Consortia et al |
| EPI_ISL_1895322 | Department of Virus and Microbiological Special Diagnostics, Statens Serum Institut, Copenhagen, Denmark | Aalborg University | Danish Covid-19 Genome Consortium et al |
| EPI_ISL_1044933 | Lighthouse Lab in Alderley Park | Wellcome Sanger Institute for the COVID-19 Genomics UK (COG-UK) Consortium | Jacquelyn Wynn et al |
| EPI_ISL_2176421 | Department of Virus and Microbiological Special Diagnostics, Statens Serum Institut, Copenhagen, Denmark | Aalborg University | Danish Covid-19 Genome Consortium et al |
| EPI_ISL_1484903 | Lighthouse Lab in Alderley Park | Wellcome Sanger Institute for the COVID-19 Genomics UK (COG-UK) Consortium | Jacquelyn Wynn et al |
| EPI_ISL_2025368 | Department of Virus and Microbiological Special Diagnostics, Statens Serum Institut, Copenhagen, Denmark | Aalborg University | Danish Covid-19 Genome Consortium et al |
| EPI_ISL_464936 | Respiratory Virus Unit, Microbiology Services Colindale, Public Health England | Respiratory Virus Unit, Microbiology Services Colindale, Public Health England | PHE Covid Sequencing Team et al |
| EPI_ISL_1413145 | Lighthouse Lab in Alderley Park | Wellcome Sanger Institute for the COVID-19 Genomics UK (COG-UK) Consortium | Jacquelyn Wynn et al |
| EPI_ISL_1891458 | Department of Virus and Microbiological Special Diagnostics, Statens Serum Institut, Copenhagen, Denmark | Aalborg University | Danish Covid-19 Genome Consortium et al |

| | | | |
|---|---|---|---|
| EPI_ISL_1891107 | Department of Virus and Microbiological Special Diagnostics, Statens Serum Institut, Copenhagen, Denmark | Aalborg University | Danish Covid-19 Genome Consortium et al |
| EPI_ISL_1245648 | Lighthouse Lab in Alderley Park | Wellcome Sanger Institute for the COVID-19 Genomics UK (COG-UK) Consortium | Jacquelyn Wynn et al |
| EPI_ISL_987014 | Lighthouse Lab in Alderley Park | Wellcome Sanger Institute for the COVID-19 Genomics UK (COG-UK) Consortium | Jacquelyn Wynn et al |
| EPI_ISL_1450361 | Lighthouse Lab in Alderley Park | Wellcome Sanger Institute for the COVID-19 Genomics UK (COG-UK) Consortium | Jacquelyn Wynn et al |
| EPI_ISL_1331357 | Lighthouse Lab in Alderley Park | Wellcome Sanger Institute for the COVID-19 Genomics UK (COG-UK) Consortium | Jacquelyn Wynn et al |
| EPI_ISL_1759038 | Lighthouse Lab in Alderley Park | Wellcome Sanger Institute for the COVID-19 Genomics UK (COG-UK) Consortium | Jacquelyn Wynn et al |
| EPI_ISL_1884752 | Department of Virus and Microbiological Special Diagnostics, Statens Serum Institut, Copenhagen, Denmark | Aalborg University | Danish Covid-19 Genome Consortium et al |
| EPI_ISL_2176585 | Department of Virus and Microbiological Special Diagnostics, Statens Serum Institut, Copenhagen, Denmark | Aalborg University | Danish Covid-19 Genome Consortium et al |
| EPI_ISL_1987354 | Lighthouse Lab in Alderley Park | Wellcome Sanger Institute for the COVID-19 Genomics UK (COG-UK) Consortium | Jacquelyn Wynn et al |
| EPI_ISL_617246 | Department of Virus and Microbiological Special Diagnostics, Statens Serum Institut, Denmark | Albertsen lab, Department of Chemistry and Bioscience, Aalborg University, Denmark | Danish Covid-19 Genome Consortia et al |
| EPI_ISL_1452044 | Lighthouse Lab in Alderley Park | Wellcome Sanger Institute for the COVID-19 Genomics UK (COG-UK) Consortium | Jacquelyn Wynn et al |
| EPI_ISL_1892369 | Department of Virus and Microbiological Special Diagnostics, Statens Serum Institut, Copenhagen, Denmark | Aalborg University | Danish Covid-19 Genome Consortium et al |
| EPI_ISL_617348 | Department of Virus and Microbiological Special Diagnostics, Statens Serum Institut, Denmark | Albertsen lab, Department of Chemistry and Bioscience, Aalborg University, Denmark | Danish Covid-19 Genome Consortia et al |
| EPI_ISL_988923 | Lighthouse Lab in Alderley Park | Wellcome Sanger Institute for the COVID-19 Genomics UK (COG-UK) Consortium | Jacquelyn Wynn et al |

| | | | |
|---|---|---|---|
| EPI_ISL_2757639 | INSPI-CRN DE INFLUENZA Y OTROS VIRUS RESPIRATORIOS | NIC-INSPI | Alfredo Bruno et al |
| EPI_ISL_2899983 | Respiratory Virus Unit, Microbiology Services Colindale, Public Health England | COVID-19 Genomics UK (COG-UK) Consortium | PHE Covid Sequencing Team et al |
| EPI_ISL_1205941 | Lighthouse Lab in Alderley Park | Wellcome Sanger Institute for the COVID-19 Genomics UK (COG-UK) Consortium | Jacquelyn Wynn et al |
| EPI_ISL_2026040 | Department of Virus and Microbiological Special Diagnostics, Statens Serum Institut, Copenhagen, Denmark | Aalborg University | Danish Covid-19 Genome Consortium et al |
| EPI_ISL_1451645 | Lighthouse Lab in Alderley Park | Wellcome Sanger Institute for the COVID-19 Genomics UK (COG-UK) Consortium | Jacquelyn Wynn et al |
| EPI_ISL_1519443 | Lighthouse Lab in Alderley Park | Wellcome Sanger Institute for the COVID-19 Genomics UK (COG-UK) Consortium | Jacquelyn Wynn et al |
| EPI_ISL_885867 | Lighthouse Lab in Alderley Park | Wellcome Sanger Institute for the COVID-19 Genomics UK (COG-UK) Consortium | Jacquelyn Wynn et al |
| EPI_ISL_466109 | Respiratory Virus Unit, Microbiology Services Colindale, Public Health England | Respiratory Virus Unit, Microbiology Services Colindale, Public Health England | PHE Covid Sequencing Team et al |
| EPI_ISL_880516 | Lighthouse Lab in Alderley Park | Wellcome Sanger Institute for the COVID-19 Genomics UK (COG-UK) Consortium | Jacquelyn Wynn et al |
| EPI_ISL_879986 | Lighthouse Lab in Alderley Park | Wellcome Sanger Institute for the COVID-19 Genomics UK (COG-UK) Consortium | Jacquelyn Wynn et al |
| EPI_ISL_1177341 | Centre for Enzyme Innovation, University of Portsmouth / Translational Research Laboratory, Portsmouth Hospitals NHS Trust | COVID-19 Genomics UK (COG-UK) Consortium | Angela Beckett et al |
| EPI_ISL_1536417 | Lighthouse Lab in Alderley Park | Wellcome Sanger Institute for the COVID-19 Genomics UK (COG-UK) Consortium | Jacquelyn Wynn et al |
| EPI_ISL_2091814 | Lighthouse Lab in Alderley Park | Wellcome Sanger Institute for the COVID-19 Genomics UK (COG-UK) Consortium | Jacquelyn Wynn et al |
| EPI_ISL_1189675 | Lighthouse Lab in Alderley Park | Wellcome Sanger Institute for the COVID-19 Genomics UK (COG-UK) Consortium | Jacquelyn Wynn et al |
| EPI_ISL_1564763 | Lighthouse Lab in Alderley Park | Wellcome Sanger Institute for the COVID-19 Genomics UK (COG-UK) Consortium | Jacquelyn Wynn et al |

| | | | |
|---|---|---|---|
| EPI_ISL_804256 | Respiratory Virus Unit, National Infection Service, Public Health England | COVID-19 Genomics UK (COG-UK) Consortium | PHE Covid Sequencing Team et al |
| EPI_ISL_2026251 | Department of Virus and Microbiological Special Diagnostics, Statens Serum Institut, Copenhagen, Denmark | Aalborg University | Danish Covid-19 Genome Consortium et al |
| EPI_ISL_874433 | Lighthouse Lab in Alderley Park | Wellcome Sanger Institute for the COVID-19 Genomics UK (COG-UK) Consortium | Jacquelyn Wynn et al |
| EPI_ISL_969520 | Lighthouse Lab in Alderley Park | Wellcome Sanger Institute for the COVID-19 Genomics UK (COG-UK) Consortium | Jacquelyn Wynn et al |
| EPI_ISL_626165 | Department of Virus and Microbiological Special Diagnostics, Statens Serum Institut, Denmark | Albertsen lab, Department of Chemistry and Bioscience, Aalborg University, Denmark | Danish Covid-19 Genome Consortia et al |
| EPI_ISL_1487418 | Lighthouse Lab in Alderley Park | Wellcome Sanger Institute for the COVID-19 Genomics UK (COG-UK) Consortium | Jacquelyn Wynn et al |
| EPI_ISL_1895979 | Department of Clinical Microbiology and Center for Genomic Medicine, Rigshospitalet, Copenhagen, Denmark | Aalborg University | Danish Covid-19 Genome Consortium et al |
| EPI_ISL_991877 | Lighthouse Lab in Alderley Park | Wellcome Sanger Institute for the COVID-19 Genomics UK (COG-UK) Consortium | Jacquelyn Wynn et al |
| EPI_ISL_1893326 | Department of Virus and Microbiological Special Diagnostics, Statens Serum Institut, Copenhagen, Denmark | Aalborg University | Danish Covid-19 Genome Consortium et al |
| EPI_ISL_2533868 | Naval Medical Research Unit No. 3 | Naval Medical Research Center Biological Defense Research Directorate | Catherine E. Arnold et al |
| EPI_ISL_1883356 | Department of Virus and Microbiological Special Diagnostics, Statens Serum Institut, Copenhagen, Denmark | Aalborg University | Danish Covid-19 Genome Consortium et al |
| EPI_ISL_2540648 | Lighthouse Lab in Alderley Park | Wellcome Sanger Institute for the COVID-19 Genomics UK (COG-UK) Consortium | Jacquelyn Wynn et al |
| EPI_ISL_3093331 | Lighthouse Lab in Alderley Park | Wellcome Sanger Institute for the COVID-19 Genomics UK (COG-UK) Consortium | Jacquelyn Wynn et al |
| EPI_ISL_3173380 | Lighthouse Lab in Alderley Park | Wellcome Sanger Institute for the COVID-19 Genomics UK (COG-UK) Consortium | Jacquelyn Wynn et al |

| | | | |
|---|---|---|---|
| EPI_ISL_2568574 | Lighthouse Lab in Alderley Park | Wellcome Sanger Institute for the COVID-19 Genomics UK (COG-UK) Consortium | Jacquelyn Wynn et al |
| EPI_ISL_2515850 | Lighthouse Lab in Alderley Park | Wellcome Sanger Institute for the COVID-19 Genomics UK (COG-UK) Consortium | Jacquelyn Wynn et al |
| EPI_ISL_3362438 | Lighthouse Lab in Alderley Park | Wellcome Sanger Institute for the COVID-19 Genomics UK (COG-UK) Consortium | Jacquelyn Wynn et al |
| EPI_ISL_2394720 | Lighthouse Lab in Alderley Park | Wellcome Sanger Institute for the COVID-19 Genomics UK (COG-UK) Consortium | Jacquelyn Wynn et al |
| EPI_ISL_3378845 | Lighthouse Lab in Alderley Park | Wellcome Sanger Institute for the COVID-19 Genomics UK (COG-UK) Consortium | Jacquelyn Wynn et al |
| EPI_ISL_2346719 | Lighthouse Lab in Alderley Park | Wellcome Sanger Institute for the COVID-19 Genomics UK (COG-UK) Consortium | Jacquelyn Wynn et al |
| EPI_ISL_2639486 | Lighthouse Lab in Alderley Park | Wellcome Sanger Institute for the COVID-19 Genomics UK (COG-UK) Consortium | Jacquelyn Wynn et al |
| EPI_ISL_3339087 | Lighthouse Lab in Alderley Park | Wellcome Sanger Institute for the COVID-19 Genomics UK (COG-UK) Consortium | Jacquelyn Wynn et al |
| EPI_ISL_3706428 | Lighthouse Lab in Alderley Park | Wellcome Sanger Institute for the COVID-19 Genomics UK (COG-UK) Consortium | Jacquelyn Wynn et al |
| EPI_ISL_3469803 | Lighthouse Lab in Alderley Park | Wellcome Sanger Institute for the COVID-19 Genomics UK (COG-UK) Consortium | Jacquelyn Wynn et al |
| EPI_ISL_4144293 | Lighthouse Lab in Alderley Park | Wellcome Sanger Institute for the COVID-19 Genomics UK (COG-UK) Consortium | Jacquelyn Wynn et al |
| EPI_ISL_4405826 | Lighthouse Lab in Alderley Park | Wellcome Sanger Institute for the COVID-19 Genomics UK (COG-UK) Consortium | Jacquelyn Wynn et al |
| EPI_ISL_2393403 | Lighthouse Lab in Alderley Park | Wellcome Sanger Institute for the COVID-19 Genomics UK (COG-UK) Consortium | Jacquelyn Wynn et al |
| EPI_ISL_3363515 | Lighthouse Lab in Alderley Park | Wellcome Sanger Institute for the COVID-19 Genomics UK (COG-UK) Consortium | Jacquelyn Wynn et al |
| EPI_ISL_3248317 | Lighthouse Lab in Alderley Park | Wellcome Sanger Institute for the COVID-19 Genomics UK (COG-UK) Consortium | Jacquelyn Wynn et al |
| EPI_ISL_3172623 | Lighthouse Lab in Alderley Park | Wellcome Sanger Institute for the COVID-19 Genomics UK (COG-UK) Consortium | Jacquelyn Wynn et al |
| EPI_ISL_3377157 | Lighthouse Lab in Alderley Park | Wellcome Sanger Institute for the COVID-19 Genomics UK (COG-UK) Consortium | Jacquelyn Wynn et al |

| | | | |
|---|---|---|---|
| EPI_ISL_3283247 | Lighthouse Lab in Alderley Park | Wellcome Sanger Institute for the COVID-19 Genomics UK (COG-UK) Consortium | Jacquelyn Wynn et al |
| EPI_ISL_3808359 | Lighthouse Lab in Alderley Park | Wellcome Sanger Institute for the COVID-19 Genomics UK (COG-UK) Consortium | Jacquelyn Wynn et al |
| EPI_ISL_2317962 | Lighthouse Lab in Alderley Park | Wellcome Sanger Institute for the COVID-19 Genomics UK (COG-UK) Consortium | Jacquelyn Wynn et al |
| EPI_ISL_2436750 | Lighthouse Lab in Alderley Park | Wellcome Sanger Institute for the COVID-19 Genomics UK (COG-UK) Consortium | Jacquelyn Wynn et al |
| EPI_ISL_2846297 | Lighthouse Lab in Alderley Park | Wellcome Sanger Institute for the COVID-19 Genomics UK (COG-UK) Consortium | Jacquelyn Wynn et al |
| EPI_ISL_2828906 | Lighthouse Lab in Alderley Park | Wellcome Sanger Institute for the COVID-19 Genomics UK (COG-UK) Consortium | Jacquelyn Wynn et al |
| EPI_ISL_3808256 | Lighthouse Lab in Alderley Park | Wellcome Sanger Institute for the COVID-19 Genomics UK (COG-UK) Consortium | Jacquelyn Wynn et al |
| EPI_ISL_2394645 | Lighthouse Lab in Alderley Park | Wellcome Sanger Institute for the COVID-19 Genomics UK (COG-UK) Consortium | Jacquelyn Wynn et al |
| EPI_ISL_3361436 | Lighthouse Lab in Alderley Park | Wellcome Sanger Institute for the COVID-19 Genomics UK (COG-UK) Consortium | Jacquelyn Wynn et al |
| EPI_ISL_2348446 | Lighthouse Lab in Alderley Park | Wellcome Sanger Institute for the COVID-19 Genomics UK (COG-UK) Consortium | Jacquelyn Wynn et al |
| EPI_ISL_2553488 | Lighthouse Lab in Alderley Park | Wellcome Sanger Institute for the COVID-19 Genomics UK (COG-UK) Consortium | Jacquelyn Wynn et al |
| EPI_ISL_3641288 | Lighthouse Lab in Alderley Park | Wellcome Sanger Institute for the COVID-19 Genomics UK (COG-UK) Consortium | Jacquelyn Wynn et al |
| EPI_ISL_2717862 | Lighthouse Lab in Alderley Park | Wellcome Sanger Institute for the COVID-19 Genomics UK (COG-UK) Consortium | Jacquelyn Wynn et al |
| EPI_ISL_4246998 | Lighthouse Lab in Alderley Park | Wellcome Sanger Institute for the COVID-19 Genomics UK (COG-UK) Consortium | Jacquelyn Wynn et al |
| EPI_ISL_3093470 | Lighthouse Lab in Alderley Park | Wellcome Sanger Institute for the COVID-19 Genomics UK (COG-UK) Consortium | Jacquelyn Wynn et al |
| EPI_ISL_2366439 | Lighthouse Lab in Alderley Park | Wellcome Sanger Institute for the COVID-19 Genomics UK (COG-UK) Consortium | Jacquelyn Wynn et al |
| EPI_ISL_2925405 | Lighthouse Lab in Alderley Park | Wellcome Sanger Institute for the COVID-19 Genomics UK (COG-UK) Consortium | Jacquelyn Wynn et al |

| | | | |
|---|---|---|---|
| EPI_ISL_2717869 | Lighthouse Lab in Alderley Park | Wellcome Sanger Institute for the COVID-19 Genomics UK (COG-UK) Consortium | Jacquelyn Wynn et al |
| EPI_ISL_2318435 | Lighthouse Lab in Alderley Park | Wellcome Sanger Institute for the COVID-19 Genomics UK (COG-UK) Consortium | Jacquelyn Wynn et al |
| EPI_ISL_4097967 | Lighthouse Lab in Alderley Park | Wellcome Sanger Institute for the COVID-19 Genomics UK (COG-UK) Consortium | Jacquelyn Wynn et al |
| EPI_ISL_4402559 | Lighthouse Lab in Alderley Park | Wellcome Sanger Institute for the COVID-19 Genomics UK (COG-UK) Consortium | Jacquelyn Wynn et al |
| EPI_ISL_3338905 | Lighthouse Lab in Alderley Park | Wellcome Sanger Institute for the COVID-19 Genomics UK (COG-UK) Consortium | Jacquelyn Wynn et al |
| EPI_ISL_2552311 | Lighthouse Lab in Alderley Park | Wellcome Sanger Institute for the COVID-19 Genomics UK (COG-UK) Consortium | Jacquelyn Wynn et al |
| EPI_ISL_4263697 | Lighthouse Lab in Alderley Park | Wellcome Sanger Institute for the COVID-19 Genomics UK (COG-UK) Consortium | Jacquelyn Wynn et al |
| EPI_ISL_4349159 | Lighthouse Lab in Alderley Park | Wellcome Sanger Institute for the COVID-19 Genomics UK (COG-UK) Consortium | Jacquelyn Wynn et al |
| EPI_ISL_3953788 | Lighthouse Lab in Alderley Park | Wellcome Sanger Institute for the COVID-19 Genomics UK (COG-UK) Consortium | Jacquelyn Wynn et al |
| EPI_ISL_2469113 | Lighthouse Lab in Alderley Park | Wellcome Sanger Institute for the COVID-19 Genomics UK (COG-UK) Consortium | Jacquelyn Wynn et al |
| EPI_ISL_2641655 | Lighthouse Lab in Alderley Park | Wellcome Sanger Institute for the COVID-19 Genomics UK (COG-UK) Consortium | Jacquelyn Wynn et al |
| EPI_ISL_4522100 | Lighthouse Lab in Alderley Park | Wellcome Sanger Institute for the COVID-19 Genomics UK (COG-UK) Consortium | Jacquelyn Wynn et al |
| EPI_ISL_3702069 | Lighthouse Lab in Alderley Park | Wellcome Sanger Institute for the COVID-19 Genomics UK (COG-UK) Consortium | Jacquelyn Wynn et al |
| EPI_ISL_2729143 | Lighthouse Lab in Alderley Park | Wellcome Sanger Institute for the COVID-19 Genomics UK (COG-UK) Consortium | Jacquelyn Wynn et al |
| EPI_ISL_4287719 | Lighthouse Lab in Alderley Park | Wellcome Sanger Institute for the COVID-19 Genomics UK (COG-UK) Consortium | Jacquelyn Wynn et al |
| EPI_ISL_2353606 | Lighthouse Lab in Alderley Park | Wellcome Sanger Institute for the COVID-19 Genomics UK (COG-UK) Consortium | Jacquelyn Wynn et al |
| EPI_ISL_4087000 | Lighthouse Lab in Alderley Park | Wellcome Sanger Institute for the COVID-19 Genomics UK (COG-UK) Consortium | Jacquelyn Wynn et al |

| | | | |
|---|---|---|---|
| EPI_ISL_2767328 | Lighthouse Lab in Alderley Park | Wellcome Sanger Institute for the COVID-19 Genomics UK (COG-UK) Consortium | Jacquelyn Wynn et al |
| EPI_ISL_4087396 | Lighthouse Lab in Alderley Park | Wellcome Sanger Institute for the COVID-19 Genomics UK (COG-UK) Consortium | Jacquelyn Wynn et al |
| EPI_ISL_3700036 | Lighthouse Lab in Alderley Park | Wellcome Sanger Institute for the COVID-19 Genomics UK (COG-UK) Consortium | Jacquelyn Wynn et al |
| EPI_ISL_4189551 | Lighthouse Lab in Alderley Park | Wellcome Sanger Institute for the COVID-19 Genomics UK (COG-UK) Consortium | Jacquelyn Wynn et al |
| EPI_ISL_2925356 | Lighthouse Lab in Alderley Park | Wellcome Sanger Institute for the COVID-19 Genomics UK (COG-UK) Consortium | Jacquelyn Wynn et al |
| EPI_ISL_3365021 | Lighthouse Lab in Alderley Park | Wellcome Sanger Institute for the COVID-19 Genomics UK (COG-UK) Consortium | Jacquelyn Wynn et al |
| EPI_ISL_2513939 | Lighthouse Lab in Alderley Park | Wellcome Sanger Institute for the COVID-19 Genomics UK (COG-UK) Consortium | Jacquelyn Wynn et al |
| EPI_ISL_3311312 | Lighthouse Lab in Alderley Park | Wellcome Sanger Institute for the COVID-19 Genomics UK (COG-UK) Consortium | Jacquelyn Wynn et al |
| EPI_ISL_4247330 | Lighthouse Lab in Alderley Park | Wellcome Sanger Institute for the COVID-19 Genomics UK (COG-UK) Consortium | Jacquelyn Wynn et al |
| EPI_ISL_2729457 | Lighthouse Lab in Alderley Park | Wellcome Sanger Institute for the COVID-19 Genomics UK (COG-UK) Consortium | Jacquelyn Wynn et al |
| EPI_ISL_2914543 | Lighthouse Lab in Alderley Park | Wellcome Sanger Institute for the COVID-19 Genomics UK (COG-UK) Consortium | Jacquelyn Wynn et al |
| EPI_ISL_2393031 | Lighthouse Lab in Alderley Park | Wellcome Sanger Institute for the COVID-19 Genomics UK (COG-UK) Consortium | Jacquelyn Wynn et al |
| EPI_ISL_3580479 | Lighthouse Lab in Alderley Park | Wellcome Sanger Institute for the COVID-19 Genomics UK (COG-UK) Consortium | Jacquelyn Wynn et al |
| EPI_ISL_3337676 | Lighthouse Lab in Alderley Park | Wellcome Sanger Institute for the COVID-19 Genomics UK (COG-UK) Consortium | Jacquelyn Wynn et al |
| EPI_ISL_2830028 | Lighthouse Lab in Alderley Park | Wellcome Sanger Institute for the COVID-19 Genomics UK (COG-UK) Consortium | Jacquelyn Wynn et al |
| EPI_ISL_3364210 | Lighthouse Lab in Alderley Park | Wellcome Sanger Institute for the COVID-19 Genomics UK (COG-UK) Consortium | Jacquelyn Wynn et al |
| EPI_ISL_4014270 | Lighthouse Lab in Alderley Park | Wellcome Sanger Institute for the COVID-19 Genomics UK (COG-UK) Consortium | Jacquelyn Wynn et al |

| | | | |
|---|---|---|---|
| EPI_ISL_2605548 | Lighthouse Lab in Alderley Park | Wellcome Sanger Institute for the COVID-19 Genomics UK (COG-UK) Consortium | Jacquelyn Wynn et al |
| EPI_ISL_2138356 | Lighthouse Lab in Alderley Park | Wellcome Sanger Institute for the COVID-19 Genomics UK (COG-UK) Consortium | Jacquelyn Wynn et al |
| EPI_ISL_2199426 | Lighthouse Lab in Alderley Park | Wellcome Sanger Institute for the COVID-19 Genomics UK (COG-UK) Consortium | Jacquelyn Wynn et al |
| EPI_ISL_2553460 | Lighthouse Lab in Alderley Park | Wellcome Sanger Institute for the COVID-19 Genomics UK (COG-UK) Consortium | Jacquelyn Wynn et al |
| EPI_ISL_3364158 | Lighthouse Lab in Alderley Park | Wellcome Sanger Institute for the COVID-19 Genomics UK (COG-UK) Consortium | Jacquelyn Wynn et al |
| EPI_ISL_3109370 | Lighthouse Lab in Alderley Park | Wellcome Sanger Institute for the COVID-19 Genomics UK (COG-UK) Consortium | Jacquelyn Wynn et al |
| EPI_ISL_3702276 | Lighthouse Lab in Alderley Park | Wellcome Sanger Institute for the COVID-19 Genomics UK (COG-UK) Consortium | Jacquelyn Wynn et al |
| EPI_ISL_4502130 | Lighthouse Lab in Alderley Park | Wellcome Sanger Institute for the COVID-19 Genomics UK (COG-UK) Consortium | Jacquelyn Wynn et al |
| EPI_ISL_4142325 | Lighthouse Lab in Alderley Park | Wellcome Sanger Institute for the COVID-19 Genomics UK (COG-UK) Consortium | Jacquelyn Wynn et al |
| EPI_ISL_4218232 | Lighthouse Lab in Alderley Park | Wellcome Sanger Institute for the COVID-19 Genomics UK (COG-UK) Consortium | Jacquelyn Wynn et al |
| EPI_ISL_4523606 | Lighthouse Lab in Alderley Park | Wellcome Sanger Institute for the COVID-19 Genomics UK (COG-UK) Consortium | Jacquelyn Wynn et al |
| EPI_ISL_2570111 | Lighthouse Lab in Alderley Park | Wellcome Sanger Institute for the COVID-19 Genomics UK (COG-UK) Consortium | Jacquelyn Wynn et al |
| EPI_ISL_3729650 | Lighthouse Lab in Alderley Park | Wellcome Sanger Institute for the COVID-19 Genomics UK (COG-UK) Consortium | Jacquelyn Wynn et al |
| EPI_ISL_4406501 | Lighthouse Lab in Alderley Park | Wellcome Sanger Institute for the COVID-19 Genomics UK (COG-UK) Consortium | Jacquelyn Wynn et al |
| EPI_ISL_4618435 | Lighthouse Lab in Alderley Park | Wellcome Sanger Institute for the COVID-19 Genomics UK (COG-UK) Consortium | Jacquelyn Wynn et al |
| EPI_ISL_2235882 | Lighthouse Lab in Alderley Park | Wellcome Sanger Institute for the COVID-19 Genomics UK (COG-UK) Consortium | Jacquelyn Wynn et al |
| EPI_ISL_2394688 | Lighthouse Lab in Alderley Park | Wellcome Sanger Institute for the COVID-19 Genomics UK (COG-UK) Consortium | Jacquelyn Wynn et al |

| | | | |
|---|---|---|---|
| EPI_ISL_3358919 | Lighthouse Lab in Alderley Park | Wellcome Sanger Institute for the COVID-19 Genomics UK (COG-UK) Consortium | Jacquelyn Wynn et al |
| EPI_ISL_3338294 | Lighthouse Lab in Alderley Park | Wellcome Sanger Institute for the COVID-19 Genomics UK (COG-UK) Consortium | Jacquelyn Wynn et al |
| EPI_ISL_2606436 | Lighthouse Lab in Alderley Park | Wellcome Sanger Institute for the COVID-19 Genomics UK (COG-UK) Consortium | Jacquelyn Wynn et al |
| EPI_ISL_3580844 | Lighthouse Lab in Alderley Park | Wellcome Sanger Institute for the COVID-19 Genomics UK (COG-UK) Consortium | Jacquelyn Wynn et al |
| EPI_ISL_2766936 | Lighthouse Lab in Alderley Park | Wellcome Sanger Institute for the COVID-19 Genomics UK (COG-UK) Consortium | Jacquelyn Wynn et al |
| EPI_ISL_4048525 | Lighthouse Lab in Alderley Park | Wellcome Sanger Institute for the COVID-19 Genomics UK (COG-UK) Consortium | Jacquelyn Wynn et al |
| EPI_ISL_2911747 | Lighthouse Lab in Alderley Park | Wellcome Sanger Institute for the COVID-19 Genomics UK (COG-UK) Consortium | Jacquelyn Wynn et al |
| EPI_ISL_4018054 | Lighthouse Lab in Alderley Park | Wellcome Sanger Institute for the COVID-19 Genomics UK (COG-UK) Consortium | Jacquelyn Wynn et al |
| EPI_ISL_3335647 | Lighthouse Lab in Alderley Park | Wellcome Sanger Institute for the COVID-19 Genomics UK (COG-UK) Consortium | Jacquelyn Wynn et al |
| EPI_ISL_4324859 | Lighthouse Lab in Alderley Park | Wellcome Sanger Institute for the COVID-19 Genomics UK (COG-UK) Consortium | Jacquelyn Wynn et al |
| EPI_ISL_6152552 | Lighthouse Lab in Alderley Park | Wellcome Sanger Institute for the COVID-19 Genomics UK (COG-UK) Consortium | Jacquelyn Wynn et al |
| EPI_ISL_4896441 | Lighthouse Lab in Alderley Park | Wellcome Sanger Institute for the COVID-19 Genomics UK (COG-UK) Consortium | Jacquelyn Wynn et al |
| EPI_ISL_7023724 | Lighthouse Lab in Alderley Park | Wellcome Sanger Institute for the COVID-19 Genomics UK (COG-UK) Consortium | Jacquelyn Wynn et al |
| EPI_ISL_575451 | Lighthouse Lab in Alderley Park | Wellcome Sanger Institute for the COVID-19 Genomics UK (COG-UK) consortium | Jacquelyn Wynn et al |
| EPI_ISL_5038050 | Lighthouse Lab in Alderley Park | Wellcome Sanger Institute for the COVID-19 Genomics UK (COG-UK) Consortium | Jacquelyn Wynn et al |
| EPI_ISL_731167 | Lighthouse Lab in Alderley Park | Wellcome Sanger Institute for the COVID-19 Genomics UK (COG-UK) Consortium | Jacquelyn Wynn et al |
| EPI_ISL_5996827 | Lighthouse Lab in Alderley Park | Wellcome Sanger Institute for the COVID-19 Genomics UK (COG-UK) Consortium | Jacquelyn Wynn et al |

| | | | |
|---|---|---|---|
| EPI_ISL_610943 | Lighthouse Lab in Alderley Park | Wellcome Sanger Institute for the COVID-19 Genomics UK (COG-UK) consortium | Jacquelyn Wynn et al |
| EPI_ISL_5189896 | Lighthouse Lab in Alderley Park | Wellcome Sanger Institute for the COVID-19 Genomics UK (COG-UK) Consortium | Jacquelyn Wynn et al |
| EPI_ISL_554833 | Lighthouse Lab in Alderley Park | Wellcome Sanger Institute for the COVID-19 Genomics UK (COG-UK) consortium | The Lighthouse Lab in Alderley Park et al |
| EPI_ISL_7032043 | Lighthouse Lab in Alderley Park | Wellcome Sanger Institute for the COVID-19 Genomics UK (COG-UK) Consortium | Jacquelyn Wynn et al |
| EPI_ISL_6152754 | Lighthouse Lab in Alderley Park | Wellcome Sanger Institute for the COVID-19 Genomics UK (COG-UK) Consortium | Jacquelyn Wynn et al |
| EPI_ISL_6808668 | Lighthouse Lab in Alderley Park | Wellcome Sanger Institute for the COVID-19 Genomics UK (COG-UK) Consortium | Jacquelyn Wynn et al |
| EPI_ISL_4753433 | Lighthouse Lab in Alderley Park | Wellcome Sanger Institute for the COVID-19 Genomics UK (COG-UK) Consortium | Jacquelyn Wynn et al |
| EPI_ISL_5011537 | Lighthouse Lab in Alderley Park | Wellcome Sanger Institute for the COVID-19 Genomics UK (COG-UK) Consortium | Jacquelyn Wynn et al |
| EPI_ISL_4976169 | Lighthouse Lab in Alderley Park | Wellcome Sanger Institute for the COVID-19 Genomics UK (COG-UK) Consortium | Jacquelyn Wynn et al |
| EPI_ISL_610510 | Lighthouse Lab in Alderley Park | Wellcome Sanger Institute for the COVID-19 Genomics UK (COG-UK) consortium | Jacquelyn Wynn et al |
| EPI_ISL_799844 | Lighthouse Lab in Alderley Park | Wellcome Sanger Institute for the COVID-19 Genomics UK (COG-UK) Consortium | Jacquelyn Wynn et al |
| EPI_ISL_6767977 | Lighthouse Lab in Alderley Park | Wellcome Sanger Institute for the COVID-19 Genomics UK (COG-UK) Consortium | Jacquelyn Wynn et al |
| EPI_ISL_5255397 | Lighthouse Lab in Alderley Park | Wellcome Sanger Institute for the COVID-19 Genomics UK (COG-UK) Consortium | Jacquelyn Wynn et al |
| EPI_ISL_6766337 | Lighthouse Lab in Alderley Park | Wellcome Sanger Institute for the COVID-19 Genomics UK (COG-UK) Consortium | Jacquelyn Wynn et al |
| EPI_ISL_6613670 | Lighthouse Lab in Alderley Park | Wellcome Sanger Institute for the COVID-19 Genomics UK (COG-UK) Consortium | Jacquelyn Wynn et al |
| EPI_ISL_6364632 | Lighthouse Lab in Alderley Park | Wellcome Sanger Institute for the COVID-19 Genomics UK (COG-UK) Consortium | Jacquelyn Wynn et al |
| EPI_ISL_555979 | Lighthouse Lab in Alderley Park | Wellcome Sanger Institute for the COVID-19 Genomics UK (COG-UK) consortium | The Lighthouse Lab in Alderley Park et al |

| | | | |
|---|---|---|---|
| EPI_ISL_6766211 | Lighthouse Lab in Alderley Park | Wellcome Sanger Institute for the COVID-19 Genomics UK (COG-UK) Consortium | Jacquelyn Wynn et al |
| EPI_ISL_5752623 | Lighthouse Lab in Alderley Park | Wellcome Sanger Institute for the COVID-19 Genomics UK (COG-UK) Consortium | Jacquelyn Wynn et al |
| EPI_ISL_558332 | Lighthouse Lab in Alderley Park | Wellcome Sanger Institute for the COVID-19 Genomics UK (COG-UK) consortium | The Lighthouse Lab in Alderley Park et al |
| EPI_ISL_4899222 | Lighthouse Lab in Alderley Park | Wellcome Sanger Institute for the COVID-19 Genomics UK (COG-UK) Consortium | Jacquelyn Wynn et al |
| EPI_ISL_5038448 | Lighthouse Lab in Alderley Park | Wellcome Sanger Institute for the COVID-19 Genomics UK (COG-UK) Consortium | Jacquelyn Wynn et al |
| EPI_ISL_6359990 | Lighthouse Lab in Alderley Park | Wellcome Sanger Institute for the COVID-19 Genomics UK (COG-UK) Consortium | Jacquelyn Wynn et al |
| EPI_ISL_4899137 | Lighthouse Lab in Alderley Park | Wellcome Sanger Institute for the COVID-19 Genomics UK (COG-UK) Consortium | Jacquelyn Wynn et al |
| EPI_ISL_5269509 | Lighthouse Lab in Alderley Park | Wellcome Sanger Institute for the COVID-19 Genomics UK (COG-UK) Consortium | Jacquelyn Wynn et al |
| EPI_ISL_5825807 | Lighthouse Lab in Alderley Park | Wellcome Sanger Institute for the COVID-19 Genomics UK (COG-UK) Consortium | Jacquelyn Wynn et al |
| EPI_ISL_555321 | Lighthouse Lab in Alderley Park | Wellcome Sanger Institute for the COVID-19 Genomics UK (COG-UK) consortium | The Lighthouse Lab in Alderley Park et al |
| EPI_ISL_4762953 | Lighthouse Lab in Alderley Park | Wellcome Sanger Institute for the COVID-19 Genomics UK (COG-UK) Consortium | Jacquelyn Wynn et al |
| EPI_ISL_822180 | Lighthouse Lab in Alderley Park | Wellcome Sanger Institute for the COVID-19 Genomics UK (COG-UK) Consortium | Jacquelyn Wynn et al |
| EPI_ISL_6805170 | Lighthouse Lab in Alderley Park | Wellcome Sanger Institute for the COVID-19 Genomics UK (COG-UK) Consortium | Jacquelyn Wynn et al |
| EPI_ISL_5024467 | Lighthouse Lab in Alderley Park | Wellcome Sanger Institute for the COVID-19 Genomics UK (COG-UK) Consortium | Jacquelyn Wynn et al |
| EPI_ISL_655589 | Lighthouse Lab in Alderley Park | Wellcome Sanger Institute for the COVID-19 Genomics UK (COG-UK) Consortium | Jacquelyn Wynn et al |
| EPI_ISL_760726 | Lighthouse Lab in Alderley Park | Wellcome Sanger Institute for the COVID-19 Genomics UK (COG-UK) Consortium | Jacquelyn Wynn et al |
| EPI_ISL_6760268 | Lighthouse Lab in Alderley Park | Wellcome Sanger Institute for the COVID-19 Genomics UK (COG-UK) Consortium | Jacquelyn Wynn et al |

| | | | |
|---|---|---|---|
| EPI_ISL_6008142 | Lighthouse Lab in Alderley Park | Wellcome Sanger Institute for the COVID-19 Genomics UK (COG-UK) Consortium | Jacquelyn Wynn et al |
| EPI_ISL_588002 | Lighthouse Lab in Alderley Park | Wellcome Sanger Institute for the COVID-19 Genomics UK (COG-UK) consortium | Jacquelyn Wynn et al |
| EPI_ISL_5949362 | Lighthouse Lab in Alderley Park | Wellcome Sanger Institute for the COVID-19 Genomics UK (COG-UK) Consortium | Jacquelyn Wynn et al |
| EPI_ISL_5527046 | Lighthouse Lab in Alderley Park | Wellcome Sanger Institute for the COVID-19 Genomics UK (COG-UK) Consortium | Jacquelyn Wynn et al |
| EPI_ISL_5920445 | Lighthouse Lab in Alderley Park | Wellcome Sanger Institute for the COVID-19 Genomics UK (COG-UK) Consortium | Jacquelyn Wynn et al |
| EPI_ISL_5992737 | Lighthouse Lab in Alderley Park | Wellcome Sanger Institute for the COVID-19 Genomics UK (COG-UK) Consortium | Jacquelyn Wynn et al |
| EPI_ISL_624217 | Lighthouse Lab in Alderley Park | Wellcome Sanger Institute for the COVID-19 Genomics UK (COG-UK) consortium | Jacquelyn Wynn et al |
| EPI_ISL_586985 | Lighthouse Lab in Alderley Park | Wellcome Sanger Institute for the COVID-19 Genomics UK (COG-UK) consortium | Jacquelyn Wynn et al |
| EPI_ISL_630510 | Lighthouse Lab in Alderley Park | Wellcome Sanger Institute for the COVID-19 Genomics UK (COG-UK) consortium | Jacquelyn Wynn et al |
| EPI_ISL_5473232 | Lighthouse Lab in Alderley Park | Wellcome Sanger Institute for the COVID-19 Genomics UK (COG-UK) Consortium | Jacquelyn Wynn et al |
| EPI_ISL_6821008 | Lighthouse Lab in Alderley Park | Wellcome Sanger Institute for the COVID-19 Genomics UK (COG-UK) Consortium | Jacquelyn Wynn et al |
| EPI_ISL_6298866 | Lighthouse Lab in Alderley Park | Wellcome Sanger Institute for the COVID-19 Genomics UK (COG-UK) Consortium | Jacquelyn Wynn et al |
| EPI_ISL_630636 | Lighthouse Lab in Alderley Park | Wellcome Sanger Institute for the COVID-19 Genomics UK (COG-UK) consortium | Jacquelyn Wynn et al |
| EPI_ISL_6552401 | Lighthouse Lab in Alderley Park | Wellcome Sanger Institute for the COVID-19 Genomics UK (COG-UK) Consortium | Jacquelyn Wynn et al |
| EPI_ISL_6364228 | Lighthouse Lab in Alderley Park | Wellcome Sanger Institute for the COVID-19 Genomics UK (COG-UK) Consortium | Jacquelyn Wynn et al |
| EPI_ISL_798969 | Lighthouse Lab in Alderley Park | Wellcome Sanger Institute for the COVID-19 Genomics UK (COG-UK) Consortium | Jacquelyn Wynn et al |
| EPI_ISL_5350753 | Lighthouse Lab in Alderley Park | Wellcome Sanger Institute for the COVID-19 Genomics UK (COG-UK) Consortium | Jacquelyn Wynn et al |

| | | | |
|---|---|---|---|
| EPI_ISL_5244334 | Lighthouse Lab in Alderley Park | Wellcome Sanger Institute for the COVID-19 Genomics UK (COG-UK) Consortium | Jacquelyn Wynn et al |
| EPI_ISL_5012244 | Lighthouse Lab in Alderley Park | Wellcome Sanger Institute for the COVID-19 Genomics UK (COG-UK) Consortium | Jacquelyn Wynn et al |
| EPI_ISL_5256515 | Lighthouse Lab in Alderley Park | Wellcome Sanger Institute for the COVID-19 Genomics UK (COG-UK) Consortium | Jacquelyn Wynn et al |
| EPI_ISL_821987 | Lighthouse Lab in Alderley Park | Wellcome Sanger Institute for the COVID-19 Genomics UK (COG-UK) Consortium | Jacquelyn Wynn et al |
| EPI_ISL_5123603 | Lighthouse Lab in Alderley Park | Wellcome Sanger Institute for the COVID-19 Genomics UK (COG-UK) Consortium | Jacquelyn Wynn et al |
| EPI_ISL_6352381 | Lighthouse Lab in Alderley Park | Wellcome Sanger Institute for the COVID-19 Genomics UK (COG-UK) Consortium | Jacquelyn Wynn et al |
| EPI_ISL_575490 | Lighthouse Lab in Alderley Park | Wellcome Sanger Institute for the COVID-19 Genomics UK (COG-UK) consortium | Jacquelyn Wynn et al |
| EPI_ISL_567836 | Lighthouse Lab in Alderley Park | Wellcome Sanger Institute for the COVID-19 Genomics UK (COG-UK) consortium | Jacquelyn Wynn et al |
| EPI_ISL_4896647 | Lighthouse Lab in Alderley Park | Wellcome Sanger Institute for the COVID-19 Genomics UK (COG-UK) Consortium | Jacquelyn Wynn et al |
| EPI_ISL_553914 | Lighthouse Lab in Alderley Park | Wellcome Sanger Institute for the COVID-19 Genomics UK (COG-UK) consortium | The Lighthouse Lab in Alderley Park et al |
| EPI_ISL_6356857 | Lighthouse Lab in Alderley Park | Wellcome Sanger Institute for the COVID-19 Genomics UK (COG-UK) Consortium | Jacquelyn Wynn et al |
| EPI_ISL_6577890 | Lighthouse Lab in Alderley Park | Wellcome Sanger Institute for the COVID-19 Genomics UK (COG-UK) Consortium | Jacquelyn Wynn et al |
| EPI_ISL_6820713 | Lighthouse Lab in Alderley Park | Wellcome Sanger Institute for the COVID-19 Genomics UK (COG-UK) Consortium | Jacquelyn Wynn et al |
| EPI_ISL_5192680 | Lighthouse Lab in Alderley Park | Wellcome Sanger Institute for the COVID-19 Genomics UK (COG-UK) Consortium | Jacquelyn Wynn et al |
| EPI_ISL_6114069 | Lighthouse Lab in Alderley Park | Wellcome Sanger Institute for the COVID-19 Genomics UK (COG-UK) Consortium | Jacquelyn Wynn et al |
| EPI_ISL_5451712 | Lighthouse Lab in Alderley Park | Wellcome Sanger Institute for the COVID-19 Genomics UK (COG-UK) Consortium | Jacquelyn Wynn et al |
| EPI_ISL_6559398 | Lighthouse Lab in Alderley Park | Wellcome Sanger Institute for the COVID-19 Genomics UK (COG-UK) Consortium | Jacquelyn Wynn et al |

| | | | |
|---|---|---|---|
| EPI_ISL_5894532 | Lighthouse Lab in Alderley Park | Wellcome Sanger Institute for the COVID-19 Genomics UK (COG-UK) Consortium | Jacquelyn Wynn et al |
| EPI_ISL_6352638 | Lighthouse Lab in Alderley Park | Wellcome Sanger Institute for the COVID-19 Genomics UK (COG-UK) Consortium | Jacquelyn Wynn et al |
| EPI_ISL_833987 | Lighthouse Lab in Alderley Park | Wellcome Sanger Institute for the COVID-19 Genomics UK (COG-UK) Consortium | Jacquelyn Wynn et al |
| EPI_ISL_5449064 | Lighthouse Lab in Alderley Park | Wellcome Sanger Institute for the COVID-19 Genomics UK (COG-UK) Consortium | Jacquelyn Wynn et al |
| EPI_ISL_761723 | Lighthouse Lab in Alderley Park | Wellcome Sanger Institute for the COVID-19 Genomics UK (COG-UK) Consortium | Jacquelyn Wynn et al |
| EPI_ISL_882352 | Lighthouse Lab in Alderley Park | Wellcome Sanger Institute for the COVID-19 Genomics UK (COG-UK) Consortium | Jacquelyn Wynn et al |
| EPI_ISL_606186 | Lighthouse Lab in Alderley Park | Wellcome Sanger Institute for the COVID-19 Genomics UK (COG-UK) consortium | Jacquelyn Wynn et al |
| EPI_ISL_731018 | Lighthouse Lab in Alderley Park | Wellcome Sanger Institute for the COVID-19 Genomics UK (COG-UK) Consortium | Jacquelyn Wynn et al |
| EPI_ISL_5352472 | Lighthouse Lab in Alderley Park | Wellcome Sanger Institute for the COVID-19 Genomics UK (COG-UK) Consortium | Jacquelyn Wynn et al |
| EPI_ISL_7031670 | Lighthouse Lab in Alderley Park | Wellcome Sanger Institute for the COVID-19 Genomics UK (COG-UK) Consortium | Jacquelyn Wynn et al |
| EPI_ISL_624068 | Lighthouse Lab in Alderley Park | Wellcome Sanger Institute for the COVID-19 Genomics UK (COG-UK) consortium | Jacquelyn Wynn et al |
| EPI_ISL_6301736 | Lighthouse Lab in Alderley Park | Wellcome Sanger Institute for the COVID-19 Genomics UK (COG-UK) Consortium | Jacquelyn Wynn et al |
| EPI_ISL_6366238 | Lighthouse Lab in Alderley Park | Wellcome Sanger Institute for the COVID-19 Genomics UK (COG-UK) Consortium | Jacquelyn Wynn et al |
| EPI_ISL_4763074 | Lighthouse Lab in Alderley Park | Wellcome Sanger Institute for the COVID-19 Genomics UK (COG-UK) Consortium | Jacquelyn Wynn et al |
| EPI_ISL_557031 | Lighthouse Lab in Alderley Park | Wellcome Sanger Institute for the COVID-19 Genomics UK (COG-UK) consortium | The Lighthouse Lab in Alderley Park et al |
| EPI_ISL_4753400 | Lighthouse Lab in Alderley Park | Wellcome Sanger Institute for the COVID-19 Genomics UK (COG-UK) Consortium | Jacquelyn Wynn et al |
| EPI_ISL_5954212 | Lighthouse Lab in Alderley Park | Wellcome Sanger Institute for the COVID-19 Genomics UK (COG-UK) Consortium | Jacquelyn Wynn et al |

| | | | |
|---|---|---|---|
| EPI_ISL_4754184 | Lighthouse Lab in Alderley Park | Wellcome Sanger Institute for the COVID-19 Genomics UK (COG-UK) Consortium | Jacquelyn Wynn et al |
| EPI_ISL_5256696 | Lighthouse Lab in Alderley Park | Wellcome Sanger Institute for the COVID-19 Genomics UK (COG-UK) Consortium | Jacquelyn Wynn et al |
| EPI_ISL_4976203 | Lighthouse Lab in Alderley Park | Wellcome Sanger Institute for the COVID-19 Genomics UK (COG-UK) Consortium | Jacquelyn Wynn et al |
| EPI_ISL_4894590 | Lighthouse Lab in Alderley Park | Wellcome Sanger Institute for the COVID-19 Genomics UK (COG-UK) Consortium | Jacquelyn Wynn et al |
| EPI_ISL_4893767 | Lighthouse Lab in Alderley Park | Wellcome Sanger Institute for the COVID-19 Genomics UK (COG-UK) Consortium | Jacquelyn Wynn et al |
| EPI_ISL_6348385 | Lighthouse Lab in Alderley Park | Wellcome Sanger Institute for the COVID-19 Genomics UK (COG-UK) Consortium | Jacquelyn Wynn et al |
| EPI_ISL_6916148 | Lighthouse Lab in Alderley Park | Wellcome Sanger Institute for the COVID-19 Genomics UK (COG-UK) Consortium | Jacquelyn Wynn et al |
| EPI_ISL_5034900 | Lighthouse Lab in Alderley Park | Wellcome Sanger Institute for the COVID-19 Genomics UK (COG-UK) Consortium | Jacquelyn Wynn et al |
| EPI_ISL_5036561 | Lighthouse Lab in Alderley Park | Wellcome Sanger Institute for the COVID-19 Genomics UK (COG-UK) Consortium | Jacquelyn Wynn et al |
| EPI_ISL_6039227 | Lighthouse Lab in Alderley Park | Wellcome Sanger Institute for the COVID-19 Genomics UK (COG-UK) Consortium | Jacquelyn Wynn et al |
| EPI_ISL_6215416 | Lighthouse Lab in Alderley Park | Wellcome Sanger Institute for the COVID-19 Genomics UK (COG-UK) Consortium | Jacquelyn Wynn et al |
| EPI_ISL_625068 | Lighthouse Lab in Alderley Park | Wellcome Sanger Institute for the COVID-19 Genomics UK (COG-UK) consortium | Jacquelyn Wynn et al |
| EPI_ISL_6223882 | Lighthouse Lab in Alderley Park | Wellcome Sanger Institute for the COVID-19 Genomics UK (COG-UK) Consortium | Jacquelyn Wynn et al |
| EPI_ISL_655382 | Lighthouse Lab in Alderley Park | Wellcome Sanger Institute for the COVID-19 Genomics UK (COG-UK) Consortium | Jacquelyn Wynn et al |
| EPI_ISL_6929667 | Lighthouse Lab in Alderley Park | Wellcome Sanger Institute for the COVID-19 Genomics UK (COG-UK) Consortium | Jacquelyn Wynn et al |
| EPI_ISL_799891 | Lighthouse Lab in Alderley Park | Wellcome Sanger Institute for the COVID-19 Genomics UK (COG-UK) Consortium | Jacquelyn Wynn et al |
| EPI_ISL_567722 | Lighthouse Lab in Alderley Park | Wellcome Sanger Institute for the COVID-19 Genomics UK (COG-UK) consortium | Jacquelyn Wynn et al |

| | | | |
|---|---|---|---|
| EPI_ISL_6591521 | Berkshire and Surrey Pathology Services Lighthouse Laboratory | Wellcome Sanger Institute for the COVID-19 Genomics UK (COG-UK) Consortium | Berkshire et al |
| EPI_ISL_1018726 | Lighthouse Lab in Cambridge | Wellcome Sanger Institute for the COVID-19 Genomics UK (COG-UK) Consortium | Rob Howes et al |
| EPI_ISL_2917649 | Berkshire and Surrey Pathology Services Lighthouse Laboratory | Wellcome Sanger Institute for the COVID-19 Genomics UK (COG-UK) Consortium | Berkshire et al |
| EPI_ISL_1033371 | Lighthouse Lab in Cambridge | Wellcome Sanger Institute for the COVID-19 Genomics UK (COG-UK) Consortium | Rob Howes et al |
| EPI_ISL_4085908 | Berkshire and Surrey Pathology Services Lighthouse Laboratory | Wellcome Sanger Institute for the COVID-19 Genomics UK (COG-UK) Consortium | Berkshire et al |
| EPI_ISL_439893 | Department of Pathology, University of Cambridge | Wellcome Sanger Institute for the COVID-19 Genomics UK (COG-UK) consortium | Luke W Meredith et al |
| EPI_ISL_4323207 | Berkshire and Surrey Pathology Services Lighthouse Laboratory | Wellcome Sanger Institute for the COVID-19 Genomics UK (COG-UK) Consortium | Berkshire et al |
| EPI_ISL_986601 | Lighthouse Lab in Cambridge | Wellcome Sanger Institute for the COVID-19 Genomics UK (COG-UK) Consortium | Rob Howes et al |
| EPI_ISL_3000847 | Department of Pathology, University of Cambridge | COVID-19 Genomics UK (COG-UK) Consortium | Aminu S. Jahun et al |
| EPI_ISL_6833293 | Berkshire and Surrey Pathology Services Lighthouse Laboratory | Wellcome Sanger Institute for the COVID-19 Genomics UK (COG-UK) Consortium | Berkshire et al |
| EPI_ISL_1187208 | Lighthouse Lab in Cambridge | Wellcome Sanger Institute for the COVID-19 Genomics UK (COG-UK) Consortium | Rob Howes et al |
| EPI_ISL_5350774 | Berkshire and Surrey Pathology Services Lighthouse Laboratory | Wellcome Sanger Institute for the COVID-19 Genomics UK (COG-UK) Consortium | Berkshire et al |
| EPI_ISL_584360 | Virology Department, Sheffield Teaching Hospitals NHS Foundation Trust/Department of Infection, Immunity and Cardiovascular Disease, The Medical School, University of Sheffield | COVID-19 Genomics UK (COG-UK) Consortium | Thushan de Silva et al |
| EPI_ISL_609160 | Lighthouse Lab in Cambridge | Wellcome Sanger Institute for the COVID-19 Genomics UK (COG-UK) consortium | Rob Howes et al |
| EPI_ISL_4525084 | Berkshire and Surrey Pathology Services Lighthouse Laboratory | Wellcome Sanger Institute for the COVID-19 Genomics UK (COG-UK) Consortium | Berkshire et al |
| EPI_ISL_1042074 | Lighthouse Lab in Cambridge | Wellcome Sanger Institute for the COVID-19 Genomics UK (COG-UK) Consortium | Rob Howes et al |

| | | | |
|---|---|---|---|
| EPI_ISL_3700455 | Berkshire and Surrey Pathology Services Lighthouse Laboratory | Wellcome Sanger Institute for the COVID-19 Genomics UK (COG-UK) Consortium | Berkshire et al |
| EPI_ISL_623679 | Lighthouse Lab in Cambridge | Wellcome Sanger Institute for the COVID-19 Genomics UK (COG-UK) consortium | Rob Howes et al |
| EPI_ISL_6303844 | Berkshire and Surrey Pathology Services Lighthouse Laboratory | Wellcome Sanger Institute for the COVID-19 Genomics UK (COG-UK) Consortium | Berkshire et al |
| EPI_ISL_3027735 | Berkshire and Surrey Pathology Services Lighthouse Laboratory | Wellcome Sanger Institute for the COVID-19 Genomics UK (COG-UK) Consortium | Berkshire et al |
| EPI_ISL_3227942 | Berkshire and Surrey Pathology Services Lighthouse Laboratory | Wellcome Sanger Institute for the COVID-19 Genomics UK (COG-UK) Consortium | Berkshire et al |
| EPI_ISL_441640 | Department of Pathology, University of Cambridge | Wellcome Sanger Institute for the COVID-19 Genomics UK (COG-UK) consortium | Luke W Meredith et al |
| EPI_ISL_4500856 | Berkshire and Surrey Pathology Services Lighthouse Laboratory | Wellcome Sanger Institute for the COVID-19 Genomics UK (COG-UK) Consortium | Berkshire et al |
| EPI_ISL_837964 | Department of Pathology, University of Cambridge | COVID-19 Genomics UK (COG-UK) Consortium | Aminu S. Jahun et al |
| EPI_ISL_5334661 | Berkshire and Surrey Pathology Services Lighthouse Laboratory | Wellcome Sanger Institute for the COVID-19 Genomics UK (COG-UK) Consortium | Berkshire et al |
| EPI_ISL_5274386 | Berkshire and Surrey Pathology Services Lighthouse Laboratory | Wellcome Sanger Institute for the COVID-19 Genomics UK (COG-UK) Consortium | Berkshire et al |
| EPI_ISL_6927601 | Berkshire and Surrey Pathology Services Lighthouse Laboratory | Wellcome Sanger Institute for the COVID-19 Genomics UK (COG-UK) Consortium | Berkshire et al |
| EPI_ISL_5350552 | Berkshire and Surrey Pathology Services Lighthouse Laboratory | Wellcome Sanger Institute for the COVID-19 Genomics UK (COG-UK) Consortium | Berkshire et al |
| EPI_ISL_3641935 | Berkshire and Surrey Pathology Services Lighthouse Laboratory | Wellcome Sanger Institute for the COVID-19 Genomics UK (COG-UK) Consortium | Berkshire et al |
| EPI_ISL_672933 | Lighthouse Lab in Cambridge | Wellcome Sanger Institute for the COVID-19 Genomics UK (COG-UK) Consortium | Rob Howes et al |
| EPI_ISL_609347 | Lighthouse Lab in Cambridge | Wellcome Sanger Institute for the COVID-19 Genomics UK (COG-UK) consortium | Rob Howes et al |
| EPI_ISL_473498 | Department of Pathology, University of Cambridge | COVID-19 Genomics UK (COG-UK) Consortium | Luke W Meredith et al |
| EPI_ISL_6618951 | Berkshire and Surrey Pathology Services Lighthouse Laboratory | Wellcome Sanger Institute for the COVID-19 Genomics UK (COG-UK) Consortium | Berkshire et al |
| EPI_ISL_1062416 | Lighthouse Lab in Cambridge | Wellcome Sanger Institute for the COVID-19 Genomics UK (COG-UK) Consortium | Rob Howes et al |

| | | | |
|---|---|---|---|
| EPI_ISL_1246217 | Lighthouse Lab in Cambridge | Wellcome Sanger Institute for the COVID-19 Genomics UK (COG-UK) Consortium | Rob Howes et al |
| EPI_ISL_501587 | PHE South West Regional Laboratory, National Infection Service | Wellcome Sanger Institute for the COVID-19 Genomics UK (COG-UK) consortium | Stephanie Hutchings et al |
| EPI_ISL_3956522 | Berkshire and Surrey Pathology Services Lighthouse Laboratory | Wellcome Sanger Institute for the COVID-19 Genomics UK (COG-UK) Consortium | Berkshire et al |
| EPI_ISL_6926355 | Berkshire and Surrey Pathology Services Lighthouse Laboratory | Wellcome Sanger Institute for the COVID-19 Genomics UK (COG-UK) Consortium | Berkshire et al |
| EPI_ISL_4452129 | Berkshire and Surrey Pathology Services Lighthouse Laboratory | Wellcome Sanger Institute for the COVID-19 Genomics UK (COG-UK) Consortium | Berkshire et al |
| EPI_ISL_3437336 | Berkshire and Surrey Pathology Services Lighthouse Laboratory | Wellcome Sanger Institute for the COVID-19 Genomics UK (COG-UK) Consortium | Berkshire et al |
| EPI_ISL_3285485 | Berkshire and Surrey Pathology Services Lighthouse Laboratory | Wellcome Sanger Institute for the COVID-19 Genomics UK (COG-UK) Consortium | Berkshire et al |
| EPI_ISL_1189960 | Lighthouse Lab in Cambridge | Wellcome Sanger Institute for the COVID-19 Genomics UK (COG-UK) Consortium | Rob Howes et al |
| EPI_ISL_1489503 | Lighthouse Lab in Cambridge | Wellcome Sanger Institute for the COVID-19 Genomics UK (COG-UK) Consortium | Rob Howes et al |
| EPI_ISL_986500 | Lighthouse Lab in Cambridge | Wellcome Sanger Institute for the COVID-19 Genomics UK (COG-UK) Consortium | Rob Howes et al |
| EPI_ISL_651578 | Department of Pathology, University of Cambridge | COVID-19 Genomics UK (COG-UK) Consortium | Aminu S. Jahun et al |
| EPI_ISL_881849 | Lighthouse Lab in Alderley Park | Wellcome Sanger Institute for the COVID-19 Genomics UK (COG-UK) Consortium | Jacquelyn Wynn et al |
| EPI_ISL_6769735 | Berkshire and Surrey Pathology Services Lighthouse Laboratory | Wellcome Sanger Institute for the COVID-19 Genomics UK (COG-UK) Consortium | Berkshire et al |
| EPI_ISL_6915404 | Berkshire and Surrey Pathology Services Lighthouse Laboratory | Wellcome Sanger Institute for the COVID-19 Genomics UK (COG-UK) Consortium | Berkshire et al |
| EPI_ISL_1007456 | Lighthouse Lab in Cambridge | Wellcome Sanger Institute for the COVID-19 Genomics UK (COG-UK) Consortium | Rob Howes et al |
| EPI_ISL_610660 | Lighthouse Lab in Cambridge | Wellcome Sanger Institute for the COVID-19 Genomics UK (COG-UK) consortium | Rob Howes et al |
| EPI_ISL_442957 | Department of Pathology, University of Cambridge | Wellcome Sanger Institute for the COVID-19 Genomics UK (COG-UK) consortium | Luke W Meredith et al |

| | | | |
|---|---|---|---|
| EPI_ISL_4500071 | Berkshire and Surrey Pathology Services Lighthouse Laboratory | Wellcome Sanger Institute for the COVID-19 Genomics UK (COG-UK) Consortium | Berkshire et al |
| EPI_ISL_1175190 | Lighthouse Lab in Cambridge | Wellcome Sanger Institute for the COVID-19 Genomics UK (COG-UK) Consortium | Rob Howes et al |
| EPI_ISL_3954117 | Berkshire and Surrey Pathology Services Lighthouse Laboratory | Wellcome Sanger Institute for the COVID-19 Genomics UK (COG-UK) Consortium | Berkshire et al |
| EPI_ISL_3249434 | Berkshire and Surrey Pathology Services Lighthouse Laboratory | Wellcome Sanger Institute for the COVID-19 Genomics UK (COG-UK) Consortium | Berkshire et al |
| EPI_ISL_2815836 | Department of Pathology, University of Cambridge | COVID-19 Genomics UK (COG-UK) Consortium | Aminu S. Jahun et al |
| EPI_ISL_3051703 | Berkshire and Surrey Pathology Services Lighthouse Laboratory | Wellcome Sanger Institute for the COVID-19 Genomics UK (COG-UK) Consortium | Berkshire et al |
| EPI_ISL_3377041 | Berkshire and Surrey Pathology Services Lighthouse Laboratory | Wellcome Sanger Institute for the COVID-19 Genomics UK (COG-UK) Consortium | Berkshire et al |
| EPI_ISL_4525058 | Berkshire and Surrey Pathology Services Lighthouse Laboratory | Wellcome Sanger Institute for the COVID-19 Genomics UK (COG-UK) Consortium | Berkshire et al |
| EPI_ISL_1653315 | Lighthouse Lab in Cambridge | Wellcome Sanger Institute for the COVID-19 Genomics UK (COG-UK) Consortium | Rob Howes et al |
| EPI_ISL_4500847 | Berkshire and Surrey Pathology Services Lighthouse Laboratory | Wellcome Sanger Institute for the COVID-19 Genomics UK (COG-UK) Consortium | Berkshire et al |
| EPI_ISL_4452107 | Berkshire and Surrey Pathology Services Lighthouse Laboratory | Wellcome Sanger Institute for the COVID-19 Genomics UK (COG-UK) Consortium | Berkshire et al |
| EPI_ISL_6219836 | Berkshire and Surrey Pathology Services Lighthouse Laboratory | Wellcome Sanger Institute for the COVID-19 Genomics UK (COG-UK) Consortium | Berkshire et al |
| EPI_ISL_469501 | PHE South West Regional Laboratory, National Infection Service | Wellcome Sanger Institute for the COVID-19 Genomics UK (COG-UK) consortium | Stephanie Hutchings et al |
| EPI_ISL_1061498 | Lighthouse Lab in Cambridge | Wellcome Sanger Institute for the COVID-19 Genomics UK (COG-UK) Consortium | Rob Howes et al |
| EPI_ISL_469728 | PHE South West Regional Laboratory, National Infection Service | Wellcome Sanger Institute for the COVID-19 Genomics UK (COG-UK) consortium | Stephanie Hutchings et al |
| EPI_ISL_3442834 | Berkshire and Surrey Pathology Services Lighthouse Laboratory | Wellcome Sanger Institute for the COVID-19 Genomics UK (COG-UK) Consortium | Berkshire et al |
| EPI_ISL_1224573 | Lighthouse Lab in Cambridge | Wellcome Sanger Institute for the COVID-19 Genomics UK (COG-UK) Consortium | Rob Howes et al |

| | | | |
|---|---|---|---|
| EPI_ISL_4453934 | Berkshire and Surrey Pathology Services Lighthouse Laboratory | Wellcome Sanger Institute for the COVID-19 Genomics UK (COG-UK) Consortium | Berkshire et al |
| EPI_ISL_3367248 | Berkshire and Surrey Pathology Services Lighthouse Laboratory | Wellcome Sanger Institute for the COVID-19 Genomics UK (COG-UK) Consortium | Berkshire et al |
| EPI_ISL_4478591 | Berkshire and Surrey Pathology Services Lighthouse Laboratory | Wellcome Sanger Institute for the COVID-19 Genomics UK (COG-UK) Consortium | Berkshire et al |
| EPI_ISL_1206343 | Lighthouse Lab in Cambridge | Wellcome Sanger Institute for the COVID-19 Genomics UK (COG-UK) Consortium | Rob Howes et al |
| EPI_ISL_6926967 | Berkshire and Surrey Pathology Services Lighthouse Laboratory | Wellcome Sanger Institute for the COVID-19 Genomics UK (COG-UK) Consortium | Berkshire et al |
| EPI_ISL_873573 | Lighthouse Lab in Cambridge | Wellcome Sanger Institute for the COVID-19 Genomics UK (COG-UK) Consortium | Rob Howes et al |
| EPI_ISL_1103766 | University of Exeter | COVID-19 Genomics UK (COG-UK) Consortium | Ben Temperton et al |
| EPI_ISL_492572 | PHE South West Regional Laboratory, National Infection Service | Wellcome Sanger Institute for the COVID-19 Genomics UK (COG-UK) consortium | Stephanie Hutchings et al |
| EPI_ISL_988453 | Lighthouse Lab in Milton Keynes | Wellcome Sanger Institute for the COVID-19 Genomics UK (COG-UK) Consortium | The Lighthouse Lab in Milton Keynes et al |
| EPI_ISL_945986 | Lighthouse Lab in Milton Keynes | Wellcome Sanger Institute for the COVID-19 Genomics UK (COG-UK) Consortium | The Lighthouse Lab in Milton Keynes et al |
| EPI_ISL_1177843 | Liverpool Clinical Laboratories | COVID-19 Genomics UK (COG-UK) Consortium | Sam Haldenby et al |
| EPI_ISL_1083006 | Lighthouse Lab in Milton Keynes | Wellcome Sanger Institute for the COVID-19 Genomics UK (COG-UK) Consortium | The Lighthouse Lab in Milton Keynes et al |
| EPI_ISL_1697565 | Lighthouse Lab in Milton Keynes | Wellcome Sanger Institute for the COVID-19 Genomics UK (COG-UK) Consortium | The Lighthouse Lab in Milton Keynes et al |
| EPI_ISL_705007 | University College London, Great Ormond Street Hospital for Children NHS Foundation Trust, Imperial College Healthcare NHS Trust | COVID-19 Genomics UK (COG-UK) Consortium | Sergi Castellano et al |
| EPI_ISL_838455 | Liverpool Clinical Laboratories | COVID-19 Genomics UK (COG-UK) Consortium | Sam Haldenby et al |
| EPI_ISL_1387017 | Quadram Institute Bioscience | COVID-19 Genomics UK (COG-UK) Consortium | Dave J. Baker et al |
| EPI_ISL_2253830 | Lighthouse Lab in Milton Keynes | Wellcome Sanger Institute for the COVID-19 Genomics UK (COG-UK) Consortium | The Lighthouse Lab in Milton Keynes et al |
| EPI_ISL_1921721 | Lighthouse Lab in Milton Keynes | Wellcome Sanger Institute for the COVID-19 Genomics UK (COG-UK) Consortium | The Lighthouse Lab in Milton Keynes et al |

| | | | |
|---|---|---|---|
| EPI_ISL_2355226 | Centre for Clinical Infection and Diagnostics Research and Genomics Innovation Unit, Guy's and St. Thomas' NHS Trust | COVID-19 Genomics UK (COG-UK) Consortium | Chloe Fisher et al |
| EPI_ISL_1248067 | University College London, Great Ormond Street Hospital for Children NHS Foundation Trust, Imperial College Healthcare NHS Trust | COVID-19 Genomics UK (COG-UK) Consortium | Sergi Castellano et al |
| EPI_ISL_920074 | University College London, Great Ormond Street Hospital for Children NHS Foundation Trust, Imperial College Healthcare NHS Trust | COVID-19 Genomics UK (COG-UK) Consortium | Sergi Castellano et al |
| EPI_ISL_1045775 | Lighthouse Lab in Milton Keynes | Wellcome Sanger Institute for the COVID-19 Genomics UK (COG-UK) Consortium | The Lighthouse Lab in Milton Keynes et al |
| EPI_ISL_1375315 | Lighthouse Lab in Milton Keynes | Wellcome Sanger Institute for the COVID-19 Genomics UK (COG-UK) Consortium | The Lighthouse Lab in Milton Keynes et al |
| EPI_ISL_3077874 | Liverpool Clinical Laboratories | COVID-19 Genomics UK (COG-UK) Consortium | Sam Haldenby et al |
| EPI_ISL_865529 | Barts Health NHS Trust | COVID-19 Genomics UK (COG-UK) Consortium | CUTINO-MOGUEL et al |
| EPI_ISL_647320 | Lighthouse Lab in Cambridge | Wellcome Sanger Institute for the COVID-19 Genomics UK (COG-UK) Consortium | Rob Howes et al |
| EPI_ISL_797581 | Lighthouse Lab in Cambridge | Wellcome Sanger Institute for the COVID-19 Genomics UK (COG-UK) Consortium | Rob Howes et al |
| EPI_ISL_958636 | Lighthouse Lab in Milton Keynes | Wellcome Sanger Institute for the COVID-19 Genomics UK (COG-UK) Consortium | The Lighthouse Lab in Milton Keynes et al |
| EPI_ISL_1637059 | University College London Hospital | COVID-19 Genomics UK (COG-UK) Consortium | Dr Judith Heaney et al |
| EPI_ISL_659281 | Lighthouse Lab in Cambridge | Wellcome Sanger Institute for the COVID-19 Genomics UK (COG-UK) Consortium | Rob Howes et al |
| EPI_ISL_704985 | Lighthouse Lab in Cambridge | Wellcome Sanger Institute for the COVID-19 Genomics UK (COG-UK) Consortium | Rob Howes et al |
| EPI_ISL_1187856 | Lighthouse Lab in Milton Keynes | Wellcome Sanger Institute for the COVID-19 Genomics UK (COG-UK) Consortium | The Lighthouse Lab in Milton Keynes et al |
| EPI_ISL_1486624 | Lighthouse Lab in Milton Keynes | Wellcome Sanger Institute for the COVID-19 Genomics UK (COG-UK) Consortium | The Lighthouse Lab in Milton Keynes et al |
| EPI_ISL_1536677 | Lighthouse Lab in Milton Keynes | Wellcome Sanger Institute for the COVID-19 Genomics UK (COG-UK) Consortium | The Lighthouse Lab in Milton Keynes et al |

| | | | |
|---|---|---|---|
| EPI_ISL_7144484 | Rosalind Franklin Laboratory | Wellcome Sanger Institute for the COVID-19 Genomics UK (COG-UK) Consortium | Donald Fraser et al |
| EPI_ISL_1537414 | Lighthouse Lab in Milton Keynes | Wellcome Sanger Institute for the COVID-19 Genomics UK (COG-UK) Consortium | The Lighthouse Lab in Milton Keynes et al |
| EPI_ISL_2118189 | Lighthouse Lab in Milton Keynes | Wellcome Sanger Institute for the COVID-19 Genomics UK (COG-UK) Consortium | The Lighthouse Lab in Milton Keynes et al |
| EPI_ISL_705404 | Liverpool Clinical Laboratories | COVID-19 Genomics UK (COG-UK) Consortium | Sam Haldenby et al |
| EPI_ISL_1207412 | Lighthouse Lab in Milton Keynes | Wellcome Sanger Institute for the COVID-19 Genomics UK (COG-UK) Consortium | The Lighthouse Lab in Milton Keynes et al |
| EPI_ISL_2237467 | Lighthouse Lab in Milton Keynes | Wellcome Sanger Institute for the COVID-19 Genomics UK (COG-UK) Consortium | The Lighthouse Lab in Milton Keynes et al |
| EPI_ISL_658321 | Lighthouse Lab in Cambridge | Wellcome Sanger Institute for the COVID-19 Genomics UK (COG-UK) Consortium | Rob Howes et al |
| EPI_ISL_1069435 | Lighthouse Lab in Milton Keynes | Wellcome Sanger Institute for the COVID-19 Genomics UK (COG-UK) Consortium | The Lighthouse Lab in Milton Keynes et al |
| EPI_ISL_920703 | University College London Hospital | COVID-19 Genomics UK (COG-UK) Consortium | Judith Heaney et al |
| EPI_ISL_865185 | Liverpool Clinical Laboratories | COVID-19 Genomics UK (COG-UK) Consortium | Sam Haldenby et al |
| EPI_ISL_444192 | University College London, Great Ormond Street Hospital for Children NHS Foundation Trust, Imperial College Healthcare NHS Trust | COVID-19 Genomics UK (COG-UK) Consortium | Sergi Castellano et al |
| EPI_ISL_1207006 | Lighthouse Lab in Milton Keynes | Wellcome Sanger Institute for the COVID-19 Genomics UK (COG-UK) Consortium | The Lighthouse Lab in Milton Keynes et al |
| EPI_ISL_718846 | Lighthouse Lab in Cambridge | Wellcome Sanger Institute for the COVID-19 Genomics UK (COG-UK) Consortium | Rob Howes et al |
| EPI_ISL_1245883 | Lighthouse Lab in Milton Keynes | Wellcome Sanger Institute for the COVID-19 Genomics UK (COG-UK) Consortium | The Lighthouse Lab in Milton Keynes et al |
| EPI_ISL_3958504 | University College London, Great Ormond Street Hospital for Children NHS Foundation Trust, Imperial College Healthcare NHS Trust | COVID-19 Genomics UK (COG-UK) Consortium | Sergi Castellano et al |
| EPI_ISL_7149316 | Rosalind Franklin Laboratory | Wellcome Sanger Institute for the COVID-19 Genomics UK (COG-UK) Consortium | Donald Fraser et al |
| EPI_ISL_2121305 | Lighthouse Lab in Milton Keynes | Wellcome Sanger Institute for the COVID-19 Genomics UK (COG-UK) Consortium | The Lighthouse Lab in Milton Keynes et al |

| | | | |
|---|---|---|---|
| EPI_ISL_1450987 | Lighthouse Lab in Milton Keynes | Wellcome Sanger Institute for the COVID-19 Genomics UK (COG-UK) Consortium | The Lighthouse Lab in Milton Keynes et al |
| EPI_ISL_1653416 | Lighthouse Lab in Milton Keynes | Wellcome Sanger Institute for the COVID-19 Genomics UK (COG-UK) Consortium | The Lighthouse Lab in Milton Keynes et al |
| EPI_ISL_2808617 | Health Services Laboratories | Wellcome Sanger Institute for the COVID-19 Genomics UK (COG-UK) Consortium | Health Services Laboratories et al |
| EPI_ISL_652084 | Liverpool Clinical Laboratories | COVID-19 Genomics UK (COG-UK) Consortium | Sam Haldenby et al |
| EPI_ISL_733629 | Lighthouse Lab in Cambridge | Wellcome Sanger Institute for the COVID-19 Genomics UK (COG-UK) Consortium | Rob Howes et al |
| EPI_ISL_991546 | Lighthouse Lab in Milton Keynes | Wellcome Sanger Institute for the COVID-19 Genomics UK (COG-UK) Consortium | The Lighthouse Lab in Milton Keynes et al |
| EPI_ISL_664752 | Virology Department, Sheffield Teaching Hospitals NHS Foundation Trust/Department of Infection, Immunity and Cardiovascular Disease, The Medical School, University of Sheffield | COVID-19 Genomics UK (COG-UK) Consortium | Thushan de Silva et al |
| EPI_ISL_5039415 | University College London, Great Ormond Street Hospital for Children NHS Foundation Trust, Imperial College Healthcare NHS Trust | COVID-19 Genomics UK (COG-UK) Consortium | Sergi Castellano et al |
| EPI_ISL_659274 | Lighthouse Lab in Cambridge | Wellcome Sanger Institute for the COVID-19 Genomics UK (COG-UK) Consortium | Rob Howes et al |
| EPI_ISL_2901148 | Liverpool Clinical Laboratories | COVID-19 Genomics UK (COG-UK) Consortium | Sam Haldenby et al |
| EPI_ISL_1986066 | Lighthouse Lab in Milton Keynes | Wellcome Sanger Institute for the COVID-19 Genomics UK (COG-UK) Consortium | The Lighthouse Lab in Milton Keynes et al |
| EPI_ISL_1033994 | Lighthouse Lab in Milton Keynes | Wellcome Sanger Institute for the COVID-19 Genomics UK (COG-UK) Consortium | The Lighthouse Lab in Milton Keynes et al |
| EPI_ISL_2704385 | Health Services Laboratories | Wellcome Sanger Institute for the COVID-19 Genomics UK (COG-UK) Consortium | Health Services Laboratories et al |
| EPI_ISL_3176665 | University College London, Great Ormond Street Hospital for Children NHS Foundation Trust, Imperial College Healthcare NHS Trust | COVID-19 Genomics UK (COG-UK) Consortium | Sergi Castellano et al |
| EPI_ISL_659307 | Lighthouse Lab in Cambridge | Wellcome Sanger Institute for the COVID-19 Genomics UK (COG-UK) Consortium | Rob Howes et al |

| | | | |
|---|---|---|---|
| EPI_ISL_611618 | University of Exeter | COVID-19 Genomics UK (COG-UK) Consortium | Ben Temperton et al |
| EPI_ISL_949685 | Liverpool Clinical Laboratories | COVID-19 Genomics UK (COG-UK) Consortium | Sam Haldenby et al |
| EPI_ISL_1326620 | Lighthouse Lab in Milton Keynes | Wellcome Sanger Institute for the COVID-19 Genomics UK (COG-UK) Consortium | The Lighthouse Lab in Milton Keynes et al |
| EPI_ISL_1187293 | Lighthouse Lab in Milton Keynes | Wellcome Sanger Institute for the COVID-19 Genomics UK (COG-UK) Consortium | The Lighthouse Lab in Milton Keynes et al |
| EPI_ISL_659292 | Lighthouse Lab in Cambridge | Wellcome Sanger Institute for the COVID-19 Genomics UK (COG-UK) Consortium | Rob Howes et al |
| EPI_ISL_3469080 | Health Services Laboratories | Wellcome Sanger Institute for the COVID-19 Genomics UK (COG-UK) Consortium | Health Services Laboratories et al |
| EPI_ISL_1084001 | Lighthouse Lab in Milton Keynes | Wellcome Sanger Institute for the COVID-19 Genomics UK (COG-UK) Consortium | The Lighthouse Lab in Milton Keynes et al |
| EPI_ISL_838826 | University College London, Great Ormond Street Hospital for Children NHS Foundation Trust, Imperial College Healthcare NHS Trust | COVID-19 Genomics UK (COG-UK) Consortium | Sergi Castellano et al |
| EPI_ISL_1274751 | Lighthouse Lab in Milton Keynes | Wellcome Sanger Institute for the COVID-19 Genomics UK (COG-UK) Consortium | The Lighthouse Lab in Milton Keynes et al |
| EPI_ISL_920416 | University College London Hospital | COVID-19 Genomics UK (COG-UK) Consortium | Judith Heaney et al |
| EPI_ISL_1099737 | Lighthouse Lab in Milton Keynes | Wellcome Sanger Institute for the COVID-19 Genomics UK (COG-UK) Consortium | The Lighthouse Lab in Milton Keynes et al |
| EPI_ISL_997747 | University College London, Great Ormond Street Hospital for Children NHS Foundation Trust, Imperial College Healthcare NHS Trust | COVID-19 Genomics UK (COG-UK) Consortium | Sergi Castellano et al |
| EPI_ISL_3580927 | Health Services Laboratories | Wellcome Sanger Institute for the COVID-19 Genomics UK (COG-UK) Consortium | Health Services Laboratories et al |
| EPI_ISL_499811 | Liverpool Clinical Laboratories | COVID-19 Genomics UK (COG-UK) Consortium | Sam Haldenby et al |
| EPI_ISL_2236602 | Lighthouse Lab in Milton Keynes | Wellcome Sanger Institute for the COVID-19 Genomics UK (COG-UK) Consortium | The Lighthouse Lab in Milton Keynes et al |
| EPI_ISL_6270401 | University College London, Great Ormond Street Hospital for Children NHS Foundation Trust, Imperial College Healthcare NHS Trust | COVID-19 Genomics UK (COG-UK) Consortium | Sergi Castellano et al |

| | | | |
|---|---|---|---|
| EPI_ISL_1071164 | Lighthouse Lab in Milton Keynes | Wellcome Sanger Institute for the COVID-19 Genomics UK (COG-UK) Consortium | The Lighthouse Lab in Milton Keynes et al |
| EPI_ISL_946015 | Lighthouse Lab in Milton Keynes | Wellcome Sanger Institute for the COVID-19 Genomics UK (COG-UK) Consortium | The Lighthouse Lab in Milton Keynes et al |
| EPI_ISL_1330738 | Lighthouse Lab in Milton Keynes | Wellcome Sanger Institute for the COVID-19 Genomics UK (COG-UK) Consortium | The Lighthouse Lab in Milton Keynes et al |
| EPI_ISL_2005676 | Lighthouse Lab in Milton Keynes | Wellcome Sanger Institute for the COVID-19 Genomics UK (COG-UK) Consortium | The Lighthouse Lab in Milton Keynes et al |
| EPI_ISL_3365325 | Lighthouse Lab in Milton Keynes | Wellcome Sanger Institute for the COVID-19 Genomics UK (COG-UK) Consortium | The Lighthouse Lab in Milton Keynes et al |
| EPI_ISL_3528980 | Lighthouse Lab in Milton Keynes | Wellcome Sanger Institute for the COVID-19 Genomics UK (COG-UK) Consortium | The Lighthouse Lab in Milton Keynes et al |
| EPI_ISL_3808747 | Lighthouse Lab in Milton Keynes | Wellcome Sanger Institute for the COVID-19 Genomics UK (COG-UK) Consortium | The Lighthouse Lab in Milton Keynes et al |
| EPI_ISL_3202907 | Lighthouse Lab in Milton Keynes | Wellcome Sanger Institute for the COVID-19 Genomics UK (COG-UK) Consortium | The Lighthouse Lab in Milton Keynes et al |
| EPI_ISL_2806529 | Lighthouse Lab in Milton Keynes | Wellcome Sanger Institute for the COVID-19 Genomics UK (COG-UK) Consortium | The Lighthouse Lab in Milton Keynes et al |
| EPI_ISL_3467464 | Lighthouse Lab in Milton Keynes | Wellcome Sanger Institute for the COVID-19 Genomics UK (COG-UK) Consortium | The Lighthouse Lab in Milton Keynes et al |
| EPI_ISL_4046635 | Lighthouse Lab in Milton Keynes | Wellcome Sanger Institute for the COVID-19 Genomics UK (COG-UK) Consortium | The Lighthouse Lab in Milton Keynes et al |
| EPI_ISL_4583826 | Lighthouse Lab in Milton Keynes | Wellcome Sanger Institute for the COVID-19 Genomics UK (COG-UK) Consortium | The Lighthouse Lab in Milton Keynes et al |
| EPI_ISL_3992202 | Lighthouse Lab in Milton Keynes | Wellcome Sanger Institute for the COVID-19 Genomics UK (COG-UK) Consortium | The Lighthouse Lab in Milton Keynes et al |
| EPI_ISL_4322766 | Lighthouse Lab in Milton Keynes | Wellcome Sanger Institute for the COVID-19 Genomics UK (COG-UK) Consortium | The Lighthouse Lab in Milton Keynes et al |
| EPI_ISL_4012771 | Lighthouse Lab in Milton Keynes | Wellcome Sanger Institute for the COVID-19 Genomics UK (COG-UK) Consortium | The Lighthouse Lab in Milton Keynes et al |
| EPI_ISL_2539510 | Lighthouse Lab in Milton Keynes | Wellcome Sanger Institute for the COVID-19 Genomics UK (COG-UK) Consortium | The Lighthouse Lab in Milton Keynes et al |
| EPI_ISL_4144556 | Lighthouse Lab in Milton Keynes | Wellcome Sanger Institute for the COVID-19 Genomics UK (COG-UK) Consortium | The Lighthouse Lab in Milton Keynes et al |

| | | | |
|---|---|---|---|
| EPI_ISL_3125626 | Lighthouse Lab in Milton Keynes | Wellcome Sanger Institute for the COVID-19 Genomics UK (COG-UK) Consortium | The Lighthouse Lab in Milton Keynes et al |
| EPI_ISL_3248876 | Lighthouse Lab in Milton Keynes | Wellcome Sanger Institute for the COVID-19 Genomics UK (COG-UK) Consortium | The Lighthouse Lab in Milton Keynes et al |
| EPI_ISL_4503028 | Lighthouse Lab in Milton Keynes | Wellcome Sanger Institute for the COVID-19 Genomics UK (COG-UK) Consortium | The Lighthouse Lab in Milton Keynes et al |
| EPI_ISL_4065229 | Lighthouse Lab in Milton Keynes | Wellcome Sanger Institute for the COVID-19 Genomics UK (COG-UK) Consortium | The Lighthouse Lab in Milton Keynes et al |
| EPI_ISL_4009730 | Lighthouse Lab in Milton Keynes | Wellcome Sanger Institute for the COVID-19 Genomics UK (COG-UK) Consortium | The Lighthouse Lab in Milton Keynes et al |
| EPI_ISL_2717177 | Lighthouse Lab in Milton Keynes | Wellcome Sanger Institute for the COVID-19 Genomics UK (COG-UK) Consortium | The Lighthouse Lab in Milton Keynes et al |
| EPI_ISL_2974226 | Lighthouse Lab in Milton Keynes | Wellcome Sanger Institute for the COVID-19 Genomics UK (COG-UK) Consortium | The Lighthouse Lab in Milton Keynes et al |
| EPI_ISL_4873297 | Lighthouse Lab in Milton Keynes | Wellcome Sanger Institute for the COVID-19 Genomics UK (COG-UK) Consortium | The Lighthouse Lab in Milton Keynes et al |
| EPI_ISL_3286281 | Lighthouse Lab in Milton Keynes | Wellcome Sanger Institute for the COVID-19 Genomics UK (COG-UK) Consortium | The Lighthouse Lab in Milton Keynes et al |
| EPI_ISL_4579111 | Lighthouse Lab in Milton Keynes | Wellcome Sanger Institute for the COVID-19 Genomics UK (COG-UK) Consortium | The Lighthouse Lab in Milton Keynes et al |
| EPI_ISL_2962783 | Lighthouse Lab in Milton Keynes | Wellcome Sanger Institute for the COVID-19 Genomics UK (COG-UK) Consortium | The Lighthouse Lab in Milton Keynes et al |
| EPI_ISL_3808583 | Lighthouse Lab in Milton Keynes | Wellcome Sanger Institute for the COVID-19 Genomics UK (COG-UK) Consortium | The Lighthouse Lab in Milton Keynes et al |
| EPI_ISL_3767535 | Lighthouse Lab in Milton Keynes | Wellcome Sanger Institute for the COVID-19 Genomics UK (COG-UK) Consortium | The Lighthouse Lab in Milton Keynes et al |
| EPI_ISL_2994498 | Lighthouse Lab in Milton Keynes | Wellcome Sanger Institute for the COVID-19 Genomics UK (COG-UK) Consortium | The Lighthouse Lab in Milton Keynes et al |
| EPI_ISL_2974345 | Lighthouse Lab in Milton Keynes | Wellcome Sanger Institute for the COVID-19 Genomics UK (COG-UK) Consortium | The Lighthouse Lab in Milton Keynes et al |
| EPI_ISL_3950458 | Lighthouse Lab in Milton Keynes | Wellcome Sanger Institute for the COVID-19 Genomics UK (COG-UK) Consortium | The Lighthouse Lab in Milton Keynes et al |
| EPI_ISL_3377620 | Lighthouse Lab in Milton Keynes | Wellcome Sanger Institute for the COVID-19 Genomics UK (COG-UK) Consortium | The Lighthouse Lab in Milton Keynes et al |

| | | | |
|---|---|---|---|
| EPI_ISL_3106501 | Lighthouse Lab in Milton Keynes | Wellcome Sanger Institute for the COVID-19 Genomics UK (COG-UK) Consortium | The Lighthouse Lab in Milton Keynes et al |
| EPI_ISL_3639738 | Lighthouse Lab in Milton Keynes | Wellcome Sanger Institute for the COVID-19 Genomics UK (COG-UK) Consortium | The Lighthouse Lab in Milton Keynes et al |
| EPI_ISL_3699657 | Lighthouse Lab in Milton Keynes | Wellcome Sanger Institute for the COVID-19 Genomics UK (COG-UK) Consortium | The Lighthouse Lab in Milton Keynes et al |
| EPI_ISL_4585531 | Lighthouse Lab in Milton Keynes | Wellcome Sanger Institute for the COVID-19 Genomics UK (COG-UK) Consortium | The Lighthouse Lab in Milton Keynes et al |
| EPI_ISL_4087210 | Lighthouse Lab in Milton Keynes | Wellcome Sanger Institute for the COVID-19 Genomics UK (COG-UK) Consortium | The Lighthouse Lab in Milton Keynes et al |
| EPI_ISL_4404359 | Lighthouse Lab in Milton Keynes | Wellcome Sanger Institute for the COVID-19 Genomics UK (COG-UK) Consortium | The Lighthouse Lab in Milton Keynes et al |
| EPI_ISL_2916497 | Lighthouse Lab in Milton Keynes | Wellcome Sanger Institute for the COVID-19 Genomics UK (COG-UK) Consortium | The Lighthouse Lab in Milton Keynes et al |
| EPI_ISL_3701807 | Lighthouse Lab in Milton Keynes | Wellcome Sanger Institute for the COVID-19 Genomics UK (COG-UK) Consortium | The Lighthouse Lab in Milton Keynes et al |
| EPI_ISL_3966982 | Lighthouse Lab in Milton Keynes | Wellcome Sanger Institute for the COVID-19 Genomics UK (COG-UK) Consortium | The Lighthouse Lab in Milton Keynes et al |
| EPI_ISL_4065149 | Lighthouse Lab in Milton Keynes | Wellcome Sanger Institute for the COVID-19 Genomics UK (COG-UK) Consortium | The Lighthouse Lab in Milton Keynes et al |
| EPI_ISL_3454534 | Lighthouse Lab in Milton Keynes | Wellcome Sanger Institute for the COVID-19 Genomics UK (COG-UK) Consortium | The Lighthouse Lab in Milton Keynes et al |
| EPI_ISL_3952274 | Lighthouse Lab in Milton Keynes | Wellcome Sanger Institute for the COVID-19 Genomics UK (COG-UK) Consortium | The Lighthouse Lab in Milton Keynes et al |
| EPI_ISL_3363526 | Lighthouse Lab in Milton Keynes | Wellcome Sanger Institute for the COVID-19 Genomics UK (COG-UK) Consortium | The Lighthouse Lab in Milton Keynes et al |
| EPI_ISL_3809608 | Lighthouse Lab in Milton Keynes | Wellcome Sanger Institute for the COVID-19 Genomics UK (COG-UK) Consortium | The Lighthouse Lab in Milton Keynes et al |
| EPI_ISL_4500512 | Lighthouse Lab in Milton Keynes | Wellcome Sanger Institute for the COVID-19 Genomics UK (COG-UK) Consortium | The Lighthouse Lab in Milton Keynes et al |
| EPI_ISL_3883865 | Lighthouse Lab in Milton Keynes | Wellcome Sanger Institute for the COVID-19 Genomics UK (COG-UK) Consortium | The Lighthouse Lab in Milton Keynes et al |
| EPI_ISL_2911406 | Lighthouse Lab in Milton Keynes | Wellcome Sanger Institute for the COVID-19 Genomics UK (COG-UK) Consortium | The Lighthouse Lab in Milton Keynes et al |

| | | | |
|---|---|---|---|
| EPI_ISL_2468935 | Lighthouse Lab in Milton Keynes | Wellcome Sanger Institute for the COVID-19 Genomics UK (COG-UK) Consortium | The Lighthouse Lab in Milton Keynes et al |
| EPI_ISL_4735142 | Lighthouse Lab in Milton Keynes | Wellcome Sanger Institute for the COVID-19 Genomics UK (COG-UK) Consortium | The Lighthouse Lab in Milton Keynes et al |
| EPI_ISL_2640571 | Lighthouse Lab in Milton Keynes | Wellcome Sanger Institute for the COVID-19 Genomics UK (COG-UK) Consortium | The Lighthouse Lab in Milton Keynes et al |
| EPI_ISL_3313139 | Lighthouse Lab in Milton Keynes | Wellcome Sanger Institute for the COVID-19 Genomics UK (COG-UK) Consortium | The Lighthouse Lab in Milton Keynes et al |
| EPI_ISL_4521728 | Lighthouse Lab in Milton Keynes | Wellcome Sanger Institute for the COVID-19 Genomics UK (COG-UK) Consortium | The Lighthouse Lab in Milton Keynes et al |
| EPI_ISL_3994911 | Lighthouse Lab in Milton Keynes | Wellcome Sanger Institute for the COVID-19 Genomics UK (COG-UK) Consortium | The Lighthouse Lab in Milton Keynes et al |
| EPI_ISL_3498564 | Lighthouse Lab in Milton Keynes | Wellcome Sanger Institute for the COVID-19 Genomics UK (COG-UK) Consortium | The Lighthouse Lab in Milton Keynes et al |
| EPI_ISL_3767686 | Lighthouse Lab in Milton Keynes | Wellcome Sanger Institute for the COVID-19 Genomics UK (COG-UK) Consortium | The Lighthouse Lab in Milton Keynes et al |
| EPI_ISL_4500293 | Lighthouse Lab in Milton Keynes | Wellcome Sanger Institute for the COVID-19 Genomics UK (COG-UK) Consortium | The Lighthouse Lab in Milton Keynes et al |
| EPI_ISL_3170644 | Lighthouse Lab in Milton Keynes | Wellcome Sanger Institute for the COVID-19 Genomics UK (COG-UK) Consortium | The Lighthouse Lab in Milton Keynes et al |
| EPI_ISL_4350399 | Lighthouse Lab in Milton Keynes | Wellcome Sanger Institute for the COVID-19 Genomics UK (COG-UK) Consortium | The Lighthouse Lab in Milton Keynes et al |
| EPI_ISL_3094965 | Lighthouse Lab in Milton Keynes | Wellcome Sanger Institute for the COVID-19 Genomics UK (COG-UK) Consortium | The Lighthouse Lab in Milton Keynes et al |
| EPI_ISL_2908825 | Lighthouse Lab in Milton Keynes | Wellcome Sanger Institute for the COVID-19 Genomics UK (COG-UK) Consortium | The Lighthouse Lab in Milton Keynes et al |
| EPI_ISL_4349569 | Lighthouse Lab in Milton Keynes | Wellcome Sanger Institute for the COVID-19 Genomics UK (COG-UK) Consortium | The Lighthouse Lab in Milton Keynes et al |
| EPI_ISL_4321139 | Lighthouse Lab in Milton Keynes | Wellcome Sanger Institute for the COVID-19 Genomics UK (COG-UK) Consortium | The Lighthouse Lab in Milton Keynes et al |
| EPI_ISL_4754561 | Lighthouse Lab in Milton Keynes | Wellcome Sanger Institute for the COVID-19 Genomics UK (COG-UK) Consortium | The Lighthouse Lab in Milton Keynes et al |
| EPI_ISL_4213706 | Lighthouse Lab in Milton Keynes | Wellcome Sanger Institute for the COVID-19 Genomics UK (COG-UK) Consortium | The Lighthouse Lab in Milton Keynes et al |

| | | | |
|---|---|---|---|
| EPI_ISL_2570283 | Lighthouse Lab in Milton Keynes | Wellcome Sanger Institute for the COVID-19 Genomics UK (COG-UK) Consortium | The Lighthouse Lab in Milton Keynes et al |
| EPI_ISL_4320504 | Lighthouse Lab in Milton Keynes | Wellcome Sanger Institute for the COVID-19 Genomics UK (COG-UK) Consortium | The Lighthouse Lab in Milton Keynes et al |
| EPI_ISL_3286878 | Lighthouse Lab in Milton Keynes | Wellcome Sanger Institute for the COVID-19 Genomics UK (COG-UK) Consortium | The Lighthouse Lab in Milton Keynes et al |
| EPI_ISL_3312011 | Lighthouse Lab in Milton Keynes | Wellcome Sanger Institute for the COVID-19 Genomics UK (COG-UK) Consortium | The Lighthouse Lab in Milton Keynes et al |
| EPI_ISL_6499037 | Lighthouse Lab in Milton Keynes | Wellcome Sanger Institute for the COVID-19 Genomics UK (COG-UK) Consortium | The Lighthouse Lab in Milton Keynes et al |
| EPI_ISL_4765780 | Lighthouse Lab in Milton Keynes | Wellcome Sanger Institute for the COVID-19 Genomics UK (COG-UK) Consortium | The Lighthouse Lab in Milton Keynes et al |
| EPI_ISL_6366304 | Lighthouse Lab in Milton Keynes | Wellcome Sanger Institute for the COVID-19 Genomics UK (COG-UK) Consortium | The Lighthouse Lab in Milton Keynes et al |
| EPI_ISL_6365108 | Lighthouse Lab in Milton Keynes | Wellcome Sanger Institute for the COVID-19 Genomics UK (COG-UK) Consortium | The Lighthouse Lab in Milton Keynes et al |
| EPI_ISL_6360728 | Lighthouse Lab in Milton Keynes | Wellcome Sanger Institute for the COVID-19 Genomics UK (COG-UK) Consortium | The Lighthouse Lab in Milton Keynes et al |
| EPI_ISL_5355515 | Lighthouse Lab in Milton Keynes | Wellcome Sanger Institute for the COVID-19 Genomics UK (COG-UK) Consortium | The Lighthouse Lab in Milton Keynes et al |
| EPI_ISL_5613098 | Lighthouse Lab in Milton Keynes | Wellcome Sanger Institute for the COVID-19 Genomics UK (COG-UK) Consortium | The Lighthouse Lab in Milton Keynes et al |
| EPI_ISL_5026907 | Lighthouse Lab in Milton Keynes | Wellcome Sanger Institute for the COVID-19 Genomics UK (COG-UK) Consortium | The Lighthouse Lab in Milton Keynes et al |
| EPI_ISL_6501634 | Lighthouse Lab in Milton Keynes | Wellcome Sanger Institute for the COVID-19 Genomics UK (COG-UK) Consortium | The Lighthouse Lab in Milton Keynes et al |
| EPI_ISL_6223628 | Lighthouse Lab in Milton Keynes | Wellcome Sanger Institute for the COVID-19 Genomics UK (COG-UK) Consortium | The Lighthouse Lab in Milton Keynes et al |
| EPI_ISL_6045984 | Lighthouse Lab in Milton Keynes | Wellcome Sanger Institute for the COVID-19 Genomics UK (COG-UK) Consortium | The Lighthouse Lab in Milton Keynes et al |
| EPI_ISL_5142339 | Lighthouse Lab in Milton Keynes | Wellcome Sanger Institute for the COVID-19 Genomics UK (COG-UK) Consortium | The Lighthouse Lab in Milton Keynes et al |
| EPI_ISL_6823138 | Lighthouse Lab in Milton Keynes | Wellcome Sanger Institute for the COVID-19 Genomics UK (COG-UK) Consortium | The Lighthouse Lab in Milton Keynes et al |

| | | | |
|---|---|---|---|
| EPI_ISL_5918768 | Lighthouse Lab in Milton Keynes | Wellcome Sanger Institute for the COVID-19 Genomics UK (COG-UK) Consortium | The Lighthouse Lab in Milton Keynes et al |
| EPI_ISL_6818284 | Lighthouse Lab in Milton Keynes | Wellcome Sanger Institute for the COVID-19 Genomics UK (COG-UK) Consortium | The Lighthouse Lab in Milton Keynes et al |
| EPI_ISL_5793751 | Lighthouse Lab in Milton Keynes | Wellcome Sanger Institute for the COVID-19 Genomics UK (COG-UK) Consortium | The Lighthouse Lab in Milton Keynes et al |
| EPI_ISL_6839601 | Lighthouse Lab in Milton Keynes | Wellcome Sanger Institute for the COVID-19 Genomics UK (COG-UK) Consortium | The Lighthouse Lab in Milton Keynes et al |
| EPI_ISL_6580748 | Lighthouse Lab in Milton Keynes | Wellcome Sanger Institute for the COVID-19 Genomics UK (COG-UK) Consortium | The Lighthouse Lab in Milton Keynes et al |
| EPI_ISL_6301087 | Lighthouse Lab in Milton Keynes | Wellcome Sanger Institute for the COVID-19 Genomics UK (COG-UK) Consortium | The Lighthouse Lab in Milton Keynes et al |
| EPI_ISL_5789515 | Lighthouse Lab in Milton Keynes | Wellcome Sanger Institute for the COVID-19 Genomics UK (COG-UK) Consortium | The Lighthouse Lab in Milton Keynes et al |
| EPI_ISL_5785344 | Lighthouse Lab in Milton Keynes | Wellcome Sanger Institute for the COVID-19 Genomics UK (COG-UK) Consortium | The Lighthouse Lab in Milton Keynes et al |
| EPI_ISL_6770549 | Lighthouse Lab in Milton Keynes | Wellcome Sanger Institute for the COVID-19 Genomics UK (COG-UK) Consortium | The Lighthouse Lab in Milton Keynes et al |
| EPI_ISL_6797801 | Lighthouse Lab in Milton Keynes | Wellcome Sanger Institute for the COVID-19 Genomics UK (COG-UK) Consortium | The Lighthouse Lab in Milton Keynes et al |
| EPI_ISL_6300419 | Lighthouse Lab in Milton Keynes | Wellcome Sanger Institute for the COVID-19 Genomics UK (COG-UK) Consortium | The Lighthouse Lab in Milton Keynes et al |
| EPI_ISL_5142196 | Lighthouse Lab in Milton Keynes | Wellcome Sanger Institute for the COVID-19 Genomics UK (COG-UK) Consortium | The Lighthouse Lab in Milton Keynes et al |
| EPI_ISL_6498582 | Lighthouse Lab in Milton Keynes | Wellcome Sanger Institute for the COVID-19 Genomics UK (COG-UK) Consortium | The Lighthouse Lab in Milton Keynes et al |
| EPI_ISL_5612955 | Lighthouse Lab in Milton Keynes | Wellcome Sanger Institute for the COVID-19 Genomics UK (COG-UK) Consortium | The Lighthouse Lab in Milton Keynes et al |
| EPI_ISL_6038438 | Lighthouse Lab in Milton Keynes | Wellcome Sanger Institute for the COVID-19 Genomics UK (COG-UK) Consortium | The Lighthouse Lab in Milton Keynes et al |
| EPI_ISL_6300957 | Lighthouse Lab in Milton Keynes | Wellcome Sanger Institute for the COVID-19 Genomics UK (COG-UK) Consortium | The Lighthouse Lab in Milton Keynes et al |
| EPI_ISL_6215130 | Lighthouse Lab in Milton Keynes | Wellcome Sanger Institute for the COVID-19 Genomics UK (COG-UK) Consortium | The Lighthouse Lab in Milton Keynes et al |

| | | | |
|---|---|---|---|
| EPI_ISL_4896945 | Lighthouse Lab in Milton Keynes | Wellcome Sanger Institute for the COVID-19 Genomics UK (COG-UK) Consortium | The Lighthouse Lab in Milton Keynes et al |
| EPI_ISL_6576556 | Lighthouse Lab in Milton Keynes | Wellcome Sanger Institute for the COVID-19 Genomics UK (COG-UK) Consortium | The Lighthouse Lab in Milton Keynes et al |
| EPI_ISL_6497131 | Lighthouse Lab in Milton Keynes | Wellcome Sanger Institute for the COVID-19 Genomics UK (COG-UK) Consortium | The Lighthouse Lab in Milton Keynes et al |
| EPI_ISL_5057063 | Lighthouse Lab in Milton Keynes | Wellcome Sanger Institute for the COVID-19 Genomics UK (COG-UK) Consortium | The Lighthouse Lab in Milton Keynes et al |
| EPI_ISL_5994255 | Lighthouse Lab in Milton Keynes | Wellcome Sanger Institute for the COVID-19 Genomics UK (COG-UK) Consortium | The Lighthouse Lab in Milton Keynes et al |
| EPI_ISL_5787558 | Lighthouse Lab in Milton Keynes | Wellcome Sanger Institute for the COVID-19 Genomics UK (COG-UK) Consortium | The Lighthouse Lab in Milton Keynes et al |
| EPI_ISL_5035012 | Lighthouse Lab in Milton Keynes | Wellcome Sanger Institute for the COVID-19 Genomics UK (COG-UK) Consortium | The Lighthouse Lab in Milton Keynes et al |
| EPI_ISL_6150502 | Lighthouse Lab in Milton Keynes | Wellcome Sanger Institute for the COVID-19 Genomics UK (COG-UK) Consortium | The Lighthouse Lab in Milton Keynes et al |
| EPI_ISL_5135290 | Lighthouse Lab in Milton Keynes | Wellcome Sanger Institute for the COVID-19 Genomics UK (COG-UK) Consortium | The Lighthouse Lab in Milton Keynes et al |
| EPI_ISL_5829867 | Lighthouse Lab in Milton Keynes | Wellcome Sanger Institute for the COVID-19 Genomics UK (COG-UK) Consortium | The Lighthouse Lab in Milton Keynes et al |
| EPI_ISL_6823529 | Lighthouse Lab in Milton Keynes | Wellcome Sanger Institute for the COVID-19 Genomics UK (COG-UK) Consortium | The Lighthouse Lab in Milton Keynes et al |
| EPI_ISL_6672336 | Lighthouse Lab in Milton Keynes | Wellcome Sanger Institute for the COVID-19 Genomics UK (COG-UK) Consortium | The Lighthouse Lab in Milton Keynes et al |
| EPI_ISL_5037498 | Lighthouse Lab in Milton Keynes | Wellcome Sanger Institute for the COVID-19 Genomics UK (COG-UK) Consortium | The Lighthouse Lab in Milton Keynes et al |
| EPI_ISL_6557673 | Lighthouse Lab in Milton Keynes | Wellcome Sanger Institute for the COVID-19 Genomics UK (COG-UK) Consortium | The Lighthouse Lab in Milton Keynes et al |
| EPI_ISL_6213126 | Lighthouse Lab in Milton Keynes | Wellcome Sanger Institute for the COVID-19 Genomics UK (COG-UK) Consortium | The Lighthouse Lab in Milton Keynes et al |
| EPI_ISL_5834577 | Lighthouse Lab in Milton Keynes | Wellcome Sanger Institute for the COVID-19 Genomics UK (COG-UK) Consortium | The Lighthouse Lab in Milton Keynes et al |
| EPI_ISL_6428764 | Lighthouse Lab in Milton Keynes | Wellcome Sanger Institute for the COVID-19 Genomics UK (COG-UK) Consortium | The Lighthouse Lab in Milton Keynes et al |

| | | | |
|---|---|---|---|
| EPI_ISL_6296250 | Lighthouse Lab in Milton Keynes | Wellcome Sanger Institute for the COVID-19 Genomics UK (COG-UK) Consortium | The Lighthouse Lab in Milton Keynes et al |
| EPI_ISL_5791498 | Lighthouse Lab in Milton Keynes | Wellcome Sanger Institute for the COVID-19 Genomics UK (COG-UK) Consortium | The Lighthouse Lab in Milton Keynes et al |
| EPI_ISL_6296620 | Lighthouse Lab in Milton Keynes | Wellcome Sanger Institute for the COVID-19 Genomics UK (COG-UK) Consortium | The Lighthouse Lab in Milton Keynes et al |
| EPI_ISL_6359613 | Lighthouse Lab in Milton Keynes | Wellcome Sanger Institute for the COVID-19 Genomics UK (COG-UK) Consortium | The Lighthouse Lab in Milton Keynes et al |
| EPI_ISL_6799283 | Lighthouse Lab in Milton Keynes | Wellcome Sanger Institute for the COVID-19 Genomics UK (COG-UK) Consortium | The Lighthouse Lab in Milton Keynes et al |
| EPI_ISL_5027776 | Lighthouse Lab in Milton Keynes | Wellcome Sanger Institute for the COVID-19 Genomics UK (COG-UK) Consortium | The Lighthouse Lab in Milton Keynes et al |
| EPI_ISL_4765303 | Lighthouse Lab in Milton Keynes | Wellcome Sanger Institute for the COVID-19 Genomics UK (COG-UK) Consortium | The Lighthouse Lab in Milton Keynes et al |
| EPI_ISL_5406930 | Lighthouse Lab in Milton Keynes | Wellcome Sanger Institute for the COVID-19 Genomics UK (COG-UK) Consortium | The Lighthouse Lab in Milton Keynes et al |
| EPI_ISL_5026205 | Lighthouse Lab in Milton Keynes | Wellcome Sanger Institute for the COVID-19 Genomics UK (COG-UK) Consortium | The Lighthouse Lab in Milton Keynes et al |
| EPI_ISL_6218738 | Lighthouse Lab in Milton Keynes | Wellcome Sanger Institute for the COVID-19 Genomics UK (COG-UK) Consortium | The Lighthouse Lab in Milton Keynes et al |
| EPI_ISL_6354592 | Lighthouse Lab in Milton Keynes | Wellcome Sanger Institute for the COVID-19 Genomics UK (COG-UK) Consortium | The Lighthouse Lab in Milton Keynes et al |
| EPI_ISL_5785750 | Lighthouse Lab in Milton Keynes | Wellcome Sanger Institute for the COVID-19 Genomics UK (COG-UK) Consortium | The Lighthouse Lab in Milton Keynes et al |
| EPI_ISL_5967651 | Lighthouse Lab in Milton Keynes | Wellcome Sanger Institute for the COVID-19 Genomics UK (COG-UK) Consortium | The Lighthouse Lab in Milton Keynes et al |
| EPI_ISL_6835597 | Lighthouse Lab in Milton Keynes | Wellcome Sanger Institute for the COVID-19 Genomics UK (COG-UK) Consortium | The Lighthouse Lab in Milton Keynes et al |
| EPI_ISL_5789330 | Lighthouse Lab in Milton Keynes | Wellcome Sanger Institute for the COVID-19 Genomics UK (COG-UK) Consortium | The Lighthouse Lab in Milton Keynes et al |
| EPI_ISL_5449516 | Lighthouse Lab in Milton Keynes | Wellcome Sanger Institute for the COVID-19 Genomics UK (COG-UK) Consortium | The Lighthouse Lab in Milton Keynes et al |
| EPI_ISL_6501457 | Lighthouse Lab in Milton Keynes | Wellcome Sanger Institute for the COVID-19 Genomics UK (COG-UK) Consortium | The Lighthouse Lab in Milton Keynes et al |

| | | | |
|---|---|---|---|
| EPI_ISL_6796053 | Lighthouse Lab in Milton Keynes | Wellcome Sanger Institute for the COVID-19 Genomics UK (COG-UK) Consortium | The Lighthouse Lab in Milton Keynes et al |
| EPI_ISL_6833884 | Lighthouse Lab in Milton Keynes | Wellcome Sanger Institute for the COVID-19 Genomics UK (COG-UK) Consortium | The Lighthouse Lab in Milton Keynes et al |
| EPI_ISL_5349451 | Lighthouse Lab in Milton Keynes | Wellcome Sanger Institute for the COVID-19 Genomics UK (COG-UK) Consortium | The Lighthouse Lab in Milton Keynes et al |
| EPI_ISL_6225666 | Lighthouse Lab in Milton Keynes | Wellcome Sanger Institute for the COVID-19 Genomics UK (COG-UK) Consortium | The Lighthouse Lab in Milton Keynes et al |
| EPI_ISL_6872286 | Lighthouse Lab in Milton Keynes | Wellcome Sanger Institute for the COVID-19 Genomics UK (COG-UK) Consortium | The Lighthouse Lab in Milton Keynes et al |
| EPI_ISL_6009700 | Lighthouse Lab in Milton Keynes | Wellcome Sanger Institute for the COVID-19 Genomics UK (COG-UK) Consortium | The Lighthouse Lab in Milton Keynes et al |
| EPI_ISL_5974987 | Lighthouse Lab in Milton Keynes | Wellcome Sanger Institute for the COVID-19 Genomics UK (COG-UK) Consortium | The Lighthouse Lab in Milton Keynes et al |
| EPI_ISL_6823336 | Lighthouse Lab in Milton Keynes | Wellcome Sanger Institute for the COVID-19 Genomics UK (COG-UK) Consortium | The Lighthouse Lab in Milton Keynes et al |
| EPI_ISL_5919099 | Lighthouse Lab in Milton Keynes | Wellcome Sanger Institute for the COVID-19 Genomics UK (COG-UK) Consortium | The Lighthouse Lab in Milton Keynes et al |
| EPI_ISL_6429492 | Lighthouse Lab in Milton Keynes | Wellcome Sanger Institute for the COVID-19 Genomics UK (COG-UK) Consortium | The Lighthouse Lab in Milton Keynes et al |
| EPI_ISL_6817721 | Lighthouse Lab in Milton Keynes | Wellcome Sanger Institute for the COVID-19 Genomics UK (COG-UK) Consortium | The Lighthouse Lab in Milton Keynes et al |
| EPI_ISL_6876326 | Lighthouse Lab in Milton Keynes | Wellcome Sanger Institute for the COVID-19 Genomics UK (COG-UK) Consortium | The Lighthouse Lab in Milton Keynes et al |
| EPI_ISL_5893303 | Lighthouse Lab in Milton Keynes | Wellcome Sanger Institute for the COVID-19 Genomics UK (COG-UK) Consortium | The Lighthouse Lab in Milton Keynes et al |
| EPI_ISL_6501032 | Lighthouse Lab in Milton Keynes | Wellcome Sanger Institute for the COVID-19 Genomics UK (COG-UK) Consortium | The Lighthouse Lab in Milton Keynes et al |
| EPI_ISL_6356346 | Lighthouse Lab in Milton Keynes | Wellcome Sanger Institute for the COVID-19 Genomics UK (COG-UK) Consortium | The Lighthouse Lab in Milton Keynes et al |
| EPI_ISL_5922883 | Lighthouse Lab in Milton Keynes | Wellcome Sanger Institute for the COVID-19 Genomics UK (COG-UK) Consortium | The Lighthouse Lab in Milton Keynes et al |
| EPI_ISL_4764149 | Lighthouse Lab in Milton Keynes | Wellcome Sanger Institute for the COVID-19 Genomics UK (COG-UK) Consortium | The Lighthouse Lab in Milton Keynes et al |

| | | | |
|---|---|---|---|
| EPI_ISL_6501697 | Lighthouse Lab in Milton Keynes | Wellcome Sanger Institute for the COVID-19 Genomics UK (COG-UK) Consortium | The Lighthouse Lab in Milton Keynes et al |
| EPI_ISL_6562859 | Lighthouse Lab in Milton Keynes | Wellcome Sanger Institute for the COVID-19 Genomics UK (COG-UK) Consortium | The Lighthouse Lab in Milton Keynes et al |
| EPI_ISL_6834467 | Lighthouse Lab in Milton Keynes | Wellcome Sanger Institute for the COVID-19 Genomics UK (COG-UK) Consortium | The Lighthouse Lab in Milton Keynes et al |
| EPI_ISL_4528512 | Northumbria University / South Tees Hospitals NHS Foundation Trust / North Cumbria Integrated Care NHS Foundation Trust / North Tees and Hartlepool NHS Foundation Trust / Newcastle Hospitals NHS Foundation Trust | COVID-19 Genomics UK (COG-UK) Consortium | Darren L Smith et al |
| EPI_ISL_7023294 | Lighthouse Lab in Milton Keynes | Wellcome Sanger Institute for the COVID-19 Genomics UK (COG-UK) Consortium | The Lighthouse Lab in Milton Keynes et al |
| EPI_ISL_2434480 | Northumbria University / South Tees Hospitals NHS Foundation Trust / North Cumbria Integrated Care NHS Foundation Trust / North Tees and Hartlepool NHS Foundation Trust / Newcastle Hospitals NHS Foundation Trust | COVID-19 Genomics UK (COG-UK) Consortium | Darren L Smith et al |
| EPI_ISL_3992207 | Integrated Covid Hub North East | Wellcome Sanger Institute for the COVID-19 Genomics UK (COG-UK) Consortium | Integrated Covid Hub North East et al |
| EPI_ISL_3969888 | Integrated Covid Hub North East | Wellcome Sanger Institute for the COVID-19 Genomics UK (COG-UK) Consortium | Integrated Covid Hub North East et al |
| EPI_ISL_3421591 | Northumbria University / South Tees Hospitals NHS Foundation Trust / North Cumbria Integrated Care NHS Foundation Trust / North Tees and Hartlepool NHS Foundation Trust / Newcastle Hospitals NHS Foundation Trust | COVID-19 Genomics UK (COG-UK) Consortium | Darren L Smith et al |
| EPI_ISL_3978246 | Integrated Covid Hub North East | Wellcome Sanger Institute for the COVID-19 Genomics UK (COG-UK) Consortium | Integrated Covid Hub North East et al |
| EPI_ISL_623315 | Lighthouse Lab in Milton Keynes | Wellcome Sanger Institute for the COVID-19 Genomics UK (COG-UK) consortium | The Lighthouse Lab in Milton Keynes et al |

| | | | |
|---|---|---|---|
| EPI_ISL_676298 | Lighthouse Lab in Milton Keynes | Wellcome Sanger Institute for the COVID-19 Genomics UK (COG-UK) Consortium | The Lighthouse Lab in Milton Keynes et al |
| EPI_ISL_3978604 | Integrated Covid Hub North East | Wellcome Sanger Institute for the COVID-19 Genomics UK (COG-UK) Consortium | Integrated Covid Hub North East et al |
| EPI_ISL_875093 | Lighthouse Lab in Milton Keynes | Wellcome Sanger Institute for the COVID-19 Genomics UK (COG-UK) Consortium | The Lighthouse Lab in Milton Keynes et al |
| EPI_ISL_4587316 | Integrated Covid Hub North East | Wellcome Sanger Institute for the COVID-19 Genomics UK (COG-UK) Consortium | Integrated Covid Hub North East et al |
| EPI_ISL_7026727 | Lighthouse Lab in Milton Keynes | Wellcome Sanger Institute for the COVID-19 Genomics UK (COG-UK) Consortium | The Lighthouse Lab in Milton Keynes et al |
| EPI_ISL_2902335 | Northumbria University / South Tees Hospitals NHS Foundation Trust / North Cumbria Integrated Care NHS Foundation Trust / North Tees and Hartlepool NHS Foundation Trust / Newcastle Hospitals NHS Foundation Trust | COVID-19 Genomics UK (COG-UK) Consortium | Darren L Smith et al |
| EPI_ISL_7146498 | Lighthouse Lab in Milton Keynes | Wellcome Sanger Institute for the COVID-19 Genomics UK (COG-UK) Consortium | The Lighthouse Lab in Milton Keynes et al |
| EPI_ISL_7143602 | Lighthouse Lab in Milton Keynes | Wellcome Sanger Institute for the COVID-19 Genomics UK (COG-UK) Consortium | The Lighthouse Lab in Milton Keynes et al |
| EPI_ISL_5794072 | Integrated Covid Hub North East | Wellcome Sanger Institute for the COVID-19 Genomics UK (COG-UK) Consortium | Integrated Covid Hub North East et al |
| EPI_ISL_735548 | Lighthouse Lab in Milton Keynes | Wellcome Sanger Institute for the COVID-19 Genomics UK (COG-UK) Consortium | The Lighthouse Lab in Milton Keynes et al |
| EPI_ISL_5244813 | Integrated Covid Hub North East | Wellcome Sanger Institute for the COVID-19 Genomics UK (COG-UK) Consortium | Integrated Covid Hub North East et al |
| EPI_ISL_7023373 | Lighthouse Lab in Milton Keynes | Wellcome Sanger Institute for the COVID-19 Genomics UK (COG-UK) Consortium | The Lighthouse Lab in Milton Keynes et al |
| EPI_ISL_606619 | Lighthouse Lab in Milton Keynes | Wellcome Sanger Institute for the COVID-19 Genomics UK (COG-UK) consortium | The Lighthouse Lab in Milton Keynes et al |
| EPI_ISL_3969125 | Integrated Covid Hub North East | Wellcome Sanger Institute for the COVID-19 Genomics UK (COG-UK) Consortium | Integrated Covid Hub North East et al |
| EPI_ISL_4100629 | Integrated Covid Hub North East | Wellcome Sanger Institute for the COVID-19 Genomics UK (COG-UK) Consortium | Integrated Covid Hub North East et al |

| | | | |
|---|---|---|---|
| EPI_ISL_3975824 | Integrated Covid Hub North East | Wellcome Sanger Institute for the COVID-19 Genomics UK (COG-UK) Consortium | Integrated Covid Hub North East et al |
| EPI_ISL_768888 | Lighthouse Lab in Milton Keynes | Wellcome Sanger Institute for the COVID-19 Genomics UK (COG-UK) Consortium | The Lighthouse Lab in Milton Keynes et al |
| EPI_ISL_5996001 | Integrated Covid Hub North East | Wellcome Sanger Institute for the COVID-19 Genomics UK (COG-UK) Consortium | Integrated Covid Hub North East et al |
| EPI_ISL_6921921 | Lighthouse Lab in Milton Keynes | Wellcome Sanger Institute for the COVID-19 Genomics UK (COG-UK) Consortium | The Lighthouse Lab in Milton Keynes et al |
| EPI_ISL_4769946 | Northumbria University / South Tees Hospitals NHS Foundation Trust / North Cumbria Integrated Care NHS Foundation Trust / North Tees and Hartlepool NHS Foundation Trust / Newcastle Hospitals NHS Foundation Trust | COVID-19 Genomics UK (COG-UK) Consortium | Darren L Smith et al |
| EPI_ISL_5612493 | Integrated Covid Hub North East | Wellcome Sanger Institute for the COVID-19 Genomics UK (COG-UK) Consortium | Integrated Covid Hub North East et al |
| EPI_ISL_4354095 | Integrated Covid Hub North East | Wellcome Sanger Institute for the COVID-19 Genomics UK (COG-UK) Consortium | Integrated Covid Hub North East et al |
| EPI_ISL_3078480 | Northumbria University / South Tees Hospitals NHS Foundation Trust / North Cumbria Integrated Care NHS Foundation Trust / North Tees and Hartlepool NHS Foundation Trust / Newcastle Hospitals NHS Foundation Trust | COVID-19 Genomics UK (COG-UK) Consortium | Darren L Smith et al |
| EPI_ISL_679653 | Oxford Viromics, NDM, University of Oxford; Oxford University Hospitals; Basingstoke and North Hampshire Hospital | COVID-19 Genomics UK (COG-UK) Consortium | Tanya Golubchik et al |
| EPI_ISL_3421190 | Northumbria University / South Tees Hospitals NHS Foundation Trust / North Cumbria Integrated Care NHS Foundation Trust / North Tees and Hartlepool NHS Foundation Trust / Newcastle Hospitals NHS Foundation Trust | COVID-19 Genomics UK (COG-UK) Consortium | Darren L Smith et al |

| | | | |
|---|---|---|---|
| EPI_ISL_4015816 | Integrated Covid Hub North East | Wellcome Sanger Institute for the COVID-19 Genomics UK (COG-UK) Consortium | Integrated Covid Hub North East et al |
| EPI_ISL_2901870 | Northumbria University / South Tees Hospitals NHS Foundation Trust / North Cumbria Integrated Care NHS Foundation Trust / North Tees and Hartlepool NHS Foundation Trust / Newcastle Hospitals NHS Foundation Trust | COVID-19 Genomics UK (COG-UK) Consortium | Darren L Smith et al |
| EPI_ISL_681365 | Lighthouse Lab in Milton Keynes | Wellcome Sanger Institute for the COVID-19 Genomics UK (COG-UK) Consortium | The Lighthouse Lab in Milton Keynes et al |
| EPI_ISL_736424 | Lighthouse Lab in Milton Keynes | Wellcome Sanger Institute for the COVID-19 Genomics UK (COG-UK) Consortium | The Lighthouse Lab in Milton Keynes et al |
| EPI_ISL_2902122 | Northumbria University / South Tees Hospitals NHS Foundation Trust / North Cumbria Integrated Care NHS Foundation Trust / North Tees and Hartlepool NHS Foundation Trust / Newcastle Hospitals NHS Foundation Trust | COVID-19 Genomics UK (COG-UK) Consortium | Darren L Smith et al |
| EPI_ISL_2240255 | Northumbria University / South Tees Hospitals NHS Foundation Trust / North Cumbria Integrated Care NHS Foundation Trust / North Tees and Hartlepool NHS Foundation Trust / Newcastle Hospitals NHS Foundation Trust | COVID-19 Genomics UK (COG-UK) Consortium | Darren L Smith et al |
| EPI_ISL_601349 | Lighthouse Lab in Milton Keynes | Wellcome Sanger Institute for the COVID-19 Genomics UK (COG-UK) consortium | The Lighthouse Lab in Milton Keynes et al |
| EPI_ISL_6923186 | Lighthouse Lab in Milton Keynes | Wellcome Sanger Institute for the COVID-19 Genomics UK (COG-UK) Consortium | The Lighthouse Lab in Milton Keynes et al |
| EPI_ISL_5184618 | Integrated Covid Hub North East | Wellcome Sanger Institute for the COVID-19 Genomics UK (COG-UK) Consortium | Integrated Covid Hub North East et al |
| EPI_ISL_6919153 | Lighthouse Lab in Milton Keynes | Wellcome Sanger Institute for the COVID-19 Genomics UK (COG-UK) Consortium | The Lighthouse Lab in Milton Keynes et al |

| | | | |
|---|---|---|---|
| EPI_ISL_3177516 | Northumbria University / South Tees Hospitals NHS Foundation Trust / North Cumbria Integrated Care NHS Foundation Trust / North Tees and Hartlepool NHS Foundation Trust / Newcastle Hospitals NHS Foundation Trust | COVID-19 Genomics UK (COG-UK) Consortium | Darren L Smith et al |
| EPI_ISL_3975798 | Integrated Covid Hub North East | Wellcome Sanger Institute for the COVID-19 Genomics UK (COG-UK) Consortium | Integrated Covid Hub North East et al |
| EPI_ISL_552018 | Lighthouse Lab in Milton Keynes | Wellcome Sanger Institute for the COVID-19 Genomics UK (COG-UK) consortium | The Lighthouse Lab in Milton Keynes et al |
| EPI_ISL_5270819 | Northumbria University / South Tees Hospitals NHS Foundation Trust / North Cumbria Integrated Care NHS Foundation Trust / North Tees and Hartlepool NHS Foundation Trust / Newcastle Hospitals NHS Foundation Trust | COVID-19 Genomics UK (COG-UK) Consortium | Darren L Smith et al |
| EPI_ISL_4619983 | Integrated Covid Hub North East | Wellcome Sanger Institute for the COVID-19 Genomics UK (COG-UK) Consortium | Integrated Covid Hub North East et al |
| EPI_ISL_3969087 | Integrated Covid Hub North East | Wellcome Sanger Institute for the COVID-19 Genomics UK (COG-UK) Consortium | Integrated Covid Hub North East et al |
| EPI_ISL_2518616 | Northumbria University / South Tees Hospitals NHS Foundation Trust / North Cumbria Integrated Care NHS Foundation Trust / North Tees and Hartlepool NHS Foundation Trust / Newcastle Hospitals NHS Foundation Trust | COVID-19 Genomics UK (COG-UK) Consortium | Darren L Smith et al |
| EPI_ISL_3977357 | Integrated Covid Hub North East | Wellcome Sanger Institute for the COVID-19 Genomics UK (COG-UK) Consortium | Integrated Covid Hub North East et al |
| EPI_ISL_557284 | Lighthouse Lab in Milton Keynes | Wellcome Sanger Institute for the COVID-19 Genomics UK (COG-UK) consortium | The Lighthouse Lab in Milton Keynes et al |
| EPI_ISL_5973312 | Integrated Covid Hub North East | Wellcome Sanger Institute for the COVID-19 Genomics UK (COG-UK) Consortium | Integrated Covid Hub North East et al |
| EPI_ISL_598764 | Lighthouse Lab in Milton Keynes | Wellcome Sanger Institute for the COVID-19 Genomics UK (COG-UK) consortium | The Lighthouse Lab in Milton Keynes et al |

| | | | |
|---|---|---|---|
| EPI_ISL_1475191 | Northumbria University / South Tees Hospitals NHS Foundation Trust / North Cumbria Integrated Care NHS Foundation Trust / North Tees and Hartlepool NHS Foundation Trust / Newcastle Hospitals NHS Foundation Trust | COVID-19 Genomics UK (COG-UK) Consortium | Darren L Smith et al |
| EPI_ISL_573385 | Northumbria University / South Tees Hospitals NHS Foundation Trust / North Cumbria Integrated Care NHS Foundation Trust / North Tees and Hartlepool NHS Foundation Trust / Newcastle Hospitals NHS Foundation Trust | COVID-19 Genomics UK (COG-UK) Consortium | Darren L Smith et al |
| EPI_ISL_4589209 | Integrated Covid Hub North East | Wellcome Sanger Institute for the COVID-19 Genomics UK (COG-UK) Consortium | Integrated Covid Hub North East et al |
| EPI_ISL_736617 | Lighthouse Lab in Milton Keynes | Wellcome Sanger Institute for the COVID-19 Genomics UK (COG-UK) Consortium | The Lighthouse Lab in Milton Keynes et al |
| EPI_ISL_2240075 | Northumbria University / South Tees Hospitals NHS Foundation Trust / North Cumbria Integrated Care NHS Foundation Trust / North Tees and Hartlepool NHS Foundation Trust / Newcastle Hospitals NHS Foundation Trust | COVID-19 Genomics UK (COG-UK) Consortium | Darren L Smith et al |
| EPI_ISL_5028346 | Integrated Covid Hub North East | Wellcome Sanger Institute for the COVID-19 Genomics UK (COG-UK) Consortium | Integrated Covid Hub North East et al |
| EPI_ISL_7026952 | Lighthouse Lab in Milton Keynes | Wellcome Sanger Institute for the COVID-19 Genomics UK (COG-UK) Consortium | The Lighthouse Lab in Milton Keynes et al |
| EPI_ISL_5267792 | Northumbria University / South Tees Hospitals NHS Foundation Trust / North Cumbria Integrated Care NHS Foundation Trust / North Tees and Hartlepool NHS Foundation Trust / Newcastle Hospitals NHS Foundation Trust | COVID-19 Genomics UK (COG-UK) Consortium | Darren L Smith et al |
| EPI_ISL_3970624 | Integrated Covid Hub North East | Wellcome Sanger Institute for the COVID-19 Genomics UK (COG-UK) Consortium | Integrated Covid Hub North East et al |

| | | | |
|---|---|---|---|
| EPI_ISL_4860126 | Integrated Covid Hub North East | Wellcome Sanger Institute for the COVID-19 Genomics UK (COG-UK) Consortium | Integrated Covid Hub North East et al |
| EPI_ISL_4596476 | Integrated Covid Hub North East | Wellcome Sanger Institute for the COVID-19 Genomics UK (COG-UK) Consortium | Integrated Covid Hub North East et al |
| EPI_ISL_4522744 | Integrated Covid Hub North East | Wellcome Sanger Institute for the COVID-19 Genomics UK (COG-UK) Consortium | Integrated Covid Hub North East et al |
| EPI_ISL_5269831 | Northumbria University / South Tees Hospitals NHS Foundation Trust / North Cumbria Integrated Care NHS Foundation Trust / North Tees and Hartlepool NHS Foundation Trust / Newcastle Hospitals NHS Foundation Trust | COVID-19 Genomics UK (COG-UK) Consortium | Darren L Smith et al |
| EPI_ISL_7026616 | Lighthouse Lab in Milton Keynes | Wellcome Sanger Institute for the COVID-19 Genomics UK (COG-UK) Consortium | The Lighthouse Lab in Milton Keynes et al |
| EPI_ISL_4403821 | Integrated Covid Hub North East | Wellcome Sanger Institute for the COVID-19 Genomics UK (COG-UK) Consortium | Integrated Covid Hub North East et al |
| EPI_ISL_5468131 | Northumbria University / South Tees Hospitals NHS Foundation Trust / North Cumbria Integrated Care NHS Foundation Trust / North Tees and Hartlepool NHS Foundation Trust / Newcastle Hospitals NHS Foundation Trust | COVID-19 Genomics UK (COG-UK) Consortium | Darren L Smith et al |
| EPI_ISL_575836 | Lighthouse Lab in Milton Keynes | Wellcome Sanger Institute for the COVID-19 Genomics UK (COG-UK) consortium | The Lighthouse Lab in Milton Keynes et al |
| EPI_ISL_7020314 | Lighthouse Lab in Milton Keynes | Wellcome Sanger Institute for the COVID-19 Genomics UK (COG-UK) Consortium | The Lighthouse Lab in Milton Keynes et al |
| EPI_ISL_3969822 | Integrated Covid Hub North East | Wellcome Sanger Institute for the COVID-19 Genomics UK (COG-UK) Consortium | Integrated Covid Hub North East et al |
| EPI_ISL_2723205 | Northumbria University / South Tees Hospitals NHS Foundation Trust / North Cumbria Integrated Care NHS Foundation Trust / North Tees and Hartlepool NHS Foundation Trust / Newcastle Hospitals NHS Foundation Trust | COVID-19 Genomics UK (COG-UK) Consortium | Darren L Smith et al |

| | | | |
|---|---|---|---|
| EPI_ISL_4522545 | Integrated Covid Hub North East | Wellcome Sanger Institute for the COVID-19 Genomics UK (COG-UK) Consortium | Integrated Covid Hub North East et al |
| EPI_ISL_3971526 | Integrated Covid Hub North East | Wellcome Sanger Institute for the COVID-19 Genomics UK (COG-UK) Consortium | Integrated Covid Hub North East et al |
| EPI_ISL_866514 | Northumbria University / South Tees Hospitals NHS Foundation Trust / North Cumbria Integrated Care NHS Foundation Trust / North Tees and Hartlepool NHS Foundation Trust / Newcastle Hospitals NHS Foundation Trust | COVID-19 Genomics UK (COG-UK) Consortium | Darren L Smith et al |
| EPI_ISL_5973075 | Integrated Covid Hub North East | Wellcome Sanger Institute for the COVID-19 Genomics UK (COG-UK) Consortium | Integrated Covid Hub North East et al |
| EPI_ISL_799874 | Lighthouse Lab in Milton Keynes | Wellcome Sanger Institute for the COVID-19 Genomics UK (COG-UK) Consortium | The Lighthouse Lab in Milton Keynes et al |
| EPI_ISL_3976011 | Integrated Covid Hub North East | Wellcome Sanger Institute for the COVID-19 Genomics UK (COG-UK) Consortium | Integrated Covid Hub North East et al |
| EPI_ISL_3970366 | Integrated Covid Hub North East | Wellcome Sanger Institute for the COVID-19 Genomics UK (COG-UK) Consortium | Integrated Covid Hub North East et al |
| EPI_ISL_733652 | Lighthouse Lab in Milton Keynes | Wellcome Sanger Institute for the COVID-19 Genomics UK (COG-UK) Consortium | The Lighthouse Lab in Milton Keynes et al |
| EPI_ISL_5994081 | Integrated Covid Hub North East | Wellcome Sanger Institute for the COVID-19 Genomics UK (COG-UK) Consortium | Integrated Covid Hub North East et al |
| EPI_ISL_638562 | Northumbria University / South Tees Hospitals NHS Foundation Trust / North Cumbria Integrated Care NHS Foundation Trust / North Tees and Hartlepool NHS Foundation Trust / Newcastle Hospitals NHS Foundation Trust | COVID-19 Genomics UK (COG-UK) Consortium | Darren L Smith et al |
| EPI_ISL_4100563 | Integrated Covid Hub North East | Wellcome Sanger Institute for the COVID-19 Genomics UK (COG-UK) Consortium | Integrated Covid Hub North East et al |
| EPI_ISL_674852 | Lighthouse Lab in Milton Keynes | Wellcome Sanger Institute for the COVID-19 Genomics UK (COG-UK) Consortium | The Lighthouse Lab in Milton Keynes et al |

| | | | |
|---|---|---|---|
| EPI_ISL_4306470 | Northumbria University / South Tees Hospitals NHS Foundation Trust / North Cumbria Integrated Care NHS Foundation Trust / North Tees and Hartlepool NHS Foundation Trust / Newcastle Hospitals NHS Foundation Trust | COVID-19 Genomics UK (COG-UK) Consortium | Darren L Smith et al |
| EPI_ISL_7029477 | Lighthouse Lab in Milton Keynes | Wellcome Sanger Institute for the COVID-19 Genomics UK (COG-UK) Consortium | The Lighthouse Lab in Milton Keynes et al |
| EPI_ISL_4851441 | Integrated Covid Hub North East | Wellcome Sanger Institute for the COVID-19 Genomics UK (COG-UK) Consortium | Integrated Covid Hub North East et al |
| EPI_ISL_6918785 | Lighthouse Lab in Milton Keynes | Wellcome Sanger Institute for the COVID-19 Genomics UK (COG-UK) Consortium | The Lighthouse Lab in Milton Keynes et al |
| EPI_ISL_2747287 | Respiratory Virus Unit, Microbiology Services Colindale, Public Health England | COVID-19 Genomics UK (COG-UK) Consortium | PHE Covid Sequencing Team et al |
| EPI_ISL_5809773 | Respiratory Virus Unit, Microbiology Services Colindale, Public Health England | COVID-19 Genomics UK (COG-UK) Consortium | PHE Covid Sequencing Team et al |
| EPI_ISL_3779148 | Respiratory Virus Unit, Microbiology Services Colindale, Public Health England | COVID-19 Genomics UK (COG-UK) Consortium | PHE Covid Sequencing Team et al |
| EPI_ISL_866799 | Quadram Institute Bioscience | COVID-19 Genomics UK (COG-UK) Consortium | Dave J. Baker et al |
| EPI_ISL_7165281 | Quadram Institute Bioscience | COVID-19 Genomics UK (COG-UK) Consortium | Dave J. Baker et al |
| EPI_ISL_4308341 | Respiratory Virus Unit, Microbiology Services Colindale, Public Health England | COVID-19 Genomics UK (COG-UK) Consortium | PHE Covid Sequencing Team et al |
| EPI_ISL_5471365 | Quadram Institute Bioscience | COVID-19 Genomics UK (COG-UK) Consortium | Dave J. Baker et al |
| EPI_ISL_1249084 | Oxford Viromics, NDM, University of Oxford; Oxford University Hospitals; Basingstoke and North Hampshire Hospital | COVID-19 Genomics UK (COG-UK) Consortium | Tanya Golubchik et al |
| EPI_ISL_866806 | Quadram Institute Bioscience | COVID-19 Genomics UK (COG-UK) Consortium | Dave J. Baker et al |
| EPI_ISL_999171 | Oxford Viromics, NDM, University of Oxford; Oxford University Hospitals; Basingstoke and North Hampshire Hospital | COVID-19 Genomics UK (COG-UK) Consortium | Tanya Golubchik et al |

| | | | |
|---|---|---|---|
| EPI_ISL_3292040 | Respiratory Virus Unit, Microbiology Services Colindale, Public Health England | COVID-19 Genomics UK (COG-UK) Consortium | PHE Covid Sequencing Team et al |
| EPI_ISL_1475570 | Oxford Viromics, NDM, University of Oxford; Oxford University Hospitals; Basingstoke and North Hampshire Hospital | COVID-19 Genomics UK (COG-UK) Consortium | Tanya Golubchik et al |
| EPI_ISL_4120054 | Respiratory Virus Unit, Microbiology Services Colindale, Public Health England | COVID-19 Genomics UK (COG-UK) Consortium | PHE Covid Sequencing Team et al |
| EPI_ISL_5275946 | Respiratory Virus Unit, Microbiology Services Colindale, Public Health England | COVID-19 Genomics UK (COG-UK) Consortium | PHE Covid Sequencing Team et al |
| EPI_ISL_3780674 | Respiratory Virus Unit, Microbiology Services Colindale, Public Health England | COVID-19 Genomics UK (COG-UK) Consortium | PHE Covid Sequencing Team et al |
| EPI_ISL_4120659 | Respiratory Virus Unit, Microbiology Services Colindale, Public Health England | COVID-19 Genomics UK (COG-UK) Consortium | PHE Covid Sequencing Team et al |
| EPI_ISL_5040636 | Quadram Institute Bioscience | COVID-19 Genomics UK (COG-UK) Consortium | Dave J. Baker et al |
| EPI_ISL_6023466 | Respiratory Virus Unit, Microbiology Services Colindale, Public Health England | COVID-19 Genomics UK (COG-UK) Consortium | PHE Covid Sequencing Team et al |
| EPI_ISL_4118488 | Respiratory Virus Unit, Microbiology Services Colindale, Public Health England | COVID-19 Genomics UK (COG-UK) Consortium | PHE Covid Sequencing Team et al |
| EPI_ISL_4306933 | Quadram Institute Bioscience | COVID-19 Genomics UK (COG-UK) Consortium | Dave J. Baker et al |
| EPI_ISL_6022813 | Respiratory Virus Unit, Microbiology Services Colindale, Public Health England | COVID-19 Genomics UK (COG-UK) Consortium | PHE Covid Sequencing Team et al |
| EPI_ISL_3778810 | Respiratory Virus Unit, Microbiology Services Colindale, Public Health England | COVID-19 Genomics UK (COG-UK) Consortium | PHE Covid Sequencing Team et al |
| EPI_ISL_3292079 | Respiratory Virus Unit, Microbiology Services Colindale, Public Health England | COVID-19 Genomics UK (COG-UK) Consortium | PHE Covid Sequencing Team et al |
| EPI_ISL_3572040 | Quadram Institute Bioscience | COVID-19 Genomics UK (COG-UK) Consortium | Dave J. Baker et al |

| | | | |
|---|---|---|---|
| EPI_ISL_2741522 | Respiratory Virus Unit, Microbiology Services Colindale, Public Health England | COVID-19 Genomics UK (COG-UK) Consortium | PHE Covid Sequencing Team et al |
| EPI_ISL_2519077 | Quadram Institute Bioscience | COVID-19 Genomics UK (COG-UK) Consortium | Dave J. Baker et al |
| EPI_ISL_6136425 | Respiratory Virus Unit, Microbiology Services Colindale, Public Health England | COVID-19 Genomics UK (COG-UK) Consortium | PHE Covid Sequencing Team et al |
| EPI_ISL_6978340 | Respiratory Virus Unit, Microbiology Services Colindale, Public Health England | COVID-19 Genomics UK (COG-UK) Consortium | PHE Covid Sequencing Team et al |
| EPI_ISL_4529251 | Quadram Institute Bioscience | COVID-19 Genomics UK (COG-UK) Consortium | Dave J. Baker et al |
| EPI_ISL_4124536 | Respiratory Virus Unit, Microbiology Services Colindale, Public Health England | COVID-19 Genomics UK (COG-UK) Consortium | PHE Covid Sequencing Team et al |
| EPI_ISL_2740388 | Respiratory Virus Unit, Microbiology Services Colindale, Public Health England | COVID-19 Genomics UK (COG-UK) Consortium | PHE Covid Sequencing Team et al |
| EPI_ISL_2746594 | Respiratory Virus Unit, Microbiology Services Colindale, Public Health England | COVID-19 Genomics UK (COG-UK) Consortium | PHE Covid Sequencing Team et al |
| EPI_ISL_1105850 | Lincolnshire Hospitals and DeepSeq Nottingham | COVID-19 Genomics UK (COG-UK) Consortium | Nichola Duckworth et al |
| EPI_ISL_7039636 | Quadram Institute Bioscience | COVID-19 Genomics UK (COG-UK) Consortium | Dave J. Baker et al |
| EPI_ISL_6976613 | Quadram Institute Bioscience | COVID-19 Genomics UK (COG-UK) Consortium | Dave J. Baker et al |
| EPI_ISL_4122327 | Respiratory Virus Unit, Microbiology Services Colindale, Public Health England | COVID-19 Genomics UK (COG-UK) Consortium | PHE Covid Sequencing Team et al |
| EPI_ISL_5271292 | Quadram Institute Bioscience | COVID-19 Genomics UK (COG-UK) Consortium | Dave J. Baker et al |
| EPI_ISL_4773355 | Respiratory Virus Unit, Microbiology Services Colindale, Public Health England | COVID-19 Genomics UK (COG-UK) Consortium | PHE Covid Sequencing Team et al |
| EPI_ISL_2519522 | Quadram Institute Bioscience | COVID-19 Genomics UK (COG-UK) Consortium | Dave J. Baker et al |
| EPI_ISL_6026354 | Respiratory Virus Unit, Microbiology Services Colindale, Public Health England | COVID-19 Genomics UK (COG-UK) Consortium | PHE Covid Sequencing Team et al |

| | | | |
|---|---|---|---|
| EPI_ISL_4123079 | Respiratory Virus Unit, Microbiology Services Colindale, Public Health England | COVID-19 Genomics UK (COG-UK) Consortium | PHE Covid Sequencing Team et al |
| EPI_ISL_4773034 | Respiratory Virus Unit, Microbiology Services Colindale, Public Health England | COVID-19 Genomics UK (COG-UK) Consortium | PHE Covid Sequencing Team et al |
| EPI_ISL_4528769 | Northumbria University / South Tees Hospitals NHS Foundation Trust / North Cumbria Integrated Care NHS Foundation Trust / North Tees and Hartlepool NHS Foundation Trust / Newcastle Hospitals NHS Foundation Trust | COVID-19 Genomics UK (COG-UK) Consortium | Darren L Smith et al |
| EPI_ISL_4308968 | Respiratory Virus Unit, Microbiology Services Colindale, Public Health England | COVID-19 Genomics UK (COG-UK) Consortium | PHE Covid Sequencing Team et al |
| EPI_ISL_3003481 | Oxford Viromics, NDM, University of Oxford; Oxford University Hospitals; Basingstoke and North Hampshire Hospital | COVID-19 Genomics UK (COG-UK) Consortium | Tanya Golubchik et al |
| EPI_ISL_4530482 | Respiratory Virus Unit, Microbiology Services Colindale, Public Health England | COVID-19 Genomics UK (COG-UK) Consortium | PHE Covid Sequencing Team et al |
| EPI_ISL_5278917 | Respiratory Virus Unit, Microbiology Services Colindale, Public Health England | COVID-19 Genomics UK (COG-UK) Consortium | PHE Covid Sequencing Team et al |
| EPI_ISL_4774590 | Respiratory Virus Unit, Microbiology Services Colindale, Public Health England | COVID-19 Genomics UK (COG-UK) Consortium | PHE Covid Sequencing Team et al |
| EPI_ISL_6136455 | Respiratory Virus Unit, Microbiology Services Colindale, Public Health England | COVID-19 Genomics UK (COG-UK) Consortium | PHE Covid Sequencing Team et al |
| EPI_ISL_3179752 | Respiratory Virus Unit, Microbiology Services Colindale, Public Health England | COVID-19 Genomics UK (COG-UK) Consortium | PHE Covid Sequencing Team et al |
| EPI_ISL_3289584 | Quadram Institute Bioscience | COVID-19 Genomics UK (COG-UK) Consortium | Dave J. Baker et al |
| EPI_ISL_4772670 | Respiratory Virus Unit, Microbiology Services Colindale, Public Health England | COVID-19 Genomics UK (COG-UK) Consortium | PHE Covid Sequencing Team et al |

| | | | |
|---|---|---|---|
| EPI_ISL_6977830 | Quadram Institute Bioscience | COVID-19 Genomics UK (COG-UK) Consortium | Dave J. Baker et al |
| EPI_ISL_6028530 | Respiratory Virus Unit, Microbiology Services Colindale, Public Health England | COVID-19 Genomics UK (COG-UK) Consortium | PHE Covid Sequencing Team et al |
| EPI_ISL_3002527 | Quadram Institute Bioscience | COVID-19 Genomics UK (COG-UK) Consortium | Dave J. Baker et al |
| EPI_ISL_2817790 | Respiratory Virus Unit, Microbiology Services Colindale, Public Health England | COVID-19 Genomics UK (COG-UK) Consortium | PHE Covid Sequencing Team et al |
| EPI_ISL_584752 | Queens Medical Centre, Clinical Microbiology Department / DeepSeq Nottingham | COVID-19 Genomics UK (COG-UK) Consortium | Gemma Clark et al |
| EPI_ISL_2746704 | Respiratory Virus Unit, Microbiology Services Colindale, Public Health England | COVID-19 Genomics UK (COG-UK) Consortium | PHE Covid Sequencing Team et al |
| EPI_ISL_2747043 | Respiratory Virus Unit, Microbiology Services Colindale, Public Health England | COVID-19 Genomics UK (COG-UK) Consortium | PHE Covid Sequencing Team et al |
| EPI_ISL_5273034 | Quadram Institute Bioscience | COVID-19 Genomics UK (COG-UK) Consortium | Dave J. Baker et al |
| EPI_ISL_4121796 | Respiratory Virus Unit, Microbiology Services Colindale, Public Health England | COVID-19 Genomics UK (COG-UK) Consortium | PHE Covid Sequencing Team et al |
| EPI_ISL_741650 | Queens Medical Centre, Clinical Microbiology Department / DeepSeq Nottingham | COVID-19 Genomics UK (COG-UK) Consortium | Gemma Clark et al |
| EPI_ISL_4116555 | Quadram Institute Bioscience | COVID-19 Genomics UK (COG-UK) Consortium | Dave J. Baker et al |
| EPI_ISL_3311749 | Lighthouse Laboratory Plymouth | Wellcome Sanger Institute for the COVID-19 Genomics UK (COG-UK) Consortium | Lighthouse Laboratory Plymouth et al |
| EPI_ISL_2850386 | Lighthouse Laboratory Plymouth | Wellcome Sanger Institute for the COVID-19 Genomics UK (COG-UK) Consortium | Lighthouse Laboratory Plymouth et al |
| EPI_ISL_3363291 | Lighthouse Laboratory Plymouth | Wellcome Sanger Institute for the COVID-19 Genomics UK (COG-UK) Consortium | Lighthouse Laboratory Plymouth et al |
| EPI_ISL_3701937 | Lighthouse Laboratory Plymouth | Wellcome Sanger Institute for the COVID-19 Genomics UK (COG-UK) Consortium | Lighthouse Laboratory Plymouth et al |
| EPI_ISL_3109781 | Lighthouse Laboratory Plymouth | Wellcome Sanger Institute for the COVID-19 Genomics UK (COG-UK) Consortium | Lighthouse Laboratory Plymouth et al |

| | | | |
|---|---|---|---|
| EPI_ISL_5449195 | Lighthouse Laboratory Plymouth | Wellcome Sanger Institute for the COVID-19 Genomics UK (COG-UK) Consortium | Lighthouse Laboratory Plymouth et al |
| EPI_ISL_3367320 | Lighthouse Laboratory Plymouth | Wellcome Sanger Institute for the COVID-19 Genomics UK (COG-UK) Consortium | Lighthouse Laboratory Plymouth et al |
| EPI_ISL_5976324 | Lighthouse Laboratory Plymouth | Wellcome Sanger Institute for the COVID-19 Genomics UK (COG-UK) Consortium | Lighthouse Laboratory Plymouth et al |
| EPI_ISL_4783087 | Respiratory Virus Unit, Microbiology Services Colindale, Public Health England | COVID-19 Genomics UK (COG-UK) Consortium | PHE Covid Sequencing Team et al |
| EPI_ISL_5042627 | Respiratory Virus Unit, Microbiology Services Colindale, Public Health England | COVID-19 Genomics UK (COG-UK) Consortium | PHE Covid Sequencing Team et al |
| EPI_ISL_6108152 | Lighthouse Laboratory Plymouth | Wellcome Sanger Institute for the COVID-19 Genomics UK (COG-UK) Consortium | Lighthouse Laboratory Plymouth et al |
| EPI_ISL_6280804 | Respiratory Virus Unit, Microbiology Services Colindale, Public Health England | COVID-19 Genomics UK (COG-UK) Consortium | PHE Covid Sequencing Team et al |
| EPI_ISL_2906081 | Originating lab: Wales Specialist Virology Centre Sequencing lab: Pathogen Genomics Unit | Public Health Wales Microbiology Cardiff Wales Specialist Virology Centre | Catherine Moore et al |
| EPI_ISL_2971835 | Lighthouse Laboratory Plymouth | Wellcome Sanger Institute for the COVID-19 Genomics UK (COG-UK) Consortium | Lighthouse Laboratory Plymouth et al |
| EPI_ISL_5043110 | Respiratory Virus Unit, Microbiology Services Colindale, Public Health England | COVID-19 Genomics UK (COG-UK) Consortium | PHE Covid Sequencing Team et al |
| EPI_ISL_6150220 | Lighthouse Laboratory Plymouth | Wellcome Sanger Institute for the COVID-19 Genomics UK (COG-UK) Consortium | Lighthouse Laboratory Plymouth et al |
| EPI_ISL_3441363 | Lighthouse Laboratory Plymouth | Wellcome Sanger Institute for the COVID-19 Genomics UK (COG-UK) Consortium | Lighthouse Laboratory Plymouth et al |
| EPI_ISL_3961986 | Respiratory Virus Unit, Microbiology Services Colindale, Public Health England | COVID-19 Genomics UK (COG-UK) Consortium | PHE Covid Sequencing Team et al |
| EPI_ISL_4532067 | Respiratory Virus Unit, Microbiology Services Colindale, Public Health England | COVID-19 Genomics UK (COG-UK) Consortium | PHE Covid Sequencing Team et al |
| EPI_ISL_3250186 | Lighthouse Laboratory Plymouth | Wellcome Sanger Institute for the COVID-19 Genomics UK (COG-UK) Consortium | Lighthouse Laboratory Plymouth et al |

| | | | |
|---|---|---|---|
| EPI_ISL_6806709 | Lighthouse Laboratory Plymouth | Wellcome Sanger Institute for the COVID-19 Genomics UK (COG-UK) Consortium | Lighthouse Laboratory Plymouth et al |
| EPI_ISL_6988516 | Respiratory Virus Unit, Microbiology Services Colindale, Public Health England | COVID-19 Genomics UK (COG-UK) Consortium | PHE Covid Sequencing Team et al |
| EPI_ISL_3950909 | Lighthouse Laboratory Plymouth | Wellcome Sanger Institute for the COVID-19 Genomics UK (COG-UK) Consortium | Lighthouse Laboratory Plymouth et al |
| EPI_ISL_5024354 | Lighthouse Laboratory Plymouth | Wellcome Sanger Institute for the COVID-19 Genomics UK (COG-UK) Consortium | Lighthouse Laboratory Plymouth et al |
| EPI_ISL_5488102 | Respiratory Virus Unit, Microbiology Services Colindale, Public Health England | COVID-19 Genomics UK (COG-UK) Consortium | PHE Covid Sequencing Team et al |
| EPI_ISL_5459142 | Lighthouse Laboratory Plymouth | Wellcome Sanger Institute for the COVID-19 Genomics UK (COG-UK) Consortium | Lighthouse Laboratory Plymouth et al |
| EPI_ISL_5813828 | Respiratory Virus Unit, Microbiology Services Colindale, Public Health England | COVID-19 Genomics UK (COG-UK) Consortium | PHE Covid Sequencing Team et al |
| EPI_ISL_4453836 | Lighthouse Laboratory Plymouth | Wellcome Sanger Institute for the COVID-19 Genomics UK (COG-UK) Consortium | Lighthouse Laboratory Plymouth et al |
| EPI_ISL_4752246 | Lighthouse Laboratory Plymouth | Wellcome Sanger Institute for the COVID-19 Genomics UK (COG-UK) Consortium | Lighthouse Laboratory Plymouth et al |
| EPI_ISL_3296560 | Wales Specialist Virology Centre Sequencing lab: Pathogen Genomics Unit | Public Health Wales Microbiology Cardiff Wales Specialist Virology Centre | Catherine Moore et al |
| EPI_ISL_3468111 | Lighthouse Laboratory Plymouth | Wellcome Sanger Institute for the COVID-19 Genomics UK (COG-UK) Consortium | Lighthouse Laboratory Plymouth et al |
| EPI_ISL_3202832 | Lighthouse Laboratory Plymouth | Wellcome Sanger Institute for the COVID-19 Genomics UK (COG-UK) Consortium | Lighthouse Laboratory Plymouth et al |
| EPI_ISL_2850075 | Lighthouse Laboratory Plymouth | Wellcome Sanger Institute for the COVID-19 Genomics UK (COG-UK) Consortium | Lighthouse Laboratory Plymouth et al |
| EPI_ISL_3765156 | Lighthouse Laboratory Plymouth | Wellcome Sanger Institute for the COVID-19 Genomics UK (COG-UK) Consortium | Lighthouse Laboratory Plymouth et al |
| EPI_ISL_3094752 | Lighthouse Laboratory Plymouth | Wellcome Sanger Institute for the COVID-19 Genomics UK (COG-UK) Consortium | Lighthouse Laboratory Plymouth et al |
| EPI_ISL_3068727 | Lighthouse Laboratory Plymouth | Wellcome Sanger Institute for the COVID-19 Genomics UK (COG-UK) Consortium | Lighthouse Laboratory Plymouth et al |

| | | | |
|---|---|---|---|
| EPI_ISL_3786684 | Originating lab: Wales Specialist Virology Centre Sequencing lab: Pathogen Genomics Unit | Public Health Wales Microbiology Cardiff Wales Specialist Virology Centre | Catherine Moore et al |
| EPI_ISL_4086758 | Lighthouse Laboratory Plymouth | Wellcome Sanger Institute for the COVID-19 Genomics UK (COG-UK) Consortium | Lighthouse Laboratory Plymouth et al |
| EPI_ISL_3224159 | Lighthouse Laboratory Plymouth | Wellcome Sanger Institute for the COVID-19 Genomics UK (COG-UK) Consortium | Lighthouse Laboratory Plymouth et al |
| EPI_ISL_6361548 | Lighthouse Laboratory Plymouth | Wellcome Sanger Institute for the COVID-19 Genomics UK (COG-UK) Consortium | Lighthouse Laboratory Plymouth et al |
| EPI_ISL_3697949 | Lighthouse Laboratory Plymouth | Wellcome Sanger Institute for the COVID-19 Genomics UK (COG-UK) Consortium | Lighthouse Laboratory Plymouth et al |
| EPI_ISL_1107669 | Centre for Enzyme Innovation, University of Portsmouth / Translational Research Laboratory, Portsmouth Hospitals NHS Trust | COVID-19 Genomics UK (COG-UK) Consortium | Angela Beckett et al |
| EPI_ISL_6772267 | Lighthouse Laboratory Plymouth | Wellcome Sanger Institute for the COVID-19 Genomics UK (COG-UK) Consortium | Lighthouse Laboratory Plymouth et al |
| EPI_ISL_3283852 | Lighthouse Laboratory Plymouth | Wellcome Sanger Institute for the COVID-19 Genomics UK (COG-UK) Consortium | Lighthouse Laboratory Plymouth et al |
| EPI_ISL_2935647 | Lighthouse Laboratory Plymouth | Wellcome Sanger Institute for the COVID-19 Genomics UK (COG-UK) Consortium | Lighthouse Laboratory Plymouth et al |
| EPI_ISL_3295335 | Wales Specialist Virology Centre Sequencing lab: Pathogen Genomics Unit | Public Health Wales Microbiology Cardiff Wales Specialist Virology Centre | Catherine Moore et al |
| EPI_ISL_3574976 | Originating lab: Wales Specialist Virology Centre Sequencing lab: Pathogen Genomics Unit | Public Health Wales Microbiology Cardiff Wales Specialist Virology Centre | Catherine Moore et al |
| EPI_ISL_3469631 | Lighthouse Laboratory Plymouth | Wellcome Sanger Institute for the COVID-19 Genomics UK (COG-UK) Consortium | Lighthouse Laboratory Plymouth et al |
| EPI_ISL_3962170 | Respiratory Virus Unit, Microbiology Services Colindale, Public Health England | COVID-19 Genomics UK (COG-UK) Consortium | PHE Covid Sequencing Team et al |
| EPI_ISL_2808469 | Lighthouse Laboratory Plymouth | Wellcome Sanger Institute for the COVID-19 Genomics UK (COG-UK) Consortium | Lighthouse Laboratory Plymouth et al |
| EPI_ISL_6765524 | Lighthouse Laboratory Plymouth | Wellcome Sanger Institute for the COVID-19 Genomics UK (COG-UK) Consortium | Lighthouse Laboratory Plymouth et al |

| | | | |
|---|---|---|---|
| EPI_ISL_3728524 | Lighthouse Laboratory Plymouth | Wellcome Sanger Institute for the COVID-19 Genomics UK (COG-UK) Consortium | Lighthouse Laboratory Plymouth et al |
| EPI_ISL_5488667 | Respiratory Virus Unit, Microbiology Services Colindale, Public Health England | COVID-19 Genomics UK (COG-UK) Consortium | PHE Covid Sequencing Team et al |
| EPI_ISL_2846291 | Lighthouse Laboratory Plymouth | Wellcome Sanger Institute for the COVID-19 Genomics UK (COG-UK) Consortium | Lighthouse Laboratory Plymouth et al |
| EPI_ISL_3728942 | Lighthouse Laboratory Plymouth | Wellcome Sanger Institute for the COVID-19 Genomics UK (COG-UK) Consortium | Lighthouse Laboratory Plymouth et al |
| EPI_ISL_3810593 | Lighthouse Laboratory Plymouth | Wellcome Sanger Institute for the COVID-19 Genomics UK (COG-UK) Consortium | Lighthouse Laboratory Plymouth et al |
| EPI_ISL_3884587 | Lighthouse Laboratory Plymouth | Wellcome Sanger Institute for the COVID-19 Genomics UK (COG-UK) Consortium | Lighthouse Laboratory Plymouth et al |
| EPI_ISL_5862502 | Lighthouse Laboratory Plymouth | Wellcome Sanger Institute for the COVID-19 Genomics UK (COG-UK) Consortium | Lighthouse Laboratory Plymouth et al |
| EPI_ISL_3765100 | Lighthouse Laboratory Plymouth | Wellcome Sanger Institute for the COVID-19 Genomics UK (COG-UK) Consortium | Lighthouse Laboratory Plymouth et al |
| EPI_ISL_4129382 | Respiratory Virus Unit, Microbiology Services Colindale, Public Health England | COVID-19 Genomics UK (COG-UK) Consortium | PHE Covid Sequencing Team et al |
| EPI_ISL_5489241 | Respiratory Virus Unit, Microbiology Services Colindale, Public Health England | COVID-19 Genomics UK (COG-UK) Consortium | PHE Covid Sequencing Team et al |
| EPI_ISL_1107142 | Wales Specialist Virology Centre Sequencing lab: Pathogen Genomics Unit | Public Health Wales Microbiology Cardiff Wales Specialist Virology Centre | Catherine Moore et al |
| EPI_ISL_6428556 | Lighthouse Laboratory Plymouth | Wellcome Sanger Institute for the COVID-19 Genomics UK (COG-UK) Consortium | Lighthouse Laboratory Plymouth et al |
| EPI_ISL_5485116 | Respiratory Virus Unit, Microbiology Services Colindale, Public Health England | COVID-19 Genomics UK (COG-UK) Consortium | PHE Covid Sequencing Team et al |
| EPI_ISL_4752175 | Lighthouse Laboratory Plymouth | Wellcome Sanger Institute for the COVID-19 Genomics UK (COG-UK) Consortium | Lighthouse Laboratory Plymouth et al |
| EPI_ISL_6806519 | Lighthouse Laboratory Plymouth | Wellcome Sanger Institute for the COVID-19 Genomics UK (COG-UK) Consortium | Lighthouse Laboratory Plymouth et al |

| | | | |
|---|---|---|---|
| EPI_ISL_4128602 | Respiratory Virus Unit, Microbiology Services Colindale, Public Health England | COVID-19 Genomics UK (COG-UK) Consortium | PHE Covid Sequencing Team et al |
| EPI_ISL_4129875 | Respiratory Virus Unit, Microbiology Services Colindale, Public Health England | COVID-19 Genomics UK (COG-UK) Consortium | PHE Covid Sequencing Team et al |
| EPI_ISL_6106732 | Lighthouse Laboratory Plymouth | Wellcome Sanger Institute for the COVID-19 Genomics UK (COG-UK) Consortium | Lighthouse Laboratory Plymouth et al |
| EPI_ISL_5838759 | Lighthouse Laboratory Plymouth | Wellcome Sanger Institute for the COVID-19 Genomics UK (COG-UK) Consortium | Lighthouse Laboratory Plymouth et al |
| EPI_ISL_3701945 | Lighthouse Laboratory Plymouth | Wellcome Sanger Institute for the COVID-19 Genomics UK (COG-UK) Consortium | Lighthouse Laboratory Plymouth et al |
| EPI_ISL_2996504 | Lighthouse Laboratory Plymouth | Wellcome Sanger Institute for the COVID-19 Genomics UK (COG-UK) Consortium | Lighthouse Laboratory Plymouth et al |
| EPI_ISL_3882808 | Lighthouse Laboratory Plymouth | Wellcome Sanger Institute for the COVID-19 Genomics UK (COG-UK) Consortium | Lighthouse Laboratory Plymouth et al |
| EPI_ISL_2917347 | Lighthouse Laboratory Plymouth | Wellcome Sanger Institute for the COVID-19 Genomics UK (COG-UK) Consortium | Lighthouse Laboratory Plymouth et al |
| EPI_ISL_6106813 | Lighthouse Laboratory Plymouth | Wellcome Sanger Institute for the COVID-19 Genomics UK (COG-UK) Consortium | Lighthouse Laboratory Plymouth et al |
| EPI_ISL_4010874 | Lighthouse Laboratory Plymouth | Wellcome Sanger Institute for the COVID-19 Genomics UK (COG-UK) Consortium | Lighthouse Laboratory Plymouth et al |
| EPI_ISL_4014683 | Lighthouse Laboratory Plymouth | Wellcome Sanger Institute for the COVID-19 Genomics UK (COG-UK) Consortium | Lighthouse Laboratory Plymouth et al |
| EPI_ISL_3728348 | Lighthouse Laboratory Plymouth | Wellcome Sanger Institute for the COVID-19 Genomics UK (COG-UK) Consortium | Lighthouse Laboratory Plymouth et al |
| EPI_ISL_2639155 | Lighthouse Laboratory Plymouth | Wellcome Sanger Institute for the COVID-19 Genomics UK (COG-UK) Consortium | Lighthouse Laboratory Plymouth et al |
| EPI_ISL_3698759 | Lighthouse Laboratory Plymouth | Wellcome Sanger Institute for the COVID-19 Genomics UK (COG-UK) Consortium | Lighthouse Laboratory Plymouth et al |
| EPI_ISL_5485216 | Respiratory Virus Unit, Microbiology Services Colindale, Public Health England | COVID-19 Genomics UK (COG-UK) Consortium | PHE Covid Sequencing Team et al |
| EPI_ISL_6137524 | Respiratory Virus Unit, Microbiology Services Colindale, Public Health England | COVID-19 Genomics UK (COG-UK) Consortium | PHE Covid Sequencing Team et al |

| | | | |
|---|---|---|---|
| EPI_ISL_3703592 | Lighthouse Laboratory Plymouth | Wellcome Sanger Institute for the COVID-19 Genomics UK (COG-UK) Consortium | Lighthouse Laboratory Plymouth et al |
| EPI_ISL_4089523 | Lighthouse Laboratory Plymouth | Wellcome Sanger Institute for the COVID-19 Genomics UK (COG-UK) Consortium | Lighthouse Laboratory Plymouth et al |
| EPI_ISL_1331841 | Lighthouse Lab in Glasgow | Wellcome Sanger Institute for the COVID-19 Genomics UK (COG-UK) Consortium | Harper VanSteenhouse et al |
| EPI_ISL_923824 | Department of Pathology, University of Cambridge | COVID-19 Genomics UK (COG-UK) Consortium | Aminu S. Jahun et al |
| EPI_ISL_6297463 | Lighthouse Lab in Glasgow | Wellcome Sanger Institute for the COVID-19 Genomics UK (COG-UK) Consortium | Harper VanSteenhouse et al |
| EPI_ISL_5793025 | Lighthouse Lab in Glasgow | Wellcome Sanger Institute for the COVID-19 Genomics UK (COG-UK) Consortium | Harper VanSteenhouse et al |
| EPI_ISL_3043241 | Lighthouse Lab in Glasgow | Wellcome Sanger Institute for the COVID-19 Genomics UK (COG-UK) Consortium | Harper VanSteenhouse et al |
| EPI_ISL_2830846 | Lighthouse Lab in Glasgow | Wellcome Sanger Institute for the COVID-19 Genomics UK (COG-UK) Consortium | Harper VanSteenhouse et al |
| EPI_ISL_3639231 | Lighthouse Lab in Glasgow | Wellcome Sanger Institute for the COVID-19 Genomics UK (COG-UK) Consortium | Harper VanSteenhouse et al |
| EPI_ISL_2847079 | Lighthouse Lab in Glasgow | Wellcome Sanger Institute for the COVID-19 Genomics UK (COG-UK) Consortium | Harper VanSteenhouse et al |
| EPI_ISL_1698839 | Lighthouse Lab in Glasgow | Wellcome Sanger Institute for the COVID-19 Genomics UK (COG-UK) Consortium | Harper VanSteenhouse et al |
| EPI_ISL_1263581 | Lighthouse Lab in Glasgow | Wellcome Sanger Institute for the COVID-19 Genomics UK (COG-UK) Consortium | Harper VanSteenhouse et al |
| EPI_ISL_1331967 | Lighthouse Lab in Glasgow | Wellcome Sanger Institute for the COVID-19 Genomics UK (COG-UK) Consortium | Harper VanSteenhouse et al |
| EPI_ISL_2539709 | Lighthouse Lab in Glasgow | Wellcome Sanger Institute for the COVID-19 Genomics UK (COG-UK) Consortium | Harper VanSteenhouse et al |
| EPI_ISL_3990587 | Lighthouse Lab in Glasgow | Wellcome Sanger Institute for the COVID-19 Genomics UK (COG-UK) Consortium | Harper VanSteenhouse et al |
| EPI_ISL_2916756 | Lighthouse Lab in Glasgow | Wellcome Sanger Institute for the COVID-19 Genomics UK (COG-UK) Consortium | Harper VanSteenhouse et al |
| EPI_ISL_1342107 | Lighthouse Lab in Glasgow | Wellcome Sanger Institute for the COVID-19 Genomics UK (COG-UK) Consortium | Harper VanSteenhouse et al |

| | | | |
|---|---|---|---|
| EPI_ISL_1519178 | Lighthouse Lab in Glasgow | Wellcome Sanger Institute for the COVID-19 Genomics UK (COG-UK) Consortium | Harper VanSteenhouse et al |
| EPI_ISL_4289224 | Lighthouse Lab in Glasgow | Wellcome Sanger Institute for the COVID-19 Genomics UK (COG-UK) Consortium | Harper VanSteenhouse et al |
| EPI_ISL_3223285 | Lighthouse Lab in Glasgow | Wellcome Sanger Institute for the COVID-19 Genomics UK (COG-UK) Consortium | Harper VanSteenhouse et al |
| EPI_ISL_3013822 | Lighthouse Lab in Glasgow | Wellcome Sanger Institute for the COVID-19 Genomics UK (COG-UK) Consortium | Harper VanSteenhouse et al |
| EPI_ISL_3094394 | Lighthouse Lab in Glasgow | Wellcome Sanger Institute for the COVID-19 Genomics UK (COG-UK) Consortium | Harper VanSteenhouse et al |
| EPI_ISL_1242560 | Lighthouse Lab in Glasgow | Wellcome Sanger Institute for the COVID-19 Genomics UK (COG-UK) Consortium | Harper VanSteenhouse et al |
| EPI_ISL_1594851 | Lighthouse Lab in Glasgow | Wellcome Sanger Institute for the COVID-19 Genomics UK (COG-UK) Consortium | Harper VanSteenhouse et al |
| EPI_ISL_1741201 | Lighthouse Lab in Glasgow | Wellcome Sanger Institute for the COVID-19 Genomics UK (COG-UK) Consortium | Harper VanSteenhouse et al |
| EPI_ISL_2352920 | Lighthouse Lab in Glasgow | Wellcome Sanger Institute for the COVID-19 Genomics UK (COG-UK) Consortium | Harper VanSteenhouse et al |
| EPI_ISL_1188483 | Lighthouse Lab in Glasgow | Wellcome Sanger Institute for the COVID-19 Genomics UK (COG-UK) Consortium | Harper VanSteenhouse et al |
| EPI_ISL_907534 | Lighthouse Lab in Glasgow | Wellcome Sanger Institute for the COVID-19 Genomics UK (COG-UK) Consortium | Harper VanSteenhouse et al |
| EPI_ISL_1489193 | Lighthouse Lab in Glasgow | Wellcome Sanger Institute for the COVID-19 Genomics UK (COG-UK) Consortium | Harper VanSteenhouse et al |
| EPI_ISL_3312227 | Lighthouse Lab in Glasgow | Wellcome Sanger Institute for the COVID-19 Genomics UK (COG-UK) Consortium | Harper VanSteenhouse et al |
| EPI_ISL_3764604 | Lighthouse Lab in Glasgow | Wellcome Sanger Institute for the COVID-19 Genomics UK (COG-UK) Consortium | Harper VanSteenhouse et al |
| EPI_ISL_1409204 | Lighthouse Lab in Glasgow | Wellcome Sanger Institute for the COVID-19 Genomics UK (COG-UK) Consortium | Harper VanSteenhouse et al |
| EPI_ISL_2569174 | Lighthouse Lab in Glasgow | Wellcome Sanger Institute for the COVID-19 Genomics UK (COG-UK) Consortium | Harper VanSteenhouse et al |
| EPI_ISL_4351200 | Lighthouse Lab in Glasgow | Wellcome Sanger Institute for the COVID-19 Genomics UK (COG-UK) Consortium | Harper VanSteenhouse et al |

| | | | |
|---|---|---|---|
| EPI_ISL_2006196 | Lighthouse Lab in Glasgow | Wellcome Sanger Institute for the COVID-19 Genomics UK (COG-UK) Consortium | Harper VanSteenhouse et al |
| EPI_ISL_5837385 | Lighthouse Lab in Glasgow | Wellcome Sanger Institute for the COVID-19 Genomics UK (COG-UK) Consortium | Harper VanSteenhouse et al |
| EPI_ISL_1264389 | Lighthouse Lab in Glasgow | Wellcome Sanger Institute for the COVID-19 Genomics UK (COG-UK) Consortium | Harper VanSteenhouse et al |
| EPI_ISL_3993881 | Lighthouse Lab in Glasgow | Wellcome Sanger Institute for the COVID-19 Genomics UK (COG-UK) Consortium | Harper VanSteenhouse et al |
| EPI_ISL_2779885 | Lighthouse Lab in Glasgow | Wellcome Sanger Institute for the COVID-19 Genomics UK (COG-UK) Consortium | Harper VanSteenhouse et al |
| EPI_ISL_2730574 | Lighthouse Lab in Glasgow | Wellcome Sanger Institute for the COVID-19 Genomics UK (COG-UK) Consortium | Harper VanSteenhouse et al |
| EPI_ISL_2702687 | Lighthouse Lab in Glasgow | Wellcome Sanger Institute for the COVID-19 Genomics UK (COG-UK) Consortium | Harper VanSteenhouse et al |
| EPI_ISL_4942640 | Lighthouse Lab in Glasgow | Wellcome Sanger Institute for the COVID-19 Genomics UK (COG-UK) Consortium | Harper VanSteenhouse et al |
| EPI_ISL_2851171 | Lighthouse Lab in Glasgow | Wellcome Sanger Institute for the COVID-19 Genomics UK (COG-UK) Consortium | Harper VanSteenhouse et al |
| EPI_ISL_1245208 | Lighthouse Lab in Glasgow | Wellcome Sanger Institute for the COVID-19 Genomics UK (COG-UK) Consortium | Harper VanSteenhouse et al |
| EPI_ISL_5991581 | Lighthouse Lab in Glasgow | Wellcome Sanger Institute for the COVID-19 Genomics UK (COG-UK) Consortium | Harper VanSteenhouse et al |
| EPI_ISL_1083095 | Lighthouse Lab in Glasgow | Wellcome Sanger Institute for the COVID-19 Genomics UK (COG-UK) Consortium | Harper VanSteenhouse et al |
| EPI_ISL_1466194 | Lighthouse Lab in Glasgow | Wellcome Sanger Institute for the COVID-19 Genomics UK (COG-UK) Consortium | Harper VanSteenhouse et al |
| EPI_ISL_2514793 | Lighthouse Lab in Glasgow | Wellcome Sanger Institute for the COVID-19 Genomics UK (COG-UK) Consortium | Harper VanSteenhouse et al |
| EPI_ISL_1274465 | Lighthouse Lab in Glasgow | Wellcome Sanger Institute for the COVID-19 Genomics UK (COG-UK) Consortium | Harper VanSteenhouse et al |
| EPI_ISL_5892551 | Lighthouse Lab in Glasgow | Wellcome Sanger Institute for the COVID-19 Genomics UK (COG-UK) Consortium | Harper VanSteenhouse et al |
| EPI_ISL_5479316 | Lighthouse Lab in Glasgow | Wellcome Sanger Institute for the COVID-19 Genomics UK (COG-UK) Consortium | Harper VanSteenhouse et al |

| | | | |
|---|---|---|---|
| EPI_ISL_3992452 | Lighthouse Lab in Glasgow | Wellcome Sanger Institute for the COVID-19 Genomics UK (COG-UK) Consortium | Harper VanSteenhouse et al |
| EPI_ISL_1452754 | Lighthouse Lab in Glasgow | Wellcome Sanger Institute for the COVID-19 Genomics UK (COG-UK) Consortium | Harper VanSteenhouse et al |
| EPI_ISL_1332371 | Lighthouse Lab in Glasgow | Wellcome Sanger Institute for the COVID-19 Genomics UK (COG-UK) Consortium | Harper VanSteenhouse et al |
| EPI_ISL_1331128 | Lighthouse Lab in Glasgow | Wellcome Sanger Institute for the COVID-19 Genomics UK (COG-UK) Consortium | Harper VanSteenhouse et al |
| EPI_ISL_2238208 | Lighthouse Lab in Glasgow | Wellcome Sanger Institute for the COVID-19 Genomics UK (COG-UK) Consortium | Harper VanSteenhouse et al |
| EPI_ISL_1857690 | Lighthouse Lab in Glasgow | Wellcome Sanger Institute for the COVID-19 Genomics UK (COG-UK) Consortium | Harper VanSteenhouse et al |
| EPI_ISL_1070517 | Lighthouse Lab in Glasgow | Wellcome Sanger Institute for the COVID-19 Genomics UK (COG-UK) Consortium | Harper VanSteenhouse et al |
| EPI_ISL_3126139 | Lighthouse Lab in Glasgow | Wellcome Sanger Institute for the COVID-19 Genomics UK (COG-UK) Consortium | Harper VanSteenhouse et al |
| EPI_ISL_6107339 | Lighthouse Lab in Glasgow | Wellcome Sanger Institute for the COVID-19 Genomics UK (COG-UK) Consortium | Harper VanSteenhouse et al |
| EPI_ISL_3123944 | Lighthouse Lab in Glasgow | Wellcome Sanger Institute for the COVID-19 Genomics UK (COG-UK) Consortium | Harper VanSteenhouse et al |
| EPI_ISL_3226004 | Lighthouse Lab in Glasgow | Wellcome Sanger Institute for the COVID-19 Genomics UK (COG-UK) Consortium | Harper VanSteenhouse et al |
| EPI_ISL_5963290 | Lighthouse Lab in Glasgow | Wellcome Sanger Institute for the COVID-19 Genomics UK (COG-UK) Consortium | Harper VanSteenhouse et al |
| EPI_ISL_1310613 | Department of Pathology, University of Cambridge | COVID-19 Genomics UK (COG-UK) Consortium | Aminu S. Jahun et al |
| EPI_ISL_1330177 | Lighthouse Lab in Glasgow | Wellcome Sanger Institute for the COVID-19 Genomics UK (COG-UK) Consortium | Harper VanSteenhouse et al |
| EPI_ISL_3993799 | Lighthouse Lab in Glasgow | Wellcome Sanger Institute for the COVID-19 Genomics UK (COG-UK) Consortium | Harper VanSteenhouse et al |
| EPI_ISL_2830986 | Lighthouse Lab in Glasgow | Wellcome Sanger Institute for the COVID-19 Genomics UK (COG-UK) Consortium | Harper VanSteenhouse et al |
| EPI_ISL_3124831 | Lighthouse Lab in Glasgow | Wellcome Sanger Institute for the COVID-19 Genomics UK (COG-UK) Consortium | Harper VanSteenhouse et al |

| | | | |
|---|---|---|---|
| EPI_ISL_4974313 | Lighthouse Lab in Glasgow | Wellcome Sanger Institute for the COVID-19 Genomics UK (COG-UK) Consortium | Harper VanSteenhouse et al |
| EPI_ISL_2487329 | Lighthouse Lab in Glasgow | Wellcome Sanger Institute for the COVID-19 Genomics UK (COG-UK) Consortium | Harper VanSteenhouse et al |
| EPI_ISL_5097485 | Lighthouse Lab in Glasgow | Wellcome Sanger Institute for the COVID-19 Genomics UK (COG-UK) Consortium | Harper VanSteenhouse et al |
| EPI_ISL_1375815 | Lighthouse Lab in Glasgow | Wellcome Sanger Institute for the COVID-19 Genomics UK (COG-UK) Consortium | Harper VanSteenhouse et al |
| EPI_ISL_2004920 | Lighthouse Lab in Glasgow | Wellcome Sanger Institute for the COVID-19 Genomics UK (COG-UK) Consortium | Harper VanSteenhouse et al |
| EPI_ISL_3360808 | Lighthouse Lab in Glasgow | Wellcome Sanger Institute for the COVID-19 Genomics UK (COG-UK) Consortium | Harper VanSteenhouse et al |
| EPI_ISL_3726237 | Lighthouse Lab in Glasgow | Wellcome Sanger Institute for the COVID-19 Genomics UK (COG-UK) Consortium | Harper VanSteenhouse et al |
| EPI_ISL_3096002 | Lighthouse Lab in Glasgow | Wellcome Sanger Institute for the COVID-19 Genomics UK (COG-UK) Consortium | Harper VanSteenhouse et al |
| EPI_ISL_3168088 | Lighthouse Lab in Glasgow | Wellcome Sanger Institute for the COVID-19 Genomics UK (COG-UK) Consortium | Harper VanSteenhouse et al |
| EPI_ISL_3766118 | Lighthouse Lab in Glasgow | Wellcome Sanger Institute for the COVID-19 Genomics UK (COG-UK) Consortium | Harper VanSteenhouse et al |
| EPI_ISL_3367730 | Lighthouse Lab in Glasgow | Wellcome Sanger Institute for the COVID-19 Genomics UK (COG-UK) Consortium | Harper VanSteenhouse et al |
| EPI_ISL_4734554 | Lighthouse Lab in Glasgow | Wellcome Sanger Institute for the COVID-19 Genomics UK (COG-UK) Consortium | Harper VanSteenhouse et al |
| EPI_ISL_6351080 | Lighthouse Lab in Glasgow | Wellcome Sanger Institute for the COVID-19 Genomics UK (COG-UK) Consortium | Harper VanSteenhouse et al |
| EPI_ISL_3764720 | Lighthouse Lab in Glasgow | Wellcome Sanger Institute for the COVID-19 Genomics UK (COG-UK) Consortium | Harper VanSteenhouse et al |
| EPI_ISL_2935923 | Lighthouse Lab in Glasgow | Wellcome Sanger Institute for the COVID-19 Genomics UK (COG-UK) Consortium | Harper VanSteenhouse et al |
| EPI_ISL_1243676 | Lighthouse Lab in Glasgow | Wellcome Sanger Institute for the COVID-19 Genomics UK (COG-UK) Consortium | Harper VanSteenhouse et al |
| EPI_ISL_2995405 | Lighthouse Lab in Glasgow | Wellcome Sanger Institute for the COVID-19 Genomics UK (COG-UK) Consortium | Harper VanSteenhouse et al |

| | | | |
|---|---|---|---|
| EPI_ISL_1188306 | Lighthouse Lab in Glasgow | Wellcome Sanger Institute for the COVID-19 Genomics UK (COG-UK) Consortium | Harper VanSteenhouse et al |
| EPI_ISL_1409144 | Lighthouse Lab in Glasgow | Wellcome Sanger Institute for the COVID-19 Genomics UK (COG-UK) Consortium | Harper VanSteenhouse et al |
| EPI_ISL_3283267 | Lighthouse Lab in Glasgow | Wellcome Sanger Institute for the COVID-19 Genomics UK (COG-UK) Consortium | Harper VanSteenhouse et al |
| EPI_ISL_1342552 | Lighthouse Lab in Glasgow | Wellcome Sanger Institute for the COVID-19 Genomics UK (COG-UK) Consortium | Harper VanSteenhouse et al |
| EPI_ISL_432849 | Virology Department, Sheffield Teaching Hospitals NHS Foundation Trust / Virology Department, Sheffield Teaching Hospitals NHS Foundation Trust | COVID-19 Genomics UK (COG-UK) Consortium | Thushan de Silva et al |
| EPI_ISL_599153 | Lighthouse Lab in Glasgow | Wellcome Sanger Institute for the COVID-19 Genomics UK (COG-UK) consortium | Harper VanSteenhouse et al |
| EPI_ISL_1504730 | Randox Laboratories | Wellcome Sanger Institute for the COVID-19 Genomics UK (COG-UK) Consortium | Randox Laboratories et al |
| EPI_ISL_6761433 | Lighthouse Lab in Glasgow | Wellcome Sanger Institute for the COVID-19 Genomics UK (COG-UK) Consortium | Harper VanSteenhouse et al |
| EPI_ISL_842170 | Oxford Viromics, NDM, University of Oxford; Oxford University Hospitals; Basingstoke and North Hampshire Hospital | COVID-19 Genomics UK (COG-UK) Consortium | Tanya Golubchik et al |
| EPI_ISL_2645239 | Seinäjoen keskussairaala | Expert Microbiology, National Institute for Health and Welfare | Soile Blomqvist et al |
| EPI_ISL_633932 | Lighthouse Lab in Glasgow | Wellcome Sanger Institute for the COVID-19 Genomics UK (COG-UK) consortium | Harper VanSteenhouse et al |
| EPI_ISL_709334 | Lighthouse Lab in Glasgow | Wellcome Sanger Institute for the COVID-19 Genomics UK (COG-UK) Consortium | Harper VanSteenhouse et al |
| EPI_ISL_3230706 | TYKS, Kliininen mikrobiologia | Expert Microbiology, National Institute for Health and Welfare | Soile Blomqvist et al |
| EPI_ISL_1371769 | CH METROPOLE SAVOIE | CNR Virus des Infections Respiratoires - France SUD | Antonin Bal et al |
| EPI_ISL_2107166 | Synlab Eesti OÜ | 1. Laboratory of Communicable Diseases (Estonia); 2. Eurofins Genomics Europe Sequencing GmbH | Liidia Dotsenko et al. |
| EPI_ISL_642087 | Lighthouse Lab in Glasgow | Wellcome Sanger Institute for the COVID-19 Genomics UK (COG-UK) Consortium | Harper VanSteenhouse et al |

| | | | |
|---|---|---|---|
| EPI_ISL_1228966 | Cerballiance | CNR Virus des Infections Respiratoires - France SUD | Antonin Bal et al |
| EPI_ISL_822265 | Lighthouse Lab in Glasgow | Wellcome Sanger Institute for the COVID-19 Genomics UK (COG-UK) Consortium | Harper VanSteenhouse et al |
| EPI_ISL_952993 | Virology Department, Sheffield Teaching Hospitals NHS Foundation Trust/Department of Infection, Immunity and Cardiovascular Disease, The Medical School, University of Sheffield | COVID-19 Genomics UK (COG-UK) Consortium | Thushan de Silva et al |
| EPI_ISL_530569 | Lighthouse Lab in Glasgow | Wellcome Sanger Institute for the COVID-19 Genomics UK (COG-UK) consortium | Harper VanSteenhouse et al |
| EPI_ISL_3577112 | Bioinformatics and Biostatistics Lab, Advanced Sequencing Facility | COVID-19 Genomics UK (COG-UK) Consortium | Aengus Stewart et al |
| EPI_ISL_567064 | Lighthouse Lab in Glasgow | Wellcome Sanger Institute for the COVID-19 Genomics UK (COG-UK) consortium | Harper VanSteenhouse et al |
| EPI_ISL_2506573 | Islab, Pohjois-Savon aluelaboratorio | Expert Microbiology, National Institute for Health and Welfare | Soile Blomqvist et al |
| EPI_ISL_797734 | Lighthouse Lab in Glasgow | Wellcome Sanger Institute for the COVID-19 Genomics UK (COG-UK) Consortium | Harper VanSteenhouse et al |
| EPI_ISL_442435 | Virology Department, Sheffield Teaching Hospitals NHS Foundation Trust/Department of Infection, Immunity and Cardiovascular Disease, The Medical School, University of Sheffield | COVID-19 Genomics UK (COG-UK) Consortium | Thushan de Silva et al |
| EPI_ISL_2643383 | SYNLAB Eesti OÜ | Department of Microbiology, Institute of Biomedicine and Translational Medicine, University of Tartu | Radko Avi et al |
| EPI_ISL_661837 | Lighthouse Lab in Glasgow | Wellcome Sanger Institute for the COVID-19 Genomics UK (COG-UK) Consortium | Harper VanSteenhouse et al |
| EPI_ISL_1673312 | Laboratorio de Investigaciones de Baney | Swiss Tropical and Public Health Institute | Salome Hosch et al |
| EPI_ISL_1672557 | Laboratorio de Investigaciones de Baney | Swiss Tropical and Public Health Institute | Carlos Cortes et al |
| EPI_ISL_532554 | Lighthouse Lab in Glasgow | Wellcome Sanger Institute for the COVID-19 Genomics UK (COG-UK) consortium | Harper VanSteenhouse et al |
| EPI_ISL_6900909 | NordLab Rovaniemi | Expert Microbiology, National Institute for Health and Welfare | Soile Blomqvist et al |

| | | | |
|---|---|---|---|
| EPI_ISL_1389073 | Virology Department, Sheffield Teaching Hospitals NHS Foundation Trust/Department of Infection, Immunity and Cardiovascular Disease, The Medical School, University of Sheffield | COVID-19 Genomics UK (COG-UK) Consortium | Thushan de Silva et al |
| EPI_ISL_1070178 | Randox Laboratories | Wellcome Sanger Institute for the COVID-19 Genomics UK (COG-UK) Consortium | Randox Laboratories et al |
| EPI_ISL_1841884 | Department of Virology and Immunology, University of Helsinki and Helsinki University Hospital, Huslab Finland | Department of Virology, Faculty of Medicine, University of Helsinki, Helsinki, Finland | Teemu Smura et al |
| EPI_ISL_648369 | Laboratorio de Investigaciones de Baney | University Hospital Basel, Clinical Bacteriology | Carlos Cortes et al |
| EPI_ISL_3006547 | Virology Department, Sheffield Teaching Hospitals NHS Foundation Trust/Department of Infection, Immunity and Cardiovascular Disease, The Medical School, University of Sheffield | COVID-19 Genomics UK (COG-UK) Consortium | Thushan de Silva et al |
| EPI_ISL_2609142 | Department of Virology and Immunology, University of Helsinki and Helsinki University Hospital, Huslab Finland | Department of Virology, Faculty of Medicine, University of Helsinki, Helsinki, Finland | Teemu Smura et al |
| EPI_ISL_7032755 | Lighthouse Lab in Glasgow | Wellcome Sanger Institute for the COVID-19 Genomics UK (COG-UK) Consortium | Harper VanSteenhouse et al |
| EPI_ISL_2642831 | SYNLAB Eesti OÜ | Department of Microbiology, Institute of Biomedicine and Translational Medicine, University of Tartu | Radko Avi et al |
| EPI_ISL_769252 | Lighthouse Lab in Glasgow | Wellcome Sanger Institute for the COVID-19 Genomics UK (COG-UK) Consortium | Harper VanSteenhouse et al |
| EPI_ISL_1497067 | Department of Virology and Immunology, University of Helsinki and Helsinki University Hospital, Huslab Finland | Department of Virology, Faculty of Medicine, University of Helsinki, Helsinki, Finland | Teemu Smura et al |
| EPI_ISL_637813 | Department of Pathology, University of Cambridge | COVID-19 Genomics UK (COG-UK) Consortium | Aminu S. Jahun et al |
| EPI_ISL_668066 | Lighthouse Lab in Glasgow | Wellcome Sanger Institute for the COVID-19 Genomics UK (COG-UK) Consortium | Harper VanSteenhouse et al |

| | | | |
|---|---|---|---|
| EPI_ISL_1242448 | Randox Laboratories | Wellcome Sanger Institute for the COVID-19 Genomics UK (COG-UK) Consortium | Randox Laboratories et al |
| EPI_ISL_1470144 | Synlab Eesti OÜ | 1. Laboratory of Communicable Diseases (Estonia); 2. Eurofins Genomics Europe Sequencing GmbH | Liidia Dotsenko et al. |
| EPI_ISL_2752960 | Respiratory Virus Unit, Microbiology Services Colindale, Public Health England | COVID-19 Genomics UK (COG-UK) Consortium | PHE Covid Sequencing Team et al |
| EPI_ISL_5160946 | Department of Virology and Immunology, University of Helsinki and Helsinki University Hospital, Huslab Finland | Department of Virology, Faculty of Medicine, University of Helsinki, Helsinki, Finland | Teemu Smura et al |
| EPI_ISL_559829 | Oxford Viromics, NDM, University of Oxford; Oxford University Hospitals; Basingstoke and North Hampshire Hospital | COVID-19 Genomics UK (COG-UK) Consortium | Tanya Golubchik et al |
| EPI_ISL_6900948 | Fimlab Laboratoriot Oy Tampere | Expert Microbiology, National Institute for Health and Welfare | Soile Blomqvist et al |
| EPI_ISL_589258 | Lighthouse Lab in Glasgow | Wellcome Sanger Institute for the COVID-19 Genomics UK (COG-UK) consortium | Harper VanSteenhouse et al |
| EPI_ISL_623872 | Lighthouse Lab in Glasgow | Wellcome Sanger Institute for the COVID-19 Genomics UK (COG-UK) consortium | Harper VanSteenhouse et al |
| EPI_ISL_2752682 | Respiratory Virus Unit, Microbiology Services Colindale, Public Health England | COVID-19 Genomics UK (COG-UK) Consortium | PHE Covid Sequencing Team et al |
| EPI_ISL_782364 | Lighthouse Lab in Glasgow | Wellcome Sanger Institute for the COVID-19 Genomics UK (COG-UK) Consortium | Harper VanSteenhouse et al |
| EPI_ISL_5641370 | Islab, Etelä-Savon aluelaboratorio | Expert Microbiology, National Institute for Health and Welfare | Soile Blomqvist et al |
| EPI_ISL_842481 | Bioinformatics and Biostatistics Lab, Advanced Sequencing Facility | COVID-19 Genomics UK (COG-UK) Consortium | Aengus Stewart et al |
| EPI_ISL_782357 | Lighthouse Lab in Glasgow | Wellcome Sanger Institute for the COVID-19 Genomics UK (COG-UK) Consortium | Harper VanSteenhouse et al |
| EPI_ISL_1344475 | Randox Laboratories | Wellcome Sanger Institute for the COVID-19 Genomics UK (COG-UK) Consortium | Randox Laboratories et al |

| | | | |
|---|---|---|---|
| EPI_ISL_1980404 | Virology Department, Sheffield Teaching Hospitals NHS Foundation Trust/Department of Infection, Immunity and Cardiovascular Disease, The Medical School, University of Sheffield | COVID-19 Genomics UK (COG-UK) Consortium | Thushan de Silva et al |
| EPI_ISL_540154 | Lighthouse Lab in Glasgow | Wellcome Sanger Institute for the COVID-19 Genomics UK (COG-UK) consortium | Harper VanSteenhouse et al |
| EPI_ISL_662828 | Lighthouse Lab in Glasgow | Wellcome Sanger Institute for the COVID-19 Genomics UK (COG-UK) Consortium | Harper VanSteenhouse et al |
| EPI_ISL_517549 | Virology Department, Sheffield Teaching Hospitals NHS Foundation Trust/Department of Infection, Immunity and Cardiovascular Disease, The Medical School, University of Sheffield | COVID-19 Genomics UK (COG-UK) Consortium | Thushan de Silva et al |
| EPI_ISL_1842120 | Department of Virology and Immunology, University of Helsinki and Helsinki University Hospital, Huslab Finland | Department of Virology, Faculty of Medicine, University of Helsinki, Helsinki, Finland | Teemu Smura et al |
| EPI_ISL_3644938 | Department of Virology and Immunology, University of Helsinki and Helsinki University Hospital, Huslab Finland | Department of Virology, Faculty of Medicine, University of Helsinki, Helsinki, Finland | Teemu Smura et al |
| EPI_ISL_1899047 | Ethiopian Biotechnology Institute (EBTI) | International Centre for Genetic Engineering and Biotechnology (ICGEB) and ARGO Open Lab for Genome Sequencing | Molalegne Bitew et al |
| EPI_ISL_481654 | Department of Virology and Immunology, University of Helsinki and Helsinki University Hospital, Huslab Finland | Department of Virology, Faculty of Medicine, University of Helsinki, Helsinki, Finland | Teemu Smura et al |
| EPI_ISL_652714 | Oxford Viromics, NDM, University of Oxford; Oxford University Hospitals; Basingstoke and North Hampshire Hospital | COVID-19 Genomics UK (COG-UK) Consortium | Tanya Golubchik et al |
| EPI_ISL_814418 | Bioinformatics and Biostatistics Lab, Advanced Sequencing Facility | COVID-19 Genomics UK (COG-UK) Consortium | Aengus Stewart et al |
| EPI_ISL_6613770 | Lighthouse Lab in Glasgow | Wellcome Sanger Institute for the COVID-19 Genomics UK (COG-UK) Consortium | Harper VanSteenhouse et al |

| | | | |
|---|---|---|---|
| EPI_ISL_1374080 | Randox Laboratories | Wellcome Sanger Institute for the COVID-19 Genomics UK (COG-UK) Consortium | Randox Laboratories et al |
| EPI_ISL_3131199 | LABORATOIRE UNILIANS DECINES | CNR Virus des Infections Respiratoires - France SUD | Antonin Bal et al |
| EPI_ISL_1406629 | Centre Hospitalier Universitaire Clermont-Ferrand | CHU Clermont-Ferrand, service de virologie | Bisseux Maxime et al |
| EPI_ISL_6900584 | NordLab Oulu | Expert Microbiology, National Institute for Health and Welfare | Soile Blomqvist et al |
| EPI_ISL_2214680 | Centre hospitalier D'AURILLAC | CHU Clermont-Ferrand, service de virologie | Bisseux Maxime et al |
| EPI_ISL_5460403 | Synlab Eesti OÜ | 1. Laboratory of Communicable Diseases (Estonia); 2. Eurofins Genomics Europe Sequencing GmbH | Liidia Dotsenko et al. |
| EPI_ISL_6817268 | Lighthouse Lab in Glasgow | Wellcome Sanger Institute for the COVID-19 Genomics UK (COG-UK) Consortium | Harper VanSteenhouse et al |
| EPI_ISL_3634117 | DYOMEDEA-LABORATOIRE DE LA SAUVEGARDE | CNR Virus des Infections Respiratoires - France SUD | Antonin Bal et al |
| EPI_ISL_663019 | Lighthouse Lab in Glasgow | Wellcome Sanger Institute for the COVID-19 Genomics UK (COG-UK) Consortium | Harper VanSteenhouse et al |
| EPI_ISL_4819839 | Fiji Centre for Communicable Disease Control | Microbiological Diagnostic Unit - Public Health Laboratory (MDU-PHL) | Sahukhan et al |
| EPI_ISL_584970 | Quadram Institute Bioscience | COVID-19 Genomics UK (COG-UK) Consortium | Dave J. Baker et al |
| EPI_ISL_782721 | Lighthouse Lab in Glasgow | Wellcome Sanger Institute for the COVID-19 Genomics UK (COG-UK) Consortium | Harper VanSteenhouse et al |
| EPI_ISL_2142632 | CENTRE HOSPITALIER DU HAUT BUGEY | CNR Virus des Infections Respiratoires - France SUD | Antonin Bal et al |
| EPI_ISL_2507724 | TYKS, Kliininen mikrobiologia | Expert Microbiology, National Institute for Health and Welfare | Soile Blomqvist et al |
| EPI_ISL_6427065 | Lighthouse Lab in Glasgow | Wellcome Sanger Institute for the COVID-19 Genomics UK (COG-UK) Consortium | Harper VanSteenhouse et al |
| EPI_ISL_799398 | Lighthouse Lab in Glasgow | Wellcome Sanger Institute for the COVID-19 Genomics UK (COG-UK) Consortium | Harper VanSteenhouse et al |
| EPI_ISL_1451204 | Randox Laboratories | Wellcome Sanger Institute for the COVID-19 Genomics UK (COG-UK) Consortium | Randox Laboratories et al |
| EPI_ISL_817108 | Bioinformatics and Biostatistics Lab, Advanced Sequencing Facility | COVID-19 Genomics UK (COG-UK) Consortium | Aengus Stewart et al |
| EPI_ISL_1242387 | Randox Laboratories | Wellcome Sanger Institute for the COVID-19 Genomics UK (COG-UK) Consortium | Randox Laboratories et al |

| | | | |
|---|---|---|---|
| EPI_ISL_660034 | Lighthouse Lab in Glasgow | Wellcome Sanger Institute for the COVID-19 Genomics UK (COG-UK) Consortium | Harper VanSteenhouse et al |
| EPI_ISL_6266177 | Ethiopian Public Health Institute | EPHI Pathogen Genomic Sequence Lab | Atsbeha G Weldemariam et al |
| EPI_ISL_2604269 | Department of Virology and Immunology, University of Helsinki and Helsinki University Hospital, Huslab Finland | Department of Virology, Faculty of Medicine, University of Helsinki, Helsinki, Finland | Teemu Smura et al |
| EPI_ISL_2152523 | Laboratoire de Virologie | Department of Virology, Henri Mondor University Hospital, Assistance Publique Hôpitaux de Paris, Université Paris-Est Créteil, INSERM U955 | Christophe Rodriguez et al |
| EPI_ISL_5694000 | LABORATOIRE SAINT-SEVER | Department of Virology, Henri Mondor University Hospital, Assistance Publique Hôpitaux de Paris, Université Paris-Est Créteil, INSERM U955 | Christophe Rodriguez et al |
| EPI_ISL_4284206 | CNR Virus des Infections Respiratoires - France SUD | CNR Virus des Infections Respiratoires - France SUD | Antonin Bal et al |
| EPI_ISL_2923189 | Labo Analyses Med | National Reference Center for Viruses of Respiratory Infections, Institut Pasteur, Paris | Marion Barbet et al |
| EPI_ISL_6963002 | Mirialis | CNR Virus des Infections Respiratoires - France SUD | Antonin Bal et al |
| EPI_ISL_3155013 | Labo Analyses Med | National Reference Center for Viruses of Respiratory Infections, Institut Pasteur, Paris | Marion Barbet et al |
| EPI_ISL_4411406 | Labo Analyses Med | National Reference Center for Viruses of Respiratory Infections, Institut Pasteur, Paris | Marion Barbet et al |
| EPI_ISL_4323957 | MIRIALIS CLUSES BECHET | CNR Virus des Infections Respiratoires - France SUD | Antonin Bal et al |
| EPI_ISL_2179638 | Hôpital Ambroise-Paré | Department of Virology, Henri Mondor University Hospital, Assistance Publique Hôpitaux de Paris, Université Paris-Est Créteil, INSERM U955 | Christophe Rodriguez et al |
| EPI_ISL_6606101 | LBM UNIBIO VALENTIN | CNR Virus des Infections Respiratoires - France SUD | Antonin Bal et al |
| EPI_ISL_6962948 | CERBALLIANCE PARIS ET IDF EST | CERBA HealthCare | Judith Zerah et al |
| EPI_ISL_3635540 | AX BIO OCEAN | CNR Virus des Infections Respiratoires - France SUD | Antonin Bal et al |
| EPI_ISL_1789107 | Labo Analyses Med | National Reference Center for Viruses of Respiratory Infections, Institut Pasteur, Paris | Marion Barbet et al |
| EPI_ISL_3472466 | LABORATOIRE BIOESTEREL | CNR Virus des Infections Respiratoires - France SUD | Antonin Bal et al |

| | | | |
|---|---|---|---|
| EPI_ISL_1583079 | Labo analyses Med | National Reference Center for Viruses of Respiratory Infections, Institut Pasteur, Paris | Marion Barbet et al |
| EPI_ISL_1297257 | BIOMNIS PARIS | CNR Virus des Infections Respiratoires - France SUD | Antonin Bal et al |
| EPI_ISL_1915138 | Hospital | National Reference Center for Viruses of Respiratory Infections, Institut Pasteur, Paris | Marion Barbet et al |
| EPI_ISL_6844723 | LABORATOIRE LABAZUR | CNR Virus des Infections Respiratoires - France SUD | Antonin Bal et al |
| EPI_ISL_5692262 | BIOMNIS EUROFINS IVRY | Department of Virology, Henri Mondor University Hospital, Assistance Publique Hôpitaux de Paris, Université Paris-Est Créteil, INSERM U955 | Christophe Rodriguez et al |
| EPI_ISL_5750248 | CHU DE DIJON | Department of Virology, Henri Mondor University Hospital, Assistance Publique Hôpitaux de Paris, Université Paris-Est Créteil, INSERM U955 | Christophe Rodriguez et al |
| EPI_ISL_6967758 | CERBALLIANCE PARIS ET IDF EST | CERBA HealthCare | Judith Zerah et al |
| EPI_ISL_1788714 | CNR Virus des Infections Respiratoires - France SUD | CNR Virus des Infections Respiratoires - France SUD | Antonin Bal et al |
| EPI_ISL_2151691 | HOPITAL NORD FRANCHE COMTE | Department of Virology, Henri Mondor University Hospital, Assistance Publique Hôpitaux de Paris, Université Paris-Est Créteil, INSERM U955 | Christophe Rodriguez et al |
| EPI_ISL_1788856 | Labo Analyses med | National Reference Center for Viruses of Respiratory Infections, Institut Pasteur, Paris | Marion Barbet et al |
| EPI_ISL_3472874 | LABORATOIRE BIOESTEREL | CNR Virus des Infections Respiratoires - France SUD | Antonin Bal et al |
| EPI_ISL_3382477 | LABORATOIRE d'ANALYSES de BIOLOGIE MEDICALES PUILBOREAU | CNR Virus des Infections Respiratoires - France SUD | Antonin Bal et al |
| EPI_ISL_5593937 | Hôpital Cochin | Department of Virology, Henri Mondor University Hospital, Assistance Publique Hôpitaux de Paris, Université Paris-Est Créteil, INSERM U955 | Christophe Rodriguez et al |
| EPI_ISL_3478022 | BIOMNIS EUROFINS IVRY | Department of Virology, Henri Mondor University Hospital, Assistance Publique Hôpitaux de Paris, Université Paris-Est Créteil, INSERM U955 | Christophe Rodriguez et al |
| EPI_ISL_4283528 | CNR Virus des Infections Respiratoires - France SUD | CNR Virus des Infections Respiratoires - France SUD | Antonin Bal et al |

| | | | |
|---|---|---|---|
| EPI_ISL_2637790 | SELAS MEDILYS | Department of Virology, Henri Mondor University Hospital, Assistance Publique Hôpitaux de Paris, Université Paris-Est Créteil, INSERM U955 | Christophe Rodriguez et al |
| EPI_ISL_4319333 | LABORATOIRE UNILIANS DECINES | CNR Virus des Infections Respiratoires - France SUD | Antonin Bal et al |
| EPI_ISL_6471541 | Labo Analyses Med | National Reference Center for Viruses of Respiratory Infections, Institut Pasteur, Paris | Marion Barbet et al |
| EPI_ISL_4281189 | CNR Virus des Infections Respiratoires - France SUD | CNR Virus des Infections Respiratoires - France SUD | Antonin Bal et al |
| EPI_ISL_1359814 | BIOGROUP MOUANS SARTOUX | CNR Virus des Infections Respiratoires - France SUD | Antonin Bal et al |
| EPI_ISL_2595905 | MEPHI, Aix Marseille University | MEPHI, Aix Marseille University | Anthony LEVASSEUR et al |
| EPI_ISL_3472799 | LABORATOIRE FORTE BIO DAX | CNR Virus des Infections Respiratoires - France SUD | Antonin Bal et al |
| EPI_ISL_2444354 | CHU LILLE | CHU Lille - Laboratoire de Virologie | AIT YAHYA Emilie et al |
| EPI_ISL_6101765 | CHU Purpan - Laboratoire de Virologie - Institut Fédératif de Biologie | CHU Purpan - Laboratoire de Virologie - Institut Fédératif de Biologie | Latour J. et al |
| EPI_ISL_1404504 | Hôpital Henri Mondor | Department of Virology, Henri Mondor University Hospital, Assistance Publique Hôpitaux de Paris, Université Paris-Est Créteil, INSERM U955 | Christophe Rodriguez et al |
| EPI_ISL_5312090 | LABORATOIRE BIOESTEREL | CNR Virus des Infections Respiratoires - France SUD | Antonin Bal et al |
| EPI_ISL_1110148 | Groupe LCD | Department of Virology, Henri Mondor University Hospital, Assistance Publique Hôpitaux de Paris, Université Paris-Est Créteil, INSERM U955 | Christophe Rodriguez et al |
| EPI_ISL_1754357 | GH A.CHENEVIER-H.MONDOR | Department of Virology, Henri Mondor University Hospital, Assistance Publique Hôpitaux de Paris, Université Paris-Est Créteil, INSERM U955 | Christophe Rodriguez et al |
| EPI_ISL_2454086 | MEPHI, Aix Marseille University | MEPHI, Aix Marseille University | Anthony LEVASSEUR et al |
| EPI_ISL_2289481 | ASTRALAB TASSIGNY | CNR Virus des Infections Respiratoires - France SUD | Antonin Bal et al |
| EPI_ISL_4323708 | LABORATOIRE CERBALLIANCE PLT VILLON | CNR Virus des Infections Respiratoires - France SUD | Antonin Bal et al |
| EPI_ISL_6651492 | LABM KUATE | CERBA HealthCare | Bénédicte Roquebert et al |
| EPI_ISL_7138751 | Labo Analyses Med | National Reference Center for Viruses of Respiratory Infections, Institut Pasteur, Paris | Marion Barbet et al |
| EPI_ISL_1910928 | CHU Lille | CHU Lille | AIT YAHYA Emilie et al |
| EPI_ISL_5464662 | MIRIALIS CLUSES BECHET | CNR Virus des Infections Respiratoires - France SUD | Antonin Bal et al |

| | | | |
|---|---|---|---|
| EPI_ISL_1297294 | BIOMNIS PARIS | CNR Virus des Infections Respiratoires - France SUD | Antonin Bal et al |
| EPI_ISL_6261852 | CERBALLIANCE PARIS ET IDF EST VILLEPINTE | CERBA HealthCare | Bénédicte Roquebert et al |
| EPI_ISL_2381160 | CHU Purpan - Laboratoire de Virologie - Institut Fédératif de Biologie | CHU Purpan - Laboratoire de Virologie - Institut Fédératif de Biologie | Latour J. et al |
| EPI_ISL_645199 | CHU Nîmes | CNR Virus des Infections Respiratoires - France SUD | Antonin Bal et al |
| EPI_ISL_2374276 | Labo Analyses Med | National Reference Center for Viruses of Respiratory Infections, Institut Pasteur, Paris | Marion Barbet et al |
| EPI_ISL_3477063 | SELAS MEDILYS | Department of Virology, Henri Mondor University Hospital, Assistance Publique Hôpitaux de Paris, Université Paris-Est Créteil, INSERM U955 | Christophe Rodriguez et al |
| EPI_ISL_6076343 | AX BIO OCEAN | CNR Virus des Infections Respiratoires - France SUD | Antonin Bal et al |
| EPI_ISL_2151783 | SELAS MEDILYS | Department of Virology, Henri Mondor University Hospital, Assistance Publique Hôpitaux de Paris, Université Paris-Est Créteil, INSERM U955 | Christophe Rodriguez et al |
| EPI_ISL_671961 | CHU Purpan - Laboratoire de Virologie - Institut Fédératif de Biologie | CHU Purpan - Laboratoire de Virologie - Institut Fédératif de Biologie | Latour J. et al |
| EPI_ISL_6254466 | CH. FLERS Jacques MONOD | Department of Virology, Henri Mondor University Hospital, Assistance Publique Hôpitaux de Paris, Université Paris-Est Créteil, INSERM U955 | Christophe Rodriguez et al |
| EPI_ISL_2612664 | Labo Analyses Med | National Reference Center for Viruses of Respiratory Infections, Institut Pasteur, Paris | Marion Barbet et al |
| EPI_ISL_6254515 | BIOSMOSE IDF | Department of Virology, Henri Mondor University Hospital, Assistance Publique Hôpitaux de Paris, Université Paris-Est Créteil, INSERM U955 | Christophe Rodriguez et al |
| EPI_ISL_4805556 | LAM BIO 86 SITE DE CHAUMONT | CNR Virus des Infections Respiratoires - France SUD | Antonin Bal et al |
| EPI_ISL_3383207 | LBM SEALAB DARRASSE ET ASSOCIES | CNR Virus des Infections Respiratoires - France SUD | Antonin Bal et al |
| EPI_ISL_2361528 | CH CALAIS | CHU Lille - Laboratoire de Virologie | AIT YAHYA Emilie et al |
| EPI_ISL_700380 | Centre hospitalier Métropole Savoie | CNR Virus des Infections Respiratoires - France SUD | Antonin Bal et al |
| EPI_ISL_754167 | Laboratoire de Microbiologie | National Reference Center for Viruses of Respiratory Infections, Institut Pasteur, Paris | Marion Barbet et al |

| | | | |
|---|---|---|---|
| EPI_ISL_5626482 | GH A.CHENEVIER-H.MONDOR | Department of Virology, Henri Mondor University Hospital, Assistance Publique Hôpitaux de Paris, Université Paris-Est Créteil, INSERM U955 | Christophe Rodriguez et al |
| EPI_ISL_1517453 | Labo Analyses Med | National Reference Center for Viruses of Respiratory Infections, Institut Pasteur, Paris | Marion Barbet et al |
| EPI_ISL_4284360 | CNR Virus des Infections Respiratoires - France SUD | CNR Virus des Infections Respiratoires - France SUD | Antonin Bal et al |
| EPI_ISL_2596330 | MEPHI, Aix Marseille University | MEPHI, Aix Marseille University | Anthony LEVASSEUR et al |
| EPI_ISL_7015784 | CERBALLIANCE IDF SUD SITE DE LISSES | CERBA HealthCare | Bénédicte Roquebert et al |
| EPI_ISL_1096295 | Hopital | National Reference Center for Viruses of Respiratory Infections, Institut Pasteur, Paris | Marion Barbet et al |
| EPI_ISL_2659942 | MEPHI, Aix Marseille University | MEPHI, Aix Marseille University | Anthony LEVASSEUR et al |
| EPI_ISL_4323834 | LABORATOIRE CERBALLIANCE PLT VILLON | CNR Virus des Infections Respiratoires - France SUD | Antonin Bal et al |
| EPI_ISL_2535277 | Labo Analyses Med | National Reference Center for Viruses of Respiratory Infections, Institut Pasteur, Paris | Marion Barbet et al |
| EPI_ISL_2228886 | Hôpital Cochin | Department of Virology, Henri Mondor University Hospital, Assistance Publique Hôpitaux de Paris, Université Paris-Est Créteil, INSERM U955 | Christophe Rodriguez et al |
| EPI_ISL_5626885 | BIO VSM LAB | Department of Virology, Henri Mondor University Hospital, Assistance Publique Hôpitaux de Paris, Université Paris-Est Créteil, INSERM U955 | Christophe Rodriguez et al |
| EPI_ISL_5905494 | SYNLAB MVZ Leinfelden-Echterdingen | Robert Koch Institute | ? |
| EPI_ISL_1351083 | SYNLAB MVZ Ettlingen | Robert Koch Institute | ? |
| EPI_ISL_1434708 | SYNLAB MVZ Leinfelden-Echterdingen | Robert Koch Institute | ? |
| EPI_ISL_2036590 | MEPHI, Aix Marseille University | MEPHI, Aix Marseille University | Anthony LEVASSEUR et al |
| EPI_ISL_2124230 | SYNLAB MVZ Berlin | Robert Koch Institute | ? |
| EPI_ISL_1364920 | München Klinik GmbH | Bundeswehr Institute of Microbiology | Markus Antwerpen et al |
| EPI_ISL_6886596 | Max von Pettenkofer Institute, Virology, National Reference Center for Retroviruses, LMU Munich | Laboratory for Functional Genome Analysis; Dept. Genomics; Gene Center of the LMU Munich | Max Muenchhoff et al |
| EPI_ISL_1355574 | Universitätsmedizin Mannheim | Robert Koch Institute | ? |
| EPI_ISL_1354954 | Universitätsklinikum Heidelberg | Robert Koch Institute | ? |

| | | | |
|---|---|---|---|
| EPI_ISL_4038652 | MVZ Labor Dr. Limbach & Kollegen GbR | Robert Koch Institute | ? |
| EPI_ISL_2263422 | SYNLAB MVZ Leinfelden-Echterdingen | Robert Koch Institute | ? |
| EPI_ISL_6467309 | SYNLAB MVZ Leinfelden-Echterdingen | Robert Koch Institute | ? |
| EPI_ISL_2264344 | SYNLAB MVZ Leinfelden-Echterdingen | Robert Koch Institute | ? |
| EPI_ISL_2261773 | SYNLAB MVZ Leinfelden-Echterdingen | Robert Koch Institute | ? |
| EPI_ISL_1914386 | Department for Virology, Molecular Biology and Genome Research, R. G. Lugar Center for Public Health Research, National Center for Disease Control and Public Health (NCDC) of Georgia. | Department for Virology, Molecular Biology and Genome Research, R. G. Lugar Center for Public Health Research, National Center for Disease Control and Public Health (NCDC) of Georgia. | Meri Pantsulaia et al |
| EPI_ISL_1351786 | SYNLAB MVZ Leinfelden-Echterdingen | Robert Koch Institute | ? |
| EPI_ISL_5399058 | SYNLAB MVZ Leinfelden-Echterdingen | Robert Koch Institute | ? |
| EPI_ISL_4227274 | MDI Limbach Berlin GmbH; MVZ Labor Berlin | Robert Koch Institute | ? |
| EPI_ISL_561046 | MRCG at LSHTM Genomics lab | MRCG at LSHTM Genomics lab | Abdul Karim sesay et al |
| EPI_ISL_1432098 | SYNLAB MVZ Ettlingen | Robert Koch Institute | ? |
| EPI_ISL_7116918 | Robert Koch-Institut ZBS1 (Zentrum für biologische Gefahren und spezielle Pathogene hochpathogene Viren) | Robert Koch Institute | ? |
| EPI_ISL_4938061 | SYNLAB MVZ Leinfelden-Echterdingen | Robert Koch Institute | ? |
| EPI_ISL_1353960 | Bioscientia MVZ Labor Karlsruhe GmbH | Robert Koch Institute | ? |
| EPI_ISL_5051002 | Max von Pettenkofer Institute, Virology, National Reference Center for Retroviruses, LMU Munich | Laboratory for Functional Genome Analysis; Dept. Genomics; Gene Center of the LMU Munich | Max Muenchhoff et al |
| EPI_ISL_5947267 | Charité Universitätsmedizin Berlin, Institut für Virologie/Labor Berlin | Charité Universitätsmedizin Berlin, Institut für Virologie | Victor M Corman et al |
| EPI_ISL_5397900 | Limbach - MVZ Humangenetik Ulm | Robert Koch Institute | ? |
| EPI_ISL_6953463 | SYNLAB MVZ Leinfelden-Echterdingen | Robert Koch Institute | ? |
| EPI_ISL_1355613 | Universitätsmedizin Mannheim | Robert Koch Institute | ? |
| EPI_ISL_5121661 | Eurofins LifeCodexx GmbH | Robert Koch Institute | ? |
| EPI_ISL_4613578 | SYNLAB MVZ Leinfelden-Echterdingen | Robert Koch Institute | ? |
| EPI_ISL_1286206 | Eurofins LifeCodexx GmbH | Robert Koch Institute | ? |

| | | | |
|---|---|---|---|
| EPI_ISL_5947081 | Charité Universitätsmedizin Berlin, Institut für Virologie/Labor Berlin | Charité Universitätsmedizin Berlin, Institut für Virologie | Victor M Corman et al |
| EPI_ISL_4928524 | SYNLAB MVZ Leinfelden-Echterdingen | Robert Koch Institute | ? |
| EPI_ISL_4703106 | Centre de Recherches Médicales de Lambaréné | Centre de Recherches Médicales de Lambaréné | Gédéon Prince Manouana et al |
| EPI_ISL_1886116 | Labor Berlin Charité Vivantes GmbH / Institut für Virologie | Charité Universitätsmedizin Berlin, Institut für Virologie/Labor Berlin | Peter Menzel et al |
| EPI_ISL_1438550 | SYNLAB MVZ Leinfelden-Echterdingen | Robert Koch Institute | ? |
| EPI_ISL_6713300 | SYNLAB MVZ Leinfelden-Echterdingen | Robert Koch Institute | ? |
| EPI_ISL_6886595 | Max von Pettenkofer Institute, Virology, National Reference Center for Retroviruses, LMU Munich | Laboratory for Functional Genome Analysis; Dept. Genomics; Gene Center of the LMU Munich | Max Muenchhoff et al |
| EPI_ISL_1216093 | MRCG at LSHTM Genomics lab | MRCG at LSHTM Genomics lab | Abdul Karim sesay et al |
| EPI_ISL_6781508 | MAINE ANJOU LBM | Department of Virology, Henri Mondor University Hospital, Assistance Publique Hôpitaux de Paris, Université Paris-Est Créteil, INSERM U955 | Christophe Rodriguez et al |
| EPI_ISL_5460193 | Labor Berlin Charité Vivantes GmbH / Institut für Virologie | Charité Universitätsmedizin Berlin, Institut für Virologie/Labor Berlin | Peter Menzel et al |
| EPI_ISL_3044374 | Corona-Testzentrum ifp Institut für Produktqualität GmbH | Robert Koch Institute | ? |
| EPI_ISL_7112501 | MVZ Labor Dr. Limbach & Kollegen GbR | Robert Koch Institute | ? |
| EPI_ISL_2313683 | Bioscientia MVZ Labor Karlsruhe GmbH | Robert Koch Institute | ? |
| EPI_ISL_6700984 | MAINE ANJOU LBM | Department of Virology, Henri Mondor University Hospital, Assistance Publique Hôpitaux de Paris, Université Paris-Est Créteil, INSERM U955 | Christophe Rodriguez et al |
| EPI_ISL_5120705 | MDI Limbach Berlin GmbH; MVZ Labor Berlin | Robert Koch Institute | ? |
| EPI_ISL_1355764 | Universitätsmedizin Mannheim | Robert Koch Institute | ? |
| EPI_ISL_2387542 | SYNLAB MVZ Leinfelden-Echterdingen | Robert Koch Institute | ? |
| EPI_ISL_6953982 | Labor Blackholm MVZ | Robert Koch Institute | ? |
| EPI_ISL_2635808 | MDI Limbach Berlin GmbH; MVZ Labor Berlin | Robert Koch Institute | ? |
| EPI_ISL_569217 | MEPHI, Aix Marseille University | MEPHI, Aix Marseille University | Anthony LEVASSEUR et al |
| EPI_ISL_3712700 | SYNLAB MVZ Leinfelden-Echterdingen | Robert Koch Institute | ? |
| EPI_ISL_5122804 | SYNLAB MVZ Leinfelden-Echterdingen | Robert Koch Institute | ? |

| | | | |
|---|---|---|---|
| EPI_ISL_6886594 | Max von Pettenkofer Institute, Virology, National Reference Center for Retroviruses, LMU Munich | Laboratory for Functional Genome Analysis; Dept. Genomics; Gene Center of the LMU Munich | Max Muenchhoff et al |
| EPI_ISL_4439761 | Limbach - MVZ Humangenetik Ulm | Robert Koch Institute | ? |
| EPI_ISL_1567019 | SYNLAB MVZ Berlin | Robert Koch Institute | ? |
| EPI_ISL_2633308 | SYNLAB MVZ Heidelberg | Robert Koch Institute | ? |
| EPI_ISL_1213817 | SYNLAB MVZ Ettlingen | Robert Koch Institute | ? |
| EPI_ISL_4035752 | Institute of Virology, Medical Center, University of Freiburg, Freiburg, Germany | Institute of Virology, Clinial Virus Genomics, Medical Center, University of Freiburg, Freiburg, Germany | Jonas Fuchs et al |
| EPI_ISL_3433329 | MEPHI, Aix Marseille University | MEPHI, Aix Marseille University | Anthony LEVASSEUR et al |
| EPI_ISL_900102 | MEPHI, Aix Marseille University | MEPHI, Aix Marseille University | Anthony LEVASSEUR et al |
| EPI_ISL_1148337 | Sonic - Dr. Staber & Kollegen GmbH München | Robert Koch Institute | ? |
| EPI_ISL_7019047 | Charité Universitätsmedizin Berlin, Institute of Virology | Charité Universitätsmedizin Berlin, Institute of Virology | Victor M Corman et al |
| EPI_ISL_1432833 | Eurofins LifeCodexx GmbH | Robert Koch Institute | ? |
| EPI_ISL_6441976 | Klinikum Ernst von Bergmann gemeinnützige GmbH - stationärer Bereich | Robert Koch Institute | ? |
| EPI_ISL_2030016 | Hospital | National Reference Center for Viruses of Respiratory Infections, Institut Pasteur, Paris | Marion Barbet et al |
| EPI_ISL_6832737 | Institute for Medical Virology Frankfurt | Institute for Medical Virology Frankfurt | Toptan T. et al |
| EPI_ISL_6886593 | Max von Pettenkofer Institute, Virology, National Reference Center for Retroviruses, LMU Munich | Laboratory for Functional Genome Analysis; Dept. Genomics; Gene Center of the LMU Munich | Max Muenchhoff et al |
| EPI_ISL_5051012 | Max von Pettenkofer Institute, Virology, National Reference Center for Retroviruses, LMU Munich | Laboratory for Functional Genome Analysis; Dept. Genomics; Gene Center of the LMU Munich | Max Muenchhoff et al |
| EPI_ISL_6971091 | Laboratoire analyse med | LABORIZON CENTRE BIOGROUP | HOLSTEIN ANNE et al |
| EPI_ISL_3712128 | Bioscientia MVZ Labor Karlsruhe GmbH | Robert Koch Institute | ? |
| EPI_ISL_912744 | Hôpital Henri Mondor | Department of Virology, Henri Mondor University Hospital, Assistance Publique Hôpitaux de Paris, Université Paris-Est Créteil, INSERM U955 | Christophe Rodriguez et al |
| EPI_ISL_5460149 | Labor Berlin Charité Vivantes GmbH / Institut für Virologie | Charité Universitätsmedizin Berlin, Institut für Virologie/Labor Berlin | Peter Menzel et al |

| | | | |
|---|---|---|---|
| EPI_ISL_1152520 | Universitätsklinikum Bonn - Institut für Medizinische Mikrobiologie Immunologie und Parasitologie + Institut für Virologie | Robert Koch Institute | ? |
| EPI_ISL_2871525 | MVZ Labor Krone GbR | Center of Medical Microbiology, Virology, and Hospital Hygiene, University of Duesseldorf | Dennis Deschka et al |
| EPI_ISL_1214990 | SYNLAB MVZ Weiden | Robert Koch Institute | ? |
| EPI_ISL_1354243 | Sonic - MVZ Medizinisches Labor Bremen GmbH | Robert Koch Institute | ? |
| EPI_ISL_3495356 | SYNLAB MVZ Weiden | Robert Koch Institute | ? |
| EPI_ISL_1722882 | SYNLAB MVZ Leverkusen | Robert Koch Institute | ? |
| EPI_ISL_1845125 | SYNLAB MVZ Leverkusen | Robert Koch Institute | ? |
| EPI_ISL_2129730 | Bernhard-Nocht-Institut fÃ¼r Tropenmedizin | Robert Koch Institute | ? |
| EPI_ISL_1641624 | SYNLAB MVZ Leverkusen | Robert Koch Institute | ? |
| EPI_ISL_7086885 | LabKom - Labor Hannover MVZ GmbH | Robert Koch Institute | ? |
| EPI_ISL_2126823 | SYNLAB MVZ Leverkusen | Robert Koch Institute | ? |
| EPI_ISL_1721300 | Sonic - MVZ Medizinisches Labor Bremen GmbH | Robert Koch Institute | ? |
| EPI_ISL_1726474 | SYNLAB MVZ Leverkusen | Robert Koch Institute | ? |
| EPI_ISL_6191139 | amedes Göttingen | Robert Koch Institute | ? |
| EPI_ISL_4458348 | ZOTZ KLIMAS MVZ Düsseldorf-Centrum GbR ÜBAG für Labormedizin, Genetik, Zytologie, Pathologie | Center of Medical Microbiology, Virology, and Hospital Hygiene, University of Duesseldorf | Maximilian Damagnez et al |
| EPI_ISL_3495428 | SYNLAB MVZ Weiden | Robert Koch Institute | ? |
| EPI_ISL_1715870 | MEDILYS Laborgesellschaft mbH | Heinrich Pette Institute, Leibniz Institute for Experimental Virology | Alexis Robitaille et al |
| EPI_ISL_1845674 | SYNLAB MVZ Leverkusen | Robert Koch Institute | ? |
| EPI_ISL_4443324 | Synlab MVZ Augsburg | Robert Koch Institute | ? |
| EPI_ISL_1215618 | Limbach - MVZ Labor Eveld & Kollegen Essen | Robert Koch Institute | ? |
| EPI_ISL_4438998 | Klinikum Fulda gAG | Robert Koch Institute | ? |
| EPI_ISL_5763360 | ZOTZ KLIMAS MVZ Düsseldorf-Centrum GbR ÜBAG für Labormedizin, Genetik, Zytologie, Pathologie | Center of Medical Microbiology, Virology, and Hospital Hygiene, University of Duesseldorf | Maximilian Damagnez et al |
| EPI_ISL_3715474 | SYNLAB Labor München Zentrum LMZ | Robert Koch Institute | ? |
| EPI_ISL_2380932 | ZOTZ KLIMAS MVZ Düsseldorf-Centrum GbR ÜBAG für Labormedizin, Genetik, Zytologie, Pathologie | Center of Medical Microbiology, Virology, and Hospital Hygiene, University of Duesseldorf | Maximilian Damagnez et al |
| EPI_ISL_6197100 | Sonic - Labor Dr. von Froreich GmbH | Robert Koch Institute | ? |
| EPI_ISL_4939246 | Labor Dr. Fenner und Kollegen | Heinrich Pette Institute, Leibniz Institute for Experimental Virology | Alexis Robitaille et al |
| EPI_ISL_4043027 | Synlab MVZ Augsburg | Robert Koch Institute | ? |
| EPI_ISL_1284398 | SYNLAB MVZ Weiden | Robert Koch Institute | ? |

| | | | |
|---|---|---|---|
| EPI_ISL_2130112 | SYNLAB MVZ Leverkusen | Robert Koch Institute | ? |
| EPI_ISL_4615258 | Laborarztpraxis Dres. med. Walther Weindel & Kollegen | Robert Koch Institute | ? |
| EPI_ISL_770761 | ZOTZ KLIMAS MVZ Düsseldorf-Centrum GbR ÜBAG für Labormedizin, Genetik, Zytologie, Pathologie | Center of Medical Microbiology, Virology, and Hospital Hygiene, University of Duesseldorf | Maximilian Damagnez et al |
| EPI_ISL_1155783 | SYNLAB MVZ Weiden | Robert Koch Institute | ? |
| EPI_ISL_1721074 | IMD Labor Frankfurt | Robert Koch Institute | ? |
| EPI_ISL_4044880 | SYNLAB MVZ Hamburg | Robert Koch Institute | ? |
| EPI_ISL_2125349 | Sonic - MVZ Medizinisches Labor Oldenburg | Robert Koch Institute | ? |
| EPI_ISL_4041394 | CENTOGENE Frankfurt Laboratory: Niederlassung Industriepark Höchst | Robert Koch Institute | ? |
| EPI_ISL_3121684 | Sonic - Dr. Staber & Kollegen GmbH München | Robert Koch Institute | ? |
| EPI_ISL_2871415 | MVZ Labor Krone GbR | Center of Medical Microbiology, Virology, and Hospital Hygiene, University of Duesseldorf | Dennis Deschka et al |
| EPI_ISL_1988670 | Center of Medical Microbiology, Virology, and Hospital Hygiene, University of Duesseldorf | Center of Medical Microbiology, Virology, and Hospital Hygiene, University of Duesseldorf | Maximilian Damagnez et al |
| EPI_ISL_7089715 | IMD - MVZ Labor Martinsried | Robert Koch Institute | ? |
| EPI_ISL_4441973 | LabKom - Labor Augsburg MVZ GmbH | Robert Koch Institute | ? |
| EPI_ISL_2117134 | SYNLAB MVZ Leverkusen | Robert Koch Institute | ? |
| EPI_ISL_7074576 | Niedersächsisches Landesgesundheitsamt (NLGA) | Robert Koch Institute | ? |
| EPI_ISL_1464497 | Labor Dr. Fenner und Kollegen | Heinrich Pette Institute, Leibniz Institute for Experimental Virology | Alexis Robitaille et al |
| EPI_ISL_1846724 | SYNLAB MVZ Leverkusen | Robert Koch Institute | ? |
| EPI_ISL_6952626 | amedes MVZ für Laboratoriumsdiagnostik Raubling GmbH | Robert Koch Institute | ? |
| EPI_ISL_1347297 | Medical Laboratories Duesseldorf | Center of Medical Microbiology, Virology, and Hospital Hygiene, University of Duesseldorf | Maximilian Damagnez et al |
| EPI_ISL_1147948 | Universitätsklinikum Leipzig - Institut für Medizinische Mikrobiologie und Virologie virologisches Labor | Robert Koch Institute | ? |
| EPI_ISL_1280208 | Limbach - MVZ Labor Eveld & Kollegen Essen | Robert Koch Institute | ? |
| EPI_ISL_1285629 | LabKom - MVZ Labor Bochum MLB GmbH | Robert Koch Institute | ? |
| EPI_ISL_776527 | University Medical Center Hamburg Eppendorf | Heinrich Pette Institute, Leibniz Institute for Experimental Virology | Alexis Robitaille et al |
| EPI_ISL_6196481 | LabKom - Labor Augsburg MVZ GmbH | Robert Koch Institute | ? |
| EPI_ISL_1729858 | SYNLAB MVZ Leverkusen | Robert Koch Institute | ? |

| | | | |
|---|---|---|---|
| EPI_ISL_1991906 | Medical Laboratories Duesseldorf | Center of Medical Microbiology, Virology, and Hospital Hygiene, University of Duesseldorf | Maximilian Damagnez et al |
| EPI_ISL_1213010 | amedes MVZ Hannover | Robert Koch Institute | ? |
| EPI_ISL_4225743 | amedes Göttingen | Robert Koch Institute | ? |
| EPI_ISL_2267862 | SYNLAB MVZ Leverkusen | Robert Koch Institute | ? |
| EPI_ISL_1433466 | Labor Becker & Kollegen (Standort München) | Robert Koch Institute | ? |
| EPI_ISL_2381451 | ZOTZ KLIMAS MVZ Düsseldorf-Centrum GbR ÜBAG für Labormedizin, Genetik, Zytologie, Pathologie | Center of Medical Microbiology, Virology, and Hospital Hygiene, University of Duesseldorf | Maximilian Damagnez et al |
| EPI_ISL_4933417 | CENTOGENE Frankfurt Laboratory: Niederlassung Industriepark Höchst | Robert Koch Institute | ? |
| EPI_ISL_6959872 | Institute for Medical Virology, Frankfurt | Institute for Medical Virology, Frankfurt | Toptan T. et al |
| EPI_ISL_4933445 | CENTOGENE Frankfurt Laboratory: Niederlassung Industriepark Höchst | Robert Koch Institute | ? |
| EPI_ISL_1729874 | SYNLAB MVZ Leverkusen | Robert Koch Institute | ? |
| EPI_ISL_5683038 | Bayerisches Landesamt für Gesundheit und Lebensmittelsicherheit (LGL) | Robert Koch Institute | ? |
| EPI_ISL_3121683 | LabKom - Labor Augsburg MVZ GmbH | Robert Koch Institute | ? |
| EPI_ISL_1723833 | SYNLAB MVZ Leverkusen | Robert Koch Institute | ? |
| EPI_ISL_1726855 | UniversitÃ¤tsklinikum MÃ¼nster Institut fÃ¼r Virologie | Robert Koch Institute | ? |
| EPI_ISL_1845584 | SYNLAB MVZ Leverkusen | Robert Koch Institute | ? |
| EPI_ISL_7078334 | IMD - MVZ Labor Greifswald | Robert Koch Institute | ? |
| EPI_ISL_1354145 | Sonic - MVZ Medizinisches Labor Bremen GmbH | Robert Koch Institute | ? |
| EPI_ISL_3947870 | Labor Dr. Fenner und Kollegen | Heinrich Pette Institute, Leibniz Institute for Experimental Virology | Alexis Robitaille et al |
| EPI_ISL_6959873 | Institute for Medical Virology, Frankfurt | Institute for Medical Virology, Frankfurt | Toptan T. et al |
| EPI_ISL_4040684 | SYNLAB MVZ Dachau | Robert Koch Institute | ? |
| EPI_ISL_1727048 | Medizinische Laboratorien DÃ¼sseldorf | Robert Koch Institute | ? |
| EPI_ISL_3044731 | SYNLAB MVZ Weiden | Robert Koch Institute | ? |
| EPI_ISL_2116386 | SYNLAB MVZ Leverkusen | Robert Koch Institute | ? |
| EPI_ISL_776109 | University Medical Center Hamburg Eppendorf | Heinrich Pette Institute, Leibniz Institute for Experimental Virology | Alexis Robitaille et al |
| EPI_ISL_1725780 | SYNLAB MVZ Leverkusen | Robert Koch Institute | ? |
| EPI_ISL_2126489 | SYNLAB MVZ Leverkusen | Robert Koch Institute | ? |
| EPI_ISL_4441610 | LabKom - Labor Hannover MVZ GmbH | Robert Koch Institute | ? |
| EPI_ISL_2384898 | Labor Dr. Fenner und Kollegen | Heinrich Pette Institute, Leibniz Institute for Experimental Virology | Alexis Robitaille et al |
| EPI_ISL_1151590 | SYNLAB MVZ Weiden | Robert Koch Institute | ? |

| | | | |
|---|---|---|---|
| EPI_ISL_1354337 | IMD - MVZ Labor Martinsried | Robert Koch Institute | ? |
| EPI_ISL_5904090 | LabKom - Labor Augsburg MVZ GmbH | Robert Koch Institute | ? |
| EPI_ISL_1565496 | IMD - MVZ Labor Martinsried | Robert Koch Institute | ? |
| EPI_ISL_5912761 | Bayerisches Landesamt für Gesundheit und Lebensmittelsicherheit (LGL) | Robert Koch Institute | ? |
| EPI_ISL_5121529 | Bayerisches Landesamt für Gesundheit und Lebensmittelsicherheit (LGL) | Robert Koch Institute | ? |
| EPI_ISL_2113953 | SYNLAB MVZ Leverkusen | Robert Koch Institute | ? |
| EPI_ISL_3239001 | SYNLAB MVZ Weiden | Robert Koch Institute | ? |
| EPI_ISL_5401168 | Labor Becker & Kollegen (Standort München) | Robert Koch Institute | ? |
| EPI_ISL_2108629 | Centogene; Dr. Bauer Laboratoriums GmbH | Robert Koch Institute | ? |
| EPI_ISL_4610561 | SYNLAB MVZ Weiden | Robert Koch Institute | ? |
| EPI_ISL_1441935 | SYNLAB MVZ Leverkusen | Robert Koch Institute | ? |
| EPI_ISL_4227897 | LabKom - Labor Hannover MVZ GmbH | Robert Koch Institute | ? |
| EPI_ISL_1845625 | SYNLAB MVZ Leverkusen | Robert Koch Institute | ? |
| EPI_ISL_3716223 | MVZ Labor Dr. Fenner und Kollegen (Standort Hamburg) | Robert Koch Institute | ? |
| EPI_ISL_2387111 | CENTOGENE Frankfurt Laboratory: Niederlassung Industriepark Höchst | Robert Koch Institute | ? |
| EPI_ISL_4223607 | SYNLAB MVZ Weiden | Robert Koch Institute | ? |
| EPI_ISL_3948064 | Labor Dr. Fenner und Kollegen | Heinrich Pette Institute, Leibniz Institute for Experimental Virology | Alexis Robitaille et al |
| EPI_ISL_2127058 | SYNLAB MVZ Leverkusen | Robert Koch Institute | ? |
| EPI_ISL_1146035 | Sonic - MVZ Medizinisches Labor Bremen GmbH | Robert Koch Institute | ? |
| EPI_ISL_2259856 | CENTOGENE Frankfurt Laboratory: Niederlassung Industriepark Höchst | Robert Koch Institute | ? |
| EPI_ISL_2390093 | Limbach - MVZ Labor Eveld & Kollegen Essen | Robert Koch Institute | ? |
| EPI_ISL_6441893 | LADR Zentrallabor DR. Kramer & Kollegen Geesthacht | Robert Koch Institute | ? |
| EPI_ISL_2266215 | amedes MVZ Halle/Leipzig | Robert Koch Institute | ? |
| EPI_ISL_1356050 | SYNLAB Jena Oncoscreen | Robert Koch Institute | ? |
| EPI_ISL_6444467 | SYNLAB MVZ Leverkusen | Robert Koch Institute | ? |
| EPI_ISL_4937601 | Uniklinikum Carl Gustav Carus an der TU Dresden; Institut für Virologie | Robert Koch Institute | ? |
| EPI_ISL_7097968 | SYNLAB MVZ Leverkusen | Robert Koch Institute | ? |
| EPI_ISL_6715673 | SYNLAB MVZ Leverkusen | Robert Koch Institute | ? |
| EPI_ISL_6957233 | Labor Mönchengladbach MVZ Dr. Stein + Kollegen GbR | Robert Koch Institute | ? |
| EPI_ISL_4042999 | SYNLAB MVZ Leverkusen | Robert Koch Institute | ? |
| EPI_ISL_3497439 | MVZ Labor Krone GbR | Robert Koch Institute | ? |

| | | | |
|---|---|---|---|
| EPI_ISL_3891504 | Eurofins MVZ Labor Gelsenkirchen | Robert Koch Institute | ? |
| EPI_ISL_7098319 | SYNLAB MVZ Leverkusen | Robert Koch Institute | ? |
| EPI_ISL_6186586 | LADR Zentrallabor DR. Kramer & Kollegen Geesthacht | Robert Koch Institute | ? |
| EPI_ISL_4444870 | Labor Dr. Wisplinghoff - Köln | Robert Koch Institute | ? |
| EPI_ISL_1141548 | Bioscientia Labor Wermsdorf | Robert Koch Institute | ? |
| EPI_ISL_1350003 | Bioscientia Labor Wermsdorf | Robert Koch Institute | ? |
| EPI_ISL_6197701 | Labor Lübeck bzw. Laborärztliche Gemeinschaftspraxis Lübeck | Robert Koch Institute | ? |
| EPI_ISL_7103128 | Sonic - Labor Staber Dresden (Klipphausen) | Robert Koch Institute | ? |
| EPI_ISL_6452720 | amedes MVZ Jena | Robert Koch Institute | ? |
| EPI_ISL_3712423 | SYNLAB MVZ Leverkusen | Robert Koch Institute | ? |
| EPI_ISL_3497769 | SYNLAB MVZ Leverkusen | Robert Koch Institute | ? |
| EPI_ISL_4443663 | SYNLAB Jena Oncoscreen | Robert Koch Institute | ? |
| EPI_ISL_2117166 | SYNLAB MVZ Trier | Robert Koch Institute | ? |
| EPI_ISL_7106244 | LADR Zentrallabor DR. Kramer & Kollegen Geesthacht | Robert Koch Institute | ? |
| EPI_ISL_2121644 | Bioscientia Labor Wermsdorf | Robert Koch Institute | ? |
| EPI_ISL_1642816 | Department fÃ¼r Labormedizin Abteilung III Bereich Molekulare Diagnostik UniversitÃ¤tsklinikum Halle(Saale) | Robert Koch Institute | ? |
| EPI_ISL_5675132 | LADR Zentrallabor DR. Kramer & Kollegen Geesthacht | Robert Koch Institute | ? |
| EPI_ISL_1366942 | Virology, Universitätsklinikum des Saarlandes | Epigenetics, Saarland University | Kathrin Kattler et al |
| EPI_ISL_4444675 | Labor Dr. Wisplinghoff - Köln | Robert Koch Institute | ? |
| EPI_ISL_1436336 | LabKom - Labor an der Salzbrücke MVZ GmbH | Robert Koch Institute | ? |
| EPI_ISL_3893449 | LADR Zentrallabor DR. Kramer & Kollegen Geesthacht | Robert Koch Institute | ? |
| EPI_ISL_7068304 | SYNLAB MVZ Leverkusen | Robert Koch Institute | ? |
| EPI_ISL_2119944 | SYNLAB MVZ Neustadt a. d. Weinstraße | Robert Koch Institute | ? |
| EPI_ISL_4438878 | SYNLAB MVZ Leverkusen | Robert Koch Institute | ? |
| EPI_ISL_1729425 | SYNLAB MVZ Trier | Robert Koch Institute | ? |
| EPI_ISL_6461346 | Bioscientia Labor Wermsdorf | Robert Koch Institute | ? |
| EPI_ISL_1436712 | amedes Medizinisches Labor Mühlhausen | Robert Koch Institute | ? |
| EPI_ISL_6954142 | MVZ Labor Dr. Quade & Kollegen GmbH | Robert Koch Institute | ? |
| EPI_ISL_3716130 | Labor Mönchengladbach MVZ Dr. Stein + Kollegen GbR | Robert Koch Institute | ? |
| EPI_ISL_1091160 | Virology, Universitätsklinikum des Saarlandes | Epigenetics, Saarland University | Kathrin Kattler et al |
| EPI_ISL_2121012 | Bioscientia Labor Wermsdorf | Robert Koch Institute | ? |
| EPI_ISL_2262052 | SYNLAB Jena Oncoscreen | Robert Koch Institute | ? |
| EPI_ISL_4928480 | SYNLAB MVZ Leverkusen | Robert Koch Institute | ? |

| | | | |
|---|---|---|---|
| EPI_ISL_1154950 | Bioscientia Labor Wermsdorf | Robert Koch Institute | ? |
| EPI_ISL_1352884 | SYNLAB MVZ Trier | Robert Koch Institute | ? |
| EPI_ISL_3239468 | SYNLAB MVZ Leverkusen | Robert Koch Institute | ? |
| EPI_ISL_707994 | Virology, Universitätsklinikum des Saarlandes | Epigenetics, Saarland University | Kathrin Kattler et al |
| EPI_ISL_4231407 | Labor Dr. Wisplinghoff - Köln | Robert Koch Institute | ? |
| EPI_ISL_4610976 | LADR Zentrallabor DR. Kramer & Kollegen Geesthacht | Robert Koch Institute | ? |
| EPI_ISL_3890991 | LADR Zentrallabor DR. Kramer & Kollegen Geesthacht | Robert Koch Institute | ? |
| EPI_ISL_6192417 | SYNLAB MVZ Leverkusen | Robert Koch Institute | ? |
| EPI_ISL_6959529 | LABOR CHEMNITZ – Zentrum für Diagnostik am Klinikum Chemnitz | Robert Koch Institute | ? |
| EPI_ISL_5904644 | LADR Zentrallabor DR. Kramer & Kollegen Geesthacht | Robert Koch Institute | ? |
| EPI_ISL_6957171 | Eurofins MVZ Labor Gelsenkirchen | Robert Koch Institute | ? |
| EPI_ISL_4227411 | MVZ für Laboratoriumsmedizin und Mikrobiologie Koblenz-Mittelrhein (Labor Koblenz) | Robert Koch Institute | ? |
| EPI_ISL_6200289 | SYNLAB MVZ Leverkusen | Robert Koch Institute | ? |
| EPI_ISL_2469761 | Med. Labor Prof. Schenk Dr. Ansorge & Kollegen | Robert Koch Institute | ? |
| EPI_ISL_1843688 | Bioscientia Labor Wermsdorf | Robert Koch Institute | ? |
| EPI_ISL_4229511 | Thüringer Landesamtes für Verbraucherschutz | Robert Koch Institute | ? |
| EPI_ISL_1212961 | amedes MVZ Jena | Robert Koch Institute | ? |
| EPI_ISL_3494937 | amedes MVZ DIAMEDIS Sennestadt | Robert Koch Institute | ? |
| EPI_ISL_6952719 | MVZ für Laboratoriumsmedizin und Mikrobiologie Koblenz-Mittelrhein (Labor Koblenz) | Robert Koch Institute | ? |
| EPI_ISL_7103814 | Bioscientia Labor Wermsdorf | Robert Koch Institute | ? |
| EPI_ISL_960939 | Landesamt für Verbraucherschutz Sachsen Anhalt, Magdeburg | Institute of Medical Microbiology and Hospital Hygiene | Prof. Dr. Achim Kaasch et al |
| EPI_ISL_7117396 | Institut für Labormedizin Mikrobiologie und Hygiene | Robert Koch Institute | ? |
| EPI_ISL_4039300 | Labor Mönchengladbach MVZ Dr. Stein + Kollegen GbR | Robert Koch Institute | ? |
| EPI_ISL_1438440 | SYNLAB Jena Oncoscreen | Robert Koch Institute | ? |
| EPI_ISL_6444716 | SYNLAB MVZ Leverkusen | Robert Koch Institute | ? |
| EPI_ISL_7099670 | SYNLAB MVZ Leverkusen | Robert Koch Institute | ? |
| EPI_ISL_4043005 | SYNLAB MVZ Leverkusen | Robert Koch Institute | ? |
| EPI_ISL_3713472 | Medizinische Laboratorien Düsseldorf | Robert Koch Institute | ? |
| EPI_ISL_7115937 | Bioscientia Labor Wermsdorf | Robert Koch Institute | ? |

| | | | |
|---|---|---|---|
| EPI_ISL_1147535 | Uniklinikum Carl Gustav Carus an der TU Dresden; Institut für Virologie | Robert Koch Institute | ? |
| EPI_ISL_3087707 | Landesamt für Verbraucherschutz Sachsen Anhalt, Magdeburg | Institute of Medical Microbiology and Hospital Hygiene | Prof. Dr. Achim Kaasch et al |
| EPI_ISL_5121928 | MVZ Labor Krone GbR | Robert Koch Institute | ? |
| EPI_ISL_3500331 | Eurofins MVZ Labor Gelsenkirchen | Robert Koch Institute | ? |
| EPI_ISL_1141286 | Bioscientia Labor Wermsdorf | Robert Koch Institute | ? |
| EPI_ISL_3713645 | amedes MVZ DIAMEDIS Sennestadt | Robert Koch Institute | ? |
| EPI_ISL_4446391 | SYNLAB MVZ Leverkusen | Robert Koch Institute | ? |
| EPI_ISL_5909996 | labopart - Medizinische Laboratorien Dresden | Robert Koch Institute | ? |
| EPI_ISL_3237502 | SYNLAB MVZ Leverkusen | Robert Koch Institute | ? |
| EPI_ISL_4444247 | Labor Mönchengladbach MVZ Dr. Stein + Kollegen GbR | Robert Koch Institute | ? |
| EPI_ISL_1573114 | SYNLAB MVZ Trier | Robert Koch Institute | ? |
| EPI_ISL_1848884 | Diagnosticum - Labor Neukirchen | Robert Koch Institute | ? |
| EPI_ISL_2388666 | Sonic - Bioscientia - MVZ Labor Saar GmbH | Robert Koch Institute | ? |
| EPI_ISL_3713354 | MVZ Dr. Eberhard & Partner Dortmund | Robert Koch Institute | ? |
| EPI_ISL_6952041 | alphaomega Labor GbR (Medizinische Kooperationsgemeinschaft Dres. Ackermann/Tennert/Zaiß) | Robert Koch Institute | ? |
| EPI_ISL_5402950 | MVZ für Laboratoriumsmedizin und Mikrobiologie Koblenz-Mittelrhein (Labor Koblenz) | Robert Koch Institute | ? |
| EPI_ISL_2263780 | SYNLAB Jena Oncoscreen | Robert Koch Institute | ? |
| EPI_ISL_3238960 | SYNLAB MVZ Leverkusen | Robert Koch Institute | ? |
| EPI_ISL_4438584 | SYNLAB MVZ Leverkusen | Robert Koch Institute | ? |
| EPI_ISL_3120971 | SYNLAB MVZ Leverkusen | Robert Koch Institute | ? |
| EPI_ISL_4045505 | LADR Zentrallabor DR. Kramer & Kollegen Geesthacht | Robert Koch Institute | ? |
| EPI_ISL_4445366 | MVZ für Laboratoriumsmedizin und Mikrobiologie Koblenz-Mittelrhein (Labor Koblenz) | Robert Koch Institute | ? |
| EPI_ISL_2879116 | INSACOG Surveillance | INSACOG at CSIR Institute of Genomics and Integrative Biology | INSACOG et al |
| EPI_ISL_6963509 | Sri Jayadeva Institute of Cardiovascular Sciences and Research / Strand Life Sciences | National Centre for Biological Sciences, TIFR - Rockefeller Foundation | Rohan Pais et al |
| EPI_ISL_6698790 | HOME QUARANTINE TASKFORCE | Hong Kong Department of Health | Alan K.L. Tsang et al |

| | | | |
|---|---|---|---|
| EPI_ISL_3066116 | Department of Microbiology | Department of Microbiology | Zigui Chen et al |
| EPI_ISL_2379419 | DSU CHAMRAJANAGARA | INSACOG-KA, NIMHANS | Chitra Pattabiraman et al |
| EPI_ISL_2370547 | Greek Genome Center, Biomedical Research Foundation of the Academy of Athens (BRFAA) | Greek Genome Center, Biomedical Research Foundation of the Academy of Athens (BRFAA) | Emmanouil Athanasiadis et al |
| EPI_ISL_6832108 | Home Quarantine Taskforce | Hong Kong Department of Health | Alan K.L. Tsang et al |
| EPI_ISL_6716902 | Hong Kong Department of Health | School of Public Health, The University of Hong Kong | Dominic N.C. Tsang et al |
| EPI_ISL_1639913 | SYNLAB MVZ Ettlingen | Robert Koch Institute | ? |
| EPI_ISL_1573473 | SYNLAB MVZ Leinfelden-Echterdingen | Robert Koch Institute | ? |
| EPI_ISL_1723066 | SYNLAB MVZ Leinfelden-Echterdingen | Robert Koch Institute | ? |
| EPI_ISL_6939058 | National Influenza Centre | National Influenza Centre | William K. Ampofo et al |
| EPI_ISL_3268037 | WACCBIP, University of Ghana, Accra, Ghana | WACCBIP, University of Ghana, Volta Road, Legon-Accra, Ghana | Collins M. Morang'a et al |
| EPI_ISL_6841980 | Department of Microbiology, The University of Hong Kong | Department of Microbiology, The University of Hong Kong | Kelvin K.W. To et al |
| EPI_ISL_4578439 | Hassan Institute of Medical Sciences | INSACOG-KA, NIMHANS | Anson Kunjumon George et al |
| EPI_ISL_1034710 | Department of Microbiology, The University of Hong Kong | Department of Microbiology, The University of Hong Kong | Kelvin K.W. To et al |
| EPI_ISL_1497354 | National Public Health Organization | National Public Health Organization | Kyriaki Tryfinopoulou et al |
| EPI_ISL_4470116 | National Centre for Disease Control (NCDC) Biotechnology Division, Delhi - INSACOG | NCDC Delhi, Biotechnology Division - INSACOG | Radhakrishnan V. S et al |
| EPI_ISL_6958280 | North Lantau Hospital | Hong Kong Department of Health | Alan K.L. Tsang et al |
| EPI_ISL_2123506 | SYNLAB MVZ Leinfelden-Echterdingen | Robert Koch Institute | ? |
| EPI_ISL_4435627 | National Centre for Disease Control (NCDC) Biotechnology Division, Delhi - INSACOG | NCDC Delhi, Biotechnology Division - INSACOG | Radhakrishnan V. S et al |
| EPI_ISL_3162078 | Government Medical College Surat | Gujarat Biotechnology Research Centre | Zarna Patel et al |
| EPI_ISL_1725977 | SYNLAB MVZ Leinfelden-Echterdingen | Robert Koch Institute | ? |
| EPI_ISL_2126396 | Landesgesundheitsamt Baden-Wurttemberg | Robert Koch Institute | ? |
| EPI_ISL_2880755 | INSACOG Surveillance | INSACOG at CSIR Institute of Genomics and Integrative Biology | INSACOG et al |
| EPI_ISL_760052 | Hong Kong Department of Health | School of Public Health, The University of Hong Kong | Daniel Chu et al |
| EPI_ISL_2365202 | Hellenic National Blood Transfusion Center - EKEA | Greek Genome Center, Biomedical Research Foundation of the Academy of Athens (BRFAA) | Emmanouil Athanasiadis et al |
| EPI_ISL_2389638 | B.J. Medical College and Civil Hospital, Ahmedabad | Gujarat Biotechnology Research Centre | Zarna Patel et al |
| EPI_ISL_5063858 | AP SSO | CSIR-Centre for Cellular and Molecular Biology-INSACOG | Priya Nurkurthy et al |
| EPI_ISL_6590782 | Home Quarantine Taskforce | Hong Kong Department of Health | Alan K.L. Tsang et al |

| | | | |
|---|---|---|---|
| EPI_ISL_6689955 | Kasturba Hospital Molecular Lab - INSACOG | Kasturba hospital WGS | Jayanthi Shastri et al |
| EPI_ISL_4444941 | National Centre for Disease Control (NCDC) Biotechnology Division, Delhi - INSACOG | NCDC Delhi, Biotechnology Division - INSACOG | Robin Marwal et al |
| EPI_ISL_5659357 | National Influenza Centre | National Influenza Centre | William K. Ampofo et al |
| EPI_ISL_2879535 | INSACOG Surveillance | INSACOG at CSIR Institute of Genomics and Integrative Biology | INSACOG et al |
| EPI_ISL_3086995 | BJ Medical College Ahmedabad | Gujarat Biotechnology Research Centre | Nitin Savaliya et al |
| EPI_ISL_1287285 | Alexianer DaKS GmbH (eigenes IT-Tochterunternehmen der Alexianer Krankenhäuser) | Robert Koch Institute | ? |
| EPI_ISL_5063898 | AP SSO | CSIR-Centre for Cellular and Molecular Biology-INSACOG | Jandhyala Sai Krishna et al |
| EPI_ISL_4749444 | Kerala State SARS-COV-2 Genome Surveillance Programme (GENESCOV2) | GENESCOV2 at CSIR Institute of Genomics and Integrative Biology | Kerala State SARS-COV-2 Genome Surveillance Programme (GENESCOV2) et al |
| EPI_ISL_6691924 | Kasturba Hospital Molecular Lab - INSACOG | Kasturba hospital WGS | Jayanthi Shastri et al |
| EPI_ISL_6690947 | Kasturba Hospital Molecular Lab - INSACOG | Kasturba hospital WGS | Jayanthi Shastri et al |
| EPI_ISL_424458 | The National University Hospital of Iceland | deCODE genetics | Daniel F Gudbjartsson et al |
| EPI_ISL_6575195 | Genepath, Pune | RF-IISER Pune | Krishanpal Karmodiya et al |
| EPI_ISL_2368759 | Greek Genome Center, Biomedical Research Foundation of the Academy of Athens (BRFAA) | Greek Genome Center, Biomedical Research Foundation of the Academy of Athens (BRFAA) | Emmanouil Athanasiadis et al |
| EPI_ISL_1572711 | SYNLAB Jena Oncoscreen | Robert Koch Institute | ? |
| EPI_ISL_862473 | Kurnool Medical College (KMC) | CSIR Institute of Genomics and Integrative Biology | Pallavali Roja Rani et al |
| EPI_ISL_3006906 | Chhattisgarh Institute of Medical Sciences (CIMS), Bilaspur | Institute of Life Sciences - INSACOG | Sunil K. Raghav et al |
| EPI_ISL_6575132 | Genepath, Pune | RF-IISER Pune | Krishanpal Karmodiya et al |
| EPI_ISL_1838411 | CSIR-Centre for Cellular and Molecular Biology | CSIR-Centre for Cellular and Molecular Biology-INSACOG | Payel Mukherjee et al |
| EPI_ISL_3929213 | The National University Hospital of Iceland | deCODE genetics | Daniel F Gudbjartsson et al |
| EPI_ISL_6841981 | Department of Microbiology, The University of Hong Kong | Department of Microbiology, The University of Hong Kong | Kelvin K.W. To et al |
| EPI_ISL_3935285 | AP SSO | CSIR-Centre for Cellular and Molecular Biology - INSACOG | Sumedha Avadhanula et al |
| EPI_ISL_3928628 | The National University Hospital of Iceland | deCODE genetics | Daniel F Gudbjartsson et al |
| EPI_ISL_3689499 | AP SSO | CSIR-Centre for Cellular and Molecular Biology - INSACOG | Sofia Banu et al |
| EPI_ISL_1586730 | The National University Hospital of Iceland | deCODE genetics | Daniel F Gudbjartsson et al |
| EPI_ISL_6939048 | National Influenza Centre | National Influenza Centre | William K. Ampofo et al |

| | | | |
|---|---|---|---|
| EPI_ISL_6938638 | GMERS Medical College, Valsad | Gujarat Biotechnology Research Centre | Apurvasinh Puvar et al |
| EPI_ISL_2461288 | National Centre for Disease Control (NCDC) Biotechnology Division, Delhi | NCDC Delhi, Biotechnology Division INSACOG | Manoj K Singh et al |
| EPI_ISL_3927996 | The National University Hospital of Iceland | deCODE genetics | Daniel F Gudbjartsson et al |
| EPI_ISL_417752 | The National University Hospital of Iceland | deCODE genetics | Daniel F Gudbjartsson et al |
| EPI_ISL_1573511 | SYNLAB MVZ Leinfelden-Echterdingen | Robert Koch Institute | ? |
| EPI_ISL_6144267 | Genepath, Pune | RF-IISER Pune | Krishanpal Karmodiya et al |
| EPI_ISL_6963510 | Auriga Research Pvt.Ltd / Strand Life Sciences | National Centre for Biological Sciences, TIFR - Rockefeller Foundation | Rohan Pais et al |
| EPI_ISL_3060802 | AP SSO | CSIR-Centre for Cellular and Molecular Biology-INSACOG | Lamuk Zaveri et al |
| EPI_ISL_1567399 | SYNLAB Jena Oncoscreen | Robert Koch Institute | ? |
| EPI_ISL_2880885 | INSACOG Surveillance | INSACOG at CSIR Institute of Genomics and Integrative Biology | INSACOG et al |
| EPI_ISL_2370725 | Greek Genome Center, Biomedical Research Foundation of the Academy of Athens (BRFAA) | Greek Genome Center, Biomedical Research Foundation of the Academy of Athens (BRFAA) | Emmanouil Athanasiadis et al |
| EPI_ISL_1720788 | MVZ Labor Dr. Limbach & Kollegen GbR | Robert Koch Institute | ? |
| EPI_ISL_2364958 | Greek Genome Center, Biomedical Research Foundation of the Academy of Athens (BRFAA) | Greek Genome Center, Biomedical Research Foundation of the Academy of Athens (BRFAA) | Emmanouil Athanasiadis et al |
| EPI_ISL_1645979 | SYNLAB MVZ Heidelberg | Robert Koch Institute | ? |
| EPI_ISL_6939060 | National Influenza Centre | National Influenza Centre | William K. Ampofo et al |
| EPI_ISL_2774832 | AP SSO | CSIR-Centre for Cellular and Molecular Biology-INSACOG | Vidhyadhari Methuku et al |
| EPI_ISL_3928725 | The National University Hospital of Iceland | deCODE genetics | Daniel F Gudbjartsson et al |
| EPI_ISL_2128120 | SYNLAB MVZ Heidelberg | Robert Koch Institute | ? |
| EPI_ISL_1851859 | SYNLAB MVZ Leinfelden-Echterdingen | Robert Koch Institute | ? |
| EPI_ISL_1850885 | SYNLAB Jena Oncoscreen | Robert Koch Institute | ? |
| EPI_ISL_4444770 | National Centre for Disease Control (NCDC) Biotechnology Division, Delhi - INSACOG | NCDC Delhi, Biotechnology Division - INSACOG | Robin Marwal et al |
| EPI_ISL_1642512 | LabKom - Labor an der SalzbrÃ¼cke MVZ GmbH | Robert Koch Institute | ? |
| EPI_ISL_3689822 | AP SSO | CSIR-Centre for Cellular and Molecular Biology - INSACOG | Tulasi Nagabandi et al |
| EPI_ISL_2502254 | National Centre for Disease Control (NCDC) Biotechnology Division, Delhi | NCDC Delhi, Biotechnology Division INSACOG | Uma Sharma et al |

| | | | |
|---|---|---|---|
| EPI_ISL_6877982 | Department of Microbiology, Papanikolaou General Hospital of Thessaloniki | Institute of Applied Biosciences, Centre for Research and Technology Hellas | Anastasia Chatzidimitriou et al. |
| EPI_ISL_6716890 | Hong Kong Department of Health | School of Public Health, The University of Hong Kong | Dominic N.C. Tsang et al |
| EPI_ISL_1570154 | Eurofins LifeCodexx GmbH | Robert Koch Institute | ? |
| EPI_ISL_2370773 | Greek Genome Center, Biomedical Research Foundation of the Academy of Athens (BRFAA) | Greek Genome Center, Biomedical Research Foundation of the Academy of Athens (BRFAA) | Emmanouil Athanasiadis et al |
| EPI_ISL_1586527 | The National University Hospital of Iceland | deCODE genetics | Daniel F Gudbjartsson et al |
| EPI_ISL_2461796 | National Centre for Disease Control (NCDC) Biotechnology Division, Delhi | NCDC Delhi, Biotechnology Division INSACOG | Meena Datta et al |
| EPI_ISL_5602611 | Israel Central Virology laboratory | Israel Central Virology laboratory | Neta Zuckerman et al |
| EPI_ISL_5210768 | National Virus Reference Laboratory | National Virus Reference Laboratory | Zoe Yandle et al |
| EPI_ISL_2502118 | National Centre for Disease Control (NCDC) Biotechnology Division, Delhi | NCDC Delhi, Biotechnology Division INSACOG | Uma Sharma et al |
| EPI_ISL_1132426 | National Virus Reference Laboratory | National Virus Reference Laboratory | Zoe Yandle et al |
| EPI_ISL_1399028 | National Virus Reference Laboratory | National Virus Reference Laboratory | Zoe Yandle et al |
| EPI_ISL_2533660 | National Virus Reference Laboratory | National Virus Reference Laboratory | Zoe Yandle et al |
| EPI_ISL_6101153 | Dr S Raju, Director of Public Health and Preventive Medicine | inStem NCBS – INSACOG | Uma Ramakrishnan et al |
| EPI_ISL_4102880 | Ispat General Hospital, Rourkela | Institute of Life Sciences-INSACOG | Sunil K. Raghav et al |
| EPI_ISL_2879804 | INSACOG Surveillance | INSACOG at CSIR Institute of Genomics and Integrative Biology | INSACOG et al |
| EPI_ISL_4460918 | National Centre for Disease Control (NCDC) Biotechnology Division, Delhi - INSACOG | NCDC Delhi, Biotechnology Division - INSACOG | Kalaiarasan Ponnusamy et al |
| EPI_ISL_6886253 | INSACOG-WB | National Institute of Biomedical Genomics – INSACOG | Arindam Maitra et al |
| EPI_ISL_6939901 | National Virus Reference Laboratory | National Virus Reference Laboratory | Zoe Yandle et al |
| EPI_ISL_5431086 | National Virus Reference Laboratory | National Virus Reference Laboratory | Zoe Yandle et al |
| EPI_ISL_5729810 | Israel Central Virology laboratory | Israel Central Virology laboratory | Neta Zuckerman et al |
| EPI_ISL_1589669 | INSACOG-WB | National Institute of Biomedical Genomics – INSACOG | Arindam Maitra et al |
| EPI_ISL_5946350 | Dr S Raju, Director of Public Health and Preventive Medicine | inStem NCBS – INSACOG | Uma Ramakrishnan et al |

| | | | |
|---|---|---|---|
| EPI_ISL_2862096 | National Virus Reference Laboratory | National Virus Reference Laboratory | Zoe Yandle et al |
| EPI_ISL_3507434 | Gandhi Medical College | Centre for DNA Fingerprinting and Diagnostics | Murali Bashyam et al |
| EPI_ISL_4102892 | Covid Laboratory, Jharsuguda | Institute of Life Sciences-INSACOG | Sunil K. Raghav et al |
| EPI_ISL_2881707 | INSACOG Surveillance | INSACOG at CSIR Institute of Genomics and Integrative Biology | INSACOG et al |
| EPI_ISL_4031705 | National Virus Reference Laboratory | National Virus Reference Laboratory | Zoe Yandle et al |
| EPI_ISL_944473 | Israel Central Virology laboratory | Israel National Consortium for SARS-CoV-2 sequencing | Neta Zuckerman et al |
| EPI_ISL_5823078 | Israel Central Virology laboratory | Israel Central Virology laboratory | Neta Zuckerman et al |
| EPI_ISL_4779816 | National Virus Reference Laboratory | National Virus Reference Laboratory | Zoe Yandle et al |
| EPI_ISL_1502308 | National Virus Reference Laboratory | National Virus Reference Laboratory | Zoe Yandle et al |
| EPI_ISL_3591064 | National Virus Reference Laboratory | National Virus Reference Laboratory | Zoe Yandle et al |
| EPI_ISL_4446981 | National Centre for Disease Control (NCDC) Biotechnology Division, Delhi - INSACOG | NCDC Delhi, Biotechnology Division - INSACOG | Mahesh S Dhar et al |
| EPI_ISL_3375073 | National Virus Reference Laboratory | National Virus Reference Laboratory | Zoe Yandle et al |
| EPI_ISL_2984498 | National Virus Reference Laboratory | National Virus Reference Laboratory | Guerrino Macori et al |
| EPI_ISL_2085261 | Israel Central Virology laboratory | Israel National Consortium for SARS-CoV-2 sequencing | Neta Zuckerman et al |
| EPI_ISL_1577501 | National Virus Reference Laboratory | National Virus Reference Laboratory | Zoe Yandle et al |
| EPI_ISL_4091745 | inDNA Life Sciences, Bhubaneswar | Institute of Life Sciences - INSACOG | Sunil K. Raghav et al |
| EPI_ISL_5620258 | Israel Central Virology laboratory | Israel Central Virology laboratory | Neta Zuckerman et al |
| EPI_ISL_3118046 | National Virus Reference Laboratory | National Virus Reference Laboratory | Zoe Yandle et al |
| EPI_ISL_4460593 | National Virus Reference Laboratory | National Virus Reference Laboratory | Zoe Yandle et al |
| EPI_ISL_2085805 | Israel Central Virology laboratory | Israel National Consortium for SARS-CoV-2 sequencing | Neta Zuckerman et al |
| EPI_ISL_3242293 | Government Fever Hospital | CDFD | Murali Bashyam et al |
| EPI_ISL_2533355 | National Virus Reference Laboratory | National Virus Reference Laboratory | Zoe Yandle et al |
| EPI_ISL_1299420 | National Virus Reference Laboratory | National Virus Reference Laboratory | Zoe Yandle et al |
| EPI_ISL_3016038 | IISER Berhampur | Institute of Life Sciences - INSACOG | Sunil K. Raghav et al |
| EPI_ISL_4091385 | FM Medical College, Balasore | Institute of Life Sciences - INSACOG | Sunil K. Raghav et al |
| EPI_ISL_500577 | National Virus Reference Laboratory | National Virus Reference Laboratory | Michael Carr et al |
| EPI_ISL_1552655 | National Virus Reference Laboratory | National Virus Reference Laboratory | Zoe Yandle et al |

| | | | |
|---|---|---|---|
| EPI_ISL_4792184 | National Virus Reference Laboratory | National Virus Reference Laboratory | Zoe Yandle et al |
| EPI_ISL_2504480 | National Centre for Disease Control (NCDC) Biotechnology Division, Delhi | NCDC Delhi, Biotechnology Division INSACOG | Mahesh S Dhar et al |
| EPI_ISL_4057140 | Banaras Hindu University | CDFD | Murali Bashyam et al |
| EPI_ISL_2968185 | National Virus Reference Laboratory | National Virus Reference Laboratory | Zoe Yandle et al |
| EPI_ISL_2158414 | National Virus Reference Laboratory | National Virus Reference Laboratory | Zoe Yandle et al |
| EPI_ISL_2504698 | National Centre for Disease Control (NCDC) Biotechnology Division, Delhi | NCDC Delhi, Biotechnology Division INSACOG | Manoj K Singh et al |
| EPI_ISL_3011240 | National Virus Reference Laboratory | National Virus Reference Laboratory | Zoe Yandle et al |
| EPI_ISL_6939819 | National Virus Reference Laboratory | National Virus Reference Laboratory | Zoe Yandle et al |
| EPI_ISL_1073512 | Israel Central Virology laboratory | Israel National Consortium for SARS-CoV-2 sequencing | Neta Zuckerman et al |
| EPI_ISL_6394295 | INSACOG-Sikkim | National Institute of Biomedical Genomics – INSACOG | Arindam Maitra et al |
| EPI_ISL_2548739 | INSACOG-WB | National Institute of Biomedical Genomics – INSACOG | Arindam Maitra et al |
| EPI_ISL_4572420 | CSIR-Central Leather Research Institute | CSIR-Centre for Cellular and Molecular Biology - INSACOG | Jandhyala Sai Krishna et al |
| EPI_ISL_2676297 | INSACOG-WB | National Institute of Biomedical Genomics – INSACOG | Arindam Maitra et al |
| EPI_ISL_5022756 | RSUP Sanglah Bali | National Institute of Health Research and Development | Triyani Soekarso et al |
| EPI_ISL_4471604 | National Virus Reference Laboratory | National Virus Reference Laboratory | Zoe Yandle et al |
| EPI_ISL_5760270 | INSACOG-Mizoram | National Institute of Biomedical Genomics – INSACOG | Arindam Maitra et al |
| EPI_ISL_3253244 | National Virus Reference Laboratory | National Virus Reference Laboratory | Zoe Yandle et al |
| EPI_ISL_2454939 | National Virus Reference Laboratory | National Virus Reference Laboratory | Zoe Yandle et al |
| EPI_ISL_2504719 | National Centre for Disease Control (NCDC) Biotechnology Division, Delhi | NCDC Delhi, Biotechnology Division INSACOG | Manoj K Singh et al |
| EPI_ISL_2101123 | Molecular Biology Laboratory, Faculty Medicine and Health Sciences, Warmadewa University | Eijkman Institute for Molecular Biology, National Agency for Research and Innovation; Molecular Biology Laboratory, Faculty Medicine and Health Sciences, Warmadewa University | Lydia V. Panggalo et al |
| EPI_ISL_5886479 | CBR, IISc | inStem NCBS – INSACOG | Uma Ramakrishnan et al |
| EPI_ISL_4126711 | INSACOG Surveillance | INSACOG at CSIR Institute of Genomics and Integrative Biology | INSACOG et al |
| EPI_ISL_6846969 | RSUD Arifin Achmad | National Institute of Health Research and Development | Vivi Setiawaty et al |

| | | | |
|---|---|---|---|
| EPI_ISL_2881675 | INSACOG Surveillance | INSACOG at CSIR Institute of Genomics and Integrative Biology | INSACOG et al |
| EPI_ISL_7055396 | CSIR-Centre for Cellular and Molecular Biology | CSIR-Centre for Cellular and Molecular Biology | Amareshwar Vodapalli et al |
| EPI_ISL_1357918 | National Virus Reference Laboratory | National Virus Reference Laboratory | Zoe Yandle et al |
| EPI_ISL_3550502 | Dinas Kesehatan Provinsi Kalimantan Timur | Eijkman Institute for Molecular Biology, National Research and Innovation Agency; National Institute of Health Research and Development | Willy Agustine et al |
| EPI_ISL_6885717 | INSACOG-WB | National Institute of Biomedical Genomics – INSACOG | Arindam Maitra et al |
| EPI_ISL_6277621 | National Virus Reference Laboratory | National Virus Reference Laboratory | Zoe Yandle et al |
| EPI_ISL_2183585 | Israel Central Virology laboratory | Israel National Consortium for SARS-CoV-2 sequencing | Neta Zuckerman et al |
| EPI_ISL_3088211 | National Virus Reference Laboratory | National Virus Reference Laboratory | Zoe Yandle et al |
| EPI_ISL_6032237 | Institute of Preventive Medicine | Centre for DNA Fingerprinting and Diagnostics | Murali Bashyam et al |
| EPI_ISL_1785446 | National Virus Reference Laboratory | National Virus Reference Laboratory | Zoe Yandle et al |
| EPI_ISL_3590817 | National Virus Reference Laboratory | National Virus Reference Laboratory | Zoe Yandle et al |
| EPI_ISL_6033345 | Dr S raju, Director of Public Health and Preventive Medicine | inStem NCBS – INSACOG | Uma Ramakrishnan et al |
| EPI_ISL_2342236 | District Headquarter Hospital, Puri | Institute of Life Sciences - INSACOG | Sunil K. Raghav et al |
| EPI_ISL_4004702 | QR Lab | National Research and Innovation Agency (BRIN-Indonesia) | Indriawati et al |
| EPI_ISL_1499793 | National Virus Reference Laboratory | National Virus Reference Laboratory | Zoe Yandle et al |
| EPI_ISL_2968183 | National Virus Reference Laboratory | National Virus Reference Laboratory | Zoe Yandle et al |
| EPI_ISL_1252276 | National Virus Reference Laboratory | National Virus Reference Laboratory | Guerrino Macori et al |
| EPI_ISL_5000651 | National Virus Reference Laboratory | National Virus Reference Laboratory | Zoe Yandle et al |
| EPI_ISL_2533229 | National Virus Reference Laboratory | National Virus Reference Laboratory | Zoe Yandle et al |

| | | | |
|---|---|---|---|
| EPI_ISL_2932610 | KOTA Yogya DKK Gondokusuman II | Genetics Working Group (Pokja Genetik) Faculty of Medicine, Public Health and Nursing Universitas Gadjah Mada (FK-KMK UGM); Disease Investigation Center Wates Ministry of Agriculture Indonesia; Department of Microbiology FK-KMK UGM; Laboratorium Diagnostik Yayasan Tahija World Mosquito Program (WMP) Yogyakarta Center for | Gunadi et al |
| EPI_ISL_1117734 | National Virus Reference Laboratory | National Virus Reference Laboratory | Zoe Yandle et al |
| EPI_ISL_1620730 | National Virus Reference Laboratory | Irish Coronavirus Sequencing Consortium - Teagasc Moorepark | Jose Sanchez-Morgado et al |
| EPI_ISL_2373593 | Banaras Hindu University | CSIR-Centre for Cellular and Molecular Biology-INSACOG | Royana Singh et al |
| EPI_ISL_5305010 | Dipartimento di Medicina di Laboratorio, Azienda sanitaria universitaria Friuli Centrale (ASU FC) | Dipartimento di Medicina di Laboratorio, Azienda sanitaria universitaria Friuli Centrale (ASU FC) | Corrado Pipan et al |
| EPI_ISL_3733994 | PO AVEZZANO - U.O.C. Laboratorio Analisi | Istituto Zooprofilattico Sperimentale dell’Abruzzo e Molise “G. Caporale” | Lorusso A et al |
| EPI_ISL_1927123 | SARS-CoV-2 testing team, National Institute of Infectious Diseases | Pathogen Genomics Center, National Institute of Infectious Diseases | Tsuyoshi Sekizuka et al |
| EPI_ISL_1287624 | Istituto Zooprofilattico Sperimentale del Mezzogiorno | TIGEM | Antonio Grimaldi Patrizia Annunziata Francesco Panariello Biancamaria Pierri Claudia Tiberio Valentina Bouche Chiara Colantuono Maria Concetta Cuomo Denise Di Concilio Lucio Di Filippo Anna Manfredi Marcello Salvi Antonio Limone Luigi Atripaldi Pellegrino Cerino Andrea Ballabio Davide Cacchiarelli et al |
| EPI_ISL_6860419 | Shamir Medical Center (Asaf Harofe) | Shamir Medical Center (Asaf Harofe) | Nir Rainy et al |
| EPI_ISL_2001309 | SIESP CHIETI - DRIVE IN LANCIANO | Istituto Zooprofilattico Sperimentale dell’Abruzzo e Molise “G. Caporale” | Lorusso A et al |
| EPI_ISL_3119801 | AOUIVR | Istituto Zooprofilattico Sperimentale delle Venezie | Adelaide Milani et al |
| EPI_ISL_1336909 | SIESP SULMONA | Istituto Zooprofilattico Sperimentale dell’Abruzzo e Molise “G. Caporale” | Lorusso A et al |

| | | | |
|---|---|---|---|
| EPI_ISL_6666228 | Azienda Ospedaliero - Universitaria di Modena Policlinico - Virologia e Microbiologia Molecolare | Istituto Zooprofilattico Sperimentale della Lombardia e dell'Emilia Romagna (IZSLER), Risk Analysis and Genomic Epidemiology Unit | Marina Morganti et al |
| EPI_ISL_4849237 | Laboratorio CQRC | CQRC_QUALITY CONTROL CHEMICAL BIOLOGICAL RISK_AOOR Villa Sofia Cervello Palermo | Di Gaudio F. et al |
| EPI_ISL_1167445 | ASL Napoli 1 Centro | AMES Centro Polidiagnostico Strumentale S.r.l. | "Giovanni Savarese et al |
| EPI_ISL_1632821 | Istituto Zooprofilattico Sperimentale del Mezzogiorno | Telethon Institute of Genetics and Medicine (TIGEM) | Antonio Grimaldi Patrizia Annunziata Francesco Panariello Biancamaria Pierri Claudia Tiberio Teresa Giuliano Valentina Bouche Chiara Colantuono Maria Concetta Cuomo Denise Di Concilio Lucio Di Filippo Anna Manfredi Marcello Salvi Antonio Limone Luigi Atripaldi Pellegrino Cerino Andrea Ballabio Davide Cacchiarelli et al |
| EPI_ISL_5879704 | Genzano | National Institute for Infectious Diseases (INMI) L. Spallanzani I.R.C.C.S | E Giombini et al |
| EPI_ISL_6205121 | Shamir Medical Center (Asaf Harofe) | Shamir Medical Center (Asaf Harofe) | Nir Rainy et al |
| EPI_ISL_882717 | 1.AO Universitaria 'S. Giovanni di Dio e Ruggi D'Aragona, Scuola Medica Salernitana' Hospital / 2.UOC di Virologia e Microbiologia, Università della Campania 'L. Vanvitelli' / 3.AO Universitaria 'Federico II' Napoli Hospital / 4.AORN 'San Giuseppe Moscati' Avellino Hospital / 5.AO 'San Pio - presidio G. Rummo' Benevento Hospital / 6.AO 'Sant'Anna e San Sebastiano' Caserta Hospital / 7.PO 'Maria Santissima Addolorata' Eboli Hospital / 8.Biogem Istituto di Ricerche Genetiche | 1. Genome Research Center for Health (CRGS) / 2. Laboratory of Molecular Medicine and Genomics(LMMGe) / 3. Center for Research in Pure and Applied Mathematics (CRMPA) | Giorgio Giurato et al |
| EPI_ISL_1557613 | ASL Napoli 1 Centro | AMES Centro Polidiagnostico Strumentale S.r.l. | "Giovanni Savarese et al |
| EPI_ISL_3253394 | A.S.L. NOVARA | Fondazione del Piemonte per l'Oncologia IRCCS | Antonino Sottile et al |

| | | | |
|---|---|---|---|
| EPI_ISL_935129 | 1.AO Universitaria 'S. Giovanni di Dio e Ruggi D'Aragona, Scuola Medica Salernitana' Hospital / 2.UOC di Virologia e Microbiologia, Università della Campania 'L. Vanvitelli' / 3.AO Universitaria 'Federico II' Napoli Hospital / 4.AORN 'San Giuseppe Moscati' Avellino Hospital / 5.AO 'San Pio - presidio G. Rummo' Benevento Hospital / 6.AO 'Sant'Anna e San Sebastiano' Caserta Hospital / 7.PO 'Maria Santissima Addolorata' Eboli Hospital / 8.Biogem Istituto di Ricerche Genetiche | 1. Genome Research Center for Health (CRGS) / 2. Laboratory of Molecular Medicine and Genomics(LMMGe) / 3. Center for Research in Pure and Applied Mathematics (CRMPA) | Giorgio Giurato (Corresponding Author) et al |
| EPI_ISL_7151452 | Lab. Microbiologia e Virologia Cotugno A.O. dei Colli | TIGEM | Antonio Grimaldi Patrizia Annunziata Francesco Panariello Claudia Tiberio Teresa Giuliano Valentina Bouche Chiara Colantuono Lucio Di Filippo Anna Manfredi Marcello Salvi Antonio Limone Luigi Atripaldi Andrea Ballabio Davide Cacchiarelli et al |
| EPI_ISL_3280880 | RM1/IRCCS IDI | National Institute for Infectious Diseases (INMI) L. Spallanzani I.R.C.C.S | E Giombini et al |
| EPI_ISL_6777160 | Policlinico San Donato | Laboratory of Clinical Microbiology, Virology and Bioemergencies, ASST Fatebenefratelli Sacco - Sacco Hospital | Valeria Micheli et al |
| EPI_ISL_6475142 | PO AVEZZANO - U.O.C. Laboratorio Analisi | Istituto Zooprofilattico Sperimentale dell’Abruzzo e Molise “G. Caporale” | Lorusso A et al |
| EPI_ISL_6854346 | Microbiologia e Virologia Cotugno | Microbiologia e Virologia Cotugno | Claudia Tiberio et al |
| EPI_ISL_2820749 | Ospedale Santissima Annunziata | Istituto Zooprofilattico Sperimentale della Puglia e della Basilicata | Parisi A. et al |
| EPI_ISL_6901618 | Shamir Medical Center (Asaf Harofe) | Shamir Medical Center (Asaf Harofe) | Nir Rainy et al |
| EPI_ISL_2084055 | Israel Central Virology laboratory | Israel National Consortium for SARS-CoV-2 sequencing | Neta Zuckerman et al |
| EPI_ISL_6781502 | Center of Advanced Studies and Technology, Molecular Genetics Laboratory | Center of Advanced Studies and Technology, Molecular Genetics Laboratory | Anaclerio Federico et al |

| | | | |
|---|---|---|---|
| EPI_ISL_2725439 | Istituto Zooprofilattico Sperimentale del Mezzogiorno | Telethon Institute of Genetics and Medicine (TIGEM) | Antonio Grimaldi Patrizia Annunziata Francesco Panariello Biancamaria Pierri Claudia Tiberio Teresa Giuliano Valentina Bouche Chiara Colantuono Maria Concetta Cuomo Denise Di Concilio Lucio Di Filippo Anna Manfredi Marcello Salvi Antonio Limone Luigi Atripaldi Pellegrino Cerino Andrea Ballabio Davide Cacchiarelli et al |
| EPI_ISL_3446800 | AULSS 3 Venezia | UOSD Genetica e Citogenetica - Azienda ULSS 3 Serenissima; Istituto Zooprofilattico Sperimentale delle Venezie | Mosé Favarato et al |
| EPI_ISL_1110368 | Azienda Sanitaria dell'Alto Adige Laboratorio Aziendale di Microbiologia e Virologia | Istituto di Genomica Applicata | Elisabetta Pagani et al |
| EPI_ISL_1048828 | Azienda Sanitaria dell’Alto Adige Laboratorio Aziendale di Microbiologia e Virologia | Istituto di Genomica Applicata | Elisabetta Pagani et al |
| EPI_ISL_1674095 | Israel Central Virology laboratory | Israel National Consortium for SARS-CoV-2 sequencing | Neta Zuckerman et al |
| EPI_ISL_1096753 | ASL Napoli 1 Centro | AMES Centro Polidiagnostico Strumentale S.r.l. | "Giovanni Savarese et al |
| EPI_ISL_1824070 | San Gallicano Dermatological Institute I.F.O. | INMI Lazzaro Spallanzani IRCCS | M Pallocca et al |
| EPI_ISL_3501754 | AULSS 3 Venezia | UOSD Genetica e Citogenetica - Azienda ULSS 3 Serenissima; Istituto Zooprofilattico Sperimentale delle Venezie | Mosé Favarato et al |
| EPI_ISL_2096784 | Hadassah Medical Center Clinical Virology Laboratory, Hadassah Ein Kerem | Hadassah Hebrew University Viral Sequencing Group, Hadassah Hebrew University Medical Center | Hadar Golan Berman et al |
| EPI_ISL_1424424 | Università Federico II - Dipartimento di scienze mediche traslazionali - Napoli | TIGEM | Antonio Grimaldi Patrizia Annunziata Francesco Panariello Michele Cennamo Valentina Bouche Chiara Colantuono Lucio Di Filippo Mariano Fiorenza Anna Manfredi Marcello Salvi Giuseppe Portella Andrea Ballabio Davide Cacchiarelli et al |
| EPI_ISL_2083470 | Israel Central Virology laboratory | Israel National Consortium for SARS-CoV-2 sequencing | Neta Zuckerman et al |
| EPI_ISL_6854348 | Microbiologia e Virologia Cotugno | Microbiologia e Virologia Cotugno | Claudia Tiberio et al |
| EPI_ISL_6854347 | Microbiologia e Virologia Cotugno | Microbiologia e Virologia Cotugno | Claudia Tiberio et al |

| | | | |
|---|---|---|---|
| EPI_ISL_944412 | Israel Central Virology laboratory | Israel National Consortium for SARS-CoV-2 sequencing | Neta Zuckerman et al |
| EPI_ISL_6795212 | Shamir Medical Center (Asaf Harofe) | Shamir Medical Center (Asaf Harofe) | Nir Rainy et al |
| EPI_ISL_3667251 | A.S.L. CUNEO 1 | Fondazione del Piemonte per l'Oncologia IRCCS | Antonino Sottile et al |
| EPI_ISL_3298668 | U.O. Microbiologia Laboratorio Unico Centro Servizi - AUSL della Romagna | U.O. Microbiologia, Laboratorio Unico Centro Servizi - AUSL della Romagna | Giorgio Dirani et al |
| EPI_ISL_3925471 | A.S.L. CN1 | Fondazione del Piemonte per l'Oncologia IRCCS | Antonino Sottile et al |
| EPI_ISL_6809092 | A.S.L. VERCELLI | Fondazione del Piemonte per l'Oncologia IRCCS | Antonino Sottile et al |
| EPI_ISL_6201738 | Laboratorio di Riferimento Regionale della Sicilia Occidentale per l'Emergenza COVID-19 | Laboratorio di Riferimento Regionale della Sicilia Occidentale per l’Emergenza COVID-19 | Fabio Tramuto et al |
| EPI_ISL_1855001 | Virology Laboratory, Scientific Department, Army Medical Center | Virology Laboratory, Scientific Department, Army Medical Center | Silvia Fillo et al |
| EPI_ISL_3014307 | Università Federico II - Dipartimento di scienze mediche traslazionali - Napoli | TIGEM | Antonio Grimaldi Patrizia Annunziata Francesco Panariello Teresa Giuliano Michele Cennamo Valentina Bouche Chiara Colantuono Lucio Di Filippo Mariano Fiorenza Anna Manfredi Marcello Salvi Giuseppe Portella Andrea Ballabio Davide Cacchiarelli et al |
| EPI_ISL_779676 | Pathogen Genomics Center, National Institute of Infectious Diseases | Pathogen Genomics Center, National Institute of Infectious Diseases | Tsuyoshi Sekizuka et al |
| EPI_ISL_1940662 | ASL Napoli 1 Centro | AMES Centro Polidiagnostico Strumentale S.r.l. | Giovanni Savarese et al |
| EPI_ISL_6200954 | Shamir Medical Center (Asaf Harofe) | Shamir Medical Center (Asaf Harofe) | Nir Rainy et al |
| EPI_ISL_1544107 | SOC Igiene ed Epidemiologia Clinica, ASUFC, Udine and SC (UCO) Igiene e Sanità Pubblica, ASUGI, Trieste | Istituto di Genomica Applicata | Pierlanfranco D'Agaro et al |
| EPI_ISL_4884804 | Laboratorio CQRC | CQRC_QUALITY CONTROL CHEMICAL BIOLOGICAL RISK_AOOR Villa Sofia Cervello Palermo | Di Gaudio F. et al |
| EPI_ISL_804119 | Israel Central Virology laboratory | Israel National Consortium for SARS-CoV-2 sequencing | Neta Zuckerman et al |
| EPI_ISL_1763158 | Israel Central Virology laboratory | Israel National Consortium for SARS-CoV-2 sequencing | Neta Zuckerman et al |
| EPI_ISL_2815364 | ASL BARI | University of Bari Biomedical Sciences and Human Oncology | Maria Chironna et al |
| EPI_ISL_3019691 | Shamir Medical Center (Asaf Harofe) | Shamir Medical Center (Asaf Harofe) | Nir Rainy et al |

| | | | |
|---|---|---|---|
| EPI_ISL_6640487 | Clinical Microbiology Laboratory, Tel Aviv Sourasky Medical Center | Clinical Microbiology Laboratory, Tel Aviv Sourasky Medical Center | Katya Levytskyi et al |
| EPI_ISL_4731601 | A.S.L. CUNEO 1 | Fondazione del Piemonte per l'Oncologia IRCCS | Antonino Sottile et al |
| EPI_ISL_6938961 | S.C. Laboratorio Analisi, ASL 3 Liguria | U.O. Igiene, Ospedale Policlinico San Martino | Bruzzone Bianca et al |
| EPI_ISL_1558358 | ASL Napoli 1 Centro | AMES Centro Polidiagnostico Strumentale S.r.l. | "Giovanni Savarese et al |
| EPI_ISL_1361607 | 1.AO Universitaria 'S. Giovanni di Dio e Ruggi D'Aragona, Scuola Medica Salernitana' Hospital / 2.UOC di Virologia e Microbiologia, Università della Campania 'L. Vanvitelli' / 3.AO Universitaria 'Federico II' Napoli Hospital / 4.AORN 'San Giuseppe Moscati' Avellino Hospital / 5.AO 'San Pio - presidio G. Rummo' Benevento Hospital / 6.AO 'Sant'Anna e San Sebastiano' Caserta Hospital / 7.PO 'Maria Santissima Addolorata' Eboli Hospital / 8.Biogem Istituto di Ricerche Genetiche / 9. U.O.C. di Genetica Medica e di Laboratorio A.O.R.N., Azienda Ospedaliera di Rilievo Nazionale Antonio Cardarelli, Napoli / 10. Centro di riferimento Oncologico della Basilicata (IRCCS-CROB), Rionero in Vulture (PZ) / 11. Presidio Ospedaliero di Agropoli, Agropoli (SA). | 1. Genome Research Center for Health (CRGS) / 2. Laboratory of Molecular Medicine and Genomics(LMMGe) / 3. Center for Research in Pure and Applied Mathematics (CRMPA) | Giorgio Giurato (Corresponding Author) et al |
| EPI_ISL_1762713 | Israel Central Virology laboratory | Israel National Consortium for SARS-CoV-2 sequencing | Neta Zuckerman et al |
| EPI_ISL_5406377 | Azienda Sanitaria dell'Alto Adige - Laboratorio Aziendale di Microbiologia e Virologia | Azienda Sanitaria dell'Alto Adige | Irene Bianconi et al |
| EPI_ISL_3249035 | ASST MONZA | ASST MONZA | Sergio Maria Ivano Malandrin et al |
| EPI_ISL_3717558 | Virology Laboratory, Scientific Department, Army Medical Center | Virology Laboratory, Scientific Department, Army Medical Center | Silvia Fillo et al |
| EPI_ISL_6859939 | Shamir Medical Center (Asaf Harofe) | Shamir Medical Center (Asaf Harofe) | Nir Rainy et al |
| EPI_ISL_6825365 | Shamir Medical Center (Asaf Harofe) | Shamir Medical Center (Asaf Harofe) | Nir Rainy et al |

| | | | |
|---|---|---|---|
| EPI_ISL_667746 | Pathogen Genomics Center, National Institute of Infectious Diseases | Pathogen Genomics Center, National Institute of Infectious Diseases | Tsuyoshi Sekizuka et al |
| EPI_ISL_2844482 | Istituto Zooprofilattico Sperimentale del Mezzogiorno | TIGEM | Antonio Grimaldi Patrizia Annunziata Francesco Panariello Biancamaria Pierri Claudia Tiberio Teresa Giuliano Valentina Bouche Chiara Colantuono Maria Concetta Cuomo Denise Di Concilio Lucio Di Filippo Anna Manfredi Marcello Salvi Antonio Limone Luigi Atripaldi Pellegrino Cerino Andrea Ballabio Davide Cacchiarelli et al |
| EPI_ISL_6206291 | Shamir Medical Center (Asaf Harofe) | Shamir Medical Center (Asaf Harofe) | Nir Rainy et al |
| EPI_ISL_2270854 | SIESP CHIETI - DRIVE IN VASTO | Istituto Zooprofilattico Sperimentale dell'Abruzzo e Molise "G. Caporale" | Lorusso A et al |
| EPI_ISL_4103982 | S.C. Laboratorio Analisi, ASL 3 Liguria | U.O. Igiene, Ospedale Policlinico San Martino | Bruzzone Bianca et al |
| EPI_ISL_4170797 | U.O. Microbiologia Laboratorio Unico Centro Servizi - AUSL della Romagna | U.O. Microbiologia, Laboratorio Unico Centro Servizi - AUSL della Romagna | Giorgio Dirani et al |
| EPI_ISL_3655439 | Genesis Institutes for Genetic Research (GIGR) | Genesis Institutes for Genetic Research (GIGR) | Mio Ogawa et al |
| EPI_ISL_1260948 | ULSS 03 Venezia | Istituto Zooprofilattico Sperimentale delle Venezie | Adelaide Milani et al |
| EPI_ISL_1647168 | DIP. PREV. AVEZZANO SERVIZIO DI IGIENE EPIDEMIOLOGIA E SANITA' PUBBLICA | Istituto Zooprofilattico Sperimentale dell'Abruzzo e Molise "G. Caporale" | Lorusso A et al |
| EPI_ISL_1260963 | ULSS 04 Veneto Orientale | Istituto Zooprofilattico Sperimentale delle Venezie | Adelaide Milani et al |
| EPI_ISL_6457129 | ASST MONZA | ASST MONZA | Sergio Maria Ivano Malandrin et al |
| EPI_ISL_751393 | IRCCS Sacro Cuore Don Calabria Hospital, Department of Infectious, Tropical Diseases & Microbiology | University of Verona, Department of Biotechnology | Antonio Mori et al |
| EPI_ISL_4055934 | The Caribbean Public Health Agency | Carrington Lab, Department of Preclinical Sciences, Faculty of Medical Sciences, The University of the West Indies, St Augustine Campus | Nikita S. D. Sahadeo et al |
| EPI_ISL_1669941 | Università degli Studi di Perugia | Istituto Zooprofilattico Sperimentale dell'Abruzzo e Molise "G. Caporale" | Mencacci A et al |

| | | | |
|---|---|---|---|
| EPI_ISL_1307960 | Lab. Microbiologia e Virologia Cotugno A.O. dei Colli | TIGEM | Antonio Grimaldi Patrizia Annunziata Francesco Panariello Biancamaria Pierri Claudia Tiberio Valentina Bouche Chiara Colantuono Maria Concetta Cuomo Denise Di Concilio Lucio Di Filippo Anna Manfredi Marcello Salvi Antonio Limone Luigi Atripaldi Pellegrino Cerino Andrea Ballabio Davide Cacchiarelli et al |
| EPI_ISL_7056614 | Shamir Medical Center (Asaf Harofe) | Shamir Medical Center (Asaf Harofe) | Nir Rainy et al |
| EPI_ISL_2978718 | A.S.L. NOVARA-BORGOMANERO | Fondazione del Piemonte per l'Oncologia IRCCS | Antonino Sottile et al |
| EPI_ISL_2604906 | AULSS 2 Marca Trevigiana | Istituto Zooprofilattico Sperimentale delle Venezie | Adelaide Milani et al |
| EPI_ISL_1759410 | Istituto Zooprofilattico Sperimentale del Mezzogiorno | TIGEM | Antonio Grimaldi Patrizia Annunziata Francesco Panariello Biancamaria Pierri Claudia Tiberio Teresa Giuliano Valentina Bouche Chiara Colantuono Maria Concetta Cuomo Denise Di Concilio Lucio Di Filippo Anna Manfredi Marcello Salvi Antonio Limone Luigi Atripaldi Pellegrino Cerino Andrea Ballabio Davide Cacchiarelli et al |
| EPI_ISL_2455812 | Istituto Zooprofilattico Sperimentale del Mezzogiorno | TIGEM | Antonio Grimaldi Patrizia Annunziata Francesco Panariello Biancamaria Pierri Claudia Tiberio Teresa Giuliano Valentina Bouche Chiara Colantuono Maria Concetta Cuomo Denise Di Concilio Lucio Di Filippo Anna Manfredi Marcello Salvi Antonio Limone Luigi Atripaldi Pellegrino Cerino Andrea Ballabio Davide Cacchiarelli et al |
| EPI_ISL_7152022 | Via G. Quagliariello 54 80131 Napoli Italy | TIGEM | Antonio Grimaldi Patrizia Annunziata Francesco Panariello Claudia Tiberio Teresa Giuliano Valentina Bouche Chiara Colantuono Lucio Di Filippo Anna Manfredi Marcello Salvi Antonio Limone Luigi Atripaldi Andrea Ballabio Davide Cacchiarelli et al |

| | | | |
|---|---|---|---|
| EPI_ISL_3840533 | AZIENDA OSP. ORDINE MAURIZIANO DI TORINO | Fondazione del Piemonte per l'Oncologia IRCCS | Antonino Sottile et al |
| EPI_ISL_2084345 | Israel Central Virology laboratory | Israel National Consortium for SARS-CoV-2 sequencing | Neta Zuckerman et al |
| EPI_ISL_3376657 | A.S.L. CUNEO 1 | Fondazione del Piemonte per l'Oncologia IRCCS | Antonino Sottile et al |
| EPI_ISL_1229612 | ASL Napoli 1 Centro | AMES Centro Polidiagnostico Strumentale S.r.l. | "Giovanni Savarese et al |
| EPI_ISL_2820958 | UOC Microbiologia e Virologia, Azienda Ospedaliera Universitaria Senese, Siena, Italy | Dipartimento di Biotecnologie Mediche | Maria Grazia Cusi et al |
| EPI_ISL_1080731 | ASL Napoli 1 Centro | AMES Centro Polidiagnostico Strumentale S.r.l. | "Giovanni Savarese et al |
| EPI_ISL_7056045 | Shamir Medical Center (Asaf Harofe) | Shamir Medical Center (Asaf Harofe) | Nir Rainy et al |
| EPI_ISL_1762751 | Israel Central Virology laboratory | Israel National Consortium for SARS-CoV-2 sequencing | Neta Zuckerman et al |
| EPI_ISL_1714981 | Istituto Zooprofilattico Sperimentale del Mezzogiorno | TIGEM | Antonio Grimaldi Patrizia Annunziata Francesco Panariello Biancamaria Pierri Claudia Tiberio Teresa Giuliano Valentina Bouche Chiara Colantuono Maria Concetta Cuomo Denise Di Concilio Lucio Di Filippo Anna Manfredi Marcello Salvi Antonio Limone Luigi Atripaldi Pellegrino Cerino Andrea Ballabio Davide Cacchiarelli et al |
| EPI_ISL_1793155 | Istituto Zooprofilattico Sperimentale del Mezzogiorno | TIGEM | Antonio Grimaldi Patrizia Annunziata Francesco Panariello Biancamaria Pierri Claudia Tiberio Teresa Giuliano Valentina Bouche Chiara Colantuono Maria Concetta Cuomo Denise Di Concilio Lucio Di Filippo Anna Manfredi Marcello Salvi Antonio Limone Luigi Atripaldi Pellegrino Cerino Andrea Ballabio Davide Cacchiarelli et al |
| EPI_ISL_5719184 | Pathogen Genomics Center, National Institute of Infectious Diseases | Pathogen Genomics Center, National Institute of Infectious Diseases | Tsuyoshi Sekizuka et al |
| EPI_ISL_4717771 | Pathogen Genomics Center, National Institute of Infectious Diseases | Pathogen Genomics Center, National Institute of Infectious Diseases | Tsuyoshi Sekizuka et al |
| EPI_ISL_2007394 | Hyogo Prefectural Institute of Public Health science | Pathogen Genomics Center, National Institute of Infectious Diseases | Kenichi Ogita et al |

| | | | |
|---|---|---|---|
| EPI_ISL_1431426 | Osaka Institute of Public Health, Morinomiya Center | Pathogen Genomics Center, National Institute of Infectious Diseases | Tsuyoshi Sekizuka et al |
| EPI_ISL_893313 | Pathogen Genomics Center, National Institute of Infectious Diseases | Pathogen Genomics Center, National Institute of Infectious Diseases | Tsuyoshi Sekizuka et al |
| EPI_ISL_1428404 | Pathogen Genomics Center, National Institute of Infectious Diseases | Pathogen Genomics Center, National Institute of Infectious Diseases | Tsuyoshi Sekizuka et al |
| EPI_ISL_1428396 | Pathogen Genomics Center, National Institute of Infectious Diseases | Pathogen Genomics Center, National Institute of Infectious Diseases | Tsuyoshi Sekizuka et al |
| EPI_ISL_5711599 | Pathogen Genomics Center, National Institute of Infectious Diseases | Pathogen Genomics Center, National Institute of Infectious Diseases | Tsuyoshi Sekizuka et al |
| EPI_ISL_1427193 | Pathogen Genomics Center, National Institute of Infectious Diseases | Pathogen Genomics Center, National Institute of Infectious Diseases | Tsuyoshi Sekizuka et al |
| EPI_ISL_5725468 | Pathogen Genomics Center, National Institute of Infectious Diseases | Pathogen Genomics Center, National Institute of Infectious Diseases | Tsuyoshi Sekizuka et al |
| EPI_ISL_2325416 | SARS-CoV-2 testing team, National Institute of Infectious Diseases | Pathogen Genomics Center, National Institute of Infectious Diseases | Tsuyoshi Sekizuka et al |
| EPI_ISL_4758357 | Pathogen Genomics Center, National Institute of Infectious Diseases | Pathogen Genomics Center, National Institute of Infectious Diseases | Tsuyoshi Sekizuka et al |
| EPI_ISL_2325325 | SARS-CoV-2 testing team, National Institute of Infectious Diseases | Pathogen Genomics Center, National Institute of Infectious Diseases | Tsuyoshi Sekizuka et al |
| EPI_ISL_4706157 | Pathogen Genomics Center, National Institute of Infectious Diseases | Pathogen Genomics Center, National Institute of Infectious Diseases | Tsuyoshi Sekizuka et al |
| EPI_ISL_5743032 | Pathogen Genomics Center, National Institute of Infectious Diseases | Pathogen Genomics Center, National Institute of Infectious Diseases | Tsuyoshi Sekizuka et al |
| EPI_ISL_5760189 | Medical Microbiology and Zoology Section, Okinawa Prefectural Institute of Health and Environment | Medical Microbiology and Zoology Section, Okinawa Prefectural Institute of Health and Environment | Tetsuya Kakita et al |
| EPI_ISL_4642801 | Hyogo Prefectural Institute of Public Health science | Pathogen Genomics Center, National Institute of Infectious Diseases | Kenichi Ogita et al |
| EPI_ISL_7066734 | Pathogen Genomics Center, National Institute of Infectious Diseases | Pathogen Genomics Center, National Institute of Infectious Diseases | Tsuyoshi Sekizuka et al |
| EPI_ISL_7051448 | Pathogen Genomics Center, National Institute of Infectious Diseases | Pathogen Genomics Center, National Institute of Infectious Diseases | Tsuyoshi Sekizuka et al |
| EPI_ISL_7083509 | Pathogen Genomics Center, National Institute of Infectious Diseases | Pathogen Genomics Center, National Institute of Infectious Diseases | Tsuyoshi Sekizuka et al |

| | | | |
|---|---|---|---|
| EPI_ISL_1426533 | Pathogen Genomics Center, National Institute of Infectious Diseases | Pathogen Genomics Center, National Institute of Infectious Diseases | Tsuyoshi Sekizuka et al |
| EPI_ISL_895509 | Pathogen Genomics Center, National Institute of Infectious Diseases | Pathogen Genomics Center, National Institute of Infectious Diseases | Tsuyoshi Sekizuka et al |
| EPI_ISL_1932905 | Pathogen Genomics Center, National Institute of Infectious Diseases | Pathogen Genomics Center, National Institute of Infectious Diseases | Tsuyoshi Sekizuka et al |
| EPI_ISL_5723302 | Pathogen Genomics Center, National Institute of Infectious Diseases | Pathogen Genomics Center, National Institute of Infectious Diseases | Tsuyoshi Sekizuka et al |
| EPI_ISL_6914908 | SARS-CoV-2 testing team, National Institute of Infectious Diseases | Pathogen Genomics Center, National Institute of Infectious Diseases | Tsuyoshi Sekizuka et al |
| EPI_ISL_1430533 | Pathogen Genomics Center, National Institute of Infectious Diseases | Pathogen Genomics Center, National Institute of Infectious Diseases | Tsuyoshi Sekizuka et al |
| EPI_ISL_4702817 | Pathogen Genomics Center, National Institute of Infectious Diseases | Pathogen Genomics Center, National Institute of Infectious Diseases | Tsuyoshi Sekizuka et al |
| EPI_ISL_2253123 | SARS-CoV-2 testing team, National Institute of Infectious Diseases | Pathogen Genomics Center, National Institute of Infectious Diseases | Tsuyoshi Sekizuka et al |
| EPI_ISL_4644797 | Kagoshima Prefectural Institute for Environmental Research and Public Health | Kagoshima Prefectural Institute for Environmental Research and Public Health | Yuka Hamada et al |
| EPI_ISL_2325475 | SARS-CoV-2 testing team, National Institute of Infectious Diseases | Pathogen Genomics Center, National Institute of Infectious Diseases | Tsuyoshi Sekizuka et al |
| EPI_ISL_5713452 | Pathogen Genomics Center, National Institute of Infectious Diseases | Pathogen Genomics Center, National Institute of Infectious Diseases | Tsuyoshi Sekizuka et al |
| EPI_ISL_1430017 | Pathogen Genomics Center, National Institute of Infectious Diseases | Pathogen Genomics Center, National Institute of Infectious Diseases | Tsuyoshi Sekizuka et al |
| EPI_ISL_5717009 | Pathogen Genomics Center, National Institute of Infectious Diseases | Pathogen Genomics Center, National Institute of Infectious Diseases | Tsuyoshi Sekizuka et al |
| EPI_ISL_4701674 | Pathogen Genomics Center, National Institute of Infectious Diseases | Pathogen Genomics Center, National Institute of Infectious Diseases | Tsuyoshi Sekizuka et al |
| EPI_ISL_7050421 | Pathogen Genomics Center, National Institute of Infectious Diseases | Pathogen Genomics Center, National Institute of Infectious Diseases | Tsuyoshi Sekizuka et al |
| EPI_ISL_7052363 | Pathogen Genomics Center, National Institute of Infectious Diseases | Pathogen Genomics Center, National Institute of Infectious Diseases | Tsuyoshi Sekizuka et al |
| EPI_ISL_7055030 | Pathogen Genomics Center, National Institute of Infectious Diseases | Pathogen Genomics Center, National Institute of Infectious Diseases | Tsuyoshi Sekizuka et al |

| | | | |
|---|---|---|---|
| EPI_ISL_4703251 | Pathogen Genomics Center, National Institute of Infectious Diseases | Pathogen Genomics Center, National Institute of Infectious Diseases | Tsuyoshi Sekizuka et al |
| EPI_ISL_480128 | Fukui Prefectural Institute of Public Health and Environmental Science | Pathogen Genomics Center, National Institute of Infectious Diseases | Tsuyoshi Sekizuka et al |
| EPI_ISL_7088178 | Ibaraki Prefectural Institute of Public Health | Ibaraki Prefectural Institute of Public Health | Keiko Goto et al |
| EPI_ISL_7042975 | Pathogen Genomics Center, National Institute of Infectious Diseases | Pathogen Genomics Center, National Institute of Infectious Diseases | Tsuyoshi Sekizuka et al |
| EPI_ISL_7093346 | Kyoto City Institute of Public Health and Environmental Sciences | Kyoto City Institute of Public Health and Environmental Sciences | Yuta Tsukamoto et al |
| EPI_ISL_4722981 | Pathogen Genomics Center, National Institute of Infectious Diseases | Pathogen Genomics Center, National Institute of Infectious Diseases | Tsuyoshi Sekizuka et al |
| EPI_ISL_1427294 | Pathogen Genomics Center, National Institute of Infectious Diseases | Pathogen Genomics Center, National Institute of Infectious Diseases | Tsuyoshi Sekizuka et al |
| EPI_ISL_7104782 | Aomori Prefectural Public Health and Environment Center | Aomori Prefectural Public Health and Environment Center | Motoko Bodaiji et al |
| EPI_ISL_1928293 | Pathogen Genomics Center, National Institute of Infectious Diseases | Pathogen Genomics Center, National Institute of Infectious Diseases | Tsuyoshi Sekizuka et al |
| EPI_ISL_894462 | Pathogen Genomics Center, National Institute of Infectious Diseases | Pathogen Genomics Center, National Institute of Infectious Diseases | Tsuyoshi Sekizuka et al |
| EPI_ISL_691002 | Pathogen Genomics Center, National Institute of Infectious Diseases | Pathogen Genomics Center, National Institute of Infectious Diseases | Tsuyoshi Sekizuka et al |
| EPI_ISL_1123564 | Gunma Prefectural Institute of Public Health and Environmental Sciences | Pathogen Genomics Center, National Institute of Infectious Diseases | Tsuyoshi Sekizuka et al |
| EPI_ISL_5769650 | Department of Microbiology, Tochigi Prefectural Institute of Public Health and Environmental Science | Department of Microbiology, Tochigi Prefectural Institute of Public Health and Environmental Science | Asumi Hirata-Saito et al |
| EPI_ISL_895270 | Pathogen Genomics Center, National Institute of Infectious Diseases | Pathogen Genomics Center, National Institute of Infectious Diseases | Tsuyoshi Sekizuka et al |
| EPI_ISL_7057963 | Pathogen Genomics Center, National Institute of Infectious Diseases | Pathogen Genomics Center, National Institute of Infectious Diseases | Tsuyoshi Sekizuka et al |
| EPI_ISL_1934251 | Pathogen Genomics Center, National Institute of Infectious Diseases | Pathogen Genomics Center, National Institute of Infectious Diseases | Tsuyoshi Sekizuka et al |
| EPI_ISL_1425377 | Department of Infectious Diseases, Kobe Institute of Health | Department of Infectious Diseases, Kobe Institute of Health | Ryohei Nomoto et al |

| | | | |
|---|---|---|---|
| EPI_ISL_5716280 | Pathogen Genomics Center, National Institute of Infectious Diseases | Pathogen Genomics Center, National Institute of Infectious Diseases | Tsuyoshi Sekizuka et al |
| EPI_ISL_685475 | Tokyo Metropolitan Institute of Public Health | Pathogen Genomics Center, National Institute of Infectious Diseases | Tsuyoshi Sekizuka et al |
| EPI_ISL_4703157 | Pathogen Genomics Center, National Institute of Infectious Diseases | Pathogen Genomics Center, National Institute of Infectious Diseases | Tsuyoshi Sekizuka et al |
| EPI_ISL_5725280 | Pathogen Genomics Center, National Institute of Infectious Diseases | Pathogen Genomics Center, National Institute of Infectious Diseases | Tsuyoshi Sekizuka et al |
| EPI_ISL_690947 | Pathogen Genomics Center, National Institute of Infectious Diseases | Pathogen Genomics Center, National Institute of Infectious Diseases | Tsuyoshi Sekizuka et al |
| EPI_ISL_7057607 | Pathogen Genomics Center, National Institute of Infectious Diseases | Pathogen Genomics Center, National Institute of Infectious Diseases | Tsuyoshi Sekizuka et al |
| EPI_ISL_1123473 | Niigata Prefectural Institute of Public Health and Environmental Sciences | Pathogen Genomics Center, National Institute of Infectious Diseases | Tsuyoshi Sekizuka et al |
| EPI_ISL_1428764 | Pathogen Genomics Center, National Institute of Infectious Diseases | Pathogen Genomics Center, National Institute of Infectious Diseases | Tsuyoshi Sekizuka et al |
| EPI_ISL_5770434 | Fukuoka Institute of Health and Environmental Sciences | Fukuoka Institute of Health and Environmental Sciences | Yoshiki Etoh et al |
| EPI_ISL_1125087 | Pathogen Genomics Center, National Institute of Infectious Diseases | Pathogen Genomics Center, National Institute of Infectious Diseases | Tsuyoshi Sekizuka et al |
| EPI_ISL_1425645 | Pathogen Genomics Center, National Institute of Infectious Diseases | Pathogen Genomics Center, National Institute of Infectious Diseases | Tsuyoshi Sekizuka et al |
| EPI_ISL_7062114 | Pathogen Genomics Center, National Institute of Infectious Diseases | Pathogen Genomics Center, National Institute of Infectious Diseases | Tsuyoshi Sekizuka et al |
| EPI_ISL_4642906 | Hyogo Prefectural Institute of Public Health science | Pathogen Genomics Center, National Institute of Infectious Diseases | Kenichi Ogita et al |
| EPI_ISL_5732975 | Pathogen Genomics Center, National Institute of Infectious Diseases | Pathogen Genomics Center, National Institute of Infectious Diseases | Tsuyoshi Sekizuka et al |
| EPI_ISL_7035311 | Pathogen Genomics Center, National Institute of Infectious Diseases | Pathogen Genomics Center, National Institute of Infectious Diseases | Tsuyoshi Sekizuka et al |
| EPI_ISL_1127875 | Pathogen Genomics Center, National Institute of Infectious Diseases | Pathogen Genomics Center, National Institute of Infectious Diseases | Tsuyoshi Sekizuka et al |
| EPI_ISL_2325647 | Chiba Prefectural Institute of Public Health | Pathogen Genomics Center, National Institute of Infectious Diseases | Tsuyoshi Sekizuka et al |

| | | | |
|---|---|---|---|
| EPI_ISL_5703504 | Pathogen Genomics Center, National Institute of Infectious Diseases | Pathogen Genomics Center, National Institute of Infectious Diseases | Tsuyoshi Sekizuka et al |
| EPI_ISL_4710888 | Department of Pathology, National Institute of Infectious Diseases | Department of Pathology, National Institute of Infectious Diseases | Shun Iida et al |
| EPI_ISL_5703798 | Pathogen Genomics Center, National Institute of Infectious Diseases | Pathogen Genomics Center, National Institute of Infectious Diseases | Tsuyoshi Sekizuka et al |
| EPI_ISL_5732957 | Pathogen Genomics Center, National Institute of Infectious Diseases | Pathogen Genomics Center, National Institute of Infectious Diseases | Tsuyoshi Sekizuka et al |
| EPI_ISL_1123453 | Pathogen Genomics Center, National Institute of Infectious Diseases | Pathogen Genomics Center, National Institute of Infectious Diseases | Tsuyoshi Sekizuka et al |
| EPI_ISL_4849590 | SARS-CoV-2 testing team, National Institute of Infectious Diseases | Pathogen Genomics Center, National Institute of Infectious Diseases | Tsuyoshi Sekizuka et al |
| EPI_ISL_4701106 | SARS-CoV-2 testing team, National Institute of Infectious Diseases | Pathogen Genomics Center, National Institute of Infectious Diseases | Tsuyoshi Sekizuka et al |
| EPI_ISL_4701487 | Pathogen Genomics Center, National Institute of Infectious Diseases | Pathogen Genomics Center, National Institute of Infectious Diseases | Tsuyoshi Sekizuka et al |
| EPI_ISL_1429421 | Pathogen Genomics Center, National Institute of Infectious Diseases | Pathogen Genomics Center, National Institute of Infectious Diseases | Tsuyoshi Sekizuka et al |
| EPI_ISL_5699938 | Pathogen Genomics Center, National Institute of Infectious Diseases | Pathogen Genomics Center, National Institute of Infectious Diseases | Tsuyoshi Sekizuka et al |
| EPI_ISL_4721360 | Pathogen Genomics Center, National Institute of Infectious Diseases | Pathogen Genomics Center, National Institute of Infectious Diseases | Tsuyoshi Sekizuka et al |
| EPI_ISL_897268 | Pathogen Genomics Center, National Institute of Infectious Diseases | Pathogen Genomics Center, National Institute of Infectious Diseases | Tsuyoshi Sekizuka et al |
| EPI_ISL_6913953 | SARS-CoV-2 testing team, National Institute of Infectious Diseases | Pathogen Genomics Center, National Institute of Infectious Diseases | Tsuyoshi Sekizuka et al |
| EPI_ISL_5738193 | Pathogen Genomics Center, National Institute of Infectious Diseases | Pathogen Genomics Center, National Institute of Infectious Diseases | Tsuyoshi Sekizuka et al |
| EPI_ISL_5713614 | Pathogen Genomics Center, National Institute of Infectious Diseases | Pathogen Genomics Center, National Institute of Infectious Diseases | Tsuyoshi Sekizuka et al |
| EPI_ISL_1302665 | "InMedica" | Lithuanian University of Health Sciences Hospital, Department of Genetics and Molecular Medicine | Rasa Ugenskiene et al |
| EPI_ISL_3877141 | Pathogen Genomics Center, National Institute of Infectious Diseases | Pathogen Genomics Center, National Institute of Infectious Diseases | Tsuyoshi Sekizuka et al |

| | | | |
|---|---|---|---|
| EPI_ISL_2335808 | Pathogen Genomics Center, National Institute of Infectious Diseases | Pathogen Genomics Center, National Institute of Infectious Diseases | Tsuyoshi Sekizuka et al |
| EPI_ISL_3199347 | Pathogen Genomics Center, National Institute of Infectious Diseases | Pathogen Genomics Center, National Institute of Infectious Diseases | Tsuyoshi Sekizuka et al |
| EPI_ISL_3197763 | Pathogen Genomics Center, National Institute of Infectious Diseases | Pathogen Genomics Center, National Institute of Infectious Diseases | Tsuyoshi Sekizuka et al |
| EPI_ISL_2602725 | KEMRI-Wellcome Trust Research Programme,Kilifi | KEMRI-Wellcome Trust Research Programme,Kilifi | Githinji G. et al |
| EPI_ISL_2329863 | SARS-CoV-2 testing team, National Institute of Infectious Diseases | Pathogen Genomics Center, National Institute of Infectious Diseases | Tsuyoshi Sekizuka et al |
| EPI_ISL_4676015 | SARS-CoV-2 testing team, National Institute of Infectious Diseases | Pathogen Genomics Center, National Institute of Infectious Diseases | Tsuyoshi Sekizuka et al |
| EPI_ISL_2772981 | Fukuoka Institute of Health and Environmental Sciences | Fukuoka Institute of Health and Environmental Sciences | Saori Ueda et al |
| EPI_ISL_3900226 | Pathogen Genomics Center, National Institute of Infectious Diseases | Pathogen Genomics Center, National Institute of Infectious Diseases | Tsuyoshi Sekizuka et al |
| EPI_ISL_689179 | Pathogen Genomics Center, National Institute of Infectious Diseases | Pathogen Genomics Center, National Institute of Infectious Diseases | Tsuyoshi Sekizuka et al |
| EPI_ISL_2765352 | SARS-CoV-2 testing team, National Institute of Infectious Diseases | Pathogen Genomics Center, National Institute of Infectious Diseases | Tsuyoshi Sekizuka et al |
| EPI_ISL_1654677 | NVMRVI | National Public Health Surveillance Laboratory | Lukas Zemaitis et al |
| EPI_ISL_2335422 | Pathogen Genomics Center, National Institute of Infectious Diseases | Pathogen Genomics Center, National Institute of Infectious Diseases | Tsuyoshi Sekizuka et al |
| EPI_ISL_3196898 | Pathogen Genomics Center, National Institute of Infectious Diseases | Pathogen Genomics Center, National Institute of Infectious Diseases | Tsuyoshi Sekizuka et al |
| EPI_ISL_1579950 | NVSPL | National Public Health Surveillance Laboratory | Lukas Zemaitis et al |
| EPI_ISL_4951455 | UAB Diagnostikos laboratorija | National Public Health Surveillance Laboratory | Lukas Zemaitis et al |
| EPI_ISL_2779346 | Homabay County Referral Hospital | USAMRD-A, Basic Science Laboratory | Gathii Kimita et al |
| EPI_ISL_5144484 | Department of Microbiology, National Institute of Public Health of Kosovo | Department of Microbiology, National Institute of Public Health of Kosovo | Xhevat Jakupi et al |
| EPI_ISL_2763358 | SARS-CoV-2 testing team, National Institute of Infectious Diseases | Pathogen Genomics Center, National Institute of Infectious Diseases | Tsuyoshi Sekizuka et al |
| EPI_ISL_855520 | KEMRI-Wellcome Trust Research Programme/KEMRI-CGMR-C Kilifi | KEMRI-Wellcome Trust Research Programme/KEMRI-CGMR-C Kilifi | Githinji et al |

| | | | |
|---|---|---|---|
| EPI_ISL_3198264 | Pathogen Genomics Center, National Institute of Infectious Diseases | Pathogen Genomics Center, National Institute of Infectious Diseases | Tsuyoshi Sekizuka et al |
| EPI_ISL_3197652 | Pathogen Genomics Center, National Institute of Infectious Diseases | Pathogen Genomics Center, National Institute of Infectious Diseases | Tsuyoshi Sekizuka et al |
| EPI_ISL_901885 | Pathogen Genomics Center, National Institute of Infectious Diseases | Pathogen Genomics Center, National Institute of Infectious Diseases | Tsuyoshi Sekizuka et al |
| EPI_ISL_3899517 | Pathogen Genomics Center, National Institute of Infectious Diseases | Pathogen Genomics Center, National Institute of Infectious Diseases | Tsuyoshi Sekizuka et al |
| EPI_ISL_3931886 | Genome Information Research Center, Research Institute for Microbial Diseases, Osaka University | Genome Information Research Center, Research Institute for Microbial Diseases, Osaka University | Daisuke Motooka et al |
| EPI_ISL_6944080 | KEMRI-Wellcome Trust Research Programme,Kilifi | KEMRI-Wellcome Trust Research Programme,Kilifi | Githinji G. et al |
| EPI_ISL_3864218 | UAB Medicina practica laboratorija | unknown | Rasa Ugenskiene et al |
| EPI_ISL_687253 | Pathogen Genomics Center, National Institute of Infectious Diseases | Pathogen Genomics Center, National Institute of Infectious Diseases | Tsuyoshi Sekizuka et al |
| EPI_ISL_3900051 | Pathogen Genomics Center, National Institute of Infectious Diseases | Pathogen Genomics Center, National Institute of Infectious Diseases | Tsuyoshi Sekizuka et al |
| EPI_ISL_3197778 | Pathogen Genomics Center, National Institute of Infectious Diseases | Pathogen Genomics Center, National Institute of Infectious Diseases | Tsuyoshi Sekizuka et al |
| EPI_ISL_3711109 | UAB "Baltic Medics" | National Public Health Surveillance Laboratory | Lukas Zemaitis et al |
| EPI_ISL_3889312 | SARS-CoV-2 testing team, National Institute of Infectious Diseases | Pathogen Genomics Center, National Institute of Infectious Diseases | Tsuyoshi Sekizuka et al |
| EPI_ISL_2337585 | Pathogen Genomics Center, National Institute of Infectious Diseases | Pathogen Genomics Center, National Institute of Infectious Diseases | Tsuyoshi Sekizuka et al |
| EPI_ISL_4694324 | Pathogen Genomics Center, National Institute of Infectious Diseases | Pathogen Genomics Center, National Institute of Infectious Diseases | Tsuyoshi Sekizuka et al |
| EPI_ISL_1613231 | LIC | Latvian Biomedical Research and Study Centre | Janis Pjalkovskis et al |
| EPI_ISL_2336524 | SARS-CoV-2 testing team, National Institute of Infectious Diseases | Pathogen Genomics Center, National Institute of Infectious Diseases | Tsuyoshi Sekizuka et al |
| EPI_ISL_2771575 | SARS-CoV-2 testing team, National Institute of Infectious Diseases | Pathogen Genomics Center, National Institute of Infectious Diseases | Tsuyoshi Sekizuka et al |
| EPI_ISL_2777034 | SARS-CoV-2 testing team, National Institute of Infectious Diseases | Pathogen Genomics Center, National Institute of Infectious Diseases | Tsuyoshi Sekizuka et al |

| | | | |
|---|---|---|---|
| EPI_ISL_4660821 | SARS-CoV-2 testing team, National Institute of Infectious Diseases | Pathogen Genomics Center, National Institute of Infectious Diseases | Tsuyoshi Sekizuka et al |
| EPI_ISL_2772997 | Fukuoka Institute of Health and Environmental Sciences | Fukuoka Institute of Health and Environmental Sciences | Saori Ueda et al |
| EPI_ISL_933876 | Vilnius university hospital Santaros Klinikos, Center of Laboratory Medicine | Vilnius university hospital Santaros Klinikos, Center of Laboratory Medicine | Ingrida Olendraite et al |
| EPI_ISL_2325749 | Pathogen Genomics Center, National Institute of Infectious Diseases | Pathogen Genomics Center, National Institute of Infectious Diseases | Tsuyoshi Sekizuka et al |
| EPI_ISL_3245760 | Department of Microbiology, Yamanashi Institute for Public Health and Environment | Genome Analysis Center, Yamanashi Central Hospital | Yosuke Hirotsu et al |
| EPI_ISL_2760765 | SARS-CoV-2 testing team, National Institute of Infectious Diseases | Pathogen Genomics Center, National Institute of Infectious Diseases | Tsuyoshi Sekizuka et al |
| EPI_ISL_568712 | KEMRI-Wellcome Trust Research Programme/KEMRI-CGMR-C Kilifi | KEMRI-Wellcome Trust Research Programme/KEMRI-CGMR-C Kilifi | Githinji et al 2020 et al |
| EPI_ISL_2357443 | Fukuoka Institute of Health and Environmental Sciences | Fukuoka Institute of Health and Environmental Sciences | Asako Nakamura et al |
| EPI_ISL_3888900 | Pathogen Genomics Center, National Institute of Infectious Diseases | Pathogen Genomics Center, National Institute of Infectious Diseases | Tsuyoshi Sekizuka et al |
| EPI_ISL_3879941 | Pathogen Genomics Center, National Institute of Infectious Diseases | Pathogen Genomics Center, National Institute of Infectious Diseases | Tsuyoshi Sekizuka et al |
| EPI_ISL_2330365 | Pathogen Genomics Center, National Institute of Infectious Diseases | Pathogen Genomics Center, National Institute of Infectious Diseases | Tsuyoshi Sekizuka et al |
| EPI_ISL_3896972 | SARS-CoV-2 testing team, National Institute of Infectious Diseases | Pathogen Genomics Center, National Institute of Infectious Diseases | Tsuyoshi Sekizuka et al |
| EPI_ISL_4691709 | Pathogen Genomics Center, National Institute of Infectious Diseases | Pathogen Genomics Center, National Institute of Infectious Diseases | Tsuyoshi Sekizuka et al |
| EPI_ISL_5628135 | Tokyo Metropolitan Institute of Public Health | Tokyo Metropolitan Institute of Public Health | Hirofumi Miyake et al |
| EPI_ISL_3896192 | SARS-CoV-2 testing team, National Institute of Infectious Diseases | Pathogen Genomics Center, National Institute of Infectious Diseases | Tsuyoshi Sekizuka et al |
| EPI_ISL_2338160 | SARS-CoV-2 testing team, National Institute of Infectious Diseases | Pathogen Genomics Center, National Institute of Infectious Diseases | Tsuyoshi Sekizuka et al |
| EPI_ISL_1341150 | Laboratory of Virology, National center of expertise | RSE "National Center of Expertise" and RSE "National center for Biotechnology" | Abdaliyev Askar et al |
| EPI_ISL_2762754 | SARS-CoV-2 testing team, National Institute of Infectious Diseases | Pathogen Genomics Center, National Institute of Infectious Diseases | Tsuyoshi Sekizuka et al |

| | | | |
|---|---|---|---|
| EPI_ISL_4232152 | National Public Health Institute of Liberia Reference Lab | Center for Infection and Immunity, Columbia University | Bode Shobayo et al |
| EPI_ISL_4847384 | Genome Information Research Center, Research Institute for Microbial Diseases, Osaka University | Genome Information Research Center, Research Institute for Microbial Diseases, Osaka University | Daisuke Motooka et al |
| EPI_ISL_3858650 | UAB Medicina practica laboratorija | Hospital of Lithuanian University of Health Sciences (LSMU) Kaunas Clinics | Rasa Ugenskiene et al |
| EPI_ISL_3890311 | Pathogen Genomics Center, National Institute of Infectious Diseases | Pathogen Genomics Center, National Institute of Infectious Diseases | Tsuyoshi Sekizuka et al |
| EPI_ISL_3464432 | UAB Diagnostikos laboratorija | National Public Health Surveillance Laboratory | Lukas Zemaitis et al |
| EPI_ISL_3876826 | SARS-CoV-2 testing team, National Institute of Infectious Diseases | Pathogen Genomics Center, National Institute of Infectious Diseases | Tsuyoshi Sekizuka et al |
| EPI_ISL_2763332 | SARS-CoV-2 testing team, National Institute of Infectious Diseases | Pathogen Genomics Center, National Institute of Infectious Diseases | Tsuyoshi Sekizuka et al |
| EPI_ISL_3193436 | Pathogen Genomics Center, National Institute of Infectious Diseases | Pathogen Genomics Center, National Institute of Infectious Diseases | Tsuyoshi Sekizuka et al |
| EPI_ISL_7027255 | Genome Information Research Center, Research Institute for Microbial Diseases, Osaka University | Genome Information Research Center, Research Institute for Microbial Diseases, Osaka University | Daisuke Motooka et al |
| EPI_ISL_2338972 | SARS-CoV-2 testing team, National Institute of Infectious Diseases | Pathogen Genomics Center, National Institute of Infectious Diseases | Tsuyoshi Sekizuka et al |
| EPI_ISL_3874835 | Fukuoka Institute of Health and Environmental Sciences | Fukuoka Institute of Health and Environmental Sciences | Yoshiki Etoh et al |
| EPI_ISL_4691739 | Pathogen Genomics Center, National Institute of Infectious Diseases | Pathogen Genomics Center, National Institute of Infectious Diseases | Tsuyoshi Sekizuka et al |
| EPI_ISL_2137825 | Life Sciences Center, Vilnius University | Institute of Biotechnology, Life Sciences Center, Vilnius University | Emilija Vasiliunaite et al |
| EPI_ISL_4878451 | UAB InMedica | Hospital of Lithuanian University of Health Sciences (LSMU) Kaunas Clinics | Rasa Ugenskiene et al |
| EPI_ISL_2765504 | SARS-CoV-2 testing team, National Institute of Infectious Diseases | Pathogen Genomics Center, National Institute of Infectious Diseases | Tsuyoshi Sekizuka et al |
| EPI_ISL_933886 | Vilnius university hospital Santaros Klinikos, Center of Laboratory Medicine | Vilnius university hospital Santaros Klinikos, Center of Laboratory Medicine | Ingrida Olendraite et al |
| EPI_ISL_1312608 | E. Gulbja laboratorija | Latvian Biomedical Research and Study Centre | Janis Pjalkovskis et al |
| EPI_ISL_3899007 | SARS-CoV-2 testing team, National Institute of Infectious Diseases | Pathogen Genomics Center, National Institute of Infectious Diseases | Tsuyoshi Sekizuka et al |

| | | | |
|---|---|---|---|
| EPI_ISL_3895704 | Pathogen Genomics Center, National Institute of Infectious Diseases | Pathogen Genomics Center, National Institute of Infectious Diseases | Tsuyoshi Sekizuka et al |
| EPI_ISL_3195052 | SARS-CoV-2 testing team, National Institute of Infectious Diseases | Pathogen Genomics Center, National Institute of Infectious Diseases | Tsuyoshi Sekizuka et al |
| EPI_ISL_2894564 | Hospital Sharp | Microbial Genomics Laboratory | Consorcio Mexicano de Vigilancia Genómica (CoViGen-Mex). Authors (in alphabetical order): Julio Elias Alvarado-Yaah et al |
| EPI_ISL_1384229 | Laboratoires d'analyses medicales - Ketterthill | Laboratoire national de sante, Microbiology, Microbial Genomics Platform | Anke Wienecke-Baldacchino et al |
| EPI_ISL_7084389 | UAB "Baltic Medics" | Vilnius University Hospital Santaros Klinikos, Center of Laboratory Medicine | Monika Katenaite et al |
| EPI_ISL_6132052 | Jabatan Forensik Hospital Sultanah Bahiyah | Department of Medical Microbiology, Faculty of Medicine, University of Malaya; University of Malaya Medical Centre | Omar Khalilur Rahman et al |
| EPI_ISL_1523184 | Laboratoire national de sante, Microbiology, Virology | Laboratoire national de sante, Microbiology, Microbial Genomics Platform | Anke Wienecke-Baldacchino et al |
| EPI_ISL_2694572 | UAB Medicina practica laboratorija | National Public Health Surveillance Laboratory | Lukas Zemaitis et al |
| EPI_ISL_4399906 | Dutch COVID-19 response team | National Institute for Public Health and the Environment (RIVM) | Adam Meijer et al |
| EPI_ISL_6586366 | Bintulu Medical Centre (Bintulu) | Institute of Health and Community Medicine | Chan Chia Jui et al |
| EPI_ISL_6775376 | Unidad de Investigacion Medica de Yucatan (UIMY) | Unidad de Genomica Avanzada | Consorcio Mexicano de Vigilancia Genomica (CoViGen-Mex). Authors (in alphabetical order): Julio Elias Alvarado-Yaah et al |
| EPI_ISL_1384256 | Laboratoires d'analyses medicales - Ketterthill | Laboratoire national de sante, Microbiology, Microbial Genomics Platform | Anke Wienecke-Baldacchino et al |
| EPI_ISL_744135 | Laboratoire national de santé, Microbiology, Virology | Laboratoire national de santé, Microbiology, Microbial Genomics Platform | Anke Wienecke-Baldacchino et al |
| EPI_ISL_2442181 | Centro de Investigação em Saúde de Manhiça (CISM) & ISGlobal, Institut de Salut Global de Barcelona | Instituto de Biomedicina de València - CSIC | Alfredo Mayor et al |
| EPI_ISL_5937731 | Nacionaline visuomenes sveikatos prieziuros laboratorija | Lithuanian University of Health Sciences | Dovydas Gecys et al |
| EPI_ISL_1792535 | Dutch COVID-19 response team | National Institute for Public Health and the Environment (RIVM) | Adam Meijer et al |
| EPI_ISL_3258539 | Dutch COVID-19 response team | National Institute for Public Health and the Environment (RIVM) | Adam Meijer et al |

| | | | |
|---|---|---|---|
| EPI_ISL_2788074 | Dutch COVID-19 response team | National Institute for Public Health and the Environment (RIVM) | Adam Meijer et al |
| EPI_ISL_1758172 | Vilnius University Hospital Santaros Klinikos, Center of Laboratory Medicine | Vilnius University Hospital Santaros Klinikos, Center of Laboratory Medicine | Monika Katenaite et al |
| EPI_ISL_1917256 | Hospital Center Emile Mayrisch | Laboratoire national de sante, Microbiology, Microbial Genomics Platform | Anke Wienecke-Baldacchino et al |
| EPI_ISL_6097704 | LESP Puebla | Instituto de Diagnostico y Referencia Epidemiologicos (INDRE) | Claudia Wong-Arambula et al |
| EPI_ISL_1661615 | UAB Medicina Practica Laboratory | Vilnius University Hospital Santaros Klinikos, Center of Laboratory Medicine | Monika Katenaite et al |
| EPI_ISL_1908585 | UAB InMedica | Vilnius University Hospital Santaros Klinikos, Center of Laboratory Medicine | Gytis Dudas et al |
| EPI_ISL_2082214 | Nacionalinis maisto ir veterinarijos rizikos vertinimo institutas | National Public Health Surveillance Laboratory | Lukas Zemaitis et al |
| EPI_ISL_1917585 | Laboratoires d'analyses medicales - Ketterthill | Laboratoire national de sante, Microbiology, Microbial Genomics Platform | Anke Wienecke-Baldacchino et al |
| EPI_ISL_1406096 | Instituto Nacional de Medicina Genomica | Instituto Nacional de Medicina Genomica | Hidalgo-Miranda A et al |
| EPI_ISL_3355554 | Centro de Investigación Biomédica del Noreste (CIBIN) | Instituto de Biotecnología de la UNAM | Consorcio Mexicano de Vigilancia Genómica (CoViGen-Mex). Authors (in alphabetical order): Julio Elias Alvarado-Yaah et al |
| EPI_ISL_2340865 | LESP Michoacan | Instituto de Diagnostico y Referencia Epidemiologicos (INDRE) | Claudia Wong-Arambula et al |
| EPI_ISL_576279 | Instituto de Diagnostico y Referencia Epidemiologicos (INDRE) | Instituto de Diagnostico y Referencia Epidemiologicos (INDRE) | Ernesto Ramirez-Gonzalez et al |
| EPI_ISL_2535735 | Borneo Medical Centre | Institute of Health and Community Medicine | Chan Chia Jui et al |
| EPI_ISL_1510561 | Klaipeda University Hospital | Vilnius University Hospital Santaros Klinikos, Center of Laboratory Medicine | Ingrida Olendraite et al |
| EPI_ISL_5391822 | BioneXt Lab | Laboratoire national de sante, Microbiology, Microbial Genomics Platform | Anke Wienecke-Baldacchino et al |
| EPI_ISL_3257776 | Dutch COVID-19 response team | National Institute for Public Health and the Environment (RIVM) | Adam Meijer et al |
| EPI_ISL_1585882 | UAB InMedica | Vilnius University Hospital Santaros Klinikos, Center of Laboratory Medicine | Dovile Ezerskyte et al |

| | | | |
|---|---|---|---|
| EPI_ISL_1662043 | Centro de Investigacion Biomedica del Noreste (CIBIN) | Unidad de Genomica Avanzada | Consorcio Mexicano de Vigilancia Genomica (CoViGen-Mex). Authors (in alphabetical order): Julio Elias Alvarado-Yaah et al |
| EPI_ISL_1398498 | National Food and Veterinary Risk Assessment Institute | Vilnius University Hospital Santaros Klinikos, Center of Laboratory Medicine | Gytis Dudas et al |
| EPI_ISL_4076428 | Dutch COVID-19 response team | National Institute for Public Health and the Environment (RIVM) | Adam Meijer et al |
| EPI_ISL_2339611 | UMC Groningen, Clinical Virology, Department of Medical Microbiology and Infection Prevention | UMC Groningen, Clinical Virology, Department of Medical Microbiology and Infection Prevention | Hubert Niesters et al |
| EPI_ISL_3864887 | UAB InMedica | Lithuanian University of Health Sciences | Arnoldas Pautienius et al |
| EPI_ISL_4171473 | Outre mer | National Reference Center for Viruses of Respiratory Infections, Institut Pasteur, Paris | Marion Barbet et al |
| EPI_ISL_1962624 | Dutch COVID-19 response team | National Institute for Public Health and the Environment (RIVM) | Adam Meijer et al |
| EPI_ISL_1301521 | Instituto de Diagnostico y Referencia Epidemiologicos InDRE_RNLSP | Instituto de Biotecnología de la UNAM | Authors from IBT et al |
| EPI_ISL_3564524 | Laboratory for Molecular Diagnostics, IPHMN | Eurofins Genomics Europe Sequencing GmbH | Danijela Vujošević et al |
| EPI_ISL_6863633 | Laboratorio Central de Epidemiología (LCE) | Instituto de Biotecnología de la UNAM | Consorcio Mexicano de Vigilancia Genómica (CoViGen-Mex). Authors (in alphabetical order): Julio Elias Alvarado-Yaah et al |
| EPI_ISL_3943411 | Instituto Nacional de Medicina Genomica | Instituto Nacional de Medicina Genomica | Cedro-Tanda A et al |
| EPI_ISL_1962557 | Dutch COVID-19 response team | National Institute for Public Health and the Environment (RIVM) | Adam Meijer et al |
| EPI_ISL_6963092 | Malawi Liverpool Wellcome Trust Clinical Research Program | Malawi Liverpool Wellcome Trust Clinical Research Program | Catherine Anscombe et al |
| EPI_ISL_7015235 | Selangor State Health Department | Institute for Medical Research, Infectious Disease Research Centre, National Institutes of Health, Ministry of Health Malaysia | Suppiah J et al |
| EPI_ISL_1262688 | Instituto Nacional de Medicina Genomica | Instituto Nacional de Medicina Genomica | Hidalgo-Miranda A et al |
| EPI_ISL_1494627 | LESP Chihuahua | Instituto de Diagnostico y Referencia Epidemiologicos (INDRE) | Claudia Wong-Arambula et al |
| EPI_ISL_6181766 | Instituto Nacional de Medicina Genomica | Instituto Nacional de Medicina Genomica | Cedro-Tanda A et al |
| EPI_ISL_2341043 | Salud Digna | Instituto Nacional de Medicina Genomica | Hidalgo-Miranda A et al |

| | | | |
|---|---|---|---|
| EPI_ISL_1522093 | Dutch COVID-19 response team | National Institute for Public Health and the Environment (RIVM) | Adam Meijer et al |
| EPI_ISL_4220124 | Instituto Nacional de Medicina Genomica | Instituto Nacional de Medicina Genomica | Cedro-Tanda A et al |
| EPI_ISL_1370516 | Dutch COVID-19 response team | National Institute for Public Health and the Environment (RIVM) | Adam Meijer et al |
| EPI_ISL_1961982 | Dutch COVID-19 response team | National Institute for Public Health and the Environment (RIVM) | Adam Meijer et al |
| EPI_ISL_4448328 | UAB "Rezus.lt" | National Public Health Surveillance Laboratory | Lukas Zemaitis et al |
| EPI_ISL_1662182 | Laboratorio Central de Epidemiologia (LCE) | Unidad de Genomica Avanzada | Consorcio Mexicano de Vigilancia Genomica (CoViGen-Mex). Authors (in alphabetical order): Julio Elias Alvarado-Yaah et al |
| EPI_ISL_910426 | Laboratoire national de sante, Microbiology, Virology | Laboratoire national de sante, Microbiology, Microbial Genomics Platform | Anke Wienecke-Baldacchino et al |
| EPI_ISL_3245942 | Outre mer | National Reference Center for Viruses of Respiratory Infections, Institut Pasteur, Paris | Marion Barbet et al |
| EPI_ISL_1596789 | Dutch COVID-19 response team | National Institute for Public Health and the Environment (RIVM) | Adam Meijer et al |
| EPI_ISL_1792394 | Dutch COVID-19 response team | National Institute for Public Health and the Environment (RIVM) | Adam Meijer et al |
| EPI_ISL_2082287 | Nacionalinis maisto ir veterinarijos rizikos vertinimo institutas | National Public Health Surveillance Laboratory | Lukas Zemaitis et al |
| EPI_ISL_4879410 | Viesoji istaiga Vilniaus universiteto ligonine Santaros klinikos | Vilnius University Hospital Santaros Klinikos, Center of Laboratory Medicine | Monika Katenaite et al |
| EPI_ISL_3023689 | UMC Groningen, Clinical Virology, Department of Medical Microbiology and Infection Prevention | UMC Groningen, Clinical Virology, Department of Medical Microbiology and Infection Prevention | Hubert Niesters et al |
| EPI_ISL_5391643 | BioneXt Lab | Laboratoire national de sante, Microbiology, Microbial Genomics Platform | Anke Wienecke-Baldacchino et al |
| EPI_ISL_5395166 | Laboratoires d'analyses medicales - Ketterthill | Laboratoire national de sante, Microbiology, Microbial Genomics Platform | Anke Wienecke-Baldacchino et al |
| EPI_ISL_1315300 | National Food and Veterinary Risk Assessment Institute | Vilnius University Hospital Santaros Klinikos, Center of Laboratory Medicine | Ingrida Olendraite et al |
| EPI_ISL_5850722 | Centro de investigacion Biomedica del Noreste (CiBiN) | Unidad de Genomica Avanzada | Consorcio Mexicano de Vigilancia Genomica (CoViGen-Mex). Authors (in alphabetical order): Julio Elias Alvarado-Yaah et al |
| EPI_ISL_1911842 | National Institute for Food and Veterinary Risk Assessment (NMVRVI) | National Public Health Surveillance Laboratory | Lukas Zemaitis et al |

| | | | |
|---|---|---|---|
| EPI_ISL_2428357 | Nacionaline visuomenes sveikatos prieziuros laboratorija | National Public Health Surveillance Laboratory | Lukas Zemaitis et al |
| EPI_ISL_1917398 | Hospital Center Luxembourg | Laboratoire national de sante, Microbiology, Microbial Genomics Platform | Anke Wienecke-Baldacchino et al |
| EPI_ISL_2928490 | Arión Genética | Instituto Nacional de Medicina Genomica | Cedro-Tanda A et al |
| EPI_ISL_2712678 | Instituto de Diagnostico y Referencia Epidemiologicos (InDRE) | Andersen lab at Scripps Research | SEARCH Alliance with Gisela Barrera Badillo et al |
| EPI_ISL_2095964 | National Food and Veterinary Risk Assessment Institute | National Public Health Surveillance Laboratory | Lukas Zemaitis et al |
| EPI_ISL_3943579 | Salud Digna | Instituto Nacional de Medicina Genomica | Cedro-Tanda A et al |
| EPI_ISL_5194118 | Laboratorio Central de Epidemiología (LCE) | Centro de Investigación en Enfermedades Infecciosas (CIENI), Instituto Nacional de Enfermedades Respiratorias (INER) | Consorcio Mexicano de Vigilancia Genómica (CoViGen-Mex). Authors (in alphabetical order): Julio Elias Alvarado-Yaah et al |
| EPI_ISL_3145793 | BioneXt Lab | Laboratoire national de sante, Microbiology, Microbial Genomics Platform | Anke Wienecke-Baldacchino et al |
| EPI_ISL_4741160 | Medical Virology and BSL3 Laboratory | Centre de Séquençage Génomique | EL Hamouchi Adil et al |
| EPI_ISL_6827843 | PathLab Bay of Plenty | Institute of Environmental Science and Research (ESR) | Sarah Cockerton et al |
| EPI_ISL_4544303 | Dutch COVID-19 response team | National Institute for Public Health and the Environment (RIVM) | Adam Meijer et al |
| EPI_ISL_1252539 | University Hospital of Northern Norway, Department for Microbiology and Infectious Disease Control | Norwegian Institute of Public Health, Department of Virology | Kathrine Stene-Johansen et al |
| EPI_ISL_6841608 | Dutch COVID-19 response team | National Institute for Public Health and the Environment (RIVM) | Adam Meijer et al |
| EPI_ISL_1522230 | Dutch COVID-19 response team | National Institute for Public Health and the Environment (RIVM) | Adam Meijer et al |
| EPI_ISL_1457694 | Dutch COVID-19 response team | National Institute for Public Health and the Environment (RIVM) | Adam Meijer et al |
| EPI_ISL_764532 | Oxford Viromics, NDM, University of Oxford; Oxford University Hospitals; Basingstoke and North Hampshire Hospital | COVID-19 Genomics UK (COG-UK) Consortium | Tanya Golubchik et al |
| EPI_ISL_1116413 | Randox Laboratories | Wellcome Sanger Institute for the COVID-19 Genomics UK (COG-UK) Consortium | Randox Laboratories et al |
| EPI_ISL_6989155 | Furst Medical Laboratory | Norwegian Institute of Public Health, Department of Virology | Kathrine Stene-Johansen et al |
| EPI_ISL_1975618 | Regional Virus Laboratory, Belfast Health and Social Care Trust | COVID-19 Genomics UK (COG-UK) Consortium | Conall McCaughey et al |

| | | | |
|---|---|---|---|
| EPI_ISL_1792290 | Dutch COVID-19 response team | National Institute for Public Health and the Environment (RIVM) | Adam Meijer et al |
| EPI_ISL_2333338 | Haukeland University Hospital, Dept. of Microbiology | Norwegian Institute of Public Health, Department of Virology | 'Kathrine Stene-Johansen et al |
| EPI_ISL_2491543 | Unilabs Laboratory Medicine | Norwegian Institute of Public Health, Department of Virology | Kathrine Stene-Johansen et al |
| EPI_ISL_4275108 | Ostfold Hospital Trust - Kalnes, Centre for Laboratory Medicine, Section for gene technology and infection serology | Norwegian Institute of Public Health, Department of Virology | Kathrine Stene-Johansen et al |
| EPI_ISL_2787298 | Dutch COVID-19 response team | National Institute for Public Health and the Environment (RIVM) | Adam Meijer et al |
| EPI_ISL_6841609 | Dutch COVID-19 response team | National Institute for Public Health and the Environment (RIVM) | Adam Meijer et al |
| EPI_ISL_547501 | Dutch COVID-19 response team | National Institute for Public Health and the Environment (RIVM) | Adam Meijer et al |
| EPI_ISL_4026558 | Biorepository and Clinical Virology Laboratory | Northwestern University - Center for Pathogen Genomics and Microbial Evolution | Ramon Lorenzo-Redondo et al |
| EPI_ISL_1054404 | Queens Medical Centre, Clinical Microbiology Department / DeepSeq Nottingham | COVID-19 Genomics UK (COG-UK) Consortium | Gemma Clark et al |
| EPI_ISL_4836165 | Dutch COVID-19 response team | National Institute for Public Health and the Environment (RIVM) | Adam Meijer et al |
| EPI_ISL_1705540 | Dutch COVID-19 response team | National Institute for Public Health and the Environment (RIVM) | Adam Meijer et al |
| EPI_ISL_3426605 | Microvida | Microvida | Suzan D. Pas et al |
| EPI_ISL_2093301 | Dutch COVID-19 response team | National Institute for Public Health and the Environment (RIVM) | Adam Meijer et al |
| EPI_ISL_3230005 | Dept. of Medical Microbiology, Stavanger University Hospital, Helse Stavanger HF | Norwegian Institute of Public Health, Department of Virology | Kathrine Stene-Johansen et al |
| EPI_ISL_2981666 | Dutch COVID-19 response team | National Institute for Public Health and the Environment (RIVM) | Adam Meijer et al |
| EPI_ISL_1311241 | Dutch COVID-19 response team | Erasmus Medical Center | Bas Oude Munnink et al |
| EPI_ISL_3258533 | Dutch COVID-19 response team | National Institute for Public Health and the Environment (RIVM) | Adam Meijer et al |
| EPI_ISL_1596313 | Dutch COVID-19 response team | National Institute for Public Health and the Environment (RIVM) | Adam Meijer et al |
| EPI_ISL_1975459 | Regional Virus Laboratory, Belfast Health and Social Care Trust | COVID-19 Genomics UK (COG-UK) Consortium | Conall McCaughey et al |
| EPI_ISL_1158693 | Dutch COVID-19 response team | Medical Microbiology, Maastricht University Medical Centre | Jozef Dingemans* et al |
| EPI_ISL_2981445 | Dutch COVID-19 response team | National Institute for Public Health and the Environment (RIVM) | Adam Meijer et al |
| EPI_ISL_2609695 | Dutch COVID-19 response team | National Institute for Public Health and the Environment (RIVM) | Adam Meijer et al |

| | | | |
|---|---|---|---|
| EPI_ISL_1264908 | Randox Laboratories | Wellcome Sanger Institute for the COVID-19 Genomics UK (COG-UK) Consortium | Randox Laboratories et al |
| EPI_ISL_6240212 | Dutch COVID-19 response team | National Institute for Public Health and the Environment (RIVM) | Adam Meijer et al |
| EPI_ISL_4483411 | Dutch COVID-19 response team | Erasmus Medical Center | Bas Oude Munnink et al |
| EPI_ISL_6971572 | Furst Medical Laboratory | Norwegian Institute of Public Health, Department of Virology | Kathrine Stene-Johansen et al |
| EPI_ISL_3229878 | University Hospital of Northern Norway, Department for Microbiology and Infectious Disease Control | Norwegian Institute of Public Health, Department of Virology | Kathrine Stene-Johansen et al |
| EPI_ISL_6367729 | PAMM | Laboratory of Medical Microbiology and Pathology, PAMM | Jeroen van de Bovenkamp et al |
| EPI_ISL_4400379 | Dutch COVID-19 response team | National Institute for Public Health and the Environment (RIVM) | Adam Meijer et al |
| EPI_ISL_5910809 | North Shore Hospital | Institute of Environmental Science and Research (ESR) | Sarah Cockerton et al |
| EPI_ISL_1678865 | Department of Medical Microbiology & Infection prevention, Amsterdam University Medical Centers location AMC | Department of Medical Microbiology & Infection prevention, Amsterdam University Medical Centers location AMC | Matthijs Welkers et al |
| EPI_ISL_3151951 | Haukeland University Hospital, Dept. of Microbiology | Norwegian Institute of Public Health, Department of Virology | Kathrine Stene-Johansen et al |
| EPI_ISL_2093800 | Dutch COVID-19 response team | National Institute for Public Health and the Environment (RIVM) | Adam Meijer et al |
| EPI_ISL_1976058 | Regional Virus Laboratory, Belfast Health and Social Care Trust | COVID-19 Genomics UK (COG-UK) Consortium | Conall McCaughey et al |
| EPI_ISL_3080668 | Respiratory Virus Unit, Microbiology Services Colindale, Public Health England | COVID-19 Genomics UK (COG-UK) Consortium | PHE Covid Sequencing Team et al |
| EPI_ISL_2302683 | Dutch COVID-19 response team | National Institute for Public Health and the Environment (RIVM) | Adam Meijer et al |
| EPI_ISL_1014075 | Dutch COVID-19 response team | National Institute for Public Health and the Environment (RIVM) | Adam Meijer et al |
| EPI_ISL_2474517 | Dutch COVID-19 response team | National Institute for Public Health and the Environment (RIVM) | Adam Meijer et al |
| EPI_ISL_6853275 | Dutch COVID-19 response team | National Institute for Public Health and the Environment (RIVM) | Adam Meijer et al |
| EPI_ISL_6002877 | Irrua Specialist Teaching Hospital (ISTH) | Africa Centre for Excellence for Genomics of Infectious Diseases (ACEGID), Redeemer’s University | Olawoye et al |
| EPI_ISL_3057957 | Dutch COVID-19 response team | National Institute for Public Health and the Environment (RIVM) | Adam Meijer et al |
| EPI_ISL_1165368 | Dutch COVID-19 response team | National Institute for Public Health and the Environment (RIVM) | Adam Meijer et al |

| | | | |
|---|---|---|---|
| EPI_ISL_6696412 | Dept. of Medical Microbiology, Stavanger University Hospital, Helse Stavanger HF | Norwegian Institute of Public Health, Department of Virology | Kathrine Stene-Johansen et al |
| EPI_ISL_2610872 | Dutch COVID-19 response team | National Institute for Public Health and the Environment (RIVM) | Adam Meijer et al |
| EPI_ISL_1705522 | Dutch COVID-19 response team | National Institute for Public Health and the Environment (RIVM) | Adam Meijer et al |
| EPI_ISL_1317824 | Furst Medical Laboratory | Norwegian Institute of Public Health, Department of Virology | Kathrine Stene-Johansen et al |
| EPI_ISL_4345365 | Middlemore Hospital | Institute of Environmental Science and Research (ESR) | Sarah Cockerton et al |
| EPI_ISL_4634781 | Akershus University Hospital, Department for Microbiology and Infectious Disease Control | Norwegian Institute of Public Health, Department of Virology | Kathrine Stene-Johansen et al |
| EPI_ISL_1456746 | Dutch COVID-19 response team | National Institute for Public Health and the Environment (RIVM) | Adam Meijer et al |
| EPI_ISL_2425749 | Akershus University Hospital, Department for Microbiology and Infectious Disease Control | Norwegian Institute of Public Health, Department of Virology | Kathrine Stene-Johansen et al |
| EPI_ISL_3058027 | Dutch COVID-19 response team | National Institute for Public Health and the Environment (RIVM) | Adam Meijer et al |
| EPI_ISL_801437 | Dutch COVID-19 response team | Erasmus Medical Center | Bas Oude Munnink et al |
| EPI_ISL_461249 | Dutch COVID-19 response team | Erasmus Medical Center | Bas Oude Munnink et al |
| EPI_ISL_549039 | Medical Microbiology Unit, Department for Laboratory Medicine, Drammen Hospital, Vestre Viken Health Trust, | Norwegian Institute of Public Health, Department of Virology | Kathrine Stene-Johansen et al |
| EPI_ISL_1715374 | Nigerian Centre for Disease Control (NCDC) | African Centre of Excellence for Genomics of Infectious Diseases, Redeemer's University | Olawoye et al |
| EPI_ISL_547519 | Dutch COVID-19 response team | National Institute for Public Health and the Environment (RIVM) | Adam Meijer et al |
| EPI_ISL_6117374 | Eurofins-NMDL | Eurofins-NMDL | Anne Vogel et al |
| EPI_ISL_1054450 | Queens Medical Centre, Clinical Microbiology Department / DeepSeq Nottingham | COVID-19 Genomics UK (COG-UK) Consortium | Gemma Clark et al |
| EPI_ISL_4416474 | Microvida | Microvida | Suzan D. Pas et al |
| EPI_ISL_6002854 | Irrua Specialist Teaching Hospital (ISTH) | Africa Centre for Excellence for Genomics of Infectious Diseases (ACEGID), Redeemer's University | Olawoye et al |
| EPI_ISL_4743207 | UNILORIN MDRL | National Reference Laboratory, Nigeria Centre for Disease Control | Dr Ndodo Nnaemeka et al |
| EPI_ISL_1289031 | Dutch COVID-19 response team | National Institute for Public Health and the Environment (RIVM) | Adam Meijer et al |
| EPI_ISL_6235234 | Dutch COVID-19 response team | National Institute for Public Health and the Environment (RIVM) | Adam Meijer et al |
| EPI_ISL_2219982 | Dutch COVID-19 response team | National Institute for Public Health and the Environment (RIVM) | Adam Meijer et al |

| | | | |
|---|---|---|---|
| EPI_ISL_2420772 | Microvida | Microvida | Suzan D. Pas et al |
| EPI_ISL_1521001 | Dutch COVID-19 response team | National Institute for Public Health and the Environment (RIVM) | Adam Meijer et al |
| EPI_ISL_1251962 | Ostfold Hospital Trust - Kalnes, Centre for Laboratory Medicine, Section for gene technology and infection serology | Norwegian Institute of Public Health, Department of Virology | Kathrine Stene-Johansen et al |
| EPI_ISL_4400384 | Dutch COVID-19 response team | National Institute for Public Health and the Environment (RIVM) | Adam Meijer et al |
| EPI_ISL_4029526 | Sanquin B.V. | Sanquin B.V. | Karin van Leeuwen et al |
| EPI_ISL_2303243 | Dutch COVID-19 response team | National Institute for Public Health and the Environment (RIVM) | Adam Meijer et al |
| EPI_ISL_2259206 | Microvida | Microvida | Suzan D. Pas et al |
| EPI_ISL_6841607 | Dutch COVID-19 response team | National Institute for Public Health and the Environment (RIVM) | Adam Meijer et al |
| EPI_ISL_2220412 | Dutch COVID-19 response team | National Institute for Public Health and the Environment (RIVM) | Adam Meijer et al |
| EPI_ISL_3057049 | Dutch COVID-19 response team | National Institute for Public Health and the Environment (RIVM) | Adam Meijer et al |
| EPI_ISL_2357778 | Medical Microbiology Unit, Department for Laboratory Medicine, Drammen Hospital, Vestre Viken Health Trust, | Norwegian Institute of Public Health, Department of Virology | Kathrine Stene-Johansen et al |
| EPI_ISL_2216996 | Dutch COVID-19 response team | National Institute for Public Health and the Environment (RIVM) | Adam Meijer et al |
| EPI_ISL_7040235 | Oslo University Hospital, Department of Medical Microbiology | Norwegian Institute of Public Health, Department of Virology | Kathrine Stene-Johansen et al |
| EPI_ISL_6841619 | Dutch COVID-19 response team | National Institute for Public Health and the Environment (RIVM) | Adam Meijer et al |
| EPI_ISL_2673077 | Dutch COVID-19 response team | National Institute for Public Health and the Environment (RIVM) | Adam Meijer et al |
| EPI_ISL_4402061 | Dutch COVID-19 response team | National Institute for Public Health and the Environment (RIVM) | Adam Meijer et al |
| EPI_ISL_1456682 | Dutch COVID-19 response team | National Institute for Public Health and the Environment (RIVM) | Adam Meijer et al |
| EPI_ISL_2365539 | Dutch COVID-19 response team | Medical Microbiology, Maastricht University Medical Centre | Jozef Dingemans* et al |
| EPI_ISL_460947 | Dutch COVID-19 response team | Erasmus Medical Center | Bas Oude Munnink et al |
| EPI_ISL_4768494 | Ekiti State Emergency Operation Centre | Africa Centre for Excellence for Genomics of Infectious Diseases (ACEGID), Redeemer's University | Olawoye et al |
| EPI_ISL_4117130 | Oslo University Hospital, Department of Microbiology | Norwegian Institute of Public Health, Department of Virology | Mona Holberg-Petersen et al |
| EPI_ISL_7171276 | ARS Algarve - Laboratorio Laura Ayres | Instituto Nacional de Saude (INSA) | Borges et al |
| EPI_ISL_960742 | Germano de sousa | Instituto Gulbenkian de Ciencia | Maria Costa et al |
| EPI_ISL_6450872 | Central Public Health Laboratory | Microbiological Diagnostic Unit - Public Health Laboratory (MDU-PHL) | Kumbu et al |

| | | | |
|---|---|---|---|
| EPI_ISL_4417266 | Laboratorio de Referencia Nacional de Virus Respiratorios. Centro Nacional de Salud Publica. Instituto Nacional de Salud Peru. | Laboratorio de Referencia Nacional de Virus Respiratorios. Centro Nacional de Salud Publica. Instituto Nacional de Salud Peru. | Carlos Padilla Rojas et al |
| EPI_ISL_4121876 | Oslo University Hospital, Department of Microbiology | Norwegian Institute of Public Health, Department of Virology | Mona Holberg-Petersen et al |
| EPI_ISL_5544367 | Kairos Diagnostics Laboratory | Philippine Genome Center | Cynthia P. Saloma et al |
| EPI_ISL_6892631 | Germano de Sousa | Instituto Nacional de Saude (INSA) | Borges et al |
| EPI_ISL_6689546 | Laboratorio de Referencia Nacional de Virus Respiratorios. Centro Nacional de Salud Publica. Instituto Nacional de Salud Peru. | Laboratorio de Referencia Nacional de Virus Respiratorios. Centro Nacional de Salud Publica. Instituto Nacional de Salud Peru. | Carlos Padilla Rojas et al |
| EPI_ISL_6012810 | UNILABS | Instituto Nacional de Saude (INSA) | Borges et al |
| EPI_ISL_2290687 | WSSE w Gdańsku | 1. National Institute of Public Health - National Institute of Hygiene, Warsaw, Poland 2. Biobank Lab, University of Lodz 3. Laboratory of Respiratory Viruses, Teaching and Clinical Center of the Medical University of Lodz | Dominik Strapagiel et al |
| EPI_ISL_6475302 | WSSE w Gdansku | Wojewodzka Stacja Sanitarno-Epidemiologiczna w Olsztynie, Laboratorium Badan Epidemiologiczno-Klinicznych | Sylwia Krzetowska et al |
| EPI_ISL_1111157 | Laboratorio de Referencia Nacional de Virus Respiratorio. Instituto Nacional de Salud Perú | Laboratorio de Referencia Nacional de Enteropatógenos. Instituto Nacional de Salud del Perú | Ronnie Gavilan Chavez et al |
| EPI_ISL_1744972 | National Institute of Public Health - National Institute of Hygiene | 1. National Institute of Public Health - National Institute of Hygiene, Warsaw, Poland 2. Biobank Lab, University of Lodz 3. Laboratory of Respiratory Viruses, Teaching and Clinical Center of the Medical University of Lodz | Dominik Strapagiel et al |
| EPI_ISL_2694948 | H Fernando Fonseca | Instituto Nacional de Saude (INSA) | Borges et al |
| EPI_ISL_1915501 | Diagnostyka. Laboratoria Medyczne. | 1. ViroGenetics - BSL3 Laboratory of Virology, Małopolska Centre of Biotechnology, Jagiellonian University; 2. genXone SA, Research & Development Laboratory | Mazur-Panasiuk et al |
| EPI_ISL_5563070 | Baguio General Hospital Medical Center (BGHMC) | Philippine Genome Center | Cynthia P. Saloma et al |
| EPI_ISL_2377220 | Laboratorium Badawczo-Rozwojowe Vitagenum | 1. National Institute of Public Health - National Institute of Hygiene; 2. Eurofins Genomics Europe Sequencing GmbH | Wołkowicz Tomasz et al |

| | | | |
|---|---|---|---|
| EPI_ISL_2319107 | Diagnostyka. Laboratoria Medyczne. | 1. ViroGenetics - BSL3 Laboratory of Virology, Małopolska Centre of Biotechnology, Jagiellonian University; 2. genXone SA, Research & Development Laboratory | Mazur-Panasiuk et al |
| EPI_ISL_1111213 | Laboratorio de Referencia Nacional de Virus Respiratorio. Instituto Nacional de Salud Perú | Laboratorio de Referencia Nacional de Enteropatógenos. Instituto Nacional de Salud del Perú | Ronnie Gavilan Chavez et al |
| EPI_ISL_5935142 | Laboratorio de Referencia Nacional de Virus Respiratorios. Centro Nacional de Salud Publica. Instituto Nacional de Salud Peru. | Laboratorio de Referencia Nacional de Virus Respiratorios. Centro Nacional de Salud Publica. Instituto Nacional de Salud Peru. | Carlos Padilla Rojas et al |
| EPI_ISL_2156698 | Research Institute for Tropical Medicine, Inc. (RITM) | Philippine Genome Center | Francis A. Tablizo et al |
| EPI_ISL_1399860 | H Vila Franca Xira | Instituto Nacional de Saude (INSA) and Instituto Gulbenkian de Ciencia (IGC) | Borges et al |
| EPI_ISL_812446 | Laboratorio de Referencia Nacional de Virus Respiratorios, Instituto Nacional de Salud Peru | Laboratorio de Genómica Microbiana, Universidad Peruana Cayetano Heredia | Pablo Tsukayama et al |
| EPI_ISL_4126796 | Oslo University Hospital, Department of Microbiology | Norwegian Institute of Public Health, Department of Virology | Mona Holberg-Petersen et al |
| EPI_ISL_1111328 | Laboratorio de Referencia Nacional de Virus Respiratorio. Instituto Nacional de Salud Perú | Laboratorio de Referencia Nacional de Enteropatógenos. Instituto Nacional de Salud del Perú | Ronnie Gavilan Chavez et al |
| EPI_ISL_2694736 | Diagnostyka. Laboratoria Medyczne. | 1. ViroGenetics - BSL3 Laboratory of Virology, Małopolska Centre of Biotechnology, Jagiellonian University; 2. Diagtron Laboratoria Łukasz Rąbalski | Rabalski L. et al |
| EPI_ISL_1117089 | ARS Algarve - Laboratorio Laura Ayres | Instituto Nacional de Saude (INSA) | Borges et al |
| EPI_ISL_4119886 | Oslo University Hospital, Department of Microbiology | Norwegian Institute of Public Health, Department of Virology | Mona Holberg-Petersen et al |
| EPI_ISL_2155490 | Manila Doctors Hospital | Philippine Genome Center | Francis A. Tablizo et al |
| EPI_ISL_1340940 | 1. Glowny Inspektorat Sanitarny; 2. Diagnostyka. Laboratoria Medyczne. | 1. ViroGenetics - BSL3 Laboratory of Virology, Małopolska Centre of Biotechnology, Jagiellonian University; 2. Diagtron Laboratoria Lukasz Rabalski | Rabalski L. et al |
| EPI_ISL_2340203 | Germano de Sousa | Instituto Nacional de Saude (INSA) | Borges et al |
| EPI_ISL_5643598 | Laboratorium Epidemiologii WSSE w Szczecinie | Wojewodzka Stacja Sanitarno-Epidemiologiczna w Gorzowie Wielkopolskim | Elzbieta Justynska et al |

| | | | |
|---|---|---|---|
| EPI_ISL_4069805 | Laboratorium Analiz Lekarskich ALAB Rzeszow | Wojewodzka Stacja Sanitarno-Epidemiologiczna w Rzeszowie, Laboratorium Diagnostyki Medycznej | Marzena Baranowska et al |
| EPI_ISL_5852997 | ABC Algarve | Instituto Nacional de Saude (INSA) | Borges et al |
| EPI_ISL_5595768 | Laboratorio de Referencia Nacional de Virus Respiratorios. Centro Nacional de Salud Publica. Instituto Nacional de Salud Peru. | Laboratorio de Referencia Nacional de Virus Respiratorios. Centro Nacional de Salud Publica. Instituto Nacional de Salud Peru. | Carlos Padilla Rojas et al |
| EPI_ISL_6442929 | Szpital Wojewódzki im. Kardynała Stefana Wyszyńskiego w Łomży | 1. Academic Center for Pathomorphological and Genetic-Molecular Diagnostics ltd, Bialystok, Poland 2. National Institute of Public Health - National Institute of Hygiene, Warsaw, Poland | Radosław Charkiewicz et al |
| EPI_ISL_732268 | CHTMAD | Instituto Nacional de Saude (INSA) and Instituto Gulbenkian de Ciencia (IGC) | Borges et al |
| EPI_ISL_2188065 | Qualimed Hospital Sta. Rosa | Philippine Genome Center | Francis A. Tablizo et al |
| EPI_ISL_5147162 | Instituto de Medicina Tropical Alexander Von Humboldt, Sede Cusco | Laboratorio de Genómica Microbiana, Universidad Peruana Cayetano Heredia | Pablo Tsukayama et al |
| EPI_ISL_3307072 | H Fernando Fonseca | Instituto Nacional de Saude (INSA) | Borges et al |
| EPI_ISL_1590606 | Wojewodzka Stacja Sanitarno-Epidemiologiczna w Olsztynie, Laboratorium Badan Epidemiologiczno-Klinicznych | Wojewodzka Stacja Sanitarno-Epidemiologiczna w Olsztynie, Laboratorium Badan Epidemiologiczno-Klinicznych | Sylwia Krzetowska et al |
| EPI_ISL_2156223 | Lung Center of the Philippines (LCP) | Philippine Genome Center | Francis A. Tablizo et al |
| EPI_ISL_2340274 | UNILABS | Instituto Nacional de Saude (INSA) | Borges et al |
| EPI_ISL_6892620 | Germano de Sousa | Instituto Nacional de Saude (INSA) | Borges et al |
| EPI_ISL_4127257 | Oslo University Hospital, Department of Microbiology | Norwegian Institute of Public Health, Department of Virology | Mona Holberg-Petersen et al |
| EPI_ISL_2859764 | Baguio General Hospital Medical Center (BGHMC) | Philippine Genome Center | Francis A. Tablizo et al |
| EPI_ISL_6826885 | H Santarem | Instituto Nacional de Saude (INSA) | Borges et al |
| EPI_ISL_6110133 | LABORATORIUM "MEDREX" | 1. Academic Center for Pathomorphological and Genetic-Molecular Diagnostics ltd, Bialystok, Poland 2. National Institute of Public Health - National Institute of Hygiene, Warsaw, Poland | Radosław Charkiewicz et al |
| EPI_ISL_5796018 | Regionalne Centrum Krwiodawstwa i Krwiolecznictwa w Białymstoku Pracownia Diagnostyki Molekularnej wirusa SARS-CoV-2 | 1. Academic Center for Pathomorphological and Genetic-Molecular Diagnostics ltd, Bialystok, Poland 2. National Institute of Public Health - National Institute of Hygiene, Warsaw, Poland | Radosław Charkiewicz et al |

| | | | |
|---|---|---|---|
| EPI_ISL_2105260 | Cagayan Valley Medical Center Molecular Laboratory | Philippine Genome Center | Francis A. Tablizo et al |
| EPI_ISL_4555847 | UNILABS | Instituto Nacional de Saude (INSA) | Borges et al |
| EPI_ISL_2142257 | Vitalabo Bydgoszcz | 1. National Institute of Public Health - National Institute of Hygiene; 2. Eurofins Genomics Europe Sequencing GmbH | Wołkowicz Tomasz et al |
| EPI_ISL_7051368 | Krotoszyn | Wojewodzka Stacja Sanitarno-Epidemiologiczna w Lodzi, Oddzial Laboratoryjny Mikrobiologii i Parazytologii | Elian Terpo et al |
| EPI_ISL_6774425 | Uniwersyteckie Centrum Kliniczne | 10. Tricity SARS-CoV-2 sequencing consortium: University of Gdansk, Medical University of Gdansk, Vaxican Ltd., Invicta Ltd. 2. National Institute of Public Health - National Institute of Hygiene, Warsaw, Poland | Maciej Kosinski et al |
| EPI_ISL_2466729 | Jamil-ur-Rahman Center for Genome Research, Dr. Panjwani Center for Molecular Medicine and Drug Research | Jamil-ur-Rahman Center for Genome Research, Dr. Panjwani Center for Molecular Medicine and Drug Research | Irfan et al |
| EPI_ISL_1040985 | Hematology Laboratory, Section of Molecular Diagnostics, University Clinical Centre, Medical University of Gdansk | Laboratory of Recombinant Vaccines | Lukasz Rabalski et al |
| EPI_ISL_940953 | Centers for Disease Control and Prevention, Dengue Branch | Centers for Disease Control and Prevention, Dengue Branch | Gilberto A. Santiago et al |
| EPI_ISL_2015741 | Centrum Medyczne MEDYK Sp. z o.o. ,Sp.k. Zakład Diagnostyki Medycznej | 1. National Institute of Public Health - National Institute of Hygiene, Warsaw, Poland 2. Biobank Lab, University of Lodz 3. Laboratory of Respiratory Viruses, Teaching and Clinical Center of the Medical University of Lodz | Dominik Strapagiel et al |
| EPI_ISL_518848 | Oman-National Influenza Center | Biotechnology & OMICs Laboratory, Natural & Medical Sciences Research Center, University of Nizwa | Abdul Latif Khan et al |
| EPI_ISL_3980914 | ARS Algarve - Laboratorio Laura Ayres | Instituto Nacional de Saude (INSA) | Borges et al |
| EPI_ISL_3798752 | SEEBMO | Instituto Nacional de Saude (INSA) | Borges et al |
| EPI_ISL_2377531 | WSSE Katowice | 1. National Institute of Public Health - National Institute of Hygiene; 2. Eurofins Genomics Europe Sequencing GmbH | Wołkowicz Tomasz et al |
| EPI_ISL_6251532 | NZOZ Meditest Diagnostyka Medyczna | Wojewodzka Stacja Sanitarno-Epidemiologiczna w Gorzowie Wielkopolskim | Elzbieta Justynska et al |

| | | | |
|---|---|---|---|
| EPI_ISL_1164332 | Centro Hospitalar de Entre o Douro e Vouga (CHEDV) | Institute of Biomedicine (IBiMED), Universidade de Aveiro | Gabriela Moura et al |
| EPI_ISL_6892644 | Germano de Sousa | Instituto Nacional de Saude (INSA) | Borges et al |
| EPI_ISL_4303334 | Laboratorio Central de Salud Pública | Laboratorio Central de Salud Pública | Sandra Gonzalez et al |
| EPI_ISL_5547060 | Western Visayas State University - Molecular Diagnostic Laboratory | Philippine Genome Center | Cynthia P. Saloma et al |
| EPI_ISL_2810656 | UNILABS | Instituto Nacional de Saude (INSA) | Borges et al |
| EPI_ISL_4036779 | Laboratorio de Referencia Nacional de Virus Respiratorios. Centro Nacional de Salud Publica. Instituto Nacional de Salud Peru. | Laboratorio de Referencia Nacional de Virus Respiratorios. Centro Nacional de Salud Publica. Instituto Nacional de Salud Peru. | Carlos Padilla Rojas et al |
| EPI_ISL_3204891 | CHTMAD | Instituto Nacional de Saude (INSA) and Institute of Biomedicine (iBiMed), Universidade de Aveiro | Borges et al |
| EPI_ISL_6226681 | Oslo University Hospital, Department of Microbiology | Norwegian Institute of Public Health, Department of Virology | Mona Holberg-Petersen et al |
| EPI_ISL_4111964 | Labolatorium Biologi Molekularnej USK | Wojewodzka Stacja Sanitarno-Epidemiologiczna w Gorzowie Wielkopolskim | Elżbieta Justyńska et al |
| EPI_ISL_2895490 | UNILABS | Instituto Nacional de Saude (INSA) | Borges et al |
| EPI_ISL_4256331 | WSSE w Olsztynie | Wojewodzka Stacja Sanitarno-Epidemiologiczna w Olsztynie, Laboratorium Badan Epidemiologiczno-Klinicznych | Sylwia Krzetowska et al |
| EPI_ISL_5315137 | Gyncentrum | Gyncentrum | Emilia Morawiec et al |
| EPI_ISL_4125486 | Oslo University Hospital, Department of Microbiology | Norwegian Institute of Public Health, Department of Virology | Mona Holberg-Petersen et al |
| EPI_ISL_4520245 | Laboratorio de Referencia Nacional de Virus Respiratorios. Centro Nacional de Salud Publica. Instituto Nacional de Salud Peru. | Laboratorio de Referencia Nacional de Virus Respiratorios. Centro Nacional de Salud Publica. Instituto Nacional de Salud Peru. | Carlos Padilla Rojas et al |
| EPI_ISL_1118127 | Dept. of Microbiology and Infection Control, Akershus University Hospital HF | Dept. of Microbiology and Infection Control, Akershus University Hospital HF | Hege Vangstein Aamot et al |
| EPI_ISL_5033478 | H Fernando Fonseca | Instituto Nacional de Saude (INSA) and Instituto Gulbenkian de Ciencia (IGC) | Borges et al |
| EPI_ISL_3023624 | Laboratorio de Referencial Nacional de Virus Respiratorios | Laboratorio de Referencial Nacional de Virus Respiratorios | Carlos Padilla Rojas et al |
| EPI_ISL_1499131 | 1. Glowny Inspektorat Sanitarny. 2. Diagnostyka. Laboratoria Medyczne. | 1. ViroGenetics - BSL3 Laboratory of Virology, Małopolska Centre of Biotechnology, Jagiellonian University; 2. genXone SA, Research & Development Laboratory | Mazur-Panasiuk et al |

| | | | |
|---|---|---|---|
| EPI_ISL_2921388 | Laboratorio de Referencial Nacional de Virus Respiratorios | Laboratorio de Referencial Nacional de Virus Respiratorios | Carlos Padilla Rojas et al |
| EPI_ISL_1317691 | Unilabs Laboratory Medicine | Norwegian Institute of Public Health, Department of Virology | Kathrine Stene-Johansen et al |
| EPI_ISL_6944767 | Zakład Mikrobiologii, WSZZ Kielce | 1. Academic Center for Pathomorphological and Genetic-Molecular Diagnostics ltd, Bialystok, Poland 2. National Institute of Public Health - National Institute of Hygiene, Warsaw, Poland | Radosław Charkiewicz et al |
| EPI_ISL_3133621 | Diagnostyka. Laboratoria Medyczne. | 1. ViroGenetics - BSL3 Laboratory of Virology, Małopolska Centre of Biotechnology, Jagiellonian University; 2. genXone SA, Research & Development Laboratory | Mazur-Panasiuk et al |
| EPI_ISL_4948938 | Central Public Health Laboratory | Microbiological Diagnostic Unit - Public Health Laboratory (MDU-PHL) | Kumbu et al |
| EPI_ISL_2695084 | H Vila Franca Xira | Instituto Nacional de Saude (INSA) | Borges et al |
| EPI_ISL_2444779 | IICS-UNA | IICS-UNA | Magaly Martinez et al |
| EPI_ISL_4543891 | WSSE w Gdansku | Wojewodzka Stacja Sanitarno-Epidemiologiczna w Olsztynie, Laboratorium Badan Epidemiologiczno-Klinicznych | Sylwia Krzetowska et al |
| EPI_ISL_511348 | SYNLAB | Instituto Nacional de Saude (INSA) | Borges et al |
| EPI_ISL_4742313 | Maria Reyna Xavier University Hospital | Philippine Genome Center | Cynthia P. Saloma et al |
| EPI_ISL_5544554 | Dr. Jose N. Rodriguez Memorial Hospital and Sanitarium (Tala) | Philippine Genome Center | Cynthia P. Saloma et al |
| EPI_ISL_5924649 | Lab. Biologii molekularnej USK | Wojewodzka Stacja Sanitarno-Epidemiologiczna w Gorzowie Wielkopolskim | Elzbieta Justynska et al |
| EPI_ISL_5334369 | Pathogenic Microorganisms Variability Laboratory | Pathogenic Microorganisms Variability Laboratory | Andrey Pochtovyy et al |
| EPI_ISL_2238601 | Lighthouse Lab in Glasgow | Wellcome Sanger Institute for the COVID-19 Genomics UK (COG-UK) Consortium | Harper VanSteenhouse et al |
| EPI_ISL_6642115 | Molecular diagnostic laboratory of Federal Budget Institution of Science "Central Research Institute of Epidemiology" of The Federal Service on Customers' Rights Protection and Human Well-being Surveillance | Group of Genomics and Postgenomic Technologies of Central Research Institute of Epidemiology | Samoilov A.E. et al |
| EPI_ISL_1413379 | Lighthouse Lab in Glasgow | Wellcome Sanger Institute for the COVID-19 Genomics UK (COG-UK) Consortium | Harper VanSteenhouse et al |

| | | | |
|---|---|---|---|
| EPI_ISL_1413259 | Lighthouse Lab in Glasgow | Wellcome Sanger Institute for the COVID-19 Genomics UK (COG-UK) Consortium | Harper VanSteenhouse et al |
| EPI_ISL_3154354 | REUNILAB | UMR PIMIT | Dr David A Wilkinson et al |
| EPI_ISL_1317070 | Lighthouse Lab in Glasgow | Wellcome Sanger Institute for the COVID-19 Genomics UK (COG-UK) Consortium | Harper VanSteenhouse et al |
| EPI_ISL_5465978 | Virology Department, Royal Infirmary of Edinburgh, NHS Lothian / School of Biological Sciences, University of Edinburgh | COVID-19 Genomics UK (COG-UK) Consortium | McHugh M et al |
| EPI_ISL_2354367 | Lighthouse Lab in Glasgow | Wellcome Sanger Institute for the COVID-19 Genomics UK (COG-UK) Consortium | Harper VanSteenhouse et al |
| EPI_ISL_2734411 | Virology Department, Royal Infirmary of Edinburgh, NHS Lothian / School of Biological Sciences, University of Edinburgh | COVID-19 Genomics UK (COG-UK) Consortium | McHugh M et al |
| EPI_ISL_4769272 | West of Scotland Specialist Virology Centre, NHSGGC / MRC-University of Glasgow Centre for Virus Research | COVID-19 Genomics UK (COG-UK) Consortium | Ana da Silva Filipe et al |
| EPI_ISL_2622293 | West of Scotland Specialist Virology Centre, NHSGGC / MRC-University of Glasgow Centre for Virus Research | COVID-19 Genomics UK (COG-UK) Consortium | Ana da Silva Filipe et al |
| EPI_ISL_2318110 | Lighthouse Lab in Glasgow | Wellcome Sanger Institute for the COVID-19 Genomics UK (COG-UK) Consortium | Harper VanSteenhouse et al |
| EPI_ISL_733097 | HELIX LLC | WHO National Influenza Centre Russian Federation | Andrey Komissarov et al |
| EPI_ISL_6132463 | Ministry of Public Health / Hamad Medical Corporation | Biomedical Research Center (BRC), Qatar University / Qatar Genome Project (QGP) | BRC: Fatiha M. Benslimane et al |
| EPI_ISL_3140323 | Lighthouse Lab in Glasgow | Wellcome Sanger Institute for the COVID-19 Genomics UK (COG-UK) Consortium | Harper VanSteenhouse et al |
| EPI_ISL_6778797 | CHU - Saint-Denis | Laboratoire de virologie, CNR arbovirus Associé, Chu de la Réunion | Etienne Frumence et al |
| EPI_ISL_2436860 | Lighthouse Lab in Glasgow | Wellcome Sanger Institute for the COVID-19 Genomics UK (COG-UK) Consortium | Harper VanSteenhouse et al |
| EPI_ISL_3109934 | Lighthouse Lab in Glasgow | Wellcome Sanger Institute for the COVID-19 Genomics UK (COG-UK) Consortium | Harper VanSteenhouse et al |
| EPI_ISL_2728732 | Lighthouse Lab in Glasgow | Wellcome Sanger Institute for the COVID-19 Genomics UK (COG-UK) Consortium | Harper VanSteenhouse et al |

| | | | |
|---|---|---|---|
| EPI_ISL_919264 | West of Scotland Specialist Virology Centre, NHSGGC / MRC-University of Glasgow Centre for Virus Research | COVID-19 Genomics UK (COG-UK) Consortium | Ana da Silva Filipe et al |
| EPI_ISL_6862897 | CERBALLIANCE | UMR PIMIT | David A Wilkinson et al |
| EPI_ISL_6016561 | Virology Department, Royal Infirmary of Edinburgh, NHS Lothian / School of Biological Sciences, University of Edinburgh | COVID-19 Genomics UK (COG-UK) Consortium | McHugh M et al |
| EPI_ISL_5917822 | Medical Genetics Laboratory, Regional Centre of Medical Genetics, Emergency County Hospital Craiova | Medical Genetics Laboratory, Regional Centre of Medical Genetics, Emergency County Hospital Craiova | Anca-Lelia (Riza) Costache et al |
| EPI_ISL_2438497 | Lighthouse Lab in Glasgow | Wellcome Sanger Institute for the COVID-19 Genomics UK (COG-UK) Consortium | Harper VanSteenhouse et al |
| EPI_ISL_2406882 | Lighthouse Lab in Glasgow | Wellcome Sanger Institute for the COVID-19 Genomics UK (COG-UK) Consortium | Harper VanSteenhouse et al |
| EPI_ISL_1257742 | Lighthouse Lab in Glasgow | Wellcome Sanger Institute for the COVID-19 Genomics UK (COG-UK) Consortium | Harper VanSteenhouse et al |
| EPI_ISL_1742083 | Lighthouse Lab in Glasgow | Wellcome Sanger Institute for the COVID-19 Genomics UK (COG-UK) Consortium | Harper VanSteenhouse et al |
| EPI_ISL_2702178 | Lighthouse Lab in Glasgow | Wellcome Sanger Institute for the COVID-19 Genomics UK (COG-UK) Consortium | Harper VanSteenhouse et al |
| EPI_ISL_2807192 | Lighthouse Lab in Glasgow | Wellcome Sanger Institute for the COVID-19 Genomics UK (COG-UK) Consortium | Harper VanSteenhouse et al |
| EPI_ISL_453188 | Virology Department, Royal Infirmary of Edinburgh, NHS Lothian / School of Biological Sciences, University of Edinburgh / Institute of Genetics and Molecular Medicine, University of Edinburgh | COVID-19 Genomics UK (COG-UK) Consortium | McHugh M et al |
| EPI_ISL_2974643 | Lighthouse Lab in Glasgow | Wellcome Sanger Institute for the COVID-19 Genomics UK (COG-UK) Consortium | Harper VanSteenhouse et al |
| EPI_ISL_594670 | West of Scotland Specialist Virology Centre, NHSGGC / MRC-University of Glasgow Centre for Virus Research | COVID-19 Genomics UK (COG-UK) Consortium | Ana da Silva Filipe et al |
| EPI_ISL_2730945 | Lighthouse Lab in Glasgow | Wellcome Sanger Institute for the COVID-19 Genomics UK (COG-UK) Consortium | Harper VanSteenhouse et al |

| | | | |
|---|---|---|---|
| EPI_ISL_5465889 | Virology Department, Royal Infirmary of Edinburgh, NHS Lothian / School of Biological Sciences, University of Edinburgh | COVID-19 Genomics UK (COG-UK) Consortium | McHugh M et al |
| EPI_ISL_4184504 | Department of Public Health Iasi | National Institute of Infectious Diseases-Prof. Dr. Matei Bals Molecular Diagnostics Laboratory | Corina Casangiu et al |
| EPI_ISL_2851222 | Lighthouse Lab in Glasgow | Wellcome Sanger Institute for the COVID-19 Genomics UK (COG-UK) Consortium | Harper VanSteenhouse et al |
| EPI_ISL_2964509 | N.F. Gamaleya Research Center for Epidemiology and Microbiology | WHO National Influenza Centre Russian Federation | Andrey Komissarov et al |
| EPI_ISL_2275406 | Lighthouse Lab in Glasgow | Wellcome Sanger Institute for the COVID-19 Genomics UK (COG-UK) Consortium | Harper VanSteenhouse et al |
| EPI_ISL_6980876 | Cerballiance, Reunion | UMR PIMIT | David A Wilkinson et al |
| EPI_ISL_1293875 | Lighthouse Lab in Glasgow | Wellcome Sanger Institute for the COVID-19 Genomics UK (COG-UK) Consortium | Harper VanSteenhouse et al |
| EPI_ISL_2899818 | N.F. Gamaleya Research Center for Epidemiology and Microbiology | WHO National Influenza Centre Russian Federation | Andrey Komissarov et al |
| EPI_ISL_2438250 | Lighthouse Lab in Glasgow | Wellcome Sanger Institute for the COVID-19 Genomics UK (COG-UK) Consortium | Harper VanSteenhouse et al |
| EPI_ISL_2385198 | WHO National Influenza Centre Russian Federation | WHO National Influenza Centre Russian Federation | Andrey Komissarov et al |
| EPI_ISL_2814937 | Lighthouse Lab in Glasgow | Wellcome Sanger Institute for the COVID-19 Genomics UK (COG-UK) Consortium | Harper VanSteenhouse et al |
| EPI_ISL_1506418 | Lighthouse Lab in Glasgow | Wellcome Sanger Institute for the COVID-19 Genomics UK (COG-UK) Consortium | Harper VanSteenhouse et al |
| EPI_ISL_3021370 | Clinical Emergency County Hospital, Craiova | Cantacuzino Institute Virology | Carmen Cherciu et al |
| EPI_ISL_5241201 | Cantacuzino National Military-Medical Institute, Viral Respiratory Infections Laboratory | Cantacuzino National Military-Medical Institute, Viral Respiratory Infections Laboratory | Luiza Ustea et al |
| EPI_ISL_560316 | UMMC-Health | WHO National Influenza Centre Russian Federation | Andrey Komissarov et al |
| EPI_ISL_1714022 | Ministry of Public Health / Hamad Medical Corporation | Weill Cornell Medical College - Qatar (WCM-Q), Genomics Core Laboratory / Qatar Genome Project (QGP) | WCMQ: Ayeda A. Ahmed et al |
| EPI_ISL_1277657 | Lighthouse Lab in Glasgow | Wellcome Sanger Institute for the COVID-19 Genomics UK (COG-UK) Consortium | Harper VanSteenhouse et al |

| | | | |
|---|---|---|---|
| EPI_ISL_6901304 | Molecular diagnostic laboratory of Federal Budget Institution of Science "Central Research Institute of Epidemiology" of The Federal Service on Customers' Rights Protection and Human Well-being Surveillance | Group of Genomics and Postgenomic Technologies of Central Research Institute of Epidemiology | Samoilov A.E. et al |
| EPI_ISL_3856802 | Department of Public Health Alba | National Institute of Infectious Diseases-Prof. Dr. Matei Bals Molecular Diagnostics Laboratory | Corina Casangiu et al |
| EPI_ISL_6195815 | Medical Genetics Laboratory, Regional Centre of Medical Genetics, Emergency County Hospital Craiova | Medical Genetics Laboratory, Regional Centre of Medical Genetics, Emergency County Hospital Craiova | Anca-Lelia (Riza) Costache et al |
| EPI_ISL_430072 | WHO National Influenza Centre Russian Federation | WHO National Influenza Centre Russian Federation | Andrey Komissarov et al |
| EPI_ISL_3367136 | Lighthouse Lab in Glasgow | Wellcome Sanger Institute for the COVID-19 Genomics UK (COG-UK) Consortium | Harper VanSteenhouse et al |
| EPI_ISL_918060 | Lighthouse Lab in Alderley Park | Wellcome Sanger Institute for the COVID-19 Genomics UK (COG-UK) Consortium | Jacquelyn Wynn et al |
| EPI_ISL_3919606 | Molecular diagnostic laboratory of Federal Budget Institution of Science "Central Research Institute of Epidemiology" of The Federal Service on Customers' Rights Protection and Human Well-being Surveillance | Group of Genomics and Postgenomic Technologies of Central Research Institute of Epidemiology | Samoilov A.E. et al |
| EPI_ISL_2852300 | Lighthouse Lab in Glasgow | Wellcome Sanger Institute for the COVID-19 Genomics UK (COG-UK) Consortium | Harper VanSteenhouse et al |
| EPI_ISL_707790 | Rwanda National Reference Laboratory | Rwanda National Reference Laboratory | Enatha Mukantwari et al |
| EPI_ISL_2852462 | Lighthouse Lab in Glasgow | Wellcome Sanger Institute for the COVID-19 Genomics UK (COG-UK) Consortium | Harper VanSteenhouse et al |
| EPI_ISL_2652196 | Cantacuzino National Military-Medical Institute, Viral Respiratory Infections Laboratory | Cantacuzino Institute Virology | Carmen Cherciu et al |
| EPI_ISL_2086633 | Kazan Federal University, Academic Consortium | FEDERAL RESEARCH AND CLINICAL CENTER OF PHYSICAL-CHEMICAL MEDICINE OF FEDERAL MEDICAL BIOLOGICAL AGENCY, Academic Consortium | Vladislav Babenko et al |
| EPI_ISL_1635098 | Lighthouse Lab in Glasgow | Wellcome Sanger Institute for the COVID-19 Genomics UK (COG-UK) Consortium | Harper VanSteenhouse et al |

| | | | |
|---|---|---|---|
| EPI_ISL_2626281 | Molecular diagnostic laboratory of Federal Budget Institution of Science "Central Research Institute of Epidemiology" of The Federal Service on Customers' Rights Protection and Human Well-being Surveillance | Group of Genomics and Postgenomic Technologies of Central Research Institute of Epidemiology | Samoilov A.E. et al |
| EPI_ISL_1316344 | Lighthouse Lab in Glasgow | Wellcome Sanger Institute for the COVID-19 Genomics UK (COG-UK) Consortium | Harper VanSteenhouse et al |
| EPI_ISL_4604566 | REUNILAB DEUX CANONS | CNR Virus des Infections Respiratoires - France SUD | Antonin Bal et al |
| EPI_ISL_888678 | Laboratory for Respiratory Viruses, Cantacuzino National Military-Medical Institute for Research and Development | Cantacuzino Institute Virology | Luiza Ustea et al |
| EPI_ISL_3141514 | Lighthouse Lab in Glasgow | Wellcome Sanger Institute for the COVID-19 Genomics UK (COG-UK) Consortium | Harper VanSteenhouse et al |
| EPI_ISL_1544973 | Lighthouse Lab in Glasgow | Wellcome Sanger Institute for the COVID-19 Genomics UK (COG-UK) Consortium | Harper VanSteenhouse et al |
| EPI_ISL_2937458 | Lighthouse Lab in Glasgow | Wellcome Sanger Institute for the COVID-19 Genomics UK (COG-UK) Consortium | Harper VanSteenhouse et al |
| EPI_ISL_2437994 | Lighthouse Lab in Glasgow | Wellcome Sanger Institute for the COVID-19 Genomics UK (COG-UK) Consortium | Harper VanSteenhouse et al |
| EPI_ISL_1714176 | Ministry of Public Health / Hamad Medical Corporation | Weill Cornell Medical College - Qatar (WCM-Q), Genomics Core Laboratory / Qatar Genome Project (QGP) | WCMQ: Ayeda A. Ahmed et al |
| EPI_ISL_3077287 | West of Scotland Specialist Virology Centre, NHSGGC / MRC-University of Glasgow Centre for Virus Research | COVID-19 Genomics UK (COG-UK) Consortium | Ana da Silva Filipe et al |
| EPI_ISL_3153653 | CERBALLIANCE | UMR PIMIT | Dr David A Wilkinson et al |
| EPI_ISL_2608152 | Lighthouse Lab in Glasgow | Wellcome Sanger Institute for the COVID-19 Genomics UK (COG-UK) Consortium | Harper VanSteenhouse et al |
| EPI_ISL_2235824 | Lighthouse Lab in Glasgow | Wellcome Sanger Institute for the COVID-19 Genomics UK (COG-UK) Consortium | Harper VanSteenhouse et al |
| EPI_ISL_5241324 | Cantacuzino National Military-Medical Institute, Viral Respiratory Infections Laboratory | Cantacuzino National Military-Medical Institute, Viral Respiratory Infections Laboratory | Luiza Ustea et al |
| EPI_ISL_1538354 | Lighthouse Lab in Glasgow | Wellcome Sanger Institute for the COVID-19 Genomics UK (COG-UK) Consortium | Harper VanSteenhouse et al |

| | | | |
|---|---|---|---|
| EPI_ISL_2734301 | West of Scotland Specialist Virology Centre, NHSGGC / MRC-University of Glasgow Centre for Virus Research | COVID-19 Genomics UK (COG-UK) Consortium | Ana da Silva Filipe et al |
| EPI_ISL_1277662 | Lighthouse Lab in Glasgow | Wellcome Sanger Institute for the COVID-19 Genomics UK (COG-UK) Consortium | Harper VanSteenhouse et al |
| EPI_ISL_2651930 | Lighthouse Lab in Glasgow | Wellcome Sanger Institute for the COVID-19 Genomics UK (COG-UK) Consortium | Harper VanSteenhouse et al |
| EPI_ISL_2899893 | N.F. Gamaleya Research Center for Epidemiology and Microbiology | WHO National Influenza Centre Russian Federation | Andrey Komissarov et al |
| EPI_ISL_3141204 | Lighthouse Lab in Glasgow | Wellcome Sanger Institute for the COVID-19 Genomics UK (COG-UK) Consortium | Harper VanSteenhouse et al |
| EPI_ISL_3077257 | West of Scotland Specialist Virology Centre, NHSGGC / MRC-University of Glasgow Centre for Virus Research | COVID-19 Genomics UK (COG-UK) Consortium | Ana da Silva Filipe et al |
| EPI_ISL_1411012 | Lighthouse Lab in Glasgow | Wellcome Sanger Institute for the COVID-19 Genomics UK (COG-UK) Consortium | Harper VanSteenhouse et al |
| EPI_ISL_2353865 | Lighthouse Lab in Glasgow | Wellcome Sanger Institute for the COVID-19 Genomics UK (COG-UK) Consortium | Harper VanSteenhouse et al |
| EPI_ISL_2354434 | Lighthouse Lab in Glasgow | Wellcome Sanger Institute for the COVID-19 Genomics UK (COG-UK) Consortium | Harper VanSteenhouse et al |
| EPI_ISL_2997442 | Lighthouse Lab in Glasgow | Wellcome Sanger Institute for the COVID-19 Genomics UK (COG-UK) Consortium | Harper VanSteenhouse et al |
| EPI_ISL_1412051 | Lighthouse Lab in Glasgow | Wellcome Sanger Institute for the COVID-19 Genomics UK (COG-UK) Consortium | Harper VanSteenhouse et al |
| EPI_ISL_6268972 | West of Scotland Specialist Virology Centre, NHSGGC / MRC-University of Glasgow Centre for Virus Research | COVID-19 Genomics UK (COG-UK) Consortium | Ana da Silva Filipe et al |
| EPI_ISL_2683237 | Lighthouse Lab in Glasgow | Wellcome Sanger Institute for the COVID-19 Genomics UK (COG-UK) Consortium | Harper VanSteenhouse et al |
| EPI_ISL_6047005 | Lighthouse Lab in Glasgow | Wellcome Sanger Institute for the COVID-19 Genomics UK (COG-UK) Consortium | Harper VanSteenhouse et al |
| EPI_ISL_710697 | Lighthouse Lab in Glasgow | Wellcome Sanger Institute for the COVID-19 Genomics UK (COG-UK) Consortium | Harper VanSteenhouse et al |
| EPI_ISL_5570871 | Public Health Authority of the Slovak Republic | Public Health Authority of the Slovak Republic | Dusan Loderer et al |

| | | | |
|---|---|---|---|
| EPI_ISL_7148719 | Lighthouse Lab in Glasgow | Wellcome Sanger Institute for the COVID-19 Genomics UK (COG-UK) Consortium | Harper VanSteenhouse et al |
| EPI_ISL_3811755 | Lighthouse Lab in Glasgow | Wellcome Sanger Institute for the COVID-19 Genomics UK (COG-UK) Consortium | Harper VanSteenhouse et al |
| EPI_ISL_5270740 | Lighthouse Lab in Glasgow | Wellcome Sanger Institute for the COVID-19 Genomics UK (COG-UK) Consortium | Harper VanSteenhouse et al |
| EPI_ISL_5114594 | Public Health Authority of the Slovak Republic | Laboratory of Genomics and Bioinformatics, Comenius University Science Park | Tatiana Sedláčková et al |
| EPI_ISL_4407461 | Lighthouse Lab in Glasgow | Wellcome Sanger Institute for the COVID-19 Genomics UK (COG-UK) Consortium | Harper VanSteenhouse et al |
| EPI_ISL_638973 | Oxford Viromics, NDM, University of Oxford; Oxford University Hospitals; Basingstoke and North Hampshire Hospital | COVID-19 Genomics UK (COG-UK) Consortium | Tanya Golubchik et al |
| EPI_ISL_1266650 | National Laboratory for Health, Environment and Food, OMM, Kranj | CISLD (Clinical Institute of Special Laboratory Diagnostics), University Children's Hospital, University Medical Center Ljubljana | Jernej Kovač et al |
| EPI_ISL_5246923 | Lighthouse Lab in Glasgow | Wellcome Sanger Institute for the COVID-19 Genomics UK (COG-UK) Consortium | Harper VanSteenhouse et al |
| EPI_ISL_3262001 | National Public Health Laboratory, National Centre for Infectious Diseases | National Public Health Laboratory, National Centre for Infectious Diseases | Tze Minn Mak et al |
| EPI_ISL_2657903 | Public Health Authority of the Slovak Republic | Laboratory of Genomics and Bioinformatics, Comenius University Science Park | Tatiana Sedláčková et al |
| EPI_ISL_6034057 | Institute of Virology, Biomedical Research Center of the Slovak Academy of Sciences, Bratislava | Faculty of Mathematics, Physics and Informatics, Comenius University, Bratislava | Viktoria Cabanova et al |
| EPI_ISL_1647680 | Public Health Authority of the Slovak Republic | Laboratory of Genomics and Bioinformatics, Comenius University Science Park | Tatiana Sedláčková et al |
| EPI_ISL_600614 | Lighthouse Lab in Glasgow | Wellcome Sanger Institute for the COVID-19 Genomics UK (COG-UK) consortium | Harper VanSteenhouse et al |
| EPI_ISL_2474474 | Dutch COVID-19 response team | National Institute for Public Health and the Environment (RIVM) | Adam Meijer et al |
| EPI_ISL_1112148 | National Laboratory for Health, Environment and Food, OMM, Celje | CISLD (Clinical Institute of Special Laboratory Diagnostics), University Children's Hospital, University Medical Center Ljubljana | Jernej Kovač et al |
| EPI_ISL_6436040 | Lighthouse Lab in Glasgow | Wellcome Sanger Institute for the COVID-19 Genomics UK (COG-UK) Consortium | Harper VanSteenhouse et al |

| | | | |
|---|---|---|---|
| EPI_ISL_7148080 | Lighthouse Lab in Glasgow | Wellcome Sanger Institute for the COVID-19 Genomics UK (COG-UK) Consortium | Harper VanSteenhouse et al |
| EPI_ISL_3726660 | Lighthouse Lab in Glasgow | Wellcome Sanger Institute for the COVID-19 Genomics UK (COG-UK) Consortium | Harper VanSteenhouse et al |
| EPI_ISL_4480841 | Lighthouse Lab in Glasgow | Wellcome Sanger Institute for the COVID-19 Genomics UK (COG-UK) Consortium | Harper VanSteenhouse et al |
| EPI_ISL_5836441 | Lighthouse Lab in Glasgow | Wellcome Sanger Institute for the COVID-19 Genomics UK (COG-UK) Consortium | Harper VanSteenhouse et al |
| EPI_ISL_6436048 | Lighthouse Lab in Glasgow | Wellcome Sanger Institute for the COVID-19 Genomics UK (COG-UK) Consortium | Harper VanSteenhouse et al |
| EPI_ISL_4330177 | Lighthouse Lab in Glasgow | Wellcome Sanger Institute for the COVID-19 Genomics UK (COG-UK) Consortium | Harper VanSteenhouse et al |
| EPI_ISL_6926451 | Lighthouse Lab in Glasgow | Wellcome Sanger Institute for the COVID-19 Genomics UK (COG-UK) Consortium | Harper VanSteenhouse et al |
| EPI_ISL_6615681 | Lighthouse Lab in Glasgow | Wellcome Sanger Institute for the COVID-19 Genomics UK (COG-UK) Consortium | Harper VanSteenhouse et al |
| EPI_ISL_6046958 | Lighthouse Lab in Glasgow | Wellcome Sanger Institute for the COVID-19 Genomics UK (COG-UK) Consortium | Harper VanSteenhouse et al |
| EPI_ISL_7035922 | Lighthouse Lab in Glasgow | Wellcome Sanger Institute for the COVID-19 Genomics UK (COG-UK) Consortium | Harper VanSteenhouse et al |
| EPI_ISL_4329828 | Lighthouse Lab in Glasgow | Wellcome Sanger Institute for the COVID-19 Genomics UK (COG-UK) Consortium | Harper VanSteenhouse et al |
| EPI_ISL_7145412 | Lighthouse Lab in Glasgow | Wellcome Sanger Institute for the COVID-19 Genomics UK (COG-UK) Consortium | Harper VanSteenhouse et al |
| EPI_ISL_6867571 | Lighthouse Lab in Glasgow | Wellcome Sanger Institute for the COVID-19 Genomics UK (COG-UK) Consortium | Harper VanSteenhouse et al |
| EPI_ISL_6929949 | The Institute of Virology, Vaccines and Sera “Torlak” | Center for forensic and applied molecular genetics, Faculty of Biology, University of Belgrade | D.Keckarević et al |
| EPI_ISL_4147072 | Lighthouse Lab in Glasgow | Wellcome Sanger Institute for the COVID-19 Genomics UK (COG-UK) Consortium | Harper VanSteenhouse et al |
| EPI_ISL_6877342 | Lighthouse Lab in Glasgow | Wellcome Sanger Institute for the COVID-19 Genomics UK (COG-UK) Consortium | Harper VanSteenhouse et al |
| EPI_ISL_648116 | Department of Laboratory Medicine, Tan Tock Seng Hospital | Department of Laboratory Medicine, Tan Tock Seng Hospital | Chen YYC et al |

| | | | |
|---|---|---|---|
| EPI_ISL_4408928 | Lighthouse Lab in Glasgow | Wellcome Sanger Institute for the COVID-19 Genomics UK (COG-UK) Consortium | Harper VanSteenhouse et al |
| EPI_ISL_4069041 | Lighthouse Lab in Glasgow | Wellcome Sanger Institute for the COVID-19 Genomics UK (COG-UK) Consortium | Harper VanSteenhouse et al |
| EPI_ISL_406593 | Shenzhen Key Laboratory of Pathogen and Immunity, National Clinical Research Center for Infectious Disease, Shenzhen Third People's Hospital | Shenzhen Key Laboratory of Pathogen and Immunity, National Clinical Research Center for Infectious Disease, Shenzhen Third People's Hospital | Yang Yang et al |
| EPI_ISL_600837 | Lighthouse Lab in Glasgow | Wellcome Sanger Institute for the COVID-19 Genomics UK (COG-UK) consortium | Harper VanSteenhouse et al |
| EPI_ISL_4146534 | Lighthouse Lab in Glasgow | Wellcome Sanger Institute for the COVID-19 Genomics UK (COG-UK) Consortium | Harper VanSteenhouse et al |
| EPI_ISL_4019830 | Lighthouse Lab in Glasgow | Wellcome Sanger Institute for the COVID-19 Genomics UK (COG-UK) Consortium | Harper VanSteenhouse et al |
| EPI_ISL_6157169 | Lighthouse Lab in Glasgow | Wellcome Sanger Institute for the COVID-19 Genomics UK (COG-UK) Consortium | Harper VanSteenhouse et al |
| EPI_ISL_6626066 | Lighthouse Lab in Glasgow | Wellcome Sanger Institute for the COVID-19 Genomics UK (COG-UK) Consortium | Harper VanSteenhouse et al |
| EPI_ISL_2093176 | Dutch COVID-19 response team | National Institute for Public Health and the Environment (RIVM) | Adam Meijer et al |
| EPI_ISL_4019424 | Lighthouse Lab in Glasgow | Wellcome Sanger Institute for the COVID-19 Genomics UK (COG-UK) Consortium | Harper VanSteenhouse et al |
| EPI_ISL_6048619 | Lighthouse Lab in Glasgow | Wellcome Sanger Institute for the COVID-19 Genomics UK (COG-UK) Consortium | Harper VanSteenhouse et al |
| EPI_ISL_6950277 | National Laboratory for Health, Environment and Food, OMM, Maribor | NLZOH (National Laboratory for Health, Environment and Food) / CISLD (Clinical Institute of Special Laboratory Diagnostics), University Children's Hospital, University Medical Center Ljubljana | Sandra Janezic et al |
| EPI_ISL_4148210 | Lighthouse Lab in Glasgow | Wellcome Sanger Institute for the COVID-19 Genomics UK (COG-UK) Consortium | Harper VanSteenhouse et al |
| EPI_ISL_6615329 | Lighthouse Lab in Glasgow | Wellcome Sanger Institute for the COVID-19 Genomics UK (COG-UK) Consortium | Harper VanSteenhouse et al |
| EPI_ISL_3381938 | Lighthouse Lab in Glasgow | Wellcome Sanger Institute for the COVID-19 Genomics UK (COG-UK) Consortium | Harper VanSteenhouse et al |

| | | | |
|---|---|---|---|
| EPI_ISL_4068523 | Lighthouse Lab in Glasgow | Wellcome Sanger Institute for the COVID-19 Genomics UK (COG-UK) Consortium | Harper VanSteenhouse et al |
| EPI_ISL_2887026 | Institute of Microbiology and Immunology, Faculty of Medicine, University of Ljubljana | Institute of Microbiology and Immunology, Faculty of Medicine, University of Ljubljana | Alen Suljič et al |
| EPI_ISL_5242902 | Lighthouse Lab in Glasgow | Wellcome Sanger Institute for the COVID-19 Genomics UK (COG-UK) Consortium | Harper VanSteenhouse et al |
| EPI_ISL_644010 | Lighthouse Lab in Glasgow | Wellcome Sanger Institute for the COVID-19 Genomics UK (COG-UK) Consortium | Harper VanSteenhouse et al |
| EPI_ISL_4685252 | Lighthouse Lab in Glasgow | Wellcome Sanger Institute for the COVID-19 Genomics UK (COG-UK) Consortium | Harper VanSteenhouse et al |
| EPI_ISL_601928 | Lighthouse Lab in Glasgow | Wellcome Sanger Institute for the COVID-19 Genomics UK (COG-UK) consortium | Harper VanSteenhouse et al |
| EPI_ISL_3582873 | Lighthouse Lab in Glasgow | Wellcome Sanger Institute for the COVID-19 Genomics UK (COG-UK) Consortium | Harper VanSteenhouse et al |
| EPI_ISL_4147233 | Lighthouse Lab in Glasgow | Wellcome Sanger Institute for the COVID-19 Genomics UK (COG-UK) Consortium | Harper VanSteenhouse et al |
| EPI_ISL_428846 | National Public Health Laboratory, National Centre for Infectious Diseases | National Public Health Laboratory, National Centre for Infectious Diseases | Mak TM et al |
| EPI_ISL_4146901 | Lighthouse Lab in Glasgow | Wellcome Sanger Institute for the COVID-19 Genomics UK (COG-UK) Consortium | Harper VanSteenhouse et al |
| EPI_ISL_7035793 | Lighthouse Lab in Glasgow | Wellcome Sanger Institute for the COVID-19 Genomics UK (COG-UK) Consortium | Harper VanSteenhouse et al |
| EPI_ISL_3968961 | Lighthouse Lab in Glasgow | Wellcome Sanger Institute for the COVID-19 Genomics UK (COG-UK) Consortium | Harper VanSteenhouse et al |
| EPI_ISL_7031141 | Lighthouse Lab in Glasgow | Wellcome Sanger Institute for the COVID-19 Genomics UK (COG-UK) Consortium | Harper VanSteenhouse et al |
| EPI_ISL_6040755 | Lighthouse Lab in Glasgow | Wellcome Sanger Institute for the COVID-19 Genomics UK (COG-UK) Consortium | Harper VanSteenhouse et al |
| EPI_ISL_6626036 | Lighthouse Lab in Glasgow | Wellcome Sanger Institute for the COVID-19 Genomics UK (COG-UK) Consortium | Harper VanSteenhouse et al |
| EPI_ISL_4767521 | Lighthouse Lab in Glasgow | Wellcome Sanger Institute for the COVID-19 Genomics UK (COG-UK) Consortium | Harper VanSteenhouse et al |
| EPI_ISL_6613540 | Lighthouse Lab in Glasgow | Wellcome Sanger Institute for the COVID-19 Genomics UK (COG-UK) Consortium | Harper VanSteenhouse et al |

| | | | |
|---|---|---|---|
| EPI_ISL_4408294 | Lighthouse Lab in Glasgow | Wellcome Sanger Institute for the COVID-19 Genomics UK (COG-UK) Consortium | Harper VanSteenhouse et al |
| EPI_ISL_909737 | Institute of Biocides and Medical Ecology, Belgrade, Serbia | Virology Department, Institute of microbiology and immunology, Faculty of Medicine University of Blegrade | Banko Ana et al |
| EPI_ISL_5423474 | Lighthouse Lab in Glasgow | Wellcome Sanger Institute for the COVID-19 Genomics UK (COG-UK) Consortium | Harper VanSteenhouse et al |
| EPI_ISL_6278702 | Institute of Virology, Biomedical Research Center of the Slovak Academy of Sciences, Bratislava | Faculty of Mathematics, Physics and Informatics, Comenius University, Bratislava | Viktoria Cabanova et al |
| EPI_ISL_4146991 | Lighthouse Lab in Glasgow | Wellcome Sanger Institute for the COVID-19 Genomics UK (COG-UK) Consortium | Harper VanSteenhouse et al |
| EPI_ISL_4476859 | Lighthouse Lab in Glasgow | Wellcome Sanger Institute for the COVID-19 Genomics UK (COG-UK) Consortium | Harper VanSteenhouse et al |
| EPI_ISL_6044897 | Lighthouse Lab in Glasgow | Wellcome Sanger Institute for the COVID-19 Genomics UK (COG-UK) Consortium | Harper VanSteenhouse et al |
| EPI_ISL_6156840 | Lighthouse Lab in Glasgow | Wellcome Sanger Institute for the COVID-19 Genomics UK (COG-UK) Consortium | Harper VanSteenhouse et al |
| EPI_ISL_6224680 | Lighthouse Lab in Glasgow | Wellcome Sanger Institute for the COVID-19 Genomics UK (COG-UK) Consortium | Harper VanSteenhouse et al |
| EPI_ISL_7029582 | Lighthouse Lab in Glasgow | Wellcome Sanger Institute for the COVID-19 Genomics UK (COG-UK) Consortium | Harper VanSteenhouse et al |
| EPI_ISL_4404712 | Lighthouse Lab in Glasgow | Wellcome Sanger Institute for the COVID-19 Genomics UK (COG-UK) Consortium | Harper VanSteenhouse et al |
| EPI_ISL_4972668 | Lighthouse Lab in Glasgow | Wellcome Sanger Institute for the COVID-19 Genomics UK (COG-UK) Consortium | Harper VanSteenhouse et al |
| EPI_ISL_2688051 | Public Health Authority of the Slovak Republic | Laboratory of Genomics and Bioinformatics, Comenius University Science Park | Tatiana Sedláčková et al |
| EPI_ISL_6038983 | Lighthouse Lab in Glasgow | Wellcome Sanger Institute for the COVID-19 Genomics UK (COG-UK) Consortium | Harper VanSteenhouse et al |
| EPI_ISL_6616878 | Lighthouse Lab in Glasgow | Wellcome Sanger Institute for the COVID-19 Genomics UK (COG-UK) Consortium | Harper VanSteenhouse et al |
| EPI_ISL_7027785 | Lighthouse Lab in Glasgow | Wellcome Sanger Institute for the COVID-19 Genomics UK (COG-UK) Consortium | Harper VanSteenhouse et al |

| | | | |
|---|---|---|---|
| EPI_ISL_6437003 | Public Health Authority of the Slovak Republic | Laboratory of Genomics and Bioinformatics, Comenius University Science Park | Tatiana Sedláčková et al |
| EPI_ISL_5855976 | Institute of Microbiology and Immunology, Faculty of Medicine, University of Ljubljana | Institute of Microbiology and Immunology, Faculty of Medicine, University of Ljubljana | Alen Suljič et al |
| EPI_ISL_2982234 | National Public Health Laboratory, National Centre for Infectious Diseases | National Public Health Laboratory, National Centre for Infectious Diseases | Tze Minn Mak et al |
| EPI_ISL_5242895 | Lighthouse Lab in Glasgow | Wellcome Sanger Institute for the COVID-19 Genomics UK (COG-UK) Consortium | Harper VanSteenhouse et al |
| EPI_ISL_6870030 | Lighthouse Lab in Glasgow | Wellcome Sanger Institute for the COVID-19 Genomics UK (COG-UK) Consortium | Harper VanSteenhouse et al |
| EPI_ISL_4079305 | National Public Health Laboratory, National Centre for Infectious Diseases | National Public Health Laboratory, National Centre for Infectious Diseases | Katherine Ching et al |
| EPI_ISL_4767175 | Lighthouse Lab in Glasgow | Wellcome Sanger Institute for the COVID-19 Genomics UK (COG-UK) Consortium | Harper VanSteenhouse et al |
| EPI_ISL_4020201 | Lighthouse Lab in Glasgow | Wellcome Sanger Institute for the COVID-19 Genomics UK (COG-UK) Consortium | Harper VanSteenhouse et al |
| EPI_ISL_5788227 | Lighthouse Lab in Glasgow | Wellcome Sanger Institute for the COVID-19 Genomics UK (COG-UK) Consortium | Harper VanSteenhouse et al |
| EPI_ISL_6043732 | Lighthouse Lab in Glasgow | Wellcome Sanger Institute for the COVID-19 Genomics UK (COG-UK) Consortium | Harper VanSteenhouse et al |
| EPI_ISL_1547879 | Institute of Virology, Biomedical Research Center of the Slovak Academy of Sciences, Bratislava | Faculty of Natural Sciences, Comenius University, Bratislava | Kristina Borsova et al |
| EPI_ISL_6613958 | Lighthouse Lab in Glasgow | Wellcome Sanger Institute for the COVID-19 Genomics UK (COG-UK) Consortium | Harper VanSteenhouse et al |
| EPI_ISL_3366628 | Lighthouse Lab in Glasgow | Wellcome Sanger Institute for the COVID-19 Genomics UK (COG-UK) Consortium | Harper VanSteenhouse et al |
| EPI_ISL_1526755 | Public Health Authority of the Slovak Republic | Laboratory of Genomics and Bioinformatics, Comenius University Science Park | Tatiana Sedláčková et al |
| EPI_ISL_5836037 | Lighthouse Lab in Glasgow | Wellcome Sanger Institute for the COVID-19 Genomics UK (COG-UK) Consortium | Harper VanSteenhouse et al |
| EPI_ISL_4147978 | Lighthouse Lab in Glasgow | Wellcome Sanger Institute for the COVID-19 Genomics UK (COG-UK) Consortium | Harper VanSteenhouse et al |

| | | | |
|---|---|---|---|
| EPI_ISL_568361 | Lighthouse Lab in Glasgow | Wellcome Sanger Institute for the COVID-19 Genomics UK (COG-UK) consortium | Harper VanSteenhouse et al |
| EPI_ISL_6927893 | Lighthouse Lab in Glasgow | Wellcome Sanger Institute for the COVID-19 Genomics UK (COG-UK) Consortium | Harper VanSteenhouse et al |
| EPI_ISL_2965745 | Public Health Authority of the Slovak Republic | Public Health Authority of the Slovak Republic | Barbora Kotvasová et al |
| EPI_ISL_2897054 | Microbiology Department, Laboratori Clínic Metropolitana Nord. Hospital Universitari Germans Trias i Pujol | Can Ruti SARS-CoV-2 Sequencing Hub (HUGTiP/IrsiCaixa/IGTP) | Marc Noguera-Julian et al |
| EPI_ISL_2001934 | National Laboratory for Health, Environment and Food, OMM, Celje | NLZOH (National Laboratory for Health, Environment and Food) / CISLD (Clinical Institute of Special Laboratory Diagnostics), University Children's Hospital, University Medical Center Ljubljana | Sandra Janezic et al |
| EPI_ISL_2273400 | National Laboratory for Health, Environment and Food, OMM, Koper | NLZOH (National Laboratory for Health, Environment and Food) / CISLD (Clinical Institute of Special Laboratory Diagnostics), University Children's Hospital, University Medical Center Ljubljana | Sandra Janezic et al |
| EPI_ISL_1123376 | Hospital Valle del Nalón | Laboratorio de Virología HUCA | Sandoval M et al |
| EPI_ISL_6746558 | Division of Emerging Infectious Diseases, Bureau of Infectious Diseases Diagnosis Control, Korea Disease Control and Prevention Agency | Division of Emerging Infectious Diseases, Bureau of Infectious Diseases Diagnosis Control, Korea Disease Control and Prevention Agency | Ae Kyung Park et al |
| EPI_ISL_2958095 | SARS-CoV-2 Sequencing Castilla y Leon-Spain Consortium | SARS-CoV-2 Sequencing Castilla y Leon-Spain Consortium | Antonio Orduña-Domingo et al |
| EPI_ISL_6699746 | National Health Laboratory Services, Virology, Charlotte Maxeke Johannesburg hospital, Parktown, Johannesburg, Gauteng | CERI, Centre for Epidemic Response and Innvoation, Stellenbosch University and KRISP, KZN Research Innovation and Sequencing Platform, UKZN. | Kathleen Subramoney et al |
| EPI_ISL_3452128 | Division of Emerging Infectious Diseases, Bureau of Infectious Diseases Diagnosis Control, Korea Disease Control and Prevention Agency | Division of Emerging Infectious Diseases, Bureau of Infectious Diseases Diagnosis Control, Korea Disease Control and Prevention Agency | Ae Kyung Park et al |
| EPI_ISL_6782055 | National Health Laboratory Services, Virology, Charlotte Maxeke Johannesburg hospital, Parktown, Johannesburg, Gauteng | CERI, Centre for Epidemic Response and Innvoation, Stellenbosch University and KRISP, KZN Research Innovation and Sequencing Platform, UKZN. | Kathleen Subramoney et al |

| | | | |
|---|---|---|---|
| EPI_ISL_6795844 | ZARV/NHLS, Department Medical Virology, University of Pretoria | CERI, Centre for Epidemic Response and Innvoation, Stellenbosch University and KRISP, KZN Research Innovation and Sequencing Platform, UKZN. | Amy Strydom et al |
| EPI_ISL_6795838 | ZARV/NHLS, Department Medical Virology, University of Pretoria | CERI, Centre for Epidemic Response and Innvoation, Stellenbosch University and KRISP, KZN Research Innovation and Sequencing Platform, UKZN. | Amy Strydom et al |
| EPI_ISL_6900143 | HELEN JOSEPH LABORATORY | National Institute for Communicable Diseases of the National Health Laboratory Service | Amoako DG et al |
| EPI_ISL_4272300 | SARS-CoV-2 Sequencing Castilla y Leon-Spain Consortium | SARS-CoV-2 Sequencing Castilla y Leon-Spain Consortium | Antonio Orduña-Domingo et al |
| EPI_ISL_498093 | NHLS-IALCH | KRISP, KZN Research Innovation and Sequencing Platform | Giandhari J et al |
| EPI_ISL_5600176 | National Laboratory for Health, Environment and Food, OMM, Kranj | NLZOH (National Laboratory for Health, Environment and Food) / CISLD (Clinical Institute of Special Laboratory Diagnostics), University Children's Hospital, University Medical Center Ljubljana | Sandra Janezic et al |
| EPI_ISL_6699728 | National Health Laboratory Services, Virology, Charlotte Maxeke Johannesburg hospital, Parktown, Johannesburg, Gauteng | CERI, Centre for Epidemic Response and Innvoation, Stellenbosch University and KRISP, KZN Research Innovation and Sequencing Platform, UKZN. | Kathleen Subramoney et al |
| EPI_ISL_6795841 | ZARV/NHLS, Department Medical Virology, University of Pretoria | CERI, Centre for Epidemic Response and Innvoation, Stellenbosch University and KRISP, KZN Research Innovation and Sequencing Platform, UKZN. | Amy Strydom et al |
| EPI_ISL_6782056 | National Health Laboratory Services, Virology, Charlotte Maxeke Johannesburg hospital, Parktown, Johannesburg, Gauteng | CERI, Centre for Epidemic Response and Innvoation, Stellenbosch University and KRISP, KZN Research Innovation and Sequencing Platform, UKZN. | Kathleen Subramoney et al |
| EPI_ISL_2380042 | Microbiology Department, Laboratori Clínic Metropolitana Nord. Hospital Universitari Germans Trias i Pujol. | Can Ruti SARS-CoV-2 Sequencing Hub (HUGTiP/IrsiCaixa/IGTP) | Marc Noguera-Julian et al |
| EPI_ISL_5007199 | Division of Emerging Infectious Diseases, Bureau of Infectious Diseases Diagnosis Control, Korea Disease Control and Prevention Agency | Division of Emerging Infectious Diseases, Bureau of Infectious Diseases Diagnosis Control, Korea Disease Control and Prevention Agency | Ae Kyung Park et al |
| EPI_ISL_2582733 | Vermaak & Partners Laboratory | National Institute for Communicable Diseases of the National Health Laboratory Service | Amoako DG et al |

| | | | |
|---|---|---|---|
| EPI_ISL_6699730 | National Health Laboratory Services, Virology, Charlotte Maxeke Johannesburg hospital, Parktown, Johannesburg, Gauteng | CERI, Centre for Epidemic Response and Innvoation, Stellenbosch University and KRISP, KZN Research Innovation and Sequencing Platform, UKZN. | Kathleen Subramoney et al |
| EPI_ISL_6699739 | National Health Laboratory Services, Virology, Charlotte Maxeke Johannesburg hospital, Parktown, Johannesburg, Gauteng | CERI, Centre for Epidemic Response and Innvoation, Stellenbosch University and KRISP, KZN Research Innovation and Sequencing Platform, UKZN. | Kathleen Subramoney et al |
| EPI_ISL_2886621 | Institute of Microbiology and Immunology, Faculty of Medicine, University of Ljubljana | Institute of Microbiology and Immunology, Faculty of Medicine, University of Ljubljana | Alen Suljič et al |
| EPI_ISL_4572818 | NHLS Charlotte Maxeke Johannesburg Academic Hospital and the University of the Witwatersrand | KRISP, KZN Research Innovation and Sequencing Platform | Florette Treurnicht et al |
| EPI_ISL_6829558 | National Health Laboratory Service, Kwazulu-Natal, South Africa | KRISP, KZn Research Innovation and Sequencing Platform | Nokukhanya M et al |
| EPI_ISL_6825390 | ZARV/NHLS, Department Medical Virology, University of Pretoria | CERI, Centre for Epidemic Response and Innvoation, Stellenbosch University and KRISP, KZN Research Innovation and Sequencing Platform, UKZN. | Amy Strydom et al |
| EPI_ISL_6647956 | LANCET LABORATORY | National Institute for Communicable Diseases of the National Health Laboratory Service | Amoako DG et al |
| EPI_ISL_2955458 | NHLS_VIRO Eastern Cape | KRISP, KZn Research Innovation and Sequencing Platform | Laguda-Akingba O et al |
| EPI_ISL_2516430 | Hospital Universitari Bellvitge | Microbiology Department | Sara Marti et al |
| EPI_ISL_1402148 | Department of Public Health Microbiology Ljubljana, National Laboratory for Health, Environment and Food | Department of Public Health Microbiology Ljubljana, National Laboratory for Health, Environment and Food | Tom Koritnik et al |
| EPI_ISL_3587881 | SARS-CoV-2 Sequencing Castilla y Leon-Spain Consortium | SARS-CoV-2 Sequencing Castilla y Leon-Spain Consortium | Antonio Orduña-Domingo et al |
| EPI_ISL_7154542 | Hospital General La Mancha Centro | Hospital General Universitario de Ciudad Real | José Martínez-Alarcón et al |
| EPI_ISL_3666731 | Division of Medical Virology, National Health Laboratory Service (NHLS), Tygerberg Hospital / Stellenbosch University | Division of Medical Virology, National Health Laboratory Service (NHLS), Tygerberg Hospital / Stellenbosch University | Susan Engelbrecht et al |
| EPI_ISL_1785995 | Servicio de Microbiología, Hospital Miguel Servet, Zaragoza | SeqCOVID-SPAIN consortium/IBV(CSIC) | Antonio Rezusta López et al |

| | | | |
|---|---|---|---|
| EPI_ISL_6699729 | National Health Laboratory Services, Virology, Charlotte Maxeke Johannesburg hospital, Parktown, Johannesburg, Gauteng | CERI, Centre for Epidemic Response and Innvoation, Stellenbosch University and KRISP, KZN Research Innovation and Sequencing Platform, UKZN. | Kathleen Subramoney et al |
| EPI_ISL_6699744 | National Health Laboratory Services, Virology, Charlotte Maxeke Johannesburg hospital, Parktown, Johannesburg, Gauteng | CERI, Centre for Epidemic Response and Innvoation, Stellenbosch University and KRISP, KZN Research Innovation and Sequencing Platform, UKZN. | Kathleen Subramoney et al |
| EPI_ISL_6782080 | National Health Laboratory Services, Virology, Charlotte Maxeke Johannesburg hospital, Parktown, Johannesburg, Gauteng | CERI, Centre for Epidemic Response and Innvoation, Stellenbosch University and KRISP, KZN Research Innovation and Sequencing Platform, UKZN. | Kathleen Subramoney et al |
| EPI_ISL_2649302 | SARS-CoV-2 Sequencing Castilla y Leon-Spain Consortium | SARS-CoV-2 Sequencing Castilla y Leon-Spain Consortium | Antonio Orduña-Domingo et al |
| EPI_ISL_3729144 | NHLS Charlotte Maxeke Johannesburg Hospital | KRISP, KZN Research Innovation and Sequencing Platform | Kathleen Subramoney et al |
| EPI_ISL_3161764 | NHLS_VIRO | KRISP, KZN Research Innovation and Sequencing Platform | Giandhari Jennifer et al |
| EPI_ISL_6782071 | National Health Laboratory Services, Virology, Charlotte Maxeke Johannesburg hospital, Parktown, Johannesburg, Gauteng | CERI, Centre for Epidemic Response and Innvoation, Stellenbosch University and KRISP, KZN Research Innovation and Sequencing Platform, UKZN. | Kathleen Subramoney et al |
| EPI_ISL_2494445 | NHLS_VIRO | KRISP, KZn Research Innovation and Sequencing Platform | Baillie Vicky et al |
| EPI_ISL_6795842 | ZARV/NHLS, Department Medical Virology, University of Pretoria | CERI, Centre for Epidemic Response and Innvoation, Stellenbosch University and KRISP, KZN Research Innovation and Sequencing Platform, UKZN. | Amy Strydom et al |
| EPI_ISL_6900141 | HELEN JOSEPH LABORATORY | National Institute for Communicable Diseases of the National Health Laboratory Service | Amoako DG et al |
| EPI_ISL_3261247 | Hospital Universitari de Bellvitge | Microbiology Department | Sara Marti et al |
| EPI_ISL_6647961 | LANCET LABORATORY | National Institute for Communicable Diseases of the National Health Laboratory Service | Amoako DG et al |
| EPI_ISL_6699731 | National Health Laboratory Services, Virology, Charlotte Maxeke Johannesburg hospital, Parktown, Johannesburg, Gauteng | CERI, Centre for Epidemic Response and Innvoation, Stellenbosch University and KRISP, KZN Research Innovation and Sequencing Platform, UKZN. | Kathleen Subramoney et al |
| EPI_ISL_6795188 | National Health Laboratory Services, Virology | CERI, Centre for Epidemic Response and Innvoation, Stellenbosch University and KRISP, KZN Research Innovation and Sequencing Platform, UKZN. | Kathleen Subramoney et al |

| | | | |
|---|---|---|---|
| EPI_ISL_6959993 | Division of Emerging Infectious Diseases, Bureau of Infectious Diseases Diagnosis Control, Korea Disease Control and Prevention Agency | Division of Emerging Infectious Diseases, Bureau of Infectious Diseases Diagnosis Control, Korea Disease Control and Prevention Agency | Ae Kyung Park et al |
| EPI_ISL_2710546 | Division of Emerging Infectious Diseases, Bureau of Infectious Diseases Diagnosis Control, Korea Disease Control and Prevention Agency | Division of Emerging Infectious Diseases, Bureau of Infectious Diseases Diagnosis Control, Korea Disease Control and Prevention Agency | Jeong-Ah Kim et al |
| EPI_ISL_1788151 | Universidad de León | SeqCOVID-SPAIN consortium/IBV(CSIC) | Ana Carvajal et al |
| EPI_ISL_2694624 | TAMBO MEMORIAL | National Institute for Communicable Diseases of the National Health Laboratory Service | Amoako DG et al |
| EPI_ISL_4253274 | National Laboratory for Health, Environment and Food, OMM, Koper | NLZOH (National Laboratory for Health, Environment and Food) / CISLD (Clinical Institute of Special Laboratory Diagnostics), University Children's Hospital, University Medical Center Ljubljana | Sandra Janezic et al |
| EPI_ISL_6795839 | ZARV/NHLS, Department Medical Virology, University of Pretoria | CERI, Centre for Epidemic Response and Innvoation, Stellenbosch University and KRISP, KZN Research Innovation and Sequencing Platform, UKZN. | Amy Strydom et al |
| EPI_ISL_6647960 | LANCET LABORATORY | National Institute for Communicable Diseases of the National Health Laboratory Service | Amoako DG et al |
| EPI_ISL_6699732 | National Health Laboratory Services, Virology, Charlotte Maxeke Johannesburg hospital, Parktown, Johannesburg, Gauteng | CERI, Centre for Epidemic Response and Innvoation, Stellenbosch University and KRISP, KZN Research Innovation and Sequencing Platform, UKZN. | Kathleen Subramoney et al |
| EPI_ISL_1486522 | Microbiology Department, Laboratori Clínic Metropolitana Nord. Hospital Universitari Germans Trias i Pujol. | Can Ruti SARS-CoV-2 Sequencing Hub (HUGTiP/IrsiCaixa/IGTP) | Marc Noguera-Julian et al |
| EPI_ISL_6959926 | Division of Emerging Infectious Diseases, Bureau of Infectious Diseases Diagnosis Control, Korea Disease Control and Prevention Agency | Division of Emerging Infectious Diseases, Bureau of Infectious Diseases Diagnosis Control, Korea Disease Control and Prevention Agency | Ae Kyung Park et al |
| EPI_ISL_6782043 | National Health Laboratory Services, Virology, Charlotte Maxeke Johannesburg hospital, Parktown, Johannesburg, Gauteng | CERI, Centre for Epidemic Response and Innvoation, Stellenbosch University and KRISP, KZN Research Innovation and Sequencing Platform, UKZN. | Kathleen Subramoney et al |

| | | | |
|---|---|---|---|
| EPI_ISL_4270872 | Institute of Microbiology and Immunology, Faculty of Medicine, University of Ljubljana | Institute of Microbiology and Immunology, Faculty of Medicine, University of Ljubljana | Alen Suljič et al |
| EPI_ISL_6900142 | HELEN JOSEPH LABORATORY | National Institute for Communicable Diseases of the National Health Laboratory Service | Amoako DG et al |
| EPI_ISL_5913542 | Microbiologia CATLAB | Can Ruti SARS-CoV-2 Sequencing Hub (HUGTiP/IrsiCaixa/IGTP) | Marc Noguera-Julian et al |
| EPI_ISL_6782092 | National Health Laboratory Services, Virology, Charlotte Maxeke Johannesburg hospital, Parktown, Johannesburg, Gauteng | CERI, Centre for Epidemic Response and Innvoation, Stellenbosch University and KRISP, KZN Research Innovation and Sequencing Platform, UKZN. | Kathleen Subramoney et al |
| EPI_ISL_660187 | NHLS-IALCH | KRISP, KZN Research Innovation and Sequencing Platform | Giandhari J et al |
| EPI_ISL_1719819 | Microbiology Department, Laboratori Clínic Metropolitana Nord. Hospital Universitari Germans Trias i Pujol. | Can Ruti SARS-CoV-2 Sequencing Hub (HUGTiP/IrsiCaixa/IGTP) | Marc Noguera-Julian et al |
| EPI_ISL_6782084 | National Health Laboratory Services, Virology, Charlotte Maxeke Johannesburg hospital, Parktown, Johannesburg, Gauteng | CERI, Centre for Epidemic Response and Innvoation, Stellenbosch University and KRISP, KZN Research Innovation and Sequencing Platform, UKZN. | Kathleen Subramoney et al |
| EPI_ISL_747383 | Division of Emerging Infectious Diseases, Bureau of Infectious Diseases Diagnosis Control, Korea Disease Control and Prevention Agency | Division of Emerging Infectious Diseases, Bureau of Infectious Diseases Diagnosis Control, Korea Disease Control and Prevention Agency | Ae Kyung Park et al |
| EPI_ISL_6699741 | National Health Laboratory Services, Virology, Charlotte Maxeke Johannesburg hospital, Parktown, Johannesburg, Gauteng | CERI, Centre for Epidemic Response and Innvoation, Stellenbosch University and KRISP, KZN Research Innovation and Sequencing Platform, UKZN. | Kathleen Subramoney et al |
| EPI_ISL_6782090 | National Health Laboratory Services, Virology, Charlotte Maxeke Johannesburg hospital, Parktown, Johannesburg, Gauteng | CERI, Centre for Epidemic Response and Innvoation, Stellenbosch University and KRISP, KZN Research Innovation and Sequencing Platform, UKZN. | Kathleen Subramoney et al |
| EPI_ISL_3026583 | Division of Emerging Infectious Diseases, Bureau of Infectious Diseases Diagnosis Control, Korea Disease Control and Prevention Agency | Division of Emerging Infectious Diseases, Bureau of Infectious Diseases Diagnosis Control, Korea Disease Control and Prevention Agency | Jeong-Ah Kim et al |
| EPI_ISL_6699745 | National Health Laboratory Services, Virology, Charlotte Maxeke Johannesburg hospital, Parktown, Johannesburg, Gauteng | CERI, Centre for Epidemic Response and Innvoation, Stellenbosch University and KRISP, KZN Research Innovation and Sequencing Platform, UKZN. | Kathleen Subramoney et al |

| | | | |
|---|---|---|---|
| EPI_ISL_6699743 | National Health Laboratory Services, Virology, Charlotte Maxeke Johannesburg hospital, Parktown, Johannesburg, Gauteng | CERI, Centre for Epidemic Response and Innvoation, Stellenbosch University and KRISP, KZN Research Innovation and Sequencing Platform, UKZN. | Kathleen Subramoney et al |
| EPI_ISL_6699735 | National Health Laboratory Services, Virology, Charlotte Maxeke Johannesburg hospital, Parktown, Johannesburg, Gauteng | CERI, Centre for Epidemic Response and Innvoation, Stellenbosch University and KRISP, KZN Research Innovation and Sequencing Platform, UKZN. | Kathleen Subramoney et al |
| EPI_ISL_6647958 | LANCET LABORATORY | National Institute for Communicable Diseases of the National Health Laboratory Service | Amoako DG et al |
| EPI_ISL_6795840 | ZARV/NHLS, Department Medical Virology, University of Pretoria | CERI, Centre for Epidemic Response and Innvoation, Stellenbosch University and KRISP, KZN Research Innovation and Sequencing Platform, UKZN. | Amy Strydom et al |
| EPI_ISL_7154531 | Hospital Virgen De Altagracia | Hospital General Universitario de Ciudad Real | José Martínez-Alarcón et al |
| EPI_ISL_6647962 | LANCET LABORATORY | National Institute for Communicable Diseases of the National Health Laboratory Service | Amoako DG et al |
| EPI_ISL_6647959 | LANCET LABORATORY | National Institute for Communicable Diseases of the National Health Laboratory Service | Amoako DG et al |
| EPI_ISL_5064406 | Institute of Microbiology and Immunology, Faculty of Medicine, University of Ljubljana | Institute of Microbiology and Immunology, Faculty of Medicine, University of Ljubljana | Alen Suljič et al |
| EPI_ISL_6751106 | Division of Emerging Infectious Diseases, Bureau of Infectious Diseases Diagnosis Control, Korea Disease Control and Prevention Agency | Division of Emerging Infectious Diseases, Bureau of Infectious Diseases Diagnosis Control, Korea Disease Control and Prevention Agency | Ae Kyung Park et al |
| EPI_ISL_6795189 | National Health Laboratory Services, Virology | CERI, Centre for Epidemic Response and Innvoation, Stellenbosch University and KRISP, KZN Research Innovation and Sequencing Platform, UKZN. | Kathleen Subramoney et al |
| EPI_ISL_862642 | Consejería de Sanidad y Asuntos Sociales | Instituto de Salud Carlos III | Iglesias-Caballero et al |
| EPI_ISL_6699734 | National Health Laboratory Services, Virology, Charlotte Maxeke Johannesburg hospital, Parktown, Johannesburg, Gauteng | CERI, Centre for Epidemic Response and Innvoation, Stellenbosch University and KRISP, KZN Research Innovation and Sequencing Platform, UKZN. | Kathleen Subramoney et al |
| EPI_ISL_6699736 | National Health Laboratory Services, Virology, Charlotte Maxeke Johannesburg hospital, Parktown, Johannesburg, Gauteng | CERI, Centre for Epidemic Response and Innvoation, Stellenbosch University and KRISP, KZN Research Innovation and Sequencing Platform, UKZN. | Kathleen Subramoney et al |

| | | | |
|---|---|---|---|
| EPI_ISL_6782079 | National Health Laboratory Services, Virology, Charlotte Maxeke Johannesburg hospital, Parktown, Johannesburg, Gauteng | CERI, Centre for Epidemic Response and Innvoation, Stellenbosch University and KRISP, KZN Research Innovation and Sequencing Platform, UKZN. | Kathleen Subramoney et al |
| EPI_ISL_6825389 | ZARV/NHLS, Department Medical Virology, University of Pretoria | CERI, Centre for Epidemic Response and Innvoation, Stellenbosch University and KRISP, KZN Research Innovation and Sequencing Platform, UKZN. | Amy Strydom et al |
| EPI_ISL_6699733 | National Health Laboratory Services, Virology, Charlotte Maxeke Johannesburg hospital, Parktown, Johannesburg, Gauteng | CERI, Centre for Epidemic Response and Innvoation, Stellenbosch University and KRISP, KZN Research Innovation and Sequencing Platform, UKZN. | Kathleen Subramoney et al |
| EPI_ISL_6745939 | Division of Emerging Infectious Diseases, Bureau of Infectious Diseases Diagnosis Control, Korea Disease Control and Prevention Agency | Division of Emerging Infectious Diseases, Bureau of Infectious Diseases Diagnosis Control, Korea Disease Control and Prevention Agency | Ae Kyung Park et al |
| EPI_ISL_6647957 | LANCET LABORATORY | National Institute for Communicable Diseases of the National Health Laboratory Service | Amoako DG et al |
| EPI_ISL_3544634 | Division of Emerging Infectious Diseases, Bureau of Infectious Diseases Diagnosis Control, Korea Disease Control and Prevention Agency | Division of Emerging Infectious Diseases, Bureau of Infectious Diseases Diagnosis Control, Korea Disease Control and Prevention Agency | Ae Kyung Park et al |
| EPI_ISL_1194635 | Universidad de León | SeqCOVID-SPAIN consortium/IBV(CSIC) | Ana Carvajal et al |
| EPI_ISL_6699738 | National Health Laboratory Services, Virology, Charlotte Maxeke Johannesburg hospital, Parktown, Johannesburg, Gauteng | CERI, Centre for Epidemic Response and Innvoation, Stellenbosch University and KRISP, KZN Research Innovation and Sequencing Platform, UKZN. | Kathleen Subramoney et al |
| EPI_ISL_4251357 | Institute of Microbiology and Immunology, Faculty of Medicine, University of Ljubljana | Institute of Microbiology and Immunology, Faculty of Medicine, University of Ljubljana | Alen Suljič et al |
| EPI_ISL_3666734 | Division of Medical Virology, National Health Laboratory Service (NHLS), Tygerberg Hospital / Stellenbosch University | Division of Medical Virology, National Health Laboratory Service (NHLS), Tygerberg Hospital / Stellenbosch University | Susan Engelbrecht et al |
| EPI_ISL_5601020 | National Laboratory for Health, Environment and Food, OMM, Maribor | NLZOH (National Laboratory for Health, Environment and Food) / CISLD (Clinical Institute of Special Laboratory Diagnostics), University Children's Hospital, University Medical Center Ljubljana | Sandra Janezic et al |

| | | | |
|---|---|---|---|
| EPI_ISL_4272636 | SARS-CoV-2 Sequencing Castilla y Leon-Spain Consortium | SARS-CoV-2 Sequencing Castilla y Leon-Spain Consortium | Antonio Orduña-Domingo et al |
| EPI_ISL_6699740 | National Health Laboratory Services, Virology, Charlotte Maxeke Johannesburg hospital, Parktown, Johannesburg, Gauteng | CERI, Centre for Epidemic Response and Innvoation, Stellenbosch University and KRISP, KZN Research Innovation and Sequencing Platform, UKZN. | Kathleen Subramoney et al |
| EPI_ISL_5918113 | Tshepong Laboratory | National Institute for Communicable Diseases of the National Health Laboratory Service | Amoako DG et al |
| EPI_ISL_5106432 | HOSPITAL UNIVERSITARI DE BELLVITGE | Microbiology Department | Sara Marti et al |
| EPI_ISL_1048491 | National Health Laboratory Service, South Africa | KRISP, KZn Research Innovation and Sequencing Platform | Giandhari J et al |
| EPI_ISL_6509092 | CAP Vic Nord | Banc de Sang i Teixits | Francisco Vidal et al |
| EPI_ISL_5262687 | Hospital General Universitario Albacete | Hospital General Universitario Albacete | Lorena Robles-Fonseca et al |
| EPI_ISL_6842165 | Division of Medical Virology, National Health Laboratory Service (NHLS), Tygerberg Hospital / Stellenbosch University | Division of Medical Virology, National Health Laboratory Service (NHLS), Tygerberg Hospital / Stellenbosch University | Susan Engelbrecht et al |
| EPI_ISL_5924264 | CLILAB | Microbiology Department | Sara Marti et al |
| EPI_ISL_6699742 | National Health Laboratory Services, Virology, Charlotte Maxeke Johannesburg hospital, Parktown, Johannesburg, Gauteng | CERI, Centre for Epidemic Response and Innvoation, Stellenbosch University and KRISP, KZN Research Innovation and Sequencing Platform, UKZN. | Kathleen Subramoney et al |
| EPI_ISL_6699737 | National Health Laboratory Services, Virology, Charlotte Maxeke Johannesburg hospital, Parktown, Johannesburg, Gauteng | CERI, Centre for Epidemic Response and Innvoation, Stellenbosch University and KRISP, KZN Research Innovation and Sequencing Platform, UKZN. | Kathleen Subramoney et al |
| EPI_ISL_6842162 | Division of Medical Virology, National Health Laboratory Service (NHLS), Tygerberg Hospital / Stellenbosch University | Division of Medical Virology, National Health Laboratory Service (NHLS), Tygerberg Hospital / Stellenbosch University | Susan Engelbrecht et al |
| EPI_ISL_1335425 | National Laboratory for Health, Environment and Food, OMM, Novo mesto | CISLD (Clinical Institute of Special Laboratory Diagnostics), University Children's Hospital, University Medical Center Ljubljana | Jernej Kovač et al |
| EPI_ISL_2649265 | SARS-CoV-2 Sequencing Castilla y Leon-Spain Consortium | SARS-CoV-2 Sequencing Castilla y Leon-Spain Consortium | Antonio Orduña-Domingo et al |
| EPI_ISL_3056388 | SARS-CoV-2 Sequencing Castilla y Leon-Spain Consortium | SARS-CoV-2 Sequencing Castilla y Leon-Spain Consortium | Antonio Orduña-Domingo et al |

| | | | |
|---|---|---|---|
| EPI_ISL_2644559 | Institute of Microbiology and Immunology, Faculty of Medicine, University of Ljubljana | Institute of Microbiology and Immunology, Faculty of Medicine, University of Ljubljana | Alen Suljič et al |
| EPI_ISL_5193002 | Institute of Microbiology and Immunology, Faculty of Medicine, University of Ljubljana | Institute of Microbiology and Immunology, Faculty of Medicine, University of Ljubljana | Alen Suljič et al |
| EPI_ISL_4949412 | Institute of Microbiology and Immunology, Faculty of Medicine, University of Ljubljana | Institute of Microbiology and Immunology, Faculty of Medicine, University of Ljubljana | Alen Suljič et al |
| EPI_ISL_6699747 | National Health Laboratory Services, Virology, Charlotte Maxeke Johannesburg hospital, Parktown, Johannesburg, Gauteng | CERI, Centre for Epidemic Response and Innvoation, Stellenbosch University and KRISP, KZN Research Innovation and Sequencing Platform, UKZN. | Kathleen Subramoney et al |
| EPI_ISL_6013704 | Division of Emerging Infectious Diseases, Bureau of Infectious Diseases Diagnosis Control, Korea Disease Control and Prevention Agency | Division of Emerging Infectious Diseases, Bureau of Infectious Diseases Diagnosis Control, Korea Disease Control and Prevention Agency | Ae Kyung Park et al |
| EPI_ISL_6829557 | National Health Laboratory Service, Kwazulu-Natal, South Africa | KRISP, KZn Research Innovation and Sequencing Platform | Nokukhanya M et al |
| EPI_ISL_5634239 | Division of Emerging Infectious Diseases, Bureau of Infectious Diseases Diagnosis Control, Korea Disease Control and Prevention Agency | Division of Emerging Infectious Diseases, Bureau of Infectious Diseases Diagnosis Control, Korea Disease Control and Prevention Agency | Ae Kyung Park et al |
| EPI_ISL_6699755 | National Health Laboratory Services, Virology, Charlotte Maxeke Johannesburg hospital, Parktown, Johannesburg, Gauteng | CERI, Centre for Epidemic Response and Innvoation, Stellenbosch University and KRISP, KZN Research Innovation and Sequencing Platform, UKZN. | Kathleen Subramoney et al |
| EPI_ISL_6699756 | National Health Laboratory Services, Virology, Charlotte Maxeke Johannesburg hospital, Parktown, Johannesburg, Gauteng | CERI, Centre for Epidemic Response and Innvoation, Stellenbosch University and KRISP, KZN Research Innovation and Sequencing Platform, UKZN. | Kathleen Subramoney et al |
| EPI_ISL_4271017 | Institute of Microbiology and Immunology, Faculty of Medicine, University of Ljubljana | Institute of Microbiology and Immunology, Faculty of Medicine, University of Ljubljana | Alen Suljič et al |
| EPI_ISL_2360776 | Vaccines and Infectious Diseases Analytics Research Unit (VIDA) | KRISP, KZn Research Innovation and Sequencing Platform | Baillie Vicky et al |
| EPI_ISL_5102924 | Institute of Microbiology and Immunology, Faculty of Medicine, University of Ljubljana | Institute of Microbiology and Immunology, Faculty of Medicine, University of Ljubljana | Alen Suljič et al |

| | | | |
|---|---|---|---|
| EPI_ISL_5062869 | Institute of Microbiology and Immunology, Faculty of Medicine, University of Ljubljana | Institute of Microbiology and Immunology, Faculty of Medicine, University of Ljubljana | Alen Suljič et al |
| EPI_ISL_6203190 | TSHEPONG LABORATORY | National Institute for Communicable Diseases of the National Health Laboratory Service | Amoako DG et al |
| EPI_ISL_3646056 | SARS-CoV-2 Sequencing Castilla y Leon-Spain Consortium | SARS-CoV-2 Sequencing Castilla y Leon-Spain Consortium | Antonio Orduña-Domingo et al |
| EPI_ISL_6959935 | Division of Emerging Infectious Diseases, Bureau of Infectious Diseases Diagnosis Control, Korea Disease Control and Prevention Agency | Division of Emerging Infectious Diseases, Bureau of Infectious Diseases Diagnosis Control, Korea Disease Control and Prevention Agency | Ae Kyung Park et al |
| EPI_ISL_3847739 | Division of Medical Virology, National Health Laboratory Service (NHLS), Tygerberg Hospital / Stellenbosch University | Division of Medical Virology, National Health Laboratory Service (NHLS), Tygerberg Hospital / Stellenbosch University | Susan Engelbrecht et al |
| EPI_ISL_2360362 | Vaccines and Infectious Diseases Analytics Research Unit (VIDA) | KRISP, KZn Research Innovation and Sequencing Platform | Baillie Vicky et al |
| EPI_ISL_760209 | Division of Emerging Infectious Diseases, Bureau of Infectious Diseases Diagnosis Control, Korea Disease Control and Prevention Agency | Division of Emerging Infectious Diseases, Bureau of Infectious Diseases Diagnosis Control, Korea Disease Control and Prevention Agency | Ae Kyung Park et al |
| EPI_ISL_4070443 | Karolinska University Hospital Solna | Karolinska University Hospital | Jan Albert et al |
| EPI_ISL_2840841 | HOSPITAL COMARCAL DE MELILLA | Instituto de Salud Carlos III | Iglesias-Caballero et al |
| EPI_ISL_1647818 | Hospital General Universitario Gregorio Marañón | Hospital General Universitario Gregorio Marañón | Sergio Buenestado Serrano et al |
| EPI_ISL_2793264 | Swedish national genomic surveillance program of SARS-CoV-2 | The Public Health Agency of Sweden | Maximilian Riess et al |
| EPI_ISL_3306376 | Hospital General Universitario Gregorio Marañón | Hospital General Universitario Gregorio Marañón | Sergio Buenestado Serrano et al |
| EPI_ISL_6045588 | Hospital Universitari Dr. Josep Trueta | Institut d'Investigació Biomèdica de Girona \| Hospital Universitari Dr. Josep Trueta | Mel·lina Pinsach et al |
| EPI_ISL_4204546 | Hospital General Universitario de Alicante - Instituto de Investigación Sanitaria y Biomédica de Alicante | SeqCOVID-SPAIN consortium/IBV(CSIC) | Maripaz Ventero Martín et al |
| EPI_ISL_2820675 | Hospital Universitari i Politècnic La Fe de València | SeqCOVID-SPAIN consortium/IBV(CSIC) | María Dolores Gómez Ruiz et al |

| | | | |
|---|---|---|---|
| EPI_ISL_2549177 | Servicio de Microbiología Clínica (Complejo Hospitalario de Navarra, Pamplona), Instituto de Investigación Sanitaria de Navarra (IdiSNA) | SeqCOVID-SPAIN consortium/IBV(CSIC) | Carmen Ezpeleta Baquedano et al |
| EPI_ISL_4070461 | Karolinska University Hospital Huddinge | Karolinska University Hospital | Jan Albert et al |
| EPI_ISL_2980138 | Swedish national genomic surveillance program of SARS-CoV-2 | The Public Health Agency of Sweden | Maximilian Riess et al |
| EPI_ISL_6324725 | Karolinska University Hospital Solna | Karolinska University Hospital | Jan Albert et al |
| EPI_ISL_3690687 | Sweden/Vastragotaland/Unilabs | Unilabs/Eskilstuna/Sweden | Emma Arvidsson et al |
| EPI_ISL_2793518 | Swedish national genomic surveillance program of SARS-CoV-2 | The Public Health Agency of Sweden | Maximilian Riess et al |
| EPI_ISL_6176729 | Swedish national genomic surveillance program of SARS-CoV-2 | The Public Health Agency of Sweden | Maximilian Riess et al |
| EPI_ISL_2803220 | Centre for Dengue Research and AICBU, Department of Immunology and Molecular Medicine | Centre for Dengue Research and AICBU, Department of Immunology and Molecular Medicine | Chandima Jeewandara et al |
| EPI_ISL_7050911 | Hospital Universitari Vall d'Hebron - Vall d'Hebron Institut de Recerca | Hospital Universitari Vall d'Hebron - Vall d'Hebron Institut de Recerca | Cristina Andrés et al |
| EPI_ISL_861774 | Hospital General Universitario Gregorio Marañón | SeqCOVID-SPAIN consortium/IBV(CSIC) | Darío García de Viedma et al |
| EPI_ISL_5334047 | Akademiska Sjukhuset, Clinical Microbiology and Hospital Hygiene | Akademiska Sjukhuset, Clinical Microbiology and Hospital Hygiene | Rene Kaden et al |
| EPI_ISL_1622673 | Servicio de Microbiología Clínica (Complejo Hospitalario de Navarra, Pamplona) | Centro de Secuenciación NASERTIC | Carmen Ezpeleta Baquedano et al |
| EPI_ISL_3047265 | Hospital Universitari Vall d'Hebron - Vall d'Hebron Institut de Recerca | Hospital Universitari Vall d'Hebron - Vall d'Hebron Institut de Recerca | Cristina Andrés et al |
| EPI_ISL_6709755 | Swedish national genomic surveillance program of SARS-CoV-2 | The Public Health Agency of Sweden | Maximilian Riess et al |
| EPI_ISL_6563525 | Karolinska University Hospital Solna | Karolinska University Hospital | Jan Albert et al |
| EPI_ISL_2417127 | Swedish national genomic surveillance program of SARS-CoV-2 | The Public Health Agency of Sweden | Maximilian Riess et al |
| EPI_ISL_450833 | Wetterhalsan | The Public Health Agency of Sweden | Anders Tengblad et al |
| EPI_ISL_2419057 | Swedish national genomic surveillance program of SARS-CoV-2 | The Public Health Agency of Sweden | Maximilian Riess et al |

| | | | |
|---|---|---|---|
| EPI_ISL_1809149 | Swedish national genomic surveillance program of SARS-CoV-2 | The Public Health Agency of Sweden | Maximilian Riess et al |
| EPI_ISL_831256 | Hospital Universitario La Paz (Madrid) | SeqCOVID-SPAIN consortium/IBV(CSIC) | Fernando Lázaro-Perona et al |
| EPI_ISL_6374918 | Karolinska University Hospital Solna | Karolinska University Hospital | Jan Albert et al |
| EPI_ISL_2794234 | Swedish national genomic surveillance program of SARS-CoV-2 | The Public Health Agency of Sweden | Maximilian Riess et al |
| EPI_ISL_2196889 | Swedish national genomic surveillance program of SARS-CoV-2 | The Public Health Agency of Sweden | Maximilian Riess et al |
| EPI_ISL_1746242 | Servicio de Microbiología Clínica (Complejo Hospitalario de Navarra, Pamplona) | Centro de Secuenciación NASERTIC | Carmen Ezpeleta Baquedano et al |
| EPI_ISL_1159206 | HOSPITAL GENERAL UNIVERSITARIO REINA SOFIA | Instituto de Salud Carlos III | Iglesias-Caballero et al |
| EPI_ISL_3907640 | Servicio de Microbiología. Hospital General Universitario de Castellón | SeqCOVID-SPAIN consortium/IBV(CSIC) | Rosario Moreno Muñoz et al |
| EPI_ISL_1110287 | Hospital de Sagunto | SeqCOVID-SPAIN consortium/IBV(CSIC) | Noelia Lozano Rodríguez et al |
| EPI_ISL_6640283 | Microbiology Department, University Hospital Donostia | Microbiology Department, University Hospital Donostia | Marimon JM et al |
| EPI_ISL_6902675 | Hospital General Universitario Gregorio Marañón | Hospital General Universitario Gregorio Marañón | Sergio Buenestado Serrano et al |
| EPI_ISL_1903490 | Swedish national genomic surveillance program of SARS-CoV-2 | The Public Health Agency of Sweden | Maximilian Riess et al |
| EPI_ISL_1617913 | Swedish national genomic surveillance program of SARS-CoV-2 | The Public Health Agency of Sweden | Swedish national genomic surveillance program of SARS-CoV-2 et al |
| EPI_ISL_3786262 | Swedish national genomic surveillance program of SARS-CoV-2 | The Public Health Agency of Sweden | Maximilian Riess et al |
| EPI_ISL_2828143 | Hospital Universitari Vall d'Hebron - Vall d'Hebron Institut de Recerca | Hospital Universitari Vall d'Hebron - Vall d'Hebron Institut de Recerca | Cristina Andrés et al |
| EPI_ISL_1510017 | HOSPITAL UNIVERSITARIO VIRGEN DE LA ARRIXACA | Instituto de Salud Carlos III | Vázquez-Morón et al |
| EPI_ISL_6204377 | Microbiology Department, University Hospital Donostia | Microbiology Department, University Hospital Donostia | Marimon JM et al |
| EPI_ISL_4535366 | Karolinska University Hospital Solna | Karolinska University Hospital | Jan Albert et al |
| EPI_ISL_2212121 | Swedish national genomic surveillance program of SARS-CoV-2 | The Public Health Agency of Sweden | Maximilian Riess et al |

| | | | |
|---|---|---|---|
| EPI_ISL_541128 | Servicio de Microbiología. Hospital Universitario Donostia. OSI Donostialdea. Área de Enfermedades Infecciosas, Grupo de Infección Respiratoria y Resistencia Antimicrobiana. Instituto de Investigación Sanitaria Biodonostia | SeqCOVID-SPAIN consortium/Institute of Biomedicine of Valencia, IBV-CSIC | Gustavo Cilla et al |
| EPI_ISL_3795484 | Swedish national genomic surveillance program of SARS-CoV-2 | The Public Health Agency of Sweden | Maximilian Riess et al |
| EPI_ISL_7042252 | Hospital General Universitario Gregorio Marañón | Hospital General Universitario Gregorio Marañón | Sergio Buenestado Serrano et al |
| EPI_ISL_4108631 | Swedish national genomic surveillance program of SARS-CoV-2 | The Public Health Agency of Sweden | Maximilian Riess et al |
| EPI_ISL_4070526 | Karolinska University Hospital Solna | Karolinska University Hospital | Jan Albert et al |
| EPI_ISL_1093120 | Centro de Investigación Biomédica de La Rioja - Hospital San Pedro Logroño | SeqCOVID-SPAIN consortium/IBV(CSIC) | María de Toro et al |
| EPI_ISL_6471034 | Akademiska Sjukhuset, Clinical Microbiology and Hospital Hygiene | Akademiska Sjukhuset, Clinical Microbiology and Hospital Hygiene | Rene Kaden et al |
| EPI_ISL_6179811 | Swedish national genomic surveillance program of SARS-CoV-2 | The Public Health Agency of Sweden | Maximilian Riess et al |
| EPI_ISL_2790431 | Servizo de Microbioloxía. Complexo Hospitalario de Santiago de Compostela | Servizo de Microbioloxía. Complexo Hospitalario de Santiago de Compostela | Xosé Costa et al |
| EPI_ISL_1322998 | Hospital Universitario La Paz (Madrid) | SeqCOVID-SPAIN consortium/IBV(CSIC) | Fernando Lázaro-Perona et al |
| EPI_ISL_6971860 | Hospital General Universitario Gregorio Marañón | Hospital General Universitario Gregorio Marañón | Sergio Buenestado Serrano et al |
| EPI_ISL_6697544 | Microbiology Department, University Hospital Donostia | Microbiology Department, University Hospital Donostia | Marimon JM et al |
| EPI_ISL_2812758 | HOSPITAL MATEU ORFILA | HOSPITAL UNIVERSITARIO SON ESPASES | Servicio de Microbiología et al |
| EPI_ISL_2211774 | Swedish national genomic surveillance program of SARS-CoV-2 | The Public Health Agency of Sweden | Maximilian Riess et al |
| EPI_ISL_732781 | Centro de Investigación Biomédica de La Rioja - Hospital San Pedro Logroño | SeqCOVID-SPAIN consortium/IBV(CSIC) | María de Toro et al |
| EPI_ISL_3275460 | Allergy, Immunology and Cell Biology Unit (AICBU) | Allergy, Immunology and Cell Biology Unit (AICBU) | Chandima Jeewandara et al |
| EPI_ISL_2438725 | HOSPITAL UNIVERSITARIO SON ESPASES | HOSPITAL UNIVERSITARIO SON ESPASES | Servicio de Microbiología et al |
| EPI_ISL_2516942 | Microbiology Department. Complexo Hospitalario Universitario de Vigo | Microbiology Department. Complexo Hospitalario Universitario de Vigo | Alfaya N et al |

| | | | |
|---|---|---|---|
| EPI_ISL_1159221 | HOSPITAL GENERAL UNIVERSITARIO SANTA LUCIA | Instituto de Salud Carlos III | Iglesias-Caballero et al |
| EPI_ISL_2957024 | Hospital Universitari Bellvitge | Microbiology Department | Sara Marti et al |
| EPI_ISL_2209987 | Swedish national genomic surveillance program of SARS-CoV-2 | The Public Health Agency of Sweden | Maximilian Riess et al |
| EPI_ISL_2414478 | Swedish national genomic surveillance program of SARS-CoV-2 | The Public Health Agency of Sweden | Maximilian Riess et al |
| EPI_ISL_1382981 | Servicio de Microbiología Clínica (Complejo Hospitalario de Navarra, Pamplona) | Centro de Secuenciación NASERTIC | Carmen Ezpeleta Baquedano et al |
| EPI_ISL_2196905 | Swedish national genomic surveillance program of SARS-CoV-2 | The Public Health Agency of Sweden | Maximilian Riess et al |
| EPI_ISL_6628705 | Servicio de Microbiologia Hospital Ramon y Cajal | Servicio de Microbiologia Hospital Ramon y Cajal | Ponce M et al |
| EPI_ISL_2888745 | Swedish national genomic surveillance program of SARS-CoV-2 | The Public Health Agency of Sweden | Maximilian Riess et al |
| EPI_ISL_3007205 | Unilabs | Karolinska University Hospital | Jan Albert et al |
| EPI_ISL_2222286 | Swedish national genomic surveillance program of SARS-CoV-2 | The Public Health Agency of Sweden | Maximilian Riess et al |
| EPI_ISL_2208251 | Swedish national genomic surveillance program of SARS-CoV-2 | The Public Health Agency of Sweden | Maximilian Riess et al |
| EPI_ISL_2208223 | Swedish national genomic surveillance program of SARS-CoV-2 | The Public Health Agency of Sweden | Maximilian Riess et al |
| EPI_ISL_3786629 | Swedish national genomic surveillance program of SARS-CoV-2 | The Public Health Agency of Sweden | Maximilian Riess et al |
| EPI_ISL_3502619 | HOSPITAL UNIVERSITARIO 12 DE OCTUBRE | HOSPITAL UNIVERSITARIO 12 DE OCTUBRE | Carmen Martín-Higuera et al |
| EPI_ISL_2271626 | Servicio de Microbiología. Consorcio Hospital General Universitario de Valencia | SeqCOVID-SPAIN consortium/IBV(CSIC) | María Dolores Ocete et al |
| EPI_ISL_6343218 | Karolinska University Hospital Solna | Karolinska University Hospital | Jan Albert et al |
| EPI_ISL_3299485 | Akademiska Sjukhuset, Clinical Microbiology and Hospital Hygiene | Akademiska Sjukhuset, Clinical Microbiology and Hospital Hygiene | Rene Kaden et al |
| EPI_ISL_4137159 | HOSPITAL CAN MISSES | HOSPITAL UNIVERSITARIO SON ESPASES | Dr. Antonio Oliver et al |
| EPI_ISL_1577109 | Hospital General Universitario Gregorio Marañón | SeqCOVID-SPAIN consortium/IBV(CSIC) | Darío García de Viedma et al |
| EPI_ISL_3785742 | Swedish national genomic surveillance program of SARS-CoV-2 | The Public Health Agency of Sweden | Maximilian Riess et al |

| | | | |
|---|---|---|---|
| EPI_ISL_6883250 | Swedish national genomic surveillance program of SARS-CoV-2 | The Public Health Agency of Sweden | Maximilian Riess et al |
| EPI_ISL_2214654 | Swedish national genomic surveillance program of SARS-CoV-2 | The Public Health Agency of Sweden | Maximilian Riess et al |
| EPI_ISL_452239 | Narhalsan Backa vardcentral | The Public Health Agency of Sweden | Mats Olsson et al |
| EPI_ISL_3158240 | Swedish national genomic surveillance program of SARS-CoV-2 | The Public Health Agency of Sweden | Maximilian Riess et al |
| EPI_ISL_2600999 | Servicio de Microbiología Clínica (Complejo Hospitalario de Navarra, Pamplona), Instituto de Investigación Sanitaria de Navarra (IdiSNA) | SeqCOVID-SPAIN consortium/IBV(CSIC) | Carmen Ezpeleta Baquedano et al |
| EPI_ISL_2211939 | Swedish national genomic surveillance program of SARS-CoV-2 | The Public Health Agency of Sweden | Maximilian Riess et al |
| EPI_ISL_1716667 | Servicio de Microbiología. Consorcio Hospital General Universitario de Valencia | SeqCOVID-SPAIN consortium/IBV(CSIC) | María Dolores Ocete et al |
| EPI_ISL_6375092 | Karolinska University Hospital Solna | Karolinska University Hospital | Jan Albert et al |
| EPI_ISL_7050918 | Hospital Universitari Vall d'Hebron - Vall d'Hebron Institut de Recerca | Hospital Universitari Vall d'Hebron - Vall d'Hebron Institut de Recerca | Cristina Andrés et al |
| EPI_ISL_3158688 | Swedish national genomic surveillance program of SARS-CoV-2 | The Public Health Agency of Sweden | Maximilian Riess et al |
| EPI_ISL_4006749 | Akademiska Sjukhuset, Clinical Microbiology and Hospital Hygiene | Akademiska Sjukhuset, Clinical Microbiology and Hospital Hygiene | Rene Kaden et al |
| EPI_ISL_2206801 | Swedish national genomic surveillance program of SARS-CoV-2 | The Public Health Agency of Sweden | Maximilian Riess et al |
| EPI_ISL_2271136 | Departamento de Microbiología, CDB, Hospital Clínic, Barcelona | SeqCOVID-SPAIN consortium/IBV(CSIC) | Andrea Vergara et al |
| EPI_ISL_6179801 | Swedish national genomic surveillance program of SARS-CoV-2 | The Public Health Agency of Sweden | Maximilian Riess et al |
| EPI_ISL_6116006 | Hospital Universitari Dr. Josep Trueta | Institut d'Investigació Biomèdica de Girona \| Hospital Universitari Dr. Josep Trueta | Mel·lina Pinsach et al |
| EPI_ISL_6376967 | Karolinska University Hospital Solna | Karolinska University Hospital | Jan Albert et al |
| EPI_ISL_871986 | Hospital Clínico San Carlos | Instituto de Salud Carlos III | Iglesias-Caballero et al |
| EPI_ISL_4820464 | HOSPITAL UNIVERSITARIO SON ESPASES | HOSPITAL UNIVERSITARIO SON ESPASES | Dr. Antonio Oliver et al |

| | | | |
|---|---|---|---|
| EPI_ISL_2416988 | Swedish national genomic surveillance program of SARS-CoV-2 | The Public Health Agency of Sweden | Maximilian Riess et al |
| EPI_ISL_2221689 | Swedish national genomic surveillance program of SARS-CoV-2 | The Public Health Agency of Sweden | Maximilian Riess et al |
| EPI_ISL_2206471 | Swedish national genomic surveillance program of SARS-CoV-2 | The Public Health Agency of Sweden | Maximilian Riess et al |
| EPI_ISL_6851526 | Hospital General Universitario Gregorio Marañón | Hospital General Universitario Gregorio Marañón | Sergio Buenestado Serrano et al |
| EPI_ISL_1600671 | Swedish national genomic surveillance program of SARS-CoV-2 | The Public Health Agency of Sweden | Swedish national genomic surveillance program of SARS-CoV-2 et al |
| EPI_ISL_3783768 | Swedish national genomic surveillance program of SARS-CoV-2 | The Public Health Agency of Sweden | Maximilian Riess et al |
| EPI_ISL_4470785 | Swedish national genomic surveillance program of SARS-CoV-2 | The Public Health Agency of Sweden | Maximilian Riess et al |
| EPI_ISL_2208137 | Swedish national genomic surveillance program of SARS-CoV-2 | The Public Health Agency of Sweden | Maximilian Riess et al |
| EPI_ISL_6135045 | Swedish national genomic surveillance program of SARS-CoV-2 | The Public Health Agency of Sweden | Maximilian Riess et al |
| EPI_ISL_1604592 | Swedish national genomic surveillance program of SARS-CoV-2 | The Public Health Agency of Sweden | Swedish national genomic surveillance program of SARS-CoV-2 et al |
| EPI_ISL_1419991 | Swedish national genomic surveillance program of SARS-CoV-2 | The Public Health Agency of Sweden | Swedish national genomic surveillance program of SARS-CoV-2 et al |
| EPI_ISL_4337954 | Swedish national genomic surveillance program of SARS-CoV-2 | The Public Health Agency of Sweden | Maximilian Riess et al |
| EPI_ISL_2227324 | Institute for Infectious Diseases, University of Bern | Institute for Infectious Diseases, University of Bern | Stefan Neuenschwander et al |
| EPI_ISL_1899449 | Swedish national genomic surveillance program of SARS-CoV-2 | The Public Health Agency of Sweden | Maximilian Riess et al |
| EPI_ISL_4108784 | Swedish national genomic surveillance program of SARS-CoV-2 | The Public Health Agency of Sweden | Maximilian Riess et al |
| EPI_ISL_3796913 | Swedish national genomic surveillance program of SARS-CoV-2 | The Public Health Agency of Sweden | Maximilian Riess et al |
| EPI_ISL_4762129 | Clinical Microbiology, Infection Prevention and Control | Section for Molecular Diagnostics | Björn Hallström et al |
| EPI_ISL_7062525 | Spital Riggisberg | Institute for Infectious Diseases, University of Bern | Stefan Neuenschwander et al |

| | | | |
|---|---|---|---|
| EPI_ISL_6134676 | Swedish national genomic surveillance program of SARS-CoV-2 | The Public Health Agency of Sweden | Maximilian Riess et al |
| EPI_ISL_1191596 | Swedish national genomic surveillance program of SARS-CoV-2 | The Public Health Agency of Sweden | Swedish national genomic surveillance program of SARS-CoV-2 et al |
| EPI_ISL_2216108 | Swedish national genomic surveillance program of SARS-CoV-2 | The Public Health Agency of Sweden | Maximilian Riess et al |
| EPI_ISL_1902635 | Swedish national genomic surveillance program of SARS-CoV-2 | The Public Health Agency of Sweden | Maximilian Riess et al |
| EPI_ISL_899280 | Viollier AG | Department of Biosystems Science and Engineering, ETH Zürich | Christian Beisel et al |
| EPI_ISL_6708598 | Swedish national genomic surveillance program of SARS-CoV-2 | The Public Health Agency of Sweden | Maximilian Riess et al |
| EPI_ISL_3787937 | Swedish national genomic surveillance program of SARS-CoV-2 | The Public Health Agency of Sweden | Maximilian Riess et al |
| EPI_ISL_2222616 | Swedish national genomic surveillance program of SARS-CoV-2 | The Public Health Agency of Sweden | Maximilian Riess et al |
| EPI_ISL_6712088 | Swedish national genomic surveillance program of SARS-CoV-2 | The Public Health Agency of Sweden | Maximilian Riess et al |
| EPI_ISL_1601470 | Swedish national genomic surveillance program of SARS-CoV-2 | The Public Health Agency of Sweden | Swedish national genomic surveillance program of SARS-CoV-2 et al |
| EPI_ISL_5261612 | Clinical Microbiology, Infection Prevention and Control | Section for Molecular Diagnostics | Björn Hallström et al |
| EPI_ISL_2584370 | Swedish national genomic surveillance program of SARS-CoV-2 | The Public Health Agency of Sweden | Maximilian Riess et al |
| EPI_ISL_6780972 | Ender Diagnostics AG | Microbiology | Nadia Wohlwend et al |
| EPI_ISL_1602311 | Swedish national genomic surveillance program of SARS-CoV-2 | The Public Health Agency of Sweden | Swedish national genomic surveillance program of SARS-CoV-2 et al |
| EPI_ISL_1901905 | Swedish national genomic surveillance program of SARS-CoV-2 | The Public Health Agency of Sweden | Maximilian Riess et al |
| EPI_ISL_1618513 | Swedish national genomic surveillance program of SARS-CoV-2 | The Public Health Agency of Sweden | Swedish national genomic surveillance program of SARS-CoV-2 et al |
| EPI_ISL_1903821 | Swedish national genomic surveillance program of SARS-CoV-2 | The Public Health Agency of Sweden | Maximilian Riess et al |
| EPI_ISL_1290773 | Swedish national genomic surveillance program of SARS-CoV-2 | The Public Health Agency of Sweden | Swedish national genomic surveillance program of SARS-CoV-2 et al |

| | | | |
|---|---|---|---|
| EPI_ISL_1677676 | Swedish national genomic surveillance program of SARS-CoV-2 | The Public Health Agency of Sweden | Maximilian Riess et al |
| EPI_ISL_2257606 | Swedish national genomic surveillance program of SARS-CoV-2 | The Public Health Agency of Sweden | Maximilian Riess et al |
| EPI_ISL_6710849 | Swedish national genomic surveillance program of SARS-CoV-2 | The Public Health Agency of Sweden | Maximilian Riess et al |
| EPI_ISL_4905438 | labor team w AG | Department of Biosystems Science and Engineering, ETH Zürich | Ivan Topolsky et al |
| EPI_ISL_2613475 | Swedish national genomic surveillance program of SARS-CoV-2 | The Public Health Agency of Sweden | Maximilian Riess et al |
| EPI_ISL_4865469 | Swedish national genomic surveillance program of SARS-CoV-2 | The Public Health Agency of Sweden | Maximilian Riess et al |
| EPI_ISL_1618301 | Swedish national genomic surveillance program of SARS-CoV-2 | The Public Health Agency of Sweden | Swedish national genomic surveillance program of SARS-CoV-2 et al |
| EPI_ISL_1418360 | Swedish national genomic surveillance program of SARS-CoV-2 | The Public Health Agency of Sweden | Swedish national genomic surveillance program of SARS-CoV-2 et al |
| EPI_ISL_4858183 | Swedish national genomic surveillance program of SARS-CoV-2 | The Public Health Agency of Sweden | Maximilian Riess et al |
| EPI_ISL_2196762 | Swedish national genomic surveillance program of SARS-CoV-2 | The Public Health Agency of Sweden | Maximilian Riess et al |
| EPI_ISL_1600667 | Swedish national genomic surveillance program of SARS-CoV-2 | The Public Health Agency of Sweden | Swedish national genomic surveillance program of SARS-CoV-2 et al |
| EPI_ISL_6855635 | labor team w AG | Department of Biosystems Science and Engineering, ETH Zürich | Ivan Topolsky et al |
| EPI_ISL_1676169 | Swedish national genomic surveillance program of SARS-CoV-2 | The Public Health Agency of Sweden | Maximilian Riess et al |
| EPI_ISL_4855068 | Swedish national genomic surveillance program of SARS-CoV-2 | The Public Health Agency of Sweden | Maximilian Riess et al |
| EPI_ISL_1199070 | Swedish national genomic surveillance program of SARS-CoV-2 | The Public Health Agency of Sweden | Swedish national genomic surveillance program of SARS-CoV-2 et al |
| EPI_ISL_1808626 | Swedish national genomic surveillance program of SARS-CoV-2 | The Public Health Agency of Sweden | Maximilian Riess et al |
| EPI_ISL_1618270 | Swedish national genomic surveillance program of SARS-CoV-2 | The Public Health Agency of Sweden | Swedish national genomic surveillance program of SARS-CoV-2 et al |
| EPI_ISL_2413040 | Swedish national genomic surveillance program of SARS-CoV-2 | The Public Health Agency of Sweden | Maximilian Riess et al |

| | | | |
|---|---|---|---|
| EPI_ISL_2613437 | Swedish national genomic surveillance program of SARS-CoV-2 | The Public Health Agency of Sweden | Maximilian Riess et al |
| EPI_ISL_3761507 | Clinical Microbiology, Infection Prevention and Control | Section for Molecular Diagnostics | Björn Hallström et al |
| EPI_ISL_1408504 | Viollier AG | Department of Biosystems Science and Engineering, ETH Zürich | Chaoran Chen et al |
| EPI_ISL_1607030 | Swedish national genomic surveillance program of SARS-CoV-2 | The Public Health Agency of Sweden | Swedish national genomic surveillance program of SARS-CoV-2 et al |
| EPI_ISL_1606821 | Swedish national genomic surveillance program of SARS-CoV-2 | The Public Health Agency of Sweden | Swedish national genomic surveillance program of SARS-CoV-2 et al |
| EPI_ISL_1198580 | Swedish national genomic surveillance program of SARS-CoV-2 | The Public Health Agency of Sweden | Swedish national genomic surveillance program of SARS-CoV-2 et al |
| EPI_ISL_2411156 | Swedish national genomic surveillance program of SARS-CoV-2 | The Public Health Agency of Sweden | Maximilian Riess et al |
| EPI_ISL_1901935 | Swedish national genomic surveillance program of SARS-CoV-2 | The Public Health Agency of Sweden | Maximilian Riess et al |
| EPI_ISL_2621659 | Clinical Microbiology, Infection Prevention and Control | Section for Molecular Diagnostics | Björn Hallström et al |
| EPI_ISL_2980200 | Swedish national genomic surveillance program of SARS-CoV-2 | The Public Health Agency of Sweden | Maximilian Riess et al |
| EPI_ISL_6116333 | Unilabs | Unilabs | Emma Arvidsson et al |
| EPI_ISL_1607041 | Swedish national genomic surveillance program of SARS-CoV-2 | The Public Health Agency of Sweden | Swedish national genomic surveillance program of SARS-CoV-2 et al |
| EPI_ISL_4337578 | Swedish national genomic surveillance program of SARS-CoV-2 | The Public Health Agency of Sweden | Maximilian Riess et al |
| EPI_ISL_2208768 | Swedish national genomic surveillance program of SARS-CoV-2 | The Public Health Agency of Sweden | Maximilian Riess et al |
| EPI_ISL_3984581 | Swedish national genomic surveillance program of SARS-CoV-2 | The Public Health Agency of Sweden | Maximilian Riess et al |
| EPI_ISL_6720525 | Viollier AG | Department of Biosystems Science and Engineering, ETH Zürich | Andrea Patrizia Salzmann et al |
| EPI_ISL_1904008 | Swedish national genomic surveillance program of SARS-CoV-2 | The Public Health Agency of Sweden | Maximilian Riess et al |
| EPI_ISL_3853933 | Viollier AG | Department of Biosystems Science and Engineering, ETH Zürich | Chaoran Chen et al |
| EPI_ISL_4317859 | Viollier AG | Department of Biosystems Science and Engineering, ETH Zürich | Christian Beisel et al |
| EPI_ISL_1364142 | Swedish national genomic surveillance program of SARS-CoV-2 | The Public Health Agency of Sweden | Swedish national genomic surveillance program of SARS-CoV-2 et al |

| | | | |
|---|---|---|---|
| EPI_ISL_6883773 | Swedish national genomic surveillance program of SARS-CoV-2 | The Public Health Agency of Sweden | Maximilian Riess et al |
| EPI_ISL_1748484 | Swedish national genomic surveillance program of SARS-CoV-2 | The Public Health Agency of Sweden | Maximilian Riess et al |
| EPI_ISL_3465108 | Viollier AG | Department of Biosystems Science and Engineering, ETH Zürich | Chaoran Chen et al |
| EPI_ISL_1900997 | Swedish national genomic surveillance program of SARS-CoV-2 | The Public Health Agency of Sweden | Maximilian Riess et al |
| EPI_ISL_6712174 | Swedish national genomic surveillance program of SARS-CoV-2 | The Public Health Agency of Sweden | Maximilian Riess et al |
| EPI_ISL_2256125 | Swedish national genomic surveillance program of SARS-CoV-2 | The Public Health Agency of Sweden | Maximilian Riess et al |
| EPI_ISL_6884183 | Swedish national genomic surveillance program of SARS-CoV-2 | The Public Health Agency of Sweden | Maximilian Riess et al |
| EPI_ISL_2410821 | Swedish national genomic surveillance program of SARS-CoV-2 | The Public Health Agency of Sweden | Maximilian Riess et al |
| EPI_ISL_2412363 | Swedish national genomic surveillance program of SARS-CoV-2 | The Public Health Agency of Sweden | Maximilian Riess et al |
| EPI_ISL_5839073 | Sweden/Vastragotaland/Unilabs | Unilabs/Eskilstuna/Sweden | Emma Arvidsson et al |
| EPI_ISL_3761415 | Clinical Microbiology, Infection Prevention and Control | Section for Molecular Diagnostics | Björn Hallström et al |
| EPI_ISL_1192627 | Swedish national genomic surveillance program of SARS-CoV-2 | The Public Health Agency of Sweden | Swedish national genomic surveillance program of SARS-CoV-2 et al |
| EPI_ISL_3159374 | Swedish national genomic surveillance program of SARS-CoV-2 | The Public Health Agency of Sweden | Maximilian Riess et al |
| EPI_ISL_913442 | Institute for Infectious Diseases, University of Bern, Switzerland | Institute for Infectious Diseases, University of Bern, Switzerland | Michel C Koch et al |
| EPI_ISL_1259984 | Viollier AG | Department of Biosystems Science and Engineering, ETH Zürich | Chaoran Chen et al |
| EPI_ISL_3854212 | Viollier AG | Department of Biosystems Science and Engineering, ETH Zürich | Andrea Patrizia Salzmann et al |
| EPI_ISL_6710858 | Swedish national genomic surveillance program of SARS-CoV-2 | The Public Health Agency of Sweden | Maximilian Riess et al |
| EPI_ISL_3431693 | Orebro University Hospital, Dept Laboratory Medicine, Clinical Microbiology | Orebro University Hospital | Sundqvist M et al |
| EPI_ISL_3794338 | Swedish national genomic surveillance program of SARS-CoV-2 | The Public Health Agency of Sweden | Maximilian Riess et al |

| | | | |
|---|---|---|---|
| EPI_ISL_1600035 | Swedish national genomic surveillance program of SARS-CoV-2 | The Public Health Agency of Sweden | Swedish national genomic surveillance program of SARS-CoV-2 et al |
| EPI_ISL_1677482 | Swedish national genomic surveillance program of SARS-CoV-2 | The Public Health Agency of Sweden | Maximilian Riess et al |
| EPI_ISL_2350324 | Clinical Microbiology, Infection Prevention and Control | Section for Molecular Diagnostics | Björn Hallström et al |
| EPI_ISL_1200299 | Swedish national genomic surveillance program of SARS-CoV-2 | The Public Health Agency of Sweden | Swedish national genomic surveillance program of SARS-CoV-2 et al |
| EPI_ISL_6100892 | Institute for Infectious Diseases | Institute for Infectious Diseases, University of Bern | Nicole Liechti et al |
| EPI_ISL_6720476 | Viollier AG | Department of Biosystems Science and Engineering, ETH Zürich | Andrea Patrizia Salzmann et al |
| EPI_ISL_3781595 | Swedish national genomic surveillance program of SARS-CoV-2 | The Public Health Agency of Sweden | Maximilian Riess et al |
| EPI_ISL_5865453 | Ender Diagnostics AG | Microbiology | Nadia Wohlwend et al |
| EPI_ISL_1676785 | Swedish national genomic surveillance program of SARS-CoV-2 | The Public Health Agency of Sweden | Maximilian Riess et al |
| EPI_ISL_4471202 | Swedish national genomic surveillance program of SARS-CoV-2 | The Public Health Agency of Sweden | Maximilian Riess et al |
| EPI_ISL_2412278 | Swedish national genomic surveillance program of SARS-CoV-2 | The Public Health Agency of Sweden | Maximilian Riess et al |
| EPI_ISL_2584343 | Swedish national genomic surveillance program of SARS-CoV-2 | The Public Health Agency of Sweden | Maximilian Riess et al |
| EPI_ISL_5532811 | labor team w AG | Department of Biosystems Science and Engineering, ETH Zürich | Ivan Topolsky et al |
| EPI_ISL_1002908 | Viollier AG | Department of Biosystems Science and Engineering, ETH Zürich | Chaoran Chen et al |
| EPI_ISL_1760982 | Viollier AG | Department of Biosystems Science and Engineering, ETH Zürich | Chaoran Chen et al |
| EPI_ISL_3906808 | Ministry of Health Turkey | Ministry of Health Turkey | Fatma Bayrakdar et al |
| EPI_ISL_5065512 | Medical Correctional Hospital | Division of Genomic Medicine and Innovation support,Department of Medical Sciences, Ministry of Public Health, Thailand | Surakameth Mahasirimongkol et al |
| EPI_ISL_3399836 | Ministry of Health Turkey | Ministry of Health Turkey | Fatma Bayrakdar et al |
| EPI_ISL_5466207 | Phatthalung Hospital | Division of Genomic Medicine and Innovation support,Department of Medical Sciences, Ministry of Public Health, Thailand | Surakameth Mahasirimongkol et al |
| EPI_ISL_847963 | University Hospitals of Geneva, Laboratory of Virology | HUG, Laboratory of Virology and the Health2030 Genome Center | Samuel Cordey et al |
| EPI_ISL_1119103 | Viollier AG | Department of Biosystems Science and Engineering, ETH Zürich | Christian Beisel et al |

| | | | |
|---|---|---|---|
| EPI_ISL_1598111 | Viollier AG | Department of Biosystems Science and Engineering, ETH Zürich | Christian Beisel et al |
| EPI_ISL_3066615 | Ministry of Health Turkey | Ministry of Health Turkey | Fatma Bayrakdar et al |
| EPI_ISL_6380320 | Viollier AG | Department of Biosystems Science and Engineering, ETH Zürich | Christian Beisel et al |
| EPI_ISL_4001063 | Ministry of Health Turkey | Ministry of Health Turkey | Fatma Bayrakdar et al |
| EPI_ISL_6329074 | Viollier AG | Department of Biosystems Science and Engineering, ETH Zürich | Chaoran Chen et al |
| EPI_ISL_2307630 | Viollier AG | Department of Biosystems Science and Engineering, ETH Zürich | Chaoran Chen et al |
| EPI_ISL_3259921 | EHNV | Laboratory of genomics and metagenomics | Trestan Pillonel et al |
| EPI_ISL_1658494 | Viollier AG | Department of Biosystems Science and Engineering, ETH Zurich | Chaoran Chen et al |
| EPI_ISL_1750021 | Viollier AG | Department of Biosystems Science and Engineering, ETH Zürich | Chaoran Chen et al |
| EPI_ISL_2918802 | Spital Limmattal | Institute of Medical Virology | Daniel Ehrsam et al |
| EPI_ISL_603369 | Viollier AG | Department of Biosystems Science and Engineering, ETH Zürich | Christian Beisel et al |
| EPI_ISL_603374 | Viollier AG | Department of Biosystems Science and Engineering, ETH Zürich | Christian Beisel et al |
| EPI_ISL_7042168 | Clinical Virology | Clinical Bacteriology, University Hospital Basel | Tim Roloff et al |
| EPI_ISL_3566066 | Ministry of Health Turkey | Ministry of Health Turkey | Fatma Bayrakdar et al |
| EPI_ISL_2629172 | Turkish Society of Internal Medicine | Gen Era Diagnostics Corporation Life Science Research and Molecular Diagnostics | Serhat Ünal et al |
| EPI_ISL_1708164 | University Hospitals of Geneva, Laboratory of Virology | HUG, Laboratory of Virology and the Health2030 Genome Center | Samuel Cordey et al |
| EPI_ISL_7042354 | Biolytix AG | Clinical Bacteriology, University Hospital Basel | Tim Roloff et al |
| EPI_ISL_603694 | Viollier AG | Department of Biosystems Science and Engineering, ETH Zürich | Christian Beisel et al |
| EPI_ISL_5148739 | labor team w AG | Department of Biosystems Science and Engineering, ETH Zürich | Ivan Topolsky et al |
| EPI_ISL_2403435 | Ministry of Health Turkey | Ministry of Health Turkey | Fatma Bayrakdar et al |
| EPI_ISL_4800929 | Synlab Bioggio | Synlab Bioggio | Cinzia ZEHNDER et al |
| EPI_ISL_1001852 | Viollier AG | Department of Biosystems Science and Engineering, ETH Zürich | Christian Beisel et al |
| EPI_ISL_4234692 | labor team w AG | Department of Biosystems Science and Engineering, ETH Zürich | Ivan Topolsky et al |
| EPI_ISL_2462493 | UniversitŠtsSpital ZŸrich | Institute of Medical Virology | Daniel Ehrsam et al |
| EPI_ISL_3666929 | Laboratorio di Microbiologia | Laboratorio di Microbiologia | Martinetti Lucchini Gladys et al |
| EPI_ISL_3938835 | CHUV | Laboratory of genomics and metagenomics | Trestan Pillonel et al |
| EPI_ISL_729106 | Viollier AG | Department of Biosystems Science and Engineering, ETH Zürich | Chaoran Chen et al |
| EPI_ISL_5655526 | King Chulalongkorn Memorial Hospital | Thai Red Cross Emerging Infectious Diseases Clinical Center and Faculty of Medicine, Chulalongkorn University | Opass Putcharoen et al |

| | | | |
|---|---|---|---|
| EPI_ISL_1130654 | Viollier AG | Department of Biosystems Science and Engineering, ETH Zürich | Christian Beisel et al |
| EPI_ISL_3717417 | Valais Hospital, Central Institute | Valais Hospital, Central Institute | Alexis Dumoulin et al |
| EPI_ISL_3613461 | Ministry of Health Turkey | Ministry of Health Turkey | Fatma Bayrakdar et al |
| EPI_ISL_3386469 | Ministry of Health Turkey | Ministry of Health Turkey | Fatma Bayrakdar et al |
| EPI_ISL_528380 | Clinical Virology | Clinical Bacteriology | Madlen Stange et al |
| EPI_ISL_6858409 | Medica | Institute of Medical Virology | Daniel Ehrsam et al |
| EPI_ISL_1408069 | Viollier AG | Department of Biosystems Science and Engineering, ETH Zürich | Chaoran Chen et al |
| EPI_ISL_1361417 | Viollier AG | Department of Biosystems Science and Engineering, ETH Zürich | Christian Beisel et al |
| EPI_ISL_1119371 | Viollier AG | Department of Biosystems Science and Engineering, ETH Zürich | Chaoran Chen et al |
| EPI_ISL_3247306 | Clinical Virology | Clinical Bacteriology | Tim Roloff et al |
| EPI_ISL_6720669 | Viollier AG | Department of Biosystems Science and Engineering, ETH Zürich | Andrea Patrizia Salzmann et al |
| EPI_ISL_4170070 | Regional Medical Sciences Center 11/1 Phuket | National Institute of Health, Department of Medical Sciences, Ministry of Public Health, Thailand | Pilailuk Okada et al |
| EPI_ISL_5884762 | Public Health Service Center 53 | Division of Genomic Medicine and Innovation support,Department of Medical Sciences, Ministry of Public Health, Thailand | Surakameth Mahasirimongkol et al |
| EPI_ISL_4025702 | Stadtspital Triemli | Institute of Medical Virology | Daniel Ehrsam et al |
| EPI_ISL_2086805 | Spital MŠnnedorf AG | Institute of Medical Virology | Daniel Ehrsam et al |
| EPI_ISL_5865300 | Viollier AG | Department of Biosystems Science and Engineering, ETH Zürich | Chaoran Chen et al |
| EPI_ISL_1273457 | Clinical Virology | Clinical Bacteriology | Tim Roloff et al |
| EPI_ISL_5644778 | Center for Laboratory Medicine | Center for Laboratory Medicine | Yannick Gerth et al |
| EPI_ISL_6720388 | Viollier AG | Department of Biosystems Science and Engineering, ETH Zürich | Andrea Patrizia Salzmann et al |
| EPI_ISL_1171961 | Ministry of Health Turkey | Ministry of Health Turkey | Fatma Bayrakdar et al |
| EPI_ISL_6971472 | Stadtspital Triemli | Institute of Medical Virology, University of Zurich | Verena Kufner et al |
| EPI_ISL_6824342 | Viollier AG | Department of Biosystems Science and Engineering, ETH Zürich | Andrea Patrizia Salzmann et al |
| EPI_ISL_3032961 | Ministry of Health Turkey | Ministry of Health Turkey | Fatma Bayrakdar et al |
| EPI_ISL_5533232 | Bioanalytica SA | Genesupport | Tanguy ARAUD et al |
| EPI_ISL_2403515 | Ministry of Health Turkey | Ministry of Health Turkey | Fatma Bayrakdar et al |
| EPI_ISL_7042161 | Clinical Virology | Clinical Bacteriology, University Hospital Basel | Tim Roloff et al |
| EPI_ISL_1533233 | University Hospitals of Geneva, Laboratory of Virology | HUG, Laboratory of Virology and the Health2030 Genome Center | Samuel Cordey et al |
| EPI_ISL_3128743 | Stadtspital Triemli | Institute of Medical Virology | Daniel Ehrsam et al |
| EPI_ISL_5327730 | Kantonsspital Winterthur | Institute of Medical Virology | Daniel Ehrsam et al |
| EPI_ISL_4368857 | Valais Hospital, Central Institute | Valais Hospital, Central Institute | Alexis Dumoulin et al |
| EPI_ISL_1598280 | Viollier AG | Department of Biosystems Science and Engineering, ETH Zürich | Christian Beisel et al |

| | | | |
|---|---|---|---|
| EPI_ISL_1293281 | Ministry of Health Turkey | Ministry of Health Turkey | Fatma Bayrakdar et al |
| EPI_ISL_796586 | Viollier AG | Department of Biosystems Science and Engineering, ETH Zürich | Chaoran Chen et al |
| EPI_ISL_3566387 | Ministry of Health Turkey | Ministry of Health Turkey | Fatma Bayrakdar et al |
| EPI_ISL_3854832 | Viollier AG | Department of Biosystems Science and Engineering, ETH Zürich | Chaoran Chen et al |
| EPI_ISL_2484659 | UniversitätsSpital Zürich | Institute of Medical Virology | Verena Kufner et al |
| EPI_ISL_1361271 | Viollier AG | Department of Biosystems Science and Engineering, ETH Zürich | Chaoran Chen et al |
| EPI_ISL_4368714 | Valais Hospital, Central Institute | Valais Hospital, Central Institute | Alexis Dumoulin et al |
| EPI_ISL_1002111 | Viollier AG | Department of Biosystems Science and Engineering, ETH Zürich | Chaoran Chen et al |
| EPI_ISL_3549440 | Ministry of Health Turkey | Ministry of Health Turkey | Fatma Bayrakdar et al |
| EPI_ISL_5841635 | Rothen Medizinische Laboratorien AG | Clinical Bacteriology | Tim Roloff et al |
| EPI_ISL_2449913 | UniversitŠtsSpital ZŸrich | Institute of Medical Virology | Daniel Ehrsam et al |
| EPI_ISL_1388583 | University Hospital Basel, Clinical Virology | University Hospital Basel, Clinical Bacteriology | Tim Roloff et al |
| EPI_ISL_7062590 | Dr. Risch Ostschweiz AG | Dr Risch Laboratory | Nadia Wohlwend et al |
| EPI_ISL_6334694 | Regional Medical Sciences Center 12 Songkhla | Division of Genomic Medicine and Innovation support,Department of Medical Sciences, Ministry of Public Health, Thailand | Surakameth Mahasirimongkol et al |
| EPI_ISL_3717386 | Valais Hospital, Central Institute | Valais Hospital, Central Institute | Alexis Dumoulin et al |
| EPI_ISL_4077427 | University Hospitals of Geneva, Laboratory of Virology | HUG, Laboratory of Virology and the Health2030 Genome Center | Samuel Cordey et al |
| EPI_ISL_6890525 | Viollier AG | Department of Biosystems Science and Engineering, ETH Zürich | Christian Beisel et al |
| EPI_ISL_3522986 | Fulgent Genetics | Centers for Disease Control and Prevention Division of Viral Diseases, Pathogen Discovery | Dakota Howard et al |
| EPI_ISL_4410458 | Arizona State University | Arizona State University | Ajeet Bains et al |
| EPI_ISL_3593474 | Ministry of Health Turkey | Ministry of Health Turkey | Fatma Bayrakdar et al |
| EPI_ISL_3655250 | Ministry of Health Turkey | Ministry of Health Turkey | Fatma Bayrakdar et al |
| EPI_ISL_4537311 | Laboratory Corporation of America | Centers for Disease Control and Prevention Division of Viral Diseases, Pathogen Discovery | Dakota Howard et al |
| EPI_ISL_4008057 | Arkansas Public Health Laboratory, Arkansas Department of Health | University of Minnesota Genomics Center | Daryl M. Gohl et al |
| EPI_ISL_3256739 | Ministry of Health Turkey | Ministry of Health Turkey | Fatma Bayrakdar et al |
| EPI_ISL_6405974 | Arkansas Public Health Laboratory, Arkansas Department of Health | University of Minnesota Genomics Center | Daryl M. Gohl et al |
| EPI_ISL_4091117 | Arizona State University | Arizona State University | Ajeet Bains et al |
| EPI_ISL_3649486 | Ministry of Health Turkey | Ministry of Health Turkey | Fatma Bayrakdar et al |
| EPI_ISL_3405745 | Ministry of Health Turkey | Ministry of Health Turkey | Fatma Bayrakdar et al |
| EPI_ISL_3648119 | Ministry of Health Turkey | Ministry of Health Turkey | Fatma Bayrakdar et al |

| | | | |
|---|---|---|---|
| EPI_ISL_1016918 | Arizona State Public Health Laboratory | Arizona State Public Health Laboratory | Trung Huynh et al |
| EPI_ISL_3554111 | Ministry of Health Turkey | Ministry of Health Turkey | Fatma Bayrakdar et al |
| EPI_ISL_4240185 | Laboratory Corporation of America | Centers for Disease Control and Prevention Division of Viral Diseases, Pathogen Discovery | Dakota Howard et al |
| EPI_ISL_4240674 | Laboratory Corporation of America | Centers for Disease Control and Prevention Division of Viral Diseases, Pathogen Discovery | Dakota Howard et al |
| EPI_ISL_3256761 | Ministry of Health Turkey | Ministry of Health Turkey | Fatma Bayrakdar et al |
| EPI_ISL_3612330 | Ministry of Health Turkey | Ministry of Health Turkey | Fatma Bayrakdar et al |
| EPI_ISL_4576096 | Ministry of Health Turkey | Ministry of Health Turkey | Fatma Bayrakdar et al |
| EPI_ISL_6934665 | Ministry of Health Turkey | Ministry of Health Turkey | Fatma Bayrakdar et al |
| EPI_ISL_3317523 | Ministry of Health Turkey | Ministry of Health Turkey | Fatma Bayrakdar et al |
| EPI_ISL_3415006 | Ministry of Health Turkey | Ministry of Health Turkey | Fatma Bayrakdar et al |
| EPI_ISL_3649726 | Ministry of Health Turkey | Ministry of Health Turkey | Fatma Bayrakdar et al |
| EPI_ISL_877273 | Eurofins Diatherix | Hudsonalpha Genome Sequencing Center | Jane Grimwood et al |
| EPI_ISL_3306730 | Ministry of Health Turkey | Ministry of Health Turkey | Fatma Bayrakdar et al |
| EPI_ISL_6059564 | Aegis Sciences Corporation | Centers for Disease Control and Prevention Division of Viral Diseases, Pathogen Discovery | Dakota Howard et al |
| EPI_ISL_3417258 | Ministry of Health Turkey | Ministry of Health Turkey | Fatma Bayrakdar et al |
| EPI_ISL_3586425 | Ministry of Health Turkey | Ministry of Health Turkey | Fatma Bayrakdar et al |
| EPI_ISL_4266615 | Ministry of Health Turkey | Ministry of Health Turkey | Fatma Bayrakdar et al |
| EPI_ISL_4008283 | Arkansas Public Health Laboratory, Arkansas Department of Health | University of Minnesota Genomics Center | Daryl M. Gohl et al |
| EPI_ISL_4166500 | Laboratory Corporation of America | Centers for Disease Control and Prevention Division of Viral Diseases, Pathogen Discovery | Dakota Howard et al |
| EPI_ISL_3760719 | Arkansas Children's Hospital | Center for Global Health, University of New Mexico Health Sciences Center | Daryl Domman et al |
| EPI_ISL_6132128 | Ministry of Health Turkey | Ministry of Health Turkey | Fatma Bayrakdar et al |
| EPI_ISL_5067074 | Ministry of Health Turkey | Ministry of Health Turkey | Fatma Bayrakdar et al |
| EPI_ISL_3559926 | Ministry of Health Turkey | Ministry of Health Turkey | Fatma Bayrakdar et al |
| EPI_ISL_5967031 | Ministry of Health Turkey | Ministry of Health Turkey | Fatma Bayrakdar et al |
| EPI_ISL_3589723 | Ministry of Health Turkey | Ministry of Health Turkey | Fatma Bayrakdar et al |
| EPI_ISL_4892557 | Ministry of Health Turkey | Ministry of Health Turkey | Fatma Bayrakdar et al |
| EPI_ISL_2686747 | Fulgent Genetics | Centers for Disease Control and Prevention Division of Viral Diseases, Pathogen Discovery | Dakota Howard et al |
| EPI_ISL_5427246 | Ministry of Health Turkey | Ministry of Health Turkey | Fatma Bayrakdar et al |
| EPI_ISL_3324771 | Aegis Sciences Corporation | Centers for Disease Control and Prevention Division of Viral Diseases, Pathogen Discovery | Dakota Howard et al |
| EPI_ISL_4812189 | Laboratory Corporation of America | Centers for Disease Control and Prevention Division of Viral Diseases, Pathogen Discovery | Dakota Howard et al |
| EPI_ISL_4744272 | Arizona State University | Arizona State University | Ajeet Bains et al |
| EPI_ISL_3412796 | Ministry of Health Turkey | Ministry of Health Turkey | Fatma Bayrakdar et al |

| | | | |
|---|---|---|---|
| EPI_ISL_4410663 | Arizona State University | Arizona State University | Ajeet Bains et al |
| EPI_ISL_3401340 | Ministry of Health Turkey | Ministry of Health Turkey | Fatma Bayrakdar et al |
| EPI_ISL_3906188 | Arizona State University | Arizona State University | Ajeet Bains et al |
| EPI_ISL_3541665 | Ministry of Health Turkey | Ministry of Health Turkey | Fatma Bayrakdar et al |
| EPI_ISL_3414355 | Ministry of Health Turkey | Ministry of Health Turkey | Fatma Bayrakdar et al |
| EPI_ISL_6067126 | Aegis Sciences Corporation | Centers for Disease Control and Prevention Division of Viral Diseases, Pathogen Discovery | Dakota Howard et al |
| EPI_ISL_4591459 | Ministry of Health Turkey | Ministry of Health Turkey | Fatma Bayrakdar et al |
| EPI_ISL_877317 | Eurofins Diatherix | Hudsonalpha Genome Sequencing Center | Jane Grimwood et al |
| EPI_ISL_3406932 | Ministry of Health Turkey | Ministry of Health Turkey | Fatma Bayrakdar et al |
| EPI_ISL_4513182 | Arizona State University | Arizona State University | Ajeet Bains et al |
| EPI_ISL_3867137 | Arkansas Public Health Laboratory, Arkansas Department of Health | University of Minnesota Genomics Center | Daryl M. Gohl et al |
| EPI_ISL_3493305 | Arizona State University | Arizona State University | Ajeet Bains et al |
| EPI_ISL_3540506 | Ministry of Health Turkey | Ministry of Health Turkey | Fatma Bayrakdar et al |
| EPI_ISL_2597732 | Arizona State Public Health Laboratory | Arizona State Public Health Laboratory | Trung Huynh et al |
| EPI_ISL_3540753 | Ministry of Health Turkey | Ministry of Health Turkey | Fatma Bayrakdar et al |
| EPI_ISL_6842432 | Ministry of Health Turkey | Ministry of Health Turkey | Fatma Bayrakdar et al |
| EPI_ISL_6816208 | Ministry of Health Turkey | Ministry of Health Turkey | Fatma Bayrakdar et al |
| EPI_ISL_1094897 | AL Dept. of Public Health Bureau of Clinical Laboratories | Respiratory Viruses Branch, Division of Viral Diseases, Centers for Disease Control and Prevention | Krista Queen et al |
| EPI_ISL_3688437 | Ministry of Health Turkey | Ministry of Health Turkey | Fatma Bayrakdar et al |
| EPI_ISL_3474021 | Arkansas Public Health Laboratory, Arkansas Department of Health | University of Minnesota Genomics Center | Daryl M. Gohl et al |
| EPI_ISL_3647364 | Ministry of Health Turkey | Ministry of Health Turkey | Fatma Bayrakdar et al |
| EPI_ISL_3401019 | Ministry of Health Turkey | Ministry of Health Turkey | Fatma Bayrakdar et al |
| EPI_ISL_5966961 | Ministry of Health Turkey | Ministry of Health Turkey | Fatma Bayrakdar et al |
| EPI_ISL_3403091 | Ministry of Health Turkey | Ministry of Health Turkey | Fatma Bayrakdar et al |
| EPI_ISL_877369 | Eurofins Diatherix | Hudsonalpha Genome Sequencing Center | Jane Grimwood et al |
| EPI_ISL_5059204 | Arizona State University | Arizona State University | Ajeet Bains et al |
| EPI_ISL_6121401 | Ministry of Health Turkey | Ministry of Health Turkey | Fatma Bayrakdar et al |
| EPI_ISL_3553490 | Arkansas Children's Hospital | Center for Global Health, University of New Mexico Health Sciences Center | Daryl Domman et al |
| EPI_ISL_3585314 | Ministry of Health Turkey | Ministry of Health Turkey | Fatma Bayrakdar et al |
| EPI_ISL_4825614 | Quest Diagnostics Incorporated | Centers for Disease Control and Prevention Division of Viral Diseases, Pathogen Discovery | Dakota Howard et al |
| EPI_ISL_2761688 | Pathology, University of Alabama at Birmingham | Pathology, University of Alabama at Birmingham | Moates et al |
| EPI_ISL_3393254 | Ministry of Health Turkey | Ministry of Health Turkey | Fatma Bayrakdar et al |
| EPI_ISL_5852936 | Ministry of Health Turkey | Ministry of Health Turkey | Fatma Bayrakdar et al |
| EPI_ISL_6074876 | Arizona State University | Arizona State University | Ajeet Bains et al |
| EPI_ISL_877403 | Eurofins Diatherix | Hudsonalpha Genome Sequencing Center | Jane Grimwood et al |

| | | | |
|---|---|---|---|
| EPI_ISL_3577185 | Ministry of Health Turkey | Ministry of Health Turkey | Fatma Bayrakdar et al |
| EPI_ISL_4206492 | Pathology, UAB | Pathology, UAB | Moates et al |
| EPI_ISL_6572765 | Arizona State University | Arizona State University | Steven C. Holland et al |
| EPI_ISL_4557409 | Arizona State University | Arizona State University | Ajeet Bains et al |
| EPI_ISL_2455422 | Alaska State Virology Laboratory | Alaska State Virology Laboratory | Stephanie DeRonde et al |
| EPI_ISL_3943747 | Mobile County Health Department | Synergy Laboratories | Megan Cornwell et al |
| EPI_ISL_3673569 | Ministry of Health Turkey | Ministry of Health Turkey | Fatma Bayrakdar et al |
| EPI_ISL_4809542 | Laboratory Corporation of America | Centers for Disease Control and Prevention Division of Viral Diseases, Pathogen Discovery | Dakota Howard et al |
| EPI_ISL_2672031 | TGen North | TGen North | Jolene Bowers et al |
| EPI_ISL_2000973 | TGen North | TGen North | "Jolene Bowers et al |
| EPI_ISL_3833041 | TGen North | TGen North | "Jolene Bowers et al |
| EPI_ISL_3349830 | Fulgent Genetics | Centers for Disease Control and Prevention Division of Viral Diseases, Pathogen Discovery | Dakota Howard et al |
| EPI_ISL_3863149 | Aegis Sciences Corporation | Centers for Disease Control and Prevention Division of Viral Diseases, Pathogen Discovery | Dakota Howard et al |
| EPI_ISL_5867656 | Scripps Medical Laboratory | Andersen lab at Scripps Research | SEARCH Alliance San Diego with Michael Quigley et al |
| EPI_ISL_3302064 | Fulgent Genetics | Centers for Disease Control and Prevention Division of Viral Diseases, Pathogen Discovery | Dakota Howard et al |
| EPI_ISL_2875964 | Aegis Sciences Corporation | Centers for Disease Control and Prevention Division of Viral Diseases, Pathogen Discovery | Dakota Howard et al |
| EPI_ISL_1445373 | Aegis Sciences Corporation | Centers for Disease Control and Prevention Division of Viral Diseases, Pathogen Discovery | Dakota Howard et al |
| EPI_ISL_2672201 | TGen North | TGen North | Jolene Bowers et al |
| EPI_ISL_6402586 | TGen North | TGen North | Jolene Bowers et al |
| EPI_ISL_1458027 | Fulgent Genetics | Centers for Disease Control and Prevention Division of Viral Diseases, Pathogen Discovery | Dakota Howard et al |
| EPI_ISL_3331468 | Fulgent Genetics | Centers for Disease Control and Prevention Division of Viral Diseases, Pathogen Discovery | Dakota Howard et al |
| EPI_ISL_3305490 | Aegis Sciences Corporation | Centers for Disease Control and Prevention Division of Viral Diseases, Pathogen Discovery | Dakota Howard et al |
| EPI_ISL_4369708 | Fulgent Genetics | Centers for Disease Control and Prevention Division of Viral Diseases, Pathogen Discovery | Dakota Howard et al |
| EPI_ISL_3661940 | TGen North | TGen North | Jolene Bowers et al |
| EPI_ISL_4809539 | Laboratory Corporation of America | Centers for Disease Control and Prevention Division of Viral Diseases, Pathogen Discovery | Dakota Howard et al |

| | | | |
|---|---|---|---|
| EPI_ISL_3813111 | Aegis Sciences Corporation | Centers for Disease Control and Prevention Division of Viral Diseases, Pathogen Discovery | Dakota Howard et al |
| EPI_ISL_2687277 | Aegis Sciences Corporation | Centers for Disease Control and Prevention Division of Viral Diseases, Pathogen Discovery | Dakota Howard et al |
| EPI_ISL_3210909 | Fulgent Genetics | Centers for Disease Control and Prevention Division of Viral Diseases, Pathogen Discovery | Dakota Howard et al |
| EPI_ISL_3850697 | Aegis Sciences Corporation | Centers for Disease Control and Prevention Division of Viral Diseases, Pathogen Discovery | Dakota Howard et al |
| EPI_ISL_6381227 | Aegis Sciences Corporation | Centers for Disease Control and Prevention Division of Viral Diseases, Pathogen Discovery | Dakota Howard et al |
| EPI_ISL_3826127 | TGen North | TGen North | Jolene Bowers et al |
| EPI_ISL_6064338 | Aegis Sciences Corporation | Centers for Disease Control and Prevention Division of Viral Diseases, Pathogen Discovery | Dakota Howard et al |
| EPI_ISL_3850318 | Aegis Sciences Corporation | Centers for Disease Control and Prevention Division of Viral Diseases, Pathogen Discovery | Dakota Howard et al |
| EPI_ISL_1556998 | Fulgent Genetics | Centers for Disease Control and Prevention Division of Viral Diseases, Pathogen Discovery | Dakota Howard et al |
| EPI_ISL_1743640 | Arizona State University | Arizona State University | Peter T. Skidmore et al |
| EPI_ISL_5874682 | Laboratory Corporation of America | Centers for Disease Control and Prevention Division of Viral Diseases, Pathogen Discovery | Dakota Howard et al |
| EPI_ISL_1560740 | Aegis Sciences Corporation | Centers for Disease Control and Prevention Division of Viral Diseases, Pathogen Discovery | Dakota Howard et al |
| EPI_ISL_426567 | AZ SPHL, Arizona Department of Health Services | TGen North | Jolene Bowers et al |
| EPI_ISL_1561866 | Aegis Sciences Corporation | Centers for Disease Control and Prevention Division of Viral Diseases, Pathogen Discovery | Dakota Howard et al |
| EPI_ISL_3457693 | Fulgent Genetics | Centers for Disease Control and Prevention Division of Viral Diseases, Pathogen Discovery | Dakota Howard et al |
| EPI_ISL_3821292 | Aegis Sciences Corporation | Centers for Disease Control and Prevention Division of Viral Diseases, Pathogen Discovery | Dakota Howard et al |
| EPI_ISL_1458920 | Fulgent Genetics | Centers for Disease Control and Prevention Division of Viral Diseases, Pathogen Discovery | Dakota Howard et al |
| EPI_ISL_3218765 | Aegis Sciences Corporation | Centers for Disease Control and Prevention Division of Viral Diseases, Pathogen Discovery | Dakota Howard et al |
| EPI_ISL_4984524 | Aegis Sciences Corporation | Centers for Disease Control and Prevention Division of Viral Diseases, Pathogen Discovery | Dakota Howard et al |

| | | | |
|---|---|---|---|
| EPI_ISL_636143 | San Diego County Public Health Laboratory | Andersen lab at Scripps Research | SEARCH Alliance San Diego with Tracy Basler et al |
| EPI_ISL_4179837 | Laboratory Corporation of America | Centers for Disease Control and Prevention Division of Viral Diseases, Pathogen Discovery | Dakota Howard et al |
| EPI_ISL_3847460 | Aegis Sciences Corporation | Centers for Disease Control and Prevention Division of Viral Diseases, Pathogen Discovery | Dakota Howard et al |
| EPI_ISL_4361943 | Aegis Sciences Corporation | Centers for Disease Control and Prevention Division of Viral Diseases, Pathogen Discovery | Dakota Howard et al |
| EPI_ISL_3460978 | Fulgent Genetics | Centers for Disease Control and Prevention Division of Viral Diseases, Pathogen Discovery | Dakota Howard et al |
| EPI_ISL_4211509 | Fulgent Genetics | Centers for Disease Control and Prevention Division of Viral Diseases, Pathogen Discovery | Dakota Howard et al |
| EPI_ISL_4961381 | Fulgent Genetics | Centers for Disease Control and Prevention Division of Viral Diseases, Pathogen Discovery | Dakota Howard et al |
| EPI_ISL_3209925 | Fulgent Genetics | Centers for Disease Control and Prevention Division of Viral Diseases, Pathogen Discovery | Dakota Howard et al |
| EPI_ISL_3088559 | TGen North | TGen North | "Jolene Bowers et al |
| EPI_ISL_3815282 | Aegis Sciences Corporation | Centers for Disease Control and Prevention Division of Viral Diseases, Pathogen Discovery | Dakota Howard et al |
| EPI_ISL_3609316 | Fulgent Genetics | Centers for Disease Control and Prevention Division of Viral Diseases, Pathogen Discovery | Dakota Howard et al |
| EPI_ISL_3149956 | Arizona State University | Arizona State University | Ajeet Bains et al |
| EPI_ISL_5553525 | Aegis Sciences Corporation | Centers for Disease Control and Prevention Division of Viral Diseases, Pathogen Discovery | Dakota Howard et al |
| EPI_ISL_3110505 | Fulgent Genetics | Centers for Disease Control and Prevention Division of Viral Diseases, Pathogen Discovery | Dakota Howard et al |
| EPI_ISL_4987249 | Aegis Sciences Corporation | Centers for Disease Control and Prevention Division of Viral Diseases, Pathogen Discovery | Dakota Howard et al |
| EPI_ISL_6063101 | Aegis Sciences Corporation | Centers for Disease Control and Prevention Division of Viral Diseases, Pathogen Discovery | Dakota Howard et al |
| EPI_ISL_3519711 | Laboratory Corporation of America | Centers for Disease Control and Prevention Division of Viral Diseases, Pathogen Discovery | Dakota Howard et al |
| EPI_ISL_3208700 | Fulgent Genetics | Centers for Disease Control and Prevention Division of Viral Diseases, Pathogen Discovery | Dakota Howard et al |
| EPI_ISL_696051 | Sonora Quest Laboratories, Laboratory Sciences of Arizona | TGen North | Jolene Bowers et al |

| | | | |
|---|---|---|---|
| EPI_ISL_1614903 | Fulgent Genetics | Centers for Disease Control and Prevention Division of Viral Diseases, Pathogen Discovery | Dakota Howard et al |
| EPI_ISL_6649872 | Aegis Sciences Corporation | Centers for Disease Control and Prevention Division of Viral Diseases, Pathogen Discovery | Dakota Howard et al |
| EPI_ISL_3815065 | Aegis Sciences Corporation | Centers for Disease Control and Prevention Division of Viral Diseases, Pathogen Discovery | Dakota Howard et al |
| EPI_ISL_2153588 | Arizona State University | Arizona State University | Peter T. Skidmore et al |
| EPI_ISL_6555095 | Aegis Sciences Corporation | Centers for Disease Control and Prevention Division of Viral Diseases, Pathogen Discovery | Dakota Howard et al |
| EPI_ISL_3396766 | Quest Diagnostics Incorporated | Centers for Disease Control and Prevention Division of Viral Diseases, Pathogen Discovery | Dakota Howard et al |
| EPI_ISL_4982942 | Aegis Sciences Corporation | Centers for Disease Control and Prevention Division of Viral Diseases, Pathogen Discovery | Dakota Howard et al |
| EPI_ISL_6787223 | Aegis Sciences Corporation | Centers for Disease Control and Prevention Division of Viral Diseases, Pathogen Discovery | Dakota Howard et al |
| EPI_ISL_3812967 | Aegis Sciences Corporation | Centers for Disease Control and Prevention Division of Viral Diseases, Pathogen Discovery | Dakota Howard et al |
| EPI_ISL_6529487 | Aegis Sciences Corporation | Centers for Disease Control and Prevention Division of Viral Diseases, Pathogen Discovery | Dakota Howard et al |
| EPI_ISL_3852228 | Aegis Sciences Corporation | Centers for Disease Control and Prevention Division of Viral Diseases, Pathogen Discovery | Dakota Howard et al |
| EPI_ISL_1458204 | Fulgent Genetics | Centers for Disease Control and Prevention Division of Viral Diseases, Pathogen Discovery | Dakota Howard et al |
| EPI_ISL_6541843 | Aegis Sciences Corporation | Centers for Disease Control and Prevention Division of Viral Diseases, Pathogen Discovery | Dakota Howard et al |
| EPI_ISL_3209900 | Fulgent Genetics | Centers for Disease Control and Prevention Division of Viral Diseases, Pathogen Discovery | Dakota Howard et al |
| EPI_ISL_648920 | San Diego County Public Health Laboratory | Andersen lab at Scripps Research | SEARCH Alliance San Diego with Tracy Basler et al |
| EPI_ISL_3397522 | Fulgent Genetics | Centers for Disease Control and Prevention Division of Viral Diseases, Pathogen Discovery | Dakota Howard et al |
| EPI_ISL_3851038 | Aegis Sciences Corporation | Centers for Disease Control and Prevention Division of Viral Diseases, Pathogen Discovery | Dakota Howard et al |
| EPI_ISL_3349301 | Fulgent Genetics | Centers for Disease Control and Prevention Division of Viral Diseases, Pathogen Discovery | Dakota Howard et al |
| EPI_ISL_1113475 | Sonora Quest Laboratories | TGen North | "Jolene Bowers et al |

| | | | |
|---|---|---|---|
| EPI_ISL_1615414 | Helix/Illumina | Centers for Disease Control and Prevention Division of Viral Diseases, Pathogen Discovery | Dakota Howard et al |
| EPI_ISL_1296883 | Sonora Quest Laboratories | TGen North | "Jolene Bowers et al |
| EPI_ISL_6170006 | Aegis Sciences Corporation | Centers for Disease Control and Prevention Division of Viral Diseases, Pathogen Discovery | Dakota Howard et al |
| EPI_ISL_1481571 | Fulgent Genetics | Centers for Disease Control and Prevention Division of Viral Diseases, Pathogen Discovery | Dakota Howard et al |
| EPI_ISL_1458527 | Fulgent Genetics | Centers for Disease Control and Prevention Division of Viral Diseases, Pathogen Discovery | Dakota Howard et al |
| EPI_ISL_1839217 | Fulgent Genetics | Centers for Disease Control and Prevention Division of Viral Diseases, Pathogen Discovery | Dakota Howard et al |
| EPI_ISL_3853864 | Aegis Sciences Corporation | Centers for Disease Control and Prevention Division of Viral Diseases, Pathogen Discovery | Dakota Howard et al |
| EPI_ISL_3430554 | Aegis Sciences Corporation | Centers for Disease Control and Prevention Division of Viral Diseases, Pathogen Discovery | Dakota Howard et al |
| EPI_ISL_2861112 | Arizona State University | Arizona State University | Ajeet Bains et al |
| EPI_ISL_1091746 | US Air Force School of Aerospace Medicine | US Air Force School of Aerospace Medicine | Anthony Fries et al |
| EPI_ISL_6065952 | Aegis Sciences Corporation | Centers for Disease Control and Prevention Division of Viral Diseases, Pathogen Discovery | Dakota Howard et al |
| EPI_ISL_3397738 | Fulgent Genetics | Centers for Disease Control and Prevention Division of Viral Diseases, Pathogen Discovery | Dakota Howard et al |
| EPI_ISL_1365550 | Arizona State University | Arizona State University | Peter T. Skidmore et al |
| EPI_ISL_2785927 | Fulgent Genetics | Centers for Disease Control and Prevention Division of Viral Diseases, Pathogen Discovery | Dakota Howard et al |
| EPI_ISL_3609356 | Fulgent Genetics | Centers for Disease Control and Prevention Division of Viral Diseases, Pathogen Discovery | Dakota Howard et al |
| EPI_ISL_3658494 | Fulgent Genetics | Centers for Disease Control and Prevention Division of Viral Diseases, Pathogen Discovery | Dakota Howard et al |
| EPI_ISL_4412832 | San Diego County Public Health Laboratory | Andersen lab at Scripps Research | SEARCH Alliance San Diego with Ashleigh Murphy et al |
| EPI_ISL_5870839 | Laboratory Corporation of America | Centers for Disease Control and Prevention Division of Viral Diseases, Pathogen Discovery | Dakota Howard et al |
| EPI_ISL_2919490 | TGen North | TGen North | "Jolene Bowers et al |
| EPI_ISL_6069412 | Aegis Sciences Corporation | Centers for Disease Control and Prevention Division of Viral Diseases, Pathogen Discovery | Dakota Howard et al |

| | | | |
|---|---|---|---|
| EPI_ISL_5551144 | Aegis Sciences Corporation | Centers for Disease Control and Prevention Division of Viral Diseases, Pathogen Discovery | Dakota Howard et al |
| EPI_ISL_2873635 | Fulgent Genetics | Centers for Disease Control and Prevention Division of Viral Diseases, Pathogen Discovery | Dakota Howard et al |
| EPI_ISL_3685602 | Laboratory Corporation of America | Centers for Disease Control and Prevention Division of Viral Diseases, Pathogen Discovery | Dakota Howard et al |
| EPI_ISL_3348074 | Fulgent Genetics | Centers for Disease Control and Prevention Division of Viral Diseases, Pathogen Discovery | Dakota Howard et al |
| EPI_ISL_2484117 | Arizona State University | Arizona State University | Peter T. Skidmore et al |
| EPI_ISL_962800 | San Diego County Public Health Laboratory | Andersen lab at Scripps Research | SEARCH Alliance San Diego with Tracy Basler et al |
| EPI_ISL_3316093 | CDPH VBL | California Department of Public Health | Emily Smith on behalf of CDPH-COVIDNet et al |
| EPI_ISL_5164914 | Laboratory Corporation of America | Centers for Disease Control and Prevention Division of Viral Diseases, Pathogen Discovery | Dakota Howard et al |
| EPI_ISL_1515020 | Laboratory Corporation of America | Centers for Disease Control and Prevention Division of Viral Diseases, Pathogen Discovery | Dakota Howard et al |
| EPI_ISL_6120606 | Fulgent Genetics | Centers for Disease Control and Prevention Division of Viral Diseases, Pathogen Discovery | Dakota Howard et al |
| EPI_ISL_3304725 | CDPH VBL | California Department of Public Health | Emily Smith on behalf of CDPH-COVIDNet et al |
| EPI_ISL_6119614 | Fulgent Genetics | Centers for Disease Control and Prevention Division of Viral Diseases, Pathogen Discovery | Dakota Howard et al |
| EPI_ISL_4203952 | Fulgent Genetics | Centers for Disease Control and Prevention Division of Viral Diseases, Pathogen Discovery | Dakota Howard et al |
| EPI_ISL_3637500 | CDPH VBL | California Department of Public Health | Emily Smith on behalf of CDPH-COVIDNet et al |
| EPI_ISL_6341661 | Fulgent Genetics | Centers for Disease Control and Prevention Division of Viral Diseases, Pathogen Discovery | Dakota Howard et al |
| EPI_ISL_5442270 | CDPH VBL | California Department of Public Health | Emily Smith on behalf of CDPH-COVIDNet et al |
| EPI_ISL_3685754 | Laboratory Corporation of America | Centers for Disease Control and Prevention Division of Viral Diseases, Pathogen Discovery | Dakota Howard et al |
| EPI_ISL_3518281 | Laboratory Corporation of America | Centers for Disease Control and Prevention Division of Viral Diseases, Pathogen Discovery | Dakota Howard et al |
| EPI_ISL_6341182 | Fulgent Genetics | Centers for Disease Control and Prevention Division of Viral Diseases, Pathogen Discovery | Dakota Howard et al |

| | | | |
|---|---|---|---|
| EPI_ISL_3101697 | California Department of Public Health Valencia Branch Laboratory (CDPH VBL) | California Department of Public Health | CDPH-COVIDNet et al |
| EPI_ISL_1668789 | Quest Diagnostics Incorporated | Centers for Disease Control and Prevention Division of Viral Diseases, Pathogen Discovery | Dakota Howard et al |
| EPI_ISL_5166145 | Laboratory Corporation of America | Centers for Disease Control and Prevention Division of Viral Diseases, Pathogen Discovery | Dakota Howard et al |
| EPI_ISL_3660139 | Fulgent Genetics | Centers for Disease Control and Prevention Division of Viral Diseases, Pathogen Discovery | Dakota Howard et al |
| EPI_ISL_5376949 | Fulgent Genetics | Centers for Disease Control and Prevention Division of Viral Diseases, Pathogen Discovery | Dakota Howard et al |
| EPI_ISL_3753806 | Fulgent Genetics | Centers for Disease Control and Prevention Division of Viral Diseases, Pathogen Discovery | Dakota Howard et al |
| EPI_ISL_6654605 | Fulgent Genetics | Centers for Disease Control and Prevention Division of Viral Diseases, Pathogen Discovery | Dakota Howard et al |
| EPI_ISL_2796688 | CDPH VBL | California Department of Public Health | CDPH-COVIDNet et al |
| EPI_ISL_6405713 | Helix | Centers for Disease Control and Prevention Division of Viral Diseases, Pathogen Discovery | Dakota Howard et al |
| EPI_ISL_6902737 | Helix | Centers for Disease Control and Prevention Division of Viral Diseases, Pathogen Discovery | Dakota Howard et al |
| EPI_ISL_3659950 | Fulgent Genetics | Centers for Disease Control and Prevention Division of Viral Diseases, Pathogen Discovery | Dakota Howard et al |
| EPI_ISL_1253001 | Quest Diagnostics Incorporated | Centers for Disease Control and Prevention Division of Viral Diseases, Pathogen Discovery | Peter W. Cook et al |
| EPI_ISL_4567758 | Fulgent Genetics | Centers for Disease Control and Prevention Division of Viral Diseases, Pathogen Discovery | Dakota Howard et al |
| EPI_ISL_1990924 | Helix/Illumina | Centers for Disease Control and Prevention Division of Viral Diseases, Pathogen Discovery | Dakota Howard et al |
| EPI_ISL_6626914 | Fulgent Genetics | Centers for Disease Control and Prevention Division of Viral Diseases, Pathogen Discovery | Dakota Howard et al |
| EPI_ISL_6628543 | Fulgent Genetics | Centers for Disease Control and Prevention Division of Viral Diseases, Pathogen Discovery | Dakota Howard et al |
| EPI_ISL_4960023 | Ambry Genetics Corporation | Fulgent Genetics | Harry Gao et al |
| EPI_ISL_2976939 | CDPH VBL | California Department of Public Health | CDPH-COVIDNet et al |
| EPI_ISL_3105560 | CDPH VBL | California Department of Public Health | CDPH-COVIDNet et al |

| | | | |
|---|---|---|---|
| EPI_ISL_4902185 | Fulgent Genetics | Centers for Disease Control and Prevention Division of Viral Diseases, Pathogen Discovery | Dakota Howard et al |
| EPI_ISL_1796250 | Helix/Illumina | Centers for Disease Control and Prevention Division of Viral Diseases, Pathogen Discovery | Dakota Howard et al |
| EPI_ISL_5232545 | Fulgent Genetics | Centers for Disease Control and Prevention Division of Viral Diseases, Pathogen Discovery | Dakota Howard et al |
| EPI_ISL_4035413 | Fulgent Genetics | Centers for Disease Control and Prevention Division of Viral Diseases, Pathogen Discovery | Dakota Howard et al |
| EPI_ISL_2873146 | Laboratory Corporation of America | Centers for Disease Control and Prevention Division of Viral Diseases, Pathogen Discovery | Dakota Howard et al |
| EPI_ISL_3750905 | Laboratory Corporation of America | Centers for Disease Control and Prevention Division of Viral Diseases, Pathogen Discovery | Dakota Howard et al |
| EPI_ISL_6893162 | Fulgent Genetics | Centers for Disease Control and Prevention Division of Viral Diseases, Pathogen Discovery | Dakota Howard et al |
| EPI_ISL_2869265 | Quest Diagnostics Incorporated | Centers for Disease Control and Prevention Division of Viral Diseases, Pathogen Discovery | Dakota Howard et al |
| EPI_ISL_5890185 | Fulgent Genetics | Centers for Disease Control and Prevention Division of Viral Diseases, Pathogen Discovery | Dakota Howard et al |
| EPI_ISL_6071788 | Fulgent Genetics | Centers for Disease Control and Prevention Division of Viral Diseases, Pathogen Discovery | Dakota Howard et al |
| EPI_ISL_6267202 | CDPH VBL | California Department of Public Health | Emily Smith on behalf of CDPH-COVIDNet et al |
| EPI_ISL_3753500 | Fulgent Genetics | Centers for Disease Control and Prevention Division of Viral Diseases, Pathogen Discovery | Dakota Howard et al |
| EPI_ISL_2825567 | CDPH VBL | California Department of Public Health | CDPH-COVIDNet et al |
| EPI_ISL_4465182 | Quest Diagnostics Incorporated | Centers for Disease Control and Prevention Division of Viral Diseases, Pathogen Discovery | Dakota Howard et al |
| EPI_ISL_6071985 | Fulgent Genetics | Centers for Disease Control and Prevention Division of Viral Diseases, Pathogen Discovery | Dakota Howard et al |
| EPI_ISL_5233987 | Fulgent Genetics | Centers for Disease Control and Prevention Division of Viral Diseases, Pathogen Discovery | Dakota Howard et al |
| EPI_ISL_1088508 | Quest Diagnostics Incorporated | Respiratory Viruses Branch, Division of Viral Diseases, Centers for Disease Control and Prevention | Peter W. Cook et al |
| EPI_ISL_6480963 | Helix | Centers for Disease Control and Prevention Division of Viral Diseases, Pathogen Discovery | Dakota Howard et al |

| | | | |
|---|---|---|---|
| EPI_ISL_4357307 | Fulgent Genetics | Centers for Disease Control and Prevention Division of Viral Diseases, Pathogen Discovery | Dakota Howard et al |
| EPI_ISL_5581334 | Fulgent Genetics | Centers for Disease Control and Prevention Division of Viral Diseases, Pathogen Discovery | Dakota Howard et al |
| EPI_ISL_1088627 | Helix / Illumina | Respiratory Viruses Branch, Division of Viral Diseases, Centers for Disease Control and Prevention | Peter W. Cook et al |
| EPI_ISL_2783260 | CDPH VBL | California Department of Public Health | CDPH-COVIDNet et al |
| EPI_ISL_4454652 | Fulgent Genetics | Centers for Disease Control and Prevention Division of Viral Diseases, Pathogen Discovery | Dakota Howard et al |
| EPI_ISL_4827112 | Fulgent Genetics | Centers for Disease Control and Prevention Division of Viral Diseases, Pathogen Discovery | Dakota Howard et al |
| EPI_ISL_2826821 | CDPH VBL | California Department of Public Health | CDPH-COVIDNet et al |
| EPI_ISL_1679529 | Helix/Illumina | Centers for Disease Control and Prevention Division of Viral Diseases, Pathogen Discovery | Dakota Howard et al |
| EPI_ISL_6511297 | Fulgent Genetics | Centers for Disease Control and Prevention Division of Viral Diseases, Pathogen Discovery | Dakota Howard et al |
| EPI_ISL_2930264 | Quest Diagnostics Incorporated | Centers for Disease Control and Prevention Division of Viral Diseases, Pathogen Discovery | Dakota Howard et al |
| EPI_ISL_6340872 | Fulgent Genetics | Centers for Disease Control and Prevention Division of Viral Diseases, Pathogen Discovery | Dakota Howard et al |
| EPI_ISL_6119566 | Fulgent Genetics | Centers for Disease Control and Prevention Division of Viral Diseases, Pathogen Discovery | Dakota Howard et al |
| EPI_ISL_4188060 | Quest Diagnostics Incorporated | Centers for Disease Control and Prevention Division of Viral Diseases, Pathogen Discovery | Dakota Howard et al |
| EPI_ISL_5225859 | Fulgent Genetics | Centers for Disease Control and Prevention Division of Viral Diseases, Pathogen Discovery | Dakota Howard et al |
| EPI_ISL_5978920 | Fulgent Genetics | Centers for Disease Control and Prevention Division of Viral Diseases, Pathogen Discovery | Dakota Howard et al |
| EPI_ISL_3661862 | Fulgent Genetics | Centers for Disease Control and Prevention Division of Viral Diseases, Pathogen Discovery | Dakota Howard et al |
| EPI_ISL_4456157 | Fulgent Genetics | Centers for Disease Control and Prevention Division of Viral Diseases, Pathogen Discovery | Dakota Howard et al |
| EPI_ISL_3750963 | Laboratory Corporation of America | Centers for Disease Control and Prevention Division of Viral Diseases, Pathogen Discovery | Dakota Howard et al |

| | | | |
|---|---|---|---|
| EPI_ISL_5175029 | Laboratory Corporation of America | Centers for Disease Control and Prevention Division of Viral Diseases, Pathogen Discovery | Dakota Howard et al |
| EPI_ISL_5796894 | Fulgent Genetics | Centers for Disease Control and Prevention Division of Viral Diseases, Pathogen Discovery | Dakota Howard et al |
| EPI_ISL_3662125 | Fulgent Genetics | Centers for Disease Control and Prevention Division of Viral Diseases, Pathogen Discovery | Dakota Howard et al |
| EPI_ISL_6906663 | Laboratory Corporation of America | Centers for Disease Control and Prevention Division of Viral Diseases, Pathogen Discovery | Dakota Howard et al |
| EPI_ISL_5092073 | Fulgent Genetics | Centers for Disease Control and Prevention Division of Viral Diseases, Pathogen Discovery | Dakota Howard et al |
| EPI_ISL_4184718 | Fulgent Genetics | Centers for Disease Control and Prevention Division of Viral Diseases, Pathogen Discovery | Dakota Howard et al |
| EPI_ISL_2826281 | CDPH VBL | California Department of Public Health | CDPH-COVIDNet et al |
| EPI_ISL_4244802 | Fulgent Genetics | Centers for Disease Control and Prevention Division of Viral Diseases, Pathogen Discovery | Dakota Howard et al |
| EPI_ISL_4535008 | Fulgent Genetics | Centers for Disease Control and Prevention Division of Viral Diseases, Pathogen Discovery | Dakota Howard et al |
| EPI_ISL_6119722 | Fulgent Genetics | Centers for Disease Control and Prevention Division of Viral Diseases, Pathogen Discovery | Dakota Howard et al |
| EPI_ISL_4028097 | Fulgent Genetics | Centers for Disease Control and Prevention Division of Viral Diseases, Pathogen Discovery | Dakota Howard et al |
| EPI_ISL_5889740 | Fulgent Genetics | Centers for Disease Control and Prevention Division of Viral Diseases, Pathogen Discovery | Dakota Howard et al |
| EPI_ISL_2797502 | CDPH VBL | California Department of Public Health | CDPH-COVIDNet et al |
| EPI_ISL_4456377 | Fulgent Genetics | Centers for Disease Control and Prevention Division of Viral Diseases, Pathogen Discovery | Dakota Howard et al |
| EPI_ISL_5581252 | Fulgent Genetics | Centers for Disease Control and Prevention Division of Viral Diseases, Pathogen Discovery | Dakota Howard et al |
| EPI_ISL_6079387 | Laboratory Corporation of America | Centers for Disease Control and Prevention Division of Viral Diseases, Pathogen Discovery | Dakota Howard et al |
| EPI_ISL_6891825 | Fulgent Genetics | Centers for Disease Control and Prevention Division of Viral Diseases, Pathogen Discovery | Dakota Howard et al |
| EPI_ISL_6081249 | Laboratory Corporation of America | Centers for Disease Control and Prevention Division of Viral Diseases, Pathogen Discovery | Dakota Howard et al |

| | | | |
|---|---|---|---|
| EPI_ISL_6399885 | Quest Diagnostics Incorporated | Centers for Disease Control and Prevention Division of Viral Diseases, Pathogen Discovery | Dakota Howard et al |
| EPI_ISL_4357476 | Fulgent Genetics | Centers for Disease Control and Prevention Division of Viral Diseases, Pathogen Discovery | Dakota Howard et al |
| EPI_ISL_1253164 | Quest Diagnostics Incorporated | Centers for Disease Control and Prevention Division of Viral Diseases, Pathogen Discovery | Peter W. Cook et al |
| EPI_ISL_1193939 | Quest Diagnostics Incorporated | Centers for Disease Control and Prevention Division of Viral Diseases, Pathogen Discovery | Peter W. Cook et al |
| EPI_ISL_6096235 | Fulgent Genetics | Centers for Disease Control and Prevention Division of Viral Diseases, Pathogen Discovery | Dakota Howard et al |
| EPI_ISL_6120185 | Fulgent Genetics | Centers for Disease Control and Prevention Division of Viral Diseases, Pathogen Discovery | Dakota Howard et al |
| EPI_ISL_3754152 | Fulgent Genetics | Centers for Disease Control and Prevention Division of Viral Diseases, Pathogen Discovery | Dakota Howard et al |
| EPI_ISL_4027389 | Fulgent Genetics | Centers for Disease Control and Prevention Division of Viral Diseases, Pathogen Discovery | Dakota Howard et al |
| EPI_ISL_1493960 | Quest Diagnostics Incorporated | Centers for Disease Control and Prevention Division of Viral Diseases, Pathogen Discovery | Peter W. Cook et al |
| EPI_ISL_6892521 | Fulgent Genetics | Centers for Disease Control and Prevention Division of Viral Diseases, Pathogen Discovery | Dakota Howard et al |
| EPI_ISL_5236763 | Scripps Medical Laboratory | Andersen lab at Scripps Research | SEARCH Alliance San Diego with Michael Quigley et al |
| EPI_ISL_1011359 | Fulgent Genetics | Fulgent Genetics | Harry Gao et al |
| EPI_ISL_5442163 | TPMG Regional Laboratory | California Department of Public Health | Emily Smith on behalf of CDPH-COVIDNet et al |
| EPI_ISL_5153396 | CDPH VBL | California Department of Public Health | Emily Smith on behalf of CDPH-COVIDNet et al |
| EPI_ISL_6692586 | Santa Clara Valley Medical Center | Santa Clara County Public Health Laboratory | Santa Clara County Public Health Department et al |
| EPI_ISL_4510757 | CDPH VBL | California Department of Public Health | Emily Smith on behalf of CDPH-COVIDNet et al |
| EPI_ISL_5485846 | Providence Saint John's Health Center | Los Angeles County Public Health Laboratories | P. Hemarajata et al. |
| EPI_ISL_2341428 | CA DPH Viral and Rickettsial Disease Laboratory | Chan-Zuckerberg Biohub | CZB Cliahub Consortium et al |
| EPI_ISL_754927 | California Department of Public Health | California Department of Public Health | CDPH IDLB COVIDNet et al |
| EPI_ISL_1184619 | Santa Clara County Public Health Laboratory | Chan-Zuckerberg Biohub | CZB Cliahub Consortium et al |
| EPI_ISL_2892533 | California Department of Public Health | California Department of Public Health | CDPH IDLB COVIDNet et al |

| | | | |
|---|---|---|---|
| EPI_ISL_5058905 | Torrance Memorial Medical Center | Los Angeles County Public Health Laboratories | P. Hemarajata et al. |
| EPI_ISL_2923667 | CDPH VBL | California Department of Public Health | CDPH-COVIDNet et al |
| EPI_ISL_5015849 | CDPH VBL | California Department of Public Health | Emily Smith on behalf of CDPH-COVIDNet et al |
| EPI_ISL_4508949 | CDPH VBL | California Department of Public Health | Emily Smith on behalf of CDPH-COVIDNet et al |
| EPI_ISL_4640564 | Good Samaritan Hospital | Santa Clara County Public Health Laboratory | Santa Clara County Public Health Department et al |
| EPI_ISL_2391789 | Flow Diagnostics Lab | Los Angeles County Public Health Laboratories | P. Hemarajata et al. |
| EPI_ISL_4268371 | Sharp HealthCare Laboratory | Andersen lab at Scripps Research | SEARCH Alliance San Diego with Aaron Harding et al |
| EPI_ISL_3757200 | Scripps Medical Laboratory | Andersen lab at Scripps Research | SEARCH Alliance San Diego with Michael Quigley et al |
| EPI_ISL_2837992 | California Department of Public Health Valencia Branch Laboratory (CDPH VBL) | California Department of Public Health | CDPH-COVIDNet et al |
| EPI_ISL_625625 | County of San Luis Obispo Public Health Laboratory | Chan-Zuckerberg Biohub | CZB Cliahub Consortium et al |
| EPI_ISL_5016407 | CDPH VBL | California Department of Public Health | Emily Smith on behalf of CDPH-COVIDNet et al |
| EPI_ISL_5696589 | CDPH VBL | California Department of Public Health | Emily Smith on behalf of CDPH-COVIDNet et al |
| EPI_ISL_2858309 | CDPH VBL | California Department of Public Health | CDPH-COVIDNet et al |
| EPI_ISL_6174051 | Orange County Public Health Laboratory | Orange County Public Health Laboratory | Angelica Torres et al |
| EPI_ISL_4926110 | CDPH VBL | California Department of Public Health | Emily Smith on behalf of CDPH-COVIDNet et al |
| EPI_ISL_4707527 | CDPH VBL | California Department of Public Health | Emily Smith on behalf of CDPH-COVIDNet et al |
| EPI_ISL_980976 | Innovative Genomics Institute, UC Berkeley | Innovative Genomics Institute, UC Berkeley | Stacia Wyman et al |
| EPI_ISL_2928421 | Humboldt County Public Health Laboratory | Humboldt County Public Health Laboratory | Jeremy Corrigan et al |
| EPI_ISL_738928 | Orange County Public Health Lab | Chan-Zuckerberg Biohub | CZB Cliahub Consortium et al |
| EPI_ISL_672088 | Santa Clara County Public Health Laboratory | Chan-Zuckerberg Biohub | CZB Cliahub Consortium et al |
| EPI_ISL_6260085 | TPMG Regional Laboratory | California Department of Public Health | Emily Smith on behalf of CDPH-COVIDNet et al |
| EPI_ISL_5869846 | CDPH VBL | California Department of Public Health | Emily Smith on behalf of CDPH-COVIDNet et al |
| EPI_ISL_6074390 | Santa Clara County Public Health Laboratory | Santa Clara County Public Health Laboratory | Santa Clara County Public Health Department et al |
| EPI_ISL_3261727 | Scripps Medical Laboratory | Andersen lab at Scripps Research | SEARCH Alliance San Diego with Michael Quigley et al |
| EPI_ISL_1811771 | EXCITE Lab | Andersen lab at Scripps Research | Nicole L Washington et al |
| EPI_ISL_1477511 | CA DPH Viral and Rickettsial Disease Laboratory | Chan-Zuckerberg Biohub | CZB Cliahub Consortium et al |

| | | | |
|---|---|---|---|
| EPI_ISL_495134 | Innovative Genomics Institute, UC Berkeley | Innovative Genomics Institute, UC Berkeley | Stacia Wyman et al |
| EPI_ISL_5013577 | CDPH VBL | California Department of Public Health | Emily Smith on behalf of CDPH-COVIDNet et al |
| EPI_ISL_6260547 | TPMG Regional Laboratory | California Department of Public Health | Emily Smith on behalf of CDPH-COVIDNet et al |
| EPI_ISL_2839970 | Curative Labs | Curative Labs | Mikhail Hanewich-Hollatz et al |
| EPI_ISL_5696641 | CDPH VBL | California Department of Public Health | Emily Smith on behalf of CDPH-COVIDNet et al |
| EPI_ISL_4509768 | CDPH VBL | California Department of Public Health | Emily Smith on behalf of CDPH-COVIDNet et al |
| EPI_ISL_1824838 | Color Genomics | Chiu Laboratory, University of California, San Francisco | Charles Chiu et al |
| EPI_ISL_2890404 | CDPH VBL | California Department of Public Health | CDPH-COVIDNet et al |
| EPI_ISL_2658669 | Orange County Public Health Lab | Chan-Zuckerberg Biohub | CZB Cliahub Consortium et al |
| EPI_ISL_1026905 | Fulgent Genetics | Fulgent Genetics | Harry Gao et al |
| EPI_ISL_1235028 | Alameda County Public Health Lab | Chan-Zuckerberg Biohub | CZB Cliahub Consortium et al |
| EPI_ISL_4869614 | UC Davis Genome Center and Healthy Davis Together | UC Davis Genome Center | UC Davis et al |
| EPI_ISL_5263045 | CDPH VBL | California Department of Public Health | Emily Smith on behalf of CDPH-COVIDNet et al |
| EPI_ISL_1011352 | Fulgent Genetics | Fulgent Genetics | Harry Gao et al |
| EPI_ISL_3273409 | Curative Labs | Curative Labs | Mikhail Hanewich-Hollatz et al |
| EPI_ISL_5442633 | TPMG Regional Laboratory | California Department of Public Health | Emily Smith on behalf of CDPH-COVIDNet et al |
| EPI_ISL_6074728 | Kaiser Santa Clara | Santa Clara County Public Health Laboratory | Santa Clara County Public Health Department et al |
| EPI_ISL_1068800 | Fulgent Genetics | Fulgent Genetics | Harry Gao et al |
| EPI_ISL_1185334 | CA DPH Viral and Rickettsial Disease Laboratory | Chan-Zuckerberg Biohub | CZB Cliahub Consortium et al |
| EPI_ISL_604505 | Quest Diagnostics | Quest Diagnostics | Rosenthal et al |
| EPI_ISL_5442614 | TPMG Regional Laboratory | California Department of Public Health | Emily Smith on behalf of CDPH-COVIDNet et al |
| EPI_ISL_979130 | Santa Clara County Public Health Laboratory | Chan-Zuckerberg Biohub | CZB Cliahub Consortium et al |
| EPI_ISL_4492526 | CDPH VBL | California Department of Public Health | Emily Smith on behalf of CDPH-COVIDNet et al |
| EPI_ISL_2892058 | CDPH VBL | California Department of Public Health | CDPH-COVIDNet et al |
| EPI_ISL_5236930 | Sharp HealthCare Laboratory | Andersen lab at Scripps Research | SEARCH Alliance San Diego with Aaron Harding et al |
| EPI_ISL_5436964 | CDPH VBL | California Department of Public Health | Emily Smith on behalf of CDPH-COVIDNet et al |
| EPI_ISL_6947612 | CDPH VBL | California Department of Public Health | Emily Smith on behalf of CDPH-COVIDNet et al |
| EPI_ISL_1379977 | USC Clinical Lab | Los Angeles County Public Health Laboratories | P. Hemarajata et al. |
| EPI_ISL_2659207 | CA DPH Viral and Rickettsial Disease Laboratory | Chan-Zuckerberg Biohub | CZB Cliahub Consortium et al |

| | | | |
|---|---|---|---|
| EPI_ISL_3261656 | Scripps Medical Laboratory | Andersen lab at Scripps Research | SEARCH Alliance San Diego with Michael Quigley et al |
| EPI_ISL_5481899 | Pomona Valley Hospital Medical Center | Los Angeles County Public Health Laboratories | P. Hemarajata et al. |
| EPI_ISL_2923517 | CDPH VBL | California Department of Public Health | CDPH-COVIDNet et al |
| EPI_ISL_2103012 | Contra Costa County Public Health Lab | Chan-Zuckerberg Biohub | CZB Cliahub Consortium et al |
| EPI_ISL_1235533 | Santa Clara County Public Health Laboratory | Chan-Zuckerberg Biohub | CZB Cliahub Consortium et al |
| EPI_ISL_1811827 | EXCITE Lab | Andersen lab at Scripps Research | Nicole L Washington et al |
| EPI_ISL_5445858 | CDPH VBL | California Department of Public Health | Emily Smith on behalf of CDPH-COVIDNet et al |
| EPI_ISL_2659030 | Santa Clara Valley Medical Center | Santa Clara County Public Health Laboratory | Santa Clara County Public Health Department et al |
| EPI_ISL_5321535 | CDPH VBL | California Department of Public Health | Emily Smith on behalf of CDPH-COVIDNet et al |
| EPI_ISL_3723285 | TPMG Regional Laboratory | California Department of Public Health | Emily Smith on behalf of CDPH-COVIDNet et al |
| EPI_ISL_6947066 | TPMG Regional Laboratory | California Department of Public Health | Emily Smith on behalf of CDPH-COVIDNet et al |
| EPI_ISL_3543709 | Sharp HealthCare Laboratory | Andersen lab at Scripps Research | SEARCH Alliance San Diego with Aaron Harding et al |
| EPI_ISL_5802741 | Pomona Valley Hospital Medical Center | Los Angeles County Public Health Laboratories | P. Hemarajata et al. |
| EPI_ISL_980853 | Innovative Genomics Institute, UC Berkeley | Innovative Genomics Institute, UC Berkeley | Stacia Wyman et al |
| EPI_ISL_6123481 | CDPH VBL | California Department of Public Health | Emily Smith on behalf of CDPH-COVIDNet et al |
| EPI_ISL_3219422 | Santa Clara Valley Medical Center | Santa Clara County Public Health Laboratory | Santa Clara County Public Health Department et al |
| EPI_ISL_2882410 | CDPH VBL | California Department of Public Health | CDPH-COVIDNet et al |
| EPI_ISL_3756705 | PAMF Palo Alto | Santa Clara County Public Health Laboratory | Santa Clara County Public Health Department et al |
| EPI_ISL_5977999 | UC Davis Genome Center | UC Davis Genome Center | UC Davis et al |
| EPI_ISL_738932 | Alameda County Public Health Lab | Chan-Zuckerberg Biohub | CZB Cliahub Consortium et al |
| EPI_ISL_6947801 | CDPH VBL | California Department of Public Health | Emily Smith on behalf of CDPH-COVIDNet et al |
| EPI_ISL_583155 | Humboldt County Public Health Laboratory | Chan-Zuckerberg Biohub | CZB Cliahub Consortium et al |
| EPI_ISL_2921668 | CDPH VBL | California Department of Public Health | CDPH-COVIDNet et al |
| EPI_ISL_6260072 | TPMG Regional Laboratory | California Department of Public Health | Emily Smith on behalf of CDPH-COVIDNet et al |
| EPI_ISL_1813111 | EXCITE Lab | Andersen lab at Scripps Research | David Pride et al |
| EPI_ISL_677869 | Innovative Genomics Institute, UC Berkeley | Innovative Genomics Institute, UC Berkeley | Stacia Wyman et al |
| EPI_ISL_5629665 | Santa Clara Valley Medical Center | Santa Clara County Public Health Laboratory | Santa Clara County Public Health Department et al |

| | | | |
|---|---|---|---|
| EPI_ISL_3722891 | TPMG Regional Laboratory | California Department of Public Health | Emily Smith on behalf of CDPH-COVIDNet et al |
| EPI_ISL_1477741 | Alameda County Public Health Lab | Chan-Zuckerberg Biohub | CZB Cliahub Consortium et al |
| EPI_ISL_4491410 | CDPH VBL | California Department of Public Health | Emily Smith on behalf of CDPH-COVIDNet et al |
| EPI_ISL_6253126 | TPMG Regional Laboratory | California Department of Public Health | Emily Smith on behalf of CDPH-COVIDNet et al |
| EPI_ISL_1172078 | Fulgent Genetics | Fulgent Genetics | Harry Gao et al |
| EPI_ISL_6253178 | TPMG Regional Laboratory | California Department of Public Health | Emily Smith on behalf of CDPH-COVIDNet et al |
| EPI_ISL_1010670 | Fulgent Genetics | Fulgent Genetics | Harry Gao et al |
| EPI_ISL_3757020 | Sharp HealthCare Laboratory | Andersen lab at Scripps Research | SEARCH Alliance San Diego with Aaron Harding et al |
| EPI_ISL_4510092 | CDPH VBL | California Department of Public Health | Emily Smith on behalf of CDPH-COVIDNet et al |
| EPI_ISL_1235054 | Alameda County Public Health Lab | Chan-Zuckerberg Biohub | CZB Cliahub Consortium et al |
| EPI_ISL_5696389 | CDPH VBL | California Department of Public Health | Emily Smith on behalf of CDPH-COVIDNet et al |
| EPI_ISL_5198672 | CDPH VBL | California Department of Public Health | Emily Smith on behalf of CDPH-COVIDNet et al |
| EPI_ISL_7131926 | CDPH VBL | California Department of Public Health | Emily Smith on behalf of CDPH-COVIDNet et al |
| EPI_ISL_5606019 | Colorado Department of Public Health and Environment | Colorado Department of Public Health and Environment | Laura Bankers et al |
| EPI_ISL_5848458 | Colorado Department of Public Health and Environment | Colorado Department of Public Health and Environment | Laura Bankers et al |
| EPI_ISL_5931869 | Colorado Department of Public Health and Environment | Colorado Department of Public Health and Environment | Laura Bankers et al |
| EPI_ISL_5260006 | Colorado Department of Public Health and Environment | Colorado Department of Public Health and Environment | Laura Bankers et al |
| EPI_ISL_5930440 | Colorado Department of Public Health and Environment | Colorado Department of Public Health and Environment | Laura Bankers et al |
| EPI_ISL_7152727 | Mako Medical | Centers for Disease Control and Prevention Division of Viral Diseases, Pathogen Discovery | Dakota Howard et al |
| EPI_ISL_1540356 | Colorado Department of Public Health and Environment | Colorado Department of Public Health and Environment | Laura Bankers et al |
| EPI_ISL_4749964 | Colorado Department of Public Health and Environment | Colorado Department of Public Health and Environment | Laura Bankers et al |
| EPI_ISL_1839794 | Fulgent Genetics | Centers for Disease Control and Prevention Division of Viral Diseases, Pathogen Discovery | Dakota Howard et al |

| | | | |
|---|---|---|---|
| EPI_ISL_5931740 | Colorado Department of Public Health and Environment | Colorado Department of Public Health and Environment | Laura Bankers et al |
| EPI_ISL_4347987 | Mako Medical | Centers for Disease Control and Prevention Division of Viral Diseases, Pathogen Discovery | Dakota Howard et al |
| EPI_ISL_6652561 | Mako Medical | Centers for Disease Control and Prevention Division of Viral Diseases, Pathogen Discovery | Dakota Howard et al |
| EPI_ISL_2647074 | Mako Medical | Centers for Disease Control and Prevention Division of Viral Diseases, Pathogen Discovery | Dakota Howard et al |
| EPI_ISL_5259891 | Colorado Department of Public Health and Environment | Colorado Department of Public Health and Environment | Laura Bankers et al |
| EPI_ISL_2646727 | Mako Medical | Centers for Disease Control and Prevention Division of Viral Diseases, Pathogen Discovery | Dakota Howard et al |
| EPI_ISL_4301509 | Colorado Department of Public Health and Environment | Colorado Department of Public Health and Environment | Laura Bankers et al |
| EPI_ISL_5709714 | Colorado Department of Public Health and Environment | Colorado Department of Public Health and Environment | Laura Bankers et al |
| EPI_ISL_5989019 | Colorado Department of Public Health and Environment | Colorado Department of Public Health and Environment | Laura Bankers et al |
| EPI_ISL_5253673 | Colorado Department of Public Health and Environment | Colorado Department of Public Health and Environment | Laura Bankers et al |
| EPI_ISL_3404319 | Colorado Department of Public Health and Environment | Colorado Department of Public Health and Environment | Laura Bankers et al |
| EPI_ISL_6256324 | Mako Medical | Centers for Disease Control and Prevention Division of Viral Diseases, Pathogen Discovery | Dakota Howard et al |
| EPI_ISL_6255584 | Mako Medical | Centers for Disease Control and Prevention Division of Viral Diseases, Pathogen Discovery | Dakota Howard et al |
| EPI_ISL_2398012 | Laboratory Corporation of America | Centers for Disease Control and Prevention Division of Viral Diseases, Pathogen Discovery | Dakota Howard et al |
| EPI_ISL_6417772 | Fulgent Genetics | Centers for Disease Control and Prevention Division of Viral Diseases, Pathogen Discovery | Dakota Howard et al |
| EPI_ISL_2928556 | Fulgent Genetics | Centers for Disease Control and Prevention Division of Viral Diseases, Pathogen Discovery | Dakota Howard et al |
| EPI_ISL_6652752 | Mako Medical | Centers for Disease Control and Prevention Division of Viral Diseases, Pathogen Discovery | Dakota Howard et al |

| | | | |
|---|---|---|---|
| EPI_ISL_5604882 | Colorado Department of Public Health and Environment | Colorado Department of Public Health and Environment | Laura Bankers et al |
| EPI_ISL_6828331 | Colorado Mesa University | Infectious Disease Program, Broad Institute of Harvard and MIT | Marshall et al |
| EPI_ISL_1665856 | Colorado Department of Public Health and Environment | Colorado Department of Public Health and Environment | Laura Bankers et al |
| EPI_ISL_4890687 | Colorado Department of Public Health and Environment | Colorado Department of Public Health and Environment | Laura Bankers et al |
| EPI_ISL_5091289 | Fulgent Genetics | Centers for Disease Control and Prevention Division of Viral Diseases, Pathogen Discovery | Dakota Howard et al |
| EPI_ISL_3603734 | Mako Medical | Centers for Disease Control and Prevention Division of Viral Diseases, Pathogen Discovery | Dakota Howard et al |
| EPI_ISL_5347714 | Colorado Department of Public Health and Environment | Colorado Department of Public Health and Environment | Laura Bankers et al |
| EPI_ISL_6473893 | "CO Dept. of Public Health and Environment, Lab Services Division" | Centers for Disease Control and Prevention Division of Viral Diseases, Pathogen Discovery | Mili Sheth et al |
| EPI_ISL_6257135 | Mako Medical | Centers for Disease Control and Prevention Division of Viral Diseases, Pathogen Discovery | Dakota Howard et al |
| EPI_ISL_6677586 | Colorado Department of Public Health and Environment | Colorado Department of Public Health and Environment | Laura Bankers et al |
| EPI_ISL_4519024 | Sharp HealthCare Laboratory | Andersen lab at Scripps Research | SEARCH Alliance San Diego with Aaron Harding et al |
| EPI_ISL_6811258 | Mako Medical | Centers for Disease Control and Prevention Division of Viral Diseases, Pathogen Discovery | Dakota Howard et al |
| EPI_ISL_6676610 | Colorado Department of Public Health and Environment | Colorado Department of Public Health and Environment | Laura Bankers et al |
| EPI_ISL_6815022 | Colorado Department of Public Health and Environment | Colorado Department of Public Health and Environment | Laura Bankers et al |
| EPI_ISL_5932023 | Colorado Department of Public Health and Environment | Colorado Department of Public Health and Environment | Laura Bankers et al |
| EPI_ISL_5848869 | Colorado Department of Public Health and Environment | Colorado Department of Public Health and Environment | Laura Bankers et al |
| EPI_ISL_5989500 | Colorado Department of Public Health and Environment | Colorado Department of Public Health and Environment | Laura Bankers et al |
| EPI_ISL_3215744 | EXCITE Lab | Andersen lab at Scripps Research | David Pride et al |

| | | | |
|---|---|---|---|
| EPI_ISL_4731015 | Colorado Department of Public Health and Environment | Colorado Department of Public Health and Environment | Laura Bankers et al |
| EPI_ISL_1812987 | EXCITE Lab | Andersen lab at Scripps Research | David Pride et al |
| EPI_ISL_1813480 | EXCITE Lab | Andersen lab at Scripps Research | Nicole L Washington et al |
| EPI_ISL_6542132 | Mako Medical | Centers for Disease Control and Prevention Division of Viral Diseases, Pathogen Discovery | Dakota Howard et al |
| EPI_ISL_6638596 | Mako Medical | Centers for Disease Control and Prevention Division of Viral Diseases, Pathogen Discovery | Dakota Howard et al |
| EPI_ISL_2277415 | Colorado Department of Public Health and Environment | Colorado Department of Public Health and Environment | Laura Bankers et al |
| EPI_ISL_6052955 | Aegis Sciences Corporation | Centers for Disease Control and Prevention Division of Viral Diseases, Pathogen Discovery | Dakota Howard et al |
| EPI_ISL_5258728 | Colorado Department of Public Health and Environment | Colorado Department of Public Health and Environment | Laura Bankers et al |
| EPI_ISL_4745527 | Colorado Department of Public Health and Environment | Colorado Department of Public Health and Environment | Laura Bankers et al |
| EPI_ISL_5930490 | Colorado Department of Public Health and Environment | Colorado Department of Public Health and Environment | Laura Bankers et al |
| EPI_ISL_5987182 | Colorado Department of Public Health and Environment | Colorado Department of Public Health and Environment | Laura Bankers et al |
| EPI_ISL_4348105 | Mako Medical | Centers for Disease Control and Prevention Division of Viral Diseases, Pathogen Discovery | Dakota Howard et al |
| EPI_ISL_3427542 | Mako Medical | Centers for Disease Control and Prevention Division of Viral Diseases, Pathogen Discovery | Dakota Howard et al |
| EPI_ISL_6257201 | Mako Medical | Centers for Disease Control and Prevention Division of Viral Diseases, Pathogen Discovery | Dakota Howard et al |
| EPI_ISL_2031781 | UCSC Genomics Institute | UCSC Genomics Institute | Mark Akeson et al |
| EPI_ISL_5849181 | Colorado Department of Public Health and Environment | Colorado Department of Public Health and Environment | Laura Bankers et al |
| EPI_ISL_3301362 | Fulgent Genetics | Centers for Disease Control and Prevention Division of Viral Diseases, Pathogen Discovery | Dakota Howard et al |
| EPI_ISL_5979066 | Fulgent Genetics | Centers for Disease Control and Prevention Division of Viral Diseases, Pathogen Discovery | Dakota Howard et al |
| EPI_ISL_6381907 | Mako Medical | Centers for Disease Control and Prevention Division of Viral Diseases, Pathogen Discovery | Dakota Howard et al |

| | | | |
|---|---|---|---|
| EPI_ISL_7122129 | Mako Medical | Centers for Disease Control and Prevention Division of Viral Diseases, Pathogen Discovery | Dakota Howard et al |
| EPI_ISL_6585577 | Colorado Department of Public Health and Environment | Colorado Department of Public Health and Environment | Laura Bankers et al |
| EPI_ISL_1612491 | Laboratory Corporation of America | Centers for Disease Control and Prevention Division of Viral Diseases, Pathogen Discovery | Dakota Howard et al |
| EPI_ISL_3916252 | Stanford Health Care | Stanford University School of Medicine, Clinical Virology Laboratory | Malaya K. Sahoo et al |
| EPI_ISL_4744921 | Colorado Department of Public Health and Environment | Colorado Department of Public Health and Environment | Laura Bankers et al |
| EPI_ISL_6784723 | Aegis Sciences Corporation | Centers for Disease Control and Prevention Division of Viral Diseases, Pathogen Discovery | Dakota Howard et al |
| EPI_ISL_2309229 | Colorado Department of Public Health and Environment | Colorado Department of Public Health and Environment | Laura Bankers et al |
| EPI_ISL_6888446 | Mako Medical | Centers for Disease Control and Prevention Division of Viral Diseases, Pathogen Discovery | Dakota Howard et al |
| EPI_ISL_5987849 | Colorado Department of Public Health and Environment | Colorado Department of Public Health and Environment | Laura Bankers et al |
| EPI_ISL_4362230 | Aegis Sciences Corporation | Centers for Disease Control and Prevention Division of Viral Diseases, Pathogen Discovery | Dakota Howard et al |
| EPI_ISL_2276221 | Colorado Department of Public Health and Environment | Colorado Department of Public Health and Environment | Laura Bankers et al |
| EPI_ISL_5609533 | Colorado Department of Public Health and Environment | Colorado Department of Public Health and Environment | Laura Bankers et al |
| EPI_ISL_4195709 | Mako Medical | Centers for Disease Control and Prevention Division of Viral Diseases, Pathogen Discovery | Dakota Howard et al |
| EPI_ISL_5987916 | Colorado Department of Public Health and Environment | Colorado Department of Public Health and Environment | Laura Bankers et al |
| EPI_ISL_5056210 | Colorado Department of Public Health and Environment | Colorado Department of Public Health and Environment | Laura Bankers et al |
| EPI_ISL_5309400 | Mako Medical | Centers for Disease Control and Prevention Division of Viral Diseases, Pathogen Discovery | Dakota Howard et al |
| EPI_ISL_6951145 | Color Genomics | Chiu Laboratory, University of California, San Francisco | Charles Chiu et al |

| | | | |
|---|---|---|---|
| EPI_ISL_3583934 | Colorado Department of Public Health and Environment | Colorado Department of Public Health and Environment | Laura Bankers et al |
| EPI_ISL_5439212 | Colorado Department of Public Health and Environment | Colorado Department of Public Health and Environment | Laura Bankers et al |
| EPI_ISL_4927173 | Colorado Department of Public Health and Environment | Colorado Department of Public Health and Environment | Laura Bankers et al |
| EPI_ISL_5415660 | Colorado Department of Public Health and Environment | Colorado Department of Public Health and Environment | Laura Bankers et al |
| EPI_ISL_2646364 | Mako Medical | Centers for Disease Control and Prevention Division of Viral Diseases, Pathogen Discovery | Dakota Howard et al |
| EPI_ISL_3988480 | Colorado Department of Public Health and Environment | Colorado Department of Public Health and Environment | Laura Bankers et al |
| EPI_ISL_6913390 | Colorado Department of Public Health and Environment | Colorado Department of Public Health and Environment | Laura Bankers et al |
| EPI_ISL_3325510 | Laboratory Corporation of America | Centers for Disease Control and Prevention Division of Viral Diseases, Pathogen Discovery | Dakota Howard et al |
| EPI_ISL_1992456 | Fulgent Genetics | Centers for Disease Control and Prevention Division of Viral Diseases, Pathogen Discovery | Dakota Howard et al |
| EPI_ISL_1668006 | Aegis Sciences Corporation | Centers for Disease Control and Prevention Division of Viral Diseases, Pathogen Discovery | Dakota Howard et al |
| EPI_ISL_3112820 | Laboratory Corporation of America | Centers for Disease Control and Prevention Division of Viral Diseases, Pathogen Discovery | Dakota Howard et al |
| EPI_ISL_5190687 | Laboratory Corporation of America | Centers for Disease Control and Prevention Division of Viral Diseases, Pathogen Discovery | Dakota Howard et al |
| EPI_ISL_5178612 | Florida Bureau of Public Health Laboratories | Florida Bureau of Public Health Laboratories | Sarah Schmedes et al |
| EPI_ISL_1017508 | Murphy Medical Associates | Grubaugh Lab - Yale School of Public Health | Mary Petrone et al |
| EPI_ISL_5170454 | Laboratory Corporation of America | Centers for Disease Control and Prevention Division of Viral Diseases, Pathogen Discovery | Dakota Howard et al |
| EPI_ISL_3509892 | Laboratory Corporation of America | Centers for Disease Control and Prevention Division of Viral Diseases, Pathogen Discovery | Dakota Howard et al |
| EPI_ISL_1482634 | DC Public Health Lab/ Dept. of Forensic Sciences | DC Public Health Lab/ Dept. of Forensic Sciences | Janis Doss et al |
| EPI_ISL_3218560 | Aegis Sciences Corporation | Centers for Disease Control and Prevention Division of Viral Diseases, Pathogen Discovery | Dakota Howard et al |

| | | | |
|---|---|---|---|
| EPI_ISL_3328987 | Laboratory Corporation of America | Centers for Disease Control and Prevention Division of Viral Diseases, Pathogen Discovery | Dakota Howard et al |
| EPI_ISL_4963193 | Laboratory Corporation of America | Centers for Disease Control and Prevention Division of Viral Diseases, Pathogen Discovery | Dakota Howard et al |
| EPI_ISL_5236011 | Aegis Sciences Corporation | Centers for Disease Control and Prevention Division of Viral Diseases, Pathogen Discovery | Dakota Howard et al |
| EPI_ISL_1562927 | Aegis Sciences Corporation | Centers for Disease Control and Prevention Division of Viral Diseases, Pathogen Discovery | Dakota Howard et al |
| EPI_ISL_1162837 | Laboratory Corporation of America | Respiratory Viruses Branch, Division of Viral Diseases, Centers for Disease Control and Prevention | Peter W. Cook et al |
| EPI_ISL_1017565 | Murphy Medical Associates | Grubaugh Lab - Yale School of Public Health | Mary Petrone et al |
| EPI_ISL_7010485 | Colorado Department of Public Health and Environment | Colorado Department of Public Health and Environment | Laura Bankers et al |
| EPI_ISL_2442456 | Florida Bureau of Public Health Laboratories | Florida Bureau of Public Health Laboratories | Sarah Schmedes et al |
| EPI_ISL_1908953 | DPHL | Delaware Public Health Lab | Rebecca Savage et al |
| EPI_ISL_1513971 | Laboratory Corporation of America | Centers for Disease Control and Prevention Division of Viral Diseases, Pathogen Discovery | Dakota Howard et al |
| EPI_ISL_7140648 | Colorado Department of Public Health and Environment | Colorado Department of Public Health and Environment | Laura Bankers et al |
| EPI_ISL_3500575 | Yale Clinical Virology Lab | Grubaugh Lab - Yale School of Public Health | Joseph Fauver et al |
| EPI_ISL_2268476 | Quest Diagnostics Incorporated | Centers for Disease Control and Prevention Division of Viral Diseases, Pathogen Discovery | Dakota Howard et al |
| EPI_ISL_2306930 | Laboratory Corporation of America | Centers for Disease Control and Prevention Division of Viral Diseases, Pathogen Discovery | Dakota Howard et al |
| EPI_ISL_3100717 | Florida Bureau of Public Health Laboratories | Florida Bureau of Public Health Laboratories | Sarah Schmedes et al |
| EPI_ISL_1531276 | Florida Bureau of Public Health Laboratories | Florida Bureau of Public Health Laboratories | Sarah Schmedes et al |
| EPI_ISL_3692668 | DPHL | Delaware Public Health Lab | Rebecca Savage et al |
| EPI_ISL_6646632 | Aegis Sciences Corporation | Centers for Disease Control and Prevention Division of Viral Diseases, Pathogen Discovery | Dakota Howard et al |
| EPI_ISL_4157451 | Laboratory Corporation of America | Centers for Disease Control and Prevention Division of Viral Diseases, Pathogen Discovery | Dakota Howard et al |
| EPI_ISL_4371421 | Quest Diagnostics Incorporated | Centers for Disease Control and Prevention Division of Viral Diseases, Pathogen Discovery | Dakota Howard et al |

| | | | |
|---|---|---|---|
| EPI_ISL_7128428 | Helix | Centers for Disease Control and Prevention Division of Viral Diseases, Pathogen Discovery | Dakota Howard et al |
| EPI_ISL_3861708 | Aegis Sciences Corporation | Centers for Disease Control and Prevention Division of Viral Diseases, Pathogen Discovery | Dakota Howard et al |
| EPI_ISL_613728 | Florida Bureau of Public Health Laboratories | Florida Bureau of Public Health Laboratories | Sarah Schmedes et al |
| EPI_ISL_4553985 | Aegis Sciences Corporation | Centers for Disease Control and Prevention Division of Viral Diseases, Pathogen Discovery | Dakota Howard et al |
| EPI_ISL_729741 | Connecticut Department of Health | Grubaugh Lab - Yale School of Public Health | Joseph Fauver et al |
| EPI_ISL_5548922 | Aegis Sciences Corporation | Centers for Disease Control and Prevention Division of Viral Diseases, Pathogen Discovery | Dakota Howard et al |
| EPI_ISL_7045293 | Colorado Department of Public Health and Environment | Colorado Department of Public Health and Environment | Laura Bankers et al |
| EPI_ISL_2383702 | DPHL | Delaware Public Health Lab | Rebecca Savage et al |
| EPI_ISL_5322613 | Florida Bureau of Public Health Laboratories | Florida Bureau of Public Health Laboratories | Sarah Schmedes et al |
| EPI_ISL_5458402 | The Jackson Laboratory | The Jackson Laboratory | Long E et al |
| EPI_ISL_3853604 | Aegis Sciences Corporation | Centers for Disease Control and Prevention Division of Viral Diseases, Pathogen Discovery | Dakota Howard et al |
| EPI_ISL_6378412 | Aegis Sciences Corporation | Centers for Disease Control and Prevention Division of Viral Diseases, Pathogen Discovery | Dakota Howard et al |
| EPI_ISL_1339945 | Quest Diagnostics Incorporated | Centers for Disease Control and Prevention Division of Viral Diseases, Pathogen Discovery | Peter W. Cook et al |
| EPI_ISL_4911042 | Aegis Sciences Corporation | Centers for Disease Control and Prevention Division of Viral Diseases, Pathogen Discovery | Dakota Howard et al |
| EPI_ISL_2149498 | Aegis Sciences Corporation | Centers for Disease Control and Prevention Division of Viral Diseases, Pathogen Discovery | Dakota Howard et al |
| EPI_ISL_1462640 | Laboratory Corporation of America | Centers for Disease Control and Prevention Division of Viral Diseases, Pathogen Discovery | Dakota Howard et al |
| EPI_ISL_2399069 | Laboratory Corporation of America | Centers for Disease Control and Prevention Division of Viral Diseases, Pathogen Discovery | Dakota Howard et al |
| EPI_ISL_2383772 | DPHL | Delaware Public Health Lab | Rebecca Savage et al |
| EPI_ISL_2089316 | Infinity Biologix | Centers for Disease Control and Prevention Division of Viral Diseases, Pathogen Discovery | Dakota Howard et al |
| EPI_ISL_3111093 | Infinity Biologix | Centers for Disease Control and Prevention Division of Viral Diseases, Pathogen Discovery | Dakota Howard et al |

| | | | |
|---|---|---|---|
| EPI_ISL_2442449 | Florida Bureau of Public Health Laboratories | Florida Bureau of Public Health Laboratories | Sarah Schmedes et al |
| EPI_ISL_3355361 | FL Bureau of Public Health Laboratories-Tampa | Centers for Disease Control and Prevention Division of Viral Diseases, Pathogen Discovery | Mili Sheth et al |
| EPI_ISL_2662757 | DPHL | Delaware Public Health Lab | Rebecca Savage et al |
| EPI_ISL_5002343 | Aegis Sciences Corporation | Centers for Disease Control and Prevention Division of Viral Diseases, Pathogen Discovery | Dakota Howard et al |
| EPI_ISL_913745 | DC Public Health Lab/ Dept. of Forensic Sciences | DC Public Health Lab/ Dept. of Forensic Sciences | Scott Nguyen et al |
| EPI_ISL_6163547 | Aegis Sciences Corporation | Centers for Disease Control and Prevention Division of Viral Diseases, Pathogen Discovery | Dakota Howard et al |
| EPI_ISL_3511261 | Laboratory Corporation of America | Centers for Disease Control and Prevention Division of Viral Diseases, Pathogen Discovery | Dakota Howard et al |
| EPI_ISL_3862754 | Aegis Sciences Corporation | Centers for Disease Control and Prevention Division of Viral Diseases, Pathogen Discovery | Dakota Howard et al |
| EPI_ISL_4373358 | Quest Diagnostics Incorporated | Centers for Disease Control and Prevention Division of Viral Diseases, Pathogen Discovery | Dakota Howard et al |
| EPI_ISL_1555434 | Fulgent Genetics | Centers for Disease Control and Prevention Division of Viral Diseases, Pathogen Discovery | Dakota Howard et al |
| EPI_ISL_2452008 | DE Public Health Laboratory | Centers for Disease Control and Prevention Division of Viral Diseases, Pathogen Discovery | Mili Sheth et al |
| EPI_ISL_1163484 | Laboratory Corporation of America | Respiratory Viruses Branch, Division of Viral Diseases, Centers for Disease Control and Prevention | Peter W. Cook et al |
| EPI_ISL_1462493 | Laboratory Corporation of America | Centers for Disease Control and Prevention Division of Viral Diseases, Pathogen Discovery | Dakota Howard et al |
| EPI_ISL_666866 | Florida Bureau of Public Health Laboratories | Florida Bureau of Public Health Laboratories | Sarah Schmedes et al |
| EPI_ISL_2228970 | DE Public Health Laboratory | Centers for Disease Control and Prevention Division of Viral Diseases, Pathogen Discovery | Mili Sheth et al |
| EPI_ISL_4157509 | Laboratory Corporation of America | Centers for Disease Control and Prevention Division of Viral Diseases, Pathogen Discovery | Dakota Howard et al |
| EPI_ISL_1447630 | Yale Clinical Virology Lab | Grubaugh Lab - Yale School of Public Health | Joseph Fauver et al |
| EPI_ISL_4297142 | Florida Bureau of Public Health Laboratories | Florida Bureau of Public Health Laboratories | Sarah Schmedes et al |
| EPI_ISL_2784931 | Aegis Sciences Corporation | Centers for Disease Control and Prevention Division of Viral Diseases, Pathogen Discovery | Dakota Howard et al |

| | | | |
|---|---|---|---|
| EPI_ISL_2242053 | Aegis Sciences Corporation | Centers for Disease Control and Prevention Division of Viral Diseases, Pathogen Discovery | Dakota Howard et al |
| EPI_ISL_5743885 | Broad Institute Clinical Research Sequencing Platform | Infectious Disease Program, Broad Institute of Harvard and MIT | Siddle et al |
| EPI_ISL_3752293 | Aegis Sciences Corporation | Centers for Disease Control and Prevention Division of Viral Diseases, Pathogen Discovery | Dakota Howard et al |
| EPI_ISL_708414 | Delaware Public Health Lab | Delaware Public Health Lab | Gregory Hovan et al |
| EPI_ISL_1447519 | Yale Clinical Virology Lab | Grubaugh Lab - Yale School of Public Health | Joseph Fauver et al |
| EPI_ISL_2153542 | Florida Bureau of Public Health Laboratories | Florida Bureau of Public Health Laboratories | Sarah Schmedes et al |
| EPI_ISL_2926230 | Laboratory Corporation of America | Centers for Disease Control and Prevention Division of Viral Diseases, Pathogen Discovery | Dakota Howard et al |
| EPI_ISL_6481011 | Helix | Centers for Disease Control and Prevention Division of Viral Diseases, Pathogen Discovery | Dakota Howard et al |
| EPI_ISL_3017161 | Laboratory Corporation of America | Centers for Disease Control and Prevention Division of Viral Diseases, Pathogen Discovery | Dakota Howard et al |
| EPI_ISL_1674921 | Florida Bureau of Public Health Laboratories | Florida Bureau of Public Health Laboratories | Sarah Schmedes et al |
| EPI_ISL_2926262 | Laboratory Corporation of America | Centers for Disease Control and Prevention Division of Viral Diseases, Pathogen Discovery | Dakota Howard et al |
| EPI_ISL_1273342 | The Jackson Laboratory | The Jackson Laboratory | Bergeron D et al |
| EPI_ISL_6398441 | Laboratory Corporation of America | Centers for Disease Control and Prevention Division of Viral Diseases, Pathogen Discovery | Dakota Howard et al |
| EPI_ISL_6858729 | Florida Bureau of Public Health Laboratories | Florida Bureau of Public Health Laboratories | Sarah Schmedes et al |
| EPI_ISL_4921150 | Aegis Sciences Corporation | Centers for Disease Control and Prevention Division of Viral Diseases, Pathogen Discovery | Dakota Howard et al |
| EPI_ISL_6935688 | Colorado Department of Public Health and Environment | Colorado Department of Public Health and Environment | Laura Bankers et al |
| EPI_ISL_2244092 | Aegis Sciences Corporation | Centers for Disease Control and Prevention Division of Viral Diseases, Pathogen Discovery | Dakota Howard et al |
| EPI_ISL_5925377 | Florida Bureau of Public Health Laboratories | Florida Bureau of Public Health Laboratories | Sarah Schmedes et al |
| EPI_ISL_6049972 | Florida Bureau of Public Health Laboratories | Florida Bureau of Public Health Laboratories | Sarah Schmedes et al |
| EPI_ISL_7145109 | DC Public Health Lab/ Dept. of Forensic Sciences | DC Public Health Lab/ Dept. of Forensic Sciences | Janis Doss et al |
| EPI_ISL_653321 | Florida Bureau of Public Health Laboratories | Florida Bureau of Public Health Laboratories | Sarah Schmedes et al |

| | | | |
|---|---|---|---|
| EPI_ISL_1560004 | Aegis Sciences Corporation | Centers for Disease Control and Prevention Division of Viral Diseases, Pathogen Discovery | Dakota Howard et al |
| EPI_ISL_5568906 | DC Public Health Lab/ Dept. of Forensic Sciences | DC Public Health Lab/ Dept. of Forensic Sciences | Janis Doss et al |
| EPI_ISL_6414787 | Quest Diagnostics Incorporated | Centers for Disease Control and Prevention Division of Viral Diseases, Pathogen Discovery | Dakota Howard et al |
| EPI_ISL_4158487 | Laboratory Corporation of America | Centers for Disease Control and Prevention Division of Viral Diseases, Pathogen Discovery | Dakota Howard et al |
| EPI_ISL_3752426 | Aegis Sciences Corporation | Centers for Disease Control and Prevention Division of Viral Diseases, Pathogen Discovery | Dakota Howard et al |
| EPI_ISL_1674809 | Florida Bureau of Public Health Laboratories | Florida Bureau of Public Health Laboratories | Sarah Schmedes et al |
| EPI_ISL_3333072 | State Hygienic Laboratory at the University of Iowa | State Hygienic Laboratory at the University of Iowa | Valerie Reeb et al |
| EPI_ISL_6337132 | Helix | Centers for Disease Control and Prevention Division of Viral Diseases, Pathogen Discovery | Dakota Howard et al |
| EPI_ISL_4368287 | Quest Diagnostics Incorporated | Centers for Disease Control and Prevention Division of Viral Diseases, Pathogen Discovery | Dakota Howard et al |
| EPI_ISL_4914706 | Aegis Sciences Corporation | Centers for Disease Control and Prevention Division of Viral Diseases, Pathogen Discovery | Dakota Howard et al |
| EPI_ISL_2160174 | Helix/Illumina | Centers for Disease Control and Prevention Division of Viral Diseases, Pathogen Discovery | Dakota Howard et al |
| EPI_ISL_2724550 | State Laboratories Division, Hawaii State Department of Health | State Laboratories Division, Hawaii State Department of Health | Pamela O'Brien et al |
| EPI_ISL_6484529 | Helix | Centers for Disease Control and Prevention Division of Viral Diseases, Pathogen Discovery | Dakota Howard et al |
| EPI_ISL_1804536 | Helix/Illumina | Centers for Disease Control and Prevention Division of Viral Diseases, Pathogen Discovery | Dakota Howard et al |
| EPI_ISL_4465221 | Quest Diagnostics Incorporated | Centers for Disease Control and Prevention Division of Viral Diseases, Pathogen Discovery | Dakota Howard et al |
| EPI_ISL_2724690 | State Laboratories Division, Hawaii State Department of Health | State Laboratories Division, Hawaii State Department of Health | Pamela O'Brien et al |
| EPI_ISL_5114783 | State Hygienic Laboratory at the University of Iowa | State Hygienic Laboratory at the University of Iowa | Valerie Reeb et al |
| EPI_ISL_2783888 | Curative Labs | Curative Labs | Mikhail Hanewich-Hollatz et al |
| EPI_ISL_4925552 | Mako Medical | Centers for Disease Control and Prevention Division of Viral Diseases, Pathogen Discovery | Dakota Howard et al |
| EPI_ISL_2784106 | Curative Labs | Curative Labs | Mikhail Hanewich-Hollatz et al |

| | | | |
|---|---|---|---|
| EPI_ISL_4815190 | Laboratory Corporation of America | Centers for Disease Control and Prevention Division of Viral Diseases, Pathogen Discovery | Dakota Howard et al |
| EPI_ISL_3840692 | GA Department of Public Health | GA Department of Public Health | Stacy Reeves et al |
| EPI_ISL_4479223 | Laboratory Corporation of America | Centers for Disease Control and Prevention Division of Viral Diseases, Pathogen Discovery | Dakota Howard et al |
| EPI_ISL_4332813 | Laboratory Corporation of America | Centers for Disease Control and Prevention Division of Viral Diseases, Pathogen Discovery | Dakota Howard et al |
| EPI_ISL_4173906 | Laboratory Corporation of America | Centers for Disease Control and Prevention Division of Viral Diseases, Pathogen Discovery | Dakota Howard et al |
| EPI_ISL_1553448 | Helix/Illumina | Centers for Disease Control and Prevention Division of Viral Diseases, Pathogen Discovery | Dakota Howard et al |
| EPI_ISL_3743665 | Laboratory Corporation of America | Centers for Disease Control and Prevention Division of Viral Diseases, Pathogen Discovery | Dakota Howard et al |
| EPI_ISL_2876677 | Quest Diagnostics Incorporated | Centers for Disease Control and Prevention Division of Viral Diseases, Pathogen Discovery | Dakota Howard et al |
| EPI_ISL_7013302 | State Laboratories Division, Hawaii State Department of Health | State Laboratories Division, Hawaii State Department of Health | Pamela O'Brien et al |
| EPI_ISL_4141756 | Aegis Sciences Corporation | Centers for Disease Control and Prevention Division of Viral Diseases, Pathogen Discovery | Dakota Howard et al |
| EPI_ISL_4481156 | Laboratory Corporation of America | Centers for Disease Control and Prevention Division of Viral Diseases, Pathogen Discovery | Dakota Howard et al |
| EPI_ISL_3355309 | GA Department of Public Health Laboratory | Centers for Disease Control and Prevention Division of Viral Diseases, Pathogen Discovery | Mili Sheth et al |
| EPI_ISL_4825329 | Mako Medical | Centers for Disease Control and Prevention Division of Viral Diseases, Pathogen Discovery | Dakota Howard et al |
| EPI_ISL_1551684 | Quest Diagnostics Incorporated | Centers for Disease Control and Prevention Division of Viral Diseases, Pathogen Discovery | Peter W. Cook et al |
| EPI_ISL_1292436 | State Laboratories Division, Hawaii State Department of Health | State Laboratories Division, Hawaii State Department of Health | Pamela O'Brien et al |
| EPI_ISL_1691518 | Quest Diagnostics Incorporated | Centers for Disease Control and Prevention Division of Viral Diseases, Pathogen Discovery | Dakota Howard et al |
| EPI_ISL_5347724 | State Hygienic Laboratory at the University of Iowa | State Hygienic Laboratory at the University of Iowa | Valerie Reeb et al |
| EPI_ISL_7013425 | State Laboratories Division, Hawaii State Department of Health | State Laboratories Division, Hawaii State Department of Health | Pamela O'Brien et al |

| | | | |
|---|---|---|---|
| EPI_ISL_6481158 | Helix | Centers for Disease Control and Prevention Division of Viral Diseases, Pathogen Discovery | Dakota Howard et al |
| EPI_ISL_6310175 | Mako Medical | Centers for Disease Control and Prevention Division of Viral Diseases, Pathogen Discovery | Dakota Howard et al |
| EPI_ISL_6336259 | Helix | Centers for Disease Control and Prevention Division of Viral Diseases, Pathogen Discovery | Dakota Howard et al |
| EPI_ISL_5083701 | Mako Medical | Centers for Disease Control and Prevention Division of Viral Diseases, Pathogen Discovery | Dakota Howard et al |
| EPI_ISL_6488553 | Helix | Centers for Disease Control and Prevention Division of Viral Diseases, Pathogen Discovery | Dakota Howard et al |
| EPI_ISL_2801465 | IVY3 Central Lab, Vanderbilt University Medical Center | Lauring Lab, University of Michigan, Department of Microbiology and Immunology | Valesano et al |
| EPI_ISL_3827527 | State Hygienic Laboratory at the University of Iowa | State Hygienic Laboratory at the University of Iowa | Valerie Reeb et al |
| EPI_ISL_5089997 | Laboratory Corporation of America | Centers for Disease Control and Prevention Division of Viral Diseases, Pathogen Discovery | Dakota Howard et al |
| EPI_ISL_1088966 | Helix / Illumina | Respiratory Viruses Branch, Division of Viral Diseases, Centers for Disease Control and Prevention | Peter W. Cook et al |
| EPI_ISL_1167957 | GA Department of Public Health Laboratory | Genomics and Discovery, Respiratory Viruses Branch, Division of Viral Diseases, Centers for Disease Control and Prevention | Brian Lynch et al |
| EPI_ISL_4728477 | State Laboratories Division, Hawaii State Department of Health | State Laboratories Division, Hawaii State Department of Health | Pamela O'Brien et al |
| EPI_ISL_1318498 | "HI Dept. of Health, State Laboratories Division" | Centers for Disease Control and Prevention Division of Viral Diseases, Pathogen Discovery | Krista Queen et al |
| EPI_ISL_1515934 | State Laboratories Division, Hawaii State Department of Health | State Laboratories Division, Hawaii State Department of Health | Pamela O'Brien et al |
| EPI_ISL_1016840 | Helix/Illumina | Respiratory Viruses Branch, Division of Viral Diseases, Centers for Disease Control and Prevention | Peter W. Cook et al |
| EPI_ISL_6628992 | Aegis Sciences Corporation | Centers for Disease Control and Prevention Division of Viral Diseases, Pathogen Discovery | Dakota Howard et al |
| EPI_ISL_1443851 | Helix/Illumina | Centers for Disease Control and Prevention Division of Viral Diseases, Pathogen Discovery | Peter W. Cook et al |
| EPI_ISL_1458712 | Fulgent Genetics | Centers for Disease Control and Prevention Division of Viral Diseases, Pathogen Discovery | Dakota Howard et al |

| | | | |
|---|---|---|---|
| EPI_ISL_1340312 | Helix/Illumina | Centers for Disease Control and Prevention Division of Viral Diseases, Pathogen Discovery | Peter W. Cook et al |
| EPI_ISL_5578539 | Mako Medical | Centers for Disease Control and Prevention Division of Viral Diseases, Pathogen Discovery | Dakota Howard et al |
| EPI_ISL_1515918 | State Laboratories Division, Hawaii State Department of Health | State Laboratories Division, Hawaii State Department of Health | Pamela O'Brien et al |
| EPI_ISL_4020973 | Quest Diagnostics Incorporated | Centers for Disease Control and Prevention Division of Viral Diseases, Pathogen Discovery | Dakota Howard et al |
| EPI_ISL_3603277 | Mako Medical | Centers for Disease Control and Prevention Division of Viral Diseases, Pathogen Discovery | Dakota Howard et al |
| EPI_ISL_3672738 | State Hygienic Laboratory at the University of Iowa | State Hygienic Laboratory at the University of Iowa | Valerie Reeb et al |
| EPI_ISL_2107350 | Emory Molecular Diagnostics Laboratory, Emory Healthcare | Piantadosi Lab, Emory Department of Pathology | Ahmed Babiker et al |
| EPI_ISL_1754177 | Tampa General Hospital Esoteric Lab | Tampa General Hospital Esoteric Research & Development Lab | Grant Vestal et al |
| EPI_ISL_4333436 | Laboratory Corporation of America | Centers for Disease Control and Prevention Division of Viral Diseases, Pathogen Discovery | Dakota Howard et al |
| EPI_ISL_1095119 | GA Department of Public Health Laboratory | Respiratory Viruses Branch, Division of Viral Diseases, Centers for Disease Control and Prevention | Krista Queen et al |
| EPI_ISL_3865144 | Mako Medical | Centers for Disease Control and Prevention Division of Viral Diseases, Pathogen Discovery | Dakota Howard et al |
| EPI_ISL_5658459 | Helix | Centers for Disease Control and Prevention Division of Viral Diseases, Pathogen Discovery | Dakota Howard et al |
| EPI_ISL_5516472 | State Laboratories Division, Hawaii State Department of Health | State Laboratories Division, Hawaii State Department of Health | Pamela O'Brien et al |
| EPI_ISL_3801945 | Gravity Diagnostics, LLC | Gravity Diagnostics, LLC | Gravity Diagnostics et al |
| EPI_ISL_2966842 | Tampa General Hospital Esoteric Lab | Tampa General Hospital Esoteric Research & Development Lab | Jon Faughn et al |
| EPI_ISL_5629015 | State Hygienic Laboratory at the University of Iowa | State Hygienic Laboratory at the University of Iowa | Valerie Reeb et al |
| EPI_ISL_1269269 | Helix/Illumina | Centers for Disease Control and Prevention Division of Viral Diseases, Pathogen Discovery | Peter W. Cook et al |
| EPI_ISL_5154491 | Mako Medical | Centers for Disease Control and Prevention Division of Viral Diseases, Pathogen Discovery | Dakota Howard et al |
| EPI_ISL_2244055 | Aegis Sciences Corporation | Centers for Disease Control and Prevention Division of Viral Diseases, Pathogen Discovery | Dakota Howard et al |

| | | | |
|---|---|---|---|
| EPI_ISL_1336030 | Quest Diagnostics Incorporated | Centers for Disease Control and Prevention Division of Viral Diseases, Pathogen Discovery | Peter W. Cook et al |
| EPI_ISL_1807216 | GA Department of Public Health | GA Department of Public Health | Stacy Reeves et al |
| EPI_ISL_1680273 | Helix/Illumina | Centers for Disease Control and Prevention Division of Viral Diseases, Pathogen Discovery | Dakota Howard et al |
| EPI_ISL_4347427 | Helix/Illumina | Centers for Disease Control and Prevention Division of Viral Diseases, Pathogen Discovery | Dakota Howard et al |
| EPI_ISL_6783238 | Quest Diagnostics Incorporated | Centers for Disease Control and Prevention Division of Viral Diseases, Pathogen Discovery | Dakota Howard et al |
| EPI_ISL_4812362 | Laboratory Corporation of America | Centers for Disease Control and Prevention Division of Viral Diseases, Pathogen Discovery | Dakota Howard et al |
| EPI_ISL_3355056 | GA Department of Public Health Laboratory | Centers for Disease Control and Prevention Division of Viral Diseases, Pathogen Discovery | Mili Sheth et al |
| EPI_ISL_4464288 | Quest Diagnostics Incorporated | Centers for Disease Control and Prevention Division of Viral Diseases, Pathogen Discovery | Dakota Howard et al |
| EPI_ISL_4813259 | Laboratory Corporation of America | Centers for Disease Control and Prevention Division of Viral Diseases, Pathogen Discovery | Dakota Howard et al |
| EPI_ISL_1089388 | Helix / Illumina | Respiratory Viruses Branch, Division of Viral Diseases, Centers for Disease Control and Prevention | Peter W. Cook et al |
| EPI_ISL_5231730 | Aegis Sciences Corporation | Centers for Disease Control and Prevention Division of Viral Diseases, Pathogen Discovery | Dakota Howard et al |
| EPI_ISL_6488826 | Helix | Centers for Disease Control and Prevention Division of Viral Diseases, Pathogen Discovery | Dakota Howard et al |
| EPI_ISL_4237632 | Mako Medical | Centers for Disease Control and Prevention Division of Viral Diseases, Pathogen Discovery | Dakota Howard et al |
| EPI_ISL_6310123 | Mako Medical | Centers for Disease Control and Prevention Division of Viral Diseases, Pathogen Discovery | Dakota Howard et al |
| EPI_ISL_5522098 | Helix | Centers for Disease Control and Prevention Division of Viral Diseases, Pathogen Discovery | Dakota Howard et al |
| EPI_ISL_2244358 | Aegis Sciences Corporation | Centers for Disease Control and Prevention Division of Viral Diseases, Pathogen Discovery | Dakota Howard et al |
| EPI_ISL_5518914 | Helix | Centers for Disease Control and Prevention Division of Viral Diseases, Pathogen Discovery | Dakota Howard et al |

| | | | |
|---|---|---|---|
| EPI_ISL_5861078 | Aegis Sciences Corporation | Centers for Disease Control and Prevention Division of Viral Diseases, Pathogen Discovery | Dakota Howard et al |
| EPI_ISL_1089376 | Helix / Illumina | Respiratory Viruses Branch, Division of Viral Diseases, Centers for Disease Control and Prevention | Peter W. Cook et al |
| EPI_ISL_6484508 | Helix | Centers for Disease Control and Prevention Division of Viral Diseases, Pathogen Discovery | Dakota Howard et al |
| EPI_ISL_3861366 | Aegis Sciences Corporation | Centers for Disease Control and Prevention Division of Viral Diseases, Pathogen Discovery | Dakota Howard et al |
| EPI_ISL_2790515 | Tampa General Hospital Esoteric Lab | Tampa General Hospital Esoteric Research & Development Lab | Jon Faughn et al |
| EPI_ISL_2784029 | Curative Labs | Curative Labs | Mikhail Hanewich-Hollatz et al |
| EPI_ISL_6398116 | Laboratory Corporation of America | Centers for Disease Control and Prevention Division of Viral Diseases, Pathogen Discovery | Dakota Howard et al |
| EPI_ISL_3748838 | Laboratory Corporation of America | Centers for Disease Control and Prevention Division of Viral Diseases, Pathogen Discovery | Dakota Howard et al |
| EPI_ISL_1549101 | Laboratory Corporation of America | Centers for Disease Control and Prevention Division of Viral Diseases, Pathogen Discovery | Dakota Howard et al |
| EPI_ISL_5849764 | State Laboratories Division, Hawaii State Department of Health | State Laboratories Division, Hawaii State Department of Health | Pamela O'Brien et al |
| EPI_ISL_5154461 | Mako Medical | Centers for Disease Control and Prevention Division of Viral Diseases, Pathogen Discovery | Dakota Howard et al |
| EPI_ISL_1480500 | Helix/Illumina | Centers for Disease Control and Prevention Division of Viral Diseases, Pathogen Discovery | Dakota Howard et al |
| EPI_ISL_1297961 | Helix/Illumina | Centers for Disease Control and Prevention Division of Viral Diseases, Pathogen Discovery | Peter W. Cook et al |
| EPI_ISL_6545853 | Aegis Sciences Corporation | Centers for Disease Control and Prevention Division of Viral Diseases, Pathogen Discovery | Dakota Howard et al |
| EPI_ISL_2444234 | Indiana State Department of Health | Quantigen Biosciences | Paul J. Chidress et al |
| EPI_ISL_6487391 | Helix | Centers for Disease Control and Prevention Division of Viral Diseases, Pathogen Discovery | Dakota Howard et al |
| EPI_ISL_1926672 | Laboratory Corporation of America | Centers for Disease Control and Prevention Division of Viral Diseases, Pathogen Discovery | Dakota Howard et al |
| EPI_ISL_2894532 | Kansas Health and Environmental Lab | Kansas Health and Environmental Lab | Katherine Wiggins et al |
| EPI_ISL_2716288 | Kansas Health and Environmental Lab | Kansas Health and Environmental Lab | Katherine Wiggins et al |

| | | | |
|---|---|---|---|
| EPI_ISL_2440584 | Quest Diagnostics Incorporated | Centers for Disease Control and Prevention Division of Viral Diseases, Pathogen Discovery | Dakota Howard et al |
| EPI_ISL_1925376 | Aegis Sciences Corporation | Centers for Disease Control and Prevention Division of Viral Diseases, Pathogen Discovery | Dakota Howard et al |
| EPI_ISL_2157611 | Reditus Laboratories | Reditus Laboratories | Joshua J. Geltz et al |
| EPI_ISL_2755706 | Curative Labs | Curative Labs | Mikhail Hanewich-Hollatz et al |
| EPI_ISL_2045279 | Laboratory Corporation of America | Centers for Disease Control and Prevention Division of Viral Diseases, Pathogen Discovery | Dakota Howard et al |
| EPI_ISL_4140444 | Aegis Sciences Corporation | Centers for Disease Control and Prevention Division of Viral Diseases, Pathogen Discovery | Dakota Howard et al |
| EPI_ISL_626424 | Northwestern Memorial Hospital | Ozer Lab | Ramon Lorenzo-Redondo et al |
| EPI_ISL_6941429 | IN State Department of Health Laboratory Services | IN State Department of Health Laboratory Services | Cassandra Campion et al |
| EPI_ISL_1686252 | Aegis Sciences Corporation | Centers for Disease Control and Prevention Division of Viral Diseases, Pathogen Discovery | Dakota Howard et al |
| EPI_ISL_5079884 | Laboratory Corporation of America | Centers for Disease Control and Prevention Division of Viral Diseases, Pathogen Discovery | Dakota Howard et al |
| EPI_ISL_1609000 | Laboratory Corporation of America | Centers for Disease Control and Prevention Division of Viral Diseases, Pathogen Discovery | Dakota Howard et al |
| EPI_ISL_5553697 | Aegis Sciences Corporation | Centers for Disease Control and Prevention Division of Viral Diseases, Pathogen Discovery | Dakota Howard et al |
| EPI_ISL_1490753 | IN State Department of Health Laboratory Services | IN State Department of Health Laboratory Services | Cassandra Campion et al |
| EPI_ISL_2159619 | Helix/Illumina | Centers for Disease Control and Prevention Division of Viral Diseases, Pathogen Discovery | Dakota Howard et al |
| EPI_ISL_1648987 | Aegis Sciences Corporation | Centers for Disease Control and Prevention Division of Viral Diseases, Pathogen Discovery | Dakota Howard et al |
| EPI_ISL_4356093 | IN State Department of Health Laboratory Services | IN State Department of Health Laboratory Services | Cassandra Campion et al |
| EPI_ISL_2300513 | Illinois Department of Public Health | Illinois Department of Public Health - Chicago Lab | Vineet K. Dhiman et al |
| EPI_ISL_4914834 | Aegis Sciences Corporation | Centers for Disease Control and Prevention Division of Viral Diseases, Pathogen Discovery | Dakota Howard et al |
| EPI_ISL_2346189 | Notre Dame COVID Response and Testing Laboratory | Notre Dame Genomics & Bioinformatics Core Facility | Michael Pfrender et al |
| EPI_ISL_2011162 | Quest Diagnostics Incorporated | Centers for Disease Control and Prevention Division of Viral Diseases, Pathogen Discovery | Dakota Howard et al |
| EPI_ISL_5411432 | Illinois Department of Public Health - Springfield Lab | Illinois Department of Public Health - Springfield Lab | Bryan Sim et al |

| | | | |
|---|---|---|---|
| EPI_ISL_2557222 | IN State Department of Health Laboratory Services | IN State Department of Health Laboratory Services | Cassandra Campion et al |
| EPI_ISL_4174633 | Laboratory Corporation of America | Centers for Disease Control and Prevention Division of Viral Diseases, Pathogen Discovery | Dakota Howard et al |
| EPI_ISL_4026234 | Northwestern Memorial Hospital | Northwestern University - Center for Pathogen Genomics and Microbial Evolution | Ramon Lorenzo-Redondo et al |
| EPI_ISL_4979864 | Aegis Sciences Corporation | Centers for Disease Control and Prevention Division of Viral Diseases, Pathogen Discovery | Dakota Howard et al |
| EPI_ISL_5990985 | Gravity Diagnostics, LLC | Gravity Diagnostics, LLC | Gravity Diagnostics et al |
| EPI_ISL_2602078 | Gravity Diagnostics, LLC | Gravity Diagnostics, LLC | Gravity Diagnostics et al |
| EPI_ISL_1823412 | Indiana Animal Disease Diagnostic Laboratory | Carpi Laboratory - Purdue University | Jack Dorman et al |
| EPI_ISL_2755661 | Curative Labs | Curative Labs | Mikhail Hanewich-Hollatz et al |
| EPI_ISL_2371711 | Quest Diagnostics Incorporated | Centers for Disease Control and Prevention Division of Viral Diseases, Pathogen Discovery | Dakota Howard et al |
| EPI_ISL_4160111 | Laboratory Corporation of America | Centers for Disease Control and Prevention Division of Viral Diseases, Pathogen Discovery | Dakota Howard et al |
| EPI_ISL_3021308 | Kansas Health and Environmental Lab | Kansas Health and Environmental Lab | Katherine Wiggins et al |
| EPI_ISL_6531124 | Laboratory Corporation of America | Centers for Disease Control and Prevention Division of Viral Diseases, Pathogen Discovery | Dakota Howard et al |
| EPI_ISL_4360136 | Aegis Sciences Corporation | Centers for Disease Control and Prevention Division of Viral Diseases, Pathogen Discovery | Dakota Howard et al |
| EPI_ISL_1799931 | Laboratory Corporation of America | Centers for Disease Control and Prevention Division of Viral Diseases, Pathogen Discovery | Dakota Howard et al |
| EPI_ISL_6583497 | Bluewater Diagnostic Laboratory, LLC | University of Louisville Sequencing Technology Center | Elizabeth Hudson et al |
| EPI_ISL_2249936 | Reditus Laboratories | Reditus Laboratories | Joshua J. Geltz et al |
| EPI_ISL_4163429 | Laboratory Corporation of America | Centers for Disease Control and Prevention Division of Viral Diseases, Pathogen Discovery | Dakota Howard et al |
| EPI_ISL_2422231 | Kansas Health and Environmental Lab | Kansas Health and Environmental Lab | Katherine Wiggins et al |
| EPI_ISL_7125727 | Kansas Health and Environmental Lab | Kansas Health and Environmental Lab | Mike Grose et al |
| EPI_ISL_4220158 | Kansas Health and Environmental Lab | Kansas Health and Environmental Lab | Mike Grose et al |
| EPI_ISL_6550255 | Aegis Sciences Corporation | Centers for Disease Control and Prevention Division of Viral Diseases, Pathogen Discovery | Dakota Howard et al |
| EPI_ISL_2148233 | Aegis Sciences Corporation | Centers for Disease Control and Prevention Division of Viral Diseases, Pathogen Discovery | Dakota Howard et al |

| | | | |
|---|---|---|---|
| EPI_ISL_5172450 | Laboratory Corporation of America | Centers for Disease Control and Prevention Division of Viral Diseases, Pathogen Discovery | Dakota Howard et al |
| EPI_ISL_6792112 | Laboratory Corporation of America | Centers for Disease Control and Prevention Division of Viral Diseases, Pathogen Discovery | Dakota Howard et al |
| EPI_ISL_2696498 | Laboratory, Gravity Diagnostics | Laboratory, Gravity Diagnostics | Diagnostics et al |
| EPI_ISL_2003665 | Kansas Health and Environmental Lab | Kansas Health and Environmental Lab | Mike Grose et al |
| EPI_ISL_3674602 | Laboratory Corporation of America | Centers for Disease Control and Prevention Division of Viral Diseases, Pathogen Discovery | Dakota Howard et al |
| EPI_ISL_2147147 | Aegis Sciences Corporation | Centers for Disease Control and Prevention Division of Viral Diseases, Pathogen Discovery | Dakota Howard et al |
| EPI_ISL_6336853 | Helix | Centers for Disease Control and Prevention Division of Viral Diseases, Pathogen Discovery | Dakota Howard et al |
| EPI_ISL_5548204 | Aegis Sciences Corporation | Centers for Disease Control and Prevention Division of Viral Diseases, Pathogen Discovery | Dakota Howard et al |
| EPI_ISL_3718561 | IN State Department of Health Laboratory Services | IN State Department of Health Laboratory Services | Cassandra Campion et al |
| EPI_ISL_3062362 | Clinical Reference Laboratory | Kansas Health and Environmental Lab | Mike Grose et al |
| EPI_ISL_1836641 | Aegis Sciences Corporation | Centers for Disease Control and Prevention Division of Viral Diseases, Pathogen Discovery | Dakota Howard et al |
| EPI_ISL_5990400 | Indiana Animal Disease Diagnostic Laboratory | Carpi Laboratory - Purdue University | Jack Dorman et al |
| EPI_ISL_1736349 | Aegis Sciences Corporation | Centers for Disease Control and Prevention Division of Viral Diseases, Pathogen Discovery | Dakota Howard et al |
| EPI_ISL_1823122 | Wichita State University - Molecular Diagnostics Lab | Kansas Health and Environmental Lab | Mike Grose et al |
| EPI_ISL_4341562 | Laboratory Corporation of America | Centers for Disease Control and Prevention Division of Viral Diseases, Pathogen Discovery | Dakota Howard et al |
| EPI_ISL_6054812 | Aegis Sciences Corporation | Centers for Disease Control and Prevention Division of Viral Diseases, Pathogen Discovery | Dakota Howard et al |
| EPI_ISL_6635495 | Helix | Centers for Disease Control and Prevention Division of Viral Diseases, Pathogen Discovery | Dakota Howard et al |
| EPI_ISL_1161207 | Laboratory Corporation of America | Respiratory Viruses Branch, Division of Viral Diseases, Centers for Disease Control and Prevention | Peter W. Cook et al |
| EPI_ISL_6070106 | Aegis Sciences Corporation | Centers for Disease Control and Prevention Division of Viral Diseases, Pathogen Discovery | Dakota Howard et al |

| | | | |
|---|---|---|---|
| EPI_ISL_4372821 | Quest Diagnostics Incorporated | Centers for Disease Control and Prevention Division of Viral Diseases, Pathogen Discovery | Dakota Howard et al |
| EPI_ISL_4987666 | Aegis Sciences Corporation | Centers for Disease Control and Prevention Division of Viral Diseases, Pathogen Discovery | Dakota Howard et al |
| EPI_ISL_4539249 | Laboratory Corporation of America | Centers for Disease Control and Prevention Division of Viral Diseases, Pathogen Discovery | Dakota Howard et al |
| EPI_ISL_3088548 | Kansas Health and Environmental Lab | Kansas Health and Environmental Lab | Katherine Wiggins et al |
| EPI_ISL_2150101 | Aegis Sciences Corporation | Centers for Disease Control and Prevention Division of Viral Diseases, Pathogen Discovery | Dakota Howard et al |
| EPI_ISL_1480286 | Helix/Illumina | Centers for Disease Control and Prevention Division of Viral Diseases, Pathogen Discovery | Dakota Howard et al |
| EPI_ISL_6388975 | Laboratory Corporation of America | Centers for Disease Control and Prevention Division of Viral Diseases, Pathogen Discovery | Dakota Howard et al |
| EPI_ISL_1550233 | Laboratory Corporation of America | Centers for Disease Control and Prevention Division of Viral Diseases, Pathogen Discovery | Dakota Howard et al |
| EPI_ISL_1550132 | Laboratory Corporation of America | Centers for Disease Control and Prevention Division of Viral Diseases, Pathogen Discovery | Dakota Howard et al |
| EPI_ISL_3863051 | Aegis Sciences Corporation | Centers for Disease Control and Prevention Division of Viral Diseases, Pathogen Discovery | Dakota Howard et al |
| EPI_ISL_5094764 | Illinois Department of Public Health - Springfield Lab | Illinois Department of Public Health - Springfield Lab | Bryan Sim et al |
| EPI_ISL_2398966 | Laboratory Corporation of America | Centers for Disease Control and Prevention Division of Viral Diseases, Pathogen Discovery | Dakota Howard et al |
| EPI_ISL_5115529 | Gravity Diagnostics, LLC | Gravity Diagnostics, LLC | Gravity Diagnostics et al |
| EPI_ISL_4990705 | Aegis Sciences Corporation | Centers for Disease Control and Prevention Division of Viral Diseases, Pathogen Discovery | Dakota Howard et al |
| EPI_ISL_4376660 | Aegis Sciences Corporation | Centers for Disease Control and Prevention Division of Viral Diseases, Pathogen Discovery | Dakota Howard et al |
| EPI_ISL_5405971 | IN State Department of Health Laboratory Services | IN State Department of Health Laboratory Services | Cassandra Campion et al |
| EPI_ISL_2180496 | Quest Diagnostics Incorporated | Centers for Disease Control and Prevention Division of Viral Diseases, Pathogen Discovery | Dakota Howard et al |
| EPI_ISL_5990306 | Indiana Animal Disease Diagnostic Laboratory | Carpi Laboratory - Purdue University | Jack Dorman et al |
| EPI_ISL_1527951 | IN State Department of Health Laboratory Services | IN State Department of Health Laboratory Services | Cassandra Campion et al |

| | | | |
|---|---|---|---|
| EPI_ISL_4368048 | Quest Diagnostics Incorporated | Centers for Disease Control and Prevention Division of Viral Diseases, Pathogen Discovery | Dakota Howard et al |
| EPI_ISL_2296004 | Simple Laboratories | Gagnon Lab, Southern Illinois University | Keith Gagnon et al |
| EPI_ISL_3905609 | Idaho Bureau of Laboratories | Center for Global Health, University of New Mexico Health Sciences Center | Daryl Domman et al |
| EPI_ISL_6387346 | Laboratory Corporation of America | Centers for Disease Control and Prevention Division of Viral Diseases, Pathogen Discovery | Dakota Howard et al |
| EPI_ISL_4403683 | IN State Department of Health Laboratory Services | IN State Department of Health Laboratory Services | Cassandra Campion et al |
| EPI_ISL_4241147 | Laboratory Corporation of America | Centers for Disease Control and Prevention Division of Viral Diseases, Pathogen Discovery | Dakota Howard et al |
| EPI_ISL_1998136 | Aegis Sciences Corporation | Centers for Disease Control and Prevention Division of Viral Diseases, Pathogen Discovery | Dakota Howard et al |
| EPI_ISL_3849943 | Aegis Sciences Corporation | Centers for Disease Control and Prevention Division of Viral Diseases, Pathogen Discovery | Dakota Howard et al |
| EPI_ISL_5411951 | Illinois Department of Public Health | Illinois Department of Public Health - Chicago Lab | Vineet K. Dhiman et al |
| EPI_ISL_848235 | Illinois Department of Public Health | Gagnon Lab, Southern Illinois University | Keith Gagnon et al |
| EPI_ISL_5573463 | Laboratory Corporation of America | Centers for Disease Control and Prevention Division of Viral Diseases, Pathogen Discovery | Dakota Howard et al |
| EPI_ISL_4455023 | Broad Institute Clinical Research Sequencing Platform | Infectious Disease Program, Broad Institute of Harvard and MIT | Siddle et al |
| EPI_ISL_4094407 | Broad Institute Clinical Research Sequencing Platform | Infectious Disease Program, Broad Institute of Harvard and MIT | Siddle et al |
| EPI_ISL_6802555 | Broad Institute Clinical Research Sequencing Platform | Infectious Disease Program, Broad Institute of Harvard and MIT | Siddle et al |
| EPI_ISL_4465965 | Willis-Knighton Medical Center Hospital Laboratory | LSUHS Emerging Viral Threat Laboratory | Lorie M. Atkins et al |
| EPI_ISL_3084707 | Broad Institute Clinical Research Sequencing Platform | Infectious Disease Program, Broad Institute of Harvard and MIT | Siddle et al |
| EPI_ISL_3404971 | Ochsner Health | BioInfoExperts | Amy Feehan et al |
| EPI_ISL_1710436 | Broad Institute Clinical Research Sequencing Platform | Infectious Disease Program, Broad Institute of Harvard and MIT | Siddle et al |
| EPI_ISL_460192 | Massachusetts General Hospital | Infectious Disease Program, Broad Institute of Harvard and MIT | Lemieux et al |
| EPI_ISL_2038185 | Fulgent Genetics | Centers for Disease Control and Prevention Division of Viral Diseases, Pathogen Discovery | Dakota Howard et al |
| EPI_ISL_5008592 | Broad Institute Clinical Research Sequencing Platform | Infectious Disease Program, Broad Institute of Harvard and MIT | Siddle et al |
| EPI_ISL_3493902 | Boston University CTL | Boston University/National Emerging Infectious Disease Laboratories | Jacquelyn Turcinovic et al |

| | | | |
|---|---|---|---|
| EPI_ISL_4729576 | Broad Institute Clinical Research Sequencing Platform | Infectious Disease Program, Broad Institute of Harvard and MIT | Siddle et al |
| EPI_ISL_6399370 | Quest Diagnostics Incorporated | Centers for Disease Control and Prevention Division of Viral Diseases, Pathogen Discovery | Dakota Howard et al |
| EPI_ISL_6754961 | Broad Institute Clinical Research Sequencing Platform | Infectious Disease Program, Broad Institute of Harvard and MIT | Siddle et al |
| EPI_ISL_5938663 | Broad Institute Clinical Research Sequencing Platform | Infectious Disease Program, Broad Institute of Harvard and MIT | Siddle et al |
| EPI_ISL_5144113 | Broad Institute Clinical Research Sequencing Platform | Infectious Disease Program, Broad Institute of Harvard and MIT | Siddle et al |
| EPI_ISL_4285360 | Broad Institute Clinical Research Sequencing Platform | Infectious Disease Program, Broad Institute of Harvard and MIT | Siddle et al |
| EPI_ISL_5454626 | Broad Institute Clinical Research Sequencing Platform | Infectious Disease Program, Broad Institute of Harvard and MIT | Siddle et al |
| EPI_ISL_5081782 | Quest Diagnostics Incorporated | Centers for Disease Control and Prevention Division of Viral Diseases, Pathogen Discovery | Dakota Howard et al |
| EPI_ISL_5571642 | Laboratory Corporation of America | Centers for Disease Control and Prevention Division of Viral Diseases, Pathogen Discovery | Dakota Howard et al |
| EPI_ISL_6514890 | Aegis Sciences Corporation | Centers for Disease Control and Prevention Division of Viral Diseases, Pathogen Discovery | Dakota Howard et al |
| EPI_ISL_1040080 | LSUHS Emerging Viral Threat Laboratory | Microbial Genome Sequencing Center | Jeremy P. Kamil et al |
| EPI_ISL_7135915 | Broad Institute Clinical Research Sequencing Platform | Infectious Disease Program, Broad Institute of Harvard and MIT | Siddle et al |
| EPI_ISL_5019598 | Broad Institute Clinical Research Sequencing Platform | Infectious Disease Program, Broad Institute of Harvard and MIT | Siddle et al |
| EPI_ISL_4488259 | Broad Institute Clinical Research Sequencing Platform | Infectious Disease Program, Broad Institute of Harvard and MIT | Siddle et al |
| EPI_ISL_3839008 | Broad Institute Clinical Research Sequencing Platform | Infectious Disease Program, Broad Institute of Harvard and MIT | Siddle et al |
| EPI_ISL_5009074 | Broad Institute Clinical Research Sequencing Platform | Infectious Disease Program, Broad Institute of Harvard and MIT | Siddle et al |
| EPI_ISL_1710454 | Broad Institute Clinical Research Sequencing Platform | Infectious Disease Program, Broad Institute of Harvard and MIT | Siddle et al |
| EPI_ISL_4729027 | Broad Institute Clinical Research Sequencing Platform | Infectious Disease Program, Broad Institute of Harvard and MIT | Siddle et al |
| EPI_ISL_6754914 | Broad Institute Clinical Research Sequencing Platform | Infectious Disease Program, Broad Institute of Harvard and MIT | Siddle et al |
| EPI_ISL_1335991 | Quest Diagnostics Incorporated | Centers for Disease Control and Prevention Division of Viral Diseases, Pathogen Discovery | Peter W. Cook et al |
| EPI_ISL_3525515 | Broad Institute Clinical Research Sequencing Platform | Infectious Disease Program, Broad Institute of Harvard and MIT | Siddle et al |
| EPI_ISL_5089115 | Laboratory Corporation of America | Centers for Disease Control and Prevention Division of Viral Diseases, Pathogen Discovery | Dakota Howard et al |
| EPI_ISL_5390806 | Baton Rouge Clinic | LSUHS Emerging Viral Threat Laboratory | Gregory L. Ware et al |

| | | | |
|---|---|---|---|
| EPI_ISL_3937727 | Broad Institute Clinical Research Sequencing Platform | Infectious Disease Program, Broad Institute of Harvard and MIT | Siddle et al |
| EPI_ISL_4095509 | Broad Institute Clinical Research Sequencing Platform | Infectious Disease Program, Broad Institute of Harvard and MIT | Siddle et al |
| EPI_ISL_7171810 | Quest Diagnostics Incorporated | Centers for Disease Control and Prevention Division of Viral Diseases, Pathogen Discovery | Dakota Howard et al |
| EPI_ISL_4729269 | Broad Institute Clinical Research Sequencing Platform | Infectious Disease Program, Broad Institute of Harvard and MIT | Siddle et al |
| EPI_ISL_5019588 | Broad Institute Clinical Research Sequencing Platform | Infectious Disease Program, Broad Institute of Harvard and MIT | Siddle et al |
| EPI_ISL_4093784 | Broad Institute Clinical Research Sequencing Platform | Infectious Disease Program, Broad Institute of Harvard and MIT | Siddle et al |
| EPI_ISL_1610325 | Laboratory Corporation of America | Centers for Disease Control and Prevention Division of Viral Diseases, Pathogen Discovery | Dakota Howard et al |
| EPI_ISL_4094970 | Broad Institute Clinical Research Sequencing Platform | Infectious Disease Program, Broad Institute of Harvard and MIT | Siddle et al |
| EPI_ISL_2840242 | Willis-Knighton Medical Center Hospital Laboratory | LSUHS Emerging Viral Threat Laboratory | Lorie M. Atkins et al |
| EPI_ISL_579002 | LSUHS Emerging Viral Threat Laboratory | Microbial Genome Sequencing Center | Jeremy P. Kamil et al |
| EPI_ISL_5193723 | Ochsner Health | BioInfoExperts | Amy Feehan et al |
| EPI_ISL_3410671 | UMass Memorial Medical Center | Center for Microbiome Research | Richard T. Ellison III et al |
| EPI_ISL_1320362 | Laboratory Corporation of America | Centers for Disease Control and Prevention Division of Viral Diseases, Pathogen Discovery | Peter W. Cook et al |
| EPI_ISL_1463275 | Laboratory Corporation of America | Centers for Disease Control and Prevention Division of Viral Diseases, Pathogen Discovery | Dakota Howard et al |
| EPI_ISL_5385896 | Broad Institute Clinical Research Sequencing Platform | Infectious Disease Program, Broad Institute of Harvard and MIT | Siddle et al |
| EPI_ISL_5518038 | Aegis Sciences Corporation | Centers for Disease Control and Prevention Division of Viral Diseases, Pathogen Discovery | Dakota Howard et al |
| EPI_ISL_4988241 | Aegis Sciences Corporation | Centers for Disease Control and Prevention Division of Viral Diseases, Pathogen Discovery | Dakota Howard et al |
| EPI_ISL_2991899 | Broad Institute Clinical Research Sequencing Platform | Infectious Disease Program, Broad Institute of Harvard and MIT | Lemieux et al |
| EPI_ISL_5582458 | Aegis Sciences Corporation | Centers for Disease Control and Prevention Division of Viral Diseases, Pathogen Discovery | Dakota Howard et al |
| EPI_ISL_5511463 | Willis-Knighton Medical Center Hospital Laboratory | LSUHS Emerging Viral Threat Laboratory | Lorie M. Atkins et al |
| EPI_ISL_2243266 | Aegis Sciences Corporation | Centers for Disease Control and Prevention Division of Viral Diseases, Pathogen Discovery | Dakota Howard et al |
| EPI_ISL_3430260 | Aegis Sciences Corporation | Centers for Disease Control and Prevention Division of Viral Diseases, Pathogen Discovery | Dakota Howard et al |

| | | | |
|---|---|---|---|
| EPI_ISL_2440717 | Quest Diagnostics Incorporated | Centers for Disease Control and Prevention Division of Viral Diseases, Pathogen Discovery | Dakota Howard et al |
| EPI_ISL_6794473 | Willis-Knighton Medical Center Hospital Laboratory | LSUHS Emerging Viral Threat Laboratory | Lorie M. Atkins et al |
| EPI_ISL_3409033 | Broad Institute Clinical Research Sequencing Platform | Infectious Disease Program, Broad Institute of Harvard and MIT | Siddle et al |
| EPI_ISL_6209522 | Broad Institute Clinical Research Sequencing Platform | Infectious Disease Program, Broad Institute of Harvard and MIT | Siddle et al |
| EPI_ISL_5744643 | Broad Institute Clinical Research Sequencing Platform | Infectious Disease Program, Broad Institute of Harvard and MIT | Siddle et al |
| EPI_ISL_6209652 | Broad Institute Clinical Research Sequencing Platform | Infectious Disease Program, Broad Institute of Harvard and MIT | Siddle et al |
| EPI_ISL_3623095 | Gravity Diagnostics, LLC | Gravity Diagnostics, LLC | Gravity Diagnostics et al |
| EPI_ISL_5952592 | Gravity Diagnostics, LLC | Gravity Diagnostics, LLC | Gravity Diagnostics et al |
| EPI_ISL_4455271 | Broad Institute Clinical Research Sequencing Platform | Infectious Disease Program, Broad Institute of Harvard and MIT | Siddle et al |
| EPI_ISL_2311878 | Broad Institute Clinical Research Sequencing Platform | Infectious Disease Program, Broad Institute of Harvard and MIT | Siddle et al |
| EPI_ISL_1011592 | Massachusetts General Hospital | Infectious Disease Program, Broad Institute of Harvard and MIT | Lemieux et al |
| EPI_ISL_4890521 | UMass Memorial Medical Center | Center for Microbiome Research | Richard T. Ellison III et al |
| EPI_ISL_693103 | Massachusetts State Public Health Laboratory | Massachusetts State Public Health Laboratory | Andrew Lang et al |
| EPI_ISL_5019757 | Broad Institute Clinical Research Sequencing Platform | Infectious Disease Program, Broad Institute of Harvard and MIT | Siddle et al |
| EPI_ISL_3007437 | Broad Institute Clinical Research Sequencing Platform | Infectious Disease Program, Broad Institute of Harvard and MIT | Siddle et al |
| EPI_ISL_3492006 | Broad Institute Clinical Research Sequencing Platform | Infectious Disease Program, Broad Institute of Harvard and MIT | Siddle et al |
| EPI_ISL_854937 | Quest Diagnostics | Quest Diagnostics | Rosenthal et al |
| EPI_ISL_3525857 | Broad Institute Clinical Research Sequencing Platform | Infectious Disease Program, Broad Institute of Harvard and MIT | Siddle et al |
| EPI_ISL_1971865 | Broad Institute Clinical Research Sequencing Platform | Infectious Disease Program, Broad Institute of Harvard and MIT | Siddle et al |
| EPI_ISL_6602954 | Broad Institute Clinical Research Sequencing Platform | Infectious Disease Program, Broad Institute of Harvard and MIT | Siddle et al |
| EPI_ISL_1918608 | Broad Institute Clinical Research Sequencing Platform | Infectious Disease Program, Broad Institute of Harvard and MIT | Siddle et al |
| EPI_ISL_6802606 | Broad Institute Clinical Research Sequencing Platform | Infectious Disease Program, Broad Institute of Harvard and MIT | Siddle et al |
| EPI_ISL_4304112 | Broad Institute Clinical Research Sequencing Platform | Infectious Disease Program, Broad Institute of Harvard and MIT | Siddle et al |
| EPI_ISL_3431442 | Broad Institute Clinical Research Sequencing Platform | Infectious Disease Program, Broad Institute of Harvard and MIT | Siddle et al |
| EPI_ISL_4381151 | Laboratory Corporation of America | Centers for Disease Control and Prevention Division of Viral Diseases, Pathogen Discovery | Dakota Howard et al |
| EPI_ISL_5878965 | Laboratory Corporation of America | Centers for Disease Control and Prevention Division of Viral Diseases, Pathogen Discovery | Dakota Howard et al |

| | | | |
|---|---|---|---|
| EPI_ISL_1838447 | Aegis Sciences Corporation | Centers for Disease Control and Prevention Division of Viral Diseases, Pathogen Discovery | Dakota Howard et al |
| EPI_ISL_2374824 | Michigan Department of Health and Human Services, Bureau of Laboratories | Michigan Department of Health and Human Services, Bureau of Laboratories | Blankenship HM et al |
| EPI_ISL_1168397 | VA Connecticut Healthcare System | Grubaugh Lab - Yale School of Public Health | Mary Petrone et al |
| EPI_ISL_5925597 | Michigan Department of Health and Human Services, Bureau of Laboratories | Michigan Department of Health and Human Services, Bureau of Laboratories | Blankenship HM et al |
| EPI_ISL_6789032 | Laboratory Corporation of America | Centers for Disease Control and Prevention Division of Viral Diseases, Pathogen Discovery | Dakota Howard et al |
| EPI_ISL_1999342 | Aegis Sciences Corporation | Centers for Disease Control and Prevention Division of Viral Diseases, Pathogen Discovery | Dakota Howard et al |
| EPI_ISL_5367757 | Michigan Department of Health and Human Services, Bureau of Laboratories | Michigan Department of Health and Human Services, Bureau of Laboratories | Blankenship HM et al |
| EPI_ISL_1008807 | Michigan Department of Health and Human Services, Bureau of Laboratories | Michigan Department of Health and Human Services, Bureau of Laboratories | Blankenship HM et al |
| EPI_ISL_4381251 | Aegis Sciences Corporation | Centers for Disease Control and Prevention Division of Viral Diseases, Pathogen Discovery | Dakota Howard et al |
| EPI_ISL_1994260 | Michigan Department of Health and Human Services, Bureau of Laboratories | Michigan Department of Health and Human Services, Bureau of Laboratories | Blankenship HM et al |
| EPI_ISL_5412357 | Johns Hopkins Hospital Department of Pathology | Johns Hopkins Hospital Department of Pathology | C. Paul Morris et al |
| EPI_ISL_1662810 | Johns Hopkins Hospital Department of Pathology | Johns Hopkins Hospital Department of Pathology | C. Paul Morris et al |
| EPI_ISL_5516471 | Aegis Sciences Corporation | Centers for Disease Control and Prevention Division of Viral Diseases, Pathogen Discovery | Dakota Howard et al |
| EPI_ISL_2481468 | Laboratory Corporation of America | Centers for Disease Control and Prevention Division of Viral Diseases, Pathogen Discovery | Dakota Howard et al |
| EPI_ISL_2959471 | Michigan Department of Health and Human Services, Bureau of Laboratories | Michigan Department of Health and Human Services, Bureau of Laboratories | Blankenship HM et al |
| EPI_ISL_1576156 | Helix/Illumina | Centers for Disease Control and Prevention Division of Viral Diseases, Pathogen Discovery | Dakota Howard et al |
| EPI_ISL_1305862 | Johns Hopkins Hospital Department of Pathology | Johns Hopkins Hospital Department of Pathology | C. Paul Morris et al |
| EPI_ISL_1444841 | Maryland Genomics, Institute for Genome Sciences, University of Maryland School of Medicine | Maryland Genomics, Institute for Genome Sciences, University of Maryland School of Medicine | Tallon et al |

| | | | |
|---|---|---|---|
| EPI_ISL_2802299 | Maine Health and Environmental Testing Laboratory | Tewhey Lab, The Jackson Laboratory | Matluk et al |
| EPI_ISL_3814533 | Johns Hopkins Hospital Department of Pathology | Johns Hopkins Hospital Department of Pathology | C. Paul Morris et al |
| EPI_ISL_507424 | Michigan Department of Health and Human Services, Bureau of Laboratories | Michigan Department of Health and Human Services, Bureau of Laboratories | Blankenship HM et al |
| EPI_ISL_3129455 | Michigan Department of Health and Human Services, Bureau of Laboratories | Michigan Department of Health and Human Services, Bureau of Laboratories | Blankenship HM et al |
| EPI_ISL_5591485 | University of Michigan Clinical Microbiology Laboratory | Lauring Lab, University of Michigan, Department of Microbiology and Immunology | Gilbert et al |
| EPI_ISL_5108333 | Aegis Sciences Corporation | Centers for Disease Control and Prevention Division of Viral Diseases, Pathogen Discovery | Dakota Howard et al |
| EPI_ISL_3425514 | Maine Health and Environmental Testing Laboratory | Tewhey Lab, The Jackson Laboratory | Matluk et al |
| EPI_ISL_1996847 | Aegis Sciences Corporation | Centers for Disease Control and Prevention Division of Viral Diseases, Pathogen Discovery | Dakota Howard et al |
| EPI_ISL_5412485 | Johns Hopkins Hospital Department of Pathology | Johns Hopkins Hospital Department of Pathology | C. Paul Morris et al |
| EPI_ISL_1838226 | Aegis Sciences Corporation | Centers for Disease Control and Prevention Division of Viral Diseases, Pathogen Discovery | Dakota Howard et al |
| EPI_ISL_1906750 | Michigan Department of Health and Human Services, Bureau of Laboratories | Michigan Department of Health and Human Services, Bureau of Laboratories | Blankenship HM et al |
| EPI_ISL_5170316 | Laboratory Corporation of America | Centers for Disease Control and Prevention Division of Viral Diseases, Pathogen Discovery | Dakota Howard et al |
| EPI_ISL_2455289 | Johns Hopkins Hospital Department of Pathology | Johns Hopkins Hospital Department of Pathology | C. Paul Morris et al |
| EPI_ISL_3922742 | Michigan Department of Health and Human Services, Bureau of Laboratories | Michigan Department of Health and Human Services, Bureau of Laboratories | Blankenship HM et al |
| EPI_ISL_1923320 | Helix/Illumina | Centers for Disease Control and Prevention Division of Viral Diseases, Pathogen Discovery | Dakota Howard et al |
| EPI_ISL_2534120 | Maryland Genomics, Institute for Genome Sciences, University of Maryland School of Medicine | Maryland Genomics, Institute for Genome Sciences, University of Maryland School of Medicine | Tallon et al |
| EPI_ISL_4169036 | University of Michigan Clinical Microbiology Laboratory | University of Michigan, Department of Microbiology and Immunology, Lauring Lab | Gilbert et al |
| EPI_ISL_5318261 | Mako Medical | Centers for Disease Control and Prevention Division of Viral Diseases, Pathogen Discovery | Dakota Howard et al |

| | | | |
|---|---|---|---|
| EPI_ISL_4760310 | Maine Health and Environmental Testing Laboratory | Tewhey Lab, The Jackson Laboratory | Matluk et al |
| EPI_ISL_955779 | University of Michigan Clinical Microbiology Laboratory | Lauring Lab, University of Michigan, Department of Microbiology and Immunology | Valesano et al |
| EPI_ISL_5413607 | Johns Hopkins Hospital Department of Pathology | Johns Hopkins Hospital Department of Pathology | C. Paul Morris et al |
| EPI_ISL_4239512 | Michigan Department of Health and Human Services, Bureau of Laboratories | Michigan Department of Health and Human Services, Bureau of Laboratories | Blankenship HM et al |
| EPI_ISL_886394 | Labcorp | Genomics and Discovery, Respiratory Viruses Branch, Division of Viral Diseases, Centers for Disease Control and Prevention | Peter W. Cook et al |
| EPI_ISL_2243970 | Aegis Sciences Corporation | Centers for Disease Control and Prevention Division of Viral Diseases, Pathogen Discovery | Dakota Howard et al |
| EPI_ISL_1994339 | Michigan Department of Health and Human Services, Bureau of Laboratories | Michigan Department of Health and Human Services, Bureau of Laboratories | Blankenship HM et al |
| EPI_ISL_2148687 | Aegis Sciences Corporation | Centers for Disease Control and Prevention Division of Viral Diseases, Pathogen Discovery | Dakota Howard et al |
| EPI_ISL_6391658 | Laboratory Corporation of America | Centers for Disease Control and Prevention Division of Viral Diseases, Pathogen Discovery | Dakota Howard et al |
| EPI_ISL_3425357 | Maine Health and Environmental Testing Laboratory | Tewhey Lab, The Jackson Laboratory | Matluk et al |
| EPI_ISL_5553798 | Aegis Sciences Corporation | Centers for Disease Control and Prevention Division of Viral Diseases, Pathogen Discovery | Dakota Howard et al |
| EPI_ISL_507925 | Michigan Department of Health and Human Services, Bureau of Laboratories | Michigan Department of Health and Human Services, Bureau of Laboratories | Blankenship HM et al |
| EPI_ISL_6648312 | Aegis Sciences Corporation | Centers for Disease Control and Prevention Division of Viral Diseases, Pathogen Discovery | Dakota Howard et al |
| EPI_ISL_5367117 | Michigan Department of Health and Human Services, Bureau of Laboratories | Michigan Department of Health and Human Services, Bureau of Laboratories | Blankenship HM et al |
| EPI_ISL_5431775 | Michigan Department of Health and Human Services, Bureau of Laboratories | Michigan Department of Health and Human Services, Bureau of Laboratories | Blankenship HM et al |
| EPI_ISL_4535795 | Aegis Sciences Corporation | Centers for Disease Control and Prevention Division of Viral Diseases, Pathogen Discovery | Dakota Howard et al |
| EPI_ISL_1995994 | Aegis Sciences Corporation | Centers for Disease Control and Prevention Division of Viral Diseases, Pathogen Discovery | Dakota Howard et al |

| | | | |
|---|---|---|---|
| EPI_ISL_1900420 | Aegis Sciences Corporation | Centers for Disease Control and Prevention Division of Viral Diseases, Pathogen Discovery | Dakota Howard et al |
| EPI_ISL_1058707 | Michigan Department of Health and Human Services, Bureau of Laboratories | Michigan Department of Health and Human Services, Bureau of Laboratories | Blankenship HM et al |
| EPI_ISL_1405479 | Johns Hopkins Hospital Department of Pathology | Johns Hopkins Hospital Department of Pathology | C. Paul Morris et al |
| EPI_ISL_6906553 | Laboratory Corporation of America | Centers for Disease Control and Prevention Division of Viral Diseases, Pathogen Discovery | Dakota Howard et al |
| EPI_ISL_4141685 | Aegis Sciences Corporation | Centers for Disease Control and Prevention Division of Viral Diseases, Pathogen Discovery | Dakota Howard et al |
| EPI_ISL_2959510 | Michigan Department of Health and Human Services, Bureau of Laboratories | Michigan Department of Health and Human Services, Bureau of Laboratories | Blankenship HM et al |
| EPI_ISL_1563553 | Aegis Sciences Corporation | Centers for Disease Control and Prevention Division of Viral Diseases, Pathogen Discovery | Dakota Howard et al |
| EPI_ISL_7128302 | Helix | Centers for Disease Control and Prevention Division of Viral Diseases, Pathogen Discovery | Dakota Howard et al |
| EPI_ISL_1675170 | Department of Laboratory Medicine, Clinical Center, National Institutes of Health | Laboratory of Parasitic Diseases, Systems Genomics Section, National Institute of Allergy and Infectious Diseases, National Institutes of Health | Allison Roder et al |
| EPI_ISL_3009141 | Maryland Genomics, Institute for Genome Sciences, University of Maryland School of Medicine | Maryland Genomics, Institute for Genome Sciences, University of Maryland School of Medicine | Tallon et al |
| EPI_ISL_3755575 | Laboratory Corporation of America | Centers for Disease Control and Prevention Division of Viral Diseases, Pathogen Discovery | Dakota Howard et al |
| EPI_ISL_3864502 | Mako Medical | Centers for Disease Control and Prevention Division of Viral Diseases, Pathogen Discovery | Dakota Howard et al |
| EPI_ISL_5090981 | Laboratory Corporation of America | Centers for Disease Control and Prevention Division of Viral Diseases, Pathogen Discovery | Dakota Howard et al |
| EPI_ISL_4239667 | Michigan Department of Health and Human Services, Bureau of Laboratories | Michigan Department of Health and Human Services, Bureau of Laboratories | Blankenship HM et al |
| EPI_ISL_2959145 | Michigan Department of Health and Human Services, Bureau of Laboratories | Michigan Department of Health and Human Services, Bureau of Laboratories | Blankenship HM et al |
| EPI_ISL_1017605 | Connecticut Veterans' Affairs Hospital | Grubaugh Lab - Yale School of Public Health | Mary Petrone et al |

| | | | |
|---|---|---|---|
| EPI_ISL_2319521 | Maryland Genomics, Institute for Genome Sciences, University of Maryland School of Medicine | Maryland Genomics, Institute for Genome Sciences, University of Maryland School of Medicine | Tallon et al |
| EPI_ISL_1796467 | Helix/Illumina | Centers for Disease Control and Prevention Division of Viral Diseases, Pathogen Discovery | Dakota Howard et al |
| EPI_ISL_5368047 | Michigan Department of Health and Human Services, Bureau of Laboratories | Michigan Department of Health and Human Services, Bureau of Laboratories | Blankenship HM et al |
| EPI_ISL_6377669 | Aegis Sciences Corporation | Centers for Disease Control and Prevention Division of Viral Diseases, Pathogen Discovery | Dakota Howard et al |
| EPI_ISL_5412155 | Johns Hopkins Hospital Department of Pathology | Johns Hopkins Hospital Department of Pathology | C. Paul Morris et al |
| EPI_ISL_6341875 | Aegis Sciences Corporation | Centers for Disease Control and Prevention Division of Viral Diseases, Pathogen Discovery | Dakota Howard et al |
| EPI_ISL_1561519 | Aegis Sciences Corporation | Centers for Disease Control and Prevention Division of Viral Diseases, Pathogen Discovery | Dakota Howard et al |
| EPI_ISL_5871812 | Laboratory Corporation of America | Centers for Disease Control and Prevention Division of Viral Diseases, Pathogen Discovery | Dakota Howard et al |
| EPI_ISL_3948811 | Maryland Genomics, Institute for Genome Sciences, University of Maryland School of Medicine | Maryland Genomics, Institute for Genome Sciences, University of Maryland School of Medicine | Tallon et al |
| EPI_ISL_1905446 | Michigan Department of Health and Human Services, Bureau of Laboratories | Michigan Department of Health and Human Services, Bureau of Laboratories | Blankenship HM et al |
| EPI_ISL_7128637 | Helix | Centers for Disease Control and Prevention Division of Viral Diseases, Pathogen Discovery | Dakota Howard et al |
| EPI_ISL_1576584 | Helix/Illumina | Centers for Disease Control and Prevention Division of Viral Diseases, Pathogen Discovery | Dakota Howard et al |
| EPI_ISL_604541 | Quest Diagnostics | Quest Diagnostics | Rosenthal et al |
| EPI_ISL_6421321 | Aegis Sciences Corporation | Centers for Disease Control and Prevention Division of Viral Diseases, Pathogen Discovery | Dakota Howard et al |
| EPI_ISL_6892988 | Fulgent Genetics | Centers for Disease Control and Prevention Division of Viral Diseases, Pathogen Discovery | Dakota Howard et al |
| EPI_ISL_4953152 | "MN PHL Division, Minnesota Department of Health" | Centers for Disease Control and Prevention Division of Viral Diseases, Pathogen Discovery | Mili Sheth et al |
| EPI_ISL_6633568 | Helix | Centers for Disease Control and Prevention Division of Viral Diseases, Pathogen Discovery | Dakota Howard et al |

| | | | |
|---|---|---|---|
| EPI_ISL_1493168 | MD PHL | MD PHL | Maryland Department of Health Laboratories Administration et al |
| EPI_ISL_2040135 | Aegis Sciences Corporation | Centers for Disease Control and Prevention Division of Viral Diseases, Pathogen Discovery | Dakota Howard et al |
| EPI_ISL_4150171 | Laboratory Corporation of America | Centers for Disease Control and Prevention Division of Viral Diseases, Pathogen Discovery | Dakota Howard et al |
| EPI_ISL_4760266 | Maine Health and Environmental Testing Laboratory | Tewhey Lab, The Jackson Laboratory | Matluk et al |
| EPI_ISL_5928663 | Quest Diagnostics Incorporated | Centers for Disease Control and Prevention Division of Viral Diseases, Pathogen Discovery | Dakota Howard et al |
| EPI_ISL_1493081 | Michigan Department of Health and Human Services, Bureau of Laboratories | Michigan Department of Health and Human Services, Bureau of Laboratories | Blankenship HM et al |
| EPI_ISL_3637588 | Michigan Department of Health and Human Services, Bureau of Laboratories | Michigan Department of Health and Human Services, Bureau of Laboratories | Blankenship HM et al |
| EPI_ISL_2090251 | Aegis Sciences Corporation | Centers for Disease Control and Prevention Division of Viral Diseases, Pathogen Discovery | Dakota Howard et al |
| EPI_ISL_2009929 | Helix/Illumina | Centers for Disease Control and Prevention Division of Viral Diseases, Pathogen Discovery | Dakota Howard et al |
| EPI_ISL_4556041 | Maryland Genomics, Institute for Genome Sciences, University of Maryland School of Medicine | Maryland Genomics, Institute for Genome Sciences, University of Maryland School of Medicine | Tallon et al |
| EPI_ISL_1372755 | Maine Health and Environmental Testing Laboratory | Tewhey Lab, The Jackson Laboratory | Matluk et al |
| EPI_ISL_5370424 | Aegis Sciences Corporation | Centers for Disease Control and Prevention Division of Viral Diseases, Pathogen Discovery | Dakota Howard et al |
| EPI_ISL_4990353 | Aegis Sciences Corporation | Centers for Disease Control and Prevention Division of Viral Diseases, Pathogen Discovery | Dakota Howard et al |
| EPI_ISL_5205207 | Helix | Centers for Disease Control and Prevention Division of Viral Diseases, Pathogen Discovery | Dakota Howard et al |
| EPI_ISL_1550997 | Aegis Sciences Corporation | Centers for Disease Control and Prevention Division of Viral Diseases, Pathogen Discovery | Dakota Howard et al |
| EPI_ISL_6902389 | Helix | Centers for Disease Control and Prevention Division of Viral Diseases, Pathogen Discovery | Dakota Howard et al |
| EPI_ISL_6786731 | Aegis Sciences Corporation | Centers for Disease Control and Prevention Division of Viral Diseases, Pathogen Discovery | Dakota Howard et al |

| | | | |
|---|---|---|---|
| EPI_ISL_1994334 | Michigan Department of Health and Human Services, Bureau of Laboratories | Michigan Department of Health and Human Services, Bureau of Laboratories | Blankenship HM et al |
| EPI_ISL_2959647 | Michigan Department of Health and Human Services, Bureau of Laboratories | Michigan Department of Health and Human Services, Bureau of Laboratories | Blankenship HM et al |
| EPI_ISL_6640205 | Missouri State Public Health Laboratory | Missouri State Public Health Laboratory | Matthew Sinn et al |
| EPI_ISL_4822979 | Infinity Biologix | Centers for Disease Control and Prevention Division of Viral Diseases, Pathogen Discovery | Dakota Howard et al |
| EPI_ISL_2014462 | Infinity Biologix | Centers for Disease Control and Prevention Division of Viral Diseases, Pathogen Discovery | Dakota Howard et al |
| EPI_ISL_1031813 | Laboratory Corporation of America | Respiratory Viruses Branch, Division of Viral Diseases, Centers for Disease Control and Prevention | Peter W. Cook et al |
| EPI_ISL_4914009 | Aegis Sciences Corporation | Centers for Disease Control and Prevention Division of Viral Diseases, Pathogen Discovery | Dakota Howard et al |
| EPI_ISL_6527313 | Aegis Sciences Corporation | Centers for Disease Control and Prevention Division of Viral Diseases, Pathogen Discovery | Dakota Howard et al |
| EPI_ISL_1693104 | Infinity Biologix | Centers for Disease Control and Prevention Division of Viral Diseases, Pathogen Discovery | Dakota Howard et al |
| EPI_ISL_5781177 | Minnesota Department of Health, Public Health Laboratory | Minnesota Department of Health, Public Health Laboratory | Alexandra Lorentz et al |
| EPI_ISL_5558619 | Infinity Biologix | Centers for Disease Control and Prevention Division of Viral Diseases, Pathogen Discovery | Dakota Howard et al |
| EPI_ISL_4385980 | Centracare Laboratory Services | Minnesota Department of Health, Public Health Laboratory | Alexandra Lorentz et al |
| EPI_ISL_6306051 | Minnesota Department of Health, Public Health Laboratory | Minnesota Department of Health, Public Health Laboratory | Alexandra Lorentz et al |
| EPI_ISL_6382994 | Infinity Biologix | Centers for Disease Control and Prevention Division of Viral Diseases, Pathogen Discovery | Dakota Howard et al |
| EPI_ISL_3019925 | Minnesota Department of Health, Public Health Laboratory | Minnesota Department of Health, Public Health Laboratory | Alexandra Lorentz et al |
| EPI_ISL_2652629 | Infinity Biologix | Centers for Disease Control and Prevention Division of Viral Diseases, Pathogen Discovery | Dakota Howard et al |
| EPI_ISL_2753816 | Missouri State Public Health Laboratory | Missouri State Public Health Laboratory | Matthew Sinn et al |
| EPI_ISL_4049587 | Allina Health Laboratory | Minnesota Department of Health, Public Health Laboratory | Alexandra Lorentz et al |

| | | | |
|---|---|---|---|
| EPI_ISL_5053041 | University of Mississippi Medical Center, Department of Pathology | University of Mississippi Medical Center, Molecular and Genomics Core Facility | Ashley C. Johnson et al |
| EPI_ISL_4367539 | Quest Diagnostics Incorporated | Centers for Disease Control and Prevention Division of Viral Diseases, Pathogen Discovery | Dakota Howard et al |
| EPI_ISL_3372484 | Minnesota Department of Health, Public Health Laboratory | Minnesota Department of Health, Public Health Laboratory | Alexandra Lorentz et al |
| EPI_ISL_4591981 | Regions Hospital | Minnesota Department of Health, Public Health Laboratory | Alexandra Lorentz et al |
| EPI_ISL_5100154 | Infinity Biologix | Centers for Disease Control and Prevention Division of Viral Diseases, Pathogen Discovery | Dakota Howard et al |
| EPI_ISL_3111025 | Infinity Biologix | Centers for Disease Control and Prevention Division of Viral Diseases, Pathogen Discovery | Dakota Howard et al |
| EPI_ISL_5209518 | Infinity Biologix | Centers for Disease Control and Prevention Division of Viral Diseases, Pathogen Discovery | Dakota Howard et al |
| EPI_ISL_4079894 | M Health Fairview | Minnesota Department of Health, Public Health Laboratory | Alexandra Lorentz et al |
| EPI_ISL_4062499 | Minnesota Department of Health, Public Health Laboratory | Minnesota Department of Health, Public Health Laboratory | Alexandra Lorentz et al |
| EPI_ISL_3396072 | Quest Diagnostics Incorporated | Centers for Disease Control and Prevention Division of Viral Diseases, Pathogen Discovery | Dakota Howard et al |
| EPI_ISL_1691615 | Infinity Biologix | Centers for Disease Control and Prevention Division of Viral Diseases, Pathogen Discovery | Dakota Howard et al |
| EPI_ISL_2241979 | Aegis Sciences Corporation | Centers for Disease Control and Prevention Division of Viral Diseases, Pathogen Discovery | Dakota Howard et al |
| EPI_ISL_3353489 | Infinity Biologix | Centers for Disease Control and Prevention Division of Viral Diseases, Pathogen Discovery | Dakota Howard et al |
| EPI_ISL_1702346 | Infinity Biologix | Centers for Disease Control and Prevention Division of Viral Diseases, Pathogen Discovery | Dakota Howard et al |
| EPI_ISL_5980656 | Infinity Biologix | Centers for Disease Control and Prevention Division of Viral Diseases, Pathogen Discovery | Dakota Howard et al |
| EPI_ISL_3523480 | Infinity Biologix | Centers for Disease Control and Prevention Division of Viral Diseases, Pathogen Discovery | Dakota Howard et al |
| EPI_ISL_1693424 | Infinity Biologix | Centers for Disease Control and Prevention Division of Viral Diseases, Pathogen Discovery | Dakota Howard et al |
| EPI_ISL_6383195 | Infinity Biologix | Centers for Disease Control and Prevention Division of Viral Diseases, Pathogen Discovery | Dakota Howard et al |

| | | | |
|---|---|---|---|
| EPI_ISL_4246359 | Infinity Biologix | Centers for Disease Control and Prevention Division of Viral Diseases, Pathogen Discovery | Dakota Howard et al |
| EPI_ISL_4821134 | University of Mississippi Medical Center, Department of Pathology | University of Mississippi Medical Center, Molecular and Genomics Core Facility | Ashley C. Johnson et al |
| EPI_ISL_2858802 | MS Public Health Laboratory | University of Mississippi Medical Center, Molecular and Genomics Core Facility | Ashley C. Johnson et al |
| EPI_ISL_1314334 | Vault Health | Minnesota Department of Health, Public Health Laboratory | Alexandra Lorentz et al |
| EPI_ISL_2014290 | Infinity Biologix | Centers for Disease Control and Prevention Division of Viral Diseases, Pathogen Discovery | Dakota Howard et al |
| EPI_ISL_5162997 | Allina Health Laboratory | Minnesota Department of Health, Public Health Laboratory | Alexandra Lorentz et al |
| EPI_ISL_1164501 | Vault Health | Minnesota Department of Health, Public Health Laboratory | Alexandra Lorentz et al |
| EPI_ISL_6105960 | Infinity Biologix | Centers for Disease Control and Prevention Division of Viral Diseases, Pathogen Discovery | Dakota Howard et al |
| EPI_ISL_6607360 | Public Health Laboratory, Minnesota Department of Health | University of Minnesota Genomics Center | Daryl M. Gohl et al |
| EPI_ISL_2014718 | Infinity Biologix | Centers for Disease Control and Prevention Division of Viral Diseases, Pathogen Discovery | Dakota Howard et al |
| EPI_ISL_1732814 | Infinity Biologix | Centers for Disease Control and Prevention Division of Viral Diseases, Pathogen Discovery | Dakota Howard et al |
| EPI_ISL_4808367 | Infinity Biologix | Centers for Disease Control and Prevention Division of Viral Diseases, Pathogen Discovery | Dakota Howard et al |
| EPI_ISL_2014851 | Infinity Biologix | Centers for Disease Control and Prevention Division of Viral Diseases, Pathogen Discovery | Dakota Howard et al |
| EPI_ISL_5235379 | Infinity Biologix | Centers for Disease Control and Prevention Division of Viral Diseases, Pathogen Discovery | Dakota Howard et al |
| EPI_ISL_4211389 | Minnesota Department of Health, Public Health Laboratory | Minnesota Department of Health, Public Health Laboratory | Alexandra Lorentz et al |
| EPI_ISL_5207030 | Infinity Biologix | Centers for Disease Control and Prevention Division of Viral Diseases, Pathogen Discovery | Dakota Howard et al |
| EPI_ISL_5347989 | Sonic Reference Laboratory | Sonic Reference Laboratory | "Robert Moller et al |
| EPI_ISL_2186290 | Infinity Biologix | Centers for Disease Control and Prevention Division of Viral Diseases, Pathogen Discovery | Dakota Howard et al |
| EPI_ISL_5420388 | Missouri State Public Health Laboratory | Missouri State Public Health Laboratory | Matthew Sinn et al |

| | | | |
|---|---|---|---|
| EPI_ISL_6410428 | Public Health Laboratory, Minnesota Department of Health | University of Minnesota Genomics Center | Daryl M. Gohl et al |
| EPI_ISL_2497888 | University of Mississippi Medical Center, Department of Pathology | University of Mississippi Medical Center, Molecular and Genomics Core Facility | Ashley C. Johnson et al |
| EPI_ISL_6906993 | Quest Diagnostics Incorporated | Centers for Disease Control and Prevention Division of Viral Diseases, Pathogen Discovery | Dakota Howard et al |
| EPI_ISL_5207239 | Infinity Biologix | Centers for Disease Control and Prevention Division of Viral Diseases, Pathogen Discovery | Dakota Howard et al |
| EPI_ISL_5560369 | Infinity Biologix | Centers for Disease Control and Prevention Division of Viral Diseases, Pathogen Discovery | Dakota Howard et al |
| EPI_ISL_1702250 | Infinity Biologix | Centers for Disease Control and Prevention Division of Viral Diseases, Pathogen Discovery | Dakota Howard et al |
| EPI_ISL_2132396 | Infinity Biologix | Centers for Disease Control and Prevention Division of Viral Diseases, Pathogen Discovery | Dakota Howard et al |
| EPI_ISL_2282975 | Infinity Biologix | Centers for Disease Control and Prevention Division of Viral Diseases, Pathogen Discovery | Dakota Howard et al |
| EPI_ISL_5209581 | Infinity Biologix | Centers for Disease Control and Prevention Division of Viral Diseases, Pathogen Discovery | Dakota Howard et al |
| EPI_ISL_6400689 | Infinity Biologix | Centers for Disease Control and Prevention Division of Viral Diseases, Pathogen Discovery | Dakota Howard et al |
| EPI_ISL_1064500 | M Health Fairview | Minnesota Department of Health, Public Health Laboratory | Alexandra Lorentz et al |
| EPI_ISL_2090537 | Quest Diagnostics Incorporated | Centers for Disease Control and Prevention Division of Viral Diseases, Pathogen Discovery | Dakota Howard et al |
| EPI_ISL_5420856 | Missouri State Public Health Laboratory | Missouri State Public Health Laboratory | Matthew Sinn et al |
| EPI_ISL_5923417 | Minnesota Department of Health, Public Health Laboratory | Minnesota Department of Health, Public Health Laboratory | Alexandra Lorentz et al |
| EPI_ISL_1733340 | Infinity Biologix | Centers for Disease Control and Prevention Division of Viral Diseases, Pathogen Discovery | Dakota Howard et al |
| EPI_ISL_4179426 | Laboratory Corporation of America | Centers for Disease Control and Prevention Division of Viral Diseases, Pathogen Discovery | Dakota Howard et al |
| EPI_ISL_4808301 | Infinity Biologix | Centers for Disease Control and Prevention Division of Viral Diseases, Pathogen Discovery | Dakota Howard et al |
| EPI_ISL_6653795 | Infinity Biologix | Centers for Disease Control and Prevention Division of Viral Diseases, Pathogen Discovery | Dakota Howard et al |

| | | | |
|---|---|---|---|
| EPI_ISL_2148763 | Aegis Sciences Corporation | Centers for Disease Control and Prevention Division of Viral Diseases, Pathogen Discovery | Dakota Howard et al |
| EPI_ISL_6640149 | Missouri State Public Health Laboratory | Missouri State Public Health Laboratory | Matthew Sinn et al |
| EPI_ISL_1372125 | M Health Fairview | Minnesota Department of Health, Public Health Laboratory | Alexandra Lorentz et al |
| EPI_ISL_2557584 | Infinity Biologix | Centers for Disease Control and Prevention Division of Viral Diseases, Pathogen Discovery | Dakota Howard et al |
| EPI_ISL_2132370 | Infinity Biologix | Centers for Disease Control and Prevention Division of Viral Diseases, Pathogen Discovery | Dakota Howard et al |
| EPI_ISL_6989662 | Vault Health | Minnesota Department of Health, Public Health Laboratory | Jacob Garfin et al |
| EPI_ISL_1733082 | Infinity Biologix | Centers for Disease Control and Prevention Division of Viral Diseases, Pathogen Discovery | Dakota Howard et al |
| EPI_ISL_3419131 | StarMed Healthcare | UNC Charlotte Environmental Monitoring Laboratory | Jessica Schlueter et al |
| EPI_ISL_6420113 | Aegis Sciences Corporation | Centers for Disease Control and Prevention Division of Viral Diseases, Pathogen Discovery | Dakota Howard et al |
| EPI_ISL_2305884 | Laboratory Corporation of America | Centers for Disease Control and Prevention Division of Viral Diseases, Pathogen Discovery | Dakota Howard et al |
| EPI_ISL_2376152 | Montana Public Health Laboratory | Montana Public Health Laboratory | Joy Ritter et al |
| EPI_ISL_2651136 | Nebraska Public Health Laboratory | NPHL COVID-19 Response Team | NPHL COVID-19 Response Team et al |
| EPI_ISL_2335039 | North Dakota Department of Health, Public Health Laboratory | North Dakota Department of Health, Public Health Laboratory | Lisa Wingerter et al |
| EPI_ISL_6062268 | Aegis Sciences Corporation | Centers for Disease Control and Prevention Division of Viral Diseases, Pathogen Discovery | Dakota Howard et al |
| EPI_ISL_5170561 | Laboratory Corporation of America | Centers for Disease Control and Prevention Division of Viral Diseases, Pathogen Discovery | Dakota Howard et al |
| EPI_ISL_4818946 | Laboratory Corporation of America | Centers for Disease Control and Prevention Division of Viral Diseases, Pathogen Discovery | Dakota Howard et al |
| EPI_ISL_4910666 | Aegis Sciences Corporation | Centers for Disease Control and Prevention Division of Viral Diseases, Pathogen Discovery | Dakota Howard et al |
| EPI_ISL_6914452 | Montana Public Health Laboratory | Montana Public Health Laboratory | Joy Ritter et al |
| EPI_ISL_4484206 | Laboratory Corporation of America | Centers for Disease Control and Prevention Division of Viral Diseases, Pathogen Discovery | Dakota Howard et al |

| | | | |
|---|---|---|---|
| EPI_ISL_978773 | Helix/Illumina | Respiratory Viruses Branch, Division of Viral Diseases, Centers for Disease Control and Prevention | Peter W. Cook et al |
| EPI_ISL_1689385 | Aegis Sciences Corporation | Centers for Disease Control and Prevention Division of Viral Diseases, Pathogen Discovery | Dakota Howard et al |
| EPI_ISL_2306314 | Laboratory Corporation of America | Centers for Disease Control and Prevention Division of Viral Diseases, Pathogen Discovery | Dakota Howard et al |
| EPI_ISL_1513393 | Aegis Sciences Corporation | Centers for Disease Control and Prevention Division of Viral Diseases, Pathogen Discovery | Dakota Howard et al |
| EPI_ISL_6208081 | Molecular Biology Laboratory, Vidant Medical Center | Brody Integrative Genomics Core, East Carolina University | Weihua Huang et al |
| EPI_ISL_5081986 | Quest Diagnostics Incorporated | Centers for Disease Control and Prevention Division of Viral Diseases, Pathogen Discovery | Dakota Howard et al |
| EPI_ISL_4196406 | Mako Medical | Centers for Disease Control and Prevention Division of Viral Diseases, Pathogen Discovery | Dakota Howard et al |
| EPI_ISL_1020819 | Laboratory Corporation of America | Respiratory Viruses Branch, Division of Viral Diseases, Centers for Disease Control and Prevention | Peter W. Cook et al |
| EPI_ISL_6174358 | Molecular Biology Laboratory, Vidant Medical Center | Brody Integrative Genomics Core, East Carolina University | Weihua Huang et al |
| EPI_ISL_1818280 | Quest Diagnostics Incorporated | Centers for Disease Control and Prevention Division of Viral Diseases, Pathogen Discovery | Dakota Howard et al |
| EPI_ISL_4888892 | US Air Force School of Aerospace Medicine | US Air Force School of Aerospace Medicine | Anthony Fries et al |
| EPI_ISL_5252444 | Broad Institute Clinical Research Sequencing Platform | Infectious Disease Program, Broad Institute of Harvard and MIT | Siddle et al |
| EPI_ISL_6908186 | Quest Diagnostics Incorporated | Centers for Disease Control and Prevention Division of Viral Diseases, Pathogen Discovery | Dakota Howard et al |
| EPI_ISL_3682639 | Laboratory Corporation of America | Centers for Disease Control and Prevention Division of Viral Diseases, Pathogen Discovery | Dakota Howard et al |
| EPI_ISL_4847025 | Clinical Molecular Microbiology Laboratory, UNC Hospitals | Jeremy Wang | Jeremy Wang et al |
| EPI_ISL_6644784 | Aegis Sciences Corporation | Centers for Disease Control and Prevention Division of Viral Diseases, Pathogen Discovery | Dakota Howard et al |
| EPI_ISL_5174192 | Laboratory Corporation of America | Centers for Disease Control and Prevention Division of Viral Diseases, Pathogen Discovery | Dakota Howard et al |
| EPI_ISL_3925326 | StarMed Healthcare | UNC Charlotte Environmental Monitoring Laboratory | Jessica Schlueter et al |
| EPI_ISL_2201444 | Aegis Sciences Corporation | Centers for Disease Control and Prevention Division of Viral Diseases, Pathogen Discovery | Dakota Howard et al |

| | | | |
|---|---|---|---|
| EPI_ISL_6415301 | Quest Diagnostics Incorporated | Centers for Disease Control and Prevention Division of Viral Diseases, Pathogen Discovery | Dakota Howard et al |
| EPI_ISL_1996737 | Aegis Sciences Corporation | Centers for Disease Control and Prevention Division of Viral Diseases, Pathogen Discovery | Dakota Howard et al |
| EPI_ISL_6383814 | Aegis Sciences Corporation | Centers for Disease Control and Prevention Division of Viral Diseases, Pathogen Discovery | Dakota Howard et al |
| EPI_ISL_2090950 | North Dakota Department of Health, Public Health Laboratory | North Dakota Department of Health, Public Health Laboratory | Lisa Wingerter et al |
| EPI_ISL_2785019 | Aegis Sciences Corporation | Centers for Disease Control and Prevention Division of Viral Diseases, Pathogen Discovery | Dakota Howard et al |
| EPI_ISL_4485335 | Laboratory Corporation of America | Centers for Disease Control and Prevention Division of Viral Diseases, Pathogen Discovery | Dakota Howard et al |
| EPI_ISL_6061998 | Aegis Sciences Corporation | Centers for Disease Control and Prevention Division of Viral Diseases, Pathogen Discovery | Dakota Howard et al |
| EPI_ISL_4564806 | Clinical Molecular Microbiology Laboratory, UNC Hospitals | Jeremy Wang | Jeremy Wang et al |
| EPI_ISL_3858122 | Aegis Sciences Corporation | Centers for Disease Control and Prevention Division of Viral Diseases, Pathogen Discovery | Dakota Howard et al |
| EPI_ISL_5586452 | Quest Diagnostics Incorporated | Centers for Disease Control and Prevention Division of Viral Diseases, Pathogen Discovery | Dakota Howard et al |
| EPI_ISL_3259663 | Montana Public Health Laboratory | Montana Public Health Laboratory | Joy Ritter et al |
| EPI_ISL_3514408 | Laboratory Corporation of America | Centers for Disease Control and Prevention Division of Viral Diseases, Pathogen Discovery | Dakota Howard et al |
| EPI_ISL_5370394 | Aegis Sciences Corporation | Centers for Disease Control and Prevention Division of Viral Diseases, Pathogen Discovery | Dakota Howard et al |
| EPI_ISL_5170975 | Altru Health System | Genomics Core University of North Dakota | Sara Strandquist et al |
| EPI_ISL_5093539 | Quest Diagnostics Incorporated | Centers for Disease Control and Prevention Division of Viral Diseases, Pathogen Discovery | Dakota Howard et al |
| EPI_ISL_3333727 | University of Montana Genomics Core (UMGC) | University of Montana Genomics Core (UMGC) | William Conner et al |
| EPI_ISL_4157637 | Laboratory Corporation of America | Centers for Disease Control and Prevention Division of Viral Diseases, Pathogen Discovery | Dakota Howard et al |
| EPI_ISL_4820929 | Screening laboratory, UNC,Chapel Hill | Dirk Dittmer | Razia Moorad et al |

| | | | |
|---|---|---|---|
| EPI_ISL_1267613 | Helix/Illumina | Centers for Disease Control and Prevention Division of Viral Diseases, Pathogen Discovery | Peter W. Cook et al |
| EPI_ISL_5080888 | Laboratory Corporation of America | Centers for Disease Control and Prevention Division of Viral Diseases, Pathogen Discovery | Dakota Howard et al |
| EPI_ISL_4554585 | Aegis Sciences Corporation | Centers for Disease Control and Prevention Division of Viral Diseases, Pathogen Discovery | Dakota Howard et al |
| EPI_ISL_6158702 | Molecular Biology Laboratory, Vidant Medical Center | Brody Integrative Genomics Core, East Carolina University | Weihua Huang et al |
| EPI_ISL_4167563 | Mako Medical | Centers for Disease Control and Prevention Division of Viral Diseases, Pathogen Discovery | Dakota Howard et al |
| EPI_ISL_2229254 | NC State Laboratory of Public Health | Centers for Disease Control and Prevention Division of Viral Diseases, Pathogen Discovery | Mili Sheth et al |
| EPI_ISL_1462329 | Laboratory Corporation of America | Centers for Disease Control and Prevention Division of Viral Diseases, Pathogen Discovery | Dakota Howard et al |
| EPI_ISL_5108547 | Aegis Sciences Corporation | Centers for Disease Control and Prevention Division of Viral Diseases, Pathogen Discovery | Dakota Howard et al |
| EPI_ISL_1692308 | Infinity Biologix | Centers for Disease Control and Prevention Division of Viral Diseases, Pathogen Discovery | Dakota Howard et al |
| EPI_ISL_4816370 | Laboratory Corporation of America | Centers for Disease Control and Prevention Division of Viral Diseases, Pathogen Discovery | Dakota Howard et al |
| EPI_ISL_984312 | NCSLPH | NCSLPH | Chase K et al |
| EPI_ISL_2854246 | NCSLPH | NCSLPH | Chase K et al |
| EPI_ISL_6421865 | Aegis Sciences Corporation | Centers for Disease Control and Prevention Division of Viral Diseases, Pathogen Discovery | Dakota Howard et al |
| EPI_ISL_3905789 | Montana DPHHS Public Health Laboratory | Center for Global Health, University of New Mexico Health Sciences Center | Daryl Domman et al |
| EPI_ISL_4398893 | Aegis Sciences Corporation | Centers for Disease Control and Prevention Division of Viral Diseases, Pathogen Discovery | Dakota Howard et al |
| EPI_ISL_5517830 | Aegis Sciences Corporation | Centers for Disease Control and Prevention Division of Viral Diseases, Pathogen Discovery | Dakota Howard et al |
| EPI_ISL_7116718 | Nebraska Public Health Laboratory | NPHL COVID-19 Response Team | NPHL COVID-19 Response Team et al |
| EPI_ISL_6071165 | Aegis Sciences Corporation | Centers for Disease Control and Prevention Division of Viral Diseases, Pathogen Discovery | Dakota Howard et al |
| EPI_ISL_1667617 | Aegis Sciences Corporation | Centers for Disease Control and Prevention Division of Viral Diseases, Pathogen Discovery | Dakota Howard et al |

| | | | |
|---|---|---|---|
| EPI_ISL_4379090 | Aegis Sciences Corporation | Centers for Disease Control and Prevention Division of Viral Diseases, Pathogen Discovery | Dakota Howard et al |
| EPI_ISL_6535524 | Laboratory Corporation of America | Centers for Disease Control and Prevention Division of Viral Diseases, Pathogen Discovery | Dakota Howard et al |
| EPI_ISL_5323760 | Molecular Biology Laboratory, Vidant Medical Center | Brody Integrative Genomics Core, East Carolina University | Weihua Huang et al |
| EPI_ISL_4955440 | Infinity Biologix | Centers for Disease Control and Prevention Division of Viral Diseases, Pathogen Discovery | Dakota Howard et al |
| EPI_ISL_7116413 | Nebraska Public Health Laboratory | NPHL COVID-19 Response Team | NPHL COVID-19 Response Team et al |
| EPI_ISL_2150060 | Aegis Sciences Corporation | Centers for Disease Control and Prevention Division of Viral Diseases, Pathogen Discovery | Dakota Howard et al |
| EPI_ISL_4150579 | Laboratory Corporation of America | Centers for Disease Control and Prevention Division of Viral Diseases, Pathogen Discovery | Dakota Howard et al |
| EPI_ISL_3752998 | Aegis Sciences Corporation | Centers for Disease Control and Prevention Division of Viral Diseases, Pathogen Discovery | Dakota Howard et al |
| EPI_ISL_2482295 | Aegis Sciences Corporation | Centers for Disease Control and Prevention Division of Viral Diseases, Pathogen Discovery | Dakota Howard et al |
| EPI_ISL_7116681 | Nebraska Public Health Laboratory | NPHL COVID-19 Response Team | NPHL COVID-19 Response Team et al |
| EPI_ISL_2869151 | Quest Diagnostics Incorporated | Centers for Disease Control and Prevention Division of Viral Diseases, Pathogen Discovery | Dakota Howard et al |
| EPI_ISL_4877449 | Nevada State Public Health Laboratory | Nevada State Public Health Laboratory | Andrew Gorzalski et al |
| EPI_ISL_1114712 | New Mexico Department of Health Scientific Laboratory | New Mexico Department of Health Scientific Laboratory | Ellie Johnson et al |
| EPI_ISL_6414814 | Nevada State Public Health Laboratory | Nevada State Public Health Laboratory | Andrew Gorzalski et al |
| EPI_ISL_4376712 | Laboratory Corporation of America | Centers for Disease Control and Prevention Division of Viral Diseases, Pathogen Discovery | Dakota Howard et al |
| EPI_ISL_5069120 | Quest Diagnostics Incorporated | Centers for Disease Control and Prevention Division of Viral Diseases, Pathogen Discovery | Dakota Howard et al |
| EPI_ISL_3825305 | Nevada State Public Health Laboratory | Nevada State Public Health Laboratory | Andrew Gorzalski et al |
| EPI_ISL_5200433 | NJDOH, Public Health and Environmental Laboratories | NJ_PHEL | Ryan Pachucki et al |
| EPI_ISL_3755278 | Laboratory Corporation of America | Centers for Disease Control and Prevention Division of Viral Diseases, Pathogen Discovery | Dakota Howard et al |
| EPI_ISL_3751654 | Laboratory Corporation of America | Centers for Disease Control and Prevention Division of Viral Diseases, Pathogen Discovery | Dakota Howard et al |

| | | | |
|---|---|---|---|
| EPI_ISL_802376 | MSHS Clinical Microbiology Laboratories | MSHS Pathogen Surveillance Program | Ana S. Gonzalez-Reiche et al |
| EPI_ISL_5998907 | New Mexico Department of Health Scientific Laboratory | Center for Global Health, University of New Mexico Health Sciences Center | Daryl Domman et al |
| EPI_ISL_5010239 | Nevada State Public Health Laboratory | Nevada State Public Health Laboratory | Andrew Gorzalski et al |
| EPI_ISL_2280774 | Aegis Sciences Corporation | Centers for Disease Control and Prevention Division of Viral Diseases, Pathogen Discovery | Dakota Howard et al |
| EPI_ISL_4371264 | Quest Diagnostics Incorporated | Centers for Disease Control and Prevention Division of Viral Diseases, Pathogen Discovery | Dakota Howard et al |
| EPI_ISL_6068346 | Aegis Sciences Corporation | Centers for Disease Control and Prevention Division of Viral Diseases, Pathogen Discovery | Dakota Howard et al |
| EPI_ISL_5611072 | Fulgent Genetics | Centers for Disease Control and Prevention Division of Viral Diseases, Pathogen Discovery | Dakota Howard et al |
| EPI_ISL_937263 | NYC HH Elmhurst Hospital Medical Center | New York City Public Health Laboratory | Jade Wang et al |
| EPI_ISL_4372567 | Quest Diagnostics Incorporated | Centers for Disease Control and Prevention Division of Viral Diseases, Pathogen Discovery | Dakota Howard et al |
| EPI_ISL_2871553 | Quest Diagnostics Incorporated | Centers for Disease Control and Prevention Division of Viral Diseases, Pathogen Discovery | Dakota Howard et al |
| EPI_ISL_5145150 | Helix | Centers for Disease Control and Prevention Division of Viral Diseases, Pathogen Discovery | Dakota Howard et al |
| EPI_ISL_4399312 | Aegis Sciences Corporation | Centers for Disease Control and Prevention Division of Viral Diseases, Pathogen Discovery | Dakota Howard et al |
| EPI_ISL_4157980 | Laboratory Corporation of America | Centers for Disease Control and Prevention Division of Viral Diseases, Pathogen Discovery | Dakota Howard et al |
| EPI_ISL_6261569 | Nevada State Public Health Laboratory | Nevada State Public Health Laboratory | Andrew Gorzalski et al |
| EPI_ISL_3232292 | Clinical Microbiology Laboratory, NewYork Presbyterian Hospital/Columbia University Irving Medical Center | Uhlemann Laboratory, Columbia University Irving Medical Center | Medini K. Annavajhala et al |
| EPI_ISL_5153075 | SUNY Upstate Medical University | SUNY Upstate Medical University | Frank Middleton et al |
| EPI_ISL_1136662 | Nevada State Public Health Laboratory | Nevada State Public Health Laboratory | Andrew Gorzalski et al |
| EPI_ISL_6930869 | NYU Langone Health | Departments of Pathology and Medicine, New York University School of Medicine | Adriana Heguy et al |

| | | | |
|---|---|---|---|
| EPI_ISL_3018913 | Quest Diagnostics Incorporated | Centers for Disease Control and Prevention Division of Viral Diseases, Pathogen Discovery | Dakota Howard et al |
| EPI_ISL_6545583 | Aegis Sciences Corporation | Centers for Disease Control and Prevention Division of Viral Diseases, Pathogen Discovery | Dakota Howard et al |
| EPI_ISL_631601 | Jamaica Hospital Medical Center | New York City Public Health Laboratory | Jade Wang et al |
| EPI_ISL_2500654 | Clinical Microbiology Laboratory, NewYork Presbyterian Hospital/Columbia University Irving Medical Center | Uhlemann Laboratory, Columbia University Irving Medical Center | Medini K. Annavajhala et al |
| EPI_ISL_5629416 | New Mexico Department of Health Scientific Laboratory | Center for Global Health, University of New Mexico Health Sciences Center | Daryl Domman et al |
| EPI_ISL_456060 | NYU Langone Health | Departments of Pathology and Medicine, New York University School of Medicine | Maria Aguero-Rosenfeld et al |
| EPI_ISL_5879779 | Nevada State Public Health Laboratory | Nevada State Public Health Laboratory | Andrew Gorzalski et al |
| EPI_ISL_6794024 | Laboratory Corporation of America | Centers for Disease Control and Prevention Division of Viral Diseases, Pathogen Discovery | Dakota Howard et al |
| EPI_ISL_3364212 | Biotia | Biotia | Dorottya Nagy-Szakal et al |
| EPI_ISL_5155919 | SUNY Upstate Medical University | 505 Irving Avenue, Syracuse, NY 13210 - Frank Middleton Lab | Frank Middleton et al |
| EPI_ISL_5364868 | Nevada State Public Health Laboratory | Nevada State Public Health Laboratory | Andrew Gorzalski et al |
| EPI_ISL_1255943 | New Mexico Department of Health Scientific Laboratory | New Mexico Department of Health Scientific Laboratory | Ellie Johnson et al |
| EPI_ISL_4498140 | Hospital of the University of Pennsylvania Molecular Pathology Lab | Bushman | John K. Everett et al |
| EPI_ISL_6492424 | MSHS Clinical Microbiology Laboratories | MSHS Pathogen Surveillance Program | Ana S. Gonzalez-Reiche et al |
| EPI_ISL_6087145 | Laboratory Corporation of America | Centers for Disease Control and Prevention Division of Viral Diseases, Pathogen Discovery | Dakota Howard et al |
| EPI_ISL_5643503 | NYU Langone Health | Departments of Pathology and Medicine, New York University School of Medicine | Adriana Heguy et al |
| EPI_ISL_3372179 | Nevada State Public Health Laboratory | Nevada State Public Health Laboratory | Andrew Gorzalski et al |
| EPI_ISL_6540682 | Laboratory Corporation of America | Centers for Disease Control and Prevention Division of Viral Diseases, Pathogen Discovery | Dakota Howard et al |
| EPI_ISL_5199694 | SUNY Upstate Medical University | SUNY Upstate Medical University - Frank Middleton | Frank Middleton et al |
| EPI_ISL_2759353 | Population Medicine and Diagnostic Sciences, Cornell University | Population Medicine and Diagnostic Sciences, Cornell University | Mitchell et al |

| | | | |
|---|---|---|---|
| EPI_ISL_1968600 | Weill Cornell Medicine | New York Genome Center | Michael Zody et al |
| EPI_ISL_671683 | DOHMH Corona | New York City Public Health Laboratory | Jade Wang et al |
| EPI_ISL_6315290 | NYU Langone Health | Departments of Pathology and Medicine, New York University School of Medicine | Adriana Heguy et al |
| EPI_ISL_4359637 | Fulgent Genetics | Centers for Disease Control and Prevention Division of Viral Diseases, Pathogen Discovery | Dakota Howard et al |
| EPI_ISL_1805364 | Curative | New Mexico Department of Health Scientific Laboratory | Ellie Johnson et al |
| EPI_ISL_2458181 | NYU Langone Health | Departments of Pathology and Medicine, New York University School of Medicine | Adriana Heguy et al |
| EPI_ISL_5364379 | Nevada State Public Health Laboratory | Nevada State Public Health Laboratory | Andrew Gorzalski et al |
| EPI_ISL_2147844 | Aegis Sciences Corporation | Centers for Disease Control and Prevention Division of Viral Diseases, Pathogen Discovery | Dakota Howard et al |
| EPI_ISL_6493510 | MSHS Clinical Microbiology Laboratories | MSHS Pathogen Surveillance Program | Ana S. Gonzalez-Reiche et al |
| EPI_ISL_5599401 | Infinity Biologix | Centers for Disease Control and Prevention Division of Viral Diseases, Pathogen Discovery | Dakota Howard et al |
| EPI_ISL_5876903 | Laboratory Corporation of America | Centers for Disease Control and Prevention Division of Viral Diseases, Pathogen Discovery | Dakota Howard et al |
| EPI_ISL_3825284 | Nevada State Public Health Laboratory | Nevada State Public Health Laboratory | Andrew Gorzalski et al |
| EPI_ISL_6644026 | Aegis Sciences Corporation | Centers for Disease Control and Prevention Division of Viral Diseases, Pathogen Discovery | Dakota Howard et al |
| EPI_ISL_5360313 | Nevada State Public Health Laboratory | Nevada State Public Health Laboratory | Andrew Gorzalski et al |
| EPI_ISL_2313617 | Respiratory Viruses Branch, Centers for Disease Control and Prevention | Respiratory Viruses Branch, Centers for Disease Control and Prevention | Howard et al |
| EPI_ISL_6474204 | NYC Department of Health and Mental Hygiene | Centers for Disease Control and Prevention Division of Viral Diseases, Pathogen Discovery | Mili Sheth et al |
| EPI_ISL_6493212 | MSHS Clinical Microbiology Laboratories | MSHS Pathogen Surveillance Program | Ana S. Gonzalez-Reiche et al |
| EPI_ISL_832334 | DOHMH Corona | New York City Public Health Laboratory | Jade Wang et al |
| EPI_ISL_1648233 | Quest Diagnostics Incorporated | Centers for Disease Control and Prevention Division of Viral Diseases, Pathogen Discovery | Dakota Howard et al |
| EPI_ISL_3692844 | LabQ Clinical Diagnostics LLC | New York City Public Health Laboratory | Jade Wang et al |
| EPI_ISL_6072494 | Laboratory Corporation of America | Centers for Disease Control and Prevention Division of Viral Diseases, Pathogen Discovery | Dakota Howard et al |

| | | | |
|---|---|---|---|
| EPI_ISL_3344922 | Fulgent Genetics | Centers for Disease Control and Prevention Division of Viral Diseases, Pathogen Discovery | Dakota Howard et al |
| EPI_ISL_1239772 | TriCore Reference Laboratories | Center for Global Health, University of New Mexico Health Sciences Center | Daryl Domman et al |
| EPI_ISL_3329223 | New Mexico Department of Health Scientific Laboratory | New Mexico Department of Health Scientific Laboratory | Ellie Johnson et al |
| EPI_ISL_6315298 | NYU Langone Health | Departments of Pathology and Medicine, New York University School of Medicine | Adriana Heguy et al |
| EPI_ISL_3299893 | Fulgent Genetics | Centers for Disease Control and Prevention Division of Viral Diseases, Pathogen Discovery | Dakota Howard et al |
| EPI_ISL_1680490 | Laboratory Corporation of America | Centers for Disease Control and Prevention Division of Viral Diseases, Pathogen Discovery | Dakota Howard et al |
| EPI_ISL_6629517 | Aegis Sciences Corporation | Centers for Disease Control and Prevention Division of Viral Diseases, Pathogen Discovery | Dakota Howard et al |
| EPI_ISL_1040889 | NYU Langone Health | Departments of Pathology and Medicine, New York University School of Medicine | Adriana Heguy et al |
| EPI_ISL_5155537 | SUNY Upstate Medical University | 505 Irving Avenue, Syracuse, NY 13210 - Frank Middleton Lab | Frank Middleton et al |
| EPI_ISL_5772735 | Aegis Sciences Corporation | Centers for Disease Control and Prevention Division of Viral Diseases, Pathogen Discovery | Dakota Howard et al |
| EPI_ISL_3755301 | Laboratory Corporation of America | Centers for Disease Control and Prevention Division of Viral Diseases, Pathogen Discovery | Dakota Howard et al |
| EPI_ISL_6639035 | Infinity Biologix | Centers for Disease Control and Prevention Division of Viral Diseases, Pathogen Discovery | Dakota Howard et al |
| EPI_ISL_2081115 | Hackensack Medical Center | New York Genome Center | Michael Zody et al |
| EPI_ISL_3751127 | Laboratory Corporation of America | Centers for Disease Control and Prevention Division of Viral Diseases, Pathogen Discovery | Dakota Howard et al |
| EPI_ISL_6079508 | Laboratory Corporation of America | Centers for Disease Control and Prevention Division of Viral Diseases, Pathogen Discovery | Dakota Howard et al |
| EPI_ISL_1511777 | Quest Diagnostics Incorporated | Centers for Disease Control and Prevention Division of Viral Diseases, Pathogen Discovery | Dakota Howard et al |
| EPI_ISL_6260949 | Pandemic Response Lab - NYC | Pandemic Response Lab, R&D | Henry Lee et al |
| EPI_ISL_3778639 | Pandemic Response Lab - NYC | Pandemic Response Lab, R&D | Henry Lee et al |
| EPI_ISL_3569121 | Pandemic Response Lab - NYC | Pandemic Response Lab, R&D | Henry Lee et al |
| EPI_ISL_3220531 | Aegis Sciences Corporation | Centers for Disease Control and Prevention Division of Viral Diseases, Pathogen Discovery | Dakota Howard et al |
| EPI_ISL_1172113 | Pandemic Response Lab - NYC | Pandemic Response Lab, R&D | Henry Lee et al |
| EPI_ISL_1097804 | Pandemic Response Lab - NYC | Pandemic Response Lab, R&D | Henry Lee et al |

| | | | |
|---|---|---|---|
| EPI_ISL_2200144 | The Ohio State University Applied Microbiology Services Laboratory | The Ohio State University Applied Microbiology Services Laboratory | Seth A. Faith PhD et al |
| EPI_ISL_3693101 | Yale Clinical Virology Lab | Grubaugh Lab - Yale School of Public Health | Joseph Fauver et al |
| EPI_ISL_4106658 | Pandemic Response Lab - NYC | Pandemic Response Lab, R&D | Henry Lee et al |
| EPI_ISL_5110782 | Aegis Sciences Corporation | Centers for Disease Control and Prevention Division of Viral Diseases, Pathogen Discovery | Dakota Howard et al |
| EPI_ISL_2229723 | Kaleida Health Laboratories | University at Buffalo Genomics and Bioinformatics Core | Jonathan Bard et al |
| EPI_ISL_5166062 | Laboratory Corporation of America | Centers for Disease Control and Prevention Division of Viral Diseases, Pathogen Discovery | Dakota Howard et al |
| EPI_ISL_3943011 | Pandemic Response Lab - NYC | Pandemic Response Lab, R&D | Henry Lee et al |
| EPI_ISL_1398305 | NYC Pandemic Response Lab | Wadsworth Center, New York State Department of Health | Kirsten St. George et al |
| EPI_ISL_5002391 | Aegis Sciences Corporation | Centers for Disease Control and Prevention Division of Viral Diseases, Pathogen Discovery | Dakota Howard et al |
| EPI_ISL_754483 | Wadsworth Center, New York State Department.of Health | Wadsworth Center, New York State Department.of Health | Kirsten St. George et al |
| EPI_ISL_5068451 | Quest Diagnostics Incorporated | Centers for Disease Control and Prevention Division of Viral Diseases, Pathogen Discovery | Dakota Howard et al |
| EPI_ISL_3942830 | Pandemic Response Lab - NYC | Pandemic Response Lab, R&D | Henry Lee et al |
| EPI_ISL_794043 | Wadsworth Center, New York State Department.of Health | Wadsworth Center, New York State Department.of Health | Kirsten St. George et al |
| EPI_ISL_4102475 | Kaiser Permanente NW Reginal Lab | OHSU MM Lab | Yun Wu et al |
| EPI_ISL_832456 | OHSU Lab Services Molecular Microbiology Lab | Oregon SARS-CoV-2 Genome Sequencing Center | Brendan L. O'Connell et al |
| EPI_ISL_1398005 | URMC LABS | Wadsworth Center, New York State Department of Health | Kirsten St. George et al |
| EPI_ISL_1610803 | Laboratory Corporation of America | Centers for Disease Control and Prevention Division of Viral Diseases, Pathogen Discovery | Dakota Howard et al |
| EPI_ISL_1582363 | SUNY UPSTATE MEDICAL UNIVERSITY | Wadsworth Center, New York State Department of Health | Kirsten St. George et al |
| EPI_ISL_5115256 | ALBANY MEDICAL CENTER | Wadsworth Center, New York State Department of Health | Kirsten St. George et al |
| EPI_ISL_4188358 | Quest Diagnostics Incorporated | Centers for Disease Control and Prevention Division of Viral Diseases, Pathogen Discovery | Dakota Howard et al |
| EPI_ISL_1386661 | Ohio Department of Health Laboratory | Ohio Department of Health Laboratory | Holmes et al |
| EPI_ISL_525857 | OHSU Lab Services Molecular Microbiology Lab | Oregon SARS-CoV-2 Genome Sequencing Center | Brendan L. O'Connell et al |
| EPI_ISL_1635704 | Pandemic Response Lab - NYC | Pandemic Response Lab, R&D | Henry Lee et al |
| EPI_ISL_1714760 | University Hospitals Translational Laboratory (UHTL), University Hospitals | University Hospitals Translational Laboratory (UHTL), University Hospitals | Sadri et al |

| | | | |
|---|---|---|---|
| EPI_ISL_3319690 | OHSU-MM Lab | OHSU-MM Lab | Jeannine Lama et al |
| EPI_ISL_1098109 | Pandemic Response Lab - NYC | Pandemic Response Lab, R&D | Henry Lee et al |
| EPI_ISL_4455579 | Fulgent Genetics | Centers for Disease Control and Prevention Division of Viral Diseases, Pathogen Discovery | Dakota Howard et al |
| EPI_ISL_3802055 | Gravity Diagnostics, LLC | Gravity Diagnostics, LLC | Gravity Diagnostics et al |
| EPI_ISL_4181900 | Laboratory Corporation of America | Centers for Disease Control and Prevention Division of Viral Diseases, Pathogen Discovery | Dakota Howard et al |
| EPI_ISL_2634338 | Pandemic Response Lab - NYC | Pandemic Response Lab, R&D | Henry Lee et al |
| EPI_ISL_1385007 | Pandemic Response Lab - NYC | Pandemic Response Lab, R&D | Henry Lee et al |
| EPI_ISL_6159090 | Aegis Sciences Corporation | Centers for Disease Control and Prevention Division of Viral Diseases, Pathogen Discovery | Dakota Howard et al |
| EPI_ISL_1542650 | Pandemic Response Lab - NYC | Pandemic Response Lab, R&D | Henry Lee et al |
| EPI_ISL_5317243 | OHSU Lab Services Molecular Microbiology Lab | Oregon SARS-CoV-2 Genome Sequencing Center | Brendan L. O'Connell et al |
| EPI_ISL_1717595 | Pandemic Response Lab - NYC | Pandemic Response Lab, R&D | Henry Lee et al |
| EPI_ISL_2097270 | NORTHWELL HEALTH LABORATORIES | Wadsworth Center, New York State Department of Health | Kirsten St. George et al |
| EPI_ISL_5546132 | Aegis Sciences Corporation | Centers for Disease Control and Prevention Division of Viral Diseases, Pathogen Discovery | Dakota Howard et al |
| EPI_ISL_4174109 | KSL Diagnostics | University at Buffalo Genomics and Bioinformatics Core | Jonathan Bard et al |
| EPI_ISL_1398023 | ALBANY MEDICAL CENTER | Wadsworth Center, New York State Department of Health | Kirsten St. George et al |
| EPI_ISL_2204558 | Aegis Sciences Corporation | Centers for Disease Control and Prevention Division of Viral Diseases, Pathogen Discovery | Dakota Howard et al |
| EPI_ISL_4107067 | Pandemic Response Lab - NYC | Pandemic Response Lab, R&D | Henry Lee et al |
| EPI_ISL_677133 | Wadsworth Center, New York State Department.of Health | Wadsworth Center, New York State Department.of Health | Kirsten St. George et al |
| EPI_ISL_4916023 | Aegis Sciences Corporation | Centers for Disease Control and Prevention Division of Viral Diseases, Pathogen Discovery | Dakota Howard et al |
| EPI_ISL_3740078 | Quest Diagnostics Incorporated | Centers for Disease Control and Prevention Division of Viral Diseases, Pathogen Discovery | Dakota Howard et al |
| EPI_ISL_2715662 | Pandemic Response Lab - NYC | Pandemic Response Lab, R&D | Henry Lee et al |
| EPI_ISL_6667528 | UW Virology Lab | UW Virology Lab | Pavitra Roychoudhury et al |
| EPI_ISL_4514257 | Pandemic Response Lab - NYC | Pandemic Response Lab, R&D | Henry Lee et al |
| EPI_ISL_1471645 | Pandemic Response Lab - NYC | Pandemic Response Lab, R&D | Henry Lee et al |
| EPI_ISL_1543842 | Pandemic Response Lab - NYC | Pandemic Response Lab, R&D | Henry Lee et al |
| EPI_ISL_1098234 | Pandemic Response Lab - NYC | Pandemic Response Lab, R&D | Henry Lee et al |
| EPI_ISL_1228515 | WESTCHESTER MEDICAL CENTER | Wadsworth Center, New York State Department of Health | Kirsten St. George et al |
| EPI_ISL_4534712 | Fulgent Genetics | Centers for Disease Control and Prevention Division of Viral Diseases, Pathogen Discovery | Dakota Howard et al |
| EPI_ISL_1398073 | NORTHWELL HEALTH LABORATORIES | Wadsworth Center, New York State Department of Health | Kirsten St. George et al |

| | | | |
|---|---|---|---|
| EPI_ISL_1098373 | Pandemic Response Lab - NYC | Pandemic Response Lab, R&D | Henry Lee et al |
| EPI_ISL_2462276 | OHSU Lab Services Molecular Microbiology Lab | Oregon SARS-CoV-2 Genome Sequencing Center | Brendan L. O'Connell et al |
| EPI_ISL_2632887 | Pandemic Response Lab - NYC | Pandemic Response Lab, R&D | Henry Lee et al |
| EPI_ISL_1755466 | NYC Pandemic Response Lab | Wadsworth Center, New York State Department of Health | Kirsten St. George et al |
| EPI_ISL_5547433 | Aegis Sciences Corporation | Centers for Disease Control and Prevention Division of Viral Diseases, Pathogen Discovery | Dakota Howard et al |
| EPI_ISL_1755446 | NYC Pandemic Response Lab | Wadsworth Center, New York State Department of Health | Kirsten St. George et al |
| EPI_ISL_6067776 | Aegis Sciences Corporation | Centers for Disease Control and Prevention Division of Viral Diseases, Pathogen Discovery | Dakota Howard et al |
| EPI_ISL_1338647 | Laboratory Corporation of America | Centers for Disease Control and Prevention Division of Viral Diseases, Pathogen Discovery | Peter W. Cook et al |
| EPI_ISL_7129869 | Pandemic Response Lab - NYC | Pandemic Response Lab, R&D | Henry Lee et al |
| EPI_ISL_1261428 | US Air Force School of Aerospace Medicine | US Air Force School of Aerospace Medicine | Anthony Fries et al |
| EPI_ISL_2100793 | The Ohio State University Applied Microbiology Services Laboratory | The Ohio State University Applied Microbiology Services Laboratory | Seth A. Faith PhD et al |
| EPI_ISL_884075 | ALBANY MEDICAL CENTER HOSPITAL CLINICAL LABORATORIES | Wadsworth Center, New York State Department of Health | Kirsten St. George et al |
| EPI_ISL_3942978 | Pandemic Response Lab - NYC | Pandemic Response Lab, R&D | Henry Lee et al |
| EPI_ISL_1634255 | Pandemic Response Lab - NYC | Pandemic Response Lab, R&D | Henry Lee et al |
| EPI_ISL_7166736 | Quadrant Laboratories, LLC | SUNY Upstate Medical University | Frank Middleton et al |
| EPI_ISL_1026851 | Laboratory Corporation of America | Respiratory Viruses Branch, Division of Viral Diseases, Centers for Disease Control and Prevention | Peter W. Cook et al |
| EPI_ISL_1907769 | Helix/Illumina | Centers for Disease Control and Prevention Division of Viral Diseases, Pathogen Discovery | Dakota Howard et al |
| EPI_ISL_4158096 | Laboratory Corporation of America | Centers for Disease Control and Prevention Division of Viral Diseases, Pathogen Discovery | Dakota Howard et al |
| EPI_ISL_6661947 | Molecular Biology Laboratory, Westchester Medical Center | Genomics Core Laboratory, New York Medical College | Weihua Huang et al |
| EPI_ISL_4194884 | CAREMOUNT MEDICAL | Wadsworth Center, New York State Department of Health | Kirsten St. George et al |
| EPI_ISL_734430 | Wadsworth Center, New York State Department.of Health | Wadsworth Center, New York State Department.of Health | Kirsten St. George et al |
| EPI_ISL_4378312 | Laboratory Corporation of America | Centers for Disease Control and Prevention Division of Viral Diseases, Pathogen Discovery | Dakota Howard et al |
| EPI_ISL_5517929 | Aegis Sciences Corporation | Centers for Disease Control and Prevention Division of Viral Diseases, Pathogen Discovery | Dakota Howard et al |
| EPI_ISL_1306597 | Pandemic Response Lab - NYC | Pandemic Response Lab, R&D | Henry Lee et al |

| | | | |
|---|---|---|---|
| EPI_ISL_667511 | OHSU Lab Services Molecular Microbiology Lab | Oregon SARS-CoV-2 Genome Sequencing Center | Brendan L. O'Connell et al |
| EPI_ISL_6644276 | Aegis Sciences Corporation | Centers for Disease Control and Prevention Division of Viral Diseases, Pathogen Discovery | Dakota Howard et al |
| EPI_ISL_2985762 | Gravity Diagnostics, LLC | Gravity Diagnostics, LLC | Gravity Diagnostics et al |
| EPI_ISL_1515177 | Laboratory Corporation of America | Centers for Disease Control and Prevention Division of Viral Diseases, Pathogen Discovery | Dakota Howard et al |
| EPI_ISL_4102587 | Kaiser Permanente NW Reginal Lab | OHSU MM Lab | Yun Wu et al |
| EPI_ISL_4174431 | Erie County Public Health (ECPHL) | University at Buffalo Genomics and Bioinformatics Core | Jonathan Bard et al |
| EPI_ISL_5021985 | Kaiser Permanente NW Reginal Lab | OHSU MM Lab | Yun Wu et al |
| EPI_ISL_4513944 | Pandemic Response Lab - NYC | Pandemic Response Lab, R&D | Henry Lee et al |
| EPI_ISL_1541751 | OHSU Lab Services Molecular Microbiology Lab | Oregon SARS-CoV-2 Genome Sequencing Center | Brendan L. O'Connell et al |
| EPI_ISL_2098548 | URMC LABS | Wadsworth Center, New York State Department of Health | Kirsten St. George et al |
| EPI_ISL_1227508 | WESTCHESTER MEDICAL CENTER | Wadsworth Center, New York State Department of Health | Kirsten St. George et al |
| EPI_ISL_3394709 | Pandemic Response Lab - NYC | Pandemic Response Lab, R&D | Henry Lee et al |
| EPI_ISL_1624410 | URMC LABS | Wadsworth Center, New York State Department of Health | Kirsten St. George et al |
| EPI_ISL_4165330 | Laboratory Corporation of America | Centers for Disease Control and Prevention Division of Viral Diseases, Pathogen Discovery | Dakota Howard et al |
| EPI_ISL_6785519 | Aegis Sciences Corporation | Centers for Disease Control and Prevention Division of Viral Diseases, Pathogen Discovery | Dakota Howard et al |
| EPI_ISL_1465821 | NYC Pandemic Response Lab | Wadsworth Center, New York State Department of Health | Kirsten St. George et al |
| EPI_ISL_4158099 | Laboratory Corporation of America | Centers for Disease Control and Prevention Division of Viral Diseases, Pathogen Discovery | Dakota Howard et al |
| EPI_ISL_3065725 | OHSU-MM Lab | OHSU-MM Lab | Rebecca O'Gara et al |
| EPI_ISL_4514185 | Pandemic Response Lab - NYC | Pandemic Response Lab, R&D | Henry Lee et al |
| EPI_ISL_6419578 | Aegis Sciences Corporation | Centers for Disease Control and Prevention Division of Viral Diseases, Pathogen Discovery | Dakota Howard et al |
| EPI_ISL_1339441 | Laboratory Corporation of America | Centers for Disease Control and Prevention Division of Viral Diseases, Pathogen Discovery | Peter W. Cook et al |
| EPI_ISL_4636953 | Kaiser Permanente NW Reginal Lab | OHSU MM Lab | Yun Wu et al |
| EPI_ISL_2106128 | Microbiology Division, SC DHEC | Microbiology Division, SC DHEC | Flores et al |
| EPI_ISL_6053517 | Aegis Sciences Corporation | Centers for Disease Control and Prevention Division of Viral Diseases, Pathogen Discovery | Dakota Howard et al |

| | | | |
|---|---|---|---|
| EPI_ISL_6527879 | Aegis Sciences Corporation | Centers for Disease Control and Prevention Division of Viral Diseases, Pathogen Discovery | Dakota Howard et al |
| EPI_ISL_5238307 | Fulgent Genetics | Centers for Disease Control and Prevention Division of Viral Diseases, Pathogen Discovery | Dakota Howard et al |
| EPI_ISL_3869356 | South Dakota Department of Health | University of Minnesota Genomics Center | Daryl M. Gohl et al |
| EPI_ISL_2481122 | Laboratory Corporation of America | Centers for Disease Control and Prevention Division of Viral Diseases, Pathogen Discovery | Dakota Howard et al |
| EPI_ISL_2398890 | Laboratory Corporation of America | Centers for Disease Control and Prevention Division of Viral Diseases, Pathogen Discovery | Dakota Howard et al |
| EPI_ISL_1588436 | Infinity Biologix | Centers for Disease Control and Prevention Division of Viral Diseases, Pathogen Discovery | Dakota Howard et al |
| EPI_ISL_1514662 | Laboratory Corporation of America | Centers for Disease Control and Prevention Division of Viral Diseases, Pathogen Discovery | Dakota Howard et al |
| EPI_ISL_5094098 | Quest Diagnostics Incorporated | Centers for Disease Control and Prevention Division of Viral Diseases, Pathogen Discovery | Dakota Howard et al |
| EPI_ISL_4395335 | UW Virology Lab | UW Virology Lab | Pavitra Roychoudhury et al |
| EPI_ISL_4914029 | Aegis Sciences Corporation | Centers for Disease Control and Prevention Division of Viral Diseases, Pathogen Discovery | Dakota Howard et al |
| EPI_ISL_1576611 | Helix/Illumina | Centers for Disease Control and Prevention Division of Viral Diseases, Pathogen Discovery | Dakota Howard et al |
| EPI_ISL_4392666 | UW Virology Lab | UW Virology Lab | Pavitra Roychoudhury et al |
| EPI_ISL_2725574 | Rhode Island State Health Laboratory | Rhode Island State Health Laboratory | Ewa King et al |
| EPI_ISL_6063200 | Aegis Sciences Corporation | Centers for Disease Control and Prevention Division of Viral Diseases, Pathogen Discovery | Dakota Howard et al |
| EPI_ISL_4485257 | Laboratory Corporation of America | Centers for Disease Control and Prevention Division of Viral Diseases, Pathogen Discovery | Dakota Howard et al |
| EPI_ISL_4057274 | Aegis Sciences Corporation | Centers for Disease Control and Prevention Division of Viral Diseases, Pathogen Discovery | Dakota Howard et al |
| EPI_ISL_1796934 | Helix/Illumina | Centers for Disease Control and Prevention Division of Viral Diseases, Pathogen Discovery | Dakota Howard et al |
| EPI_ISL_2320641 | Helix/Illumina | Centers for Disease Control and Prevention Division of Viral Diseases, Pathogen Discovery | Dakota Howard et al |
| EPI_ISL_1501765 | Oregon State University TRACE Project | OSU Center for Genome Research and Biocomputing | Oregon State University TRACE Project et al |

| | | | |
|---|---|---|---|
| EPI_ISL_1581075 | Helix/Illumina | Centers for Disease Control and Prevention Division of Viral Diseases, Pathogen Discovery | Dakota Howard et al |
| EPI_ISL_2422463 | TN Division of Laboratory Services | Genomics and Discovery, Respiratory Viruses Branch, Division of Viral Diseases, Centers for Disease Control and Prevention | Anna Kelleher et al |
| EPI_ISL_2184251 | Laboratory Corporation of America | Centers for Disease Control and Prevention Division of Viral Diseases, Pathogen Discovery | Dakota Howard et al |
| EPI_ISL_1320803 | Laboratory Corporation of America | Centers for Disease Control and Prevention Division of Viral Diseases, Pathogen Discovery | Peter W. Cook et al |
| EPI_ISL_1320974 | Laboratory Corporation of America | Centers for Disease Control and Prevention Division of Viral Diseases, Pathogen Discovery | Peter W. Cook et al |
| EPI_ISL_876593 | TN Division of Laboratory Services | Pathogen Discovery, Respiratory Viruses Branch, Division of Viral Diseases, Centers for Disease Control and Prevention | Ying Tao et al |
| EPI_ISL_1736397 | Aegis Sciences Corporation | Centers for Disease Control and Prevention Division of Viral Diseases, Pathogen Discovery | Dakota Howard et al |
| EPI_ISL_1924441 | Quest Diagnostics Incorporated | Centers for Disease Control and Prevention Division of Viral Diseases, Pathogen Discovery | Dakota Howard et al |
| EPI_ISL_6171031 | Aegis Sciences Corporation | Centers for Disease Control and Prevention Division of Viral Diseases, Pathogen Discovery | Dakota Howard et al |
| EPI_ISL_4484251 | Laboratory Corporation of America | Centers for Disease Control and Prevention Division of Viral Diseases, Pathogen Discovery | Dakota Howard et al |
| EPI_ISL_4386358 | Aegis Sciences Corporation | Centers for Disease Control and Prevention Division of Viral Diseases, Pathogen Discovery | Dakota Howard et al |
| EPI_ISL_5859973 | Aegis Sciences Corporation | Centers for Disease Control and Prevention Division of Viral Diseases, Pathogen Discovery | Dakota Howard et al |
| EPI_ISL_2147241 | Aegis Sciences Corporation | Centers for Disease Control and Prevention Division of Viral Diseases, Pathogen Discovery | Dakota Howard et al |
| EPI_ISL_1667870 | Aegis Sciences Corporation | Centers for Disease Control and Prevention Division of Viral Diseases, Pathogen Discovery | Dakota Howard et al |
| EPI_ISL_876213 | Massachusetts State Public Health Laboratory | Massachusetts State Public Health Laboratory | Andrew Lang et al |
| EPI_ISL_4996424 | Aegis Sciences Corporation | Centers for Disease Control and Prevention Division of Viral Diseases, Pathogen Discovery | Dakota Howard et al |
| EPI_ISL_5372667 | Aegis Sciences Corporation | Centers for Disease Control and Prevention Division of Viral Diseases, Pathogen Discovery | Dakota Howard et al |

| | | | |
|---|---|---|---|
| EPI_ISL_2427242 | South Dakota Public Health Laboratory | South Dakota Public Health Laboratory | Jacob Garfin et al |
| EPI_ISL_3451840 | Pennsylvania Department of Health Bureau of Laboratories | Pennsylvania Department of Health Bureau of Laboratories | Dongxiang Xia et al |
| EPI_ISL_1512316 | Helix/Illumina | Centers for Disease Control and Prevention Division of Viral Diseases, Pathogen Discovery | Dakota Howard et al |
| EPI_ISL_4398846 | Aegis Sciences Corporation | Centers for Disease Control and Prevention Division of Viral Diseases, Pathogen Discovery | Dakota Howard et al |
| EPI_ISL_6536957 | Aegis Sciences Corporation | Centers for Disease Control and Prevention Division of Viral Diseases, Pathogen Discovery | Dakota Howard et al |
| EPI_ISL_2716504 | UW Virology Lab | UW Virology Lab | Pavitra Roychoudhury et al |
| EPI_ISL_5571492 | Laboratory Corporation of America | Centers for Disease Control and Prevention Division of Viral Diseases, Pathogen Discovery | Dakota Howard et al |
| EPI_ISL_1575819 | Helix/Illumina | Centers for Disease Control and Prevention Division of Viral Diseases, Pathogen Discovery | Dakota Howard et al |
| EPI_ISL_6401173 | Infinity Biologix | Centers for Disease Control and Prevention Division of Viral Diseases, Pathogen Discovery | Dakota Howard et al |
| EPI_ISL_1702020 | Aegis Sciences Corporation | Centers for Disease Control and Prevention Division of Viral Diseases, Pathogen Discovery | Dakota Howard et al |
| EPI_ISL_5583484 | Aegis Sciences Corporation | Centers for Disease Control and Prevention Division of Viral Diseases, Pathogen Discovery | Dakota Howard et al |
| EPI_ISL_6405882 | Helix | Centers for Disease Control and Prevention Division of Viral Diseases, Pathogen Discovery | Dakota Howard et al |
| EPI_ISL_4384833 | Aegis Sciences Corporation | Centers for Disease Control and Prevention Division of Viral Diseases, Pathogen Discovery | Dakota Howard et al |
| EPI_ISL_6310777 | Mako Medical | Centers for Disease Control and Prevention Division of Viral Diseases, Pathogen Discovery | Dakota Howard et al |
| EPI_ISL_2320082 | Helix/Illumina | Centers for Disease Control and Prevention Division of Viral Diseases, Pathogen Discovery | Dakota Howard et al |
| EPI_ISL_6405216 | South Dakota Department of Health | University of Minnesota Genomics Center | Daryl M. Gohl et al |
| EPI_ISL_2016497 | Rhode Island Department of Health | Infectious Disease Program, Broad Institute of Harvard and MIT | Siddle et al |
| EPI_ISL_1683690 | Laboratory Corporation of America | Centers for Disease Control and Prevention Division of Viral Diseases, Pathogen Discovery | Dakota Howard et al |
| EPI_ISL_1513412 | Aegis Sciences Corporation | Centers for Disease Control and Prevention Division of Viral Diseases, Pathogen Discovery | Dakota Howard et al |

| | | | |
|---|---|---|---|
| EPI_ISL_2528544 | Aegis Sciences Corporation | Centers for Disease Control and Prevention Division of Viral Diseases, Pathogen Discovery | Dakota Howard et al |
| EPI_ISL_6650346 | Aegis Sciences Corporation | Centers for Disease Control and Prevention Division of Viral Diseases, Pathogen Discovery | Dakota Howard et al |
| EPI_ISL_2839011 | OHSU Lab Services Molecular Microbiology Lab | Ginkgo Bioworks Clinical Laboratory | VLOOKUP(C206 et al |
| EPI_ISL_3607431 | Laboratory Corporation of America | Centers for Disease Control and Prevention Division of Viral Diseases, Pathogen Discovery | Dakota Howard et al |
| EPI_ISL_4388351 | Aegis Sciences Corporation | Centers for Disease Control and Prevention Division of Viral Diseases, Pathogen Discovery | Dakota Howard et al |
| EPI_ISL_1482733 | MUSC Molecular Pathology Laboratory | MUSC Molecular Pathology Laboratory | Julie W. Hirschhorn et al |
| EPI_ISL_2016565 | Rhode Island Department of Health | Infectious Disease Program, Broad Institute of Harvard and MIT | Siddle et al |
| EPI_ISL_1493263 | Oregon State Public Health Laboratory | Oregon State Public Health Laboratory | Rafia Razzaque et al |
| EPI_ISL_5110497 | Aegis Sciences Corporation | Centers for Disease Control and Prevention Division of Viral Diseases, Pathogen Discovery | Dakota Howard et al |
| EPI_ISL_7156386 | Research Education in Disease Diagnosis and Intervention (REDDI) Lab, Clemson University | Research Education in Disease Diagnosis and Intervention (REDDI) Lab, Clemson University | Justin Napolitano et al |
| EPI_ISL_6491538 | Pennsylvania Department of Health Bureau of Laboratories | Pennsylvania Department of Health Bureau of Laboratories | Dongxiang Xia et al |
| EPI_ISL_5084117 | Mako Medical | Centers for Disease Control and Prevention Division of Viral Diseases, Pathogen Discovery | Dakota Howard et al |
| EPI_ISL_5860241 | Aegis Sciences Corporation | Centers for Disease Control and Prevention Division of Viral Diseases, Pathogen Discovery | Dakota Howard et al |
| EPI_ISL_6792886 | Laboratory Corporation of America | Centers for Disease Control and Prevention Division of Viral Diseases, Pathogen Discovery | Dakota Howard et al |
| EPI_ISL_2000549 | Aegis Sciences Corporation | Centers for Disease Control and Prevention Division of Viral Diseases, Pathogen Discovery | Dakota Howard et al |
| EPI_ISL_1972271 | Baylor Scott & White-Temple | Baylor Scott & White-Temple | Ari Rao et al |
| EPI_ISL_1623708 | Oregon State Public Health Laboratory | Oregon State Public Health Laboratory | Rafia Razzaque et al |
| EPI_ISL_4391348 | Aegis Sciences Corporation | Centers for Disease Control and Prevention Division of Viral Diseases, Pathogen Discovery | Dakota Howard et al |
| EPI_ISL_2242005 | Aegis Sciences Corporation | Centers for Disease Control and Prevention Division of Viral Diseases, Pathogen Discovery | Dakota Howard et al |

| | | | |
|---|---|---|---|
| EPI_ISL_4817139 | Infinity Biologix | Centers for Disease Control and Prevention Division of Viral Diseases, Pathogen Discovery | Dakota Howard et al |
| EPI_ISL_3491579 | RI State Health Laboratories | Centers for Disease Control and Prevention Division of Viral Diseases, Pathogen Discovery | Mili Sheth et al |
| EPI_ISL_1511539 | Hospital of the University of Pennsylvania Molecular Pathology Lab | Bushman Lab - University of Pennsylvania | John Everett et al |
| EPI_ISL_5586912 | Fulgent Genetics | Centers for Disease Control and Prevention Division of Viral Diseases, Pathogen Discovery | Dakota Howard et al |
| EPI_ISL_1609400 | Laboratory Corporation of America | Centers for Disease Control and Prevention Division of Viral Diseases, Pathogen Discovery | Dakota Howard et al |
| EPI_ISL_3769494 | Oregon State Public Health Laboratory | Oregon State Public Health Laboratory | Rafia Razzaque et al |
| EPI_ISL_3841432 | Oregon State Public Health Laboratory | Oregon State Public Health Laboratory | Rafia Razzaque et al |
| EPI_ISL_2242003 | Aegis Sciences Corporation | Centers for Disease Control and Prevention Division of Viral Diseases, Pathogen Discovery | Dakota Howard et al |
| EPI_ISL_2660387 | Infectious Disease Program, Broad Institute of Harvard and MIT | Infectious Disease Program, Broad Institute of Harvard and MIT | Lemieux et al |
| EPI_ISL_3801366 | Rhode Island State Health Laboratory | Rhode Island State Health Laboratory | Ewa King et al |
| EPI_ISL_2010439 | Helix/Illumina | Centers for Disease Control and Prevention Division of Viral Diseases, Pathogen Discovery | Dakota Howard et al |
| EPI_ISL_2086697 | Oregon State Public Health Laboratory | Oregon State Public Health Laboratory | Rafia Razzaque et al |
| EPI_ISL_3819289 | Aegis Sciences Corporation | Centers for Disease Control and Prevention Division of Viral Diseases, Pathogen Discovery | Dakota Howard et al |
| EPI_ISL_7156143 | Broad Institute Clinical Research Sequencing Platform | Infectious Disease Program, Broad Institute of Harvard and MIT | Siddle et al |
| EPI_ISL_2146383 | Aegis Sciences Corporation | Centers for Disease Control and Prevention Division of Viral Diseases, Pathogen Discovery | Dakota Howard et al |
| EPI_ISL_5236678 | Aegis Sciences Corporation | Centers for Disease Control and Prevention Division of Viral Diseases, Pathogen Discovery | Dakota Howard et al |
| EPI_ISL_6624369 | Fulgent Genetics | Centers for Disease Control and Prevention Division of Viral Diseases, Pathogen Discovery | Dakota Howard et al |
| EPI_ISL_1238583 | Houston Methodist Hospital | Houston Methodist Hospital | S. Wesley Long et al |
| EPI_ISL_3855958 | Aegis Sciences Corporation | Centers for Disease Control and Prevention Division of Viral Diseases, Pathogen Discovery | Dakota Howard et al |
| EPI_ISL_2616762 | Houston Health Dept. | Houston Health Dept. | Ryker Penn et al |
| EPI_ISL_547394 | Houston Methodist Hospital | Houston Methodist Hospital | S. Wesley Long et al |

| | | | |
|---|---|---|---|
| EPI_ISL_3608186 | Fulgent Genetics | Centers for Disease Control and Prevention Division of Viral Diseases, Pathogen Discovery | Dakota Howard et al |
| EPI_ISL_785641 | Houston Methodist Hospital | Houston Methodist Hospital | S. Wesley Long et al |
| EPI_ISL_1038563 | Laboratory Corporation of America | Respiratory Viruses Branch, Division of Viral Diseases, Centers for Disease Control and Prevention | Peter W. Cook et al |
| EPI_ISL_1201363 | Texas Department of State Health Services | Texas Department of State Health Services | Rashmi Tuladhar et al |
| EPI_ISL_1077198 | Houston Methodist Hospital | Houston Methodist Hospital | S. Wesley Long et al |
| EPI_ISL_784492 | Houston Methodist Hospital | Houston Methodist Hospital | S. Wesley Long et al |
| EPI_ISL_3822464 | Aegis Sciences Corporation | Centers for Disease Control and Prevention Division of Viral Diseases, Pathogen Discovery | Dakota Howard et al |
| EPI_ISL_788611 | Houston Methodist Hospital | Houston Methodist Hospital | S. Wesley Long et al |
| EPI_ISL_6163783 | TXDSHS | TXDSHS | Rashmi Tuladhar et al |
| EPI_ISL_2368300 | Aegis Sciences Corporation | Centers for Disease Control and Prevention Division of Viral Diseases, Pathogen Discovery | Dakota Howard et al |
| EPI_ISL_5239230 | TXDSHS | TXDSHS | Rashmi Tuladhar et al |
| EPI_ISL_546875 | Houston Methodist Hospital | Houston Methodist Hospital | S. Wesley Long et al |
| EPI_ISL_1078099 | Houston Methodist Hospital | Houston Methodist Hospital | S. Wesley Long et al |
| EPI_ISL_5773170 | Aegis Sciences Corporation | Centers for Disease Control and Prevention Division of Viral Diseases, Pathogen Discovery | Dakota Howard et al |
| EPI_ISL_2941835 | Curative Labs | Curative Labs | Mikhail Hanewich-Hollatz et al |
| EPI_ISL_1581204 | Helix/Illumina | Centers for Disease Control and Prevention Division of Viral Diseases, Pathogen Discovery | Dakota Howard et al |
| EPI_ISL_3344230 | Fulgent Genetics | Centers for Disease Control and Prevention Division of Viral Diseases, Pathogen Discovery | Dakota Howard et al |
| EPI_ISL_2205833 | Houston Methodist Hospital | Houston Methodist Hospital | Randall J. Olsen et al |
| EPI_ISL_6629055 | Aegis Sciences Corporation | Centers for Disease Control and Prevention Division of Viral Diseases, Pathogen Discovery | Dakota Howard et al |
| EPI_ISL_4367585 | Fulgent Genetics | Centers for Disease Control and Prevention Division of Viral Diseases, Pathogen Discovery | Dakota Howard et al |
| EPI_ISL_790042 | Houston Methodist Hospital | Houston Methodist Hospital | S. Wesley Long et al |
| EPI_ISL_1684599 | Laboratory Corporation of America | Centers for Disease Control and Prevention Division of Viral Diseases, Pathogen Discovery | Dakota Howard et al |
| EPI_ISL_543103 | Houston Methodist Hospital | Houston Methodist Hospital | S. Wesley Long et al |
| EPI_ISL_788538 | Houston Methodist Hospital | Houston Methodist Hospital | S. Wesley Long et al |
| EPI_ISL_1238257 | Houston Methodist Hospital | Houston Methodist Hospital | S. Wesley Long et al |
| EPI_ISL_1760571 | TXDSHS | TXDSHS | Rashmi Tuladhar et al |
| EPI_ISL_5659016 | Helix | Centers for Disease Control and Prevention Division of Viral Diseases, Pathogen Discovery | Dakota Howard et al |
| EPI_ISL_3455711 | TXDSHS | TXDSHS | Rashmi Tuladhar et al |
| EPI_ISL_1237527 | Houston Methodist Hospital | Houston Methodist Hospital | S. Wesley Long et al |

| | | | |
|---|---|---|---|
| EPI_ISL_5240669 | Helix | Centers for Disease Control and Prevention Division of Viral Diseases, Pathogen Discovery | Dakota Howard et al |
| EPI_ISL_1238660 | Houston Methodist Hospital | Houston Methodist Hospital | S. Wesley Long et al |
| EPI_ISL_5092505 | Fulgent Genetics | Centers for Disease Control and Prevention Division of Viral Diseases, Pathogen Discovery | Dakota Howard et al |
| EPI_ISL_2190291 | Houston Methodist Hospital | Houston Methodist Hospital | Randall J. Olsen et al |
| EPI_ISL_2190468 | Houston Methodist Hospital | Houston Methodist Hospital | Randall J. Olsen et al |
| EPI_ISL_4908794 | Aegis Sciences Corporation | Centers for Disease Control and Prevention Division of Viral Diseases, Pathogen Discovery | Dakota Howard et al |
| EPI_ISL_785467 | Houston Methodist Hospital | Houston Methodist Hospital | S. Wesley Long et al |
| EPI_ISL_1303698 | Houston Methodist Hospital | Houston Methodist Hospital | S. Wesley Long et al |
| EPI_ISL_543380 | Houston Methodist Hospital | Houston Methodist Hospital | S. Wesley Long et al |
| EPI_ISL_4240743 | Laboratory Corporation of America | Centers for Disease Control and Prevention Division of Viral Diseases, Pathogen Discovery | Dakota Howard et al |
| EPI_ISL_5875639 | Laboratory Corporation of America | Centers for Disease Control and Prevention Division of Viral Diseases, Pathogen Discovery | Dakota Howard et al |
| EPI_ISL_6378728 | Aegis Sciences Corporation | Centers for Disease Control and Prevention Division of Viral Diseases, Pathogen Discovery | Dakota Howard et al |
| EPI_ISL_2203469 | Houston Methodist Hospital | Houston Methodist Hospital | Randall J. Olsen et al |
| EPI_ISL_5109483 | Aegis Sciences Corporation | Centers for Disease Control and Prevention Division of Viral Diseases, Pathogen Discovery | Dakota Howard et al |
| EPI_ISL_3064635 | Curative Labs | Curative Labs | Mikhail Hanewich-Hollatz et al |
| EPI_ISL_3846211 | Aegis Sciences Corporation | Centers for Disease Control and Prevention Division of Viral Diseases, Pathogen Discovery | Dakota Howard et al |
| EPI_ISL_1444288 | TXDSHS | TXDSHS | Rashmi Tuladhar et al |
| EPI_ISL_5875007 | Laboratory Corporation of America | Centers for Disease Control and Prevention Division of Viral Diseases, Pathogen Discovery | Dakota Howard et al |
| EPI_ISL_4391477 | Aegis Sciences Corporation | Centers for Disease Control and Prevention Division of Viral Diseases, Pathogen Discovery | Dakota Howard et al |
| EPI_ISL_6420617 | Aegis Sciences Corporation | Centers for Disease Control and Prevention Division of Viral Diseases, Pathogen Discovery | Dakota Howard et al |
| EPI_ISL_3815922 | Aegis Sciences Corporation | Centers for Disease Control and Prevention Division of Viral Diseases, Pathogen Discovery | Dakota Howard et al |
| EPI_ISL_5113191 | Aegis Sciences Corporation | Centers for Disease Control and Prevention Division of Viral Diseases, Pathogen Discovery | Dakota Howard et al |
| EPI_ISL_3821172 | Aegis Sciences Corporation | Centers for Disease Control and Prevention Division of Viral Diseases, Pathogen Discovery | Dakota Howard et al |
| EPI_ISL_1076085 | Houston Methodist Hospital | Houston Methodist Hospital | S. Wesley Long et al |

| | | | |
|---|---|---|---|
| EPI_ISL_1078333 | Houston Methodist Hospital | Houston Methodist Hospital | S. Wesley Long et al |
| EPI_ISL_4358918 | Fulgent Genetics | Centers for Disease Control and Prevention Division of Viral Diseases, Pathogen Discovery | Dakota Howard et al |
| EPI_ISL_3661755 | Fulgent Genetics | Centers for Disease Control and Prevention Division of Viral Diseases, Pathogen Discovery | Dakota Howard et al |
| EPI_ISL_2873941 | Quest Diagnostics Incorporated | Centers for Disease Control and Prevention Division of Viral Diseases, Pathogen Discovery | Dakota Howard et al |
| EPI_ISL_4183610 | Fulgent Genetics | Centers for Disease Control and Prevention Division of Viral Diseases, Pathogen Discovery | Dakota Howard et al |
| EPI_ISL_2340594 | GHRC | TxGen | Benjamin W. Neuman et al |
| EPI_ISL_4911620 | Aegis Sciences Corporation | Centers for Disease Control and Prevention Division of Viral Diseases, Pathogen Discovery | Dakota Howard et al |
| EPI_ISL_1079587 | Houston Methodist Hospital | Houston Methodist Hospital | S. Wesley Long et al |
| EPI_ISL_3686275 | Laboratory Corporation of America | Centers for Disease Control and Prevention Division of Viral Diseases, Pathogen Discovery | Dakota Howard et al |
| EPI_ISL_1391520 | Quest Diagnostics Incorporated | Centers for Disease Control and Prevention Division of Viral Diseases, Pathogen Discovery | Peter W. Cook et al |
| EPI_ISL_1687576 | Aegis Sciences Corporation | Centers for Disease Control and Prevention Division of Viral Diseases, Pathogen Discovery | Dakota Howard et al |
| EPI_ISL_2134519 | Fulgent Genetics | Centers for Disease Control and Prevention Division of Viral Diseases, Pathogen Discovery | Dakota Howard et al |
| EPI_ISL_1203072 | Houston Methodist Hospital | Houston Methodist Hospital | S. Wesley Long et al |
| EPI_ISL_4023524 | Fulgent Genetics | Centers for Disease Control and Prevention Division of Viral Diseases, Pathogen Discovery | Dakota Howard et al |
| EPI_ISL_4023078 | Fulgent Genetics | Centers for Disease Control and Prevention Division of Viral Diseases, Pathogen Discovery | Dakota Howard et al |
| EPI_ISL_1392728 | TXDSHS | TXDSHS | Rashmi Tuladhar et al |
| EPI_ISL_1203533 | Houston Methodist Hospital | Houston Methodist Hospital | S. Wesley Long et al |
| EPI_ISL_2206386 | Houston Methodist Hospital | Houston Methodist Hospital | Randall J. Olsen et al |
| EPI_ISL_7118422 | Fulgent Genetics | Centers for Disease Control and Prevention Division of Viral Diseases, Pathogen Discovery | Dakota Howard et al |
| EPI_ISL_2215401 | Houston Methodist Hospital | Houston Methodist Hospital | Randall J. Olsen et al |
| EPI_ISL_2785411 | Aegis Sciences Corporation | Centers for Disease Control and Prevention Division of Viral Diseases, Pathogen Discovery | Dakota Howard et al |
| EPI_ISL_5553736 | Aegis Sciences Corporation | Centers for Disease Control and Prevention Division of Viral Diseases, Pathogen Discovery | Dakota Howard et al |
| EPI_ISL_546670 | Houston Methodist Hospital | Houston Methodist Hospital | S. Wesley Long et al |

| | | | |
|---|---|---|---|
| EPI_ISL_4368044 | Quest Diagnostics Incorporated | Centers for Disease Control and Prevention Division of Viral Diseases, Pathogen Discovery | Dakota Howard et al |
| EPI_ISL_3518626 | Laboratory Corporation of America | Centers for Disease Control and Prevention Division of Viral Diseases, Pathogen Discovery | Dakota Howard et al |
| EPI_ISL_785191 | Houston Methodist Hospital | Houston Methodist Hospital | S. Wesley Long et al |
| EPI_ISL_1422375 | Quest Diagnostics Incorporated | Centers for Disease Control and Prevention Division of Viral Diseases, Pathogen Discovery | Peter W. Cook et al |
| EPI_ISL_3865827 | TXDSHS | TXDSHS | Rashmi Tuladhar et al |
| EPI_ISL_4138346 | Aegis Sciences Corporation | Centers for Disease Control and Prevention Division of Viral Diseases, Pathogen Discovery | Dakota Howard et al |
| EPI_ISL_3852888 | Aegis Sciences Corporation | Centers for Disease Control and Prevention Division of Viral Diseases, Pathogen Discovery | Dakota Howard et al |
| EPI_ISL_3113884 | Quest Diagnostics Incorporated | Centers for Disease Control and Prevention Division of Viral Diseases, Pathogen Discovery | Dakota Howard et al |
| EPI_ISL_6481112 | Helix | Centers for Disease Control and Prevention Division of Viral Diseases, Pathogen Discovery | Dakota Howard et al |
| EPI_ISL_4914007 | Aegis Sciences Corporation | Centers for Disease Control and Prevention Division of Viral Diseases, Pathogen Discovery | Dakota Howard et al |
| EPI_ISL_1305265 | Houston Methodist Hospital | Houston Methodist Hospital | S. Wesley Long et al |
| EPI_ISL_5520906 | Aegis Sciences Corporation | Centers for Disease Control and Prevention Division of Viral Diseases, Pathogen Discovery | Dakota Howard et al |
| EPI_ISL_1077757 | Houston Methodist Hospital | Houston Methodist Hospital | S. Wesley Long et al |
| EPI_ISL_1992932 | Fulgent Genetics | Centers for Disease Control and Prevention Division of Viral Diseases, Pathogen Discovery | Dakota Howard et al |
| EPI_ISL_3719795 | Fulgent Genetics | Centers for Disease Control and Prevention Division of Viral Diseases, Pathogen Discovery | Dakota Howard et al |
| EPI_ISL_4496360 | Aegis Sciences Corporation | Centers for Disease Control and Prevention Division of Viral Diseases, Pathogen Discovery | Dakota Howard et al |
| EPI_ISL_2220500 | Houston Methodist Hospital | Houston Methodist Hospital | Randall J. Olsen et al |
| EPI_ISL_2320115 | Helix/Illumina | Centers for Disease Control and Prevention Division of Viral Diseases, Pathogen Discovery | Dakota Howard et al |
| EPI_ISL_2398031 | Laboratory Corporation of America | Centers for Disease Control and Prevention Division of Viral Diseases, Pathogen Discovery | Dakota Howard et al |
| EPI_ISL_7136766 | Utah Public Health Laboratory | Utah Public Health Laboratory | Erin L. Young et al |
| EPI_ISL_5339969 | Houston Methodist Hospital | Houston Methodist Hospital | Randall J. Olsen et al |
| EPI_ISL_6372151 | Utah Public Health Laboratory | Utah Public Health Laboratory | Erin L. Young et al |
| EPI_ISL_5085842 | Houston Methodist Hospital | Houston Methodist Hospital | Randall J. Olsen et al |

| | | | |
|---|---|---|---|
| EPI_ISL_1552827 | Quest Diagnostics Incorporated | Centers for Disease Control and Prevention Division of Viral Diseases, Pathogen Discovery | Dakota Howard et al |
| EPI_ISL_5878183 | Laboratory Corporation of America | Centers for Disease Control and Prevention Division of Viral Diseases, Pathogen Discovery | Dakota Howard et al |
| EPI_ISL_2225318 | Houston Methodist Hospital | Houston Methodist Hospital | Randall J. Olsen et al |
| EPI_ISL_5110616 | Aegis Sciences Corporation | Centers for Disease Control and Prevention Division of Viral Diseases, Pathogen Discovery | Dakota Howard et al |
| EPI_ISL_2856865 | Utah Public Health Laboratory | Utah Public Health Laboratory | Olinto Linares-Perdomo et al |
| EPI_ISL_3268613 | Utah Public Health Laboratory | Utah Public Health Laboratory | Olinto Linares-Perdomo et al |
| EPI_ISL_3756021 | Utah Public Health Laboratory | Utah Public Health Laboratory | Erin L. Young et al |
| EPI_ISL_6063123 | Aegis Sciences Corporation | Centers for Disease Control and Prevention Division of Viral Diseases, Pathogen Discovery | Dakota Howard et al |
| EPI_ISL_5339052 | Houston Methodist Hospital | Houston Methodist Hospital | Randall J. Olsen et al |
| EPI_ISL_4177014 | Utah Public Health Laboratory | Utah Public Health Laboratory | Erin L. Young et al |
| EPI_ISL_5235222 | Laboratory Corporation of America | Centers for Disease Control and Prevention Division of Viral Diseases, Pathogen Discovery | Dakota Howard et al |
| EPI_ISL_4987126 | Aegis Sciences Corporation | Centers for Disease Control and Prevention Division of Viral Diseases, Pathogen Discovery | Dakota Howard et al |
| EPI_ISL_5375043 | Houston Methodist Hospital | Houston Methodist Hospital | Randall J. Olsen et al |
| EPI_ISL_5214044 | Laboratory Corporation of America | Centers for Disease Control and Prevention Division of Viral Diseases, Pathogen Discovery | Dakota Howard et al |
| EPI_ISL_487214 | Utah Public Health Laboratory | Utah Public Health Laboratory | Heidi Butz et al |
| EPI_ISL_1113003 | Baylor College of Medicine/ GCID | Baylor College of Medicine/ GCID | Vasanthi Avadhanula et al |
| EPI_ISL_450542 | Utah Public Health Laboratory | Utah Public Health Laboratory | Erin Young et al |
| EPI_ISL_1649180 | Aegis Sciences Corporation | Centers for Disease Control and Prevention Division of Viral Diseases, Pathogen Discovery | Dakota Howard et al |
| EPI_ISL_5373205 | Houston Methodist Hospital | Houston Methodist Hospital | Randall J. Olsen et al |
| EPI_ISL_5345273 | Houston Methodist Hospital | Houston Methodist Hospital | Randall J. Olsen et al |
| EPI_ISL_5373780 | Houston Methodist Hospital | Houston Methodist Hospital | Randall J. Olsen et al |
| EPI_ISL_2214509 | Houston Methodist Hospital | Houston Methodist Hospital | Randall J. Olsen et al |
| EPI_ISL_2456141 | Utah Public Health Laboratory | Utah Public Health Laboratory | Erin L. Young et al |
| EPI_ISL_2213083 | Houston Methodist Hospital | Houston Methodist Hospital | Randall J. Olsen et al |
| EPI_ISL_5088012 | Houston Methodist Hospital | Houston Methodist Hospital | Randall J. Olsen et al |
| EPI_ISL_5384868 | Houston Methodist Hospital | Houston Methodist Hospital | Randall J. Olsen et al |
| EPI_ISL_3755042 | Laboratory Corporation of America | Centers for Disease Control and Prevention Division of Viral Diseases, Pathogen Discovery | Dakota Howard et al |
| EPI_ISL_4467588 | Utah Public Health Laboratory | Utah Public Health Laboratory | Erin L. Young et al |
| EPI_ISL_4157819 | Laboratory Corporation of America | Centers for Disease Control and Prevention Division of Viral Diseases, Pathogen Discovery | Dakota Howard et al |
| EPI_ISL_2214494 | Houston Methodist Hospital | Houston Methodist Hospital | Randall J. Olsen et al |
| EPI_ISL_5346327 | Houston Methodist Hospital | Houston Methodist Hospital | Randall J. Olsen et al |

| | | | |
|---|---|---|---|
| EPI_ISL_2677944 | Utah Public Health Laboratory | Utah Public Health Laboratory | Erin L. Young et al |
| EPI_ISL_2679002 | Utah Public Health Laboratory | Utah Public Health Laboratory | Erin L. Young et al |
| EPI_ISL_5368807 | Houston Methodist Hospital | Houston Methodist Hospital | Randall J. Olsen et al |
| EPI_ISL_4003861 | Utah Public Health Laboratory | Utah Public Health Laboratory | Erin L. Young et al |
| EPI_ISL_418964 | Utah Public Health Laboratory | Utah Public Health Laboratory | Erin Young et al |
| EPI_ISL_5085090 | Houston Methodist Hospital | Houston Methodist Hospital | Randall J. Olsen et al |
| EPI_ISL_6312910 | Aegis Sciences Corporation | Centers for Disease Control and Prevention Division of Viral Diseases, Pathogen Discovery | Dakota Howard et al |
| EPI_ISL_2496443 | Utah Public Health Laboratory | Utah Public Health Laboratory | Erin L. Young et al |
| EPI_ISL_5382809 | Houston Methodist Hospital | Houston Methodist Hospital | Randall J. Olsen et al |
| EPI_ISL_5083375 | Houston Methodist Hospital | Houston Methodist Hospital | Randall J. Olsen et al |
| EPI_ISL_5074747 | Houston Methodist Hospital | Houston Methodist Hospital | Randall J. Olsen et al |
| EPI_ISL_1800893 | Laboratory Corporation of America | Centers for Disease Control and Prevention Division of Viral Diseases, Pathogen Discovery | Dakota Howard et al |
| EPI_ISL_3450212 | Utah Public Health Laboratory | Utah Public Health Laboratory | Erin L. Young et al |
| EPI_ISL_6158827 | Aegis Sciences Corporation | Centers for Disease Control and Prevention Division of Viral Diseases, Pathogen Discovery | Dakota Howard et al |
| EPI_ISL_5084751 | Houston Methodist Hospital | Houston Methodist Hospital | Randall J. Olsen et al |
| EPI_ISL_4204822 | Utah Public Health Laboratory | Utah Public Health Laboratory | Erin L. Young et al |
| EPI_ISL_7136523 | Utah Public Health Laboratory | Utah Public Health Laboratory | Erin L. Young et al |
| EPI_ISL_2676921 | Utah Public Health Laboratory | Utah Public Health Laboratory | Erin L. Young et al |
| EPI_ISL_2223995 | Houston Methodist Hospital | Houston Methodist Hospital | Randall J. Olsen et al |
| EPI_ISL_5112266 | Aegis Sciences Corporation | Centers for Disease Control and Prevention Division of Viral Diseases, Pathogen Discovery | Dakota Howard et al |
| EPI_ISL_575182 | Utah Public Health Laboratory | Utah Public Health Laboratory | Erin Young et al |
| EPI_ISL_5342739 | Houston Methodist Hospital | Houston Methodist Hospital | Randall J. Olsen et al |
| EPI_ISL_4485600 | Laboratory Corporation of America | Centers for Disease Control and Prevention Division of Viral Diseases, Pathogen Discovery | Dakota Howard et al |
| EPI_ISL_4486604 | Laboratory Corporation of America | Centers for Disease Control and Prevention Division of Viral Diseases, Pathogen Discovery | Dakota Howard et al |
| EPI_ISL_941915 | Virginia DCLS | Virginia DCLS | Virginia DCLS et al |
| EPI_ISL_5214075 | Laboratory Corporation of America | Centers for Disease Control and Prevention Division of Viral Diseases, Pathogen Discovery | Dakota Howard et al |
| EPI_ISL_3353976 | VA-Division of Consolidated Laboratory Services | Centers for Disease Control and Prevention Division of Viral Diseases, Pathogen Discovery | Mili Sheth et al |
| EPI_ISL_4165185 | Laboratory Corporation of America | Centers for Disease Control and Prevention Division of Viral Diseases, Pathogen Discovery | Dakota Howard et al |
| EPI_ISL_3144715 | Utah Public Health Laboratory | Utah Public Health Laboratory | Olinto Linares-Perdomo et al |
| EPI_ISL_2149968 | Aegis Sciences Corporation | Centers for Disease Control and Prevention Division of Viral Diseases, Pathogen Discovery | Dakota Howard et al |
| EPI_ISL_780098 | Houston Methodist Hospital | Houston Methodist Hospital | S. Wesley Long et al |
| EPI_ISL_4204845 | Utah Public Health Laboratory | Utah Public Health Laboratory | Erin L. Young et al |

| | | | |
|---|---|---|---|
| EPI_ISL_5339565 | Houston Methodist Hospital | Houston Methodist Hospital | Randall J. Olsen et al |
| EPI_ISL_5849596 | Utah Public Health Laboratory | Utah Public Health Laboratory | Erin L. Young et al |
| EPI_ISL_5373290 | Houston Methodist Hospital | Houston Methodist Hospital | Randall J. Olsen et al |
| EPI_ISL_5384155 | Houston Methodist Hospital | Houston Methodist Hospital | Randall J. Olsen et al |
| EPI_ISL_2224761 | Houston Methodist Hospital | Houston Methodist Hospital | Randall J. Olsen et al |
| EPI_ISL_2857198 | Utah Public Health Laboratory | Utah Public Health Laboratory | Olinto Linares-Perdomo et al |
| EPI_ISL_2292060 | Utah Public Health Laboratory | Utah Public Health Laboratory | Erin L. Young et al |
| EPI_ISL_5085082 | Houston Methodist Hospital | Houston Methodist Hospital | Randall J. Olsen et al |
| EPI_ISL_4177734 | Utah Public Health Laboratory | Utah Public Health Laboratory | Erin L. Young et al |
| EPI_ISL_5585959 | Quest Diagnostics Incorporated | Centers for Disease Control and Prevention Division of Viral Diseases, Pathogen Discovery | Dakota Howard et al |
| EPI_ISL_1183726 | Utah Public Health Laboratory | Utah Public Health Laboratory | Erin L. Young et al |
| EPI_ISL_3263188 | Utah Public Health Laboratory | Utah Public Health Laboratory | Erin L. Young et al |
| EPI_ISL_5078284 | Utah Public Health Laboratory | Utah Public Health Laboratory | Erin L. Young et al |
| EPI_ISL_5338953 | Houston Methodist Hospital | Houston Methodist Hospital | Randall J. Olsen et al |
| EPI_ISL_2857755 | Utah Public Health Laboratory | Utah Public Health Laboratory | Olinto Linares-Perdomo et al |
| EPI_ISL_5013931 | Utah Public Health Laboratory | Utah Public Health Laboratory | Erin L. Young et al |
| EPI_ISL_745965 | Ginkgo Bioworks Clinical Laboratory | Utah Public Health Laboratory | Erin L. Young et al |
| EPI_ISL_4819206 | Laboratory Corporation of America | Centers for Disease Control and Prevention Division of Viral Diseases, Pathogen Discovery | Dakota Howard et al |
| EPI_ISL_5382881 | Houston Methodist Hospital | Houston Methodist Hospital | Randall J. Olsen et al |
| EPI_ISL_2224789 | Houston Methodist Hospital | Houston Methodist Hospital | Randall J. Olsen et al |
| EPI_ISL_780322 | Houston Methodist Hospital | Houston Methodist Hospital | S. Wesley Long et al |
| EPI_ISL_5341486 | Houston Methodist Hospital | Houston Methodist Hospital | Randall J. Olsen et al |
| EPI_ISL_5380840 | Houston Methodist Hospital | Houston Methodist Hospital | Randall J. Olsen et al |
| EPI_ISL_5344786 | Houston Methodist Hospital | Houston Methodist Hospital | Randall J. Olsen et al |
| EPI_ISL_5381653 | Houston Methodist Hospital | Houston Methodist Hospital | Randall J. Olsen et al |
| EPI_ISL_2677031 | Utah Public Health Laboratory | Utah Public Health Laboratory | Erin L. Young et al |
| EPI_ISL_2291441 | Utah Public Health Laboratory | Utah Public Health Laboratory | Erin L. Young et al |
| EPI_ISL_5338630 | Houston Methodist Hospital | Houston Methodist Hospital | Randall J. Olsen et al |
| EPI_ISL_4818479 | Laboratory Corporation of America | Centers for Disease Control and Prevention Division of Viral Diseases, Pathogen Discovery | Dakota Howard et al |
| EPI_ISL_2291206 | Utah Public Health Laboratory | Utah Public Health Laboratory | Erin L. Young et al |
| EPI_ISL_5072675 | Houston Methodist Hospital | Houston Methodist Hospital | Randall J. Olsen et al |
| EPI_ISL_6511244 | Fulgent Genetics | Centers for Disease Control and Prevention Division of Viral Diseases, Pathogen Discovery | Dakota Howard et al |
| EPI_ISL_6904649 | Laboratory Corporation of America | Centers for Disease Control and Prevention Division of Viral Diseases, Pathogen Discovery | Dakota Howard et al |
| EPI_ISL_3049889 | State Testing Facility | Altius Institute for Biomedical Research | Daniel Bates et al |
| EPI_ISL_3149643 | UW Virology Lab | UW Virology Lab | Pavitra Roychoudhury et al |
| EPI_ISL_1445063 | Virginia Division of Consolidated Laboratory Services | Virginia Division of Consolidated Laboratory Services | Virginia DCLS et al |
| EPI_ISL_5001783 | UW Virology Lab | UW Virology Lab | Pavitra Roychoudhury et al |
| EPI_ISL_5415742 | UW Virology Lab | UW Virology Lab | Pavitra Roychoudhury et al |

| | | | |
|---|---|---|---|
| EPI_ISL_5889591 | Fulgent Genetics | Centers for Disease Control and Prevention Division of Viral Diseases, Pathogen Discovery | Dakota Howard et al |
| EPI_ISL_4000283 | Virginia Division of Consolidated Laboratory Services | Virginia Division of Consolidated Laboratory Services | Virginia Division of Consolidated Laboratory Services et al |
| EPI_ISL_5064747 | Virginia Division of Consolidated Laboratory Services | Virginia Division of Consolidated Laboratory Services | Virginia Division of Consolidated Laboratory Services et al |
| EPI_ISL_5458069 | Broad Institute Clinical Research Sequencing Platform | Infectious Disease Program, Broad Institute of Harvard and MIT | Siddle et al |
| EPI_ISL_4169745 | Virginia Division of Consolidated Laboratory Services | Virginia Division of Consolidated Laboratory Services | Virginia Division of Consolidated Laboratory Services et al |
| EPI_ISL_2544958 | UW Virology Lab | UW Virology Lab | Pavitra Roychoudhury et al |
| EPI_ISL_4411086 | Washington State Department of Health Public Health Laboratories | Washington State Department of Health Public Health Laboratories | Drew MacKellar et al |
| EPI_ISL_6079093 | Laboratory Corporation of America | Centers for Disease Control and Prevention Division of Viral Diseases, Pathogen Discovery | Dakota Howard et al |
| EPI_ISL_4036155 | UW Virology Lab | UW Virology Lab | Pavitra Roychoudhury et al |
| EPI_ISL_1608941 | Laboratory Corporation of America | Centers for Disease Control and Prevention Division of Viral Diseases, Pathogen Discovery | Dakota Howard et al |
| EPI_ISL_5533015 | UW Virology Lab | UW Virology Lab | Pavitra Roychoudhury et al |
| EPI_ISL_5170491 | Laboratory Corporation of America | Centers for Disease Control and Prevention Division of Viral Diseases, Pathogen Discovery | Dakota Howard et al |
| EPI_ISL_1785629 | UW Virology Lab | UW Virology Lab | Pavitra Roychoudhury et al |
| EPI_ISL_5530243 | UW Virology Lab | UW Virology Lab | Pavitra Roychoudhury et al |
| EPI_ISL_1448355 | UW Virology Lab | UW Virology Lab | Pavitra Roychoudhury et al |
| EPI_ISL_3346906 | Northwest Laboratory | Seattle Flu Study | Deborah A. Nickerson et al |
| EPI_ISL_5628423 | Washington State Department of Health Public Health Laboratories | Washington State Department of Health Public Health Laboratories | Drew MacKellar et al |
| EPI_ISL_4476615 | Laboratory Corporation of America | Centers for Disease Control and Prevention Division of Viral Diseases, Pathogen Discovery | Dakota Howard et al |
| EPI_ISL_2161862 | UW Virology Lab | UW Virology Lab | Pavitra Roychoudhury et al |
| EPI_ISL_4825786 | Quest Diagnostics Incorporated | Centers for Disease Control and Prevention Division of Viral Diseases, Pathogen Discovery | Dakota Howard et al |
| EPI_ISL_6259123 | Washington State Department of Health Public Health Laboratories | Washington State Department of Health Public Health Laboratories | Drew MacKellar et al |
| EPI_ISL_4455399 | Fulgent Genetics | Centers for Disease Control and Prevention Division of Viral Diseases, Pathogen Discovery | Dakota Howard et al |
| EPI_ISL_5869297 | UW Virology Lab | UW Virology Lab | Pavitra Roychoudhury et al |
| EPI_ISL_2161989 | UW Virology Lab | UW Virology Lab | Pavitra Roychoudhury et al |
| EPI_ISL_1849786 | UW Virology Lab | UW Virology Lab | Pavitra Roychoudhury et al |

| | | | |
|---|---|---|---|
| EPI_ISL_2201277 | Aegis Sciences Corporation | Centers for Disease Control and Prevention Division of Viral Diseases, Pathogen Discovery | Dakota Howard et al |
| EPI_ISL_2194131 | UW Virology Lab | UW Virology Lab | Pavitra Roychoudhury et al |
| EPI_ISL_6173973 | Virginia Division of Consolidated Laboratory Services | Virginia Commonwealth University | Gregory A Buck et al |
| EPI_ISL_3372818 | Northwest Laboratory | Seattle Flu Study | Deborah A. Nickerson et al |
| EPI_ISL_570520 | UW Virology Lab | UW Virology Lab | Pavitra Roychoudhury et al |
| EPI_ISL_5082686 | Fulgent Genetics | Centers for Disease Control and Prevention Division of Viral Diseases, Pathogen Discovery | Dakota Howard et al |
| EPI_ISL_5252842 | Broad Institute Clinical Research Sequencing Platform | Infectious Disease Program, Broad Institute of Harvard and MIT | Siddle et al |
| EPI_ISL_5583951 | Aegis Sciences Corporation | Centers for Disease Control and Prevention Division of Viral Diseases, Pathogen Discovery | Dakota Howard et al |
| EPI_ISL_1400395 | UW Virology Lab | UW Virology Lab | Pavitra Roychoudhury et al |
| EPI_ISL_1448334 | UW Virology Lab | UW Virology Lab | Pavitra Roychoudhury et al |
| EPI_ISL_6541120 | Aegis Sciences Corporation | Centers for Disease Control and Prevention Division of Viral Diseases, Pathogen Discovery | Dakota Howard et al |
| EPI_ISL_2405028 | UW Virology Lab | UW Virology Lab | Pavitra Roychoudhury et al |
| EPI_ISL_6414850 | Quest Diagnostics Incorporated | Centers for Disease Control and Prevention Division of Viral Diseases, Pathogen Discovery | Dakota Howard et al |
| EPI_ISL_5416002 | UW Virology Lab | UW Virology Lab | Pavitra Roychoudhury et al |
| EPI_ISL_5900142 | UW Virology Lab | UW Virology Lab | Pavitra Roychoudhury et al |
| EPI_ISL_3666583 | UW Virology Lab | UW Virology Lab | Pavitra Roychoudhury et al |
| EPI_ISL_495672 | Washington State Department of Health | Seattle Flu Study | Deborah A. Nickerson et al |
| EPI_ISL_3847573 | Aegis Sciences Corporation | Centers for Disease Control and Prevention Division of Viral Diseases, Pathogen Discovery | Dakota Howard et al |
| EPI_ISL_849622 | Washington State Department of Health | Seattle Flu Study | Deborah A. Nickerson et al |
| EPI_ISL_4742598 | UW Virology Lab | UW Virology Lab | Pavitra Roychoudhury et al |
| EPI_ISL_2184739 | Laboratory Corporation of America | Centers for Disease Control and Prevention Division of Viral Diseases, Pathogen Discovery | Dakota Howard et al |
| EPI_ISL_5161732 | UW Virology Lab | UW Virology Lab | Pavitra Roychoudhury et al |
| EPI_ISL_1826178 | UW Virology Lab | UW Virology Lab | Pavitra Roychoudhury et al |
| EPI_ISL_1498032 | UW Virology Lab | UW Virology Lab | Pavitra Roychoudhury et al |
| EPI_ISL_463651 | Washington State Department of Health | Seattle Flu Study | Chu et al |
| EPI_ISL_2432714 | UW Virology Lab | UW Virology Lab | Pavitra Roychoudhury et al |
| EPI_ISL_1621112 | UW Virology Lab | UW Virology Lab | Pavitra Roychoudhury et al |
| EPI_ISL_6931364 | Washington State Department of Health Public Health Laboratories | Washington State Department of Health Public Health Laboratories | Drew MacKellar et al |
| EPI_ISL_4301919 | UW Virology Lab | UW Virology Lab | Pavitra Roychoudhury et al |
| EPI_ISL_5238863 | UW Virology Lab | UW Virology Lab | Pavitra Roychoudhury et al |

| | | | |
|---|---|---|---|
| EPI_ISL_4810846 | Laboratory Corporation of America | Centers for Disease Control and Prevention Division of Viral Diseases, Pathogen Discovery | Dakota Howard et al |
| EPI_ISL_5876173 | Laboratory Corporation of America | Centers for Disease Control and Prevention Division of Viral Diseases, Pathogen Discovery | Dakota Howard et al |
| EPI_ISL_2502987 | UW Virology Lab | UW Virology Lab | Pavitra Roychoudhury et al |
| EPI_ISL_3666221 | UW Virology Lab | UW Virology Lab | Pavitra Roychoudhury et al |
| EPI_ISL_3123459 | Virginia Division of Consolidated Laboratory Services | Virginia Division of Consolidated Laboratory Services | Virginia Division of Consolidated Laboratory Services et al |
| EPI_ISL_1222611 | Seattle Flu Study | Seattle Flu Study | Deborah A. Nickerson et al |
| EPI_ISL_2003568 | Virginia Division of Consolidated Laboratory Services | Virginia Division of Consolidated Laboratory Services | Virginia DCLS et al |
| EPI_ISL_4071370 | Atlas Genomics | Seattle Flu Study | Deborah A. Nickerson et al |
| EPI_ISL_4410975 | Washington State Department of Health Public Health Laboratories | Washington State Department of Health Public Health Laboratories | Drew MacKellar et al |
| EPI_ISL_477705 | UW Virology Lab | UW Virology Lab | Pavitra Roychoudhury et al |
| EPI_ISL_6340993 | Fulgent Genetics | Centers for Disease Control and Prevention Division of Viral Diseases, Pathogen Discovery | Dakota Howard et al |
| EPI_ISL_1222459 | WA State PHL | Seattle Flu Study | Deborah A. Nickerson et al |
| EPI_ISL_6608155 | Washington State Department of Health Public Health Laboratories | Washington State Department of Health Public Health Laboratories | Drew MacKellar et al |
| EPI_ISL_5903461 | Washington State Department of Health Public Health Laboratories | Washington State Department of Health Public Health Laboratories | Drew MacKellar et al |
| EPI_ISL_5990690 | Washington State Department of Health Public Health Laboratories | Washington State Department of Health Public Health Laboratories | Drew MacKellar et al |
| EPI_ISL_3759015 | UW Virology Lab | UW Virology Lab | Pavitra Roychoudhury et al |
| EPI_ISL_6786737 | Aegis Sciences Corporation | Centers for Disease Control and Prevention Division of Viral Diseases, Pathogen Discovery | Dakota Howard et al |
| EPI_ISL_2273738 | UW Virology Lab | UW Virology Lab | Pavitra Roychoudhury et al |
| EPI_ISL_3724423 | Broad Institute Clinical Research Sequencing Platform | Infectious Disease Program, Broad Institute of Harvard and MIT | Siddle et al |
| EPI_ISL_5169878 | Laboratory Corporation of America | Centers for Disease Control and Prevention Division of Viral Diseases, Pathogen Discovery | Dakota Howard et al |
| EPI_ISL_4299407 | Marshfield Clinic Labs | Marshfield Clinic Research Institute - Integrated Research and Development Laboratory | Jen K. Meece et al |
| EPI_ISL_5161543 | UW Virology Lab | UW Virology Lab | Pavitra Roychoudhury et al |
| EPI_ISL_5169230 | Laboratory Corporation of America | Centers for Disease Control and Prevention Division of Viral Diseases, Pathogen Discovery | Dakota Howard et al |

| | | | |
|---|---|---|---|
| EPI_ISL_520729 | Mohammed Bin Rashid University of Medicine and Health Sciences | Al Jalila Genomics Center | Ahmad Abou Tayoun et al |
| EPI_ISL_6436120 | Lighthouse Lab in Milton Keynes | Wellcome Sanger Institute for the COVID-19 Genomics UK (COG-UK) Consortium | The Lighthouse Lab in Milton Keynes et al |
| EPI_ISL_769906 | Wyoming Public Health Laboratory | Wyoming Public Health Laboratory | Noah Hull et al |
| EPI_ISL_1421656 | Wisconsin State Laboratory of Hygiene Communicable Disease Division | Wisconsin State Laboratory of Hygiene Communicable Disease Division | Kelsey R. Florek et al |
| EPI_ISL_726258 | Wales Specialist Virology Centre Sequencing lab: Pathogen Genomics Unit | COVID-19 Genomics UK (COG-UK) Consortium | Catherine Moore et al |
| EPI_ISL_1182683 | University of Wisconsin-Madison AIDS Vaccine Research Laboratories | University of Wisconsin-Madison AIDS Vaccine Research Laboratories | Gage Moreno et al |
| EPI_ISL_859583 | BTC, Khalifa University | BTC, Khalifa University | Al Safar et al |
| EPI_ISL_814298 | Wales Specialist Virology Centre Sequencing lab: Pathogen Genomics Unit | COVID-19 Genomics UK (COG-UK) Consortium | Catherine Moore et al |
| EPI_ISL_650914 | Wales Specialist Virology Centre Sequencing lab: Pathogen Genomics Unit | COVID-19 Genomics UK (COG-UK) Consortium | Catherine Moore et al |
| EPI_ISL_5868070 | Wisconsin State Laboratory of Hygiene Communicable Disease Division | Wisconsin State Laboratory of Hygiene Communicable Disease Division | Abigail C. Shockey et al |
| EPI_ISL_3751943 | Laboratory Corporation of America | Centers for Disease Control and Prevention Division of Viral Diseases, Pathogen Discovery | Dakota Howard et al |
| EPI_ISL_1995644 | Aegis Sciences Corporation | Centers for Disease Control and Prevention Division of Viral Diseases, Pathogen Discovery | Dakota Howard et al |
| EPI_ISL_4212715 | Wyoming Public Health Laboratory | Wyoming Public Health Laboratory | Jim Mildenberger et al |
| EPI_ISL_550041 | Lighthouse Lab in Milton Keynes | Wellcome Sanger Institute for the COVID-19 Genomics UK (COG-UK) consortium | The Lighthouse Lab in Alderley Park et al |
| EPI_ISL_4691213 | QLabs | WVU and Marshall University Combined Genomics Core Facilities | James Denvir et al |
| EPI_ISL_1166146 | University of Wisconsin-Madison AIDS Vaccine Research Laboratories | University of Wisconsin-Madison AIDS Vaccine Research Laboratories | Gage Moreno et al |
| EPI_ISL_1269382 | Wisconsin State Laboratory of Hygiene Communicable Disease Division | Ginkgo Bioworks Clinical Laboratory | Malaika Mckenzie-Bennett et al |
| EPI_ISL_767130 | Lighthouse Lab in Alderley Park | Wellcome Sanger Institute for the COVID-19 Genomics UK (COG-UK) Consortium | Jacquelyn Wynn et al |
| EPI_ISL_1835184 | Aegis Sciences Corporation | Centers for Disease Control and Prevention Division of Viral Diseases, Pathogen Discovery | Dakota Howard et al |

| | | | |
|---|---|---|---|
| EPI_ISL_5018883 | Wisconsin State Laboratory of Hygiene Communicable Disease Division | Wisconsin State Laboratory of Hygiene Communicable Disease Division | Abigail C. Shockey et al |
| EPI_ISL_5078996 | Laboratory Corporation of America | Centers for Disease Control and Prevention Division of Viral Diseases, Pathogen Discovery | Dakota Howard et al |
| EPI_ISL_840525 | Originating lab: Wales Specialist Virology Centre Sequencing lab: Pathogen Genomics Unit | Public Health Wales Microbiology Cardiff Wales Specialist Virology Centre | Catherine Moore et al |
| EPI_ISL_539795 | Wyoming Public Health Laboratory | Wyoming Public Health Laboratory | Noah Hull et al |
| EPI_ISL_4695758 | Summersville Regional Medical Center | WVU and Marshall University Combined Genomics Core Facilities | James Denvir et al |
| EPI_ISL_922754 | Wales Specialist Virology Centre Sequencing lab: Pathogen Genomics Unit | Public Health Wales Microbiology Cardiff Wales Specialist Virology Centre | Catherine Moore et al |
| EPI_ISL_3234914 | Wisconsin State Laboratory of Hygiene Communicable Disease Division | Wisconsin State Laboratory of Hygiene Communicable Disease Division | Abigail C. Shockey et al |
| EPI_ISL_4920051 | Wyoming Public Health Laboratory | Wyoming Public Health Laboratory | Jim Mildenberger et al |
| EPI_ISL_1337604 | Wyoming Public Health Laboratory | Wyoming Public Health Laboratory | Jim Mildenberger et al |
| EPI_ISL_7125157 | Quest Diagnostics Incorporated | Centers for Disease Control and Prevention Division of Viral Diseases, Pathogen Discovery | Dakota Howard et al |
| EPI_ISL_6433062 | Lighthouse Lab in Alderley Park | Wellcome Sanger Institute for the COVID-19 Genomics UK (COG-UK) Consortium | Jacquelyn Wynn et al |
| EPI_ISL_5076242 | Laboratory Corporation of America | Centers for Disease Control and Prevention Division of Viral Diseases, Pathogen Discovery | Dakota Howard et al |
| EPI_ISL_6520165 | Aegis Sciences Corporation | Centers for Disease Control and Prevention Division of Viral Diseases, Pathogen Discovery | Dakota Howard et al |
| EPI_ISL_471265 | Wisconsin State Laboratory of Hygiene Communicable Disease Division | Wisconsin State Laboratory of Hygiene Communicable Disease Division | Kelsey R. Florek et al |
| EPI_ISL_859334 | Lighthouse Lab in Alderley Park | Wellcome Sanger Institute for the COVID-19 Genomics UK (COG-UK) Consortium | Jacquelyn Wynn et al |
| EPI_ISL_1592245 | City of Milwaukee Health Department Laboratory | City of Milwaukee Health Department Laboratory | Sanjib Bhattacharyya et al |
| EPI_ISL_823456 | Wales Specialist Virology Centre Sequencing lab: Pathogen Genomics Unit | COVID-19 Genomics UK (COG-UK) Consortium | Catherine Moore et al |
| EPI_ISL_1454342 | Lighthouse Lab in Alderley Park | Wellcome Sanger Institute for the COVID-19 Genomics UK (COG-UK) Consortium | Jacquelyn Wynn et al |
| EPI_ISL_2455495 | National Hospital for Tropical Diseases | Oxford University Clinical Research Unit, Hanoi, Vietnam | Pham Ngoc Thach et al |

| | | | |
|---|---|---|---|
| EPI_ISL_2594127 | Wyoming Public Health Laboratory | Wyoming Public Health Laboratory | Jim Mildenberger et al |
| EPI_ISL_742925 | Wales Specialist Virology Centre Sequencing lab: Pathogen Genomics Unit | COVID-19 Genomics UK (COG-UK) Consortium | Catherine Moore et al |
| EPI_ISL_858581 | Lighthouse Lab in Alderley Park | Wellcome Sanger Institute for the COVID-19 Genomics UK (COG-UK) Consortium | Jacquelyn Wynn et al |
| EPI_ISL_6263671 | Berkeley Medical Center | WVU and Marshall University Combined Genomics Core Facilities | James Denvir et al |
| EPI_ISL_750430 | Sanatorio Americano | Institut Pasteur de Montevideo | Daiana Mir et al |
| EPI_ISL_1012702 | Lighthouse Lab in Alderley Park | Wellcome Sanger Institute for the COVID-19 Genomics UK (COG-UK) Consortium | Jacquelyn Wynn et al |
| EPI_ISL_666508 | Wales Specialist Virology Centre Sequencing lab: Pathogen Genomics Unit | COVID-19 Genomics UK (COG-UK) Consortium | Catherine Moore et al |
| EPI_ISL_891978 | Lighthouse Lab in Alderley Park | Wellcome Sanger Institute for the COVID-19 Genomics UK (COG-UK) Consortium | Jacquelyn Wynn et al |
| EPI_ISL_582660 | Sheikh Khalifa Medical City | Molecular/Surveillance lab Sheikh Khalifa Medical City | Amirtharaj Francis et al |
| EPI_ISL_706535 | Wales Specialist Virology Centre Sequencing lab: Pathogen Genomics Unit | COVID-19 Genomics UK (COG-UK) Consortium | Catherine Moore et al |
| EPI_ISL_4504008 | National Hospital for Tropical Diseases | Oxford University Clinical Research Unit, Hanoi, Vietnam | Pham Ngoc Thach et al |
| EPI_ISL_767325 | Lighthouse Lab in Alderley Park | Wellcome Sanger Institute for the COVID-19 Genomics UK (COG-UK) Consortium | Jacquelyn Wynn et al |
| EPI_ISL_1060205 | Wyoming Public Health Laboratory | Wyoming Public Health Laboratory | Jim Mildenberger et al |
| EPI_ISL_5690124 | Helix | Centers for Disease Control and Prevention Division of Viral Diseases, Pathogen Discovery | Dakota Howard et al |
| EPI_ISL_4061926 | Wyoming Public Health Laboratory | Wyoming Public Health Laboratory | Jim Mildenberger et al |
| EPI_ISL_514572 | Wales Specialist Virology Centre Sequencing lab: Pathogen Genomics Unit | COVID-19 Genomics UK (COG-UK) Consortium | Catherine Moore et al |
| EPI_ISL_954292 | MRC/UVRI & LSHTM Uganda Research Unit | Where sequence data have been generated and submitted to GISAID | Matthew Cotten et al |
| EPI_ISL_2279285 | Wisconsin State Laboratory of Hygiene Communicable Disease Division | Wisconsin State Laboratory of Hygiene Communicable Disease Division | Abigail C. Shockey et al |
| EPI_ISL_6304381 | Lighthouse Lab in Alderley Park | Wellcome Sanger Institute for the COVID-19 Genomics UK (COG-UK) Consortium | Jacquelyn Wynn et al |
| EPI_ISL_536729 | University of Wisconsin-Madison AIDS Vaccine Research Laboratories | University of Wisconsin-Madison AIDS Vaccine Research Laboratories | Gage Moreno et al |

| | | | |
|---|---|---|---|
| EPI_ISL_4244073 | Fulgent Genetics | Centers for Disease Control and Prevention Division of Viral Diseases, Pathogen Discovery | Dakota Howard et al |
| EPI_ISL_3556356 | Wisconsin State Laboratory of Hygiene Communicable Disease Division | Wisconsin State Laboratory of Hygiene Communicable Disease Division | Abigail C. Shockey et al |
| EPI_ISL_2728144 | Wyoming Public Health Laboratory | Wyoming Public Health Laboratory | Jim Mildenberger et al |
| EPI_ISL_698588 | Group 42 (G42) Healthcare, Abu Dhabi, United Arab Emirates; Department of Health, The United Arab Emirates | G42 Healthcare | Rong Liu et al |
| EPI_ISL_2615637 | J.W. Ruby Memorial Hospital | WVU and Marshall University Combined Genomics Core Facilities | James Denvir et al |
| EPI_ISL_484953 | University of Wisconsin-Madison AIDS Vaccine Research Laboratories | University of Wisconsin-Madison AIDS Vaccine Research Laboratories | Gage Moreno et al |
| EPI_ISL_4691698 | War Memorial Hospital | WVU and Marshall University Combined Genomics Core Facilities | James Denvir et al |
| EPI_ISL_1484435 | Wyoming Public Health Laboratory | Wyoming Public Health Laboratory | Jim Mildenberger et al |
| EPI_ISL_5868020 | Wisconsin State Laboratory of Hygiene Communicable Disease Division | Wisconsin State Laboratory of Hygiene Communicable Disease Division | Abigail C. Shockey et al |
| EPI_ISL_3383329 | Wyoming Public Health Laboratory | Wyoming Public Health Laboratory | Jim Mildenberger et al |
| EPI_ISL_432446 | Wales Specialist Virology Centre | Public Health Wales Microbiology Cardiff | Catherine Moore et al |
| EPI_ISL_6061947 | Aegis Sciences Corporation | Centers for Disease Control and Prevention Division of Viral Diseases, Pathogen Discovery | Dakota Howard et al |
| EPI_ISL_822810 | Wales Specialist Virology Centre Sequencing lab: Pathogen Genomics Unit | COVID-19 Genomics UK (COG-UK) Consortium | Catherine Moore et al |
| EPI_ISL_516443 | University of Wisconsin-Madison AIDS Vaccine Research Laboratories | University of Wisconsin-Madison AIDS Vaccine Research Laboratories | Gage Moreno et al |
| EPI_ISL_5868506 | Wisconsin State Laboratory of Hygiene Communicable Disease Division | Wisconsin State Laboratory of Hygiene Communicable Disease Division | Abigail C. Shockey et al |
| EPI_ISL_2833986 | Wyoming Public Health Laboratory | Wyoming Public Health Laboratory | Jim Mildenberger et al |
| EPI_ISL_6264262 | United Hospital Center | WVU and Marshall University Combined Genomics Core Facilities | James Denvir et al |
| EPI_ISL_4175207 | Laboratory Corporation of America | Centers for Disease Control and Prevention Division of Viral Diseases, Pathogen Discovery | Dakota Howard et al |
| EPI_ISL_4692289 | WVU Rapid Development Lab | WVU and Marshall University Combined Genomics Core Facilities | James Denvir et al |

| | | | |
|---|---|---|---|
| EPI_ISL_4809221 | Laboratory Corporation of America | Centers for Disease Control and Prevention Division of Viral Diseases, Pathogen Discovery | Dakota Howard et al |
| EPI_ISL_710870 | Lighthouse Lab in Alderley Park | Wellcome Sanger Institute for the COVID-19 Genomics UK (COG-UK) Consortium | Jacquelyn Wynn et al |
| EPI_ISL_6102584 | Infinity Biologix | Centers for Disease Control and Prevention Division of Viral Diseases, Pathogen Discovery | Dakota Howard et al |
| EPI_ISL_706832 | Wales Specialist Virology Centre Sequencing lab: Pathogen Genomics Unit | COVID-19 Genomics UK (COG-UK) Consortium | Catherine Moore et al |
| EPI_ISL_6506835 | MRC/UVRI & LSHTM Uganda Research Unit, Uganda Virus Research Institute | CERI, Centre for Epidemic Response and Innvoation, Stellenbosch University and KRISP, KZN Research Innovation and Sequencing Platform, UKZN. | Deogratius Ssemwanga et al |
| EPI_ISL_1624899 | QLabs | WVU and Marshall University Combined Genomics Core Facilities | James Denvir et al |
| EPI_ISL_5347344 | Pasteur Institute in Ho Chi Minh city | Department of Microbiology and Immunology | Vũ Phạm Hồng Nhung et al |
| EPI_ISL_1277982 | Lighthouse Lab in Alderley Park | Wellcome Sanger Institute for the COVID-19 Genomics UK (COG-UK) Consortium | Jacquelyn Wynn et al |
| EPI_ISL_2294598 | University of Wisconsin-Madison AIDS Vaccine Research Laboratories | University of Wisconsin-Madison AIDS Vaccine Research Laboratories | Gage Moreno et al |
| EPI_ISL_6264175 | Berkeley Medical Center | WVU and Marshall University Combined Genomics Core Facilities | James Denvir et al |
| EPI_ISL_767207 | Lighthouse Lab in Alderley Park | Wellcome Sanger Institute for the COVID-19 Genomics UK (COG-UK) Consortium | Jacquelyn Wynn et al |
| EPI_ISL_5323839 | Wyoming Public Health Laboratory | Wyoming Public Health Laboratory | Jim Mildenberger et al |
| EPI_ISL_1140066 | Wyoming Public Health Laboratory | Wyoming Public Health Laboratory | Jim Mildenberger et al |
| EPI_ISL_422071 | Wales Specialist Virology Centre | Public Health Wales Microbiology Cardiff | Catherine Moore et al |
| EPI_ISL_4389410 | Infinity Biologix | Centers for Disease Control and Prevention Division of Viral Diseases, Pathogen Discovery | Dakota Howard et al |
| EPI_ISL_3099824 | Wyoming Public Health Laboratory | Wyoming Public Health Laboratory | Jim Mildenberger et al |
| EPI_ISL_1295285 | Wisconsin State Laboratory of Hygiene Communicable Disease Division | Wisconsin State Laboratory of Hygiene Communicable Disease Division | Kelsey R. Florek et al |
| EPI_ISL_6673737 | Lighthouse Lab in Milton Keynes | Wellcome Sanger Institute for the COVID-19 Genomics UK (COG-UK) Consortium | The Lighthouse Lab in Milton Keynes et al |
| EPI_ISL_422151 | Wales Specialist Virology Centre | Public Health Wales Microbiology Cardiff | Catherine Moore et al |

| | | | |
|---|---|---|---|
| EPI_ISL_6264209 | Princeton Community Hospital | WVU and Marshall University Combined Genomics Core Facilities | James Denvir et al |
| EPI_ISL_2137771 | Wyoming Public Health Laboratory | Wyoming Public Health Laboratory | Jim Mildenberger et al |
| EPI_ISL_1012694 | Lighthouse Lab in Alderley Park | Wellcome Sanger Institute for the COVID-19 Genomics UK (COG-UK) Consortium | Jacquelyn Wynn et al |
| EPI_ISL_652617 | Wales Specialist Virology Centre Sequencing lab: Pathogen Genomics Unit | COVID-19 Genomics UK (COG-UK) Consortium | Catherine Moore et al |
| EPI_ISL_2750552 | Originating lab: Wales Specialist Virology Centre Sequencing lab: Pathogen Genomics Unit | Public Health Wales Microbiology Cardiff Wales Specialist Virology Centre | Catherine Moore et al |
| EPI_ISL_4134785 | Originating lab: Wales Specialist Virology Centre Sequencing lab: Pathogen Genomics Unit | Public Health Wales Microbiology Cardiff Wales Specialist Virology Centre | Catherine Moore et al |
| EPI_ISL_1546947 | Originating lab: Wales Specialist Virology Centre Sequencing lab: Pathogen Genomics Unit | Public Health Wales Microbiology Cardiff Wales Specialist Virology Centre | Catherine Moore et al |
| EPI_ISL_447623 | Virology, Wageningen Bioveterinary Research | Virology, Wageningen Bioveterinary Research | van der Poel et al |
| EPI_ISL_2624587 | Originating lab: Wales Specialist Virology Centre Sequencing lab: Pathogen Genomics Unit | Public Health Wales Microbiology Cardiff Wales Specialist Virology Centre | Catherine Moore et al |
| EPI_ISL_4794433 | Originating lab: Wales Specialist Virology Centre Sequencing lab: Pathogen Genomics Unit | Public Health Wales Microbiology Cardiff Wales Specialist Virology Centre | Catherine Moore et al |
| EPI_ISL_1106954 | Wales Specialist Virology Centre Sequencing lab: Pathogen Genomics Unit | Public Health Wales Microbiology Cardiff Wales Specialist Virology Centre | Catherine Moore et al |
| EPI_ISL_5284490 | Originating lab: Wales Specialist Virology Centre Sequencing lab: Pathogen Genomics Unit | Public Health Wales Microbiology Cardiff Wales Specialist Virology Centre | Catherine Moore et al |
| EPI_ISL_6142827 | Originating lab: Wales Specialist Virology Centre Sequencing lab: Pathogen Genomics Unit | Public Health Wales Microbiology Cardiff Wales Specialist Virology Centre | Catherine Moore et al |
| EPI_ISL_3081448 | Originating lab: Wales Specialist Virology Centre Sequencing lab: Pathogen Genomics Unit | Public Health Wales Microbiology Cardiff Wales Specialist Virology Centre | Catherine Moore et al |
| EPI_ISL_4134007 | Originating lab: Wales Specialist Virology Centre Sequencing lab: Pathogen Genomics Unit | Public Health Wales Microbiology Cardiff Wales Specialist Virology Centre | Catherine Moore et al |

| | | | |
|---|---|---|---|
| EPI_ISL_2906576 | Originating lab: Wales Specialist Virology Centre Sequencing lab: Pathogen Genomics Unit | Public Health Wales Microbiology Cardiff Wales Specialist Virology Centre | Catherine Moore et al |
| EPI_ISL_3422753 | Originating lab: Wales Specialist Virology Centre Sequencing lab: Pathogen Genomics Unit | Public Health Wales Microbiology Cardiff Wales Specialist Virology Centre | Catherine Moore et al |
| EPI_ISL_6285850 | Originating lab: Wales Specialist Virology Centre Sequencing lab: Pathogen Genomics Unit | Public Health Wales Microbiology Cardiff Wales Specialist Virology Centre | Catherine Moore et al |
| EPI_ISL_5303298 | Wales Specialist Virology Centre Sequencing lab: Pathogen Genomics Unit | Public Health Wales Microbiology Cardiff Wales Specialist Virology Centre | Catherine Moore et al |
| EPI_ISL_4315047 | Originating lab: Wales Specialist Virology Centre Sequencing lab: Pathogen Genomics Unit | Public Health Wales Microbiology Cardiff Wales Specialist Virology Centre | Catherine Moore et al |
| EPI_ISL_6292899 | Originating lab: Wales Specialist Virology Centre Sequencing lab: Pathogen Genomics Unit | Public Health Wales Microbiology Cardiff Wales Specialist Virology Centre | Catherine Moore et al |
| EPI_ISL_4136425 | Originating lab: Wales Specialist Virology Centre Sequencing lab: Pathogen Genomics Unit | Public Health Wales Microbiology Cardiff Wales Specialist Virology Centre | Catherine Moore et al |
| EPI_ISL_3963723 | Originating lab: Wales Specialist Virology Centre Sequencing lab: Pathogen Genomics Unit | Public Health Wales Microbiology Cardiff Wales Specialist Virology Centre | Catherine Moore et al |
| EPI_ISL_4532823 | Originating lab: Wales Specialist Virology Centre Sequencing lab: Pathogen Genomics Unit | Public Health Wales Microbiology Cardiff Wales Specialist Virology Centre | Catherine Moore et al |
| EPI_ISL_4134563 | Originating lab: Wales Specialist Virology Centre Sequencing lab: Pathogen Genomics Unit | Public Health Wales Microbiology Cardiff Wales Specialist Virology Centre | Catherine Moore et al |
| EPI_ISL_5300712 | Originating lab: Wales Specialist Virology Centre Sequencing lab: Pathogen Genomics Unit | Public Health Wales Microbiology Cardiff Wales Specialist Virology Centre | Catherine Moore et al |
| EPI_ISL_3293652 | Wales Specialist Virology Centre Sequencing lab: Pathogen Genomics Unit | Public Health Wales Microbiology Cardiff Wales Specialist Virology Centre | Catherine Moore et al |
| EPI_ISL_1547159 | Originating lab: Wales Specialist Virology Centre Sequencing lab: Pathogen Genomics Unit | Public Health Wales Microbiology Cardiff Wales Specialist Virology Centre | Catherine Moore et al |

| | | | |
|---|---|---|---|
| EPI_ISL_4789925 | Wales Specialist Vir Pathogen Genomics Unit | Public Health Wales Microbiology Cardiff Wales Specialist Virology Centre | Catherine Moore et al |
| EPI_ISL_4788182 | Wales Specialist Vir Pathogen Genomics Unit | Public Health Wales Microbiology Cardiff Wales Specialist Virology Centre | Catherine Moore et al |
| EPI_ISL_5285414 | Originating lab: Wales Specialist Virology Centre Sequencing lab: Pathogen Genomics Unit | Public Health Wales Microbiology Cardiff Wales Specialist Virology Centre | Catherine Moore et al |
| EPI_ISL_5507215 | Originating lab: Wales Specialist Virology Centre Sequencing lab: Pathogen Genomics Unit | Public Health Wales Microbiology Cardiff Wales Specialist Virology Centre | Catherine Moore et al |
| EPI_ISL_3422465 | Originating lab: Wales Specialist Virology Centre Sequencing lab: Pathogen Genomics Unit | Public Health Wales Microbiology Cardiff Wales Specialist Virology Centre | Catherine Moore et al |
| EPI_ISL_1388039 | Wales Specialist Virology Centre Sequencing lab: Pathogen Genomics Unit | Public Health Wales Microbiology Cardiff Wales Specialist Virology Centre | Catherine Moore et al |
| EPI_ISL_5295346 | Originating lab: Wales Specialist Virology Centre Sequencing lab: Pathogen Genomics Unit | Public Health Wales Microbiology Cardiff Wales Specialist Virology Centre | Catherine Moore et al |
| EPI_ISL_3128559 | Wisconsin Veterinary Diagnostic Laboratory | Diagnostic Virology Laboratory, National Veterinary Services Laboratories, USDA1920 Dayton Avenue, Ames, IA 50010, USA | Kerrie M. Franzen et al |
| EPI_ISL_5296479 | Originating lab: Wales Specialist Virology Centre Sequencing lab: Pathogen Genomics Unit | Public Health Wales Microbiology Cardiff Wales Specialist Virology Centre | Catherine Moore et al |
| EPI_ISL_5289411 | Originating lab: Wales Specialist Virology Centre Sequencing lab: Pathogen Genomics Unit | Public Health Wales Microbiology Cardiff Wales Specialist Virology Centre | Catherine Moore et al |
| EPI_ISL_4313045 | Originating lab: Wales Specialist Virology Centre Sequencing lab: Pathogen Genomics Unit | Public Health Wales Microbiology Cardiff Wales Specialist Virology Centre | Catherine Moore et al |
| EPI_ISL_1106109 | Wales Specialist Virology Centre Sequencing lab: Pathogen Genomics Unit | Public Health Wales Microbiology Cardiff Wales Specialist Virology Centre | Catherine Moore et al |
| EPI_ISL_2818896 | Originating lab: Wales Specialist Virology Centre Sequencing lab: Pathogen Genomics Unit | Public Health Wales Microbiology Cardiff Wales Specialist Virology Centre | Catherine Moore et al |
| EPI_ISL_6141803 | Originating lab: Wales Specialist Virology Centre Sequencing lab: Pathogen Genomics Unit | Public Health Wales Microbiology Cardiff Wales Specialist Virology Centre | Catherine Moore et al |

| | | | |
|---|---|---|---|
| EPI_ISL_4316040 | Originating lab: Wales Specialist Virology Centre Sequencing lab: Pathogen Genomics Unit | Public Health Wales Microbiology Cardiff Wales Specialist Virology Centre | Catherine Moore et al |
| EPI_ISL_3791875 | Originating lab: Wales Specialist Virology Centre Sequencing lab: Pathogen Genomics Unit | Public Health Wales Microbiology Cardiff Wales Specialist Virology Centre | Catherine Moore et al |
| EPI_ISL_3424034 | Originating lab: Wales Specialist Virology Centre Sequencing lab: Pathogen Genomics Unit | Public Health Wales Microbiology Cardiff Wales Specialist Virology Centre | Catherine Moore et al |
| EPI_ISL_4317360 | Originating lab: Wales Specialist Virology Centre Sequencing lab: Pathogen Genomics Unit | Public Health Wales Microbiology Cardiff Wales Specialist Virology Centre | Catherine Moore et al |
| EPI_ISL_3182108 | Wales Specialist Virology Centre Sequencing lab: Pathogen Genomics Unit | Public Health Wales Microbiology Cardiff Wales Specialist Virology Centre | Catherine Moore et al |
| EPI_ISL_5492870 | Originating lab: Wales Specialist Virology Centre Sequencing lab: Pathogen Genomics Unit | Public Health Wales Microbiology Cardiff Wales Specialist Virology Centre | Catherine Moore et al |
| EPI_ISL_4317318 | Originating lab: Wales Specialist Virology Centre Sequencing lab: Pathogen Genomics Unit | Public Health Wales Microbiology Cardiff Wales Specialist Virology Centre | Catherine Moore et al |
| EPI_ISL_5288534 | Originating lab: Wales Specialist Virology Centre Sequencing lab: Pathogen Genomics Unit | Public Health Wales Microbiology Cardiff Wales Specialist Virology Centre | Catherine Moore et al |
| EPI_ISL_2232940 | Institute for Water Quality and Resource Management, Technical University Vienna | Bergthaler laboratory, CeMM Research Center for Molecular Medicine of the Austrian Academy of Sciences | Lukas Endler et al |
| EPI_ISL_3004501 | Originating lab: Wales Specialist Virology Centre Sequencing lab: Pathogen Genomics Unit | Public Health Wales Microbiology Cardiff Wales Specialist Virology Centre | Catherine Moore et al |
| EPI_ISL_4136209 | Originating lab: Wales Specialist Virology Centre Sequencing lab: Pathogen Genomics Unit | Public Health Wales Microbiology Cardiff Wales Specialist Virology Centre | Catherine Moore et al |
| EPI_ISL_4314691 | Originating lab: Wales Specialist Virology Centre Sequencing lab: Pathogen Genomics Unit | Public Health Wales Microbiology Cardiff Wales Specialist Virology Centre | Catherine Moore et al |
| EPI_ISL_4878316 | The Ohio State University Infectious Diseases Institute | USDA National Veterinary Services Laboratory | Andrew S. Bowman et al |

| | | | |
|---|---|---|---|
| EPI_ISL_6293475 | Originating lab: Wales Specialist Virology Centre Sequencing lab: Pathogen Genomics Unit | Public Health Wales Microbiology Cardiff Wales Specialist Virology Centre | Catherine Moore et al |
| EPI_ISL_3082981 | Originating lab: Wales Specialist Virology Centre Sequencing lab: Pathogen Genomics Unit | Public Health Wales Microbiology Cardiff Wales Specialist Virology Centre | Catherine Moore et al |
| EPI_ISL_3965441 | Originating lab: Wales Specialist Virology Centre Sequencing lab: Pathogen Genomics Unit | Public Health Wales Microbiology Cardiff Wales Specialist Virology Centre | Catherine Moore et al |
| EPI_ISL_5285535 | Originating lab: Wales Specialist Virology Centre Sequencing lab: Pathogen Genomics Unit | Public Health Wales Microbiology Cardiff Wales Specialist Virology Centre | Catherine Moore et al |
| EPI_ISL_5285206 | Originating lab: Wales Specialist Virology Centre Sequencing lab: Pathogen Genomics Unit | Public Health Wales Microbiology Cardiff Wales Specialist Virology Centre | Catherine Moore et al |
| EPI_ISL_5294644 | Originating lab: Wales Specialist Virology Centre Sequencing lab: Pathogen Genomics Unit | Public Health Wales Microbiology Cardiff Wales Specialist Virology Centre | Catherine Moore et al |
| EPI_ISL_5044463 | Originating lab: Wales Specialist Virology Centre Sequencing lab: Pathogen Genomics Unit | Public Health Wales Microbiology Cardiff Wales Specialist Virology Centre | Catherine Moore et al |
| EPI_ISL_7011411 | Originating lab: Wales Specialist Virology Centre Sequencing lab: Pathogen Genomics Unit | Public Health Wales Microbiology Cardiff Wales Specialist Virology Centre | Catherine Moore et al |
| EPI_ISL_5296790 | Originating lab: Wales Specialist Virology Centre Sequencing lab: Pathogen Genomics Unit | Public Health Wales Microbiology Cardiff Wales Specialist Virology Centre | Catherine Moore et al |
| EPI_ISL_6998778 | Originating lab: Wales Specialist Virology Centre Sequencing lab: Pathogen Genomics Unit | Public Health Wales Microbiology Cardiff Wales Specialist Virology Centre | Catherine Moore et al |
| EPI_ISL_7003303 | Originating lab: Wales Specialist Virology Centre Sequencing lab: Pathogen Genomics Unit | Public Health Wales Microbiology Cardiff Wales Specialist Virology Centre | Catherine Moore et al |
| EPI_ISL_4802234 | Wales Specialist Virology Centre Sequencing lab: Pathogen Genomics Unit | Public Health Wales Microbiology Cardiff Wales Specialist Virology Centre | Catherine Moore et al |

| | | | |
|---|---|---|---|
| EPI_ISL_406798 | General Hospital of Central Theater Command of People's Liberation Army of China | BGI & Institute of Microbiology, Chinese Academy of Sciences & Shandong First Medical University & Shandong Academy of Medical Sciences & General Hospital of Central Theater Command of People's Liberation Army of China | Weijun Chen et al |
| EPI_ISL_6285234 | Originating lab: Wales Specialist Virology Centre Sequencing lab: Pathogen Genomics Unit | Public Health Wales Microbiology Cardiff Wales Specialist Virology Centre | Catherine Moore et al |
| EPI_ISL_402125 | National Institute for Communicable Disease Control and Prevention (ICDC) Chinese Center for Disease Control and Prevention (China CDC) | National Institute for Communicable Disease Control and Prevention (ICDC) Chinese Center for Disease Control and Prevention (China CDC) | Zhang et al |
| EPI_ISL_5817711 | Originating lab: Wales Specialist Virology Centre Sequencing lab: Pathogen Genomics Unit | Public Health Wales Microbiology Cardiff Wales Specialist Virology Centre | Catherine Moore et al |
| EPI_ISL_3791917 | Originating lab: Wales Specialist Virology Centre Sequencing lab: Pathogen Genomics Unit | Public Health Wales Microbiology Cardiff Wales Specialist Virology Centre | Catherine Moore et al |
| EPI_ISL_7003822 | Originating lab: Wales Specialist Virology Centre Sequencing lab: Pathogen Genomics Unit | Public Health Wales Microbiology Cardiff Wales Specialist Virology Centre | Catherine Moore et al |
| EPI_ISL_7012690 | Originating lab: Wales Specialist Virology Centre Sequencing lab: Pathogen Genomics Unit | Public Health Wales Microbiology Cardiff Wales Specialist Virology Centre | Catherine Moore et al |
| EPI_ISL_5299317 | Originating lab: Wales Specialist Virology Centre Sequencing lab: Pathogen Genomics Unit | Public Health Wales Microbiology Cardiff Wales Specialist Virology Centre | Catherine Moore et al |
| EPI_ISL_2904989 | Originating lab: Wales Specialist Virology Centre Sequencing lab: Pathogen Genomics Unit | Public Health Wales Microbiology Cardiff Wales Specialist Virology Centre | Catherine Moore et al |
| EPI_ISL_6292663 | Originating lab: Wales Specialist Virology Centre Sequencing lab: Pathogen Genomics Unit | Public Health Wales Microbiology Cardiff Wales Specialist Virology Centre | Catherine Moore et al |
| EPI_ISL_4797435 | Wales Specialist Virology Centre Sequencing lab: Pathogen Genomics Unit | Public Health Wales Microbiology Cardiff Wales Specialist Virology Centre | Catherine Moore et al |
| EPI_ISL_3004920 | Originating lab: Wales Specialist Virology Centre Sequencing lab: Pathogen Genomics Unit | Public Health Wales Microbiology Cardiff Wales Specialist Virology Centre | Catherine Moore et al |

| | | | |
|---|---|---|---|
| EPI_ISL_6997000 | Originating lab: Wales Specialist Virology Centre Sequencing lab: Pathogen Genomics Unit | Public Health Wales Microbiology Cardiff Wales Specialist Virology Centre | Catherine Moore et al |
